# Optimization and functionalization of red-shifted rhodamine dyes

**DOI:** 10.1101/2019.12.20.881227

**Authors:** Jonathan B. Grimm, Ariana N. Tkachuk, Heejun Choi, Boaz Mohar, Natalie Falco, Ronak Patel, Jennifer Lippincott-Schwartz, Timothy A. Brown, Luke D. Lavis

## Abstract

Expanding the palette of fluorescent dyes is vital for pushing the frontier of biological imaging. Although rhodamine dyes remain the premier type of small-molecule fluorophore due to their bioavailability and brightness, variants excited with far-red or near-infrared light suffer from poor performance due to their propensity to adopt a lipophilic, nonfluorescent form. We report a general chemical modification for rhodamines that optimizes long-wavelength variants and enables facile functionalization with different chemical groups.

---

The development of hybrid small-molecule:protein labeling strategies enable the use of chemical fluorophores in living cells and *in vivo*^1^. Optimizing small-molecule dyes for these complex biological environments is important, as synthetic fluorophores are often brighter and more photostable than fluorescent proteins^2^. We recently developed general methods to improve^3^ and fine-tune^4^ rhodamine fluorophores by incorporating four-membered azetidines into the structure, yielding the ‘Janelia Fluor’ dyes. Although our existing tuning strategies allow optimization of short-wavelength rhodamines, we discovered these methods cannot be applied to analogs excited with far-red and near-infrared (NIR) light because such dyes strongly favor formation of a colorless species. Here, we report a new complementary tuning strategy that allows rational optimization of a broader palette of fluorophores. This general method also serves as a basis for facile functionalization, enabling the synthesis of novel cell- and tissue-permeable rhodamine labels for biological imaging experiments.

The bioavailability and behavior of rhodamine dyes is dictated by a key property—the equilibrium between the nonfluorescent lactone and the fluorescent zwitterion (**Fig. 1a**). Our previous work on rhodamine dyes^3–6^ allows definition of a general rubric that directly correlates the lactone–zwitterion equilibrium constant (*K*_L–Z_) of the free dye to the cellular performance of HaloTag^7^ ligand derivatives (**Fig. 1b**). Dyes with high *K*_L–Z_ exist almost exclusively in the zwitterionic form, making them useful as environmentally insensitive biomolecule labels^8^. Rhodamines with intermediate *K*_L–Z_ values exhibit improved cell and tissue permeability due to the modestly higher propensity of the molecule to adopt the lipophilic lactone form and rapidly traverse biological membranes^4,6^. Dyes exhibiting even smaller *K*_L–Z_ values (*K*_L–Z_ = 10^−2^–10^−3^) preferentially adopt the closed lactone form, which can be exploited to create ‘fluorogenic’ ligands^6,9,10^ as binding to the HaloTag protein shifts the equilibrium to the fluorescent form. This property also decreases *in vivo* utility, however, due to problems with solubility and sequestration in membranes. Finally, dyes with extremely small *K*_L–Z_ values (< 10^−3^) exist completely in the nonfluorescent lactone form, rendering them effectively unusable in biological experiments.

**Figure 1.**
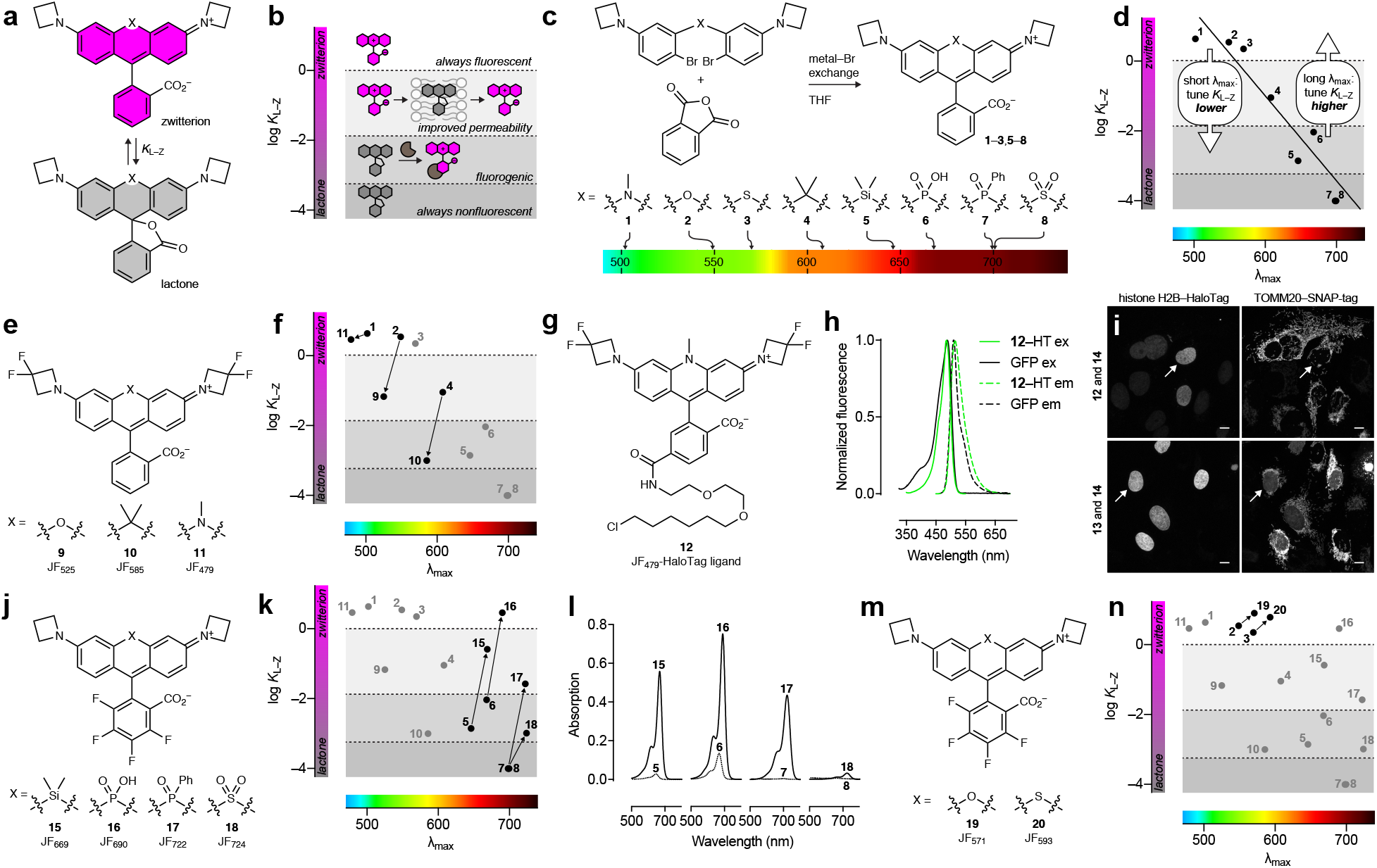
(**a**) The lactone–zwitterion equilibrium of Janelia Fluor rhodamine dyes and the equilibrium constant: *K*_L–Z_. (**b**) Phenomenological plot detailing the properties of different dyes based on *K*_L–Z_. (**c**) Synthesis, structures, and λ_max_ of dyes **1**–**8**. (**d**) Plot of *K*_L–Z_ *vs*. *λ*_max_ for dyes **1**–**8** and general tuning strategies for dyes with short or long λ_max_. (**e**) Structures of dyes **9**–**11**. (**f**) Plot of *K*_L–Z_ *vs*. λ_max_ showing decreased *K*_L–Z_ for dyes **9**–**11**. (**g**) Structure of JF_479_–HaloTag ligand (**12**). (**h**) Fluorescence excitation (ex) and emission (em) spectra of **12**–HaloTag conjugate and GFP. (**i**) Two-color confocal imaging experiment of U2OS cells expressing histone H2B–HaloTag fusion protein labeled with either **12** or JF_503_–HaloTag ligand (**13**) excited with 488 nm and TOMM20–SNAP-tag fusion protein labeled with JF_525_–SNAP-tag ligand (**14**) excited with 532 nm; scale bars: 10 μm. (**j**) Structures of dyes **15**–**18**. (**k**) Plot of *K*_L–Z_ *vs*. λ_max_ showing increased *K*_L–Z_ for dyes **15**–**18**. (**l**) Absorption spectra of **5**–**8** and fluorinated analogs **15**–**18**. (**m**) Structures of dyes **19**–**20**. (**n**) Plot of *K*_L–Z_ *vs. λ*_max_ showing increased *K*_L–Z_ for dyes **19**–**20**.

We compared a series of Janelia Fluor rhodamine analogs with different fluorophoric systems (**1–8**, **Fig. 1c**). Compounds **2, 4**, and **5** were described previously and include the azetidine-containing rhodamine (**2**), which we termed ‘Janelia Fluor 549’ (JF_549_), and the carbo- and Si-rhodamine analogs **4** and **5** (JF_608_ and JF_646_, respectively; **Fig. 1c**)^3^. We expanded the wavelength range of the JF dyes by synthesizing new analogs based on known rhodamine structures containing nitrogen^11,12^ (NCH_3_; **1**), sulfur^13,14^ (S; **3** and SO_2_; **8**), and phosphorous^15,16^ (PO_2_H; 6 and P(O)Ph; 7) starting from *bis*-arylbromides^5^ (**Fig. 1c, Supplementary Note**). We evaluated the spectral and chemical properties of these dyes: absorption maximum (*λ*_max_), extinction coefficient at *λ*_max_ (*ε*), fluorescence emission maximum *λ*_em_, fluorescence quantum yield (Φ), and *K*_L–Z_ (**Table 1**). Comparing *K*_L–Z_ and *λ*_max_ uncovered an inverse correlation (**Fig. 1d**), with the short wavelength NCH3-containing JF_502_ exhibiting a high *K*_L–Z_ = 5.5 and the near-infrared (NIR) dyes containing SO_2_ and P(O)Ph showing a low *K*_L–Z_ ≈ 10^−4^.

**Table 1.** Properties of Janelia Fluor dyes **1**–**11**, **15**–**20**, and **43**, and **45**.

| Compound | Name | X | Y | NR<sub>2</sub> | $\lambda_{\text{max}}$ (nm) | $\epsilon_w$ (M<sup>-1</sup>cm<sup>-1</sup>) | $\lambda_{\text{em}}$ (nm) | $\Phi$ | $K_{\text{L-z}}$ |
| --- | --- | --- | --- | --- | --- | --- | --- | --- | --- |
| 1 | JF<sub>502</sub> |  | H |  | 502 | 57,800 | 533 | 0.71 | 4.33 |
| 2 | JF<sub>549</sub> |  | H |  | 549 | 101,000 | 571 | 0.88 | 3.47 |
| 3 | JF<sub>570</sub> |  | H |  | 570 | 83,600 | 593 | 0.63 | 2.24 |
| 4 | JF<sub>608</sub> |  | H |  | 608 | 99,000 | 631 | 0.67 | 0.091 |
| 5 | JF<sub>646</sub> |  | H |  | 646 | 5,600 | 664 | 0.54 | 0.0014 |
| 6 | JF<sub>668</sub> |  | H |  | 668 | 26,700 | 687 | 0.34 | 0.0093 |
| 7 | - |  | H |  | ~704 | <200 | ~723 | - | <0.0001 |
| 8 | - |  | H |  | - | <100 | - | - | <0.0001 |
| 9 | JF<sub>525</sub> |  | H |  | 525 | 94,000 | 549 | 0.91 | 0.068 |
| 10 | JF<sub>585</sub> |  | H |  | 585 | 1,500 | 609 | 0.78 | ~0.001 |
| 11 | JF<sub>479</sub> |  | H |  | 479 | 47,900 | 517 | 0.62 | 2.88 |
| 15 | JF<sub>669</sub> |  | F |  | 669 | 112,000 | 682 | 0.37 | 0.262 |
| 16 | JF<sub>690</sub> |  | F |  | 690 | 150,000 | 707 | 0.24 | 2.90 |
| 17 | JF<sub>722</sub> |  | F |  | 722 | 87,200 | 743 | 0.11 | 0.026 |
| 18 | JF<sub>724</sub> |  | F |  | 724 | 6,600 | 748 | 0.05 | ~0.001 |
| 19 | JF<sub>571</sub> |  | F |  | 571 | 101,000 | 590 | 0.78 | 7.93 |
| 20 | JF<sub>593</sub> |  | F |  | 593 | 90,300 | 612 | 0.55 | 6.06 |
| 43 | JF<sub>559</sub> |  | F |  | 559 | 106,000 | 579 | 0.85 | 6.22 |
| 45 | JF<sub>711</sub> |  | F |  | 711 | 12,400 | 732 | 0.17 | ~0.001 |

Due to the inverse correlation of *K*_L–Z_ and *λ*_max_, short-wavelength dyes would typically require tuning *K*_L–Z_ *lower* to improve tissue permeability and create fluorogenic HaloTag ligands (**Fig. 1d**) ^4,6,10^. In previous work^4^, we found incorporation of 3,3-difluoroazetidines into the Janelia Fluor structure decreases *K*_L–Z_ and elicits a hypsochromic shift of ~25 nm. This strategy transformed JF_549_ (**2**; *K*_L–Z_ = 3.5) into JF_525_ (**9**; *K*_L–Z_ = 0.68, **Fig. 1e,f**, **Table 1**); the JF_525_–HaloTag ligand shows improved *in vivo* bioavailability and is an excellent component of the hybrid voltage indicator Voltron^2^. This modification also transformed JF_608_ (**4**; *K*_L–Z_ = 0.091) into the highly fluorogenic JF_585_ (**10**; *K*_L–Z_ = 0.001, **Fig. 1e,f**, **Table 1**). Given the relatively high *K*_L–Z_ = 4.33 observed for JF_502_ (**1**) we applied this same tuning strategy to yield the fluorinated JF_479_ (**11**) that showed the expected decrease in *λ*_max_ and *K*_L–Z_ = 2.88 (**Fig. 1e,f, Table 1**, **Supplementary Fig. 1a–c**). The JF_479_–HaloTag ligand (**12**, **Fig. 1g**) exhibits similar spectral properties to enhanced green fluorescent protein (GFP) when attached to the HaloTag protein (**Fig. 1h**) allowing imaging with low bleed-through compared to the similarly cell-permeable—albeit brighter—JF_503_ ligand (**13**) when multiplexed with JF_525_-SNAP-tag ligand (**14**; **Fig. 1i**, **Supplementary Fig. 1d–i**)^4,7^.

For the far-red and NIR rhodamines (**5**–**8**, **Fig. 1c**) the *K*_L–Z_ *vs*. λ_max_ relationship reveals the need for alternative tuning strategy to *increase K*_L–Z_ (**Fig. 1d**). This would improve the *in vivo* performance of Si-rhodamine dyes such as JF_646_ and rescue the NIR P(O)Ph- and SO_2_-containing dyes (**7**–**8**). We previously showed that halogenation of the pendant phenyl ring system can substantially increase the *K*_L–Z_ of Si-rhodamine dyes^5^, presumably by lowering the p*K*a of the benzoic acid moiety; this substitution also elicits a bathochromic shift^17^. For example, JF_646_ (**5**; *λ*_max_/*λ*_em_ = 646 nm/664 nm) exhibits a *K*_L–Z_ = 0.0014 but the fluorinated analog, JF_669_ (**15**; *λ*_max_/*λ*em = 669 nm/682 nm), is higher with *K*_L–Z_ = 0.262 (**Fig. 1j,k**, **Supplementary Fig. 1j,k**). This manifests in a higher absorptivity in aqueous solution with **5** exhibiting ε = 5,600 M^−1^cm^−1^ but the fluorinated analog **15** showing ε= 112,000 M^−1^cm^−1^ (**Fig. 1l**, **Table 1**). We surmised this strategy would be general and prepared the fluorinated PO_2_H-, P(O)Ph-, and SO_2_-containing rhodamines (**16**–**18**, **Fig. 1j**) by replacing phthalic anhydride with tetrafluorophthalic anhydride in our synthetic scheme (**Supplementary Fig. 1j**, **Supplementary Note**). This modification increased *K*_L–Z_ and *ε* and elicited a ~23 nm shift in *λ*_max_ (**Fig. 1j–l**, **Table 1, Supplementary Fig. l–n**). In particular, the fluorinated P(O)Ph-derivative **17** strongly absorbs visible light in aqueous solution (*ε*= 87,000 M^−1^cm^−1^; *λ*_max_ = 722 nm) compared to the parent compound **7** (*ε* < 200 M^−1^cm^−1^; **Fig. 1l**). This trend was generalizable to oxygen- and sulfur-containing rhodamines based on **2** and **3** where the fluorine substitution on the pendant phenyl ring also increased *K*_L–Z_ and *λ*_max_ (**19**–**20**, **Fig. 1m,n**, **Table 1, Supplementary Fig. 1o,p**).

We then explored conjugatable versions of these new far-red and NIR dyes. In addition to increasing *λ*_max_ and *K*_L–Z_, the halogenated phenyl ring motif can also serve as an electrophile for attack by thiols through a nucleophilic aromatic substitution reaction (SNAr), an established strategy for preparing xanthene dye derivatives^8,18^. The reactivity of other nucleophiles was unknown, however; we discovered that N_3_^−^, CN^−^, NH_3_, and NH_2_OH could react with JF_669_ (**15**) to provide derivatives **21**–**24**(**Fig. 2a**). This reaction type was generalizable to other fluorinated rhodamines and regioselective at the 6-position (**Supplementary Note**). Although beyond the scope of this report, we briefly investigated some of these derivatives, finding azide **21** was an excellent reactant in strain-promoted ‘click chemistry’ with cyclic alkynes^19^ **25** and **26** to form triazole adducts **27** and **28** (**Supplementary Fig. 2a**), validating the regiochemistry of the amine addition to form **22** using intermediates **21** and **29** (**Supplementary Fig. 2b**), and testing the reactivity of amine-containing ion-chelating groups **30** and **31** which generated novel prototype far-red indicators for K^+^ and Zn^2+^ (**32**–**33**, **Supplementary Fig. 2c**–**e**).

**Figure 2.**
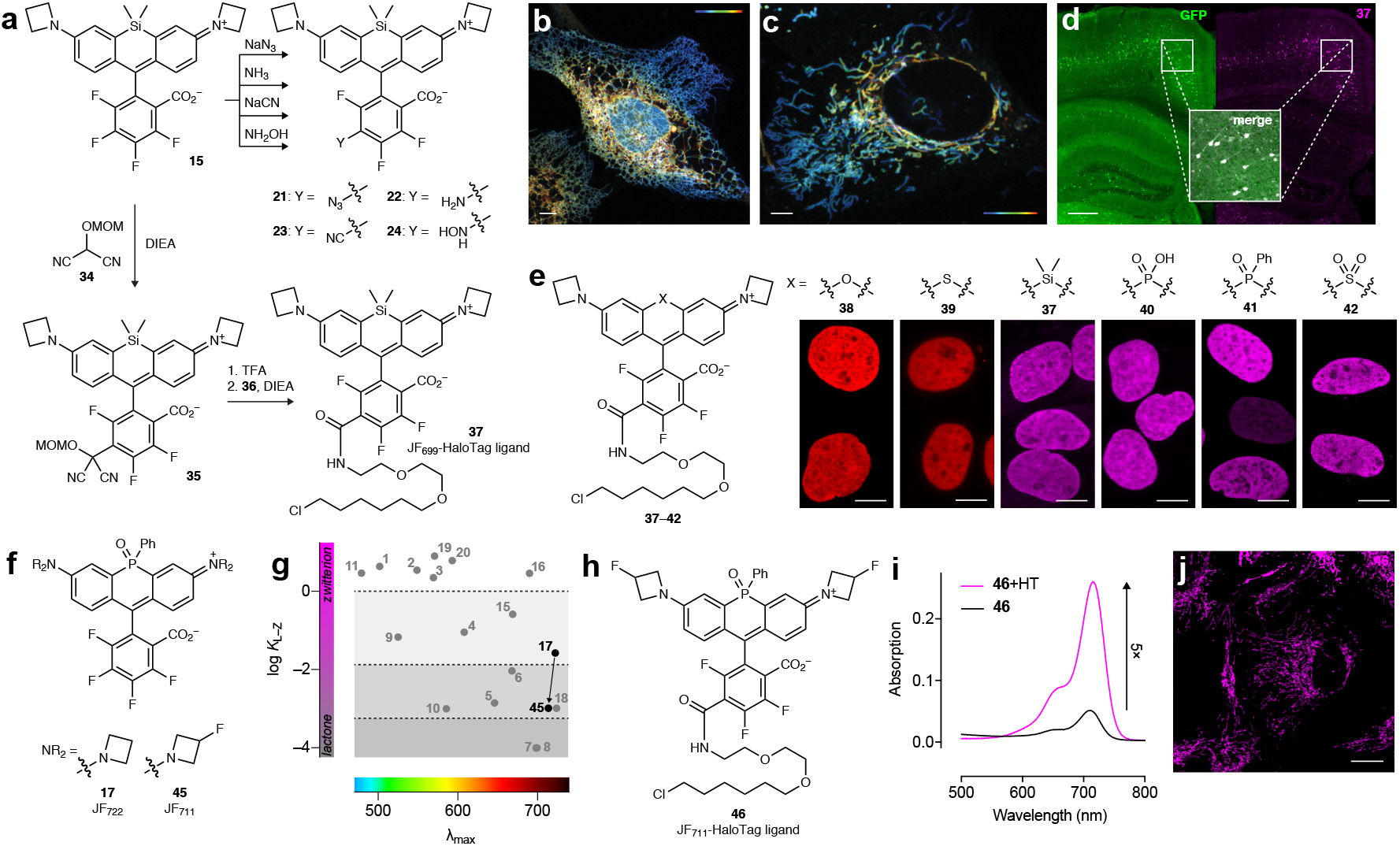
(**a**) Nucleophilic aromatic substitution (SNAr) of JF_669_ (**15**) to form derivatives **21**–**24** and **35**; subsequent synthesis of JF_669_–HaloTag ligand (**37**). (**b**) Airyscan image of U2OS cell expressing endoplasmic reticulum-localized Sec61β–HaloTag labeled with **37**. (**c**) Airyscan image of U2OS cell expressing mitochondria-localized TOMM20–HaloTag labeled with **37**; color scale in **b** and **c** indicates z-depth; scale bars: 5 μm. (**d**) Image of fixed coronal mouse brain slice from animal expressing GFP– HaloTag fusion protein in neurons after *intravenous* injection of **37**; scale bar = 500 μm. (**e**) Structures of HaloTag ligands **37**–**42** and high-magnification images of U2OS cell nuclei expressing histone H2B–HaloTag fusion protein and labeled **37**–**42**; scale bar = 10 μm. (**f**) Structures of dyes **17** and **45**. (**g**) Plot of *K*_L–Z_ *vs. λ*_max_ showing decreased *K*_L–Z_ for dye **45**. (**h**) Structure of JF_711_–HaloTag ligand (**46**). (**i**) Absorption spectra of **46** in the absence (-HT) or presence (+HT) of excess HaloTag protein. (**j**) Confocal imaging experiment of U2OS cells expressing TOMM20–HaloTag fusion protein labeled with **46**; scale bar = 20 μm.

We then sought to create carboxy derivatives compatible with different labeling strategies such as the HaloTag. We explored the reactivity of malonates and related carbon nucleophiles, all of which showed reaction with fluorinated rhodamines with regioselectivity at the 6-position (**Supplementary Note**). In particular, the addition of masked acyl cyanide^20^ reagent **34**, an *umpolung*-type acyl anion equivalent, to JF_669_ resulted in intermediate **35**, which could be deprotected to yield a reactive acyl cyanide suitable for conjugation with the HaloTag ligand amine (**36**) to form JF_669_–HaloTag ligand **37** (**Fig. 2a**). This late-stage, regioselective introduction of the carboxy handle has distinct advantages over previous rhodamine syntheses that generate isomeric mixtures^21^. Compound **37** was an excellent label for cell biological experiments (**Fig. 2b,c**) and was also blood–brain-barrier permeable, labeling HaloTag-expressing neurons throughout the mouse brain after *intravenous* injection (**Fig. 2d**, **Supplementary Fig. 3a**). This chemistry was generalizable across fluorinated rhodamines, allowing facile synthesis of HaloTag ligands **37**–**42** from dyes **15**–**20** with *λ*_max_ ranging from the green to NIR (**Supplementary Fig. 3b,c**, **Supplementary Note**). These compounds selectively labeled HaloTag fusions in cells, demonstrating that fluorination on the pendant phenyl ring does not prohibit HaloTag labeling (**Fig. 2e**).

Finally, we sought to create a bright, fluorogenic NIR HaloTag ligand suitable for biological imaging. The SO_2_-containing rhodamine JF_724_ (**18**) possessed a promising *K*_L–Z_ = 10^−3^ for creating fluorogenic compounds; the JF_724_–HaloTag ligand (**42**) showed a 15-fold increase upon reaction with HaloTag protein *in vitro* (**Supplementary Fig. 3d**). Nevertheless, this dye was plagued with a low *Φ* = 0.05 (**Table 1**), making it suboptimal for imaging experiments. In contrast, the P(O)Ph-containing fluorophore JF_722_ (**17**) exhibits a larger *Φ* = 0.11 but also a relatively high *K*L– Z = 0.026; the JF_722_–HaloTag ligand (**41**) was not fluorogenic (**Supplementary Fig. 3e**). We investigated whether our two tuning strategies could work synergistically, using JF_571_ (**19**, **Fig. 1m**) as a proof-of-concept. We introduced a fluorine substituent on each azetidine ring to create JF_559_ (**43**; **Supplementary Fig. 3f**) and found this dye exhibits *K*_L–Z_ = 6.22, intermediate between JF_549_ (**2**; *K*_L–Z_ = 3.5) and JF_571_ (**19**; *K*_L–Z_ = 7.93; **Table 1**, **Supplementary Fig. 3g–i**), demonstrating the compatibility of these strategies. The JF_559_–HaloTag ligand (**44**) could be used in live cell labeling (**Supplementary Fig. 3j,k**). We then applied this modification to JF_722_ by synthesizing derivatives with fluorine substituents on the azetidine ring, yielding JF_711_ (**45**, **Fig. 2f**, **Table 1**, **Supplementary Fig. 3l**). Compound **45** exhibited a further improvement in *Φ* = 0.17 (**Table 1**)and was predicted to yield fluorogenic ligands based on its *K*_L–Z_ = 10^−3^ (**Fig. 2g**). The JF_711_–HaloTag ligand **46** (**Fig. 2h**) showed a 5-fold increase upon binding HaloTag (**Fig. 2i**) with excellent performance in live-cell imaging experiments (**Fig. 2j**).

In summary, we expanded the palette of Janelia Fluor dyes by replacing the central oxygen in JF_549_ (**2**) with nitrogen (NCH3, **1**), sulfur (S and SO_2_, **3,8**), and phosphorous (PO_2_H and P(O)Ph, **6**–**7**; **Fig. 1c**). As the nitrogen-containing dye JF_502_ (**1**) exhibited relatively high *K*_L–Z_ and long *λ*_max_, we applied our established tuning strategy^4^ to transform this dye into the GFP-like JF_479_ (**11**; **Fig. 1e–i**). The NIR-excited dyes **7** and **8** exhibited the opposite problem with low *K*_L–Z_ values that rendered the compounds unusable in biological environments (**Fig. 1d**). We therefore established a complementary general method to increase both *λ*_max_ and *K*_L–Z_ by incorporating fluorines on the pendant phenyl ring of rhodamine dyes (**Fig. 1j,k**) followed by facile, generalizable SNAr chemistry to install groups for bioconjugation (**Fig. 2a**). This strategy yielded the bioavailable JF_669_–HaloTag ligand (**37**, **Fig. 2b**–**e**) along with other new fluorophores (**38**–**42**, **Fig. 2e**) and could be combined with our previous tuning method to create the fluorogenic NIR-excited JF_711_ HaloTag ligand (**46**, **Fig 2f**–**j**). Although we have focused here on HaloTag ligands and mammalian cells, we expect this general rubric relating *K*_L–Z_ to cellular performance (**Fig. 1b**) to be applicable to other ligands and biological systems^6^. We also anticipate this expanded fluorophore palette to enable the rational design of finely tuned labels for biological imaging experiments in cells or animals and the new derivatization chemistry to facilitate the synthesis of novel ligands, labels, stains, and indicators, particularly those excited by far-red or NIR light.

## ACKNOWLEDGEMENTS

We thank: S. Sternson (Janelia) for initial discussions on *umpolung* reagents; K. Svoboda (Janelia) for advice on *in vivo* labeling experiments; C. Deo and E. Schreiter (Janelia) for purified HaloTag protein, contributive discussions, and a critical reading of the manuscript; the Janelia Cell and Tissue Culture, Anatomy and Histology, and Vivarium teams for assistance with biological experiments. This work was supported by the Howard Hughes Medical Institute.

## AUTHOR CONTRIBUTIONS

L.D.L. and J.B.G. conceived the project. J.B.G. contributed organic synthesis and 1-photon spectroscopy measurements. A.N.T. and H.C. contributed cultured cell imaging experiments. B.M. contributed *in vivo* labeling and tissue imaging experiments. N.F. contributed organic synthesis. R.P. contributed 2-photon spectroscopy measurements. J.L-S and T.A.B. directed the project. L.D.L. directed the project and wrote the paper with input from the other authors.

---

**Supplementary Figure 1.**
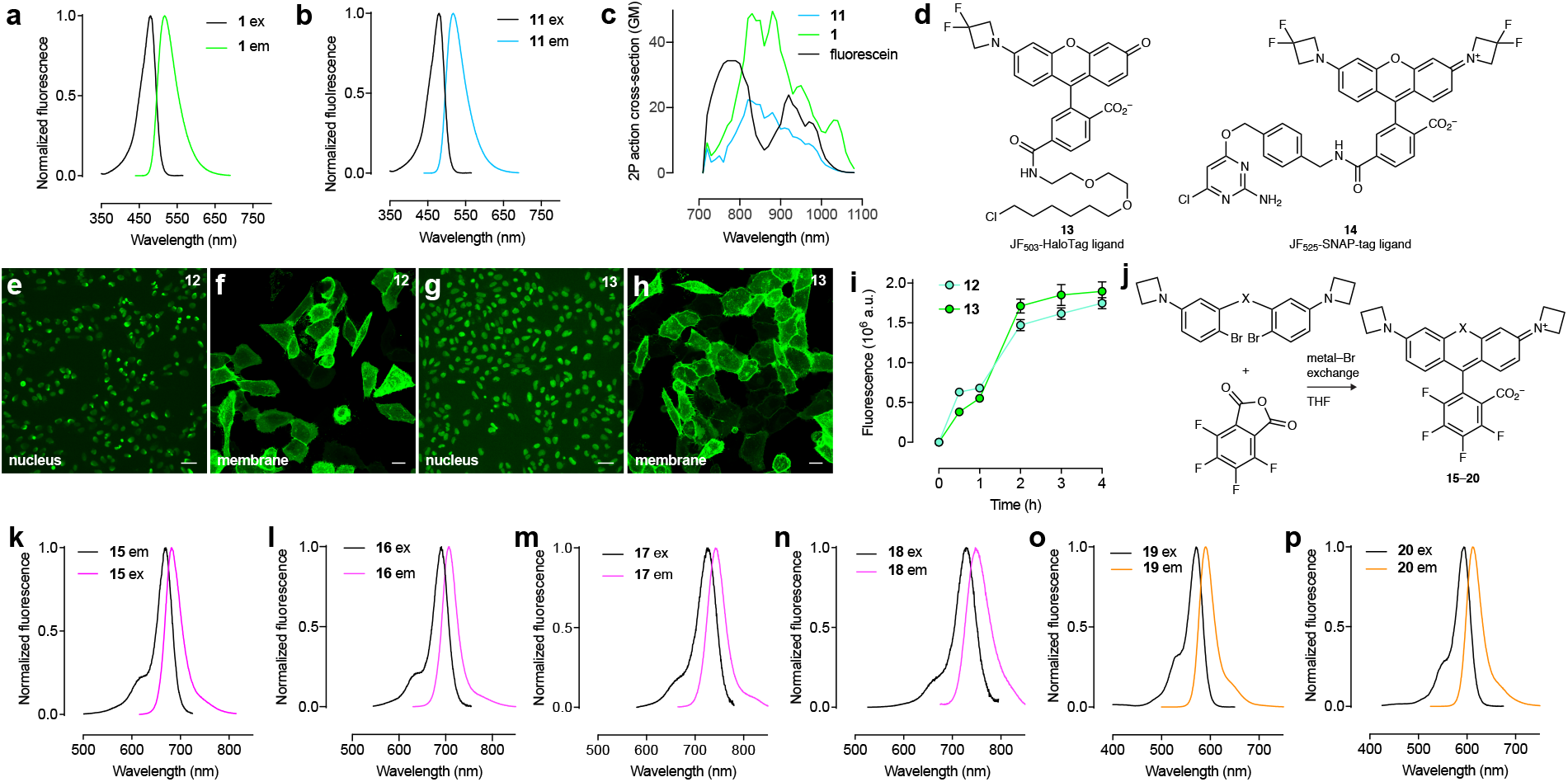
(**a**–**b**) Fluorescence excitation (ex) and emission (em) spectra of **1** (**a**) and **11** (**b**). (**c**) Two-photon absorption spectra of **1**, **11**, and fluorescein. (**d**) Structures of JF_503_–HaloTag ligand (**13**) and JF_525_–SNAP-tag ligand (**14**). (**e**–**h**) Widefield imaging experiment of U2OS cells expressing either histone H2B–HaloTag fusion protein (**e**,**g**; scale bars: 51 μm) and confocal imaging of PDGFR transmembrane domain (TMD)–HaloTag fusion protein (**f**,**h**; scale bars: 21 μm) labeled with JF_479_–HaloTag ligand (**12**; **e**,**f**) or JF_503_–HaloTag ligand (**13**; **g**,**h**) (**i**) Nuclear fluorescence *vs*. time upon addition of ligands **12** or **13** to cells expressing histone H2B–HaloTag fusion protein. (**j**) Synthesis of Janelia Fluor dyes **15**–**20**. (**k**–**p**) Fluorescence excitation (ex) and emission (em) spectra of **15** (**k**), **16** (**l**), **17** (**m**), **18** (**n**), **19** (**o**), and **20** (**p**).

**Supplementary Figure 2.**
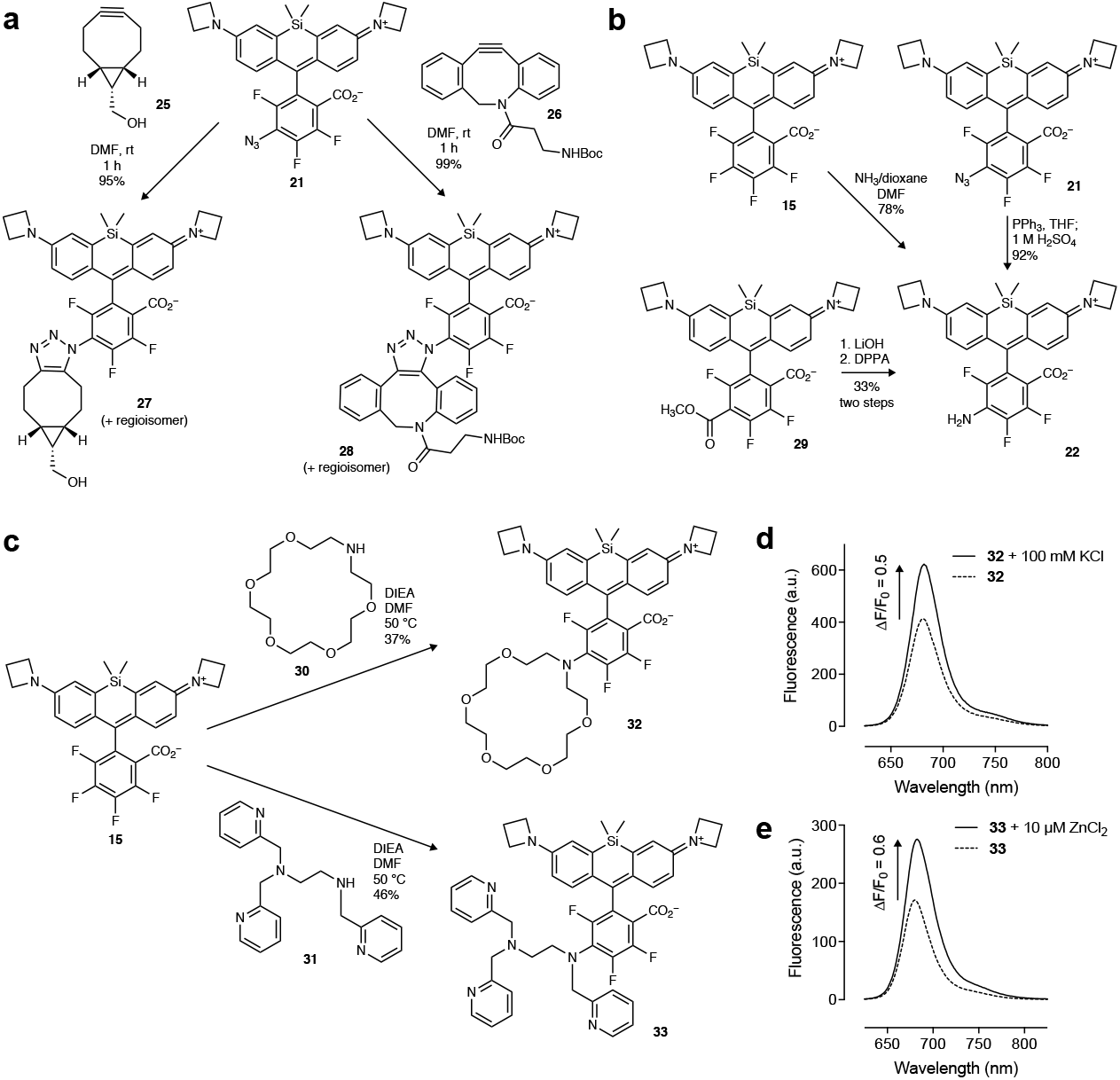
(**a**) Reaction of azide **21** with strained alkynes **25** or **26** to form triazole adducts **27** or **28**. (**b**) Synthesis of amine **22** via reaction of JF_669_ (**15**) with NH_3_, reduction of azide **21**, or Curtius rearrangement starting from ester **29** showing consistent regioselectivity of S_N_Ar. (**c**) Reaction of **15** with amine-containing chelator groups **30** and **31** to form far-red K^+^ indicator **32** and far-red Zn^2+^ indicator **33**. (**d**) Fluorescence emission spectra of **32** in the absence or presence of 100 mM K^+^. (**e**) Fluorescence emission spectra of **30** in the absence or presence of 10 μM Zn^2+^.

**Supplementary Figure 3.**
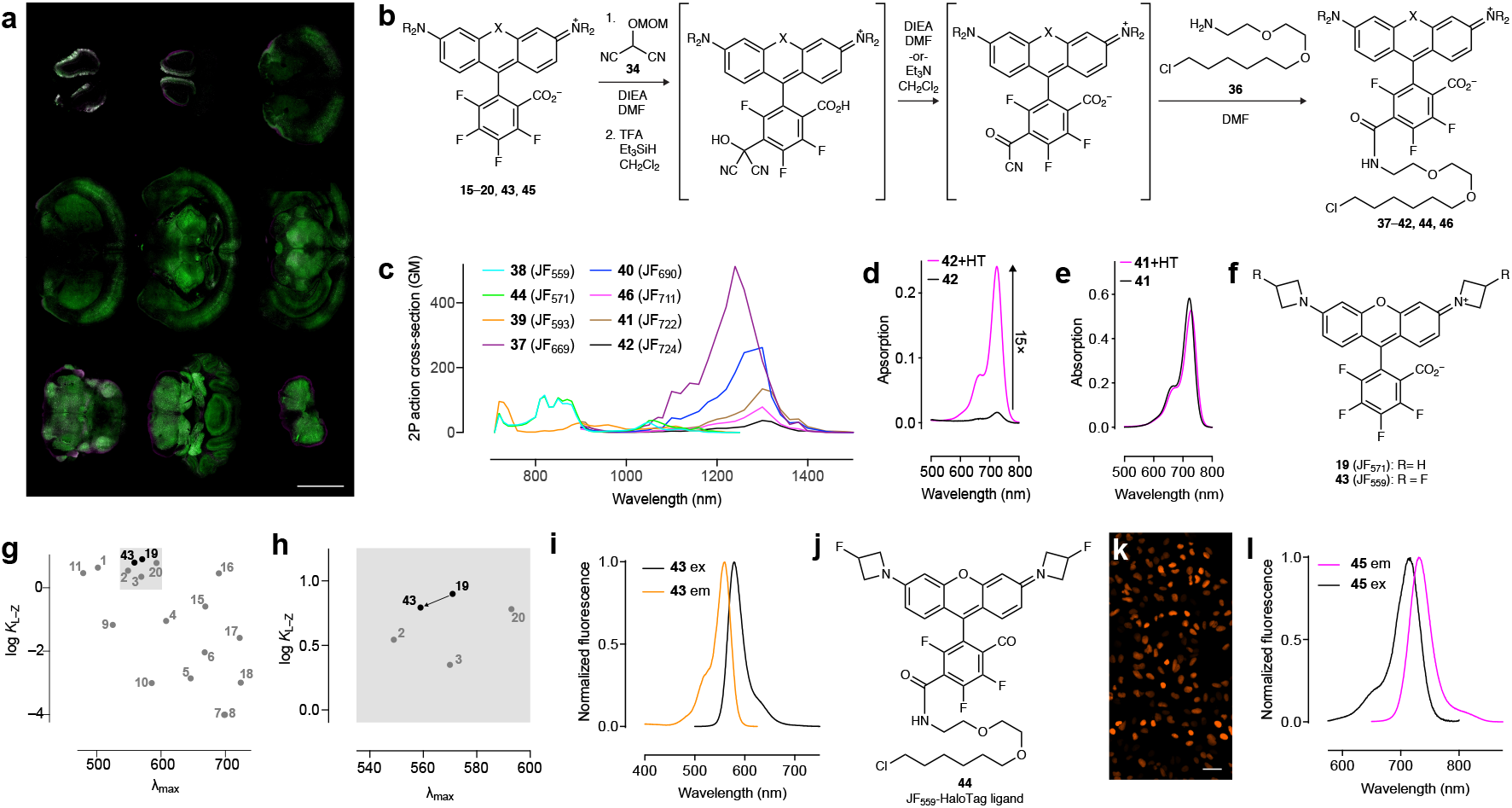
(**a**) Two-color mosaic image of fixed coronal slices from a mouse expressing HaloTag–GFP (green) in neurons transduced by *intravenous* injection of the viral vector PHP-eB-Syn- HaloTag–GFP followed by *intravenous* injection of **37** (magenta), perfusion, and slicing; scale bar = 3 mm. (**b**) Expanded synthetic scheme of HaloTag ligands **37**–**42**, **44** and **46** starting with nucleophilic aromatic substitution (SNAr) with **34**. (**c**) Two-photon absorption spectra of the HaloTag conjugates of HaloTag ligands **37**–**42**, **44** and **46**. (**d**) Absorption spectra of **42** in the absence (-HT) or presence (+HT) of excess HaloTag protein. (**e**) Absorption spectra of **41** in the absence (-HT) or presence (+HT) of excess HaloTag protein. (**f**) Structure of JF_571_ (**19**) and JF_559_ (**43**). (**g**–**h**) Full plot of *K*_L–Z_ *vs. λ*_max_ (**g**) and zoom (**h**) showing decreased *K*_L–Z_ for dye **43**. (**i**) Fluorescence excitation (ex) and emission (em) spectra of **43**. (**j**) Structure of JF_559_–HaloTag ligand (**44**). (**k**) Widefield imaging experiment of U2OS cells expressing histone H2B– HaloTag fusion protein labeled with **44**; scale bar: 51 μm. (**q**) Fluorescence excitation (ex) and emission (em) spectra of JF_711_ (**45**).

## ONLINE METHODS

### Chemical synthesis

Methods for chemical synthesis, full characterization of all novel compounds, and crystallographic confirmation of regioselective SNAr can be found in the Supplementary Note.

### UV–vis and fluorescence spectroscopy

Fluorescent and fluorogenic molecules for spectroscopy were prepared as stock solutions in DMSO and diluted such that the DMSO concentration did not exceed 1% v/v. Spectroscopy was performed using 1-cm path length, 3.5-mL quartz cuvettes or 1-cm path length, 1.0-mL quartz microcuvettes from Starna Cells. All measurements were taken at ambient temperature (22 ± 2 °C). Absorption spectra were recorded on a Cary Model 100 spectrometer (Agilent). Fluorescence spectra were recorded on a Cary Eclipse fluorometer (Varian). Maximum absorption wavelength (*λ*_abs_), extinction coefficient (*ε*), and maximum emission wavelength (*λ*_em_) were taken in 10 mM HEPES, pH 7.3 buffer unless otherwise noted; reported values for *ε* are averages (*n* = 3). Normalized spectra are shown for clarity. For prototype ion indicators **32** and **33** (**Supplementary Fig. 2c**) the compounds were dissolved in 10 mM HEPES, pH 7.3 buffer alone or with either 100 mM KCl or 10 μM ZnCl_2_; the fluorescence emission spectra of these solutions were recorded using *λ*_ex_ = 575 nm and *λ*_em_ = 625–825 nm.

### Determination *K*_L–Z_ and *ε*_max_

We calculated *K*_L–Z_ using the following equation^22^: *K*_L–Z_ = (*ε*_dw_/*ε*_max_)/(1 – *ε*_dw_/*ε*_max_). *ε*_dw_ is the extinction coefficient of the dyes in a 1:1 v/v dioxane:water solvent mixture; this dioxane–water mixture was chosen to give the maximum spread of *K*_L–Z_ values^4^. *ε*_max_ refers to the maximal extinction coefficients measured in different solvent mixtures empirically determined depending on dye type: 0.1% v/v TFA in ethanol for the Si-rhodamines (**5** and **15**); 0.1% v/v trifluoroacetic acid (TFA) in 2,2,2-trifluoroethanol (TFE) for all the other rhodamines. We note that accurate determination of low *K*_L–Z_ values (≤ 10^−3^) is complicated by the relatively poor sensitivity of absorbance measurements. We estimated *K*_L–Z_ = 10^−3^ when we observed a small but significant absorbance signal in 1:1 v/v dioxane:water solvent mixture over the dye-free control, and *K*_L–Z_ ≈10^−4^ when we observed no significant absorbance of the dye solution.

### Quantum yield determination

All reported absolute fluorescence quantum yield values (Φ) were measured in our laboratory under identical conditions using a Quantaurus-QY spectrometer (model C11374, Hamamatsu). This instrument uses an integrating sphere to determine photons absorbed and emitted by a sample. Measurements were carried out using dilute samples (*A* < 0.1) and self-absorption corrections^23^ were performed using the instrument software. Reported values are averages (*n* = 3).

### 1-Photon spectroscopy of HaloTag conjugates

HaloTag protein was used as a 100 μM solution in 75 mM NaCl, 50 mM TRIS·HCl, pH 7.4 with 50% v/v glycerol (TBS–glycerol). Absorbance measurements were performed in 1-mL quartz cuvettes. HaloTag ligands **12**, **41**–**42**, **46**(5 μM) were dissolved in 10 mM HEPES, pH 7.3 containing 0.1 mg·mL^−1^ CHAPS. An aliquot of HaloTag protein (1.5 equiv) was added and the resulting mixture was incubated until consistent absorbance signal was observed (60–120 min). To measure the fold-increase of absorbance upon HaloTag binding (**41**, **42**, **46**) an equivalent volume of TBS–glycerol blank was added in place of enzyme to record the ‘-HT’ absorbance. Absorbance scans are averages (*n* = 2).

### Multiphoton spectroscopy

For the environmentally insensitive compounds **1**, **11**, and a fluorescein control (**Supplementary Fig. 1c**) we prepared 5 μM solutions of the free dyes in 10 mM HEPES buffer, pH 7.3. For other rhodamines (**Supplementary Fig. 3c**), we measured spectra of the HaloTag conjugates: compounds **37**–**42**, **44**, and **46**(5 μM) were incubated with excess purified HaloTag protein (1.5 equiv) in 10 mM HEPES, pH 7.3 containing 0.1 mg·mL^−1^ CHAPS as above for 24 h at 4 °C. These solutions were then diluted to 1 μM in 10 mM HEPES buffer, pH 7.3 and the two-photon excitation spectra were measured as previously described^48,49^. Briefly, measurements were taken on an inverted microscope (IX81, Olympus) equipped with a 60×, 1.2NA water objective (Olympus). Dye–protein samples were excited with pulses from an 80 MHz Ti-Sapphire laser (Chameleon Ultra II, Coherent) for 710-1080 nm and with an OPO (Chameleon Compact OPO, Coherent) for 1000-1300 nm. Fluorescence collected by the objective was passed through a dichroic filter (675DCSXR, Omega) and a short pass filter (720SP, Semrock) and detected by a fiber-coupled Avalanche Photodiode (SPCM_AQRH-14, Perkin Elmer). For reference, a two-photon excitation spectrum was also obtained for the red fluorescent protein mCherry (1 μM), in the same HEPES buffer. All excitation spectra are corrected for the wavelength-dependent transmission of the dichroic and band-pass filters, and quantum efficiency of the detector.

### General cell culture and fluorescence microscopy

U2OS cells (ATCC) were cultured in Dulbecco’s modified Eagle medium (DMEM, phenol red-free; Life Technologies) supplemented with 10% (v/v) fetal bovine serum (Life Technologies), 1 mM GlutaMAX (Life Technologies) and maintained at 37 °C in a humidified 5% (v/v) CO_2_ environment. For confocal imaging of cell nuclei (**Fig. 1i**, **Fig. 2e**, **Supplementary Fig. 1e,g**, **Supplementary Fig. 3k**), we used U2OS cells with an integrated a histone H2B–HaloTag expressing plasmid via the piggyback transposase. For confocal imaging of the cell surface (**Supplementary Fig. 1f,h**), we used U2OS cells transiently transfected with a plasmid expressing a C-terminal transmembrane anchoring domain from platelet-derived growth factor receptor (PDGFR) fused to the HaloTag protein (PDGFR– HaloTag); nucleofection (Lonza) was used for transfection. For the Airyscan imaging experiments (**Fig. 2b,c**) we used U2OS cells transiently transfected with Sec61β–HaloTag expressing plasmid or TOMM20–HaloTag expressing plasmid using FuGENE HD (Promega). Sec61β is an endoplasmic reticulum membrane protein translocator protein and TOMM20 is an outer mitochondrial membrane protein as part of a protein translocase complex. For confocal imaging of mitochondria (**Fig. 2j**), we used U2OS cells with an integrated a TOMM20–HaloTag expressing plasmid via the piggyback transposase. These cell lines were kept under the selection of 500 μg/mL Geneticin (Life Technologies). Cell lines undergo regular mycoplasma testing by the Janelia Cell Culture Facility. Unless otherwise noted, cells were imaged on a Leica SP8 Falcon confocal microscope with an HC PL-APO 86×/1.20 water objective, a Zeiss LSM 800 confocal microscope with a Plan APO 20×/0.8 air M27 objective or Plan APO 63×/1.4 oil DIC M27 objective, a Zeiss LSM 880 with a C-APO 40×/1.2 W Corr FCS M27, or a Zeiss LSM 880 with Airyscan and a plan-apochromatic 63× oil objective (NA=1.4). The airyscan images were processed using the Zen software from Zeiss, the Leica and Zeiss LSM 800 confocal images were processed using FIJI.

### Dye loading kinetics

For the dye loading comparison (**Supplementary Fig. 1i**), U2OS cells stably expressing histone H2B–HaloTag were stained over a time course of 0–4 h with 200 nM of either JF_479_–HaloTag ligand **12** or JF_503_–HaloTag ligand **13**. Cells were washed 2× with dye-free media and imaged live using widefield microscopy (Nikon Eclipse Ti, Plan APO l 20×/0.75; 470nm Ex/FITC (515/30) Em. Fluorescence was quantified from the average of the summed intensity of nuclear signals in single plane widefield analyzed using Nikon NIS-Elements AR software (**Supplementary Fig. 1e,g**).

### Multiplexed imaging comparison JF_503_ and JF_479_

U2OS cells stably expressing histone H2B– HaloTag fusion protein were transiently transfected with TOMM20–pSNAPf plasmids using nucleofection (Lonza). Live cells were simultaneously incubated with JF_479_–HaloTag ligand (**12**, 500 nM) or JF_503_–HaloTag ligand (**13**, 500 nM) for 3 h, adding JF_525_–SNAP-tag ligand (**14**, 1 μM, **Supplementary Fig. 2d**) for 60 min. These cells were then washed and imaged (**Fig. 1i**) using tunable white light laser excitation at 488 nm and 532 nm on a Leica SP8 Falcon confocal microscope with an HC PL-APO 86×/1.20 water objective. The images are displayed as maximum intensity projections of confocal image stacks using FIJI.

### Mouse *in vivo* labeling experiments

Adult C57/BL6 male mice were used to express a GFP HaloTag fusion protein throughout the brain by systemic injection using the viral vector: PHP-eB-Syn-HaloTag-GFP (~5 × 10^11^ infectious units per ml, 100 μl). The virus was injected using a 0.5ml 27G syringe to the retro-orbital sinus. JF_669_–HaloTag ligand (**37**) was administered to mice 3–4 weeks after the viral injection. Dye solution was prepared by first dissolving 100 nmol (76 μg) of **37** in 20 μL DMSO. After vortexing, 20 μL of a Pluronic F-127 solution (20% w/w in DMSO) was added and this stock solution was diluted into 200 μL sterile saline for IV (retro-orbital) injection. All experimental protocols were conducted according to the National Institutes of Health guidelines for animal research and were approved by the Institutional Animal Care and Use Committee at the Janelia Research Campus, HHMI.

### Statistics

For spectroscopy measurements (**Fig. 1l**, **Table 1**, **Supplementary Fig. 1a,b,k–p, and Supplementary Fig. 1i,l**) reported n values for absorption spectra, extinction coefficient (*ε*) and quantum yield (Φ) represent measurements of different samples prepared from the same dye DMSO stock solution. For the cell loading experiment (**Supplementary Fig. 1i**) the following reported n values represent the number of intensity values measured from three fields of view for the time points at 30 s, 1 min, 2 min, 3 min, and 4 minutes, respectively: JF_479_–HaloTag ligand **12**: n= 112, 120, 130, 114, 128; JF_503_– HaloTag ligand **13**: n = 135, 129, 135, 158, 161.

### Data availability

The data that support the findings of this study are provided in the Source Data files or available from the corresponding author upon request.

