## Supplementary Note for "Optimization and functionalization of red-shifted rhodamine dyes"

### EXPERIMENTAL INFORMATION

| Page | Contents |
|---|---|
| S2 | Schemes S1–S4 |
| S6 | General Experimental Information for Synthesis |
| S7–S63 | Experimentals and Characterization Data for All Compounds |
| S7 | Preparation of Dibromide Intermediates |
| S22 | Rhodamine Synthesis via Metal-Bromide Exchange of Dibromides |
| S33 | Aza-Rhodamine HaloTag Ligands |
| S35 | MAC Substitution of 4,5,6,7-Tetrafluororhodamines |
| S41 | Conversion of MAC Substitution Products to Amides and Esters |
| S52 | Substitution of 4,5,6,7-Tetrafluororhodamines with Other Nucleophiles |
| S63 | Cycloadditions of 6-Azido-$JF_{669}$ |
| S64 | X-Ray Crystallography |
| S67 | NMR and HPLC/MS |

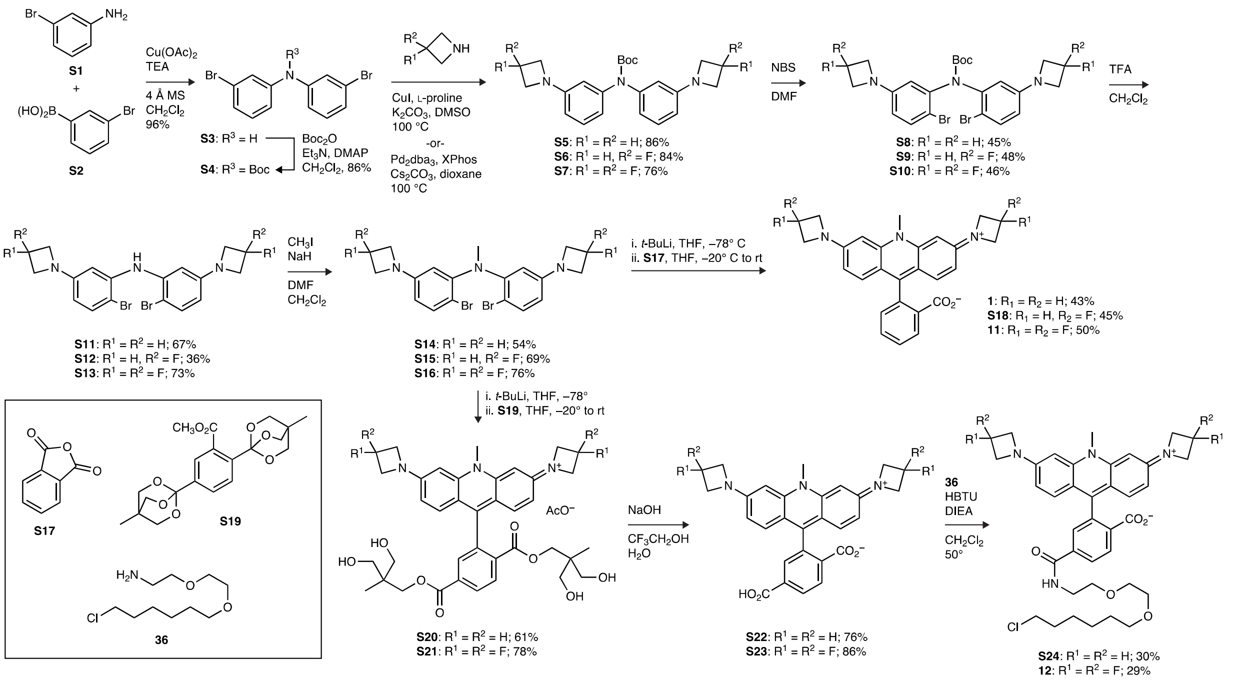


**Scheme S1.** Synthesis of the nitrogen rhodamines $JF_{502}$ (**1**), $JF_{479}$ (**11**), and their HaloTag ligand derivatives (**12**, **S24**).

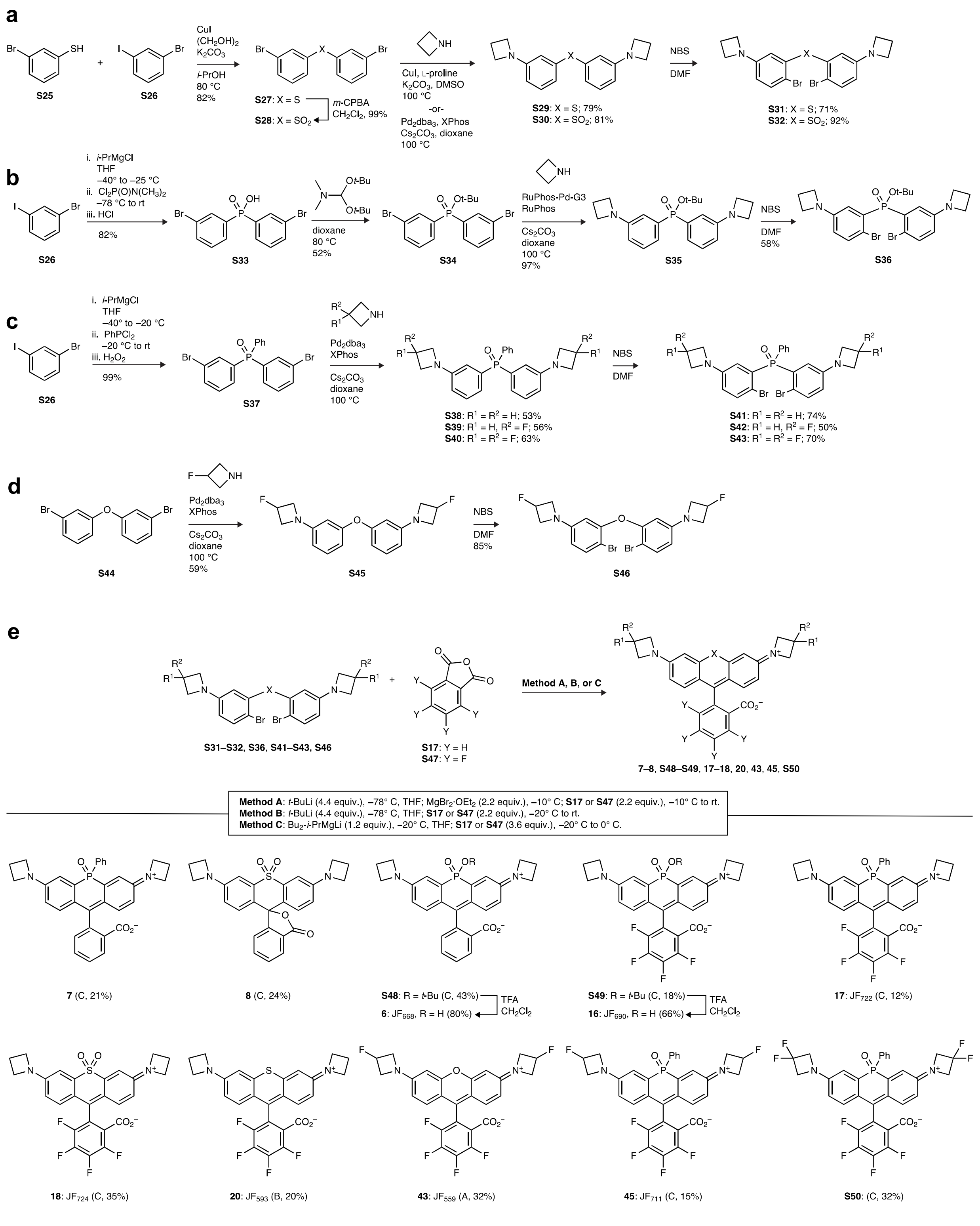


**Scheme S2.** (**a**) Synthesis of sulfide and sulfone dibromides **S31** and **S32**. (**b**) Synthesis of phosphinate dibromide **S36**. (**c**) Synthesis of phosphine oxide dibromides **S41**–**S43**. (**d**) Synthesis of ether dibromide **S46**. (**e**) Synthesis of heteroatom-containing rhodamines **7**–**8**, **S48**–**S49**, **17**–**18**, **20**, **43**, **45**, and **S50** via metal-bromide exchange of the dibromides described in (**a**–**d**) and addition to anhydrides **S17** or **S47**; TFA-mediated *tert*-butyl phosphinate deprotection of **S48** and **S49** yielded phosphinic acids **6** and **16**.

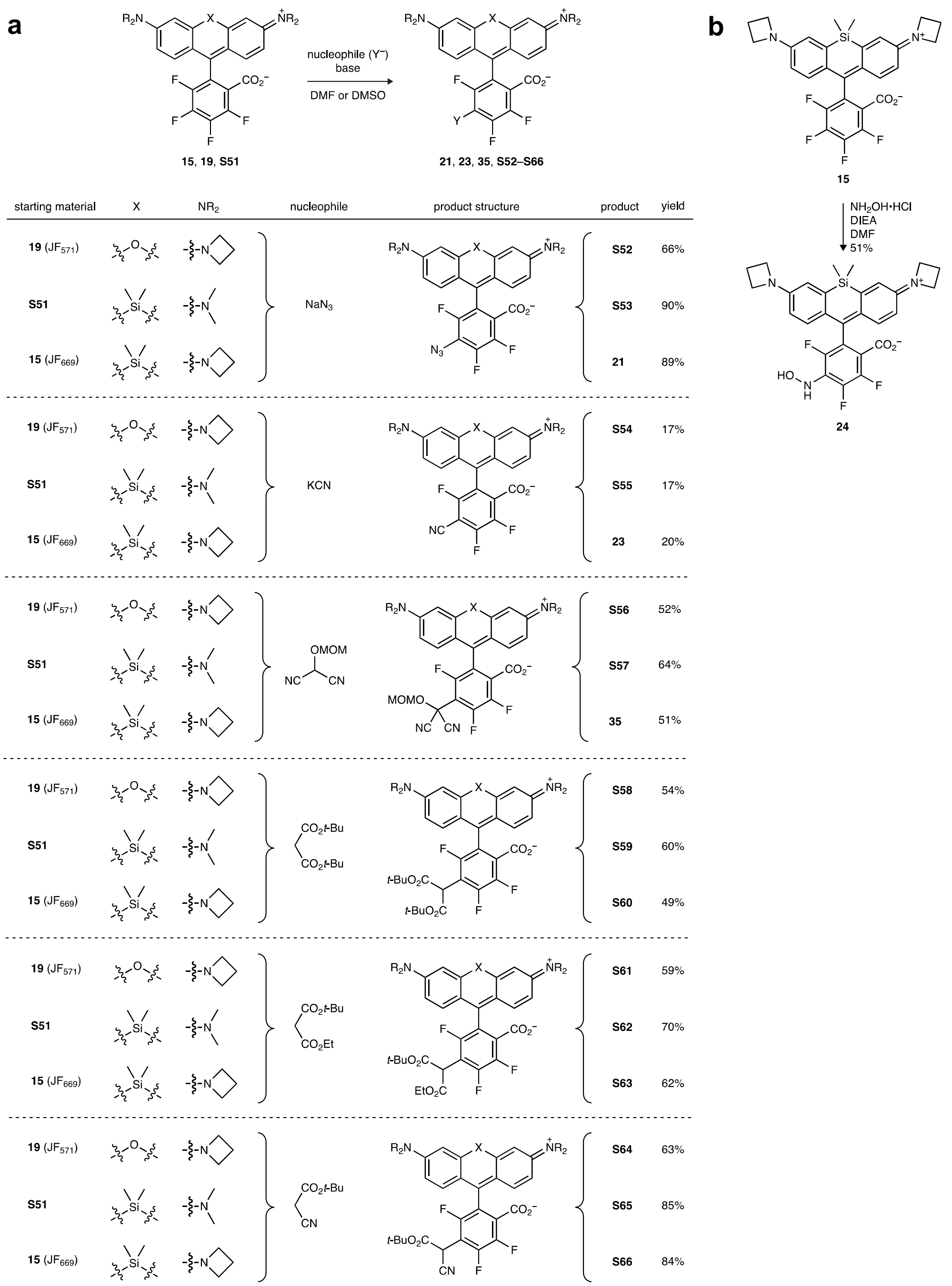


**Scheme S3.** (**a**) Nucleophilic substitution ($S_NAr$) reactions of 4,5,6,7-tetrafluororhodamines (**15**, **19**, and **S51**) with various nucleophiles. (**b**) Synthesis of hydroxylamine **24** from $JF_{669}$ (**15**).

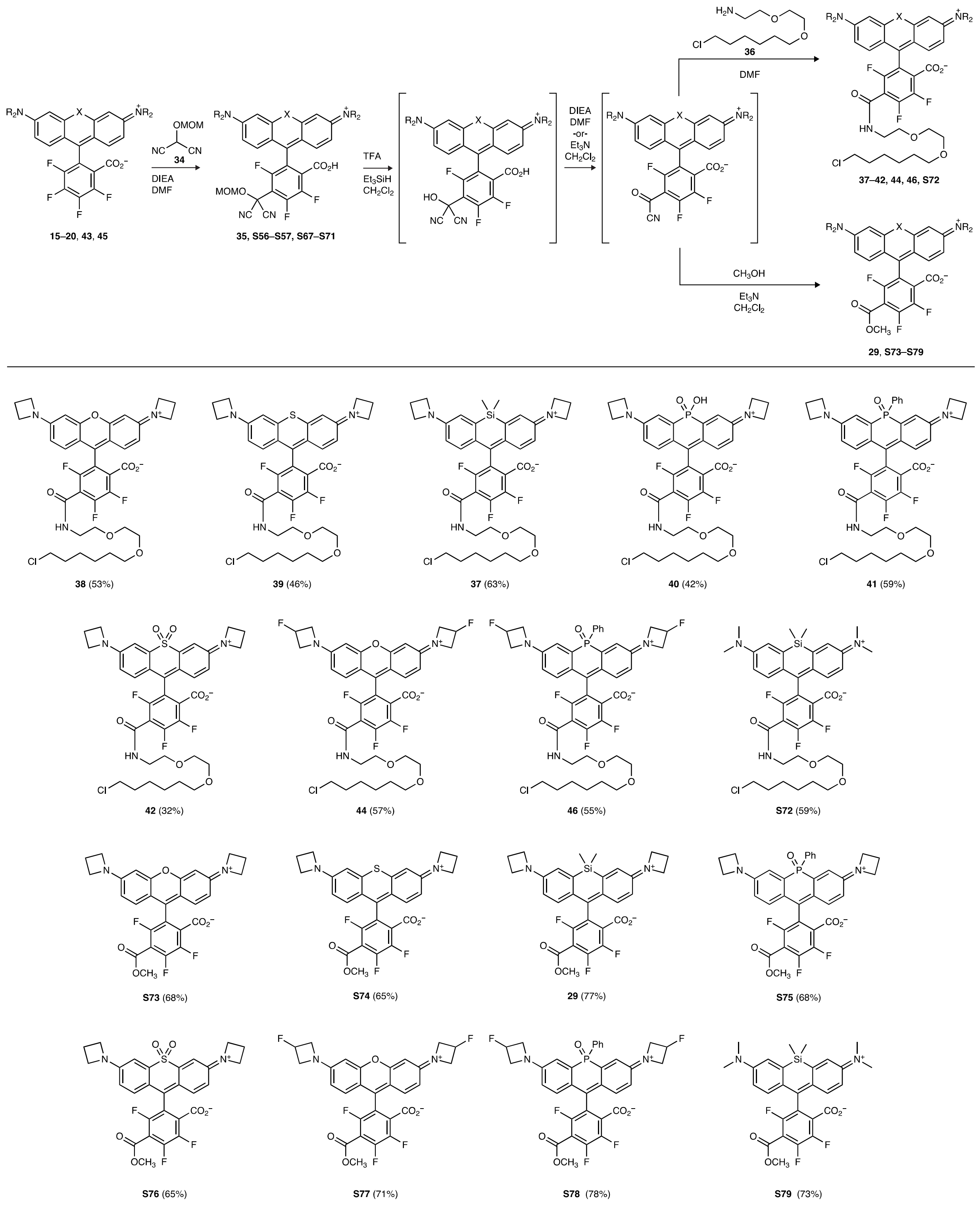


**Scheme S4.** Substitution of 4,5,6,7-tetrafluororhodamines with masked acyl cyanide (MAC) reagent **34** and subsequent derivatization to access HaloTag ligand labels and methyl ester analogs.

### General Experimental Information for Synthesis

Commercial reagents were obtained from reputable suppliers and used as received. All solvents were purchased in septum-sealed bottles stored under an inert atmosphere. All reactions were sealed with septa through which a nitrogen atmosphere was introduced unless otherwise noted. Reactions were conducted in round-bottomed flasks or septum-capped crimp-top vials containing Teflon-coated magnetic stir bars. Heating of reactions was accomplished with a silicon oil bath or an aluminum reaction block on top of a stirring hotplate equipped with an electronic contact thermometer to maintain the indicated temperatures.

Reactions were monitored by thin layer chromatography (TLC) on precoated TLC glass plates (silica gel 60 $F_{254}$, 250 μm thickness) or by LC/MS (Phenomenex Kinetex 2.1 mm × 30 mm 2.6 μm C18 column; 5 μL injection; 5–98% $MeCN/H_2O$, linear gradient, with constant 0.1% v/v $HCO_2H$ additive; 6 min run; 0.5 mL/min flow; ESI; positive ion mode). TLC chromatograms were visualized by UV illumination or developed with *p*-anisaldehyde, ceric ammonium molybdate, or $KMnO_4$ stain. Reaction products were purified by flash chromatography on an automated purification system using pre-packed silica gel columns or by preparative HPLC (Phenomenex Gemini–NX 30 × 150 mm 5 μm C18 column). Analytical HPLC analysis was performed with an Agilent Eclipse XDB 4.6 × 150 mm 5 μm C18 column under the indicated conditions. High-resolution mass spectrometry was performed by the High Resolution Mass Spectrometry Facility at the University of Iowa.

NMR spectra were recorded on a 400 MHz spectrometer. $^{1}H$ and $^{13}C$ chemical shifts were referenced to TMS or residual solvent peaks, and $^{19}F$ chemical shifts were referenced to $CFCl_3$. Data for $^{1}H$ NMR spectra are reported as follows: chemical shift (δ ppm), multiplicity (s = singlet, d = doublet, t = triplet, q = quartet, dd = doublet of doublets, m = multiplet), coupling constant (Hz), integration. Data for $^{13}C$ NMR spectra are reported by chemical shift (δ ppm) with hydrogen multiplicity (C, CH, $CH_2$, $CH_3$) information obtained from DEPT spectra. The $^{13}C$ NMR spectra are not reported for compounds containing trifluoro- or tetrafluoro-substituted aryl rings, as the numerous distinct fluorine couplings confounded interpretation of the spectra.

### PREPARATION OF DIBROMIDE INTERMEDIATES

We previously described a new strategy for the efficient synthesis of Si-fluoresceins and Si-rhodamines through metal–bromide exchange of bis(2-bromophenyl)silanes and double addition of the resulting bis(arylmetal) species to anhydrides.[1] Given the success of this approach, we sought to apply the same methodology to the synthesis of the rhodamines with heteroatom substitution at the 10-position described in this report (**Fig. 1c**). **Schemes S1**–**S2** depict the routes used to access the necessary dibromides. In each case, a bis(3-bromophenyl) species (**S3**, **Scheme S1**; **S27**, **S33**, **S37**, **Scheme S2**) was prepared through either a Chan–Lam coupling (nitrogen, **S3**; oxygen, **S44**),[1] an Ullmann-type coupling (sulfur, **S27**), or the addition of 3-bromophenyl Grignard to a phosphoryl dichloride (phosphinate, **S33**) or dichlorophosphine (phosphine oxide, **S37**). The azetidine substituents were then installed through Pd- or Cu-catalyzed C–N cross-coupling of these bis(3-bromophenyl) intermediates. Dibromination of the bis(3-azetidin-1-yl) cross-coupling products **S5**–**S7**, **S29**–**S30**, **S34**, **S38**–**S40**, and **S45** was achieved with good to excellent regioselectivity using NBS in DMF. For the sulfur, phosphorus, and oxygen analogs (**Scheme S2a–d**), this directly provided the desired dibromides (**S31**–**S32**, **S36**, **S41**–**S43**, and **S46**) for metal–bromide exchange. In the case of the nitrogen analogs (**Scheme S1**), acceptable regioselectivity in the NBS bromination was only achieved when the central nitrogen was acylated (*i.e.*, Boc-protected). Following bromination, the Boc group was removed and the nitrogen alkylated to afford the necessary dibromides **S14**–**S16**.

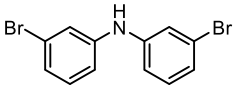


**Bis(3-bromophenyl)amine (S3):** An oven-dried round-bottom flask was charged with 3-bromophenylboronic acid (**S2**; 4.67 g, 23.3 mmol, 2 eq), $Cu(OAc)_2$ (2.11 g, 11.6 mmol, 1 eq), and 4 Å molecular sieves (10.00 g). After adding $CH_2Cl_2$ (50 mL), 3-bromoaniline (**S1**; 2.00 g, 11.6 mmol), and $Et_3N$ (3.24 mL, 23.3 mmol, 2 eq), the reaction was stirred at room temperature under ambient atmosphere for 48 h. The reaction mixture was filtered through Celite with $CH_2Cl_2$ and concentrated to dryness. The resultant residue was partitioned between saturated $NH_4Cl$ and $Et_2O$, and the aqueous layer was washed a second time with $Et_2O$. The combined organics were dried over anhydrous $MgSO_4$, filtered, and evaporated. Flash chromatography on silica gel (0–15% EtOAc/hexanes, linear gradient) provided dibromide **S3** as a yellow oil (3.64 g, 96%). $^1H$ NMR ($CDCl_3$, 400 MHz) δ 7.20 (t, $J$ = 2.0 Hz, 2H), 7.13 (t, $J$ = 7.9 Hz, 2H), 7.10 – 7.05 (m, 2H), 7.00 – 6.95 (m, 2H), 5.69 (s, 1H); $^{13}C$ NMR ($CDCl_3$, 101 MHz) δ 143.9 (C), 130.9 (CH), 124.7 (CH), 123.3 (C), 121.0 (CH), 116.8 (CH); HRMS (ESI) calcd for $C_{12}H_{10}Br_2N$ $[M+H]^+$ 325.9175, found 325.9188.

[1] Grimm, J. B.; Brown, T. A.; Tkachuk, A. N.; Lavis, L. D. *ACS Cent. Sci.* **2017**, *3*, 975–985.

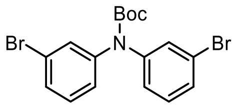


***tert*-Butyl bis(3-bromophenyl)carbamate (S4):** Aniline **S3** (3.35 g, 10.2 mmol) was taken up in $CH_2Cl_2$ (50 mL); $Et_3N$ (2.86 mL, 20.5 mmol, 2 eq), $Boc_2O$ (3.35 g, 15.4 mmol, 1.5 eq), and DMAP (125 mg, 1.02 mmol, 0.1 eq) were added, and the reaction was stirred at room temperature for 18 h. It was subsequently diluted with water and extracted with $CH_2Cl_2$ (2×). The combined organic extracts were washed with brine, dried over anhydrous $MgSO_4$, filtered, and concentrated *in vacuo*. Silica gel chromatography (0–5% $Et_2O$/hexanes, linear gradient) provided 3.78 g (86%) of carbamate **S4** as a pale yellow gum. $^1$H NMR ($CDCl_3$, 400 MHz) δ 7.37 (t, $J$ = 1.9 Hz, 2H), 7.34 (ddd, $J$ = 7.9, 1.9, 1.1 Hz, 2H), 7.20 (t, $J$ = 7.9 Hz, 2H), 7.12 (ddd, $J$ = 8.1, 2.1, 1.1 Hz, 2H), 1.45 (s, 9H); $^{13}$C NMR ($CDCl_3$, 101 MHz) δ 153.2 (C), 143.8 (C), 130.18 (CH), 130.16 (CH), 129.2 (CH), 125.8 (CH), 122.3 (C), 82.3 (C), 28.3 ($CH_3$); HRMS (ESI) calcd for $C_{17}H_{17}Br_2NO_2Na$ $[M+Na]^+$ 447.9518, found 447.9533.

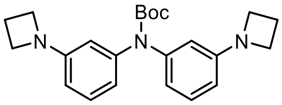


***tert*-Butyl bis(3-(azetidin-1-yl)phenyl)carbamate (S5):** An oven-dried round-bottom flask was charged with CuI (259 mg, 1.36 mmol, 0.2 eq), L-proline (313 mg, 2.72 mmol, 0.4 eq), and $K_2CO_3$ (3.75 g, 27.2 mmol, 4 eq). The flask was sealed and evacuated/backfilled with nitrogen (3×). A solution of dibromide **S4** (2.90 g, 6.79 mmol) in DMSO (27 mL) was added, and the reaction was flushed again with nitrogen (3×). Following the addition of azetidine (2.75 mL, 40.7 mmol, 6 eq), the reaction was stirred at 100 °C for 18 h. It was then cooled to room temperature, diluted with saturated $NH_4Cl$, and extracted with EtOAc (2×). The combined organic extracts were washed with brine, dried over anhydrous $MgSO_4$, filtered, and concentrated *in vacuo*. Purification by flash chromatography on silica gel (0–50% EtOAc/hexanes, linear gradient) afforded **S5** (2.21 g, 86%) as a yellow gum. $^1$H NMR ($CDCl_3$, 400 MHz) δ 7.09 (t, $J$ = 8.0 Hz, 2H), 6.55 (ddd, $J$ = 7.9, 2.0, 0.9 Hz, 2H), 6.31 (t, $J$ = 2.2 Hz, 2H), 6.24 (ddd, $J$ = 8.1, 2.3, 0.9 Hz, 2H), 3.82 (t, $J$ = 7.2 Hz, 8H), 2.32 (p, $J$ = 7.2 Hz, 4H), 1.45 (s, 9H); $^{13}$C NMR ($CDCl_3$, 101 MHz) δ 154.1 (C), 152.7 (C), 143.9 (C), 128.9 (CH), 116.2 (CH), 110.2 (CH), 108.8 (CH), 80.8 (C), 52.5 ($CH_2$), 28.5 ($CH_3$), 17.0 ($CH_2$); HRMS (ESI) calcd for $C_{23}H_{30}N_3O_2$ $[M+H]^+$ 380.2333, found 380.2333.

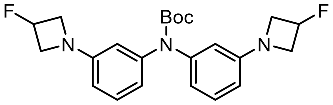


***tert*-Butyl bis(3-(3-fluoroazetidin-1-yl)phenyl)carbamate (S6):** An oven-dried round-bottom flask was charged with 3-fluoroazetidine hydrochloride (4.23 g, 37.9 mmol, 2.4 eq), $Pd_2dba_3$ (1.45 g, 1.58 mmol, 0.1 eq), XPhos (2.26 g, 4.74 mmol, 0.3 eq), and $Cs_2CO_3$ (24.72 g, 75.9 mmol, 4.8 eq). The flask was sealed and evacuated/backfilled with nitrogen (3×). A solution of dibromide **S4** (6.75 g, 15.8 mmol) in dioxane (80 mL) was added, and after flushing the reaction again with nitrogen (3×), it was stirred at 100 °C for 18 h. It was then cooled to room temperature, filtered through Celite with $CH_2Cl_2$, and concentrated to dryness. Purification by flash chromatography (0–50%

EtOAc/hexanes, linear gradient) provided **S6** as a yellow gum (5.50 g, 84%). $^1$H NMR ($CDCl_3$, 400 MHz) δ 7.13 (t, *J* = 8.0 Hz, 2H), 6.60 (ddd, *J* = 7.9, 1.9, 0.8 Hz, 2H), 6.34 (t, *J* = 2.1 Hz, 2H), 6.28 (ddd, *J* = 8.1, 2.3, 0.7 Hz, 2H), 5.38 (dtt, $^2J_{HF}$ = 57.0 Hz, *J* = 5.9, 3.7 Hz, 2H), 4.20 – 4.07 (m, 4H), 3.98 – 3.84 (m, 4H), 1.45 (s, 9H); $^{19}$F NMR ($CDCl_3$, 376 MHz) δ −180.65 (dtt, $J_{FH}$ = 57.1, 24.3, 18.1 Hz); $^{13}$C NMR ($CDCl_3$, 101 MHz) δ 153.9 (C), 151.6 (d, $^4J_{CF}$ = 1.2 Hz, C), 144.0 (C), 129.2 (CH), 117.1 (CH), 110.7 (CH), 109.4 (CH), 82.9 (d, $^1J_{CF}$ = 204.7 Hz, CHF), 81.1 (C), 59.7 (d, $^2J_{CF}$ = 23.5 Hz, $CH_2$), 28.5 ($CH_3$); HRMS (ESI) calcd for $C_{23}H_{27}F_2N_3O_2Na$ $[M+Na]^+$ 438.1964, found 438.1968.

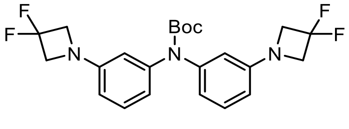


***tert*-Butyl bis(3-(3,3-difluoroazetidin-1-yl)phenyl)carbamate (S7):** An oven-dried round-bottom flask was charged with 3,3-difluoroazetidine hydrochloride (4.88 g, 37.7 mmol, 2.4 eq), $Pd_2dba_3$ (1.44 g, 1.57 mmol, 0.1 eq), XPhos (2.24 g, 4.71 mmol, 0.3 eq), and $Cs_2CO_3$ (24.53 g, 75.3 mmol, 4.8 eq). The flask was sealed and evacuated/backfilled with nitrogen (3×). A solution of dibromide **S4** (6.70 g, 15.7 mmol) in dioxane (80 mL) was added, and after flushing the reaction again with nitrogen (3×), it was stirred at 100 °C for 18 h. It was then cooled to room temperature, filtered through Celite with $CH_2Cl_2$, and concentrated to dryness. Purification by flash chromatography (0–50% $Et_2O$/hexanes, linear gradient) provided **S7** as a yellow solid (5.41 g, 76%). $^1$H NMR ($CDCl_3$, 400 MHz) δ 7.17 (t, *J* = 8.0 Hz, 2H), 6.66 (ddd, *J* = 8.0, 1.9, 0.7 Hz, 2H), 6.37 (t, *J* = 2.2 Hz, 2H), 6.32 (ddd, *J* = 8.1, 2.3, 0.7 Hz, 2H), 4.18 (t, $^3J_{HF}$ = 11.8 Hz, 8H), 1.45 (s, 9H); $^{19}$F NMR ($CDCl_3$, 376 MHz) δ −99.96 (p, $^3J_{FH}$ = 11.7 Hz); $^{13}$C NMR ($CDCl_3$, 101 MHz) δ 153.8 (C), 150.21 (t, $^4J_{CF}$ = 2.7 Hz, C), 144.0 (C), 129.4 (CH), 117.9 (CH), 116.0 (t, $^1J_{CF}$ = 274.6 Hz, $CF_2$), 111.1 (CH), 109.9 (CH), 81.3 (C), 63.5 (t, $^2J_{CF}$ = 25.7 Hz, $CH_2$), 28.5 ($CH_3$); HRMS (ESI) calcd for $C_{23}H_{25}F_4N_3O_2Na$ $[M+Na]^+$ 474.1775, found 474.1785.

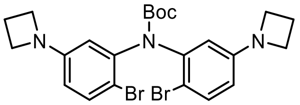


***tert*-Butyl bis(5-(azetidin-1-yl)-2-bromophenyl)carbamate (S8):** Carbamate **S5** (425 mg, 1.12 mmol) was taken up in DMF (10 mL). *N*-Bromosuccinimide (399 mg, 2.24 mmol, 2 eq) was added portion-wise over 1–2 min, and the reaction was then stirred at room temperature for 2 h. The reaction mixture was subsequently diluted with water and extracted with EtOAc (2×). The combined organic extracts were washed with water and brine, dried over anhydrous $MgSO_4$, filtered, and concentrated *in vacuo*. Silica gel chromatography (0–50% $Et_2O$/hexanes, linear gradient) afforded 269 mg (45%) of dibromide **S8** as a white solid. $^1$H NMR (DMSO-$d_6$, 400 MHz, 350 K) δ 7.41 (d, *J* = 8.7 Hz, 2H), 6.44 (d, *J* = 2.8 Hz, 2H), 6.27 (dd, *J* = 8.7, 2.8 Hz, 2H), 3.75 (t, *J* = 7.2 Hz, 8H), 2.28 (p, *J* = 7.2 Hz, 4H), 1.40 (s, 9H); $^{13}$C NMR (DMSO-$d_6$, 101 MHz, 350 K) δ 151.4 (C), 150.8 (C), 141.7 (C), 132.6 (CH), 111.3 (CH), 110.6 (CH), 108.1 (C), 79.9 (C), 51.4 ($CH_2$), 27.5 ($CH_3$), 15.8 ($CH_2$); HRMS (ESI) calcd for $C_{23}H_{28}Br_2N_3O_2$ $[M+H]^+$ 536.0543, found 536.0556.

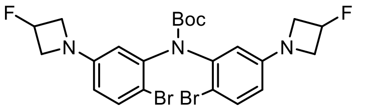


***tert*-Butyl bis(2-bromo-5-(3-fluoroazetidin-1-yl)phenyl)carbamate (S9):** Carbamate **S6** (5.35 g, 12.9 mmol) was taken up in DMF (130 mL) and cooled to 0 °C. *N*-Bromosuccinimide (4.58 g, 25.8 mmol, 2 eq) was added portion-wise over 10 min. The reaction was stirred at 0 °C for 30 min, then warmed to room temperature and stirred 1 h. The DMF was evaporated; the resulting residue was diluted with water and extracted with EtOAc (2×). The combined organic extracts were washed with water and brine, dried over anhydrous $MgSO_4$, filtered, and concentrated *in vacuo*. The crude was purified twice by silica gel chromatography (5–75% $Et_2O$/hexanes, linear gradient; then, 0–50% EtOAc/hexanes, linear gradient) to afford 3.51 g (48%) of dibromide **S9** as an off-white solid. $^1$H NMR (DMSO-$d_6$, 400 MHz, 350 K) δ 7.46 (d, $J$ = 8.6 Hz, 2H), 6.50 (d, $J$ = 2.8 Hz, 2H), 6.36 (dd, $J$ = 8.7, 2.8 Hz, 2H), 5.43 (dtt, $^2J_{HF}$ = 57.4 Hz, $J$ = 6.0, 3.2 Hz, 2H), 4.16 – 4.02 (m, 4H), 3.89 – 3.76 (m, 4H), 1.40 (s, 9H); $^{19}$F NMR (DMSO-$d_6$, 376 MHz, 350 K) δ −179.66 (dtt, $J_{FH}$ = 57.5, 23.8, 20.4 Hz).; $^{13}$C NMR (DMSO-$d_6$, 101 MHz, 350 K) δ 150.8 (C), 150.4 (d, $^4J_{CF}$ = 1.3 Hz, C), 141.7 (C), 132.8 (CH), 112.1 (CH), 111.3 (CH), 109.2 (C), 82.7 (d, $^1J_{CF}$ = 201.1 Hz, CHF), 80.1 (C), 58.7 (d, $^2J_{CF}$ = 23.7 Hz, $CH_2$), 27.5 ($CH_3$); HRMS (ESI) calcd for $C_{23}H_{25}Br_2F_2N_3O_2Na$ $[M+Na]^+$ 594.0174, found 594.0193.

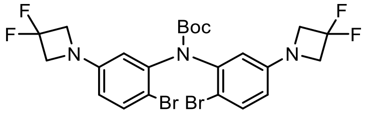


***tert*-Butyl bis(2-bromo-5-(3,3-difluoroazetidin-1-yl)phenyl)carbamate (S10):** Carbamate **S7** (5.50 g, 12.2 mmol) was taken up in DMF (120 mL) and cooled to 0 °C. *N*-Bromosuccinimide (4.34 g, 24.4 mmol, 2 eq) was added portion-wise over 10 min. The reaction was stirred at 0 °C for 30 min, then warmed to room temperature and stirred 1 h. The DMF was evaporated; the resulting residue was diluted with water and extracted with EtOAc (2×). The combined organic extracts were washed with water and brine, dried over anhydrous $MgSO_4$, filtered, and concentrated *in vacuo*. Purification by silica gel chromatography (0–40% EtOAc/hexanes, linear gradient) afforded 3.41 g (46%) of dibromide **S10** as a yellow foam. $^1$H NMR (DMSO-$d_6$, 400 MHz, 350 K) δ 7.52 (d, $J$ = 8.7 Hz, 2H), 6.58 (d, $J$ = 2.9 Hz, 2H), 6.47 (dd, $J$ = 8.7, 2.8 Hz, 2H), 4.21 (t, $^3J_{HF}$ = 12.1 Hz, 8H), 1.41 (s, 9H); $^{19}$F NMR (DMSO-$d_6$, 376 MHz, 350 K) δ −99.12 (p, $^3J_{FH}$ = 12.3 Hz); $^{13}$C NMR (DMSO-$d_6$, 101 MHz, 350 K) δ 150.8 (C), 149.2 (t, $^4J_{CF}$ = 3.1 Hz, C), 141.6 (C), 133.0 (CH), 115.8 (t, $^1J_{CF}$ = 273.5 Hz, $CF_2$), 113.0 (CH), 112.2 (CH), 110.5 (C), 80.3 (C), 62.4 (t, $^2J_{CF}$ = 25.7 Hz, $CH_2$), 27.5 ($CH_3$); HRMS (ESI) calcd for $C_{23}H_{23}Br_2F_4N_3O_2Na$ $[M+Na]^+$ 629.9985, found 629.9999.

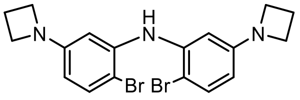


**Bis(5-(azetidin-1-yl)-2-bromophenyl)amine (S11):** Carbamate **S8** (2.10 g, 3.91 mmol) was taken up in $CH_2Cl_2$ (40 mL), and trifluoroacetic acid (8 mL) was added. The reaction was stirred at room temperature for 18 h. Toluene (40 mL) was added, and the reaction mixture was concentrated to dryness. The resulting residue was partitioned between saturated $NaHCO_3$ and $CH_2Cl_2$, and the aqueous layer was extracted a second time with $CH_2Cl_2$. The combined organic extracts were dried over anhydrous $MgSO_4$, filtered, deposited onto Celite, and concentrated *in vacuo*. The crude material was purified twice by flash chromatography on silica gel (0–50% EtOAc/hexanes, linear gradient; then, 0–40% EtOAc/hexanes, linear gradient, with 20% constant v/v $CH_2Cl_2$ additive; dry loaded in both instances with Celite) to provide 1.14 g (67%) of **S11** as a white solid. $^1$H NMR ($CDCl_3$, 400 MHz) δ 7.33 (d, $J$ = 8.6 Hz, 2H), 6.44 (d, $J$ = 2.6 Hz, 2H), 6.36 (s, 1H), 5.95 (dd, $J$ = 8.6, 2.7 Hz, 2H), 3.81 (t, $J$ = 7.2 Hz, 8H), 2.34 (p, $J$ = 7.2 Hz, 4H); $^{13}$C NMR ($CDCl_3$, 101 MHz) δ 152.2 (C), 140.2 (C), 133.2 (CH), 106.1 (CH), 101.6 (C), 100.9 (CH), 52.6 ($CH_2$), 16.9 ($CH_2$); HRMS (ESI) calcd for $C_{18}H_{20}Br_2N_3$ $[M+H]^+$ 436.0018, found 436.0033.

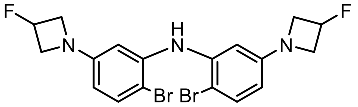


**Bis(2-bromo-5-(3-fluoroazetidin-1-yl)phenyl)amine (S12):** Carbamate **S9** (3.35 g, 5.84 mmol) was taken up in $CH_2Cl_2$ (50 mL), and trifluoroacetic acid (10 mL) was added. The reaction was stirred at room temperature for 18 h. Toluene (50 mL) was added, and the reaction mixture was concentrated to dryness. The resulting residue was partitioned between saturated $NaHCO_3$ and $CH_2Cl_2$, and the aqueous layer was extracted a second time with $CH_2Cl_2$. The combined organic extracts were dried over anhydrous $MgSO_4$, filtered, and concentrated *in vacuo*. The crude material was purified by flash chromatography on silica gel (0–40% EtOAc/hexanes, linear gradient, with 20% constant v/v $CH_2Cl_2$ additive) to provide 995 mg (36%) of **S12** as a white solid. $^1$H NMR ($CDCl_3$, 400 MHz) δ 7.38 (d, $J$ = 8.6 Hz, 2H), 6.43 (d, $J$ = 2.7 Hz, 2H), 6.35 (s, 1H), 6.00 (dd, $J$ = 8.6, 2.7 Hz, 2H), 5.39 (dtt, $^2J_{HF}$ = 56.8 Hz, $J$ = 5.8, 3.7 Hz, 2H), 4.19 – 4.05 (m, 4H), 3.96 – 3.82 (m, 4H); $^{19}$F NMR ($CDCl_3$, 376 MHz) δ −180.51 (dtt, $J_{FH}$ = 56.6, 23.8, 17.8 Hz); $^{13}$C NMR ($CDCl_3$, 101 MHz) δ 151.1 (d, $^4J_{CF}$ = 1.4 Hz, C), 140.3 (C), 133.4 (CH), 106.7 (CH), 102.7 (C), 101.4 (CH), 82.5 (d, $^1J_{CF}$ = 205.0 Hz, CHF), 59.7 (d, $^2J_{CF}$ = 23.7 Hz, $CH_2$); HRMS (ESI) calcd for $C_{18}H_{18}Br_2F_2N_3$ $[M+H]^+$ 471.9830, found 471.9850.

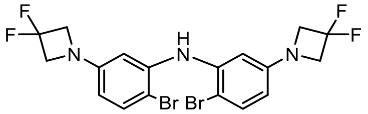


**Bis(2-bromo-5-(3,3-difluoroazetidin-1-yl)phenyl)amine (S13):** Carbamate **S10** (1.65 g, 2.71 mmol) was taken up in $CH_2Cl_2$ (25 mL), and trifluoroacetic acid (5 mL) was added. The reaction was stirred at room temperature for 6 h. Toluene (25 mL) was added, and the reaction mixture was concentrated to dryness. The resulting residue was partitioned between saturated $NaHCO_3$ and $CH_2Cl_2$, and the aqueous layer was extracted a second time with $CH_2Cl_2$. The combined organic extracts were washed with brine, dried over anhydrous $MgSO_4$, filtered, and concentrated *in vacuo*. Purification of the crude material by flash chromatography on silica gel (0–20% EtOAc/hexanes, linear gradient, with 20% constant v/v $CH_2Cl_2$ additive) yielded 1.00 g (73%) of **S13** as a colorless, viscous oil. $^1$H NMR ($CDCl_3$, 400 MHz) δ 7.41 (d, $J$ = 8.6 Hz, 2H), 6.39 (d, $J$ = 2.7 Hz, 2H), 6.34 (s, 1H), 6.03 (dd, $J$ = 8.6, 2.7 Hz, 2H), 4.15 (t, $^3J_{HF}$ = 11.7 Hz, 8H); $^{19}$F NMR ($CDCl_3$, 376 MHz) δ −99.94 (p, $^3J_{FH}$ = 11.7 Hz); $^{13}$C NMR ($CDCl_3$, 101 MHz)

δ 149.8 (t, $^4J_{CF}$ = 2.8 Hz, C), 140.5 (C), 133.7 (CH), 115.7 (t, $^1J_{CF}$ = 274.7 Hz, $CF_2$), 107.3 (CH), 103.8 (C), 102.1 (CH), 63.5 (t, $^2J_{CF}$ = 25.9 Hz, $CH_2$); HRMS (ESI) calcd for $C_{18}H_{16}Br_2F_4N_3$ $[M+H]^+$ 507.9642, found 507.9654.

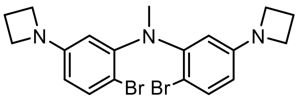

**5-(Azetidin-1-yl)-*N*-(5-(azetidin-1-yl)-2-bromophenyl)-2-bromo-*N*-methylaniline (S14):** Sodium hydride (60% dispersion in mineral oil, 210 mg, 5.26 mmol, 2 eq) was suspended in DMF (25 mL). A solution of aniline **S11** (1.15 g, 2.63 mmol) in $CH_2Cl_2$ (25 mL) was added dropwise, and the reaction was stirred at room temperature for 1 h. Iodomethane (491 μL, 7.89 mmol, 3 eq) was then added, and the reaction was stirred at room temperature for an additional 45 min. It was subsequently quenched with saturated $NH_4Cl$, diluted with water, and extracted with $CH_2Cl_2$ (2×). The combined organic extracts were washed with water and brine, dried over anhydrous $MgSO_4$, filtered, and evaporated. Flash chromatography on silica gel (0–25% EtOAc/hexanes, linear gradient, with 20% constant v/v $CH_2Cl_2$ additive) provided **S14** as a white solid (645 mg, 54%). $^1$H NMR ($CDCl_3$, 400 MHz) δ 7.32 – 7.28 (m, 2H), 6.06 – 6.01 (m, 4H), 3.80 (t, *J* = 7.2 Hz, 8H), 3.16 (s, 3H), 2.32 (p, *J* = 7.2 Hz, 4H); $^{13}$C NMR ($CDCl_3$, 101 MHz) δ 152.3 (C), 149.1 (C), 134.2 (CH), 108.0 (CH), 107.5 (C), 106.9 (CH), 52.6 ($CH_2$), 41.4 ($CH_3$), 16.9 ($CH_2$); HRMS (ESI) calcd for $C_{19}H_{22}Br_2N_3$ $[M+H]^+$ 450.0175, found 450.0177.

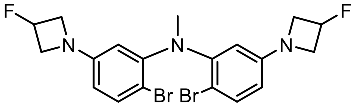

**2-Bromo-*N*-(2-bromo-5-(3-fluoroazetidin-1-yl)phenyl)-5-(3-fluoroazetidin-1-yl)-*N*-methylaniline (S15):** Sodium hydride (60% dispersion in mineral oil, 152 mg, 3.80 mmol, 2 eq) was suspended in DMF (15 mL). A solution of aniline **S12** (900 mg, 1.90 mmol) in $CH_2Cl_2$ (15 mL) was added dropwise, and the reaction was stirred at room temperature for 1 h. Iodomethane (355 μL, 5.71 mmol, 3 eq) was then added, and the reaction was stirred at room temperature for an additional 30 min. It was subsequently quenched with saturated $NH_4Cl$, diluted with water, and extracted with $CH_2Cl_2$ (2×). The combined organic extracts were washed with water and brine, dried over anhydrous $MgSO_4$, filtered, and evaporated. Flash chromatography on silica gel (0–20% EtOAc/hexanes, linear gradient, with 20% constant v/v $CH_2Cl_2$ additive) provided **S15** as a white solid (644 mg, 69%). $^1$H NMR ($CDCl_3$, 400 MHz) δ 7.34 (d, *J* = 8.2 Hz, 2H), 6.11 – 6.04 (m, 4H), 5.38 (dtt, $^2J_{HF}$ = 57.0 Hz, *J* = 6.1, 3.7 Hz, 2H), 4.18 – 4.06 (m, 4H), 3.96 – 3.83 (m, 4H), 3.17 (s, 3H); $^{19}$F NMR ($CDCl_3$, 376 MHz) δ −180.68 (dtt, $J_{FH}$ = 57.1, 23.9, 18.1 Hz); $^{13}$C NMR ($CDCl_3$, 101 MHz) δ 151.2 (C), 149.1 (C), 134.5 (CH), 108.54 (CH), 108.46 (C), 107.3 (CH), 82.6 (d, $^1J_{CF}$ = 204.7 Hz, CHF), 59.7 (d, $^2J_{CF}$ = 23.7 Hz, $CH_2$), 41.4 ($CH_3$); HRMS (ESI) calcd for $C_{19}H_{20}Br_2F_2N_3$ $[M+H]^+$ 485.9987, found 485.9999.

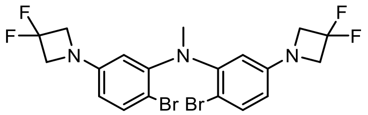


**2-Bromo-*N*-(2-bromo-5-(3,3-difluoroazetidin-1-yl)phenyl)-5-(3,3-difluoroazetidin-1-yl)-*N*-methylaniline (S16):** Sodium hydride (60% dispersion in mineral oil, 220 mg, 5.50 mmol, 2 eq) was suspended in DMF (25 mL). A solution of aniline **S13** (1.40 g, 2.75 mmol) in $CH_2Cl_2$ (25 mL) was added dropwise, and the reaction was stirred at room temperature for 1 h. Iodomethane (514 μL, 8.25 mmol, 3 eq) was then added, and the reaction was stirred at room temperature for an additional 1 h. It was subsequently quenched with saturated $NH_4Cl$, diluted with water, and extracted with $CH_2Cl_2$ (2×). The combined organic extracts were dried over anhydrous $MgSO_4$, filtered, and evaporated. Silica gel chromatography (0–20% EtOAc/hexanes, linear gradient) provided **S16** as a white solid (1.09 g, 76%). $^1$H NMR ($CDCl_3$, 400 MHz) δ 7.37 (d, $J$ = 8.4 Hz, 2H), 6.10 (dd, $J$ = 8.4, 2.8 Hz, 2H), 6.07 (d, $J$ = 2.7 Hz, 2H), 4.15 (t, $^3J_{HF}$ = 11.7 Hz, 8H), 3.18 (s, 3H); $^{19}$F NMR ($CDCl_3$, 376 MHz) δ −100.02 (p, $^3J_{FH}$ = 11.7 Hz); $^{13}$C NMR ($CDCl_3$, 101 MHz) δ 149.9 (t, $^4J_{CF}$ = 2.8 Hz, C), 149.1 (C), 134.7 (CH), 115.8 (t, $^1J_{CF}$ = 274.5 Hz, $CF_2$), 109.3 (C), 109.0 (CH), 107.8 (CH), 63.5 (t, $^2J_{CF}$ = 25.9 Hz, $CH_2$), 41.4 ($CH_3$); HRMS (ESI) calcd for $C_{19}H_{18}Br_2F_4N_3$ $[M+H]^+$ 521.9798, found 521.9809.

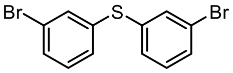


**Bis(3-bromophenyl)sulfane (S27):** An oven-dried round-bottom flask was charged with CuI (604 mg, 3.17 mmol, 0.1 eq) and $K_2CO_3$ (8.77 g, 63.5 mmol, 2 eq). The flask was sealed and evacuated/backfilled with nitrogen (3×). Isopropanol (125 mL) was added, followed by ethylene glycol (3.54 mL, 63.5 mmol, 2 eq), 3-bromothiophenol (**S25**; 3.28 mL, 31.7 mmol), and 3-bromoiodobenzene (**S26**; 4.45 mL, 34.9 mmol, 1.1 eq). The reaction mixture was stirred at 80 °C for 18 h. It was then diluted with saturated $NH_4Cl$ (200 mL) and EtOAc (200 mL), vigorously stirred for 30 min, and filtered through Celite. The filtrate was separated, and the aqueous layer was extracted again with EtOAc. The combined organics were washed with brine, dried over anhydrous $MgSO_4$, filtered, and evaporated. Flash chromatography (100% hexanes, linear gradient) afforded 8.93 g (82%) of dibromide **S27** as a colorless oil. $^1$H NMR ($CDCl_3$, 400 MHz) δ 7.48 (t, $J$ = 1.8 Hz, 2H), 7.40 (ddd, $J$ = 7.8, 1.9, 1.2 Hz, 2H), 7.28 – 7.23 (m, 2H), 7.18 (t, $J$ = 7.8 Hz, 2H); $^{13}$C NMR ($CDCl_3$, 101 MHz) δ 137.3 (C), 133.8 (CH), 130.79 (CH), 130.75 (CH), 129.8 (CH), 123.3 (C); HRMS (EI) calcd for $C_{12}H_8Br_2S$ $[M]^{+\bullet}$ 341.8708, found 341.8732.

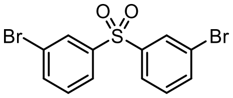


**3,3'-Sulfonylbis(bromobenzene) (S28):** To a solution of sulfide **S27** (6.45 g, 18.7 mmol) in $CH_2Cl_2$ (100 mL) was added *m*-CPBA (9.71 g, 56.2 mmol, 3 eq). After stirring the reaction at room temperature for 4 h, it was diluted with 10% $Na_2S_2O_3$ and extracted with $CH_2Cl_2$ (2×). The combined organic extracts were washed with 10% $Na_2S_2O_3$, saturated $NaHCO_3$, and brine, dried over anhydrous $MgSO_4$, filtered, and evaporated to provide sulfone **S28** as a white solid (7.01 g, 99%). Analytical HPLC and NMR indicated that the material was >95% pure and did not require further purification prior to cross-coupling with azetidine. $^{1}$H NMR ($CDCl_3$, 400 MHz) δ 8.08 (t, *J* = 1.9 Hz, 2H), 7.87 (ddd, *J* = 7.9, 1.8, 1.0 Hz, 2H), 7.72 (ddd, *J* = 8.0, 1.9, 1.0 Hz, 2H), 7.41 (t, *J* = 7.9 Hz, 2H); $^{13}$C NMR ($CDCl_3$, 101 MHz) δ 142.9 (C), 136.8 (CH), 131.1 (CH), 130.8 (CH), 126.5 (CH), 123.6 (C); HRMS (ESI) calcd for $C_{12}H_8Br_2O_2SNa$ $[M+Na]^+$ 396.8504, found 396.8514.

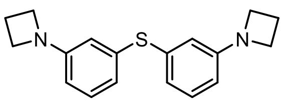


**Bis(3-(azetidin-1-yl)phenyl)sulfane (S29):** An oven-dried round-bottom flask was charged with CuI (985 mg, 5.17 mmol, 0.2 eq), L-proline (1.19 g, 10.4 mmol, 0.4 eq), and $K_2CO_3$ (14.30 g, 103.5 mmol, 4 eq). The flask was sealed and evacuated/backfilled with nitrogen (3×). A solution of dibromide **S27** (8.90 g, 25.9 mmol) in DMSO (100 mL) was added, and the reaction was flushed again with nitrogen (3×). Following the addition of azetidine (10.46 mL, 155.2 mmol, 6 eq), the reaction was stirred at 100 °C for 18 h. It was then cooled to room temperature, diluted with saturated $NH_4Cl$, and extracted with EtOAc (2×). The combined organic extracts were washed with water and brine, dried over anhydrous $MgSO_4$, filtered, and concentrated *in vacuo*. Purification by flash chromatography on silica gel (0–30% EtOAc/hexanes, linear gradient) afforded **S29** (6.08 g, 79%) as a white solid. $^{1}$H NMR ($CDCl_3$, 400 MHz) δ 7.11 (t, *J* = 7.9 Hz, 2H), 6.68 (ddd, *J* = 7.7, 1.7, 1.0 Hz, 2H), 6.45 (t, *J* = 2.0 Hz, 2H), 6.30 (ddd, *J* = 8.1, 2.3, 0.9 Hz, 2H), 3.83 (t, *J* = 7.2 Hz, 8H), 2.33 (p, *J* = 7.3 Hz, 4H); $^{13}$C NMR ($CDCl_3$, 101 MHz) δ 152.8 (C), 136.3 (C), 129.5 (CH), 119.9 (CH), 113.6 (CH), 110.1 (CH), 52.5 ($CH_2$), 17.1 ($CH_2$); HRMS (ESI) calcd for $C_{18}H_{21}N_2S$ $[M+H]^+$ 297.1420, found 297.1428.

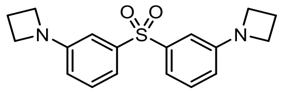


**1,1'-(Sulfonylbis(3,1-phenylene))bis(azetidine) (S30):** An oven-dried round-bottom flask was charged with sulfone **S28** (2.80 g, 7.45 mmol), $Pd_2dba_3$ (682 mg, 0.745 mmol, 0.1 eq), XPhos (1.06 g, 2.23 mmol, 0.3 eq), and $Cs_2CO_3$ (6.79 g, 20.9 mmol, 2.8 eq). The flask was sealed and evacuated/backfilled with nitrogen (3×). Dioxane (35 mL) was added, and the reaction was flushed again with nitrogen (3×). Following the addition of azetidine (1.51 mL, 22.3 mmol, 3 eq), the reaction was stirred at 100 °C for 4 h. It was then cooled to room temperature, filtered through Celite with $CH_2Cl_2$, and concentrated to dryness. The resulting residue was purified by flash chromatography (0–40% EtOAc/hexanes, linear gradient, with constant 40% v/v $CH_2Cl_2$ additive) to provide a brown-orange solid. The solid was triturated with $Et_2O$, sonicated, and filtered; the filter cake was washed with additional $Et_2O$ and dried to yield 1.97 g (81%) of **S30** as an off-white solid. $^{1}$H NMR ($CDCl_3$, 400 MHz) δ 7.26 (t, *J* = 7.9 Hz, 2H), 7.22 – 7.16 (m, 2H), 6.94 (t, *J* = 2.1 Hz, 2H), 6.52 (ddd, *J* = 8.0, 2.3, 1.0 Hz, 2H), 3.90 (t, *J* = 7.3 Hz, 8H), 2.38 (p, *J* = 7.2 Hz, 4H); $^{13}$C NMR ($CDCl_3$, 101 MHz) δ 152.2 (C), 142.4 (C), 129.7 (CH), 115.9 (CH), 115.3 (CH), 109.4 (CH), 52.4 ($CH_2$), 16.9 ($CH_2$); HRMS (ESI) calcd for $C_{18}H_{20}N_2O_2SNa$ $[M+Na]^+$ 351.1138, found 351.1137.

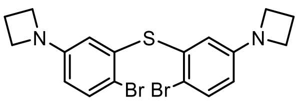

**Bis(5-(azetidin-1-yl)-2-bromophenyl)sulfane (S31):** Sulfide **S29** (6.00 g, 20.2 mmol) was taken up in DMF (100 mL). *N*-Bromosuccinimide (7.20 g, 40.5 mmol, 2 eq) was added portion-wise over 5 min, and the reaction was then stirred at room temperature for 2 h. The reaction mixture was concentrated *in vacuo*; the resulting residue was diluted with water and extracted with EtOAc (2×). The combined organic extracts were washed with water and brine, dried over anhydrous $MgSO_4$, filtered, and concentrated *in vacuo*. The crude product was triturated with $Et_2O$, sonicated, and filtered. The filter cake was washed with $Et_2O$ and dried to provide the title compound as a white solid. The filtrate was concentrated, chromatographed on silica gel (0–50% $Et_2O$/hexanes, linear gradient), and triturated as before to yield additional dibromide product. The two crops of white powder were combined, affording 6.51 g (71%) of dibromide **S31**. $^1$H NMR ($CDCl_3$, 400 MHz) δ 7.40 – 7.34 (m, 2H), 6.23 – 6.18 (m, 4H), 3.76 (t, *J* = 7.3 Hz, 8H), 2.31 (p, *J* = 7.2 Hz, 4H); $^{13}$C NMR ($CDCl_3$, 101 MHz) δ 151.9 (C), 135.6 (C), 133.3 (CH), 115.0 (CH), 112.3 (C), 112.1 (CH), 52.4 ($CH_2$), 16.9 ($CH_2$); HRMS (ESI) calcd for $C_{18}H_{19}Br_2N_2S$ $[M+H]^+$ 452.9630, found 452.9632.

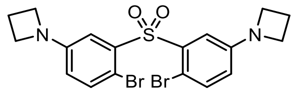

**1,1'-(Sulfonylbis(4-bromo-3,1-phenylene))bis(azetidine) (S32):** Sulfone **S30** (1.82 g, 5.54 mmol) was taken up in DMF (200 mL). *N*-Bromosuccinimide (1.97 g, 11.1 mmol, 2 eq) was added portion-wise over 10 min, and the reaction was then stirred at room temperature for 72 h. The reaction mixture was concentrated *in vacuo*; the resulting residue was diluted with saturated $NaHCO_3$ and extracted with $CH_2Cl_2$ (2×). The combined organic extracts were washed with brine, dried over anhydrous $MgSO_4$, filtered, and evaporated. Silica gel chromatography (0–10% EtOAc/toluene, linear gradient) afforded 2.47 g (92%) of dibromide **S32** as a white solid. $^1$H NMR ($CDCl_3$, 400 MHz) δ 7.51 (d, *J* = 2.9 Hz, 2H), 7.35 (d, *J* = 8.6 Hz, 2H), 6.42 (dd, *J* = 8.6, 2.9 Hz, 2H), 3.95 (t, *J* = 7.3 Hz, 8H), 2.42 (p, *J* = 7.3 Hz, 4H); $^{13}$C NMR ($CDCl_3$, 101 MHz) δ 150.8 (C), 139.1 (C), 135.3 (CH), 116.8 (CH), 115.9 (CH), 105.8 (C), 52.6 ($CH_2$), 16.9 ($CH_2$); HRMS (ESI) calcd for $C_{18}H_{18}Br_2N_2O_2SNa$ $[M+Na]^+$ 506.9348, found 506.9362.

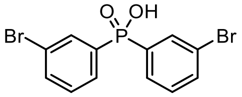

**Bis(3-bromophenyl)phosphinic acid (S33):** To a solution of 3-bromoiodobenzene (**S26**; 11.32 g, 40.0 mmol, 3 eq) in THF (100 mL) at −40 °C was added *i*-PrMgCl (2.0 M in THF, 20.00 mL, 40.0 mmol, 3 eq). The reaction was then gradually warmed to −25 °C while stirring for 2 h. After cooling the solution to −78 °C, *N,N*-dimethylphosphoramic dichloride (1.59 mL, 13.3 mmol) was added dropwise. The reaction was allowed to warm to room temperature overnight (18 h). After adding 6 N HCl (10 mL), the resulting mixture was vigorously stirred at room temperature for 4 h. It was then diluted with water and extracted with EtOAc (2×). The organics were washed with brine, dried over anhydrous $MgSO_4$, filtered, and concentrated *in vacuo*. The crude product was dissolved in $CH_2Cl_2$ (~100 mL), and hexanes (~100 mL) were slowly added. This solution was gently concentrated with rotary evaporation (~250 Torr)

until a pale yellow solid precipitated. The solid was isolated by filtration, washed with 2:1 hexanes/$CH_2Cl_2$ and $Et_2O$, and dried to afford 4.13 g (82%) of phosphinic acid **S33** as a yellow powder. $^1$H NMR (DMSO-$d_6$, 400 MHz) δ 7.86 (dt, $J$ = 12.1, 1.7 Hz, 2H), 7.78 – 7.69 (m, 4H), 7.46 (td, $J$ = 7.8, 3.8 Hz, 2H); $^{13}$C NMR (DMSO-$d_6$, 101 MHz) δ 137.4 (d, $J_{CP}$ = 133.3 Hz, C), 134.6 (d, $J_{CP}$ = 2.5 Hz, CH), 133.2 (d, $J_{CP}$ = 10.7 Hz, CH), 131.1 (d, $J_{CP}$ = 13.4 Hz, CH), 130.1 (d, $J_{CP}$ = 9.8 Hz, CH), 122.1 (d, $J_{CP}$ = 16.4 Hz, C); HRMS (ESI) calcd for $C_{12}H_{10}Br_2O_2P$ [M+H]$^+$ 374.8780, found 374.8791.

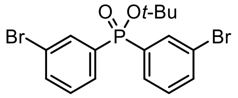


***tert*-Butyl bis(3-bromophenyl)phosphinate (S34):** A suspension of phosphinic acid **S33** (3.00 g, 7.98 mmol) in dioxane (25 mL) was heated to 80 °C, and *N*,*N*-dimethylformamide di-*tert*-butyl acetal (9.57 mL, 39.9 mmol, 5 eq) was added dropwise over 5 min. The reaction was stirred at 80 °C for 1 h. After cooling the mixture to room temperature, it was diluted with saturated $NaHCO_3$ and extracted with EtOAc (2×). The combined organic extracts were washed with water and brine, dried over anhydrous $MgSO_4$, filtered, and evaporated. Flash chromatography (0–50% EtOAc/hexanes, linear gradient) provided phosphinate **S34** as a white solid (1.78 g, 52%). $^1$H NMR ($CDCl_3$, 400 MHz) δ 7.92 (dt, $J$ = 12.4, 1.6 Hz, 2H), 7.69 (ddt, $J$ = 12.0, 7.6, 1.2 Hz, 2H), 7.62 (ddt, $J$ = 8.0, 1.9, 0.9 Hz, 2H), 7.31 (td, $J$ = 7.8, 4.0 Hz, 2H), 1.53 (s, 9H); $^{13}$C NMR ($CDCl_3$, 101 MHz) δ 136.8 (d, $J_{CP}$ = 137.5 Hz, C), 135.0 (d, $J_{CP}$ = 2.6 Hz, CH), 134.1 (d, $J_{CP}$ = 10.7 Hz, CH), 130.3 (d, $J_{CP}$ = 14.2 Hz, CH), 129.9 (d, $J_{CP}$ = 9.9 Hz, CH), 123.1 (d, $J_{CP}$ = 16.9 Hz, C), 85.3 (d, $J_{CP}$ = 8.1 Hz, C), 31.1 (d, $J_{CP}$ = 4.0 Hz, $CH_3$); HRMS (ESI) calcd for $C_{16}H_{17}Br_2O_2PNa$ [M+Na]$^+$ 452.9225, found 452.9243.

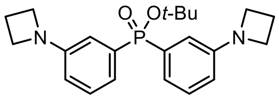


***tert*-Butyl bis(3-(azetidin-1-yl)phenyl)phosphinate (S35):** A vial was charged with dibromide **S34** (1.25 g, 2.89 mmol), RuPhos-G3-palladacycle (484 mg, 0.579 mmol, 0.2 eq), RuPhos (270 mg, 0.579 mmol, 0.2 eq), and $Cs_2CO_3$ (2.83 mg, 8.68 mmol, 3 eq). The vial was sealed and evacuated/backfilled with nitrogen (3×). Dioxane (11 mL) was added, and the reaction was flushed again with nitrogen (3×). Following the addition of azetidine (585 μL, 8.68 mmol, 3 eq), the reaction was stirred at 100 °C for 18 h. It was then cooled to room temperature, filtered through Celite with $CH_2Cl_2$, and concentrated to dryness. Purification by silica gel chromatography (0–40% acetone/$CH_2Cl_2$, linear gradient) afforded **S35** (1.08 g, 97%) as a pale yellow gum. $^1$H NMR ($CDCl_3$, 400 MHz) δ 7.20 (td, $J$ = 7.7, 4.2 Hz, 2H), 7.07 (ddt, $J$ = 12.0, 7.5, 1.2 Hz, 2H), 6.89 (ddd, $J$ = 13.7, 2.5, 1.3 Hz, 2H), 6.49 (ddt, $J$ = 8.1, 2.4, 1.1 Hz, 2H), 3.87 (t, $J$ = 7.2 Hz, 8H), 2.34 (p, $J$ = 7.2 Hz, 4H), 1.50 (s, 9H); $^{13}$C NMR ($CDCl_3$, 101 MHz) δ 151.8 (d, $J_{CP}$ = 15.0 Hz, C), 135.3 (d, $J_{CP}$ = 136.9 Hz, C), 128.8 (d, $J_{CP}$ = 14.9 Hz, CH), 120.1 (d, $J_{CP}$ = 9.9 Hz, CH), 114.1 (d, $J_{CP}$ = 2.9 Hz, CH), 113.9 (d, $J_{CP}$ = 11.6 Hz, CH), 83.1 (d, $J_{CP}$ = 8.3 Hz, C), 52.5 ($CH_2$), 31.0 (d, $J_{CP}$ = 4.0 Hz, $CH_3$), 17.0 ($CH_2$); HRMS (ESI) calcd for $C_{22}H_{29}N_2O_2PNa$ [M+Na]$^+$ 407.1859, found 407.1864.

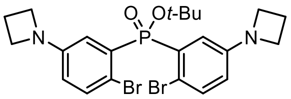


***tert*-Butyl bis(5-(azetidin-1-yl)-2-bromophenyl)phosphinate (S36):** Phosphinate **S35** (1.00 g, 2.60 mmol) was taken up in DMF (13 mL) and cooled to 0 °C. *N*-Bromosuccinimide (926 mg, 5.20 mmol, 2 eq) was added portion-wise over 10 min. The reaction was stirred at 0 °C for 1 h, then warmed to room temperature and stirred 1 h. It was subsequently diluted with water and extracted with EtOAc (2×). The combined organic extracts were washed with water and brine, dried over anhydrous $MgSO_4$, filtered, and concentrated *in vacuo*. Purification by silica gel chromatography (0–40% $MeCN/CH_2Cl_2$, linear gradient) provided dibromide **S36** as an off-white solid (825 mg, 58%). $^1H$ NMR ($CDCl_3$, 400 MHz) δ 7.33 – 7.27 (m, 4H), 6.41 – 6.34 (m, 2H), 3.89 (t, $J$ = 7.2 Hz, 8H), 2.38 (p, $J$ = 7.2 Hz, 4H), 1.53 (s, 9H); $^{13}C$ NMR ($CDCl_3$, 101 MHz) δ 150.7 (d, $J_{CP}$ = 13.1 Hz, C), 134.3 (d, $J_{CP}$ = 11.3 Hz, CH), 133.7 (d, $J_{CP}$ = 144.4 Hz, C), 119.9 (d, $J_{CP}$ = 8.5 Hz, CH), 115.9 (d, $J_{CP}$ = 2.7 Hz, CH), 111.3 (d, $J_{CP}$ = 6.7 Hz, C), 84.8 (d, $J_{CP}$ = 8.3 Hz, C), 52.6 ($CH_2$), 30.8 (d, $J_{CP}$ = 4.0 Hz, $CH_3$), 17.0 ($CH_2$); HRMS (ESI) calcd for $C_{22}H_{27}Br_2N_2O_2PNa$ $[M+Na]^+$ 563.0069, found 563.0080.

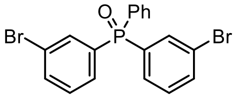


**Bis(3-bromophenyl)(phenyl)phosphine oxide (S37):** To a solution of 3-bromoiodobenzene (**S26**; 21.22 g, 75.0 mmol, 3 eq) in THF (200 mL) at −40 °C was added *i*-PrMgCl (2.0 M in THF, 37.50 mL, 75.0 mmol, 3 eq). The reaction was then gradually warmed to −20 °C over 2 h while stirring. Dichlorophenylphosphine (3.39 mL, 25.0 mmol) was added; the reaction was then warmed to room temperature and stirred for 2 h. It was subsequently quenched with saturated $NH_4Cl$ and diluted with water (~100 mL). After adding $H_2O_2$ (30%, 100 mL), the mixture was vigorously stirred at room temperature for 30 min and then extracted with EtOAc (2×). The combined organic extracts were washed with brine, dried over anhydrous $MgSO_4$, filtered, and evaporated. Flash chromatography on silica gel (10–100% EtOAc/toluene, linear gradient) afforded 10.81 g (99%) of phosphine oxide **S37** as a colorless gum. $^1H$ NMR ($CDCl_3$, 400 MHz) δ 7.82 (dt, $J$ = 12.1, 1.6 Hz, 2H), 7.73 – 7.47 (m, 9H), 7.36 (td, $J$ = 7.8, 3.4 Hz, 2H); $^{13}C$ NMR ($CDCl_3$, 101 MHz) δ 135.5 (d, $J_{CP}$ = 2.6 Hz, CH), 134.78 (d, $J_{CP}$ = 101.6 Hz, C), 134.76 (d, $J_{CP}$ = 10.6 Hz, CH), 132.7 (d, $J_{CP}$ = 2.8 Hz, CH), 132.1 (d, $J_{CP}$ = 10.1 Hz, CH), 131.2 (d, $J_{CP}$ = 106.0 Hz, C), 130.6 (d, $J_{CP}$ = 9.6 Hz, CH), 130.4 (d, $J_{CP}$ = 13.0 Hz, CH), 129.0 (d, $J_{CP}$ = 12.4 Hz, CH), 123.5 (d, $J_{CP}$ = 15.4 Hz, C); HRMS (ESI) calcd for $C_{18}H_{14}Br_2OP$ $[M+H]^+$ 434.9144, found 434.9150.

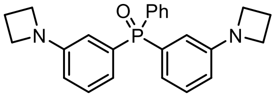


**Bis(3-(azetidin-1-yl)phenyl)(phenyl)phosphine oxide (S38):** A vial was charged with $Pd_2dba_3$ (210 mg, 0.229 mmol, 0.1 eq), XPhos (328 mg, 0.668 mmol, 0.3 eq), and $Cs_2CO_3$ (2.09 g, 6.42 mmol, 2.8 eq). The vial was sealed and evacuated/backfilled with nitrogen (3×). A solution of dibromide **S37** (1.00 g, 2.29 mmol) in dioxane (10 mL) was added, and the reaction was flushed again with nitrogen (3×). Following the addition of azetidine (371 µL, 5.50

mmol, 2.4 eq), the reaction was stirred at 100 °C for 18 h. It was then cooled to room temperature, filtered through Celite with $CH_2Cl_2$, and concentrated to dryness. The crude product was purified by flash chromatography (0–75% acetone/$CH_2Cl_2$, linear gradient) to yield 475 mg (53%) of **S38** as a pale yellow solid. $^{1}$H NMR ($CDCl_3$, 400 MHz) δ 7.70 – 7.62 (m, 2H), 7.53 – 7.46 (m, 1H), 7.45 – 7.38 (m, 2H), 7.21 (td, $J$ = 7.8, 3.7 Hz, 2H), 6.92 – 6.84 (m, 2H), 6.83 – 6.75 (m, 2H), 6.58 – 6.51 (m, 2H), 3.86 (t, $J$ = 7.2 Hz, 8H), 2.34 (p, $J$ = 7.3 Hz, 4H); $^{13}$C NMR ($CDCl_3$, 101 MHz) δ 152.0 (d, $J_{CP}$ = 13.3 Hz, C), 133.4 (d, $J_{CP}$ = 103.2 Hz, C), 133.1 (d, $J_{CP}$ = 103.0 Hz, C), 132.2 (d, $J_{CP}$ = 9.9 Hz, CH), 131.7 (d, $J_{CP}$ = 2.7 Hz, CH), 128.8 (d, $J_{CP}$ = 14.2 Hz, CH), 128.3 (d, $J_{CP}$ = 12.0 Hz, CH), 120.8 (d, $J_{CP}$ = 10.7 Hz, CH), 114.53 (d, $J_{CP}$ = 8.8 Hz, CH), 114.47 (d, $J_{CP}$ = 0.8 Hz, CH), 52.4 ($CH_2$), 17.0 ($CH_2$); HRMS (ESI) calcd for $C_{24}H_{26}N_2OP$ [M+H]$^+$ 389.1777, found 389.1779.

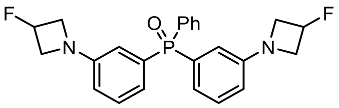

**Bis(3-(3-fluoroazetidin-1-yl)phenyl)(phenyl)phosphine oxide (S39):** An oven-dried round-bottom flask was charged with 3-fluoroazetidine hydrochloride (2.70 g, 24.2 mmol, 2.4 eq), $Pd_2dba_3$ (924 mg, 1.01 mmol, 0.1 eq), XPhos (1.44 g, 3.03 mmol, 0.3 eq), and $Cs_2CO_3$ (15.78 g, 48.4 mmol, 4.8 eq). The flask was sealed and evacuated/backfilled with nitrogen (3×). A solution of dibromide **S37** (4.40 g, 10.1 mmol) in dioxane (50 mL) was added, and after flushing the reaction again with nitrogen (3×), it was stirred at 100 °C for 18 h. It was then cooled to room temperature, filtered through Celite with $CH_2Cl_2$, and concentrated to dryness. The crude product was purified by flash chromatography (0–60% acetone/$CH_2Cl_2$, linear gradient) to yield 2.41 g (56%) of **S39** as a pale yellow solid. $^{1}$H NMR ($CDCl_3$, 400 MHz) δ 7.69 – 7.61 (m, 2H), 7.56 – 7.49 (m, 1H), 7.48 – 7.40 (m, 2H), 7.25 (td, $J$ = 8.0, 3.7 Hz, 2H), 6.93 (ddd, $J$ = 13.2, 2.2, 1.4 Hz, 2H), 6.88 – 6.79 (m, 2H), 6.63 – 6.56 (m, 2H), 5.39 (dtt, $^{2}J_{HF}$ = 57.0 Hz, $J$ = 5.9, 3.7 Hz, 2H), 4.24 – 4.11 (m, 4H), 4.03 – 3.88 (m, 4H); $^{19}$F NMR ($CDCl_3$, 376 MHz) δ −180.52 (dtt, $J_{FH}$ = 56.7, 23.8, 18.4 Hz); $^{13}$C NMR ($CDCl_3$, 101 MHz) δ 150.9 (dd, $^{3}J_{CP}$ = 13.4 Hz, $^{4}J_{CF}$ = 1.4 Hz, C), 133.3 (d, $J_{CP}$ = 103.0 Hz, C), 132.9 (d, $J_{CP}$ = 103.8 Hz, C), 132.2 (d, $J_{CP}$ = 9.9 Hz, CH), 131.9 (d, $J_{CP}$ = 2.9 Hz, CH), 129.0 (d, $J_{CP}$ = 14.0 Hz, CH), 128.5 (d, $J_{CP}$ = 12.1 Hz, CH), 121.6 (d, $J_{CP}$ = 10.7 Hz, CH), 115.2 (d, $J_{CP}$ = 2.8 Hz, CH), 115.0 (d, $J_{CP}$ = 10.3 Hz, CH), 82.8 (d, $^{1}J_{CF}$ = 204.7 Hz, CFH), 59.6 (d, $^{2}J_{CF}$ = 23.9 Hz, $CH_2$); HRMS (ESI) calcd for $C_{24}H_{24}F_2N_2OP$ [M+H]$^+$ 425.1589, found 425.1596.

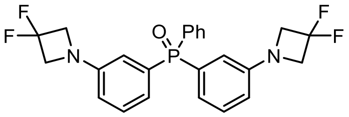

**Bis(3-(3,3-difluoroazetidin-1-yl)phenyl)(phenyl)phosphine oxide (S40):** An oven-dried round-bottom flask was charged with 3,3-difluoroazetidine hydrochloride (3.14 g, 24.2 mmol, 2.4 eq), $Pd_2dba_3$ (924 mg, 1.01 mmol, 0.1 eq), XPhos (1.44 g, 3.03 mmol, 0.3 eq), and $Cs_2CO_3$ (15.78 g, 48.4 mmol, 4.8 eq). The flask was sealed and evacuated/backfilled with nitrogen (3×). A solution of dibromide **S37** (4.40 g, 10.1 mmol) in dioxane (50 mL) was added, and after flushing the reaction again with nitrogen (3×), it was stirred at 100 °C for 18 h. It was then cooled to room temperature, filtered through Celite with $CH_2Cl_2$, and concentrated to dryness. The crude product was purified by flash chromatography (0–40% acetone/$CH_2Cl_2$, linear gradient) to yield 2.91 g (63%) of **S40** as a yellow foam. $^1$H NMR ($CDCl_3$, 400 MHz) δ 7.69 – 7.61 (m, 2H), 7.57 – 7.51 (m, 1H), 7.49 – 7.42 (m, 2H), 7.29 (td, $J$ = 7.8, 3.7 Hz, 2H), 6.97 (ddd, $J$ = 13.1, 2.3, 1.3 Hz, 2H), 6.93 – 6.85 (m, 2H), 6.66 – 6.61 (m, 2H), 4.22 (t, $^3J_{HF}$ = 11.7 Hz, 8H); $^{19}$F NMR ($CDCl_3$, 376 MHz) δ −99.92 (p, $^3J_{FH}$ = 11.9 Hz); $^{13}$C NMR ($CDCl_3$, 101 MHz) δ 149.7 (dt, $^3J_{CP}$ = 13.6 Hz, $^4J_{CF}$ = 2.8 Hz, C), 133.5 (d, $J_{CP}$ = 103.0 Hz, C), 132.6 (d, $J_{CP}$ = 104.2 Hz, C), 132.13 (d, $J_{CP}$ = 9.9 Hz, CH), 132.11 (d, $J_{CP}$ = 2.7 Hz, CH), 129.2 (d, $J_{CP}$ = 13.9 Hz, CH), 128.6 (d, $J_{CP}$ = 12.2 Hz, CH), 122.4 (d, $J_{CP}$ = 10.7 Hz, CH), 115.8 (t, $^1J_{CF}$ = 274.6 Hz, $CF_2$), 115.7 (d, $J_{CP}$ = 2.8 Hz, CH), 115.5 (d, $J_{CP}$ = 10.2 Hz, CH), 63.4 (t, $^2J_{CF}$ = 26.0 Hz, $CH_2$); HRMS (ESI) calcd for $C_{24}H_{22}F_4N_2OP$ $[M+H]^+$ 461.1400, found 461.1403.

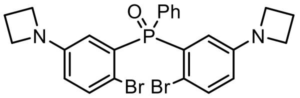


**Bis(5-(azetidin-1-yl)-2-bromophenyl)(phenyl)phosphine oxide (S41):** Phosphine oxide **S38** (3.00 g, 7.72 mmol) was taken up in DMF (250 mL). *N*-Bromosuccinimide (2.75 g, 15.5 mmol, 2 eq) was added portion-wise over 5 min, and the reaction was then stirred at room temperature for 18 h. The reaction mixture was concentrated *in vacuo*; the resulting residue was diluted with water and extracted with $CH_2Cl_2$ (2×). The combined organic extracts were washed with brine, dried over anhydrous $MgSO_4$, filtered, and evaporated to dryness. Silica gel chromatography (0–75% EtOAc/$CH_2Cl_2$, linear gradient) yielded 3.13 g (74%) of dibromide **S41** as a white solid. $^1$H NMR ($CDCl_3$, 400 MHz) δ 7.86 – 7.79 (m, 2H), 7.56 – 7.50 (m, 1H), 7.48 – 7.43 (m, 2H), 7.41 (dd, $J$ = 8.5, 4.7 Hz, 2H), 6.73 (dd, $J$ = 14.7, 2.9 Hz, 2H), 6.40 (ddd, $J$ = 8.6, 2.9, 0.7 Hz, 2H), 3.83 – 3.72 (m, 8H), 2.32 (p, $J$ = 7.3 Hz, 4H); $^{13}$C NMR ($CDCl_3$, 101 MHz) δ 150.8 (d, $J_{CP}$ = 12.8 Hz, C), 134.8 (d, $J_{CP}$ = 9.1 Hz, CH), 132.9 (d, $J_{CP}$ = 9.9 Hz, CH), 132.4 (d, $J_{CP}$ = 108.2 Hz, C), 131.9 (d, $J_{CP}$ = 2.9 Hz, CH), 131.5 (d, $J_{CP}$ = 109.7 Hz, C), 128.3 (d, $J_{CP}$ = 12.6 Hz, CH), 119.1 (d, $J_{CP}$ = 11.2 Hz, CH), 115.8 (d, $J_{CP}$ = 2.7 Hz, CH), 112.1 (d, $J_{CP}$ = 4.6 Hz, C), 52.3 ($CH_2$), 16.8 ($CH_2$); HRMS (ESI) calcd for $C_{24}H_{24}Br_2N_2OP$ $[M+H]^+$ 544.9988, found 545.0004.

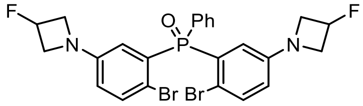


**Bis(2-bromo-5-(3-fluoroazetidin-1-yl)phenyl)(phenyl)phosphine oxide (S42):** Phosphine oxide **S39** (2.25 g, 5.30 mmol) was taken up in DMF (100 mL). *N*-Bromosuccinimide (1.89 g, 10.6 mmol, 2 eq) was added portion-wise over 5 min, and the reaction was then stirred at room temperature for 2 h. The reaction mixture was concentrated *in vacuo*; the resulting residue was diluted with water and extracted with $CH_2Cl_2$ (2×). The combined organic extracts were washed with brine, dried over anhydrous $MgSO_4$, filtered, and evaporated to dryness. Silica gel chromatography (0–60% EtOAc/$CH_2Cl_2$, linear gradient) afforded 1.53 g (50%) of dibromide **S42** as a white solid. $^1$H NMR ($CDCl_3$, 400 MHz) δ 7.88 – 7.79 (m, 2H), 7.60 – 7.52 (m, 1H), 7.50 – 7.42 (m, 4H), 6.79 (dd, $J$ = 14.5, 2.9 Hz, 2H), 6.46 (ddd, $J$ = 8.6, 2.9, 0.8 Hz, 2H), 5.36 (dtt, $^2J_{HF}$ = 56.9 Hz, $J$ = 5.9, 3.6 Hz, 2H), 4.15 – 4.03 (m, 4H), 3.95 – 3.82 (m, 4H); $^{19}$F NMR ($CDCl_3$, 376 MHz) δ −180.60 (dtt, $J_{FH}$ = 57.1, 23.8, 18.4 Hz); $^{13}$C NMR ($CDCl_3$, 101 MHz) δ 149.8 (dd, $^3J_{CP}$ = 12.9 Hz, $^4J_{CF}$ = 1.4 Hz, C), 135.0 (d, $J_{CP}$ = 9.0 Hz, CH), 132.9 (d, $J_{CP}$ = 9.9 Hz, CH), 132.8 (d, $J_{CP}$ = 108.1 Hz, C),

132.2 (d, $J_{CP}$ = 2.9 Hz, CH), 131.1 (d, $J_{CP}$ = 110.2 Hz, C), 128.4 (d, $J_{CP}$ = 12.6 Hz, CH), 119.6 (d, $J_{CP}$ = 11.2 Hz, CH), 116.5 (d, $J_{CP}$ = 2.5 Hz, CH), 113.1 (d, $J_{CP}$ = 4.7 Hz, C), 82.5 (d, $^1J_{CF}$ = 205.2 Hz, CFH), 59.5 (d, $^2J_{CF}$ = 24.0 Hz, $CH_2$); HRMS (ESI) calcd for $C_{24}H_{22}Br_2F_2N_2OP$ $[M+H]^+$ 580.9799, found 580.9811.

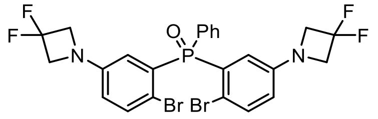

**Bis(2-bromo-5-(3,3-difluoroazetidin-1-yl)phenyl)(phenyl)phosphine oxide (S43):** Phosphine oxide **S40** (3.50 g, 7.60 mmol) was taken up in DMF (50 mL). *N*-Bromosuccinimide (2.71 g, 15.2 mmol, 2 eq) was added portion-wise over 5 min, and the reaction was then stirred at room temperature for 2 h. The reaction mixture was concentrated *in vacuo*; the resulting residue was diluted with water and extracted with $CH_2Cl_2$ (2×). The combined organic extracts were washed with brine, dried over anhydrous $MgSO_4$, filtered, and evaporated to dryness. Silica gel chromatography (10–100% EtOAc/toluene, linear gradient) afforded an off-white solid that was subsequently triturated with $Et_2O$ (~100 mL), sonicated, and filtered. The filter cake was washed with additional $Et_2O$ and dried to yield 3.31 g (70%) of dibromide **S43** as a white solid. $^1$H NMR ($CDCl_3$, 400 MHz) δ 7.89 – 7.80 (m, 2H), 7.62 – 7.54 (m, 1H), 7.53 – 7.45 (m, 4H), 6.84 (dd, $J$ = 14.4, 3.0 Hz, 2H), 6.50 (ddd, $J$ = 8.6, 3.0, 0.8 Hz, 2H), 4.21 – 4.06 (m, 8H); $^{19}$F NMR ($CDCl_3$, 376 MHz) δ −99.93 (p, $^3J_{HF}$ = 11.7 Hz); $^{13}$C NMR ($CDCl_3$, 101 MHz) δ 148.53 (dt, $^3J_{CP}$ = 12.9 Hz, $^4J_{CF}$ = 3.0 Hz, C), 135.2 (d, $J_{CP}$ = 9.1 Hz, CH), 133.1 (d, $J_{CP}$ = 107.9 Hz, C), 132.8 (d, $J_{CP}$ = 10.0 Hz, CH), 132.4 (d, $J_{CP}$ = 2.9 Hz, CH), 130.8 (d, $J_{CP}$ = 110.4 Hz, C), 128.5 (d, $J_{CP}$ = 12.7 Hz, CH), 120.0 (d, $J_{CP}$ = 11.1 Hz, CH), 117.0 (d, $J_{CP}$ = 2.6 Hz, CH), 115.6 (t, $^1J_{CF}$ = 274.7 Hz, $CF_2$), 114.1 (d, $J_{CP}$ = 4.7 Hz, C), 63.4 (t, $^2J_{CF}$ = 26.3 Hz, $CH_2$); HRMS (ESI) calcd for $C_{24}H_{20}Br_2F_4N_2OP$ $[M+H]^+$ 616.9611, found 616.9615.

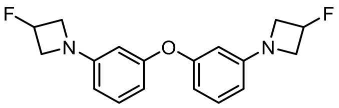

**1,1'-(Oxybis(3,1-phenylene))bis(3-fluoroazetidine) (S45):** An oven-dried round-bottom flask was charged with 3-fluoroazetidine hydrochloride (2.04 g, 18.3 mmol, 2.4 eq), $Pd_2dba_3$ (698 mg, 0.762 mmol, 0.1 eq), XPhos (1.09 g, 2.29 mmol, 0.3 eq), and $Cs_2CO_3$ (11.92 g, 36.6 mmol, 4.8 eq). The flask was sealed and evacuated/backfilled with nitrogen (3×). A solution of dibromide **S44** (2.50 g, 7.62 mmol) in dioxane (40 mL) was added, and after flushing the reaction again with nitrogen (3×), it was stirred at 100 °C for 18 h. It was then cooled to room temperature, filtered through Celite with $CH_2Cl_2$, and concentrated to dryness. Purification by flash chromatography (0–50% $Et_2O$/hexanes, linear gradient) yielded 1.43 g (59%) of ether **S45** as a yellow gum. $^1$H NMR ($CDCl_3$, 400 MHz) δ 7.15 (t, $J$ = 8.1 Hz, 2H), 6.41 (ddd, $J$ = 8.1, 2.2, 0.7 Hz, 2H), 6.21 (ddd, $J$ = 8.0, 2.2, 0.7 Hz, 2H), 6.15 (t, $J$ = 2.3 Hz, 2H), 5.39 (dtt, $^2J_{HF}$ = 57.0 Hz, $J$ = 5.9, 3.8 Hz, 2H), 4.21 – 4.09 (m, 4H), 3.99 – 3.87 (m, 4H); $^{19}$F NMR ($CDCl_3$, 376 MHz) δ −180.65 (dtt, $J_{FH}$ = 57.2, 23.9, 18.0 Hz); $^{13}$C NMR ($CDCl_3$, 101 MHz) δ 158.2 (C), 152.64 (d, $^4J_{CF}$ = 1.1 Hz, C), 130.2 (CH), 108.7 (CH), 107.0 (CH), 102.8 (CH), 82.8 (d, $^1J_{CF}$ = 204.5 Hz, CFH), 59.7 (d, $^2J_{CF}$ = 23.7 Hz, $CH_2$); HRMS (ESI) calcd for $C_{18}H_{19}F_2N_2O$ $[M+H]^+$ 317.1460, found 317.1456.

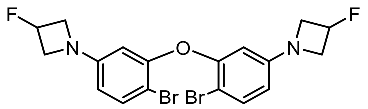

**1,1'-(Oxybis(4-bromo-3,1-phenylene))bis(3-fluoroazetidine) (S46):** Ether **S45** (1.50 g, 4.74 mmol) was taken up in DMF (30 mL). *N*-Bromosuccinimide (1.69 g, 9.48 mmol, 2 eq) was added portion-wise over 5 min, and the reaction was then stirred at room temperature for 2 h. The reaction mixture was concentrated *in vacuo*; the resulting residue was diluted with water and extracted with EtOAc (2×). The combined organic extracts were washed with brine, dried over anhydrous $MgSO_4$, filtered, and evaporated to dryness. Silica gel chromatography (0–10% EtOAc/toluene, linear gradient; mixed fractions repurified with 0–50% EtOAc/hexanes, linear gradient) provided 1.92 g (85%) of dibromide **S46** as a white solid. $^{1}$H NMR ($CDCl_3$, 400 MHz) δ 7.41 (d, $J$ = 8.6 Hz, 2H), 6.13 (dd, $J$ = 8.6, 2.6 Hz, 2H), 5.93 (d, $J$ = 2.7 Hz, 2H), 5.36 (dtt, $^{2}J_{HF}$ = 56.9 Hz, $J$ = 5.9, 3.6 Hz, 2H), 4.15 – 4.04 (m, 4H), 3.96 – 3.82 (m, 4H); $^{19}$F NMR ($CDCl_3$, 376 MHz) δ −180.71 (dtt, $J_{FH}$ = 57.1, 23.8, 18.3 Hz); $^{13}$C NMR ($CDCl_3$, 101 MHz) δ 153.8 (C), 151.6 (d, $^{4}J_{CF}$ = 1.3 Hz, C), 133.9 (CH), 108.9 (CH), 103.2 (CH), 101.9 (C), 82.5 (d, $^{1}J_{CF}$ = 205.0 Hz, CFH), 59.7 (d, $^{2}J_{CF}$ = 23.9 Hz, $CH_2$); HRMS (ESI) calcd for $C_{18}H_{17}Br_2F_2N_2O$ $[M+H]^+$ 472.9670, found 472.9677.

### Rhodamine Synthesis via Metal–Bromide Exchange of Dibromides

In applying the bis(arylmetal)-anhydride approach[1] to the synthesis of these new rhodamines (**Scheme S1**, **Scheme S2e**), we found that some variation of the reaction conditions was necessary depending on the dibromide substrate. For the nitrogen (**S14**–**S16**), sulfide (**S31**), and ether (**S46**) dibromides, the previously described conditions (*t*-BuLi alone or *t*-BuLi/MgBr·$OEt_2$) were typically adequate. Lithium–bromide exchange of **S14**–**S16** with *t*-BuLi and addition to phthalic anhydride yielded nitrogen rhodamines **1** ($JF_{502}$), **11** ($JF_{479}$), and **S18** ($JF_{490}$) in moderate yield. Alternatively, use of methyl ester **S19**[1] as the electrophile provided the 6-carboxy analogs of $JF_{502}$ (**S22**) and $JF_{479}$ (**S23**) in excellent yield following ester hydrolysis (**Scheme S1**). Similar reaction of sulfide **S31** and ether **S46** with tetrafluorophthalic anhydride afforded the 4,5,6,7-tetrafluororhodamines **20** ($JF_{593}$) and **43** ($JF_{559}$) in moderate yield (**Scheme S2e**). For sulfone **S32**, phosphinate **S36**, and phosphine oxides **S41**–**S43**, however, we observed poor yields, complex mixtures of products, and/or decomposition when these transformations were attempted with *t*-BuLi or *t*-BuLi/MgBr·$OEt_2$. We instead found that the milder magnesate $Bu_2$-*i*-PrMgLi was able to achieve reasonably clean magnesium–bromide exchange with these dibromides; subsequent addition to phthalic or tetrafluorophthalic anhydride allowed access to the sulfone (**8**, **18**), phosphine oxide (**7**, **17**, **45**, **S50**), and phosphinate (**S48**, **S49**) rhodamines, albeit in modest yields. Straightforward TFA deprotection of the *tert*-butyl phosphinates generated the phosphinic acids **6** ($JF_{668}$) and **16** ($JF_{690}$).

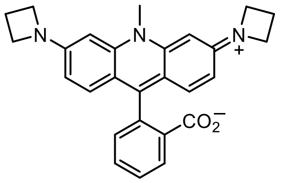


**$JF_{502}$ (1):** A solution of dibromide **S14** (150 mg, 0.332 mmol) in THF (10 mL) was cooled to −78 °C under nitrogen. *tert*-Butyllithium (1.7 M in pentane, 860 µL, 1.46 mmol, 4.4 eq) was added, and the reaction was stirred at −78 °C for 30 min. It was then warmed to −20 °C, and a solution of phthalic anhydride (**S17**; 108 mg, 0.731 mmol, 2.2 eq) in THF (10 mL) was added dropwise over 30 min via addition funnel. The reaction was allowed to warm to room temperature overnight (18 h). Following the addition of AcOH (100 µL), the mixture was diluted with MeOH, deposited onto Celite, and concentrated to dryness. Silica gel chromatography (0–10% MeOH (2 M $NH_3$)/$CH_2Cl_2$, linear gradient) followed by reverse phase HPLC (30–60% MeCN/$H_2O$, linear gradient, with constant 0.1% v/v TFA additive) afforded 77.6 mg (43%, TFA salt) of $JF_{502}$ (**1**) as a bright orange solid. $^1H$ NMR ($CD_3OD$, 400 MHz) δ 8.14 – 8.10 (m, 1H), 7.69 – 7.59 (m, 2H), 7.33 (d, $J$ = 9.2 Hz, 2H), 7.21 – 7.16 (m, 1H), 6.66 (dd, $J$ = 9.3, 2.0 Hz, 2H), 6.35 (d, $J$ = 2.0 Hz, 2H), 4.27 – 4.13 (m, 8H), 3.94 (s, 3H), 2.51 (p, $J$ = 7.6 Hz, 4H); $^{13}C$ NMR ($CD_3OD$, 101 MHz) δ 158.8 (C), 156.3 (C), 144.7 (C), 135.8 (C), 133.2 (CH), 131.5 (CH), 131.13 (CH), 131.06 (CH), 131.04 (C), 130.3 (CH), 117.9 (C), 113.1 (CH), 91.8 (CH), 52.5 ($CH_2$), 36.3 ($CH_3$), 16.9 ($CH_2$); Analytical HPLC: $t_R$ = 12.0 min, >99% purity (10–95% MeCN/$H_2O$, linear gradient, with constant 0.1% v/v TFA additive; 20 min run; 1 mL/min flow; ESI; positive ion mode; detection at 500 nm); HRMS (ESI) calcd for $C_{27}H_{26}N_3O_2$ $[M+H]^+$ 424.2020, found 424.2021.

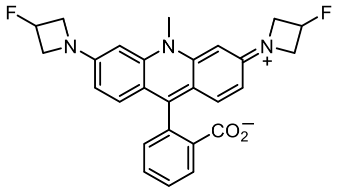

**$JF_{490}$ (S18):** A solution of dibromide **S15** (200 mg, 0.411 mmol) in THF (10 mL) was cooled to −78 °C under nitrogen. *tert*-Butyllithium (1.7 M in pentane, 1.06 mL, 1.81 mmol, 4.4 eq) was added, and the reaction was stirred at −78 °C for 30 min. It was then warmed to −20 °C, and a solution of phthalic anhydride (**S17**; 134 mg, 0.903 mmol, 2.2 eq) in THF (10 mL) was added dropwise over 30 min via addition funnel. The reaction was allowed to warm to room temperature overnight (18 h). Following the addition of AcOH (100 μL), the mixture was diluted with MeOH, deposited onto Celite, and concentrated to dryness. Silica gel chromatography (0–20% MeOH (2 M $NH_3$)/$CH_2Cl_2$, linear gradient; dry load with Celite) followed by reverse phase HPLC (10–75% MeCN/$H_2O$, linear gradient, with constant 0.1% v/v TFA additive) afforded 105 mg (45%, TFA salt) of $JF_{490}$ (**S18**) as a bright orange solid. $^1$H NMR ($CD_3OD$, 400 MHz) δ 8.39 – 8.34 (m, 1H), 7.84 (td, $J$ = 7.5, 1.7 Hz, 1H), 7.79 (td, $J$ = 7.6, 1.6 Hz, 1H), 7.36 – 7.31 (m, 1H), 7.27 (d, $J$ = 9.2 Hz, 2H), 6.78 (dd, $J$ = 9.2, 2.0 Hz, 2H), 6.61 (d, $J$ = 2.0 Hz, 2H), 5.55 (dtt, $^2J_{HF}$ = 57.1 Hz, $J$ = 6.0, 3.1 Hz, 2H), 4.61 – 4.48 (m, 4H), 4.39 – 4.23 (m, 4H), 4.13 (s, 3H); $^{19}$F NMR ($CD_3OD$, 376 MHz) δ −75.59 (s, 3F), -180.26 (dtt, $J_{FH}$ = 56.8, 23.7, 20.3 Hz, 2F); $^{13}$C NMR ($CD_3OD$, 101 MHz) δ 168.2 (C), 158.2 (C), 156.1 (d, $^4J_{CF}$ = 2.3 Hz, C), 144.8 (C), 137.5 (C), 133.9 (CH), 132.62 (CH), 132.58 (CH), 132.21 (C), 132.17 (CH), 131.1 (CH), 118.3 (C), 114.3 (CH), 93.0 (CH), 83.7 (d, $^1J_{CF}$ = 203.4 Hz, CHF), 60.2 (d, $^2J_{CF}$ = 26.0 Hz, $CH_2$), 36.6 ($CH_3$); Analytical HPLC: $t_R$ = 10.9 min, >99% purity (10–75% MeCN/$H_2O$, linear gradient, with constant 0.1% v/v TFA additive; 20 min run; 1 mL/min flow; ESI; positive ion mode; detection at 500 nm); HRMS (ESI) calcd for $C_{27}H_{24}F_2N_3O_2$ $[M+H]^+$ 460.1831, found 460.1835.

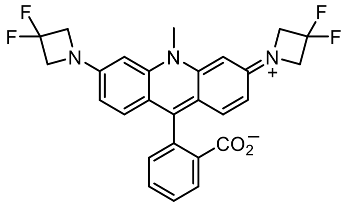

**$JF_{479}$ (11):** A solution of dibromide **S16** (200 mg, 0.382 mmol) in THF (10 mL) was cooled to −78 °C under nitrogen. *tert*-Butyllithium (1.7 M in pentane, 989 μL, 1.68 mmol, 4.4 eq) was added, and the reaction was stirred at −78 °C for 30 min. It was then warmed to −20 °C, and a solution of phthalic anhydride (**S17**; 125 mg, 0.841 mmol, 2.2 eq) in THF (10 mL) was added dropwise over 30 min via addition funnel. The reaction was allowed to warm to room temperature overnight (18 h). Following the addition of AcOH (100 μL), the mixture was diluted with MeOH, deposited onto Celite, and concentrated to dryness. Silica gel chromatography (0–20% MeOH (2 M $NH_3$)/$CH_2Cl_2$, linear gradient; dry load with Celite) afforded 95 mg (50%) of $JF_{479}$ (**11**) as a bright orange solid. $^1$H NMR ($CD_3OD$, 400 MHz) δ 8.19 – 8.13 (m, 1H), 7.69 (td, $J$ = 7.6, 1.5 Hz, 1H), 7.64 (td, $J$ = 7.4, 1.6 Hz, 1H), 7.49 (d, $J$ = 9.2 Hz, 2H), 7.23 – 7.18 (m, 1H), 6.85 (dd, $J$ = 9.2, 2.0 Hz, 2H), 6.73 (d, $J$ = 2.0 Hz, 2H), 4.65 – 4.56 (m, 8H), 4.15 (s, 3H); $^{19}$F NMR ($CD_3OD$, 376 MHz) δ −100.58 – −100.76 (m); Analytical HPLC: $t_R$ = 11.4 min, >99% purity (10–75%

$MeCN/H_2O$, linear gradient, with constant 0.1% v/v TFA additive; 20 min run; 1 mL/min flow; ESI; positive ion mode; detection at 500 nm); HRMS (ESI) calcd for $C_{27}H_{22}F_4N_3O_2$ $[M+H]^+$ 496.1643, found 496.1651.

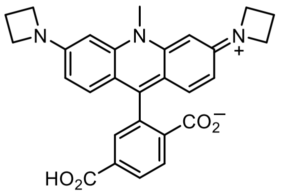

**6-Carboxy-JF$_{502}$ (S22):** A solution of dibromide **S14** (325 mg, 0.720 mmol, 1.5 eq) in THF (20 mL) was cooled to −78 °C under nitrogen. *tert*-Butyllithium (1.7 M in pentane, 1.69 mL, 2.88 mmol, 6 eq) was added, and the reaction was stirred at −78 °C for 30 min. It was then warmed to −20 °C, and a solution of ester **S19**[1] (188 mg, 0.480 mmol) in THF (10 mL) was added dropwise over 30 min via addition funnel. The reaction was allowed to warm to room temperature overnight (18 h). It was subsequently diluted with saturated $NH_4Cl$ and water and extracted with 15% *i*-PrOH/$CHCl_3$ (3×). The combined organic extracts were dried over anhydrous $MgSO_4$, filtered, and concentrated *in vacuo*. The resulting residue was redissolved in MeOH (10 mL), and 1 M HCl (0.5 mL) was added. After stirring the solution at room temperature for 30 min, it was diluted with toluene (10 mL), deposited onto Celite, concentrated to dryness, and purified by flash chromatography (0–20% MeOH/$CH_2Cl_2$, linear gradient, with constant 1% v/v AcOH additive; dry load with Celite) to provide the bis(2,2-bis(hydroxymethyl)propyl) diester intermediate **S20** (213 mg, 61%, acetate salt).

Diester **S20** (213 mg, 0.291 mmol) was taken up in 2,2,2-trifluoroethanol (9 mL), and 25% w/w NaOH (3 mL) was added. The reaction was stirred at room temperature for 7 days. It was then acidified with AcOH (3 mL), diluted with water, and extracted with 15% *i*-PrOH/$CHCl_3$ (3×). The combined organic extracts were dried over anhydrous $MgSO_4$, filtered, and concentrated *in vacuo*. Purification by flash chromatography on silica gel (0–20% MeOH/$CH_2Cl_2$, linear gradient, with constant 1% v/v AcOH additive) afforded 6-carboxy-JF$_{502}$ (**S22**) as an orange solid (116 mg, 76%, acetate salt). $^1$H NMR ($CD_3OD$, 400 MHz) δ 8.41 (d, $J$ = 8.1 Hz, 1H), 8.37 (dd, $J$ = 8.2, 1.6 Hz, 1H), 7.89 (d, $J$ = 1.6 Hz, 1H), 7.18 (d, $J$ = 9.2 Hz, 2H), 6.70 (dd, $J$ = 9.2, 2.0 Hz, 2H), 6.45 (d, $J$ = 2.0 Hz, 2H), 4.30 – 4.21 (m, 8H), 4.08 (s, 3H), 2.54 (p, $J$ = 7.5 Hz, 4H); $^{13}$C NMR ($CD_3OD$, 101 MHz) δ 167.9 (C), 167.6 (C), 156.5 (C), 155.9 (C), 144.9 (C), 137.7 (C), 136.2 (C), 135.8 (C), 133.0 (CH), 132.7 (CH), 132.2 (CH), 131.8 (CH), 117.6 (C), 113.7 (CH), 91.9 (CH), 52.6 ($CH_2$), 36.4 ($CH_3$), 16.9 ($CH_2$); Analytical HPLC: $t_R$ = 9.7 min, >99% purity (10–95% $MeCN/H_2O$, linear gradient, with constant 0.1% v/v TFA additive; 20 min run; 1 mL/min flow; ESI; positive ion mode; detection at 500 nm); HRMS (ESI) calcd for $C_{28}H_{26}N_3O_4$ $[M+H]^+$ 468.1918, found 468.1921.

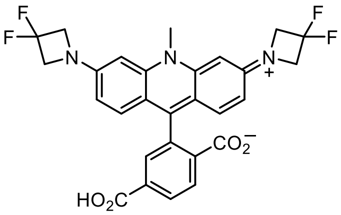


**6-Carboxy-JF$_{479}$ (S23):** A solution of dibromide **S16** (500 mg, 0.956 mmol, 1.5 eq) in THF (25 mL) was cooled to −78 °C under nitrogen. *tert*-Butyllithium (1.7 M in pentane, 2.25 mL, 3.82 mmol, 6 eq) was added, and the reaction was stirred at −78 °C for 30 min. It was then warmed to −20 °C, and a solution of ester **S19** (250 mg, 0.637 mmol, 1 eq) in THF (15 mL) was added dropwise over 30 min via addition funnel. The reaction was allowed to warm to room temperature overnight (18 h). It was subsequently diluted with saturated $NH_4Cl$ and water and extracted with EtOAc (1×) and 15% *i*-PrOH/$CHCl_3$ (3×). The combined organic extracts were dried over anhydrous $MgSO_4$, filtered, and concentrated *in vacuo*. The resulting residue was redissolved in MeOH (10 mL), and 1 M HCl (0.5 mL) was added. After stirring the solution at room temperature for 30 min, it was diluted with toluene (10 mL), concentrated to dryness, and purified by flash chromatography (0–20% MeOH/$CH_2Cl_2$, linear gradient, with constant 1% v/v AcOH additive) to provide the bis(2,2-bis(hydroxymethyl)propyl) diester intermediate **S21** (397 mg, 78%, acetate salt).

Diester **S21** (397 mg, 0.494 mmol) was taken up in 2,2,2-trifluoroethanol (15 mL), and 25% w/w NaOH (5 mL) was added. The reaction was stirred at room temperature for 6 days. It was then acidified with AcOH (5 mL), diluted with water, and extracted with 15% *i*-PrOH/$CHCl_3$ (3×) and 10% MeOH/$CH_2Cl_2$ (3×). The aqueous layer—which was still an orange suspension after extensive extraction—was filtered; the filter cake was washed with water and $CH_2Cl_2$ and dried to provide clean product as an orange powder. The combined organic extracts were dried over anhydrous $MgSO_4$, filtered, and evaporated to dryness. The resultant residue was triturated with $CH_2Cl_2$ and filtered. The isolated solid was washed with $CH_2Cl_2$ and water and dried, affording additional clean product. The two crops of orange solid were combined, providing a total yield of 229 mg (86%) of 6-carboxy-JF$_{479}$ (**S23**). $^1$H NMR ($CD_3OD$, 400 MHz) δ 8.45 (d, *J* = 8.2 Hz, 1H), 8.40 (dd, *J* = 8.2, 1.6 Hz, 1H), 7.93 (d, *J* = 1.6 Hz, 1H), 7.35 (d, *J* = 9.2 Hz, 2H), 6.89 (dd, *J* = 9.3, 2.0 Hz, 2H), 6.83 (d, *J* = 2.0 Hz, 2H), 4.66 (t, $^3J_{HF}$ = 11.8 Hz, 8H), 4.31 (s, 3H); $^{19}$F NMR ($CD_3OD$, 376 MHz) δ -100.74 (p, $^3J_{FH}$ = 11.7 Hz); Analytical HPLC: $t_R$ = 9.0 min, >99% purity (10–95% MeCN/$H_2O$, linear gradient, with constant 0.1% v/v TFA additive; 20 min run; 1 mL/min flow; ESI; positive ion mode; detection at 475 nm); HRMS (ESI) calcd for $C_{28}H_{22}F_4N_3O_4$ $[M+H]^+$ 540.1541, found 540.1544.

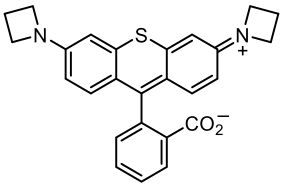


**JF$_{570}$ (3):** A solution of dibromide **S31** (200 mg, 0.440 mmol) in THF (10 mL) was cooled to −78 °C under nitrogen. *tert*-Butyllithium (1.7 M in pentane, 1.14 mL, 1.94 mmol, 4.4 eq) was added, and the reaction was stirred at −78 °C for 30 min. It was then warmed to −20 °C, and a solution of phthalic anhydride (**S17**; 143 mg, 0.969 mmol, 2.2 eq) in

THF (10 mL) was added dropwise over 30 min via addition funnel. The reaction was allowed to warm to room temperature overnight (18 h). Following the addition of AcOH (100 μL), the mixture was diluted with MeOH, deposited onto Celite, and concentrated to dryness. Silica gel chromatography (0–10% MeOH (2 M $NH_3$)/$CH_2Cl_2$, linear gradient; dry load with Celite) afforded 87 mg (46%) of $JF_{570}$ (**3**) as a dark purple solid. $^1$H NMR ($CD_3OD$, 400 MHz) δ 8.14 – 8.08 (m, 1H), 7.68 – 7.59 (m, 2H), 7.25 (d, $J$ = 9.3 Hz, 2H), 7.21 – 7.17 (m, 1H), 6.79 (d, $J$ = 2.3 Hz, 2H), 6.55 (dd, $J$ = 9.3, 2.3 Hz, 2H), 4.22 (t, $J$ = 7.6 Hz, 8H), 2.51 (p, $J$ = 7.6 Hz, 4H); $^{13}$C NMR ($CD_3OD$, 101 MHz) δ 172.6 (C), 158.9 (C), 154.0 (C), 144.0 (C), 139.14 (C), 139.10 (C), 136.7 (CH), 131.3 (CH), 130.8 (CH), 130.4 (CH), 130.3 (CH), 120.7 (C), 113.9 (CH), 104.6 (CH), 52.6 ($CH_2$), 16.9 ($CH_2$); Analytical HPLC: $t_R$ = 11.8 min, >99% purity (10–95% MeCN/$H_2O$, linear gradient, with constant 0.1% v/v TFA additive; 20 min run; 1 mL/min flow; ESI; positive ion mode; detection at 550 nm); HRMS (ESI) calcd for $C_{26}H_{23}N_2O_2S$ $[M+H]^+$ 427.1475, found 427.1489.

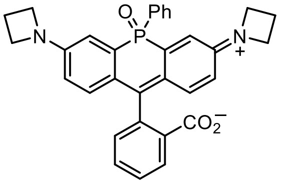

**2-(3,7-Di(azetidin-1-yl)-5-oxido-5-phenyl-10*H*-acridophosphin-10-ylium-10-yl)benzoate (7):** A solution of dibromide **S41** (250 mg, 0.458 mmol) in THF (40 mL) was cooled to −20 °C under nitrogen. Lithium dibutyl(isopropyl)magnesate (0.7 M in $Et_2O$/hexanes, 785 μL, 0.549 mmol, 1.2 eq) was added, and the reaction was stirred at -20 °C for 20 min. A solution of phthalic anhydride (**S17**; 244 mg, 1.65 mmol, 3.6 eq) in THF (5 mL) was added dropwise over 5 min, and the reaction was gradually warmed to 0 °C over 4 h. It was subsequently quenched with saturated $NH_4Cl$, diluted with water, and extracted with EtOAc (2×). The combined organic extracts were washed with saturated $NaHCO_3$ and brine, dried over anhydrous $MgSO_4$, filtered, and evaporated. Flash chromatography on silica gel (0–75% acetone/$CH_2Cl_2$, linear gradient) yielded 50 mg (21%) of phosphine oxide rhodamine **7** as a white solid. $^1$H NMR ($CDCl_3$, 400 MHz) δ 7.94 (dt, $J$ = 7.8, 1.0 Hz, 1H), 7.68 – 7.58 (m, 2H), 7.54 – 7.36 (m, 4H), 7.22 (td, $J$ = 7.7, 1.2 Hz, 1H), 7.13 (dd, $J$ = 13.2, 2.6 Hz, 2H), 6.92 (dd, $J$ = 8.8, 6.0 Hz, 2H), 6.49 (dd, $J$ = 8.8, 2.6 Hz, 2H), 6.22 (d, $J$ = 7.8 Hz, 1H), 4.01 – 3.88 (m, 8H), 2.38 (p, $J$ = 7.3 Hz, 4H); $^{13}$C NMR ($CDCl_3$, 101 MHz) δ 171.0 (C), 155.9 (d, $J_{CP}$ = 0.8 Hz, C), 151.4 (d, $J_{CP}$ = 11.9 Hz, C), 136.5 (d, $J_{CP}$ = 108.0 Hz, C), 134.7 (CH), 131.6 (d, $J_{CP}$ = 2.8 Hz, CH), 131.3 (d, $J_{CP}$ = 10.0 Hz, CH), 129.0 (CH), 128.9 (d, $J_{CP}$ = 2.9 Hz, CH), 128.7 (d, $J_{CP}$ = 12.2 Hz, CH), 127.5 (d, $J_{CP}$ = 100.2 Hz, C), 127.1 (d, $J_{CP}$ = 7.6 Hz, C), 125.6 (CH), 124.8 (C), 122.4 (CH), 115.5 (d, $J_{CP}$ = 2.4 Hz, CH), 112.1 (d, $J_{CP}$ = 6.6 Hz, CH), 85.9 (d, $J_{CP}$ = 7.5 Hz, C), 52.2 ($CH_2$), 16.8 ($CH_2$); Analytical HPLC: $t_R$ = 12.4 min, >99% purity (10–95% MeCN/$H_2O$, linear gradient, with constant 0.1% v/v TFA additive; 20 min run; 1 mL/min flow; ESI; positive ion mode; detection at 675 nm); HRMS (ESI) calcd for $C_{32}H_{28}N_2O_3P$ $[M+H]^+$ 519.1832, found 519.1837.

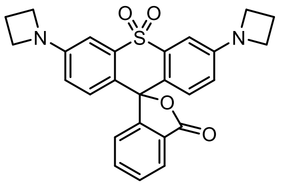


**3',6'-Di(azetidin-1-yl)-3*H*-spiro[isobenzofuran-1,9'-thioxanthen]-3-one 10',10'-dioxide (8):** A solution of dibromide **S32** (250 mg, 0.514 mmol) in THF (40 mL) was cooled to −20 °C under nitrogen. Lithium dibutyl(isopropyl)magnesate (0.7 M in $Et_2O$/hexanes, 881 µL, 0.617 mmol, 1.2 eq) was added, and the reaction was stirred at -20 °C for 20 min. A solution of phthalic anhydride (**S17**; 274 mg, 1.85 mmol, 3.6 eq) in THF (5 mL) was then added dropwise over 10 min. The reaction was gradually warmed to 0 °C over 4 h while stirring. It was subsequently quenched with saturated $NH_4Cl$, diluted with water, and extracted with EtOAc (1×) and 15% *i*-PrOH/$CHCl_3$ (2×). The combined organic extracts were dried over anhydrous $MgSO_4$, filtered, and evaporated. The crude residue was triturated with $CH_2Cl_2$, sonicated, and filtered; the filter cake was washed with additional $CH_2Cl_2$ and dried to yield 57 mg (24%) of $SO_2$-rhodamine **8** as an off-white solid. $^1H$ NMR ($CDCl_3$, 400 MHz) δ 7.95 – 7.91 (m, 1H), 7.82 – 7.79 (m, 1H), 7.52 (td, $J$ = 7.5, 1.3 Hz, 1H), 7.46 (td, $J$ = 7.4, 1.0 Hz, 1H), 7.13 (d, $J$ = 8.7 Hz, 2H), 7.10 (d, $J$ = 2.5 Hz, 2H), 6.47 (dd, $J$ = 8.8, 2.6 Hz, 2H), 3.97 (t, $J$ = 7.4 Hz, 8H), 2.42 (p, $J$ = 7.3 Hz, 4H); $^{13}C$ NMR ($CDCl_3$, 101 MHz) δ 170.9 (C), 154.6 (C), 152.0 (C), 136.6 (C), 135.5 (CH), 129.5 (CH), 127.5 (CH), 125.9 (CH), 123.8 (CH), 123.6 (C), 123.3 (C), 115.4 (CH), 104.7 (CH), 83.1 (C), 52.2 ($CH_2$), 16.8 ($CH_2$); Analytical HPLC: $t_R$ = 15.5 min, >99% purity (10–95% MeCN/$H_2O$, linear gradient, with constant 0.1% v/v TFA additive; 20 min run; 1 mL/min flow; ESI; positive ion mode; detection at 280 nm); HRMS (ESI) calcd for $C_{26}H_{22}N_2O_4SNa$ $[M+Na]^+$ 481.1192, found 481.1201.

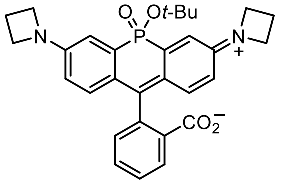


***tert*-Butyl-JF$_{668}$ (S48):** A solution of dibromide **S36** (100 mg, 0.184 mmol) in THF (4 mL) was cooled to −20 °C under nitrogen. Lithium dibutyl(isopropyl)magnesate (0.7 M in $Et_2O$/hexanes, 316 µL, 0.221 mmol, 1.2 eq) was added, and the reaction was stirred at -20 °C for 20 min. A solution of phthalic anhydride (**S17**; 98 mg, 0.664 mmol, 3.6 eq) in THF (2 mL) was added dropwise over 10 min, and the reaction was allowed to warm to room temperature overnight (18 h). It was subsequently quenched with saturated $NH_4Cl$, diluted with water, and extracted with EtOAc (2×). The combined organic extracts were washed with saturated $NaHCO_3$ and brine, dried over anhydrous $MgSO_4$, filtered, and evaporated. Flash chromatography on silica gel (0–50% MeCN/$CH_2Cl_2$, linear gradient) afforded 41 mg (43%) of *tert*-butyl-JF$_{668}$ (**S48**) as a tan solid. $^1H$ NMR ($CDCl_3$, 400 MHz) δ 7.93 – 7.88 (m, 1H), 7.72 – 7.66 (m, 1H), 7.44 – 7.37 (m, 2H), 7.13 (dd, $J$ = 8.8, 6.9 Hz, 2H), 7.09 (dd, $J$ = 14.4, 2.6 Hz, 2H), 6.47 (dd, $J$ = 8.8, 2.6 Hz, 2H), 3.99 – 3.88 (m, 8H), 2.38 (p, $J$ = 7.3 Hz, 4H), 1.66 (s, 9H); $^{13}C$ NMR ($CDCl_3$, 101 MHz) δ 171.7 (C), 156.92 (d, $J_{CP}$ = 1.2 Hz, C), 151.4 (d, $J_{CP}$ = 13.1 Hz, C), 134.6 (CH), 129.4 (d, $J_{CP}$= 8.6 Hz, C), 129.0 (d, $J_{CP}$ = 130.5 Hz, C), 128.8 (CH), 127.0 (d, $J_{CP}$ = 11.7 Hz, CH), 125.9 (CH), 123.13 (CH), 123.12 (C), 115.2 (d, $J_{CP}$ = 2.3 Hz, CH), 111.1 (d, $J_{CP}$ = 6.4

Hz, CH), 85.8 (d, $J_{CP}$ = 9.9 Hz, C), 84.8 (d, $J_{CP}$ = 8.5 Hz, C), 52.4 ($CH_2$), 31.4 (d, $J_{CP}$ = 3.9 Hz, $CH_3$), 17.0 ($CH_2$); Analytical HPLC: $t_R$ = 14.8 min, >99% purity (10–95% MeCN/$H_2O$, linear gradient, with constant 0.1% v/v TFA additive; 20 min run; 1 mL/min flow; ESI; positive ion mode; detection at 700 nm); HRMS (ESI) calcd for $C_{30}H_{31}N_2O_4PNa$ $[M+Na]^+$ 537.1914, found 537.1930.

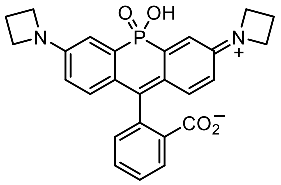

**JF$_{668}$ (6):** Phosphinate **S48** (28 mg, 54.4 µmol) was taken up in $CH_2Cl_2$ (2 mL); triethylsilane (200 µL) was added, followed by trifluoroacetic acid (200 µL). The reaction was stirred at room temperature for 1 h. Toluene (2 mL) was added, and the reaction mixture was concentrated to dryness. The crude material was purified by reverse phase HPLC (10–50% MeCN/$H_2O$, linear gradient, with constant 0.1% v/v TFA additive). The pooled HPLC product fractions were partially concentrated to remove MeCN and extracted with 15% *i*-PrOH/$CHCl_3$ (3×). The organic extracts were dried over anhydrous $MgSO_4$, filtered, and evaporated to yield 20 mg (80%) of JF$_{668}$ (**6**) as a blue solid. $^1H$ NMR ($CD_3OD$, 400 MHz) δ 8.07 (d, $J$ = 7.7 Hz, 1H), 7.68 (td, $J$ = 7.4, 1.2 Hz, 1H), 7.62 (td, $J$ = 7.5, 1.0 Hz, 1H), 7.43 (d, $J$ = 7.6 Hz, 1H), 7.14 (dd, $J$ = 14.6, 2.6 Hz, 2H), 6.97 (dd, $J$ = 9.0, 6.5 Hz, 2H), 6.58 (dd, $J$ = 8.9, 2.3 Hz, 2H), 4.17 (t, $J$ = 7.5 Hz, 8H), 2.49 (p, $J$ = 7.5 Hz, 4H); Analytical HPLC: $t_R$ = 9.5 min, >99% purity (10–95% MeCN/$H_2O$, linear gradient, with constant 0.1% v/v TFA additive; 20 min run; 1 mL/min flow; ESI; positive ion mode; detection at 675 nm); HRMS (ESI) calcd for $C_{26}H_{23}N_2O_4PNa$ $[M+Na]^+$ 481.1288, found 481.1292.

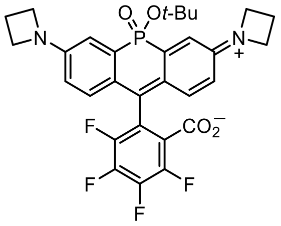

***tert*-Butyl-JF$_{690}$ (S49):** A solution of dibromide **S36** (200 mg, 0.369 mmol) in THF (8 mL) was cooled to −20 °C under nitrogen. Lithium dibutyl(isopropyl)magnesate (0.7 M in $Et_2O$/hexanes, 632 µL, 0.443 mmol, 1.2 eq) was added, and the reaction was stirred at −20 °C for 20 min. A solution of tetrafluorophthalic anhydride (**S47**; 292 mg, 1.33 mmol, 3.6 eq) in THF (4 mL) was added dropwise over 10 min, and the reaction was allowed to warm to room temperature over ~8 h. It was subsequently quenched with saturated $NH_4Cl$, diluted with water, and extracted with EtOAc (2×). The combined organic extracts were washed with saturated $NaHCO_3$ and brine, dried over anhydrous $MgSO_4$, filtered, and evaporated. Flash chromatography on silica gel (0–50% MeCN/$CH_2Cl_2$, linear gradient) afforded 38 mg (18%) of *tert*-butyl-JF$_{690}$ (**S49**) as an off-white solid. $^1H$ NMR ($CDCl_3$, 400 MHz) δ 7.02 (dd, $J$ = 14.5, 2.6 Hz, 2H), 6.88 (dd, $J$ = 8.8, 6.9 Hz, 2H), 6.48 (dd, $J$ = 8.8, 2.6 Hz, 2H), 4.03 – 3.91 (m, 8H), 2.41 (p, $J$ = 7.3 Hz, 4H), 1.59 (s, 9H); $^{19}F$ NMR ($CDCl_3$, 376 MHz) δ −138.63 (td, $J$ = 19.9, 8.8 Hz, 1F), −139.62 – −139.78 (m, 1F), −142.54 – −142.73 (m, 1F), −151.67 – −151.86 (m, 1F); Analytical HPLC: $t_R$ = 15.3 min, 97.1% purity (10–95% MeCN/$H_2O$, linear gradient, with constant 0.1% v/v TFA additive; 20 min run; 1 mL/min flow; ESI; positive ion mode; detection at 700 nm); HRMS (ESI) calcd for $C_{30}H_{27}F_4N_2O_4PNa$ $[M+Na]^+$ 609.1537, found 609.1546.

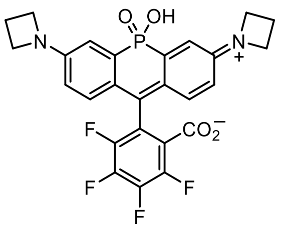

**$JF_{690}$ (16):** Phosphinate **S49** (27 mg, 46.0 μmol) was taken up in $CH_2Cl_2$ (2 mL); triethylsilane (200 μL) was added, followed by trifluoroacetic acid (200 μL). The reaction was stirred at room temperature for 1 h. Toluene (2 mL) was added, and the reaction mixture was concentrated to dryness. The crude material was purified by reverse phase HPLC (10–50% $MeCN/H_2O$, linear gradient, with constant 0.1% v/v TFA additive). The pooled HPLC product fractions were partially concentrated to remove MeCN and extracted with 15% *i*-PrOH/$CHCl_3$ (3×). The organic extracts were dried over anhydrous $MgSO_4$, filtered, and evaporated to yield 16 mg (66%) of $JF_{690}$ (**16**) as a dark blue solid. $^1H$ NMR (DMSO-$d_6$, 400 MHz) δ 6.89 (dd, $J$ = 8.9, 6.4 Hz, 2H), 6.87 (dd, $J$ = 14.0, 2.6 Hz, 2H), 6.43 (dd, $J$ = 8.7, 2.7 Hz, 2H), 3.92 (t, $J$ = 7.3 Hz, 8H), 2.36 (p, $J$ = 7.2 Hz, 4H); $^{19}F$ NMR (DMSO-$d_6$, 376 MHz) δ −73.16, −139.66 – −139.94 (m, 1F), −140.06 – −140.43 (m, 1F), −143.22 – −143.50 (m, 1F), −151.11 – −151.43 (m, 1F); Analytical HPLC: $t_R$ = 10.8 min, >99% purity (10–95% $MeCN/H_2O$, linear gradient, with constant 0.1% v/v TFA additive; 20 min run; 1 mL/min flow; ESI; positive ion mode; detection at 675 nm); HRMS (ESI) calcd for $C_{26}H_{19}F_4N_2O_4PNa$ $[M+Na]^+$ 553.0911, found 553.0921.

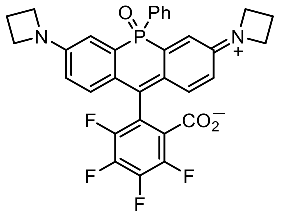

**$JF_{722}$ (17):** A solution of dibromide **S41** (500 mg, 0.915 mmol) in THF (100 mL) was cooled to −20 °C under nitrogen. Lithium dibutyl(isopropyl)magnesate (0.7 M in $Et_2O$/hexanes, 1.57 mL, 1.10 mmol, 1.2 eq) was added, and the reaction was stirred at −20 °C for 20 min. A solution of tetrafluorophthalic anhydride (**S47**; 725 mg, 3.30 mmol, 3.6 eq) in THF (5 mL) was added dropwise over 5 min, and the reaction was gradually warmed to 0 °C over 4 h. It was subsequently quenched with saturated $NH_4Cl$, diluted with water, and extracted with EtOAc (2×). The combined organic extracts were washed with saturated $NaHCO_3$ and brine, dried over anhydrous $MgSO_4$, filtered, and evaporated. The crude material was purified by silica gel chromatography (0–75% acetone/$CH_2Cl_2$, linear gradient) followed by reverse phase HPLC (10–95% $MeCN/H_2O$, linear gradient, with constant 0.1% v/v TFA additive). The pooled HPLC product fractions were partially concentrated to remove MeCN, diluted with saturated $NaHCO_3$, and extracted with $CH_2Cl_2$ (2×). The organic extracts were dried over anhydrous $MgSO_4$, filtered, and evaporated to yield 63 mg (12%) of $JF_{722}$ (**17**) as an off-white solid. $^1H$ NMR ($CDCl_3$, 400 MHz) δ 7.75 – 7.67 (m, 2H), 7.53 – 7.46 (m,

1H), 7.46 – 7.39 (m, 2H), 6.78 (dd, *J* = 13.8, 2.6 Hz, 2H), 6.77 (d, *J* = 8.8, 5.9 Hz, 2H), 6.45 (dd, *J* = 8.8, 2.5 Hz, 2H), 3.98 – 3.82 (m, 8H), 2.36 (p, *J* = 7.3 Hz, 4H); $^{19}$F NMR ($CDCl_3$, 376 MHz) δ −138.39 (td, *J* = 19.8, 9.0 Hz, 1F), −141.27 (td, *J* = 20.3, 3.9 Hz, 1F), −142.40 (ddd, *J* = 20.5, 18.4, 9.1 Hz, 1F), −150.60 – −150.76 (m, 1F); Analytical HPLC: $t_R$ = 12.2 min, >99% purity (10–95% $MeCN/H_2O$, linear gradient, with constant 0.1% v/v TFA additive; 20 min run; 1 mL/min flow; ESI; positive ion mode; detection at 725 nm); HRMS (ESI) calcd for $C_{32}H_{24}F_4N_2O_3P$ $[M+H]^+$ 591.1455, found 591.1464.

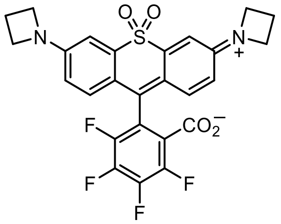


**$JF_{724}$ (18):** A solution of dibromide **S32** (800 mg, 1.65 mmol) in THF (125 mL) was cooled to −20 °C under nitrogen. Lithium dibutyl(isopropyl)magnesate (0.7 M in $Et_2O$/hexanes, 2.82 mL, 1.97 mmol, 1.2 eq) was added, and the reaction was stirred at -20 °C for 20 min. A solution of tetrafluorophthalic anhydride (**S47**; 1.30 g, 5.92 mmol, 3.6 eq) in THF (10 mL) was then added dropwise over 10 min. The reaction was gradually warmed to 0 °C over 4 h while stirring. It was subsequently quenched with saturated $NH_4Cl$, diluted with water, and extracted with EtOAc (2×). The combined organic extracts were washed with saturated $NaHCO_3$ and brine, dried over anhydrous $MgSO_4$, filtered, and evaporated. Flash chromatography on silica gel (0–40% EtOAc/hexanes, linear gradient, with constant 40% v/v $CH_2Cl_2$ additive) afforded 307 mg (35%) of $JF_{724}$ (**18**) as an off-white solid. $^1$H NMR ($CDCl_3$, 400 MHz) δ 7.09 (d, *J* = 2.5 Hz, 2H), 6.74 (d, *J* = 8.7 Hz, 2H), 6.40 (dd, *J* = 8.7, 2.5 Hz, 2H), 4.01 (t, *J* = 7.4 Hz, 8H), 2.44 (p, *J* = 7.3 Hz, 4H); $^{19}$F NMR ($CDCl_3$, 376 MHz) δ −137.58 (td, *J* = 20.0, 9.3 Hz), −138.15 (td, *J* = 20.0, 4.7 Hz), −141.77 (ddd, *J* = 20.6, 18.4, 9.2 Hz), −149.37 (ddd, *J* = 20.4, 18.4, 4.7 Hz); Analytical HPLC: $t_R$ = 15.7 min, 98.4% purity (10–95% $MeCN/H_2O$, linear gradient, with constant 0.1% v/v TFA additive; 20 min run; 1 mL/min flow; ESI; positive ion mode; detection at 280 nm); HRMS (ESI) calcd for $C_{26}H_{19}F_4N_2O_4S$ $[M+H]^+$ 531.0996, found 531.1001.

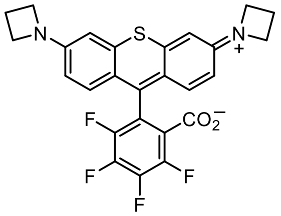


**$JF_{593}$ (20):** A solution of dibromide **S31** (2.00 g, 4.40 mmol) in THF (100 mL) was cooled to −78 °C under nitrogen. *tert*-Butyllithium (1.7 M in pentane, 11.40 mL, 19.4 mmol, 4.4 eq) was added, and the reaction was stirred at −78 °C for 30 min. It was then warmed to −20 °C, and a solution of tetrafluorophthalic anhydride (**S47**; 2.13 g, 9.69 mmol, 2.2 eq) in THF (25 mL) was added dropwise over 30 min via addition funnel. The reaction was allowed to warm to room temperature overnight (18 h). Following the addition of AcOH (1 mL), the mixture was diluted with MeOH, deposited onto Celite, and concentrated to dryness. Silica gel chromatography (0–10% $MeOH/CH_2Cl_2$, linear gradient, with constant 1% v/v AcOH additive; dry load with Celite) afforded 484 mg (20%) of the acetate salt of $JF_{593}$ (**20**) as a dark purple solid. $^1H$ NMR ($CD_3OD$, 400 MHz) δ 7.40 (d, $J$ = 9.3 Hz, 2H), 6.83 (d, $J$ = 2.3 Hz, 2H), 6.68 (dd, $J$ = 9.4, 2.3 Hz, 2H), 4.35 – 4.22 (m, 8H), 2.54 (p, $J$ = 7.6 Hz, 4H), 1.99 (s, 3H); $^{19}F$ NMR ($CD_3OD$, 376 MHz) δ −139.39 – −139.59 (m, 1F), −140.40 – −140.62 (m, 1F), −153.89 – −154.12 (m, 1F), −157.07 – −157.39 (m, 1F); Analytical HPLC: $t_R$ = 12.3 min, >99% purity (10–95% $MeCN/H_2O$, linear gradient, with constant 0.1% v/v TFA additive; 20 min run; 1 mL/min flow; ESI; positive ion mode; detection at 600 nm); HRMS (ESI) calcd for $C_{26}H_{19}F_4N_2O_2S$ $[M+H]^+$ 499.1098, found 499.1102.

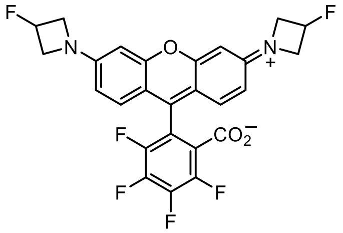

**$JF_{559}$ (43):** A solution of dibromide **S46** (500 mg, 1.05 mmol) in THF (20 mL) was cooled to −78 °C under nitrogen. *tert*-Butyllithium (1.7 M in pentane, 2.73 mL, 4.64 mmol, 4.4 eq) was added, and the reaction was stirred at −78 °C for 30 min. It was then warmed to −10 °C before adding a solution of $MgBr_2 \cdot OEt_2$ (599 mg, 2.32 mmol, 2.2 eq) in THF (10 mL). After an additional 30 min at −10 °C, a solution of tetrafluorophthalic anhydride (**S47**; 511 mg, 2.32 mmol, 2.2 eq) in THF (10 mL) was added dropwise over 30 min via addition funnel. The reaction was then allowed to warm to room temperature overnight (18 h). Following the addition of AcOH (100 μL), the mixture was diluted with MeOH, deposited onto Celite, and concentrated to dryness. Silica gel chromatography (0–10% MeOH (2 M $NH_3$)/$CH_2Cl_2$, linear gradient; dry load with Celite) afforded 177 mg (32%) of $JF_{559}$ (**43**) as a dark red-purple solid. $^1H$ NMR ($CD_3OD$, 400 MHz) δ 7.37 (dd, $J$ = 9.1, 0.5 Hz, 2H), 6.72 (dd, $J$ = 9.2, 2.2 Hz, 2H), 6.58 (d, $J$ = 2.2 Hz, 2H), 5.56 (dtt, $^2J_{HF}$ = 57.0 Hz, $J$ = 6.0, 3.0 Hz, 2H), 4.66 – 4.53 (m, 4H), 4.44 – 4.30 (m, 4H); $^{19}F$ NMR ($CD_3OD$, 376 MHz) δ −139.12 (ddd, $J$ = 21.0, 12.7, 3.9 Hz, 1F), −140.70 (ddd, $J$ = 22.1, 12.8, 4.0 Hz, 1F), −153.21 (ddd, $J$ = 22.9, 19.1, 4.2 Hz, 1F), −157.02 – −157.18 (m, 1F), −180.54 (dtt, $J_{FH}$ = 57.1, 23.7, 20.4 Hz, 2F); Analytical HPLC: $t_R$ = 11.1 min, >99% purity (10–95% $MeCN/H_2O$, linear gradient, with constant 0.1% v/v TFA additive; 20 min run; 1 mL/min flow; ESI; positive ion mode; detection at 575 nm); HRMS (ESI) calcd for $C_{26}H_{17}F_6N_2O_3$ $[M+H]^+$ 519.1138, found 519.1138.

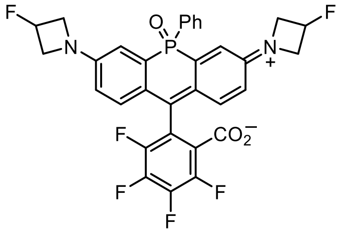

**$JF_{711}$ (45):** A solution of dibromide **S42** (1.35 g, 2.32 mmol) in THF (90 mL) was cooled to −20 °C under nitrogen. Lithium dibutyl(isopropyl)magnesate (0.7 M in $Et_2O$/hexanes, 3.97 mL, 2.78 mmol, 1.2 eq) was added, and the reaction was stirred at −20 °C for 20 min. A solution of tetrafluorophthalic anhydride (**S47**; 1.84 g, 8.35 mmol, 3.6

eq) in THF (10 mL) was added dropwise over 5 min, and the reaction was gradually warmed to 0 °C over 4 h. It was subsequently quenched with saturated $NH_4Cl$, diluted with water, and extracted with EtOAc (2×). The combined organic extracts were washed with saturated $NaHCO_3$ and brine, dried over anhydrous $MgSO_4$, filtered, and evaporated. The crude was purified twice by silica gel chromatography (0–50% acetone/$CH_2Cl_2$, linear gradient; then, 25–100% EtOAc/hexanes, linear gradient) to afford 223 mg (15%) of $JF_{711}$ (**45**) as an off-white solid. $^1$H NMR ($CDCl_3$, 400 MHz) δ 7.76 – 7.66 (m, 2H), 7.56 – 7.49 (m, 1H), 7.49 – 7.41 (m, 2H), 6.83 (dd, $J$ = 13.6, 2.7 Hz, 2H), 6.82 (dd, $J$ = 8.9, 5.8 Hz, 2H), 6.51 (dd, $J$ = 8.9, 2.6 Hz, 2H), 5.38 (dtt, $^2J_{HF}$ = 56.7 Hz, $J$ = 6.1, 3.3 Hz, 2H), 4.29 – 4.11 (m, 4H), 4.09 – 3.89 (m, 4H); $^{19}$F NMR ($CDCl_3$, 376 MHz) δ −137.88 (td, $J$ = 19.8, 9.2 Hz, 1F), −141.19 (td, $J$ = 20.7, 4.1 Hz, 1F), −141.89 (ddd, $J$ = 20.5, 18.5, 9.3 Hz, 1F), −150.00 – −150.17 (m, 1F), −180.55 (dtt, $J_{FH}$ = 56.6, 23.4, 18.8 Hz, 2F); Analytical HPLC: $t_R$ = 12.3 min, 97.6% purity (30–95% MeCN/$H_2O$, linear gradient, with constant 0.1% v/v TFA additive; 20 min run; 1 mL/min flow; ESI; positive ion mode; detection at 254 nm); HRMS (ESI) calcd for $C_{32}H_{22}F_6N_2O_3P$ $[M+H]^+$ 627.1267, found 627.1276.

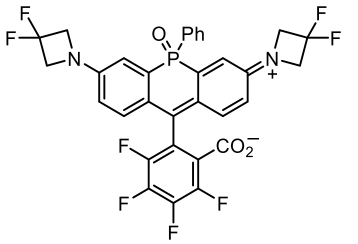


**2-(3,7-Bis(3,3-difluoroazetidin-1-yl)-5-oxido-5-phenyl-10*H*-acridophosphin-10-ylium-10-yl)-3,4,5,6-tetrafluorobenzoate (S50):** A solution of dibromide **S43** (500 mg, 0.809 mmol) in THF (30 mL) was cooled to −20 °C under nitrogen. Lithium dibutyl(isopropyl)magnesate (0.7 M in $Et_2O$/hexanes, 1.39 mL, 0.971 mmol, 1.2 eq) was added, and the reaction was stirred at −20 °C for 20 min. A solution of tetrafluorophthalic anhydride (**S47**; 641 mg, 2.91 mmol, 3.6 eq) in THF (5 mL) was added dropwise over 5 min, and the reaction was gradually warmed to 0 °C over 4 h. It was subsequently quenched with saturated $NH_4Cl$, diluted with water, and extracted with EtOAc (2×). The combined organic extracts were washed with saturated $NaHCO_3$ and brine, dried over anhydrous $MgSO_4$, filtered, and evaporated. Flash chromatography on silica gel (0–100% EtOAc/toluene, linear gradient) afforded 172 mg (32%) of **S50** as an off-white solid. $^1$H NMR ($CDCl_3$, 400 MHz) δ 7.76 – 7.67 (m, 2H), 7.59 – 7.52 (m, 1H), 7.52 – 7.44 (m, 2H), 6.90 (dd, $J$ = 14.0, 2.6 Hz, 2H), 6.88 (dd, $J$ = 9.0, 5.6 Hz, 2H), 6.58 (dd, $J$ = 8.8, 2.7 Hz, 2H), 4.34 – 4.16 (m, 8H); $^{19}$F NMR ($CDCl_3$, 376 MHz) δ −100.01 (p, $J$ = 11.6 Hz, 4F), −137.35 (td, $J$ = 19.9, 9.7 Hz, 1F), −141.18 (td, $J$ = 20.5, 19.7, 4.1 Hz, 1F), −141.46 (ddd, $J$ = 20.8, 18.4, 9.6 Hz, 1F), −149.46 – −149.62 (m, 1F); Analytical HPLC: $t_R$ = 13.0 min, >99% purity (30–95% MeCN/$H_2O$, linear gradient, with constant 0.1% v/v TFA additive; 20 min run; 1 mL/min flow; ESI; positive ion mode; detection at 254 nm); HRMS (ESI) calcd for $C_{32}H_{20}F_8N_2O_3P$ $[M+H]^+$ 663.1078, found 663.1088.

### AZA-RHODAMINE HALOTAG LIGANDS

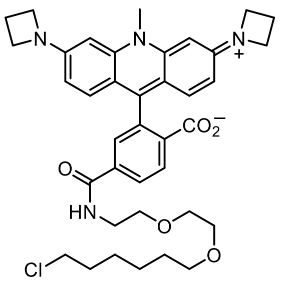

**$JF_{502}$–HaloTag ligand (S24):** A suspension of acid **S22** (acetate salt; 25 mg, 47.4 μmol) in DMF (10 mL) was heated to 50 °C. DIEA (41.3 μL, 0.237 mmol, 5 eq) and HBTU (27.0 mg, 71.1 μmol, 1.5 eq) were added, and the reaction was stirred for 10 min at 50 °C. A solution of HaloTag(O2)amine (HTL-$NH_2$, **36**; TFA salt; 24.0 mg, 71.1 μmol, 1.5 eq) in DMF (500 μL) was then added. After stirring the reaction for 1 h at 50 °C, it was cooled to room temperature and concentrated to remove DMF. Reverse phase HPLC of the crude material (30–60% MeCN/$H_2O$, linear gradient, with constant 0.1% v/v TFA additive) afforded $JF_{502}$–HaloTag ligand (**S24**) as an orange solid (11.2 mg, 30%, TFA salt). $^1H$ NMR ($CD_3OD$, 400 MHz) δ 8.77 (t, $J$ = 5.2 Hz, 1H), 8.39 (d, $J$ = 8.3 Hz, 1H), 8.19 (dd, $J$ = 8.2, 1.8 Hz, 1H), 7.75 (d, $J$ = 1.8 Hz, 1H), 7.19 (d, $J$ = 9.2 Hz, 2H), 6.69 (dd, $J$ = 9.3, 2.0 Hz, 2H), 6.47 (d, $J$ = 2.0 Hz, 2H), 4.27 (t, $J$ = 7.7 Hz, 8H), 4.12 (s, 3H), 3.69 – 3.54 (m, 8H), 3.52 (t, $J$ = 6.6 Hz, 2H), 3.41 (t, $J$ = 6.5 Hz, 2H), 2.55 (p, $J$ = 7.5 Hz, 4H), 1.75 – 1.66 (m, 2H), 1.52 – 1.44 (m, 2H), 1.43 – 1.27 (m, 4H); Analytical HPLC: $t_R$ = 12.5 min, >99% purity (10–95% MeCN/$H_2O$, linear gradient, with constant 0.1% v/v TFA additive; 20 min run; 1 mL/min flow; ESI; positive ion mode; detection at 500 nm); HRMS (ESI) calcd for $C_{38}H_{46}ClN_4O_5$ $[M+H]^+$ 673.3151, found 673.3158.

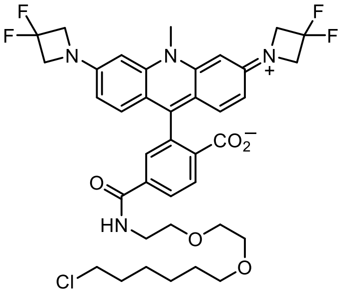

**$JF_{479}$–HaloTag ligand (12):** A suspension of acid **S23** (25 mg, 46.3 μmol) in DMF (10 mL) was heated to 50 °C. DIEA (40.4 μL, 0.232 mmol, 5 eq) and HBTU (21.1 mg, 55.6 μmol, 1.2 eq) were added, and the reaction was stirred for 15 min at 50 °C. A solution of HaloTag(O2)amine (HTL-$NH_2$, **36**; TFA salt; 18.8 mg, 55.6 μmol, 1.2 eq) in DMF (500 μL) was then added. After stirring the reaction for 2 h at 50 °C, it was cooled to room temperature and concentrated to remove DMF. Reverse phase HPLC of the crude material (35–55% MeCN/$H_2O$, linear gradient, with constant 0.1% v/v TFA additive) afforded $JF_{479}$–HaloTag ligand (**12**) as a yellow-orange solid (11.4 mg, 29%, TFA salt). $^1H$ NMR ($CD_3OD$, 400 MHz) δ 8.76 (t, $J$ = 5.3 Hz, 1H), 8.43 (d, $J$ = 8.2 Hz, 1H), 8.22 (dd, $J$ = 8.2, 1.8 Hz, 1H), 7.79 (d, $J$ = 1.8 Hz, 1H), 7.36 (d, $J$ = 9.2 Hz, 2H), 6.89 (dd, $J$ = 9.2, 2.1 Hz, 2H), 6.85 (d, $J$ = 2.0 Hz, 2H), 4.67 (t, $^3J_{HF}$ = 11.7 Hz, 8H), 4.34 (s, 3H), 3.69 – 3.54 (m, 8H), 3.52 (t, $J$ = 6.6 Hz, 2H), 3.43 (t, $J$ = 6.5 Hz, 2H), 1.76 – 1.67 (m,

2H), 1.55 – 1.46 (m, 2H), 1.45 – 1.28 (m, 4H); $^{19}$F NMR ($CD_3OD$, 376 MHz) δ −75.26 (s, 3F), −100.72 (p, $^3J_{FH}$ = 12.0 Hz, 4F); Analytical HPLC: $t_R$ = 12.6 min, >99% purity (10–95% MeCN/$H_2O$, linear gradient, with constant 0.1% v/v TFA additive; 20 min run; 1 mL/min flow; ESI; positive ion mode; detection at 475 nm); HRMS (ESI) calcd for $C_{38}H_{42}ClF_4N_4O_5$ $[M+H]^+$ 745.2774, found 745.2787.

### MAC SUBSTITUTION OF 4,5,6,7-TETRAFLUORORHODAMINES

Given the generality of the pendant aryl ring fluorination as a strategy to tune the lactone–zwitterion equilibria and spectral properties of various rhodamine dyes, we then sought a convenient method to install the functionality necessary for labeling with and bioconjugation of 4,5,6,7-tetrafluororhodamines. It had been well-established that thiols undergo efficient substitution reactions with 4,5,6,7-tetrafluoroxanthene fluorophores—including fluoresceins and rhodamines—to form aryl sulfides at the 6-position.[2,3] Nevertheless, the *general* nucleophile scope for the $S_NAr$ of fluorinated xanthenes was less explored. We aimed to briefly investigate the reactivity of fluorinated rhodamines as substrates for nucleophilic aromatic substitution with a variety of common nucleophiles, including carbon-centered anions (*e.g.*, malonates) and heteroatom-based species (*e.g.*, azide, amine; see **Substitution of 4,5,6,7-Tetrafluororhodamines with Other Nucleophiles**).

Our overriding interest, however, was to develop an efficient approach to install a carboxyl group—or related functional surrogate—onto the fluorinated aryl ring. This would allow for the application of more traditional methods of bioconjugation, which typically involve the formation of an amide through a 5- or 6-carboxy substituent on the xanthene fluorophore. More specifically, it would enable the synthesis of self-labeling tag ligands (*e.g.*, HaloTag ligands) directly comparable to the existing des-fluorinated analogs, which already demonstrate ideal, optimized labeling kinetics. We required, therefore, a mild transformation that substituted one fluoride substituent with a carboxyl group; this suggested the use of an *umpolung*-type acyl anion equivalent. The elegant (albeit underused) masked acyl cyanide (MAC) chemistry developed by Nemoto and coworkers seemed most promising.[4,5]

With this in mind, the known MOM-protected MAC reagent **34** (2-(methoxymethoxy)malononitrile)[6,7] was reacted with a selection of 4,5,6,7-tetrafluororhodamines, including the yellow/orange oxygen- (**19**, **43**) and sulfide- (**20**) derivatives, the red silicon analogs (**15**, **S51**), and the near-IR sulfone (**18**) and phosphine oxide (**17**, **45**) dyes (**Fig. 2, Schemes S3–S4**).[1] In the presence of an amine base (DIEA), each rhodamine underwent clean displacement of one fluoride substituent with **34** to provide a single regioisomeric substitution product in moderate to good yield. Although we expected—based on the thiol precedent and the probable $S_NAr$ mechanism—that the substitution was most likely occurring at the position *para* to the lactone (*i.e.*, the 6-position), the regiochemistry of the reaction was definitively confirmed by single crystal X-ray diffraction (see **X-Ray Crystallography**). Compound **S57** was scaled up for structural determination by SC-XRD simply out of convenience, as it entailed the shortest synthetic route.

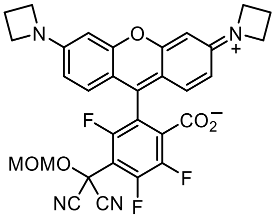


**6-(MOM-MAC)-JF$_{571}$ (S56):** JF$_{571}$[1] (**19**; 155 mg, 0.321 mmol) and 2-(methoxymethoxy)malononitrile (**34**; 40.5 mg, 0.321 mmol, 1 eq) were combined in DMF (5 mL), and DIEA (112 μL, 0.643 mmol, 2 eq) was added. After stirring the reaction at room temperature for 2 h, it was evaporated to dryness. Flash chromatography on silica gel (0–15% MeOH/$CH_2Cl_2$, linear gradient, with constant 1% v/v AcOH additive) afforded **S56** as a dark red-purple solid (108 mg, 52%, acetate salt). $^{1}$H NMR ($CDCl_3$, 400 MHz) δ 7.30 (d, *J* = 9.1 Hz, 2H), 6.46 (dd, *J* = 9.1, 2.2 Hz, 2H), 6.27 (d, *J* = 2.2 Hz, 2H), 5.15 (s, 2H), 4.22 (t, *J* = 7.6 Hz, 8H), 3.55 (s, 3H), 2.56 (p, *J* = 7.6 Hz, 4H), 2.05 (s, 3H); $^{19}$F NMR ($CDCl_3$, 376 MHz) δ −114.38 (d, *J* = 16.5 Hz, 1F), −127.29 (d, *J* = 22.6 Hz, 1F), −138.86 (dd, *J* = 22.5, 16.6 Hz, 1F); Analytical HPLC: $t_R$ = 10.4 min, 99.0% purity (10–95% MeCN/$H_2O$, linear gradient, with constant 0.1% v/v TFA additive; 20 min run; 1 mL/min flow; ESI; positive ion mode; detection at 575 nm); HRMS (ESI) calcd for $C_{31}H_{24}F_3N_4O_5$ $[M+H]^+$ 589.1693, found 589.1698.

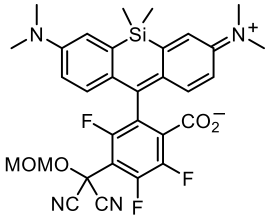


**4,5,7-Trifluoro-6-(MOM-MAC)-SiTMR (S57):** 4,5,6,7-Tetrafluoro-SiTMR[1] (**S51**; 1.50 g, 3.00 mmol) and 2-(methoxymethoxy)malononitrile (**34**; 378 mg, 3.00 mmol, 1 eq) were combined in DMF (30 mL), and DIEA (1.04 mL, 5.99 mmol, 2 eq) was added. After stirring the reaction at room temperature for 1 h, it was evaporated to dryness. Flash chromatography on silica gel (10–100% EtOAc/hexanes, linear gradient) afforded **S57** as a blue-green solid (1.17 g, 64%). $^{1}$H NMR ($CDCl_3$, 400 MHz) δ 6.94 (d, *J* = 2.9 Hz, 2H), 6.74 (d, *J* = 8.8 Hz, 2H), 6.62 (dd, *J* = 8.9, 2.9 Hz, 2H), 5.16 (s, 2H), 3.53 (s, 3H), 3.00 (s, 12H), 0.59 (s, 3H), 0.56 (s, 3H); $^{19}$F NMR ($CDCl_3$, 376 MHz) δ −113.66 (d, *J* = 22.7 Hz, 1F), −127.80 (d, *J* = 20.4 Hz, 1F), −139.94 (dd, *J* = 22.6, 20.3 Hz, 1F); Analytical HPLC: $t_R$ = 12.5 min, >99% purity (10–95% MeCN/$H_2O$, linear gradient, with constant 0.1% v/v TFA additive; 20 min run; 1 mL/min flow; ESI; positive ion mode; detection at 675 nm); HRMS (ESI) calcd for $C_{31}H_{30}F_3N_4O_4Si$ $[M+H]^+$ 607.1983, found 607.1990.

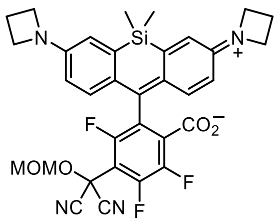


**6-(MOM-MAC)-$JF_{669}$ (35):** $JF_{669}$[1] (**15**; 72 mg, 0.137 mmol) and DIEA (47.8 µL, 0.275 mmol, 2 eq) were combined in DMF (3 mL), and a solution of 2-(methoxymethoxy)malononitrile (**34**; 17.3 mg, 0.137 mmol, 1 eq) in DMF (1 mL) was added dropwise. After stirring the reaction at room temperature for 4 h, it was concentrated to dryness. Flash chromatography on silica gel (10–100% EtOAc/hexanes, linear gradient) afforded 44.4 mg (51%) of **35** as a green foam. $^1$H NMR ($CDCl_3$, 400 MHz) δ 6.70 (d, $J$ = 8.7 Hz, 2H), 6.64 (d, $J$ = 2.6 Hz, 2H), 6.32 (dd, $J$ = 8.7, 2.6 Hz, 2H), 5.16 (s, 2H), 3.93 (t, $J$ = 7.2 Hz, 8H), 3.53 (s, 3H), 2.39 (p, $J$ = 7.3 Hz, 4H), 0.55 (s, 3H), 0.53 (s, 3H); $^{19}$F NMR ($CDCl_3$, 376 MHz) δ −113.41 (d, $J$ = 22.6 Hz, 1F), −127.72 (d, $J$ = 20.3 Hz, 1F), −139.78 (dd, $J$ = 22.7, 20.2 Hz, 1F); Analytical HPLC: $t_R$ = 13.0 min, >99% purity (10–95% MeCN/$H_2O$, linear gradient, with constant 0.1% v/v TFA additive; 20 min run; 1 mL/min flow; ESI; positive ion mode; detection at 675 nm); HRMS (ESI) calcd for $C_{33}H_{30}F_3N_4O_4Si$ $[M+H]^+$ 631.1983, found 631.1989.

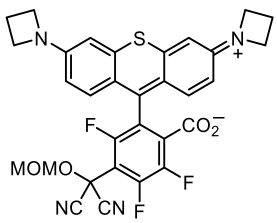


**6-(MOM-MAC)-$JF_{593}$ (S67):** $JF_{593}$ (**20**, acetate salt; 250 mg, 0.448 mmol) and 2-(methoxymethoxy)malononitrile (**34**; 56 mg, 0.448 mmol, 1 eq) were combined in DMF (10 mL), and DIEA (234 µL, 1.34 mmol, 3 eq) was added. After stirring the reaction at room temperature for 3 h, it was concentrated *in vacuo*. The residue was redissolved in MeOH/$CH_2Cl_2$, deposited onto Celite, and evaporated to dryness. Flash chromatography on silica gel (0–15% MeOH/$CH_2Cl_2$, linear gradient, with constant 1% v/v AcOH additive; dry load with Celite) afforded **S67** as a dark purple solid (164 mg, 55%, acetate salt). $^1$H NMR ($CD_3OD$, 400 MHz) δ 7.38 (d, $J$ = 9.4 Hz, 2H), 6.87 (d, $J$ = 2.2 Hz, 2H), 6.71 (dd, $J$ = 9.4, 2.3 Hz, 2H), 5.21 (s, 2H), 4.31 (t, $J$ = 7.7 Hz, 8H), 3.53 (s, 3H), 2.55 (p, $J$ = 7.7 Hz, 4H); $^{19}$F NMR ($CD_3OD$, 376 MHz) δ −114.32 (d, $J$ = 15.0 Hz, 1F), −130.76 (d, $J$ = 21.5 Hz, 1F), −143.57 (dd, $J$ = 21.8, 15.2 Hz, 1F); Analytical HPLC: $t_R$ = 12.2 min, >99% purity (10–95% MeCN/$H_2O$, linear gradient, with constant 0.1% v/v TFA additive; 20 min run; 1 mL/min flow; ESI; positive ion mode; detection at 600 nm); HRMS (ESI) calcd for $C_{31}H_{24}F_3N_4O_4S$ $[M+H]^+$ 605.1465, found 605.1464.

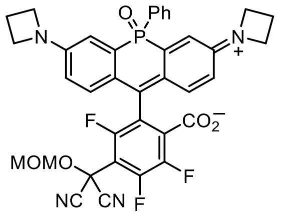

**6-(MOM-MAC)-JF$_{722}$ (S68):** JF$_{722}$ (**17**; 145 mg, 0.246 mmol) and 2-(methoxymethoxy)malononitrile (**34**; 31.0 mg, 0.246 mmol, 1 eq) were combined in DMF (8 mL), and DIEA (85.5 µL, 0.491 mmol, 2 eq) was added. After stirring the reaction at room temperature for 2 h, it was evaporated to dryness. Flash chromatography on silica gel (0–75% acetone/$CH_2Cl_2$, linear gradient) afforded **S68** as a pale yellow solid (75.4 mg, 44%). $^1$H NMR ($CDCl_3$, 400 MHz) δ 7.74 – 7.65 (m, 2H), 7.51 – 7.37 (m, 3H), 6.83 (dd, $J$ = 13.7, 2.6 Hz, 2H), 6.76 (dd, $J$ = 8.8, 5.9 Hz, 2H), 6.47 (dd, $J$ = 8.8, 2.6 Hz, 2H), 5.10 (s, 2H), 4.01 – 3.83 (m, 8H), 3.48 (s, 3H), 2.37 (p, $J$ = 7.3 Hz, 4H); $^{19}$F NMR ($CDCl_3$, 376 MHz) δ −116.06 (d, $J$ = 22.6 Hz, 1F), −126.27 (d, $J$ = 20.2 Hz, 1F), −138.85 (dd, $J$ = 22.5, 20.3 Hz, 1F); Analytical HPLC: $t_R$ = 13.8 min, >99% purity (10–95% MeCN/$H_2O$, linear gradient, with constant 0.1% v/v TFA additive; 20 min run; 1 mL/min flow; ESI; positive ion mode; detection at 725 nm); HRMS (ESI) calcd for $C_{37}H_{29}F_3N_4O_5P$ [M+H]$^+$ 697.1822, found 697.1825.

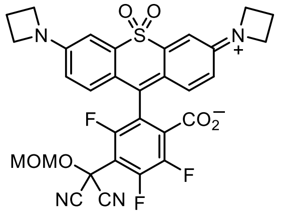

**6-(MOM-MAC)-JF$_{724}$ (S69):** To a solution of JF$_{724}$ (**18**; 150 mg, 0.283 mmol) in DMF (5 mL) were added DIEA (99 µL, 0.566 mmol, 2 eq) and 2-(methoxymethoxy)malononitrile (**34**; 35.7 mg, 0.283 mmol, 1 eq). After stirring the reaction at room temperature for 2 h, it was evaporated to dryness. Flash chromatography on silica gel (0–50% EtOAc/hexanes, linear gradient, with constant 40% v/v $CH_2Cl_2$ additive) afforded **S69** as a yellow-green solid (76.4 mg, 42%). $^1$H NMR ($CDCl_3$, 400 MHz) δ 7.10 (d, $J$ = 2.5 Hz, 2H), 6.67 (d, $J$ = 8.7 Hz, 2H), 6.40 (dd, $J$ = 8.7, 2.5 Hz, 2H), 5.17 (s, 2H), 4.02 (t, $J$ = 7.4 Hz, 8H), 3.54 (s, 3H), 2.45 (p, $J$ = 7.4 Hz, 4H); $^{19}$F NMR ($CDCl_3$, 376 MHz) δ −112.54 (d, $J$ = 22.8 Hz, 1F), −124.88 (d, $J$ = 20.2 Hz, 1F), −138.16 (dd, $J$ = 23.0, 20.3 Hz, 1F); Analytical HPLC: $t_R$ = 13.0 min, >99% purity (10–95% MeCN/$H_2O$, linear gradient, with constant 0.1% v/v TFA additive; 20 min run; 1 mL/min flow; ESI; positive ion mode; detection at 725 nm); HRMS (ESI) calcd for $C_{31}H_{24}F_3N_4O_6S$ [M+H]$^+$ 637.1363, found 637.1365.

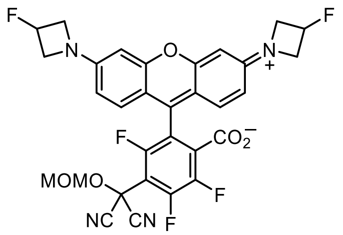

**6-(MOM-MAC)-JF$_{559}$ (S70):** JF$_{559}$ (**43**; 250 mg, 0.482 mmol) and 2-(methoxymethoxy)malononitrile (**34**; 60.8 mg, 0.482 mmol, 1 eq) were combined in DMF (10 mL), and DIEA (168 µL, 0.964 mmol, 2 eq) was added. After stirring the reaction at room temperature for 3 h, it was concentrated *in vacuo*. The crude material was purified by silica gel chromatography (0–15% MeOH/$CH_2Cl_2$, linear gradient, with constant 1% v/v AcOH additive) followed by reverse phase HPLC (10–75% MeCN/$H_2O$, linear gradient, with constant 0.1% v/v TFA additive). The pooled HPLC product fractions were partially concentrated to remove MeCN, diluted with saturated $NaHCO_3$, and extracted with $CH_2Cl_2$ (2×). The organic extracts were dried over anhydrous $MgSO_4$, filtered, and evaporated to yield 153 mg (51%) of **S70** as a dark red-purple solid. $^1$H NMR ($CDCl_3$, 400 MHz) δ 6.75 (d, $J$ = 8.6 Hz, 2H), 6.26 (d, $J$ = 2.3 Hz, 2H), 6.22 (dd, $J$ = 8.6, 2.3 Hz, 2H), 5.45 (dtt, $^2J_{HF}$ = 56.7 Hz, $J$ = 6.1, 3.5 Hz, 2H), 5.14 (s, 2H), 4.31 – 4.19 (m, 4H), 4.12 – 4.00 (m, 4H), 3.52 (s, 3H); $^{19}$F NMR ($CDCl_3$, 376 MHz) δ −116.80 (d, $J$ = 22.5 Hz, 1F), −126.23 (d, $J$ = 20.3 Hz, 1F), −139.30 (dd, $J$ = 22.5, 20.5 Hz, 1F), −180.63 (dtt, $J_{FH}$ = 56.6, 23.8, 18.6 Hz, 2F); Analytical HPLC: $t_R$ = 11.9 min, 97.2% purity (10–95% MeCN/$H_2O$, linear gradient, with constant 0.1% v/v TFA additive; 20 min run; 1 mL/min flow; ESI; positive ion mode; detection at 575 nm); HRMS (ESI) calcd for $C_{31}H_{22}F_5N_4O_5$ $[M+H]^+$ 625.1505, found 625.1514.

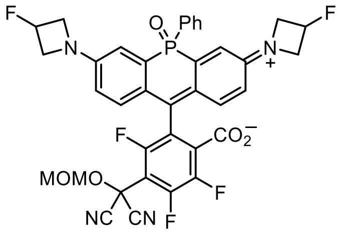

**6-(MOM-MAC)-JF$_{711}$ (S71):** JF$_{711}$ (**45**; 190 mg, 0.303 mmol) and 2-(methoxymethoxy)malononitrile (**34**; 38.3 mg, 0.303 mmol, 1 eq) were combined in DMF (6 mL), and DIEA (106 µL, 0.607 mmol, 2 eq) was added. After stirring the reaction at room temperature for 2 h, it was concentrated *in vacuo*. The crude material was purified by silica gel chromatography (0–40% acetone/$CH_2Cl_2$, linear gradient) followed by reverse phase HPLC (20–80% MeCN/$H_2O$, linear gradient, with constant 0.1% v/v TFA additive). The pooled HPLC product fractions were partially concentrated to remove MeCN, diluted with saturated $NaHCO_3$, and extracted with $CH_2Cl_2$ (2×). The organic extracts were dried over anhydrous $MgSO_4$, filtered, and evaporated to yield 88 mg (40%) of **S71** as a blue-green solid. $^1$H NMR ($CDCl_3$, 400 MHz) δ 7.74 – 7.65 (m, 2H), 7.54 – 7.48 (m, 1H), 7.47 – 7.40 (m, 2H), 6.88 (dd, $J$ = 13.6, 2.6 Hz, 2H), 6.81 (dd, $J$ = 8.8, 5.8 Hz, 2H), 6.54 (dd, $J$ = 8.7, 2.6 Hz, 2H), 5.40 (dtt, $^2J_{HF}$ = 56.8 Hz, $J$ = 6.3, 3.5 Hz, 2H), 5.10 (s, 2H), 4.31 – 4.14 (m, 4H), 4.13 – 3.92 (m, 4H), 3.49 (s, 3H); $^{19}$F NMR ($CDCl_3$, 376 MHz) δ −116.15 (d, $J$ = 22.3 Hz, 1F), −125.67 (d, $J$ = 20.1 Hz, 1F), −138.35 (dd, $J$ = 22.6, 20.1 Hz, 1F), −180.54 (dtt, $J_{FH}$ = 56.5, 23.7, 18.9 Hz, 2F); Analytical HPLC: $t_R$ = 14.5 min, 97.3% purity (10–95% MeCN/$H_2O$, linear gradient, with constant 0.1% v/v TFA additive; 20

min run; 1 mL/min flow; ESI; positive ion mode; detection at 280 nm); HRMS (ESI) calcd for $C_{37}H_{27}F_5N_4O_5P$ $[M+H]^+$ 733.1634, found 733.1651.

### CONVERSION OF MAC SUBSTITUTION PRODUCTS TO AMIDES AND ESTERS

With the MAC-rhodamine products in hand, we then applied a variation of the protocols previously described by Nemoto[4,5] and Rawal[7] to convert these masked acyl groups into amides and esters (**Fig. 2a**, **Supplementary Fig. 3b**, **Scheme S4**). Removal of the MOM protecting group on the MAC oxygen is achieved with TFA to yield a gem-dicyano alcohol. Following the removal of acid and solvent, this crude intermediate is reacted with an amine or alcohol in the presence of TEA or DIEA to directly afford an amide or ester. This conversion is presumed to proceed through an acyl cyanide (an activated ester equivalent) formed through a base-mediated cyanide elimination (**Supplementary Fig. 3b**). This approach permitted direct access to the HaloTag ligands without formation and isolation of an intermediate 6-carboxyrhodamine. Although not detailed here this transformation is quite general in scope and allows for the efficient amidation of fluorinated rhodamines with large and functionally diverse amines via this activated acyl cyanide intermediate. We also note that the structure of ester **S79** as determined by SC-XRD was consistent with the regiochemistry confirmed by X-ray diffraction of its precursor, MAC product **S57** (see **X-Ray Crystallography**).

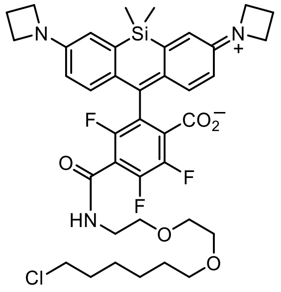

**$JF_{669}$–HaloTag ligand (37):** Ether **35** (151 mg, 0.239 mmol) was taken up in $CH_2Cl_2$ (10 mL); triethylsilane (1 mL) was added, followed by trifluoroacetic acid (2 mL). The reaction was stirred at room temperature for 6 h. Toluene (10 mL) was added, and the reaction mixture was concentrated to dryness. The residue was combined with a premixed solution of HaloTag(O2)amine (HTL-$NH_2$, **36**; TFA salt; 162 mg, 0.479 mmol, 2 eq) and DIEA (417 µL, 2.39 mmol, 10 eq) in $CH_2Cl_2$ (10 mL), and the reaction was stirred at room temperature for 18 h. The solvent was removed by rotary evaporation, and the crude material was purified by reverse phase HPLC (10–95% MeCN/$H_2O$, linear gradient, with constant 0.1% v/v TFA additive). The pooled HPLC product fractions were partially concentrated to remove MeCN, diluted with saturated $NaHCO_3$, and extracted with $CH_2Cl_2$ (2×). The organic extracts were dried over anhydrous $MgSO_4$, filtered, and evaporated to yield 114 mg (63%) of $JF_{669}$–HaloTag ligand (**37**) as a blue solid. $^1H$ NMR ($CDCl_3$, 400 MHz) δ 6.78 (d, $J$ = 8.6 Hz, 2H), 6.74 (bs, 1H), 6.63 (d, $J$ = 2.7 Hz, 2H), 6.32 (dd, $J$ = 8.7, 2.6 Hz, 2H), 3.92 (t, $J$ = 7.3 Hz, 8H), 3.66 – 3.59 (m, 6H), 3.55 – 3.48 (m, 4H), 3.37 (t, $J$ = 6.7 Hz, 2H), 2.38 (p, $J$ = 7.3 Hz, 4H), 1.78 – 1.69 (m, 2H), 1.53 – 1.45 (m, 2H), 1.43 – 1.25 (m, 4H), 0.53 (s, 6H); $^{19}F$ NMR ($CDCl_3$, 376 MHz) δ −118.50 (d, $J$ = 22.5 Hz, 1F), −133.47 (d, $J$ = 21.6 Hz, 1F), −142.24 (t, $J$ = 22.1 Hz, 1F); Analytical HPLC: $t_R$ = 13.6 min, >99% purity (10–95% MeCN/$H_2O$, linear gradient, with constant 0.1% v/v TFA additive; 20 min run; 1 mL/min flow; ESI; positive ion mode; detection at 675 nm); HRMS (ESI) calcd for $C_{39}H_{46}ClF_3N_3O_5Si$ $[M+H]^+$ 756.2842, found 756.2856.

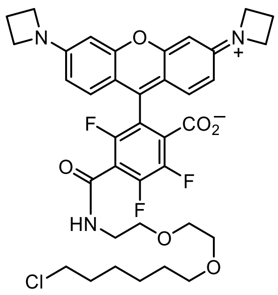

**$JF_{571}$–HaloTag ligand (38):** Ether **S56** (50 mg, 77.1 μmol) was taken up in $CH_2Cl_2$ (5 mL); triethylsilane (500 μL) was added, followed by trifluoroacetic acid (1 mL). The reaction was stirred at room temperature for 6 h. Toluene (5 mL) was added, and the reaction mixture was concentrated to dryness. The residue was combined with a premixed solution of HaloTag(O2)amine (HTL-$NH_2$, **36**; TFA salt; 52.1 mg, 0.154 mmol, 2 eq) and DIEA (134 μL, 0.771 mmol, 10 eq) in $CH_2Cl_2$ (4 mL), and the reaction was stirred at room temperature for 18 h. The solvent was removed by rotary evaporation, and the crude material was purified by reverse phase HPLC (30–60% MeCN/$H_2O$, linear gradient, with constant 0.1% v/v TFA additive) to yield 33.7 mg (53%, TFA salt) of $JF_{571}$–HaloTag ligand (**38**) as a dark red-purple solid. $^1H$ NMR ($CD_3OD$, 400 MHz) δ 9.08 (t, *J* = 5.3 Hz, 1H), 7.28 (d, *J* = 9.2 Hz, 2H), 6.67 (dd, *J* = 9.2, 2.2 Hz, 2H), 6.53 (d, *J* = 2.2 Hz, 2H), 4.34 (t, *J* = 7.6 Hz, 8H), 3.67 – 3.55 (m, 8H), 3.52 (t, *J* = 6.7 Hz, 2H), 3.43 (t, *J* = 6.5 Hz, 2H), 2.57 (p, *J* = 7.6 Hz, 4H), 1.76 – 1.67 (m, 2H), 1.54 – 1.46 (m, 2H), 1.44 – 1.29 (m, 4H); $^{19}F$ NMR ($CD_3OD$, 376 MHz) δ −75.40 (s, 3F), −116.71 (d, *J* = 15.3 Hz, 1F), −132.53 (d, *J* = 22.4 Hz, 1F), −140.20 (dd, *J* = 22.6, 15.3 Hz, 1F); Analytical HPLC: $t_R$ = 12.8 min, >99% purity (10–95% MeCN/$H_2O$, linear gradient, with constant 0.1% v/v TFA additive; 20 min run; 1 mL/min flow; ESI; positive ion mode; detection at 575 nm); HRMS (ESI) calcd for $C_{37}H_{40}ClF_3N_3O_6$ $[M+H]^+$ 714.2552, found 714.2561.

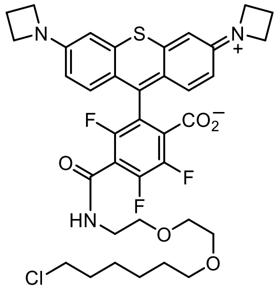

**$JF_{593}$–HaloTag ligand (39):** Ether **S67** (acetate salt; 50 mg, 75.2 μmol) was taken up in $CH_2Cl_2$ (4 mL); triethylsilane (400 μL) was added, followed by trifluoroacetic acid (800 μL). The reaction was stirred at room temperature for 6 h. Toluene (5 mL) was added, and the reaction mixture was concentrated to dryness. The residue was combined with a premixed solution of HaloTag(O2)amine (HTL-$NH_2$, **36**; TFA salt; 50.8 mg, 0.150 mmol, 2 eq) and DIEA (131 μL, 0.752 mmol, 10 eq) in $CH_2Cl_2$ (3 mL), and the reaction was stirred at room temperature for 18 h. The solvent was removed by rotary evaporation, and the crude material was purified by reverse phase HPLC (30–60% MeCN/$H_2O$, linear gradient, with constant 0.1% v/v TFA additive) to afford $JF_{593}$–HaloTag ligand (**39**) as a dark purple solid (28.9 mg, 46%, TFA salt). $^1H$ NMR ($CD_3OD$, 400 MHz) δ 9.10 (t, *J* = 5.4 Hz, 1H), 7.39 (d, *J* = 9.3 Hz, 2H), 6.89 (d, *J* = 2.3 Hz, 2H), 6.70 (dd, *J* = 9.4, 2.3 Hz, 2H), 4.32 (t, *J* = 7.7 Hz, 8H), 3.67 – 3.54 (m, 8H), 3.52 (t, *J* = 6.6 Hz, 2H), 3.42

(t, $J$ = 6.5 Hz, 2H), 2.56 (p, $J$ = 7.6 Hz, 4H), 1.76 – 1.67 (m, 2H), 1.54 – 1.45 (m, 2H), 1.44 – 1.27 (m, 4H); $^{19}$F NMR ($CD_3OD$, 376 MHz) δ −75.42 (s, 3F), −117.12 (d, $J$ = 15.4 Hz, 1F), −133.08 (d, $J$ = 22.2 Hz, 1F), −139.93 (dd, $J$ = 21.7, 15.8 Hz, 1F); Analytical HPLC: $t_R$ = 12.5 min, >99% purity (10–95% MeCN/$H_2O$, linear gradient, with constant 0.1% v/v TFA additive; 20 min run; 1 mL/min flow; ESI; positive ion mode; detection at 600 nm); HRMS (ESI) calcd for $C_{37}H_{40}ClF_3N_3O_5S$ $[M+H]^+$ 730.2324, found 730.2333.

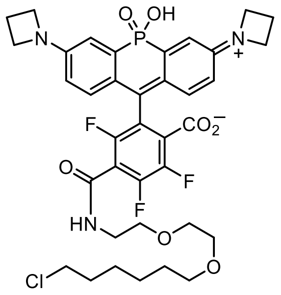

**$JF_{690}$–HaloTag ligand (40):** Phosphinate **S49** (110 mg, 0.188 mmol) and 2-(methoxymethoxy)malononitrile (**34**; 23.7 mg, 0.188 mmol, 1 eq) were combined in DMF (4 mL), and DIEA (65.3 µL, 0.375 mmol, 2 eq) was added. After stirring the reaction at room temperature for 2 h, it was evaporated to dryness. Flash chromatography on silica gel (0–40% acetone/$CH_2Cl_2$, linear gradient) afforded the MOM-MAC-substituted *tert*-butyl phosphinate adduct as a yellow solid (31 mg, 24%). Analytical HPLC: $t_R$ = 14.0 min, 96.1% purity (10–95% MeCN/$H_2O$, linear gradient, with constant 0.1% v/v TFA additive; 20 min run; 1 mL/min flow; ESI; positive ion mode; detection at 725 nm); MS (ESI) calcd for $C_{35}H_{33}F_3N_4O_6P$ $[M+H]^+$ 693.2, found 692.8.

This ether (30 mg, 43.3 µmol) was taken up in $CH_2Cl_2$ (3 mL); triethylsilane (300 µL) was added, followed by trifluoroacetic acid (600 µL). The reaction was stirred at room temperature for 6 h. Toluene (3 mL) was added, and the reaction mixture was concentrated to dryness. The residue was combined with a premixed solution of HaloTag(O2)amine (HTL-$NH_2$, **36**; TFA salt; 29.3 mg, 86.6 µmol, 2 eq) and DIEA (75.4 µL, 0.433 mmol, 10 eq) in $CH_2Cl_2$ (3 mL), and the reaction was stirred at room temperature for 18 h. The solvent was removed by rotary evaporation, and the crude material was purified by reverse phase HPLC (10–75% MeCN/$H_2O$, linear gradient, with constant 0.1% v/v TFA additive). The pooled HPLC product fractions were partially concentrated to remove MeCN and extracted with $CH_2Cl_2$ (2×). The organic extracts were dried over anhydrous $MgSO_4$, filtered, and evaporated to yield 14.0 mg (42%) of $JF_{690}$–HaloTag ligand (**40**) as a dark blue-green solid. $^1$H NMR ($CD_3OD$, 400 MHz) δ 7.08 (dd, $J$ = 14.5, 2.5 Hz, 2H), 6.98 (dd, $J$ = 9.1, 6.0 Hz, 2H), 6.47 (dd, $J$ = 9.1, 2.5 Hz, 2H), 4.25 (t, $J$ = 7.6 Hz, 8H), 3.67 – 3.55 (m, 8H), 3.53 (d, $J$ = 6.6 Hz, 2H), 3.44 (t, $J$ = 6.5 Hz, 2H), 2.52 (p, $J$ = 7.6 Hz, 4H), 1.79 – 1.68 (m, 2H), 1.59 – 1.49 (m, 2H), 1.47 – 1.31 (m, 4H); $^{19}$F NMR ($CD_3OD$, 376 MHz) δ −117.71 – −118.09 (m, 1F), −133.70 – −133.93 (m, 1F), −141.50 – −141.84 (m, 1F); Analytical HPLC: $t_R$ = 12.3 min, >99% purity (10–95% MeCN/$H_2O$, linear gradient, with constant 0.1% v/v TFA additive; 20 min run; 1 mL/min flow; ESI; positive ion mode; detection at 700 nm); HRMS (ESI) calcd for $C_{37}H_{41}ClF_3N_3O_7P$ $[M+H]^+$ 762.2317, found 762.2334.

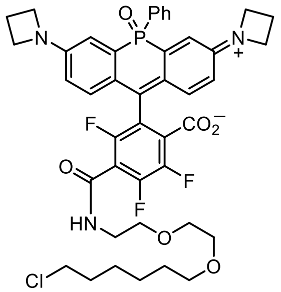


**$JF_{722}$–HaloTag ligand (41):** Ether **S68** (35 mg, 50.2 μmol) was taken up in $CH_2Cl_2$ (3 mL); triethylsilane (300 μL) was added, followed by trifluoroacetic acid (600 μL). The reaction was stirred at room temperature for 6 h. Toluene (5 mL) was added, and the reaction mixture was concentrated to dryness. The residue was combined with a premixed solution of HaloTag(O2)amine (HTL-$NH_2$, **36**; TFA salt; 33.9 mg, 0.100 mmol, 2 eq) and DIEA (87.5 μL, 0.502 mmol, 10 eq) in $CH_2Cl_2$ (3 mL), and the reaction was stirred at room temperature for 18 h. The solvent was removed by rotary evaporation, and the crude material was purified by reverse phase HPLC (10–95% MeCN/$H_2O$, linear gradient, with constant 0.1% v/v TFA additive). The pooled HPLC product fractions were partially concentrated to remove MeCN, diluted with saturated $NaHCO_3$, and extracted with $CH_2Cl_2$ (2×). The organic extracts were dried over anhydrous $MgSO_4$, filtered, and evaporated to yield 24.3 mg (59%) of $JF_{722}$–HaloTag ligand (**41**) as a blue-green solid. $^1H$ NMR ($CDCl_3$, 400 MHz) δ 7.76 – 7.65 (m, 2H), 7.50 – 7.37 (m, 3H), 6.76 (dd, $J$ = 8.8, 6.0 Hz, 2H), 6.74 (dd, $J$ = 13.9, 2.6 Hz, 2H), 6.66 (s, 1H), 6.44 (dd, $J$ = 8.8, 2.5 Hz, 2H), 3.96 – 3.81 (m, 8H), 3.64 – 3.56 (m, 6H), 3.53 – 3.47 (m, 4H), 3.36 (t, $J$ = 6.7 Hz, 2H), 2.35 (p, $J$ = 7.3 Hz, 4H), 1.78 – 1.69 (m, 2H), 1.54 – 1.37 (m, 4H), 1.35 – 1.27 (m, 2H); $^{19}F$ NMR ($CDCl_3$, 376 MHz) δ −120.45 (d, $J$ = 22.7 Hz, 1F), −132.28 (d, $J$ = 21.4 Hz, 1F), −141.55 (t, $J$ = 22.2 Hz, 1F); Analytical HPLC: $t_R$ = 14.8 min, >99% purity (10–95% MeCN/$H_2O$, linear gradient, with constant 0.1% v/v TFA additive; 20 min run; 1 mL/min flow; ESI; positive ion mode; detection at 725 nm); HRMS (ESI) calcd for $C_{43}H_{45}ClF_3N_3O_6P$ $[M+H]^+$ 822.2681, found 822.2688.

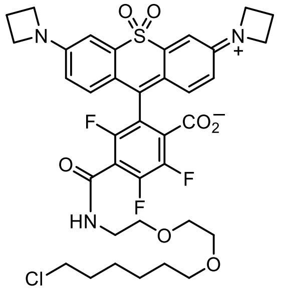


**$JF_{724}$–HaloTag ligand (42):** Ether **S69** (40 mg, 62.8 μmol) was taken up in $CH_2Cl_2$ (4 mL); triethylsilane (400 μL) was added, followed by trifluoroacetic acid (800 μL). The reaction was stirred at room temperature for 18 h. Toluene (5 mL) was added, and the reaction mixture was concentrated to dryness. The residue was combined with a premixed solution of HaloTag(O2)amine (HTL-$NH_2$, **36**; TFA salt; 42 mg, 0.126 mmol, 2 eq) and DIEA (109 μL, 0.628 mmol, 10 eq) in $CH_2Cl_2$ (3 mL), and the reaction was stirred at room temperature for 2 h. The solvent was removed by rotary evaporation, and the crude material was purified by reverse phase HPLC (40–70% MeCN/$H_2O$, linear gradient, with constant 0.1% v/v TFA additive). The pooled HPLC product fractions were partially concentrated to remove MeCN, diluted with saturated $NaHCO_3$, and extracted with $CH_2Cl_2$ (2×). The organic extracts were dried over anhydrous $MgSO_4$, filtered, and evaporated to yield 15.3 mg (32%) of $JF_{724}$–HaloTag ligand (**42**) as an off-white solid. $^1$H NMR ($CDCl_3$, 400 MHz) δ 7.07 (d, $J$ = 2.5 Hz, 2H), 6.77 (d, $J$ = 8.8 Hz, 2H), 6.69 (bs, 1H), 6.42 (dd, $J$ = 8.7, 2.5 Hz, 2H), 4.00 (t, $J$ = 7.3 Hz, 8H), 3.66 – 3.57 (m, 6H), 3.56 – 3.48 (m, 4H), 3.38 (t, $J$ = 6.6 Hz, 2H), 2.44 (p, $J$ = 7.3 Hz, 4H), 1.79 – 1.69 (m, 2H), 1.55 – 1.36 (m, 4H), 1.36 – 1.27 (m, 2H); $^{19}$F NMR ($CDCl_3$, 376 MHz) δ −117.76 (d, $J$ = 22.3 Hz, 1F), −130.89 (d, $J$ = 22.2 Hz, 1F), −140.68 (t, $J$ = 22.1 Hz, 1F); Analytical HPLC: $t_R$ = 13.6 min, >99% purity (10–95% MeCN/$H_2O$, linear gradient, with constant 0.1% v/v TFA additive; 20 min run; 1 mL/min flow; ESI; positive ion mode; detection at 254 nm); HRMS (ESI) calcd for $C_{37}H_{40}ClF_3N_3O_7S$ $[M+H]^+$ 762.2222, found 762.2238.

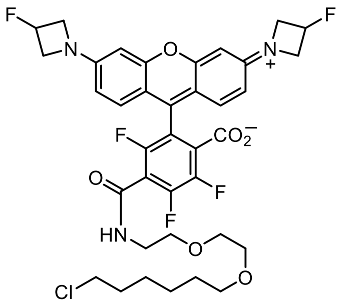


**$JF_{559}$–HaloTag ligand (44):** Ether **S70** (80 mg, 0.128 mmol) was taken up in $CH_2Cl_2$ (5 mL); triethylsilane (500 μL) was added, followed by trifluoroacetic acid (1 mL). The reaction was stirred at room temperature for 6 h. Toluene (5 mL) was added, and the reaction mixture was concentrated to dryness. The residue was combined with a premixed solution of HaloTag(O2)amine (HTL-$NH_2$, **36**; TFA salt; 87 mg, 0.256 mmol, 2 eq) and DIEA (223 μL, 1.28 mmol, 10 eq) in DMF (4 mL), and the reaction was stirred at room temperature for 18 h. The solvent was removed by rotary evaporation, and the crude material was purified by reverse phase HPLC (10–75% MeCN/$H_2O$, linear gradient, with constant 0.1% v/v TFA additive). The pooled HPLC product fractions were partially concentrated to remove MeCN, diluted with saturated $NaHCO_3$, and extracted with 10% MeOH/$CH_2Cl_2$ (3×). The organic extracts were dried over anhydrous $MgSO_4$, filtered, and evaporated to yield 54.3 mg (57%) of $JF_{559}$–HaloTag ligand (**44**) as a dark red-purple solid. $^1$H NMR ($CD_3OD$, 400 MHz) δ 7.43 (d, $J$ = 9.1 Hz, 2H), 6.76 (dd, $J$ = 9.2, 2.2 Hz, 2H), 6.62 (d, $J$ = 2.2 Hz, 2H), 5.59 (dtt, $^2J_{HF}$ = 57.0 Hz, $J$ = 6.0, 3.0 Hz, 2H), 4.70 – 4.56 (m, 4H), 4.47 – 4.34 (m, 4H), 3.69 – 3.57 (m, 8H), 3.54 (t, $J$ = 6.6 Hz, 2H), 3.45 (t, $J$ = 6.5 Hz, 2H), 1.78 – 1.69 (m, 2H), 1.58 – 1.49 (m, 2H), 1.47 – 1.30 (m, 4H); $^{19}$F NMR ($CD_3OD$, 376 MHz) δ −117.57 (d, $J$ = 15.7 Hz, 1F), −133.24 (d, $J$ = 23.1 Hz, 1F), −143.33 (dd, $J$ = 23.4, 15.7 Hz, 1F), −180.48 (dtt, $J_{FH}$ = 56.8, 23.3, 20.3 Hz, 2F); Analytical HPLC: $t_R$ = 12.4 min, >99% purity (10–95% MeCN/$H_2O$, linear gradient, with constant 0.1% v/v TFA additive; 20 min run; 1 mL/min flow; ESI; positive ion mode; detection at 550 nm); HRMS (ESI) calcd for $C_{37}H_{38}ClF_5N_3O_6$ $[M+H]^+$ 750.2364, found 750.2378.

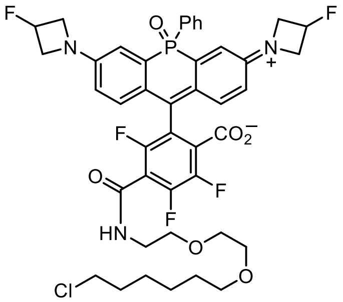


**JF$_{711}$–HaloTag ligand (46):** Ether **S71** (50 mg, 68.3 μmol) was taken up in $CH_2Cl_2$ (4 mL); triethylsilane (400 μL) was added, followed by trifluoroacetic acid (800 μL). The reaction was stirred at room temperature for 6 h. Toluene (4 mL) was added, and the reaction mixture was concentrated to dryness. The residue was combined with a premixed solution of HaloTag(O2)amine (HTL-$NH_2$, **36**; TFA salt; 46.1 mg, 0.137 mmol, 2 eq) and DIEA (119 μL, 0.683 mmol, 10 eq) in $CH_2Cl_2$ (3 mL), and the reaction was stirred at room temperature for 18 h. The solvent was removed by rotary evaporation, and the crude material was purified by reverse phase HPLC (30–70% MeCN/$H_2O$, linear gradient, with constant 0.1% v/v TFA additive). The pooled HPLC product fractions were partially concentrated to remove MeCN, diluted with saturated $NaHCO_3$, and extracted with $CH_2Cl_2$ (2×). The organic extracts were dried over anhydrous $MgSO_4$, filtered, and evaporated to yield 32 mg (55%) of JF$_{711}$–HaloTag ligand (**46**) as a pale blue-green solid. $^1$H NMR ($CDCl_3$, 400 MHz) δ 7.76 – 7.67 (m, 2H), 7.54 – 7.47 (m, 1H), 7.47 – 7.40 (m, 2H), 6.82 (dd, $J$ = 8.8, 5.7 Hz, 2H), 6.79 (dd, $J$ = 13.7, 2.6 Hz, 2H), 6.68 (s, 1H), 6.51 (dd, $J$ = 8.8, 2.6 Hz, 2H), 5.38 (dtt, $^2J_{HF}$ = 56.7 Hz, $J$ = 6.0, 3.4 Hz, 2H), 4.28 – 4.10 (m, 4H), 4.08 – 3.87 (m, 4H), 3.65 – 3.55 (m, 6H), 3.54 – 3.47 (m, 4H), 3.37 (t, $J$ = 6.7 Hz, 2H), 1.78 – 1.70 (m, 2H), 1.55 – 1.46 (m, 2H), 1.46 – 1.37 (m, 2H), 1.35 – 1.26 (m, 2H); $^{19}$F NMR ($CDCl_3$, 376 MHz) δ −120.42 (d, $J$ = 22.4 Hz, 1F), −131.74 (d, $J$ = 21.8 Hz, 1F), −141.07 (t, $J$ = 22.2 Hz, 1F), −180.54 (dtt, $J_{FH}$ = 56.6, 23.7, 18.8 Hz, 2F); Analytical HPLC: $t_R$ = 15.1 min, >99% purity (10–95% MeCN/$H_2O$, linear gradient, with constant 0.1% v/v TFA additive; 20 min run; 1 mL/min flow; ESI; positive ion mode; detection at 254 nm); HRMS (ESI) calcd for $C_{43}H_{43}ClF_5N_3O_6P$ $[M+H]^+$ 858.2493, found 858.2518.

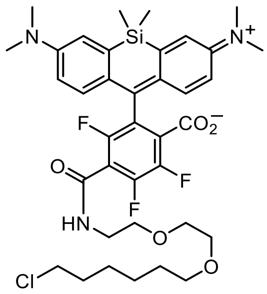


**4,5,7-Trifluoro-SiTMR–HaloTag ligand (S72):** Ether **S57** (70 mg, 0.115 mmol) was taken up in $CH_2Cl_2$ (4 mL); triethylsilane (400 μL) was added, followed by trifluoroacetic acid (800 μL). The reaction was stirred at room temperature for 18 h. Toluene (5 mL) was added, and the reaction mixture was concentrated to dryness. The residue was combined with a premixed solution of HaloTag(O2)amine (HTL-$NH_2$, **36**; TFA salt; 77.9 mg, 0.231 mmol, 2 eq) and DIEA (201 μL, 1.15 mmol, 10 eq) in $CH_2Cl_2$ (3 mL), and the reaction was stirred at room temperature for 2 h. The solvent was removed by rotary evaporation, and the crude material was purified by reverse phase HPLC (30–60%

MeCN/$H_2O$, linear gradient, with constant 0.1% v/v TFA additive). The pooled HPLC product fractions were partially concentrated to remove MeCN, diluted with saturated $NaHCO_3$, and extracted with $CH_2Cl_2$ (2×). The organic extracts were dried over anhydrous $MgSO_4$, filtered, and evaporated to yield 49.5 mg (59%) of 4,5,7-trifluoro-SiTMR–HaloTag ligand (**S72**) as a blue-green solid. $^1$H NMR ($CDCl_3$, 400 MHz) δ 6.93 (d, $J$ = 2.9 Hz, 2H), 6.81 (d, $J$ = 8.9 Hz, 2H), 6.69 (s, 1H), 6.61 (dd, $J$ = 8.9, 2.9 Hz, 2H), 3.66 – 3.58 (m, 6H), 3.54 – 3.47 (m, 4H), 3.37 (t, $J$ = 6.7 Hz, 2H), 2.99 (s, 12H), 1.77 – 1.68 (m, 2H), 1.53 – 1.44 (m, 2H), 1.43 – 1.24 (m, 4H), 0.565 (s, 3H), 0.555 (s, 3H); $^{19}$F NMR ($CDCl_3$, 376 MHz) δ −118.73 (d, $J$ = 22.7 Hz, 1F), −133.63 (d, $J$ = 21.3 Hz, 1F), −142.40 (t, $J$ = 22.1 Hz, 1F); Analytical HPLC: $t_R$ = 13.2 min, >99% purity (10–95% MeCN/$H_2O$, linear gradient, with constant 0.1% v/v TFA additive; 20 min run; 1 mL/min flow; ESI; positive ion mode; detection at 675 nm); HRMS (ESI) calcd for $C_{37}H_{46}ClF_3N_3O_5Si$ $[M+H]^+$ 732.2842, found 732.2848.

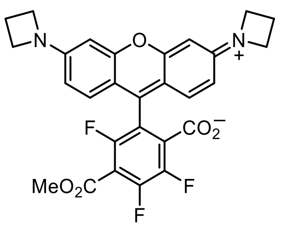


**6-Methoxycarbonyl-JF$_{571}$ (S73):** Ether **S56** (40 mg, 61.7 µmol) was taken up in $CH_2Cl_2$ (4 mL); triethylsilane (400 µL) was added, followed by trifluoroacetic acid (800 µL). The reaction was stirred at room temperature for 6 h. Toluene (5 mL) was added, and the reaction mixture was concentrated to dryness. The residue was combined with a premixed solution of MeOH (62.4 µL, 1.54 mmol, 25 eq) and $Et_3N$ (86.0 µL, 0.617 mmol, 10 eq) in $CH_2Cl_2$ (3 mL), and the reaction was stirred at room temperature for 18 h. The solvent was removed by rotary evaporation, and the crude material was purified by reverse phase HPLC (20–50% MeCN/$H_2O$, linear gradient, with constant 0.1% v/v TFA additive) to afford 6-methoxycarbonyl-JF$_{571}$ (**S73**) as a dark purple solid (26.8 mg, 68%, TFA salt). $^1$H NMR ($CD_3OD$, 400 MHz) δ 7.27 (d, $J$ = 9.2 Hz, 2H), 6.67 (dd, $J$ = 9.2, 2.2 Hz, 2H), 6.52 (d, $J$ = 2.2 Hz, 2H), 4.34 (t, $J$ = 7.7 Hz, 8H), 3.98 (s, 3H), 2.57 (p, $J$ = 7.7 Hz, 4H); $^{19}$F NMR ($CD_3OD$, 376 MHz) δ −75.52 (s, 3F), −114.14 (d, $J$ = 15.4 Hz, 1F), −129.35 (d, $J$ = 21.2 Hz, 1F), −139.46 (dd, $J$ = 21.5, 15.4 Hz, 1F); Analytical HPLC: $t_R$ = 11.2 min, >99% purity (10–95% MeCN/$H_2O$, linear gradient, with constant 0.1% v/v TFA additive; 20 min run; 1 mL/min flow; ESI; positive ion mode; detection at 575 nm); HRMS (ESI) calcd for $C_{28}H_{22}F_3N_2O_5$ $[M+H]^+$ 523.1475, found 523.1480.

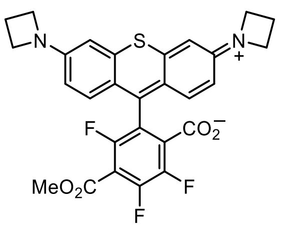


**6-Methoxycarbonyl-JF$_{593}$ (S74):** Ether **S67** (acetate salt; 50 mg, 75.2 µmol) was taken up in $CH_2Cl_2$ (4 mL); triethylsilane (400 µL) was added, followed by trifluoroacetic acid (800 µL). The reaction was stirred at room temperature for 6 h. Toluene (5 mL) was added, and the reaction mixture was concentrated to dryness. The residue was combined with a premixed solution of MeOH (76.1 μL, 1.88 mmol, 25 eq) and $Et_3N$ (105 μL, 0.752 mmol, 10 eq) in $CH_2Cl_2$ (3 mL), and the reaction was stirred at room temperature for 18 h. The solvent was removed by rotary evaporation, and the crude material was purified by reverse phase HPLC (30–50% $MeCN/H_2O$, linear gradient, with constant 0.1% v/v TFA additive) to afford 6-methoxycarbonyl-$JF_{593}$ (**S74**) as a dark purple solid (32.1 mg, 65%, TFA salt). $^1$H NMR ($CD_3OD$, 400 MHz) δ 7.38 (d, $J$ = 9.4 Hz, 2H), 6.88 (d, $J$ = 2.3 Hz, 2H), 6.71 (dd, $J$ = 9.4, 2.3 Hz, 2H), 4.31 (t, $J$ = 7.7 Hz, 8H), 3.98 (s, 3H), 2.55 (p, $J$ = 7.6 Hz, 4H); $^{19}$F NMR ($CD_3OD$, 376 MHz) δ −75.45 (s, 3F), −114.79 (d, $J$ = 15.4 Hz, 1F), −130.14 (d, $J$ = 21.7 Hz, 1F), −139.89 (dd, $J$ = 21.3, 15.6 Hz, 1F); Analytical HPLC: $t_R$ = 11.4 min, >99% purity (10–95% $MeCN/H_2O$, linear gradient, with constant 0.1% v/v TFA additive; 20 min run; 1 mL/min flow; ESI; positive ion mode; detection at 600 nm); HRMS (ESI) calcd for $C_{28}H_{22}F_3N_2O_4S$ $[M+H]^+$ 539.1247, found 539.1254.

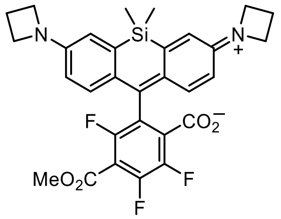

**6-Methoxycarbonyl-$JF_{669}$ (29):** Ether **35** (140 mg, 0.222 mmol) was taken up in $CH_2Cl_2$ (10 mL); triethylsilane (1 mL) was added, followed by trifluoroacetic acid (2 mL). The reaction was stirred at room temperature for 18 h. Toluene (10 mL) was added, and the reaction mixture was concentrated to dryness. The residue was combined with a premixed solution of MeOH (224 μL, 5.55 mmol, 25 eq) and $Et_3N$ (309 μL, 2.22 mmol, 10 eq) in $CH_2Cl_2$ (6 mL), and the reaction was stirred at room temperature for 30 min. The solvent was removed by rotary evaporation, and the crude material was purified by silica gel chromatography (10–100% EtOAc/hexanes, linear gradient) to yield 97 mg (77%) of 6-methoxycarbonyl-$JF_{669}$ (**29**) as a blue-green solid. $^1$H NMR ($CDCl_3$, 400 MHz) δ 6.74 (d, $J$ = 8.6 Hz, 2H), 6.64 (d, $J$ = 2.7 Hz, 2H), 6.31 (dd, $J$ = 8.7, 2.6 Hz, 2H), 3.95 (s, 3H), 3.92 (t, $J$ = 7.2 Hz, 8H), 2.38 (p, $J$ = 7.3 Hz, 4H), 0.545 (s, 3H), 0.538 (s, 3H); $^{19}$F NMR ($CDCl_3$, 376 MHz) δ −116.04 (d, $J$ = 22.6 Hz, 1F), −131.67 (d, $J$ = 20.9 Hz, 1F), −142.29 (dd, $J$ = 22.9, 20.9 Hz, 1F); Analytical HPLC: $t_R$ = 12.6 min, >99% purity (10–95% $MeCN/H_2O$, linear gradient, with constant 0.1% v/v TFA additive; 20 min run; 1 mL/min flow; ESI; positive ion mode; detection at 675 nm); HRMS (ESI) calcd for $C_{30}H_{28}F_3N_2O_4Si$ $[M+H]^+$ 565.1765, found 565.1774.

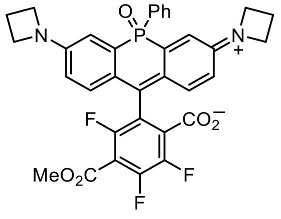

**6-Methoxycarbonyl-$JF_{722}$ (S75):** Ether **S68** (30 mg, 43.1 μmol) was taken up in $CH_2Cl_2$ (3 mL); triethylsilane (300 μL) was added, followed by trifluoroacetic acid (600 μL). The reaction was stirred at room temperature for 6 h.

Toluene (4 mL) was added, and the reaction mixture was concentrated to dryness. The residue was combined with a premixed solution of MeOH (43.6 μL, 1.08 mmol, 25 eq) and $Et_3N$ (60.0 μL, 0.431 mmol, 10 eq) in $CH_2Cl_2$ (3 mL), and the reaction was stirred at room temperature for 18 h. The solvent was removed by rotary evaporation, and the crude material was purified by reverse phase HPLC (40–50% MeCN/$H_2O$, linear gradient, with constant 0.1% v/v TFA additive). The pooled HPLC product fractions were partially concentrated to remove MeCN, diluted with saturated $NaHCO_3$, and extracted with $CH_2Cl_2$ (2×). The organic extracts were dried over anhydrous $MgSO_4$, filtered, and evaporated to yield 18.6 mg (68%) of 6-methoxycarbonyl-$JF_{722}$ (**S75**) as an off-white solid. $^1H$ NMR ($CDCl_3$, 400 MHz) δ 7.76 – 7.67 (m, 2H), 7.51 – 7.38 (m, 3H), 6.76 (dd, $J$ = 8.8, 6.0 Hz, 2H), 6.75 (dd, $J$ = 13.8, 2.6 Hz, 2H), 6.45 (dd, $J$ = 8.8, 2.5 Hz, 2H), 3.96 – 3.81 (m, 8H), 3.91 (s, 3Hz), 2.35 (p, $J$ = 7.3 Hz, 4H); $^{19}F$ NMR ($CDCl_3$, 376 MHz) δ −118.57 (d, $J$ = 22.6 Hz, 1F), −130.21 (d, $J$ = 20.9 Hz, 1F), −141.33 – −141.52 (m, 1F); Analytical HPLC: $t_R$ = 13.5 min, >99% purity (10–95% MeCN/$H_2O$, linear gradient, with constant 0.1% v/v TFA additive; 20 min run; 1 mL/min flow; ESI; positive ion mode; detection at 725 nm); HRMS (ESI) calcd for $C_{34}H_{27}F_3N_2O_5P$ $[M+H]^+$ 631.1604, found 631.1608.

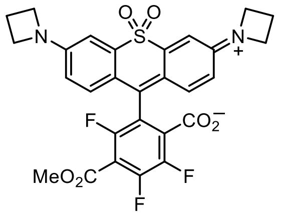


**6-Methoxycarbonyl-$JF_{724}$ (S76):** Ether **S69** (20 mg, 31.4 μmol) was taken up in $CH_2Cl_2$ (2 mL); triethylsilane (200 μL) was added, followed by trifluoroacetic acid (400 μL). The reaction was stirred at room temperature for 6 h. Toluene (3 mL) was added, and the reaction mixture was concentrated to dryness. The residue was combined with a premixed solution of MeOH (31.8 μL, 0.785 mmol, 25 eq) and $Et_3N$ (43.8 μL, 0.314 mmol, 10 eq) in $CH_2Cl_2$ (3 mL), and the reaction was stirred at room temperature for 18 h. The solvent was removed by rotary evaporation, and the crude material was purified by reverse phase HPLC (30–70% MeCN/$H_2O$, linear gradient, with constant 0.1% v/v TFA additive). The pooled product fractions were partially concentrated to remove MeCN, diluted with saturated $NaHCO_3$, and extracted with $CH_2Cl_2$ (2×). The organic extracts were dried over anhydrous $MgSO_4$, filtered, and evaporated to yield 11.6 mg (65%) of 6-methoxycarbonyl-$JF_{724}$ (**S76**) as a pale yellow solid. $^1H$ NMR ($CDCl_3$, 400 MHz) δ 7.09 (d, $J$ = 2.5 Hz, 2H), 6.74 (d, $J$ = 8.7 Hz, 2H), 6.41 (dd, $J$ = 8.7, 2.6 Hz, 2H), 4.00 (t, $J$ = 7.4 Hz, 8H), 3.93 (s, 3H), 2.44 (p, $J$ = 7.3 Hz, 4H); $^{19}F$ NMR ($CDCl_3$, 376 MHz) δ −115.63 (d, $J$ = 22.7 Hz, 1F), −129.24 (d, $J$ = 20.7 Hz, 1F), −140.60 (t, $J$ = 21.8 Hz, 1F); Analytical HPLC: $t_R$ = 15.2 min, >99% purity (10–95% MeCN/$H_2O$, linear gradient, with constant 0.1% v/v TFA additive; 20 min run; 1 mL/min flow; ESI; positive ion mode; detection at 254 nm); HRMS (ESI) calcd for $C_{28}H_{22}F_3N_2O_6S$ $[M+H]^+$ 571.1145, found 571.1156.

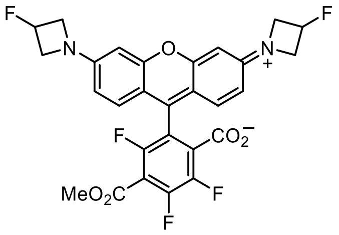


**6-Methoxycarbonyl-JF$_{559}$ (S77):** Ether **S70** (40 mg, 64.0 μmol) was taken up in $CH_2Cl_2$ (2.5 mL); triethylsilane (250 μL) was added, followed by trifluoroacetic acid (500 μL). The reaction was stirred at room temperature for 6 h. Toluene (3 mL) was added, and the reaction mixture was concentrated to dryness. The residue was combined with a premixed solution of MeOH (64.9 μL, 1.60 mmol, 25 eq) and $Et_3N$ (89.3 μL, 0.640 mmol, 10 eq) in $CH_2Cl_2$ (3 mL), and the reaction was stirred at room temperature for 18 h. The solvent was removed by rotary evaporation, and the crude material was purified by reverse phase HPLC (20–60% MeCN/$H_2O$, linear gradient, with constant 0.1% v/v TFA additive) to yield 30.7 mg (71%, TFA salt) of 6-methoxycarbonyl-JF$_{559}$ (**S77**) as a dark red-purple solid. $^1$H NMR ($CD_3OD$, 400 MHz) δ 7.37 (d, $J$ = 9.1 Hz, 2H), 6.77 (dd, $J$ = 9.2, 2.2 Hz, 2H), 6.67 (d, $J$ = 2.2 Hz, 2H), 5.57 (dtt, $^2J_{HF}$ = 56.9 Hz, $J$ = 5.9, 2.9 Hz, 2H), 4.70 – 4.57 (m, 4H), 4.49 – 4.35 (m, 4H), 3.98 (s, 3H); $^{19}$F NMR ($CD_3OD$, 376 MHz) δ −75.50 (s, 3F), −114.05 (d, $J$ = 15.4 Hz, 1F), −129.03 (d, $J$ = 21.5 Hz, 1F), −139.23 (dd, $J$ = 21.5, 15.4 Hz, 1F), −180.58 (dtt, $J_{FH}$ = 56.7, 23.2, 20.1 Hz, 2F); Analytical HPLC: $t_R$ = 11.0 min, >99% purity (10–95% MeCN/$H_2O$, linear gradient, with constant 0.1% v/v TFA additive; 20 min run; 1 mL/min flow; ESI; positive ion mode; detection at 550 nm); HRMS (ESI) calcd for $C_{28}H_{20}F_5N_2O_5$ $[M+H]^+$ 559.1287, found 559.1296.

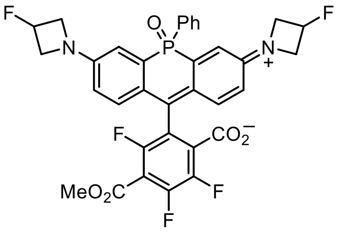


**6-Methoxycarbonyl-JF$_{711}$ (S78):** Ether **S71** (25 mg, 34.1 μmol) was taken up in $CH_2Cl_2$ (2 mL); triethylsilane (200 μL) was added, followed by trifluoroacetic acid (400 μL). The reaction was stirred at room temperature for 6 h. Toluene (3 mL) was added, and the reaction mixture was concentrated to dryness. The residue was combined with a premixed solution of MeOH (34.6 μL, 0.853 mmol, 25 eq) and $Et_3N$ (47.6 μL, 0.341 mmol, 10 eq) in $CH_2Cl_2$ (3 mL), and the reaction was stirred at room temperature for 18 h. The solvent was removed by rotary evaporation, and the crude material was purified by reverse phase HPLC (20–70% MeCN/$H_2O$, linear gradient, with constant 0.1% v/v TFA additive). The pooled HPLC product fractions were partially concentrated to remove MeCN, diluted with saturated $NaHCO_3$, and extracted with $CH_2Cl_2$ (2×). The organic extracts were dried over anhydrous $MgSO_4$, filtered, and evaporated to yield 17.7 mg (78%) of 6-methoxycarbonyl-JF$_{711}$ (**S78**) as an off-white solid. $^1$H NMR ($CDCl_3$, 400 MHz) δ 7.72 (dd, $J$ = 12.7, 7.4 Hz, 2H), 7.55 – 7.48 (m, 1H), 7.48 – 7.41 (m, 2H), 6.81 (dd, $J$ = 8.7, 5.9 Hz, 2H), 6.80 (dd, $J$ = 13.5, 2.7 Hz, 2H), 6.52 (dd, $J$ = 8.8, 2.6 Hz, 2H), 5.38 (dtt, $^2J_{HF}$ = 56.8 Hz, $J$ = 6.1, 3.4 Hz, 2H), 4.29 – 4.10 (m, 4H), 4.09 – 3.92 (m, 4H), 3.92 (s, 3H); $^{19}$F NMR ($CDCl_3$, 376 MHz) δ −118.65 (d, $J$ = 22.5 Hz, 1F), −129.59 (d, $J$ = 20.7 Hz, 1F), −140.88 (t, $J$ = 21.8 Hz, 1F), −180.52 (dtt, $J_{FH}$ = 57.0, 23.8, 18.8 Hz, 2H); Analytical HPLC: $t_R$ =

14.1 min, >99% purity (10–95% MeCN/$H_2O$, linear gradient, with constant 0.1% v/v TFA additive; 20 min run; 1 mL/min flow; ESI; positive ion mode; detection at 280 nm); HRMS (ESI) calcd for $C_{34}H_{25}F_5N_2O_5P$ $[M+H]^+$ 667.1416, found 667.1423.

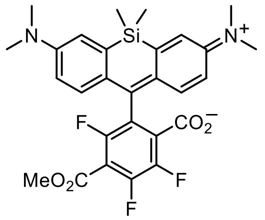


**4,5,7-Trifluoro-6-methoxycarbonyl-SiTMR (S79):** Ether **S57** (600 mg, 0.989 mmol) was taken up in $CH_2Cl_2$ (30 mL); triethylsilane (3 mL) was added, followed by trifluoroacetic acid (6 mL). The reaction was stirred at room temperature for 18 h. Toluene (15 mL) was added, and the reaction mixture was concentrated to dryness. The residue was combined with a premixed solution of MeOH (1.00 mL, 24.7 mmol, 25 eq) and $Et_3N$ (1.38 mL, 9.89 mmol, 10 eq) in $CH_2Cl_2$ (20 mL), and the reaction was stirred at room temperature for 30 min. The solvent was removed by rotary evaporation, and the crude material was purified by silica gel chromatography (10–100% EtOAc/hexanes, linear gradient) to yield 392 mg (73%) of **S79** as a yellow-green solid. $^1$H NMR ($CDCl_3$, 400 MHz) δ 6.94 (d, $J$ = 2.9 Hz, 2H), 6.78 (d, $J$ = 9.0 Hz, 2H), 6.61 (dd, $J$ = 8.9, 2.9 Hz, 2H), 3.94 (s, 3H), 2.99 (s, 12H), 0.58 (s, 3H), 0.56 (s, 3H); $^{19}$F NMR ($CDCl_3$, 376 MHz) δ −116.26 (d, $J$ = 22.8 Hz, 1F), −131.82 (d, $J$ = 20.9 Hz, 1F), −142.43 (dd, $J$ = 22.7, 21.0 Hz, 1F); Analytical HPLC: $t_R$ = 10.3 min, >99% purity (10–95% MeCN/$H_2O$, linear gradient, with constant 0.1% v/v TFA additive; 20 min run; 1 mL/min flow; ESI; positive ion mode; detection at 675 nm); HRMS (ESI) calcd for $C_{28}H_{28}F_3N_2O_4Si$ $[M+H]^+$ 541.1765, found 541.1769.

### SUBSTITUTION OF 4,5,6,7-TETRAFLUORORHODAMINES WITH OTHER NUCLEOPHILES

During our efforts to develop a method for the installation of amides and esters on the bottom aryl ring of fluorinated rhodamines, we also explored the scope of fluoride substitution with nucleophiles beyond MAC reagents. As shown in **Figure 2a**, **Supplementary Figure 2**, and **Scheme S3**, the range of tolerated nucleophiles was quite broad, encompassing azide ($NaN_3$), cyanide (KCN), malonates, cyanoacetates, amines (including ammonia and secondary amines), hydroxylamine, and the previously described MAC reagent; thiols are also excellent reagents for this process, but their utility was well-known prior to this work.[2,3] In each case, the rhodamine was simply combined with the nucleophile in DMF or DMSO; a mild base (DIEA or $K_2CO_3$) was added for nucleophiles other than azide and cyanide; heating was only required for larger secondary amines (**30**–**31**, **Supplementary Fig. 2c**) and *tert*-butyl malonates. Si-rhodamines ($JF_{669}$, **15**; $F_4$SiTMR, **S51**) and traditional oxygen rhodamines (*e.g.*, $JF_{571}$, **19**) showed similar reactivity for the different nucleophile types tested (**Scheme S3**).

We hypothesized these $S_NAr$ substitutions would proceed with the same regioselectivity as observed for the MAC reagent nucleophiles. Indeed, we observed one predominant product in every case with many reactions showing clean conversion to a single rhodamine derivative. In order to more definitively confirm the regiochemical preferences of these reactions, we converted several different substituted $JF_{669}$ products to a common derivative (**22**, **Supplementary Fig. 2**). The crystal structures of **S57** and **S79** demonstrated that the MAC chemistry provided the 6-isomer; hence, the structures of MAC-$JF_{669}$ **35** and $JF_{669}$ methyl ester **29** were as shown. Hydrolysis and Curtius rearrangement of methyl ester **29** afforded 6-amino-$JF_{669}$ (**22**). Because reduction of the azide substitution product (**21**) *and* direct substitution of $JF_{669}$ with ammonia provided material analytically identical ($^1H$/$^{13}C$/$^{19}F$ NMR, HPLC, HRMS) to the Curtius product **22**, the azide and amine substitution products were similarly assigned to be the 6-isomers.

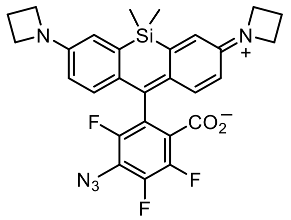


**6-Azido-$JF_{669}$ (21):** To a solution of $JF_{669}$ (**15**; 200 mg, 0.381 mmol) in DMSO (4 mL) was added $NaN_3$ (24.8 mg, 0.381 mmol, 1 eq). After stirring the reaction at room temperature for 2 h, it was diluted with water and extracted with EtOAc (2×). The combined organic extracts were washed with water and brine, dried over anhydrous $MgSO_4$, filtered, and evaporated. Silica gel chromatography (0–50% EtOAc/hexanes, linear gradient) afforded azide **21** as a blue-green solid (186 mg, 89%). $^1H$ NMR ($CDCl_3$, 400 MHz) δ 6.77 (dd, $J$ = 8.7, 0.8 Hz, 2H), 6.63 (d, $J$ = 2.6 Hz, 2H), 6.32 (dd, $J$ = 8.7, 2.7 Hz, 2H), 3.92 (t, $J$ = 7.3 Hz, 8H), 2.38 (p, $J$ = 7.2 Hz, 4H), 0.54 (s, 3H), 0.52 (s, 3H); $^{19}F$ NMR ($CDCl_3$, 376 MHz) δ −129.29 (dd, $J$ = 19.8, 4.7 Hz, 1F), −141.30 (t, $J$ = 19.8 Hz, 1F), −142.83 (dd, $J$ = 19.6, 4.8 Hz, 1F); Analytical HPLC: $t_R$ = 13.8 min, >99% purity (10–95% MeCN/$H_2O$, linear gradient, with constant 0.1% v/v TFA additive; 20 min run; 1 mL/min flow; ESI; positive ion mode; detection at 675 nm); HRMS (ESI) calcd for $C_{28}H_{25}F_3N_5O_2Si$ $[M+H]^+$ 548.1724, found 548.1722.

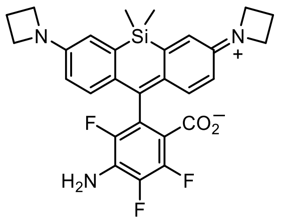


**6-Amino-JF$_{669}$ (22):** The title compound was prepared from three different substrates via three different approaches: (a) $S_NAr$ of JF$_{669}$ (**15**) with ammonia; (b) Staudinger reduction of 6-azido-JF$_{669}$ (**21**); and (c) hydrolysis and Curtius rearrangement of 6-methoxycarbonyl-JF$_{669}$ (**29**). The NMR spectra and LC/MS analyses of the three products were identical. Because the crystal structures of **S57** and **S79** had confirmed the regioselectivity of the MAC substitution (*vide infra*), this result also corroborated the regiochemical outcomes of the azide and amine substitutions.

*Via $S_NAr$:* To a solution of JF$_{669}$ (**15**; 100 mg, 0.191 mmol) in DMF (2.5 mL) was added $NH_3$ in dioxane (0.5 M, 2.29 mL, 1.14 mmol, 6 eq). After stirring the sealed reaction at room temperature for 72 h, it was concentrated *in vacuo* and purified by flash chromatography on silica gel (0–75% EtOAc/hexanes, linear gradient) to provide 78 mg (78%) of **22** as a pale blue solid.

*Via Staudinger reduction:* To a solution of 6-azido-JF$_{669}$ (**21**; 25 mg, 45.7 μmol) in THF (2 mL) was added $PPh_3$ (23.9 mg, 91.3 μmol, 2 eq). The reaction was stirred at room temperature for 30 min, at which point LC/MS analysis indicated complete conversion to the iminophosphorane. Following the addition of 1 M $H_2SO_4$ (500 μL), the mixture was vigorously stirred at room temperature for 3 h. It was subsequently diluted with saturated $NaHCO_3$ and extracted with EtOAc (2×). The combined organic extracts were washed with brine, dried over anhydrous $MgSO_4$, filtered, and concentrated *in vacuo*. Purification of the crude product by silica gel chromatography (0–75% EtOAc/hexanes, linear gradient) yielded 21.9 mg (92%) of **22** as a pale blue solid.

*Via Curtius rearrangement:* To a solution of 6-methoxycarbonyl-JF$_{669}$ (**29**; 525 mg, 0.930 mmol) in THF (19 mL) was added 1 M LiOH (4.65 mL, 4.65 mmol, 5 eq). After stirring the reaction at room temperature for 18 h, it was acidified with 1 M HCl (5 mL), diluted with water, and extracted with EtOAc (2×). The combined organic extracts were dried over anhydrous $MgSO_4$, filtered, and evaporated to provide 6-carboxy-JF$_{669}$ as a dark blue solid (382 mg, 75%). An analytically pure sample of 6-carboxy-JF$_{669}$ for spectral characterization was obtained by reverse phase HPLC (20–60% $MeCN/H_2O$, linear gradient, with constant 0.1% TFA). $^1H$ NMR ($CD_3OD$, 400 MHz) δ 7.04 (d, $J$ = 9.2 Hz, 2H), 6.87 (d, $J$ = 2.6 Hz, 2H), 6.41 (dd, $J$ = 9.2, 2.6 Hz, 2H), 4.26 (t, $J$ = 7.6 Hz, 8H), 2.51 (p, $J$ = 7.6 Hz, 4H), 0.56 (s, 3H), 0.49 (s, 3H); $^{19}F$ NMR ($CD_3OD$, 376 MHz) δ −116.97 – −117.17 (m, 1F), −133.81 – −134.08 (m, 1F), −141.65 – −141.92 (m, 1F); Analytical HPLC: $t_R$ = 10.9 min, >99% purity (10–95% $MeCN/H_2O$, linear gradient, with constant 0.1% v/v TFA additive; 20 min run; 1 mL/min flow; ESI; positive ion mode; detection at 675 nm); HRMS (ESI) calcd for $C_{29}H_{26}F_3N_2O_4Si$ $[M+H]^+$ 551.1608, found 551.1616.

A vial was charged with 6-carboxy-$JF_{669}$ (20 mg, 36.3 μmol), sealed, and evacuated/backfilled with nitrogen (3×). After suspending the starting material in dry *tert*-butanol (2 mL), DPPA (15.7 μL, 72.6 μmol, 2 eq) and $Et_3N$ (15.2 μL, 109 μmol, 3 eq) were added. The sealed reaction was then stirred at 100 ° C for 18 h. It was subsequently cooled to room temperature and concentrated *in vacuo*. The resulting residue was redissolved in $CH_2Cl_2$ (3 mL); triethylsilane (300 μL) was added, followed by trifluoroacetic acid (600 μL). The reaction was stirred at room temperature for 2 h. Toluene (5 mL) was added, and the reaction mixture was concentrated to dryness. The crude material was diluted with saturated $NaHCO_3$ and extracted with $CH_2Cl_2$ (2×). The combined organic extracts were dried over anhydrous $MgSO_4$, filtered, and evaporated. Flash chromatography on silica gel (0–75% EtOAc/hexanes, linear gradient) afforded 8.4 mg (44%) of **22** as a pale blue solid.

$^1H$ NMR ($CDCl_3$, 400 MHz) δ 6.84 (dd, *J* = 8.6, 0.8 Hz, 2H), 6.64 (d, *J* = 2.6 Hz, 2H), 6.31 (dd, *J* = 8.6, 2.6 Hz, 2H), 4.36 (s, 2H), 3.91 (t, *J* = 7.2 Hz, 8H), 2.37 (p, *J* = 7.3 Hz, 4H), 0.54 (s, 3H), 0.53 (s, 3H); $^{19}F$ NMR ($CDCl_3$, 376 MHz) δ −140.63 (dd, *J* = 18.1, 11.5 Hz, 1F), −143.85 (dd, *J* = 20.3, 18.1 Hz, 1F), −154.70 (dd, *J* = 20.3, 11.6 Hz, 1F); Analytical HPLC: $t_R$ = 12.3 min, >99% purity (10–95% $MeCN/H_2O$, linear gradient, with constant 0.1% v/v TFA additive; 20 min run; 1 mL/min flow; ESI; positive ion mode; detection at 675 nm); HRMS (ESI) calcd for $C_{28}H_{27}F_3N_3O_2Si$ $[M+H]^+$ 522.1819, found 522.1828.

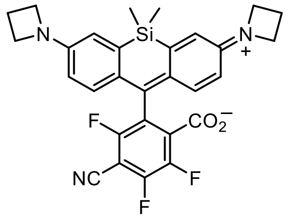


**6-Cyano-$JF_{669}$ (23):** To a solution of $JF_{669}$ (**15**; 200 mg, 0.381 mmol) in DMSO (4 mL) was added NaCN (28.0 mg, 0.572 mmol, 1.5 eq). After stirring the reaction at room temperature for 2 h, it was diluted with water and extracted with EtOAc (2×). The combined organic extracts were washed with brine, dried over anhydrous $MgSO_4$, filtered, and evaporated. Silica gel chromatography (25–100% EtOAc/hexanes, linear gradient) afforded nitrile **23** as a dark green solid (40.2 mg, 20%). $^1H$ NMR ($CDCl_3$, 400 MHz) δ 6.69 (dd, *J* = 8.7, 0.8 Hz, 2H), 6.63 (d, *J* = 2.7 Hz, 2H), 6.31 (dd, *J* = 8.7, 2.6 Hz, 2H), 3.93 (t, *J* = 7.3 Hz, 8H), 2.39 (p, *J* = 7.2 Hz, 4H), 0.56 (s, 3H), 0.53 (s, 3H); $^{19}F$ NMR ($CDCl_3$, 376 MHz, $^1H$ decoupled) δ −109.23 (dd, *J* = 22.6, 2.2 Hz, 1F), −124.52 (dd, *J* = 20.0, 2.1 Hz, 1F), −140.61 (dd, *J* = 22.7, 20.1 Hz, 1F); Analytical HPLC: $t_R$ = 12.5 min, 98.9% purity (10–95% $MeCN/H_2O$, linear gradient, with constant 0.1% v/v TFA additive; 20 min run; 1 mL/min flow; ESI; positive ion mode; detection at 675 nm); HRMS (ESI) calcd for $C_{29}H_{25}F_3N_3O_2Si$ $[M+H]^+$ 532.1663, found 532.1667.

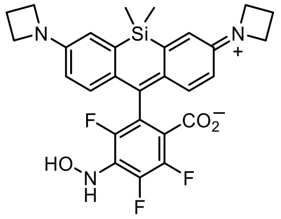

**6-Hydroxylamino-JF$_{669}$ (24):** JF$_{669}$ (**15**; 100 mg, 0.191 mmol) was dissolved in DMF (2 mL); DIEA (99.6 μL, 0.572 mmol, 3 eq) and hydroxylamine hydrochloride (14.6 mg, 0.210 mmol, 1.1 eq) were added, and the reaction was stirred at room temperature for 4 h. It was then concentrated to dryness and purified by flash chromatography on silica gel (0–50% EtOAc/toluene, linear gradient) to provide 52 mg (51%) of **24** as a blue solid. $^1$H NMR ($CDCl_3$, 400 MHz) δ 6.98 (s, 1H), 6.79 (dd, *J* = 8.6, 1.0 Hz, 2H), 6.64 (d, *J* = 2.7 Hz, 2H), 6.31 (dd, *J* = 8.6, 2.6 Hz, 2H), 5.87 (s, 1H), 3.91 (t, *J* = 7.2 Hz, 8H), 2.37 (p, *J* = 7.2 Hz, 4H), 0.55 (s, 3H), 0.54 (s, 3H); $^{19}$F NMR ($CDCl_3$, 376 MHz, $^1$H decoupled) δ −132.16 (dd, *J* = 19.5, 6.9 Hz, 1F), −142.58 (t, *J* = 20.1 Hz, 1F), −145.42 (dd, *J* = 20.5, 6.9 Hz, 1F); Analytical HPLC: $t_R$ = 10.0 min, 96.8% purity (10–95% MeCN/$H_2O$, linear gradient, with constant 0.1% v/v TFA additive; 20 min run; 1 mL/min flow; ESI; positive ion mode; detection at 675 nm); HRMS (ESI) calcd for $C_{28}H_{27}F_3N_3O_3Si$ $[M+H]^+$ 538.1768, found 538.1779.

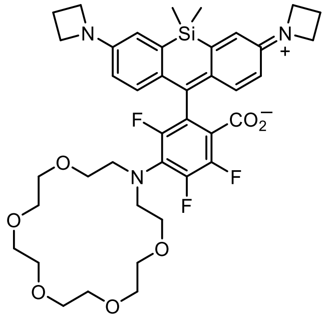

**JF$_{669}$-18-crown-6 (32):** JF$_{669}$ (**15**; 100 mg, 0.191 mmol) and 1-aza-18-crown-6 (**30**; 100 mg, 0.381 mmol, 2 eq) were combined in DMF (2 mL). After adding DIEA (99.6 μL, 0.572 mmol, 3 eq), the reaction was stirred at 50 °C for 72 h. The crude reaction mixture was directly purified by reverse phase HPLC (30–50% MeCN/$H_2O$, linear gradient, with constant 0.1% v/v TFA additive). The pooled product fractions were partially concentrated to remove MeCN, diluted with saturated $NaHCO_3$, and extracted with $CH_2Cl_2$ (2×). The organic extracts were dried over anhydrous $MgSO_4$, filtered, and evaporated to provide 54.0 mg of **32** (37%) as a light blue solid. $^1$H NMR ($CDCl_3$, 400 MHz) δ 6.79 (dd, *J* = 8.6, 0.9 Hz, 2H), 6.64 (d, *J* = 2.6 Hz, 2H), 6.28 (dd, *J* = 8.6, 2.6 Hz, 2H), 3.91 (t, *J* = 7.3 Hz, 8H), 3.70 (t, *J* = 5.5 Hz, 4H), 3.68 – 3.52 (m, 20H), 2.37 (p, *J* = 7.2 Hz, 4H), 0.54 (s, 3H), 0.54 (s, 3H); $^{19}$F NMR ($CDCl_3$, 376 MHz) δ −126.05 (dd, *J* = 18.0, 8.1 Hz, 1F), −141.75 (dd, *J* = 20.1, 8.1 Hz, 1F), −142.95 (dd, *J* = 19.9, 18.2 Hz, 1F); Analytical HPLC: $t_R$ = 12.9 min, >99% purity (10–95% MeCN/$H_2O$, linear gradient, with constant 0.1% v/v TFA additive; 20 min run; 1 mL/min flow; ESI; positive ion mode; detection at 675 nm); HRMS (ESI) calcd for $C_{40}H_{49}F_3N_3O_7Si$ $[M+H]^+$ 768.3286, found 768.3298.

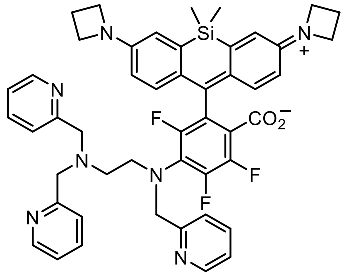

**$JF_{669}$-TPMED (33):** $JF_{669}$ (**15**; 75 mg, 0.143 mmol) and $N^1$,$N^1$,$N^2$-**t**ris(**p**yridin-2-yl**m**ethyl)**e**thane-1,2-**d**iamine[8,9] (**31**; 95.3 mg, 0.286 mmol, 2 eq) were combined in DMF (2 mL). After adding DIEA (74.7 μL, 0.429 mmol, 3 eq), the reaction was stirred at 50 °C for 96 h. The reaction mixture was concentrated to dryness and purified by reverse phase HPLC (10–75% MeCN/$H_2O$, linear gradient, with constant 0.1% v/v TFA additive). The pooled product fractions were partially concentrated to remove MeCN, diluted with saturated $NaHCO_3$, and extracted with $CH_2Cl_2$ (2×). The organic extracts were dried over anhydrous $MgSO_4$, filtered, and evaporated to provide 55.6 mg of **33** (46%) as a purple solid. $^1$H NMR ($CD_3OD$, 400 MHz) δ 8.41 – 8.36 (m, 3H), 7.72 – 7.64 (m, 3H), 7.45 (dt, $J$ = 7.9, 1.1 Hz, 2H), 7.29 – 7.24 (m, 2H), 7.23 (ddd, $J$ = 7.5, 4.9, 1.2 Hz, 2H), 6.70 (d, $J$ = 2.6 Hz, 2H), 6.54 (dd, $J$ = 8.7, 1.0 Hz, 2H), 6.23 (dd, $J$ = 8.7, 2.7 Hz, 2H), 4.49 (s, 2H), 3.89 (t, $J$ = 7.3 Hz, 8H), 3.75 (s, 4H), 3.46 (t, $J$ = 6.3 Hz, 2H), 2.76 (t, $J$ = 6.2 Hz, 2H), 2.38 (p, $J$ = 7.2 Hz, 4H), 0.53 (s, 3H), 0.46 (s, 3H); $^{19}$F NMR ($CD_3OD$, 376 MHz) δ −123.75 (dd, $J$ = 17.9, 7.1 Hz, 1F), −139.44 (dd, $J$ = 19.5, 7.3 Hz, 1F), −143.27 (t, $J$ = 18.8 Hz, 1F); Analytical HPLC: $t_R$ = 9.4 min, 97.7% purity (10–95% MeCN/$H_2O$, linear gradient, with constant 0.1% v/v TFA additive; 20 min run; 1 mL/min flow; ESI; positive ion mode; detection at 675 nm); HRMS (ESI) calcd for $C_{48}H_{47}F_3N_7O_2Si$ $[M+H]^+$ 838.3507, found 838.3526.

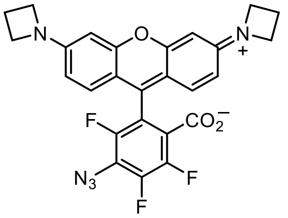

**6-Azido-$JF_{571}$ (S52):** To a solution of $JF_{571}$ (**19**; 20 mg, 41.5 μmol) in DMSO (1 mL) was added $NaN_3$ (3.0 mg, 45.6 μmol, 1.1 eq). After stirring the reaction at room temperature for 2 h, it was directly purified by reverse phase HPLC (30–60% MeCN/$H_2O$, linear gradient, with constant 0.1% v/v TFA additive) to provide 17.0 mg (66%, TFA salt) of azide **S52** as a dark red-purple solid. $^1$H NMR ($CD_3OD$, 400 MHz) δ 7.29 (d, $J$ = 9.2 Hz, 2H), 6.67 (dd, $J$ = 9.2, 2.2 Hz, 2H), 6.54 (d, $J$ = 2.2 Hz, 2H), 4.34 (t, $J$ = 7.6 Hz, 8H), 2.57 (p, $J$ = 7.7 Hz, 4H); $^{19}$F NMR ($CD_3OD$, 376 MHz) δ −75.41 (s, 3F), −126.37 (dd, $J$ = 12.6, 6.2 Hz, 1F), −137.79 (dd, $J$ = 19.6, 12.6 Hz, 1F), −142.08 (dd, $J$ = 20.0, 6.3 Hz, 1F); Analytical HPLC: $t_R$ = 11.7 min, >99% purity (10–95% MeCN/$H_2O$, linear gradient, with constant 0.1% v/v TFA additive; 20 min run; 1 mL/min flow; ESI; positive ion mode; detection at 575 nm); HRMS (ESI) calcd for $C_{26}H_{19}F_3N_5O_3$ $[M+H]^+$ 506.1435, found 506.1439.

[8] Que, E. L.; Bleher, R.; Duncan, F. E.; Kong, B. Y.; Gleber, S. C.; Vogt, S.; Chen, S.; Garwin, S. A.; Bayer, A. R.; Dravid, V. P.; Woodruff, T. K.; O'Halloran, T. V. *Nat. Chem.* **2015**, *7*, 130–139.
[9] Hureau, C.; Groni, S.; Guillot, R.; Blondin, G.; Duboc, C.; Anxolabéhère-Mallart, E. *Inorg. Chem.* **2008**, *47*, 9238–9247.

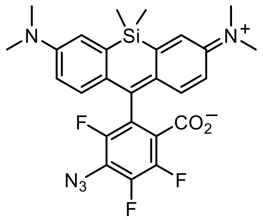


**6-Azido-4,5,7-trifluoro-SiTMR (S53):** To a solution of 4,5,6,7-tetrafluoro-SiTMR (**S51**; 100 mg, 0.200 mmol) in DMSO (2 mL) was added $NaN_3$ (14.3 mg, 0.220 mmol, 1.1 eq). After stirring the reaction at room temperature for 1 h, it was diluted with water and extracted with EtOAc (2×). The combined organic extracts were washed with brine, dried over anhydrous $MgSO_4$, filtered, and evaporated. Silica gel chromatography (0–50% EtOAc/hexanes, linear gradient) afforded azide **S53** as a blue-green solid (94 mg, 90%). $^1H$ NMR ($CDCl_3$, 400 MHz) δ 6.93 (d, *J* = 2.9 Hz, 2H), 6.82 (dd, *J* = 8.9, 0.8 Hz, 2H), 6.62 (dd, *J* = 8.9, 2.9 Hz, 2H), 2.99 (s, 12H), 0.58 (s, 3H), 0.55 (s, 3H); $^{19}F$ NMR ($CDCl_3$, 376 MHz) δ −129.60 (dd, *J* = 20.3, 4.7 Hz, 1F), −141.41 (t, *J* = 19.8 Hz, 1F), −142.94 (dd, *J* = 19.7, 4.7 Hz, 1F); Analytical HPLC: $t_R$ = 13.2 min, 98.1% purity (10–95% $MeCN/H_2O$, linear gradient, with constant 0.1% v/v TFA additive; 20 min run; 1 mL/min flow; ESI; positive ion mode; detection at 675 nm); HRMS (ESI) calcd for $C_{26}H_{25}F_3N_5O_2Si$ $[M+H]^+$ 524.1724, found 524.1731.

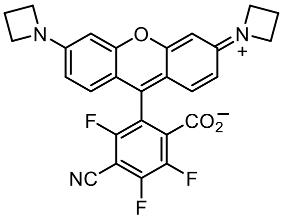


**6-Cyano-JF$_{571}$ (S54):** To a solution of JF$_{571}$ (**19**; 75 mg, 0.155 mmol) in DMSO (1.5 mL) was added NaCN (11.4 mg, 0.233 mmol, 1.5 eq). After stirring the reaction at room temperature for 18 h, it was directly purified by reverse phase HPLC (30–50% $MeCN/H_2O$, linear gradient, with constant 0.1% v/v TFA additive) to provide 16.1 mg (17%, TFA salt) of nitrile **S54** as a dark red solid. $^1H$ NMR ($CD_3OD$, 400 MHz) δ 7.29 (d, *J* = 9.2 Hz, 2H), 6.67 (dd, *J* = 9.2, 2.2 Hz, 2H), 6.52 (d, *J* = 2.2 Hz, 2H), 4.34 (t, *J* = 7.6 Hz, 8H), 2.57 (p, *J* = 7.7 Hz, 4H); $^{19}F$ NMR ($CD_3OD$, 376 MHz) δ −75.50 (s, 3F), −109.22 (d, *J* = 14.9 Hz, 1F), −123.89 (d, *J* = 21.3 Hz, 1F), −139.51 (dd, *J* = 21.2, 15.0 Hz, 1F); Analytical HPLC: $t_R$ = 11.6 min, 99.0% purity (10–95% $MeCN/H_2O$, linear gradient, with constant 0.1% v/v TFA additive; 20 min run; 1 mL/min flow; ESI; positive ion mode; detection at 575 nm); HRMS (ESI) calcd for $C_{27}H_{19}F_3N_3O_3$ $[M+H]^+$ 490.1373, found 490.1376.

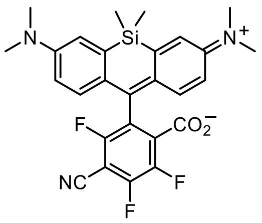

**6-Cyano-4,5,7-trifluoro-SiTMR (S55):** To a solution of 4,5,6,7-tetrafluoro-SiTMR (**S51**; 100 mg, 0.200 mmol) in DMSO (2 mL) was added NaCN (14.7 mg, 0.300 mmol, 1.5 eq). After stirring the reaction at room temperature for 4 h, it was diluted with water and extracted with EtOAc (2×). The combined organic extracts were washed with brine, dried over anhydrous $MgSO_4$, filtered, and evaporated. Silica gel chromatography (25–100% EtOAc/hexanes, linear gradient) afforded nitrile **S55** as a dark green solid (17.0 mg, 17%). $^1H$ NMR ($CDCl_3$, 400 MHz) δ 6.94 (d, $J$ = 2.8 Hz, 2H), 6.74 (dd, $J$ = 8.9, 0.8 Hz, 2H), 6.61 (dd, $J$ = 8.9, 2.9 Hz, 2H), 3.00 (s, 12H), 0.59 (s, 3H), 0.55 (s, 3H); $^{19}F$ NMR ($CDCl_3$, 376 MHz, $^1H$ decoupled) δ −109.43 (d, $J$ = 22.8, 2.2 Hz, 1F), −124.68 (dd, $J$ = 20.1, 2.2 Hz, 1F), −140.75 (dd, $J$ = 22.7, 20.1 Hz, 1F); Analytical HPLC: $t_R$ = 12.0 min, 98.7% purity (10–95% $MeCN/H_2O$, linear gradient, with constant 0.1% v/v TFA additive; 20 min run; 1 mL/min flow; ESI; positive ion mode; detection at 675 nm); HRMS (ESI) calcd for $C_{27}H_{25}F_3N_3O_2Si$ $[M+H]^+$ 508.1663, found 508.1667.

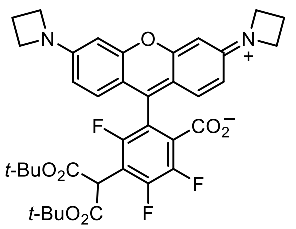

**6-(Di-*tert*-butyl malonate)-JF$_{571}$ (S58):** JF$_{571}$ (**19**; 75 mg, 0.155 mmol) and di-*tert*-butyl malonate (41.8 µL, 0.187 mmol, 1.2 eq) were combined in DMF (2 mL), and $K_2CO_3$ (51.6 mg, 0.373 mmol, 2.4 eq) was added. After stirring the reaction at 50 °C for 18 h, it was directly purified by reverse phase HPLC (30–60% $MeCN/H_2O$, linear gradient, with constant 0.1% v/v TFA additive). The pooled HPLC product fractions were partially concentrated to remove MeCN, diluted with saturated $NaHCO_3$, and extracted with $CH_2Cl_2$ (2×). The organic extracts were dried over anhydrous $MgSO_4$, filtered, and evaporated to yield 57 mg (54%) of malonate **S58** as a dark red-purple solid. $^1H$ NMR ($CDCl_3$, 400 MHz) δ 6.82 (d, $J$ = 8.6 Hz, 2H), 6.20 (d, $J$ = 2.2 Hz, 2H), 6.17 (dd, $J$ = 8.6, 2.3 Hz, 2H), 4.75 (s, 1H), 3.96 (t, $J$ = 7.4 Hz, 8H), 2.41 (p, $J$ = 7.3 Hz, 4H), 1.38 (s, 18H); $^{19}F$ NMR ($CDCl_3$, 376 MHz) δ −119.24 – −119.66 (m, 1F), −131.54 (d, $J$ = 21.0 Hz, 1F), −142.90 – −143.40 (m, 1F); Analytical HPLC: $t_R$ = 10.9 min, >99% purity (30–95% $MeCN/H_2O$, linear gradient, with constant 0.1% v/v TFA additive; 20 min run; 1 mL/min flow; ESI; positive ion mode; detection at 575 nm); HRMS (ESI) calcd for $C_{37}H_{38}F_3N_2O_7$ $[M+H]^+$ 679.2626, found 679.2623.

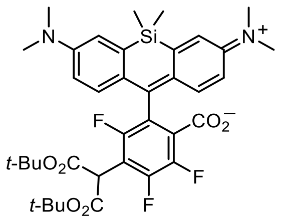

**6-(Di-*tert*-butyl malonate)-4,5,7-trifluoro-SiTMR (S59):** 4,5,6,7-Tetrafluoro-SiTMR (**S51**; 100 mg, 0.200 mmol) and di-*tert*-butyl malonate (49.2 μL, 0.220 mmol, 1.1 eq) were combined in DMF (2 mL), and $K_2CO_3$ (55.2 mg, 0.400 mmol, 2 eq) was added. After stirring the reaction at room temperature for 18 h, additional di-*tert*-butyl malonate (49.2 μL, 0.220 mmol, 1.1 eq) and $K_2CO_3$ (55.2 mg, 0.400 mmol, 2 eq) were added. The mixture was stirred at 50 °C for 24 h. It was then cooled to room temperature, diluted with water, and extracted with EtOAc (2×). The combined organic extracts were washed with brine, dried over anhydrous $MgSO_4$, filtered, and evaporated. Purification of the crude by silica gel chromatography (0–50% EtOAc/hexanes, linear gradient) yielded 83 mg (60%) of malonate **S59** as a blue-green solid. $^1$H NMR ($CDCl_3$, 400 MHz) δ 6.96 (d, $J$ = 2.9 Hz, 2H), 6.79 (dd, $J$ = 8.8, 1.4 Hz, 2H), 6.55 (dd, $J$ = 8.9, 2.9 Hz, 2H), 4.83 (s, 1H), 2.97 (s, 12H), 1.41 (s, 18H), 0.583 (s, 3H), 0.581 (s, 3H); $^{19}$F NMR ($CDCl_3$, 376 MHz, $^1$H decoupled) δ −115.39 (d, $J$ = 21.5 Hz, 1F), −132.72 (d, $J$ = 20.7 Hz, 1F), −143.43 (t, $J$ = 21.3 Hz, 1F); Analytical HPLC: $t_R$ = 12.8 min, >99% purity (30–95% MeCN/$H_2O$, linear gradient, with constant 0.1% v/v TFA additive; 20 min run; 1 mL/min flow; ESI; positive ion mode; detection at 675 nm); HRMS (ESI) calcd for $C_{37}H_{44}F_3N_2O_6Si$ $[M+H]^+$ 697.2915, found 697.2920.

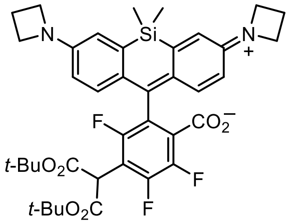

**6-(Di-*tert*-butyl malonate)-JF$_{669}$ (S60):** JF$_{669}$ (**15**; 150 mg, 0.286 mmol) and di-*tert*-butyl malonate (76.8 μL, 0.343 mmol, 1.2 eq) were combined in DMF (3 mL), and $K_2CO_3$ (94.8 mg, 0.686 mmol, 2.4 eq) was added. After stirring the reaction at 50 °C for 48 h, it was cooled to room temperature, diluted with water, and extracted with EtOAc (2×). The combined organic extracts were washed with brine, dried over anhydrous $MgSO_4$, filtered, and evaporated. Purification of the crude by silica gel chromatography (0–50% EtOAc/hexanes, linear gradient) yielded 101 mg (49%) of malonate **S60** as a blue-green solid. $^1$H NMR ($CDCl_3$, 400 MHz) δ 6.75 (dd, $J$ = 8.6, 1.7 Hz, 2H), 6.66 (d, $J$ = 2.6 Hz, 2H), 6.25 (dd, $J$ = 8.6, 2.6 Hz, 2H), 4.84 (s, 1H), 3.90 (t, $J$ = 7.2 Hz, 8H), 2.37 (p, $J$ = 7.2 Hz, 4H), 1.42 (s, 18H), 0.56 (s, 3H), 0.55 (s, 3H); $^{19}$F NMR ($CDCl_3$, 376 MHz, $^1$H decoupled) δ −114.87 (d, $J$ = 21.3 Hz, 1F), −132.66 (d, $J$ = 20.8 Hz, 1F), −143.30 (t, $J$ = 21.2 Hz, 1F); Analytical HPLC: $t_R$ = 13.4 min, 98.0% purity (30–95% MeCN/$H_2O$, linear gradient, with constant 0.1% v/v TFA additive; 20 min run; 1 mL/min flow; ESI; positive ion mode; detection at 675 nm); HRMS (ESI) calcd for $C_{39}H_{44}F_3N_2O_6Si$ $[M+H]^+$ 721.2915, found 721.2924.

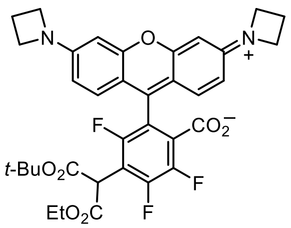


**6-(*tert*-Butyl ethyl malonate)-JF$_{571}$ (S61):** JF$_{571}$ (**19**; 75 mg, 0.155 mmol) and *tert*-butyl ethyl malonate (35.3 µL, 0.187 mmol, 1.2 eq) were combined in DMF (2 mL), and $K_2CO_3$ (51.6 mg, 0.373 mmol, 2.4 eq) was added. After stirring the reaction at 50 °C for 18 h, it was directly purified by reverse phase HPLC (30–60% MeCN/$H_2O$, linear gradient, with constant 0.1% v/v TFA additive). The pooled HPLC product fractions were partially concentrated to remove MeCN, diluted with saturated $NaHCO_3$, and extracted with $CH_2Cl_2$ (2×). The organic extracts were dried over anhydrous $MgSO_4$, filtered, and evaporated to yield 60 mg (59%) of malonate **S61** as a dark red-purple solid. $^1$H NMR ($CDCl_3$, 400 MHz) δ 6.90 – 6.82 (m, 2H), 6.22 – 6.17 (m, 4H), 4.83 (s, 1H), 4.24 – 4.12 (m, 2H), 3.98 (t, $J$ = 7.3 Hz, 8H), 2.42 (p, $J$ = 7.2 Hz, 4H), 1.38 (s, 9H), 1.20 (t, $J$ = 7.1 Hz, 3H); $^{19}$F NMR ($CDCl_3$, 376 MHz) δ −119.43 – −119.90 (m, 1F), −131.51 (d, $J$ = 21.1 Hz, 1F), −142.52 – −143.06 (m, 1F); Analytical HPLC: $t_R$ = 9.9 min, >99% purity (30–95% MeCN/$H_2O$, linear gradient, with constant 0.1% v/v TFA additive; 20 min run; 1 mL/min flow; ESI; positive ion mode; detection at 575 nm); HRMS (ESI) calcd for $C_{35}H_{34}F_3N_2O_7$ [M+H]$^+$ 651.2313, found 651.2315.

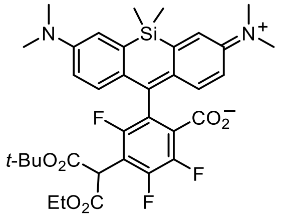


**6-(*tert*-Butyl ethyl malonate)-4,5,7-trifluoro-SiTMR (S62):** 4,5,6,7-Tetrafluoro-SiTMR (**S51**; 90 mg, 0.180 mmol) and *tert*-butyl ethyl malonate (40.9 µL, 0.216 mmol, 1.2 eq) were combined in DMF (2 mL), and $K_2CO_3$ (59.6 mg, 0.432 mmol, 2.4 eq) was added. After stirring the reaction at 50 °C for 18 h, it was cooled to room temperature, diluted with water, and extracted with EtOAc (2×). The combined organic extracts were washed with brine, dried over anhydrous $MgSO_4$, filtered, and evaporated. Purification of the crude by silica gel chromatography (0–50% EtOAc/hexanes, linear gradient) yielded 84 mg (70%) of malonate **S62** as a blue-green solid. $^1$H NMR ($CDCl_3$, 400 MHz) δ 6.96 (d, $J$ = 2.9 Hz, 2H), 6.81 – 6.74 (m, 2H), 6.58 – 6.52 (m, 2H), 4.93 (s, 1H), 4.28 – 4.19 (m, 2H), 2.98 (s, 12H), 1.42 (s, 9H), 1.25 (t, $J$ = 7.1 Hz, 3H), 0.59 (s, 3H), 0.58 (s, 3H); $^{19}$F NMR ($CDCl_3$, 376 MHz) δ −115.61 (d, $J$ = 21.9 Hz, 1F), −132.63 (d, $J$ = 20.6 Hz, 1F), −143.27 (t, $J$ = 21.3 Hz, 1F); Analytical HPLC: $t_R$ = 11.6 min, >99% purity (30–95% MeCN/$H_2O$, linear gradient, with constant 0.1% v/v TFA additive; 20 min run; 1 mL/min flow; ESI; positive ion mode; detection at 675 nm); HRMS (ESI) calcd for $C_{35}H_{40}F_3N_2O_6Si$ [M+H]$^+$ 669.2602, found 669.2610.

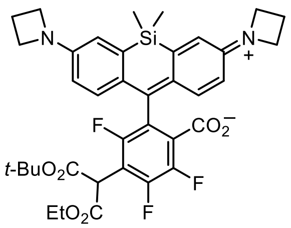

**6-(*tert*-Butyl ethyl malonate)-JF$_{669}$ (S63):** JF$_{669}$ (**15**; 150 mg, 0.286 mmol) and *tert*-butyl ethyl malonate (65.0 μL, 0.343 mmol, 1.2 eq) were combined in DMF (3 mL), and $K_2CO_3$ (94.8 mg, 0.686 mmol, 2.4 eq) was added. After stirring the reaction at 50 °C for 18 h, it was cooled to room temperature, diluted with water, and extracted with EtOAc (2×). The combined organic extracts were washed with brine, dried over anhydrous $MgSO_4$, filtered, and evaporated. Purification of the crude by silica gel chromatography (0–50% EtOAc/hexanes, linear gradient) yielded 122 mg (62%) of malonate **S63** as a blue-green solid. $^1H$ NMR ($CDCl_3$, 400 MHz) δ 6.77 – 6.71 (m, 2H), 6.66 (d, $J$ = 2.6 Hz, 2H), 6.29 – 6.23 (m, 2H), 4.93 (s, 1H), 4.28 – 4.19 (m, 2H), 3.90 (t, $J$ = 7.2 Hz, 8H), 2.37 (p, $J$ = 7.2 Hz, 4H), 1.43 (s, 9H), 1.25 (t, $J$ = 7.1 Hz, 3H), 0.57 (s, 3H), 0.55 (s, 3H); $^{19}F$ NMR ($CDCl_3$, 376 MHz) δ −115.07 (d, $J$ = 21.7 Hz, 1F), −132.59 (d, $J$ = 20.7 Hz, 1F), −143.13 (t, $J$ = 21.2 Hz, 1F); Analytical HPLC: $t_R$ = 12.1 min, >99% purity (30–95% MeCN/$H_2O$, linear gradient, with constant 0.1% v/v TFA additive; 20 min run; 1 mL/min flow; ESI; positive ion mode; detection at 675 nm); HRMS (ESI) calcd for $C_{37}H_{40}F_3N_2O_6Si$ $[M+H]^+$ 693.2602, found 693.2608.

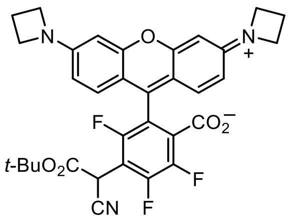

**6-(*tert*-Butyl cyanoacetate)-JF$_{571}$ (S64):** JF$_{571}$ (**19**; 75 mg, 0.155 mmol) and *tert*-butyl cyanoacetate (26.7 μL, 0.187 mmol, 1.2 eq) were combined in DMF (2 mL), and $K_2CO_3$ (51.6 mg, 0.373 mmol, 2.4 eq) was added. After stirring the reaction at room temperature for 18 h, it was directly purified by reverse phase HPLC (30–60% MeCN/$H_2O$, linear gradient, with constant 0.1% v/v TFA additive). The pooled HPLC product fractions were partially concentrated to remove MeCN, diluted with saturated $NaHCO_3$, and extracted with $CH_2Cl_2$ (2×). The organic extracts were dried over anhydrous $MgSO_4$, filtered, and evaporated to yield 59 mg (63%) of **S64** as a dark red-purple solid. $^1H$ NMR ($CDCl_3$, 400 MHz) δ 6.96 (d, $J$ = 8.7 Hz, 2H), 6.26 (dd, $J$ = 8.8, 2.3 Hz, 2H), 6.22 (d, $J$ = 2.2 Hz, 2H), 4.97 (s, 1H), 4.04 (t, $J$ = 7.4 Hz, 8H), 2.46 (p, $J$ = 7.4 Hz, 4H), 1.44 (s, 9H); $^{19}F$ NMR ($CDCl_3$, 376 MHz) δ −120.49 (d, $J$ = 20.0 Hz, 1F), −131.85 (d, $J$ = 21.6 Hz, 1F), −141.12 (t, $J$ = 20.8 Hz, 1F); Analytical HPLC: $t_R$ = 12.2 min, >99% purity (10–95% MeCN/$H_2O$, linear gradient, with constant 0.1% v/v TFA additive; 20 min run; 1 mL/min flow; ESI; positive ion mode; detection at 575 nm); HRMS (ESI) calcd for $C_{33}H_{29}F_3N_3O_5$ $[M+H]^+$ 604.2054, found 604.2050.

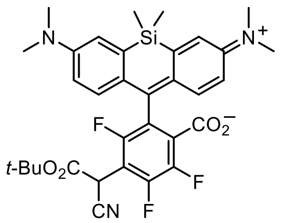

**6-(*tert*-Butyl cyanoacetate)-4,5,7-trifluoro-SiTMR (S65):** 4,5,6,7-Tetrafluoro-SiTMR (**S51**; 90 mg, 0.180 mmol) and *tert*-butyl cyanoacetate (30.8 μL, 0.216 mmol, 1.2 eq) were combined in DMF (2 mL), and $K_2CO_3$ (59.6 mg, 0.432 mmol, 2.4 eq) was added. After stirring the reaction at room temperature for 18 h, it was diluted with water and extracted with EtOAc (2×). The combined organic extracts were washed with brine, dried over anhydrous $MgSO_4$, filtered, and evaporated. Purification of the crude by silica gel chromatography (10–75% EtOAc/hexanes, linear gradient) yielded 95 mg (85%) of **S65** as a blue-green foam. $^1$H NMR ($CDCl_3$, 400 MHz) δ 6.97 – 6.93 (m, 2H), 6.77 – 6.71 (m, 2H), 6.61 – 6.55 (m, 2H), 5.03 (s, 1H), 2.99 (s, 6H), 2.99 (s, 6H), 1.46 (s, 9H), 0.59 (s, 3H), 0.57 (s, 3H); $^{19}$F NMR ($CDCl_3$, 376 MHz) δ −117.92 (d, $J$ = 22.3 Hz, 1F), −132.35 (d, $J$ = 20.4 Hz, 1F), −141.60 (dd, $J$ = 22.4, 20.4 Hz, 1F); Analytical HPLC: $t_R$ = 10.4 min, >99% purity (30–95% MeCN/$H_2O$, linear gradient, with constant 0.1% v/v TFA additive; 20 min run; 1 mL/min flow; ESI; positive ion mode; detection at 675 nm); HRMS (ESI) calcd for $C_{33}H_{35}F_3N_3O_4Si$ $[M+H]^+$ 622.2343, found 622.2345.

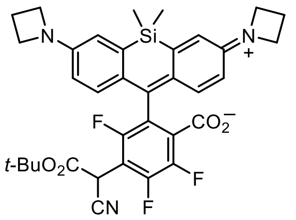

**6-(*tert*-Butyl cyanoacetate)-JF$_{669}$ (S66):** JF$_{669}$ (**15**; 150 mg, 0.286 mmol) and *tert*-butyl cyanoacetate (49.0 μL, 0.343 mmol, 1.2 eq) were combined in DMF (3 mL), and $K_2CO_3$ (94.8 mg, 0.686 mmol, 2.4 eq) was added. After stirring the reaction at room temperature for 48 h, it was diluted with water and extracted with EtOAc (2×). The combined organic extracts were washed with brine, dried over anhydrous $MgSO_4$, filtered, and evaporated. Purification of the crude by silica gel chromatography (10–75% EtOAc/hexanes, linear gradient) yielded 155 mg (84%) of **S66** as a blue-green solid. $^1$H NMR ($CDCl_3$, 400 MHz) δ 6.73 – 6.67 (m, 2H), 6.67 – 6.63 (m, 2H), 6.29 (dd, $J$ = 8.7, 2.7 Hz, 2H), 5.03 (s, 1H), 3.92 (t, $J$ = 7.3 Hz, 8H), 2.44 – 2.33 (m, 4H), 1.46 (s, 9H), 0.56 (s, 3H), 0.55 (s, 3H); $^{19}$F NMR ($CDCl_3$, 376 MHz) δ −117.51 (d, $J$ = 22.4 Hz, 1F), −132.27 (d, $J$ = 20.5 Hz, 1F), −141.37 – −141.55 (m, 1F); Analytical HPLC: $t_R$ = 10.9 min, 99.0% purity (30–95% MeCN/$H_2O$, linear gradient, with constant 0.1% v/v TFA additive; 20 min run; 1 mL/min flow; ESI; positive ion mode; detection at 675 nm); HRMS (ESI) calcd for $C_{35}H_{35}F_3N_3O_4Si$ $[M+H]^+$ 646.2343, found 646.2350.

### CYCLOADDITIONS OF 6-AZIDO-JF$_{669}$

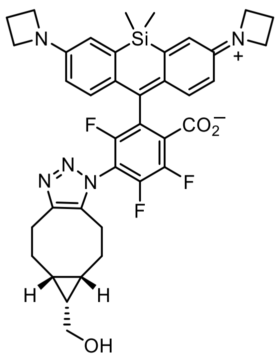


**JF$_{669}$-BCN triazole (27):** 6-Azido-JF$_{669}$ (**21**; 40 mg, 73.0 μmol) and (1*R*,8*S*,9*s*)-bicyclo[6.1.0]non-4-yn-9-ylmethanol (**25**; 14.3 mg, 95.0 μmol, 1.3 eq) were combined in DMF (1.5 mL) and stirred at room temperature for 1 h. The reaction was concentrated to dryness and purified by flash chromatography on silica gel (25–100% EtOAc/$CH_2Cl_2$, linear gradient) to afford **27** as a dark blue-green solid (48.2 mg, 95%). $^1H$ NMR (DMSO-$d_6$, 400 MHz, 350 K) δ 6.87 (d, *J* = 8.6 Hz, 2H), 6.71 (d, *J* = 2.6 Hz, 2H), 6.38 – 6.32 (m, 2H), 4.04 – 3.99 (m, 1H), 3.89 (t, *J* = 7.3 Hz, 8H), 3.56 – 3.43 (m, 2H), 3.14 – 3.07 (m, 1H), 2.93 – 2.77 (m, 2H), 2.69 – 2.55 (m, 1H), 2.34 (p, *J* = 7.2 Hz, 4H), 2.19 – 2.09 (m, 1H), 2.07 – 1.98 (m, 1H), 1.64 – 1.49 (m, 2H), 1.07 – 0.96 (m, 1H), 0.94 – 0.75 (m, 2H), 0.54 (s, 3H), 0.48 (s, 3H); $^{19}F$ NMR (DMSO-$d_6$, 376 MHz, 350 K) δ −124.27 (d, *J* = 20.8 Hz, 1F), −137.21 (d, *J* = 22.1 Hz, 1F), −140.63 (t, *J* = 21.3 Hz, 1F); Analytical HPLC: $t_R$ = 11.7 min, >99% purity (10–95% MeCN/$H_2O$, linear gradient, with constant 0.1% v/v TFA additive; 20 min run; 1 mL/min flow; ESI; positive ion mode; detection at 675 nm); HRMS (ESI) calcd for $C_{38}H_{39}F_3N_5O_3Si$ $[M+H]^+$ 698.2769, found 698.2779.

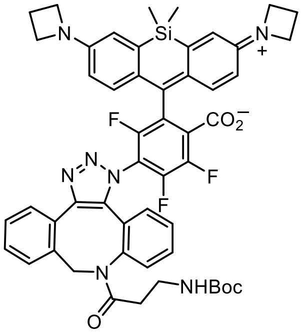


**JF$_{669}$-DBCO triazole (28):** 6-Azido-JF$_{669}$ (**21**; 30 mg, 54.8 μmol) and DBCO-NH-Boc (**26**; 24.8 mg, 65.7 μmol, 1.2 eq) were combined in DMF (1 mL) and stirred at room temperature for 1 h. The reaction was concentrated to dryness and purified by flash chromatography on silica gel (25–100% EtOAc/hexanes, linear gradient) to provide the desired product **28** as a mixture of regioisomers (pale green solid, 50.1 mg, 99%). Although the NMR spectra were not interpretable, HPLC and HRMS analyses were consistent with the expected product mixture. Analytical HPLC: $t_R$ (major isomer) = 11.5 min, 60.3% by peak integration; $t_R$ (minor isomer) = 11.9 min, 39.7% by peak integration (45–55% MeCN/$H_2O$, linear gradient, with constant 0.1% v/v TFA additive; 20 min run; 1 mL/min flow; ESI; positive ion mode; detection at 675 nm); HRMS (ESI) calcd for $C_{51}H_{49}F_3N_7O_5Si$ $[M+H]^+$ 924.3511, found 924.3531.

### X-RAY CRYSTALLOGRAPHY

#### Confirmation of the regioselectivity of masked acyl cyanide (MAC) substitution

*Single crystal X-ray diffraction (SC-XRD) of* **S57** *and* **S79**

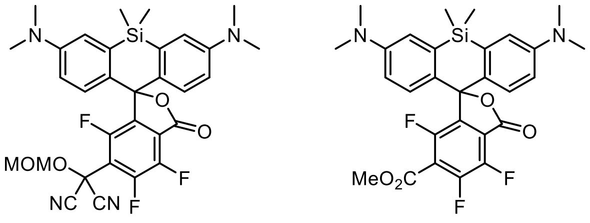

**S57** **S79**

**MAC substitution product S57:** Crystallization and SC-XRD of **S57** were performed by Ardena in Amsterdam, The Netherlands. Crystals of adequate quality for SC-XRD were obtained by vapor diffusion crystallization from 1,4-dioxane (solvent) and water (anti-solvent). The single crystal measurements were performed on a Nonius Kappa-CCD. The data were collected at 296 K. The full sphere data were collected up to θ = 32.7° (20840 reflections). Data reduction was performed using HKL Scalepack and cell parameters were obtained using Denzo and Scalepak from 11297 reflections within θ range 1 to 32.7°.[10] The structure was solved using direct methods by SHELXT-2014/7.[11] The structure was refined by least square full matrix refinement using SHELXL-2014/7.[12] All H-atoms were included from the geometry and kept with fixed thermal parameters. The obtained crystal structure (**Figure S1**) confirmed that substitution by the MAC reagent occurred exclusively at the position *para* to the carbonyl.

**Methyl ester S79:** Crystallization and SC-XRD of **S79** were performed by Ardena in Amsterdam, The Netherlands. Crystals of adequate quality for SC-XRD were obtained by vapor diffusion crystallization from EtOAc (solvent) and heptane (anti-solvent). The single crystal measurements were performed on a Nonius Kappa-CCD. The data were collected at 296 K. The full sphere data were collected up to θ = 32.6° (18406 reflections). Data reduction was performed using HKL Scalepack and cell parameters were obtained using Denzo and Scalepak from 9087 reflections within θ range 1 to 32.6°.[10] The structure was solved using direct methods by SHELXT-2014/7.[11] The structure was refined by least square full matrix refinement using SHELXL-2014/7.[12] All H-atoms were included from the geometry and kept with fixed thermal parameters. During the refinement the relatively high peak ~2.4 e/$Å^3$ was observed in the symmetry center associated with peaks of lower intensities. Attempts to model these peaks as solvent were unsuccessful (blue balls in the figure). During the final cycles it was found out that carbonyl O atom from the methyl ester group is disordered in two positions with following occupancy factors 0.65(5) and 0.35(5). Regardless, the obtained crystal structure (**Figure S1**) was consistent with the regiochemistry of the structure assigned to **S57**, from which **S79** was derived.

[10] Otwinowski, Z.; Minor, W. *Methods Enzymol.* **1997**, *276*, 307–326.
[11] Sheldrick, G. M. *Acta Crystallogr., Sect. A: Found. Adv.* **2015**, *71*, 3–8.
[12] Sheldrick, G. M. *Acta Crystallogr., Sect. C: Struct. Chem.* **2015**, *71*, 3–8.

**Table S1.** Crystal data and structure refinement for **S57** and **S79**.

| | MAC product **S57** | Methyl ester **S79** |
|---|---|---|
| Empirical Formula | $C_{31}H_{29}F_3N_4O_4Si \cdot 1.5(C_4H_8O_2)$ | $C_{28}H_{27}F_3N_2O_4Si \cdot 1/6(C_4H_8O_2)$ |
| Formula Weight | 738.83 | 540.62 + 14.66 |
| Temperature | 296(2) K | 296(2) K |
| λ | 0.71073 Å | 0.71073 Å |
| Crystal System, Space Group | Triclinic, P-1 | Trigonal, R-3 |
| Unit Cell Dimensions | a = 9.4514(6) Å | a = 30.3536(10) Å |
| | b = 12.9790(9) Å | |
| | c = 15.4757(12) Å | c = 15.4361(3) Å |
| | α = 88.026(1)° | |
| | β = 81.890(1)° | |
| | γ = 87.402(1)° | |
| Volume | 1876.7(2) $Å^3$ | 12316.5(8) $Å^3$ |
| Z | 2 | 18 |
| $D_c$ | 1.307 g/$cm^3$ | 1.357 g/$cm^3$ |
| μ | 0.130 $mm^{-1}$ | 0.145 $mm^{-1}$ |
| F(000) | 776 | 5244 |
| Crystal Size | 0.35 × 0.22 × 0.20 $mm^3$ | 0.45 × 0.30 × 0.22 $mm^3$ |
| θ Range for Data Collection | 2.7–32.7° | 2.0–32.6° |
| Reflections Collected | 20840 | 18406 |
| Independent Reflections | 13660 [$R_{int}$ = 0.0258] | 9910 [$R_{int}$ = 0.0267] |
| Completeness to θ = 25.242° | 99.7 | 99.3 |
| Absorption Correction | Integration | Integration |
| Max. and Min. Transmission | 0.990 and 0.958 | 0.978 and 0.953 |
| Data / Restraint / Parameters | 13660 / 0 / 476 | 9910 / 2 / 367 |
| Goodness-of-Fit on $F^2$ | 1.023 | 1.021 |
| Final R Indices [I>2σ(I)] | R1 = 0.0673, wR2 = 0.1801 | R1 = 0.0582, wR2 = 0.1478 |
| R Indices (All Data) | R1 = 0.1248, wR2 = 0.2170 | R1 = 1.077, wR2 = 0.1894 |
| Largest Diff. Peak and Hole | 0.490 and -0.324 e/$Å^3$ | 0.302 and -0.458 e/$Å^3$ |

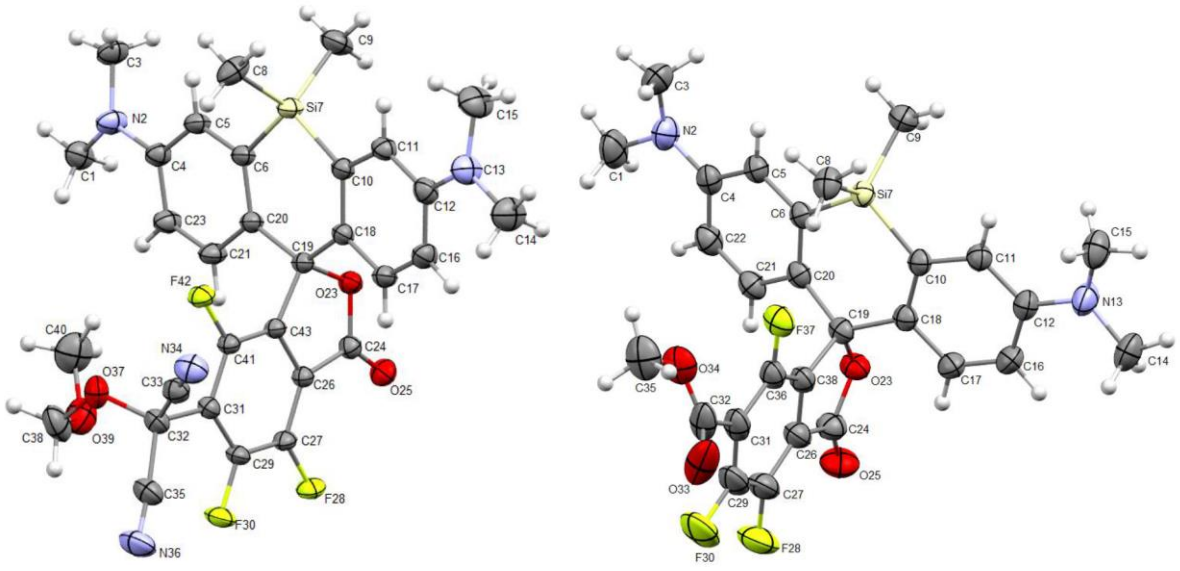


**Figure S1.** Molecular structure and atoms numbering scheme for **S57** (left) and **S79** (right); displacement ellipsoids at the 30% probability level.

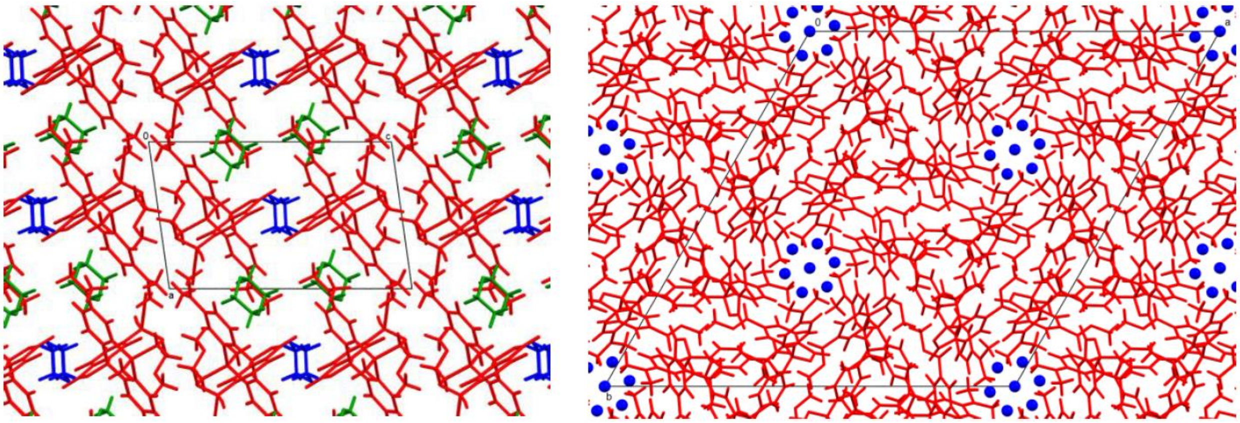

**Figure S2.** Packing along ***b*** axis in the crystal of **S57**-1,4-dioxane solvate (left) and along ***c*** axis in the crystal of **S79**-EtOAc solvate (right). In both images, **S57** and **S79** are shown in red while the solvent molecules are blue and green (left) and blue (right).

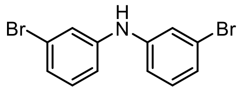

Br
H
N
Br


**S3**

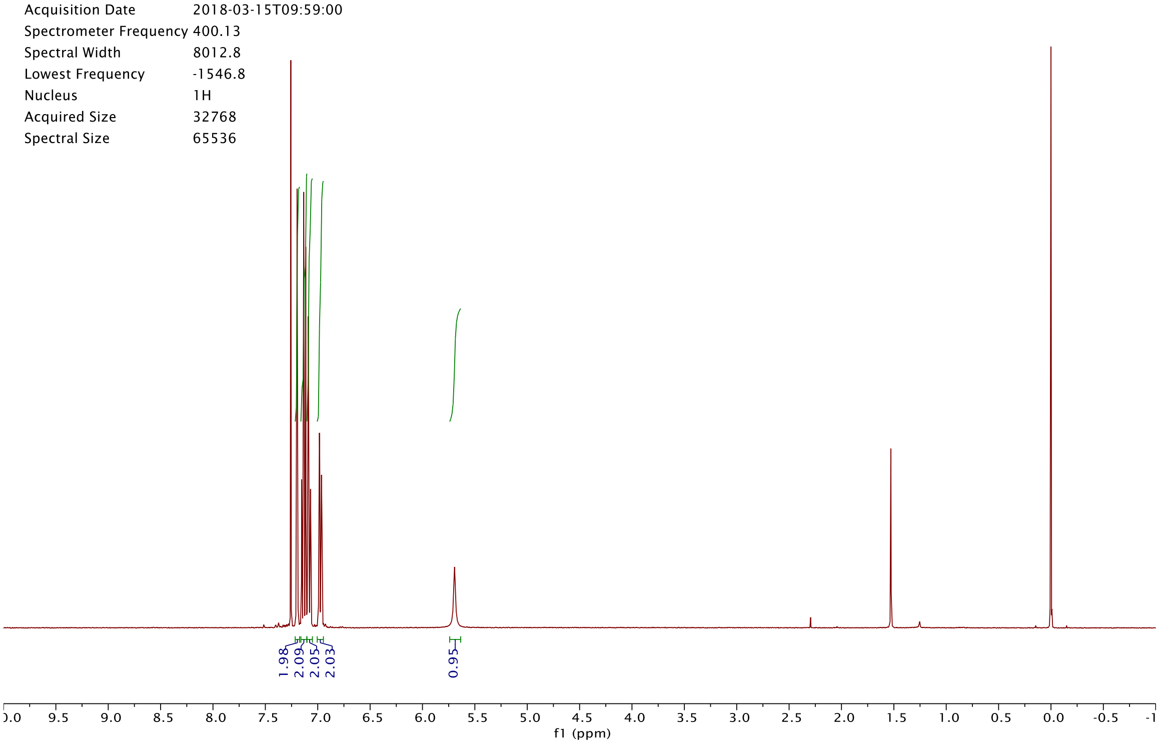

Origin Bruker BioSpin GmbH
Solvent CDCl3
Temperature 300.0
Pulse Sequence zg30
Experiment 1D
Number of Scans 16
Acquisition Date 2018-03-15T09:59:00
Spectrometer Frequency 400.13
Spectral Width 8012.8
Lowest Frequency -1546.8
Nucleus 1H
Acquired Size 32768
Spectral Size 65536
1.98
2.09
2.05
2.03
0.95
f1 (ppm)


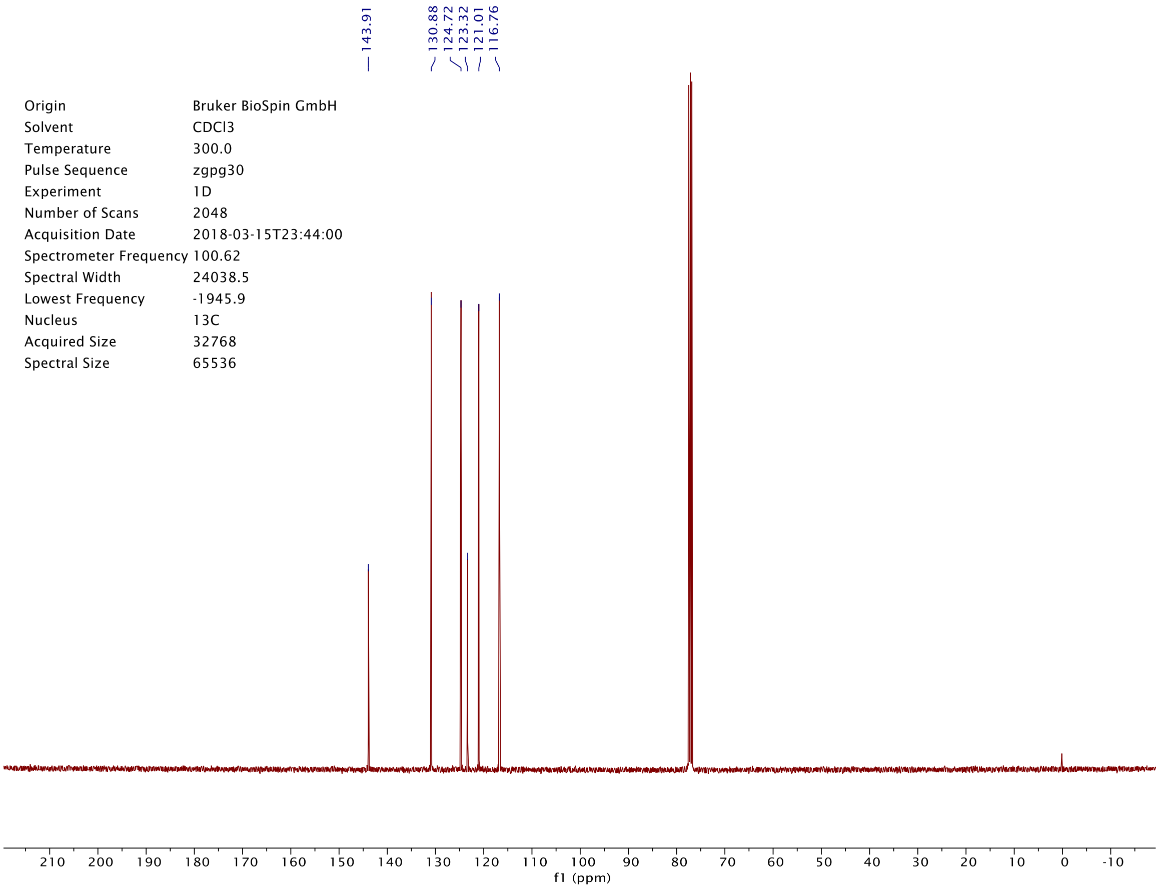

143.91
130.88
124.72
123.32
121.01
116.76
Origin Bruker BioSpin GmbH
Solvent CDCl3
Temperature 300.0
Pulse Sequence zgpg30
Experiment 1D
Number of Scans 2048
Acquisition Date 2018-03-15T23:44:00
Spectrometer Frequency 100.62
Spectral Width 24038.5
Lowest Frequency -1945.9
Nucleus 13C
Acquired Size 32768
Spectral Size 65536
f1 (ppm)

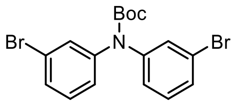
Boc
N
Br
Br

**S4**

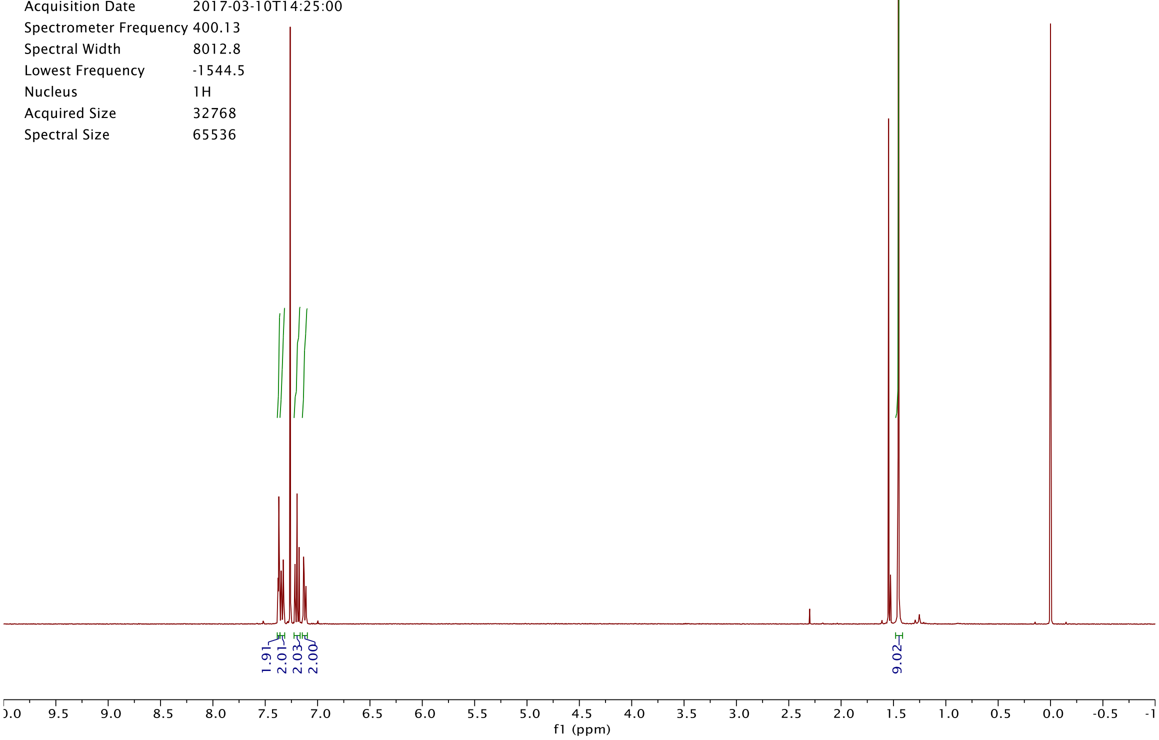
Origin Bruker BioSpin GmbH
Solvent CDCl3
Temperature 295.4
Pulse Sequence zg30
Experiment 1D
Number of Scans 16
Acquisition Date 2017-03-10T14:25:00
Spectrometer Frequency 400.13
Spectral Width 8012.8
Lowest Frequency -1544.5
Nucleus 1H
Acquired Size 32768
Spectral Size 65536
1.91
2.01
2.03
2.00
9.02
f1 (ppm)

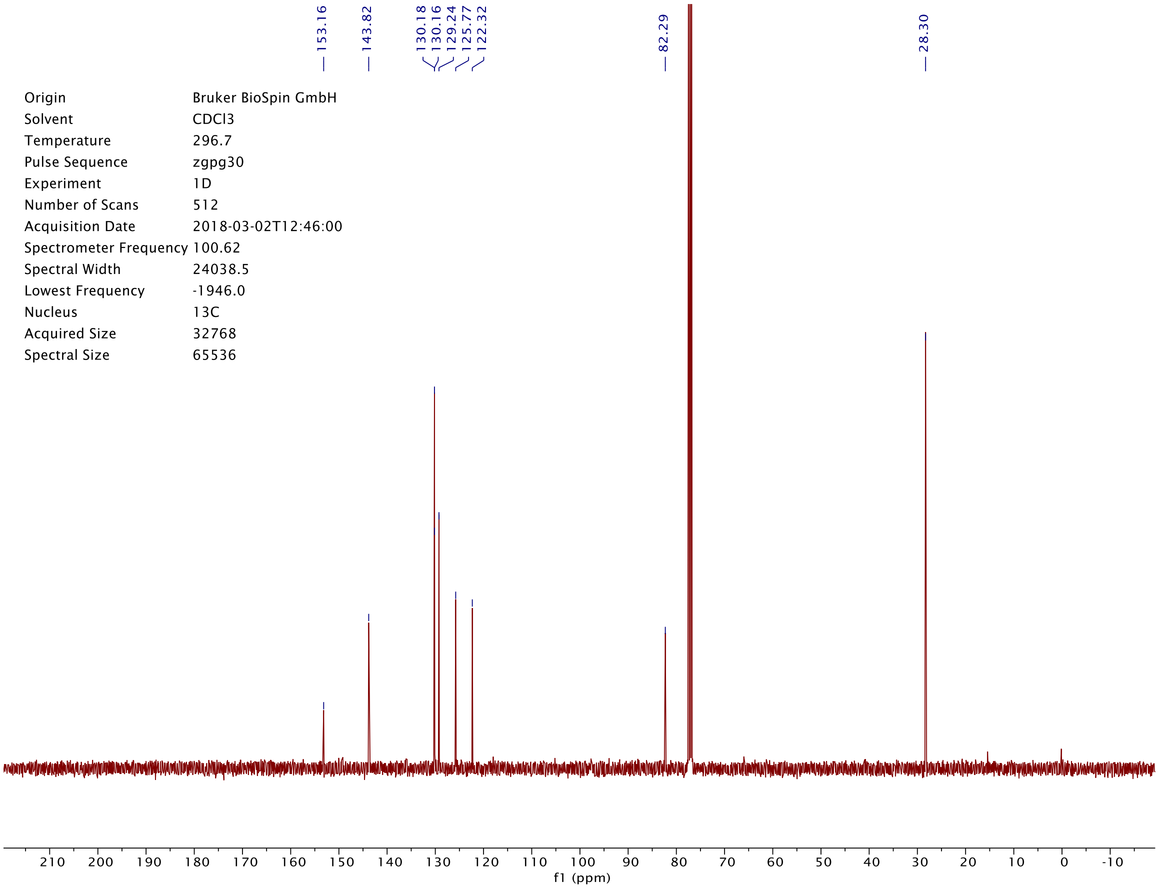
153.16
143.82
130.18
130.16
129.24
125.77
122.32
82.29
28.30
Origin Bruker BioSpin GmbH
Solvent CDCl3
Temperature 296.7
Pulse Sequence zgpg30
Experiment 1D
Number of Scans 512
Acquisition Date 2018-03-02T12:46:00
Spectrometer Frequency 100.62
Spectral Width 24038.5
Lowest Frequency -1946.0
Nucleus 13C
Acquired Size 32768
Spectral Size 65536
f1 (ppm)

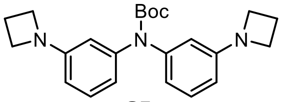

**S5**

| | |
|---|---|
| Origin | Bruker BioSpin GmbH |
| Solvent | CDCl3 |
| Temperature | 300.0 |
| Pulse Sequence | zg30 |
| Experiment | 1D |
| Number of Scans | 16 |
| Acquisition Date | 2018-03-15T19:05:00 |
| Spectrometer Frequency | 400.13 |
| Spectral Width | 8012.8 |
| Lowest Frequency | -1547.9 |
| Nucleus | 1H |
| Acquired Size | 32768 |
| Spectral Size | 65536 |

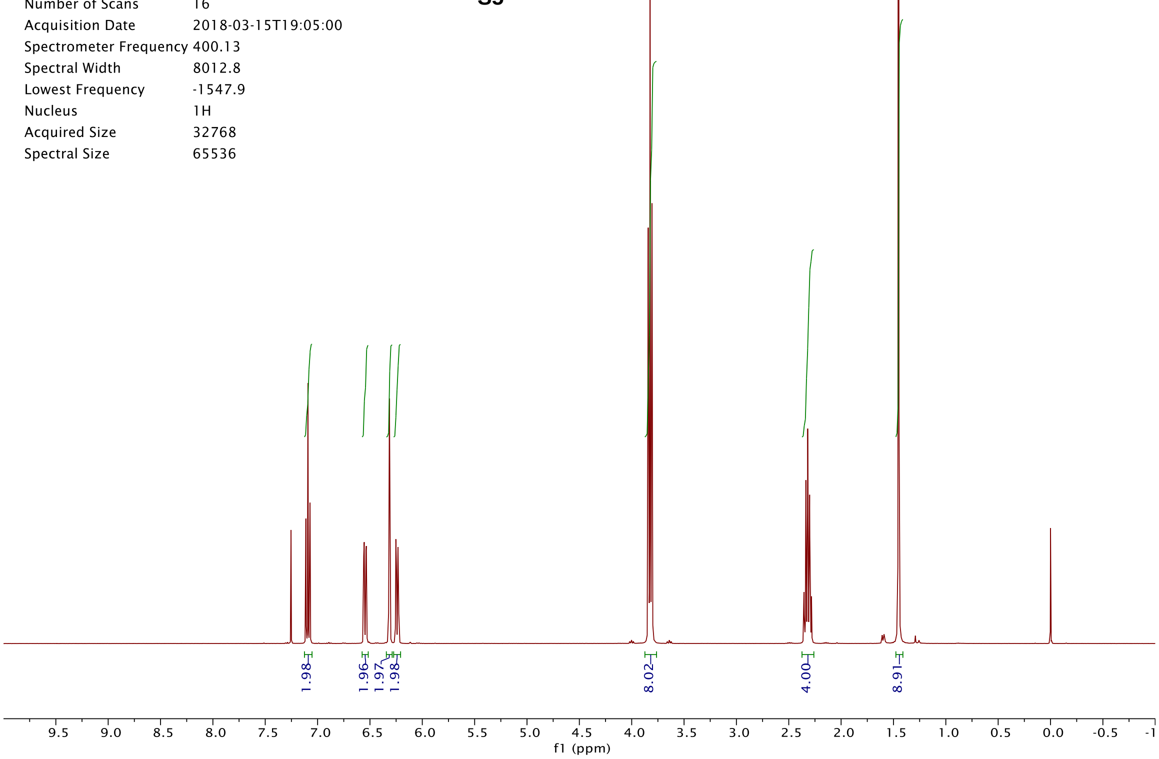


| | |
|---|---|
| Origin | Bruker BioSpin GmbH |
| Solvent | CDCl3 |
| Temperature | 300.0 |
| Pulse Sequence | zgpg30 |
| Experiment | 1D |
| Number of Scans | 2048 |
| Acquisition Date | 2018-03-15T21:04:00 |
| Spectrometer Frequency | 100.62 |
| Spectral Width | 24038.5 |
| Lowest Frequency | -1946.5 |
| Nucleus | 13C |
| Acquired Size | 32768 |
| Spectral Size | 65536 |

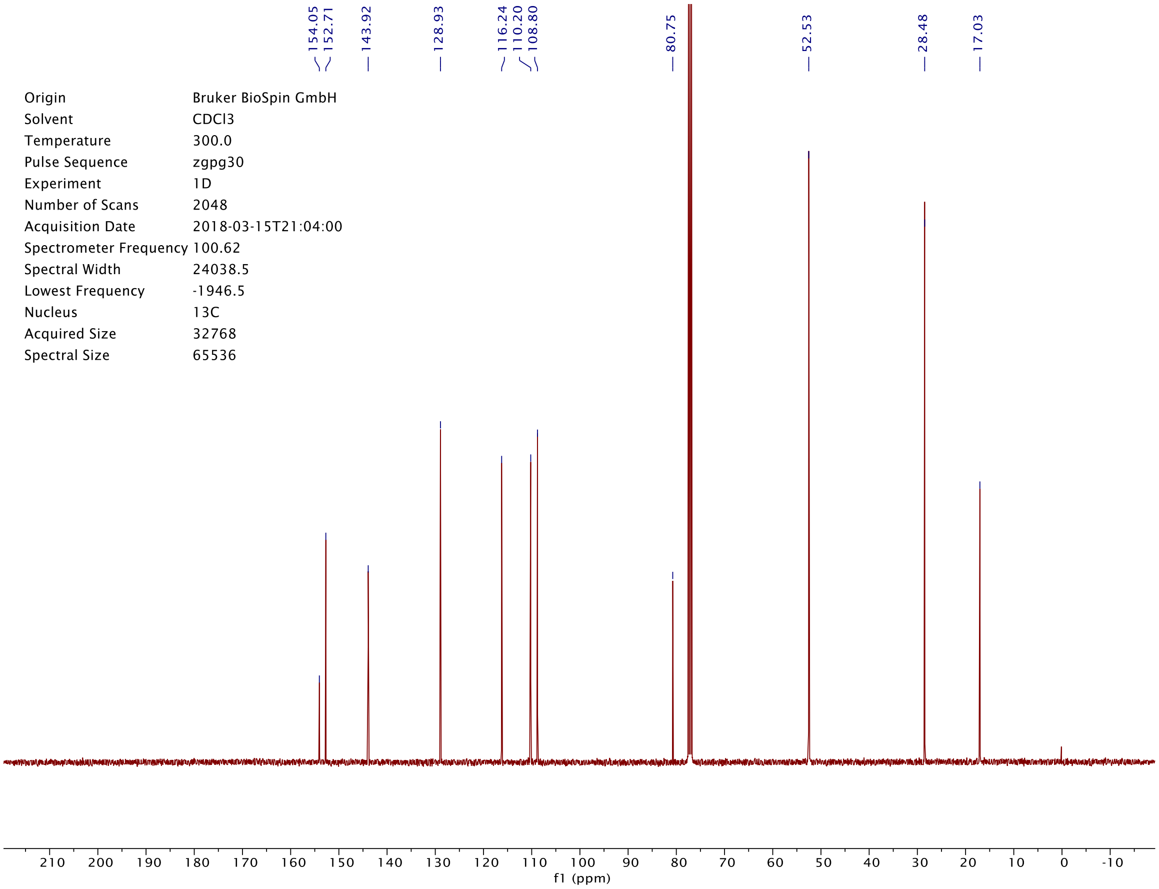

| | |
|---|---|
| Origin | Bruker BioSpin GmbH |
| Solvent | CDCl3 |
| Temperature | 300.0 |
| Pulse Sequence | zg30 |
| Experiment | 1D |
| Number of Scans | 16 |
| Acquisition Date | 2018-03-16T00:25:00 |
| Spectrometer Frequency | 400.13 |
| Spectral Width | 8012.8 |
| Lowest Frequency | -1547.2 |
| Nucleus | 1H |
| Acquired Size | 32768 |
| Spectral Size | 65536 |

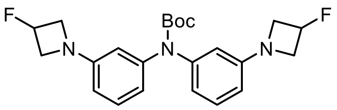


**S6**

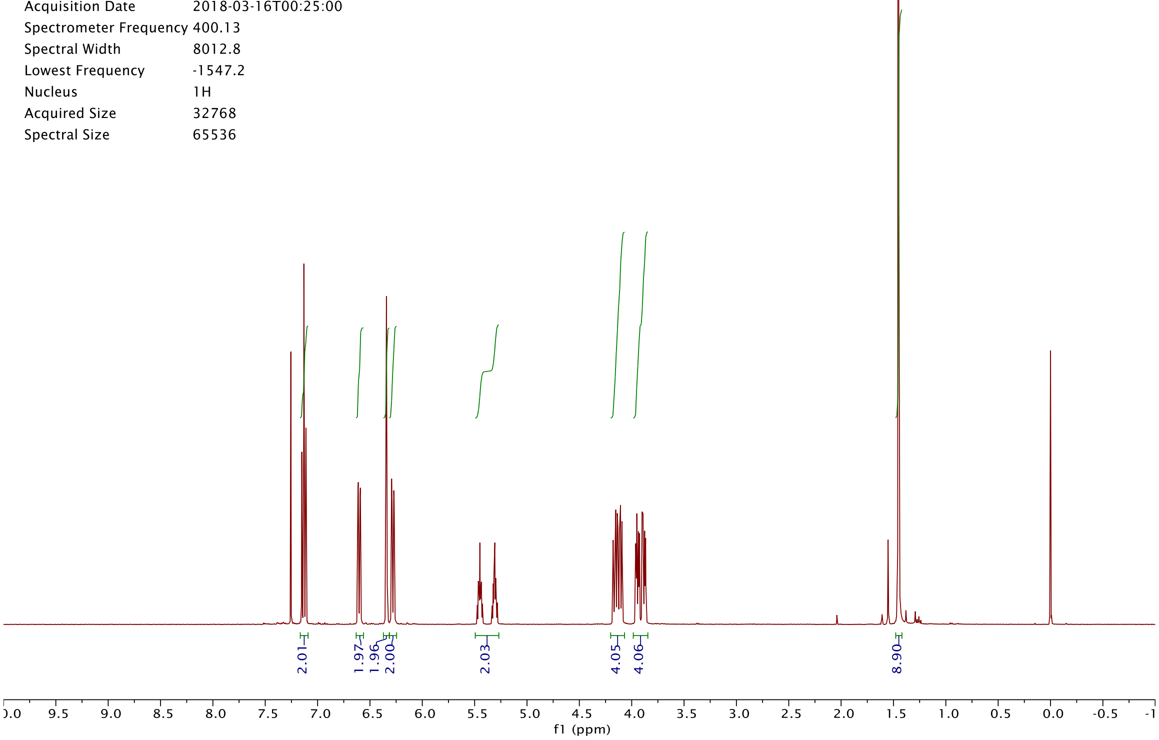


| | |
|---|---|
| Origin | Bruker BioSpin GmbH |
| Solvent | CDCl3 |
| Temperature | 300.0 |
| Pulse Sequence | zgpg30 |
| Experiment | 1D |
| Number of Scans | 2048 |
| Acquisition Date | 2018-03-16T02:24:00 |
| Spectrometer Frequency | 100.62 |
| Spectral Width | 24038.5 |
| Lowest Frequency | -1945.0 |
| Nucleus | 13C |
| Acquired Size | 32768 |
| Spectral Size | 65536 |

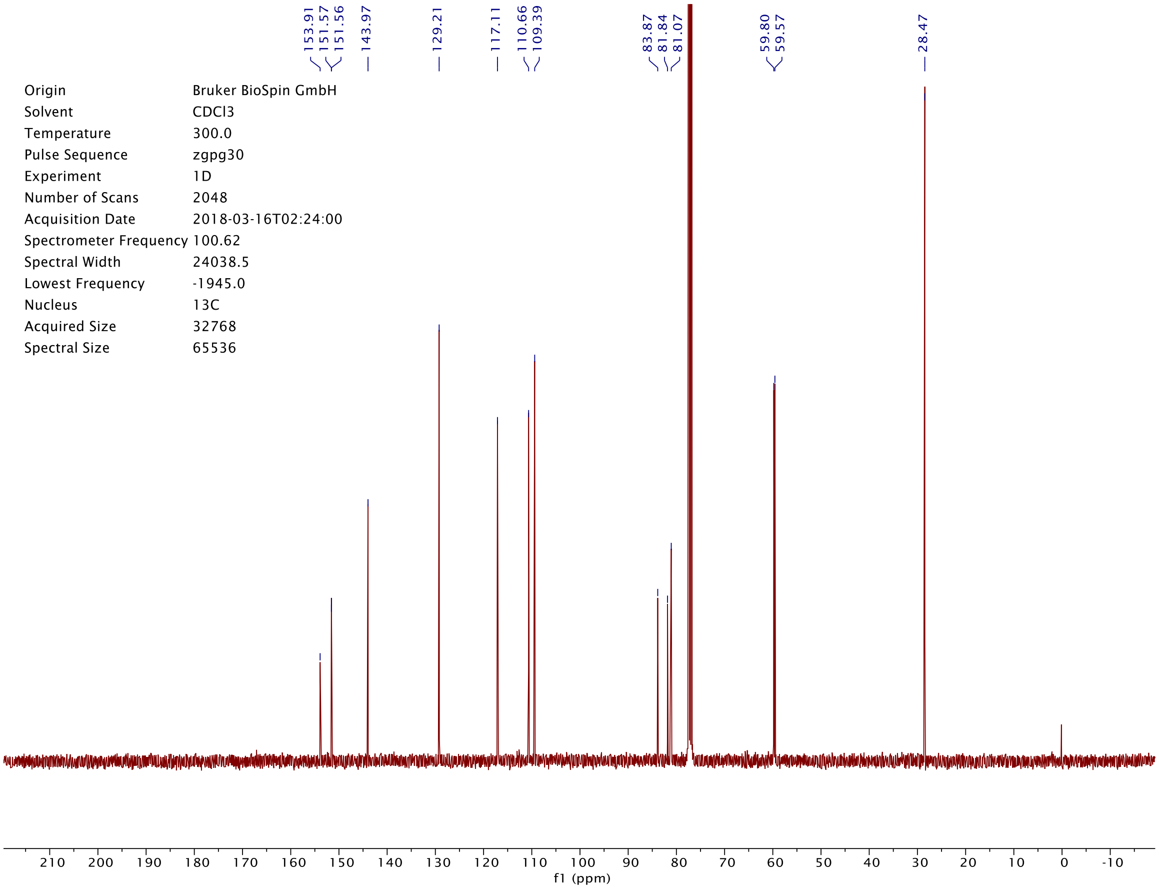

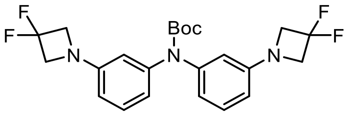


**S7**

| | |
|---|---|
| Origin | Bruker BioSpin GmbH |
| Solvent | CDCl3 |
| Temperature | 295.4 |
| Pulse Sequence | zg30 |
| Experiment | 1D |
| Number of Scans | 16 |
| Acquisition Date | 2018-01-04T11:07:00 |
| Spectrometer Frequency | 400.13 |
| Spectral Width | 8012.8 |
| Lowest Frequency | -1545.7 |
| Nucleus | 1H |
| Acquired Size | 32768 |
| Spectral Size | 65536 |

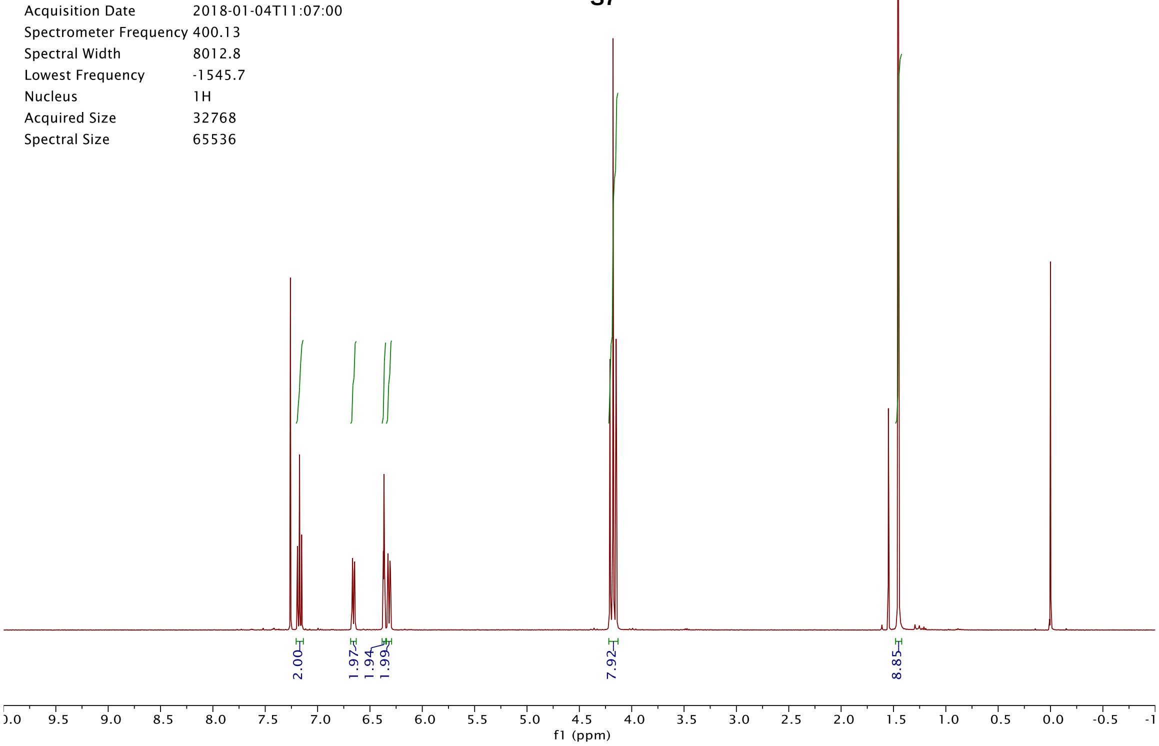


| | |
|---|---|
| Origin | Bruker BioSpin GmbH |
| Solvent | CDCl3 |
| Temperature | 300.0 |
| Pulse Sequence | zgpg30 |
| Experiment | 1D |
| Number of Scans | 2048 |
| Acquisition Date | 2018-03-02T01:24:00 |
| Spectrometer Frequency | 100.62 |
| Spectral Width | 24038.5 |
| Lowest Frequency | -1945.0 |
| Nucleus | 13C |
| Acquired Size | 32768 |
| Spectral Size | 65536 |

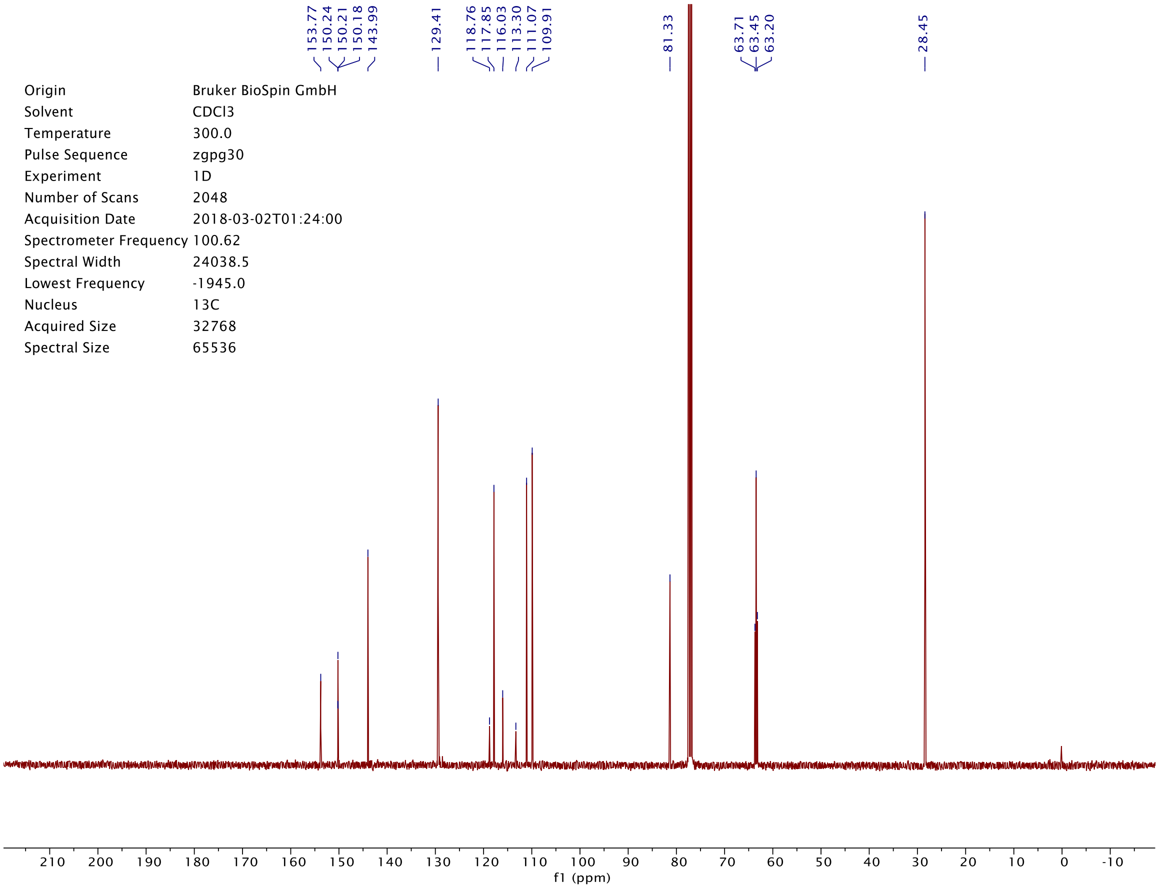

| | |
|---|---|
| Origin | Bruker BioSpin GmbH |
| Solvent | DMSO |
| Temperature | 350.2 |
| Pulse Sequence | zg30 |
| Experiment | 1D |
| Number of Scans | 16 |
| Acquisition Date | 2018-03-26T17:02:00 |
| Spectrometer Frequency | 400.13 |
| Spectral Width | 8012.8 |
| Lowest Frequency | -1538.4 |
| Nucleus | 1H |
| Acquired Size | 32768 |
| Spectral Size | 65536 |

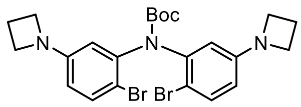


**S8**

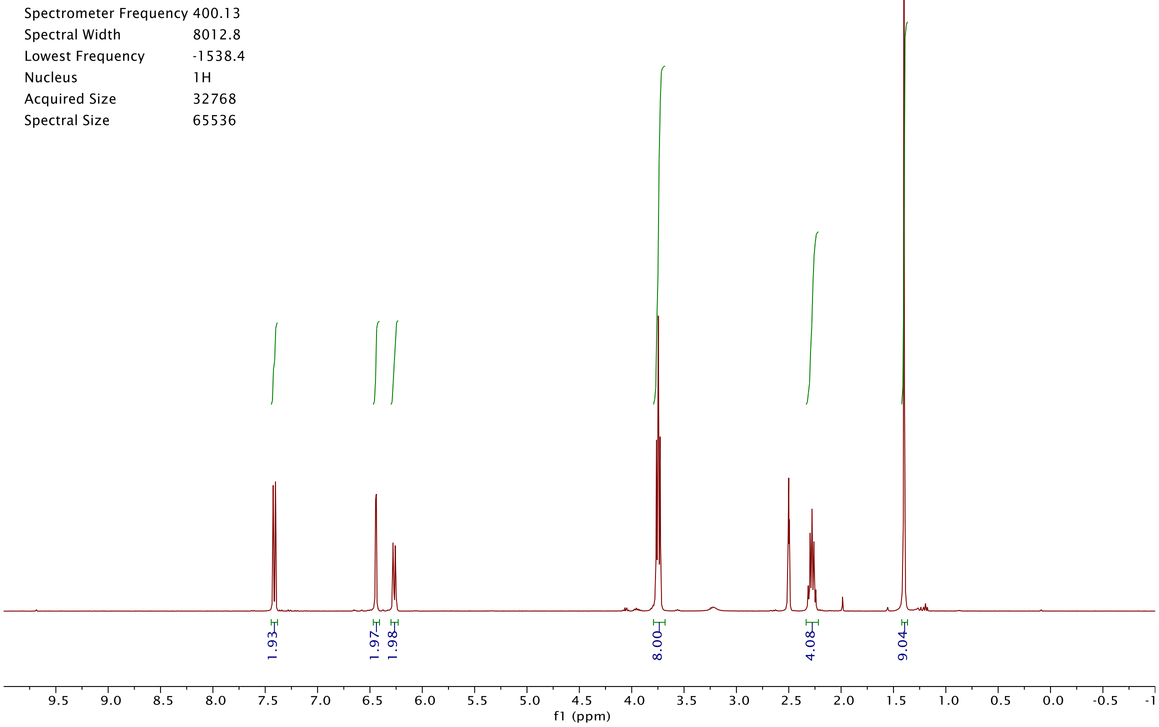


| | |
|---|---|
| Origin | Bruker BioSpin GmbH |
| Solvent | DMSO |
| Temperature | 350.0 |
| Pulse Sequence | zgpg30 |
| Experiment | 1D |
| Number of Scans | 2048 |
| Acquisition Date | 2018-03-26T19:01:00 |
| Spectrometer Frequency | 100.62 |
| Spectral Width | 24038.5 |
| Lowest Frequency | -2047.1 |
| Nucleus | 13C |
| Acquired Size | 32768 |
| Spectral Size | 65536 |

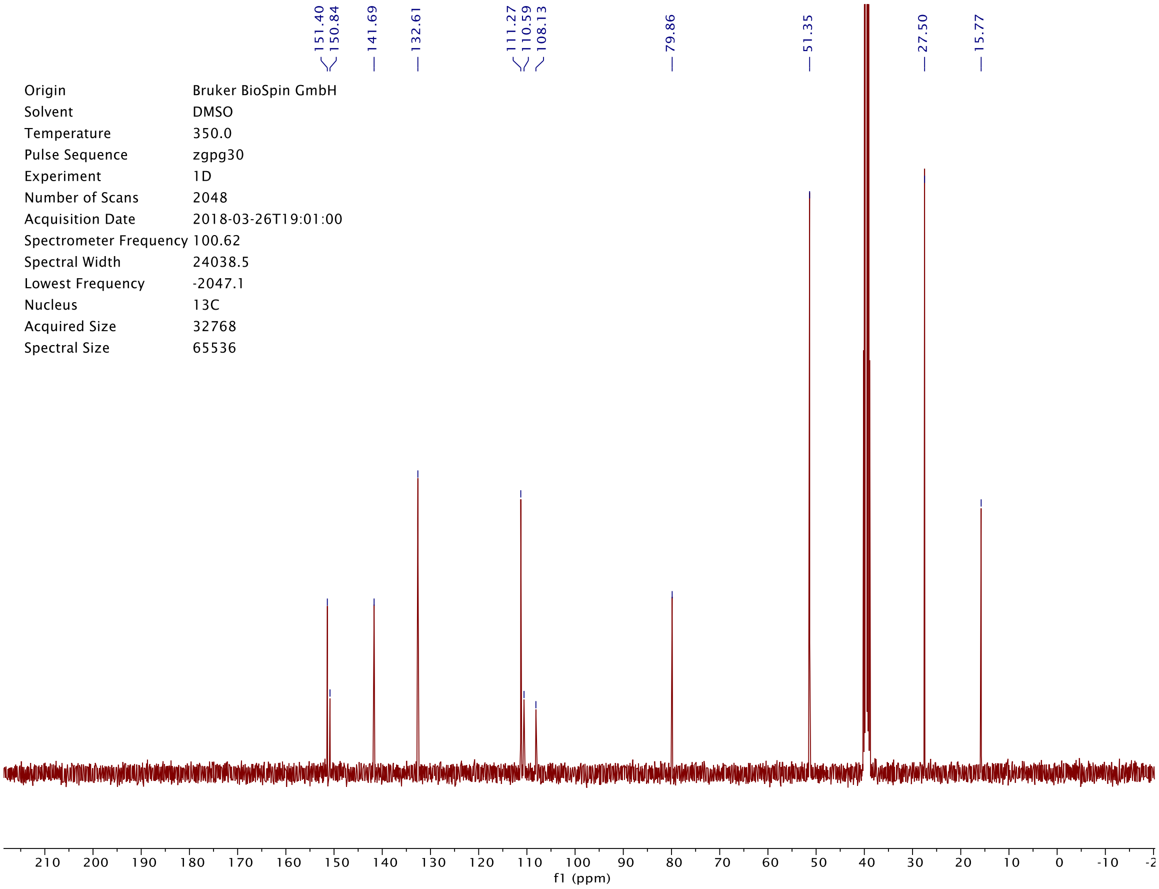

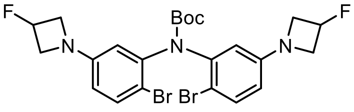
F
Boc
N
N
N
F
Br Br

**S9**

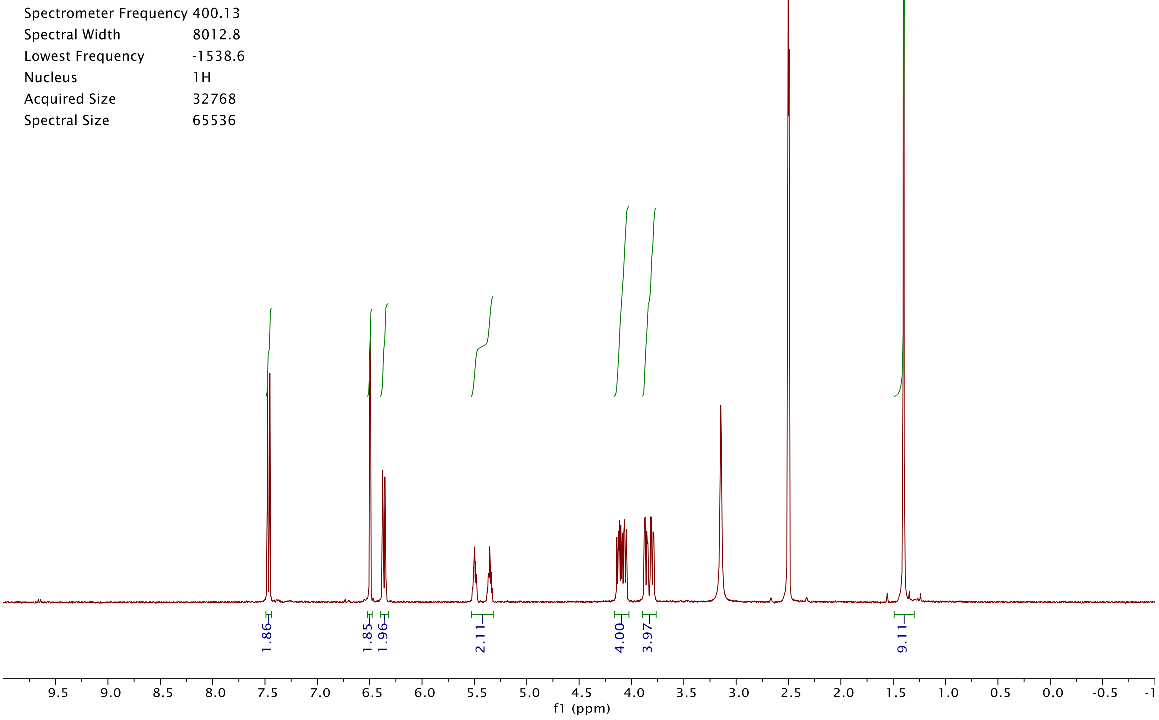
Origin Bruker BioSpin GmbH
Solvent DMSO
Temperature 350.2
Pulse Sequence zg30
Experiment 1D
Number of Scans 16
Acquisition Date 2018-03-26T13:30:00
Spectrometer Frequency 400.13
Spectral Width 8012.8
Lowest Frequency -1538.6
Nucleus 1H
Acquired Size 32768
Spectral Size 65536
1.86
1.85
1.96
2.11
4.00
3.97
9.11
9.5 9.0 8.5 8.0 7.5 7.0 6.5 6.0 5.5 5.0 4.5 4.0 3.5 3.0 2.5 2.0 1.5 1.0 0.5 0.0 -0.5 -1
f1 (ppm)

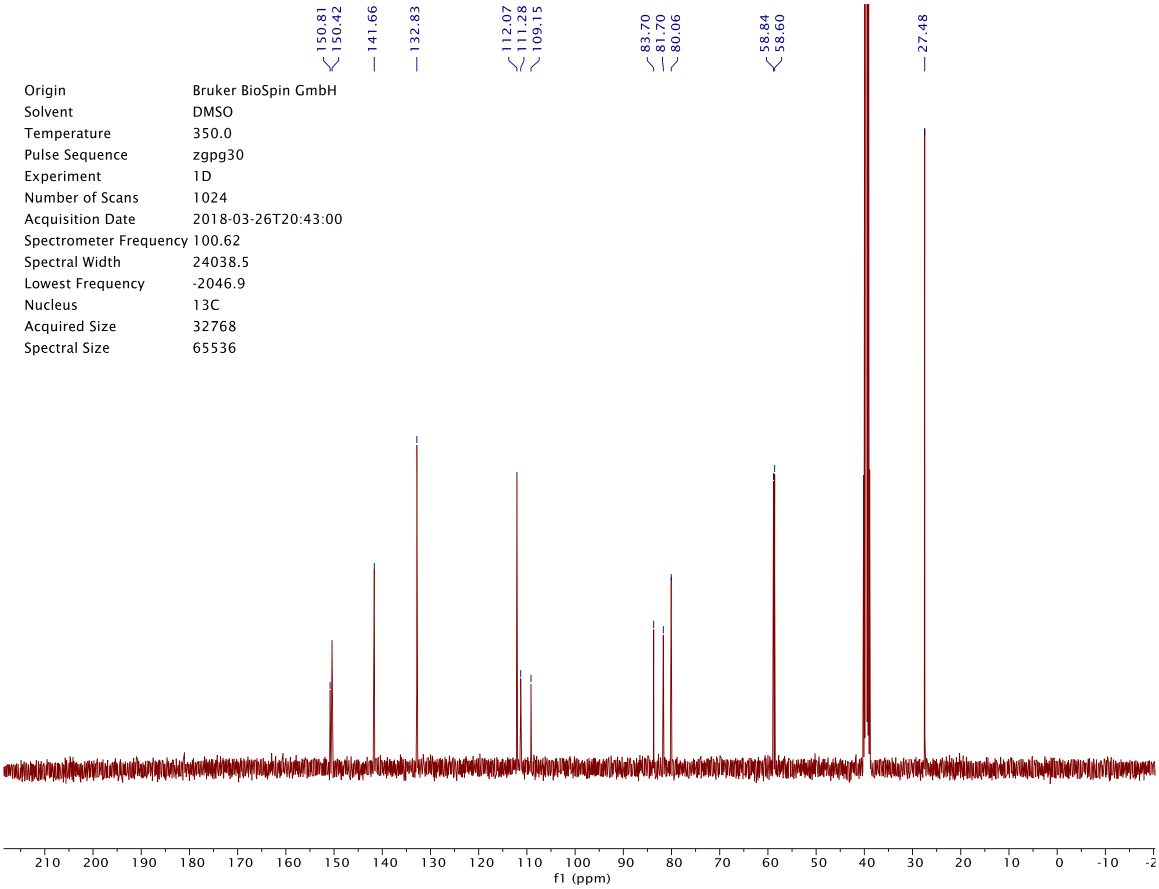
150.81
150.42
141.66
132.83
112.07
111.28
109.15
83.70
81.70
80.06
58.84
58.60
27.48
Origin Bruker BioSpin GmbH
Solvent DMSO
Temperature 350.0
Pulse Sequence zgpg30
Experiment 1D
Number of Scans 1024
Acquisition Date 2018-03-26T20:43:00
Spectrometer Frequency 100.62
Spectral Width 24038.5
Lowest Frequency -2046.9
Nucleus 13C
Acquired Size 32768
Spectral Size 65536
210 200 190 180 170 160 150 140 130 120 110 100 90 80 70 60 50 40 30 20 10 0 -10
f1 (ppm)

| | |
|---|---|
| Origin | Bruker BioSpin GmbH |
| Solvent | DMSO |
| Temperature | 350.2 |
| Pulse Sequence | zg30 |
| Experiment | 1D |
| Number of Scans | 16 |
| Acquisition Date | 2018-03-16T11:52:00 |
| Spectrometer Frequency | 400.13 |
| Spectral Width | 8012.8 |
| Lowest Frequency | -1538.6 |
| Nucleus | 1H |
| Acquired Size | 32768 |
| Spectral Size | 65536 |

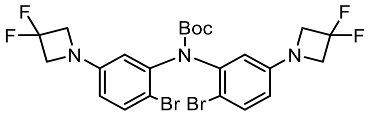


**S10**

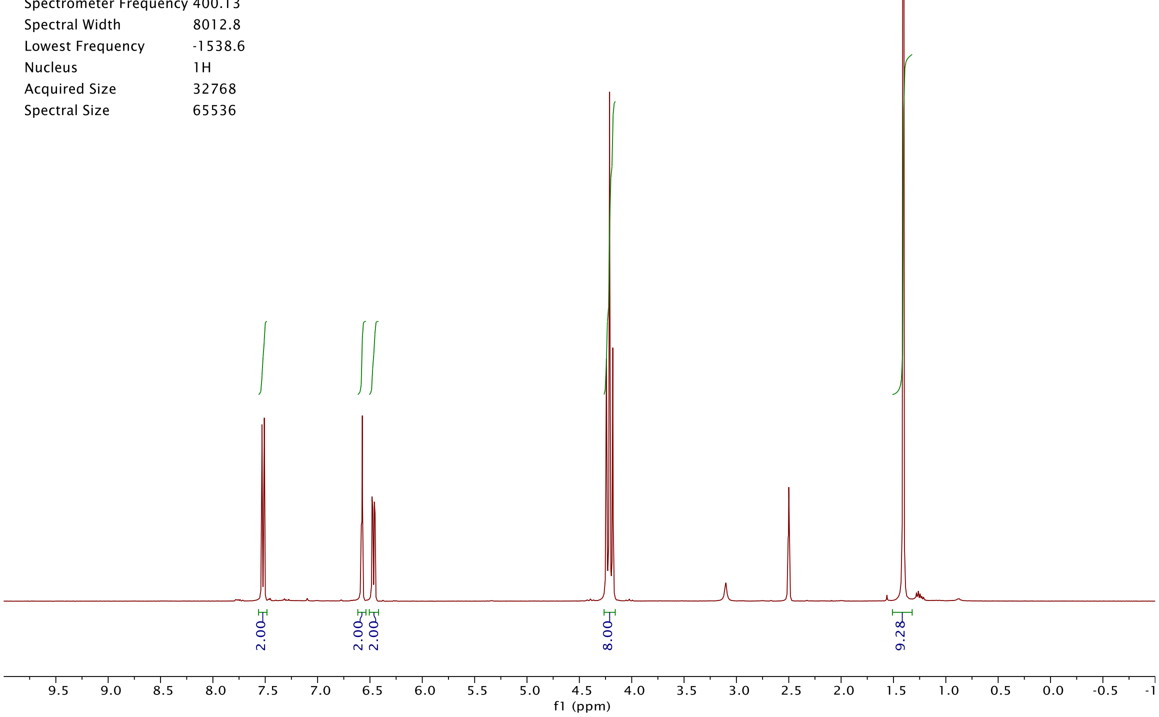


| | |
|---|---|
| Origin | Bruker BioSpin GmbH |
| Solvent | DMSO |
| Temperature | 350.0 |
| Pulse Sequence | zgpg30 |
| Experiment | 1D |
| Number of Scans | 1024 |
| Acquisition Date | 2018-03-16T12:53:00 |
| Spectrometer Frequency | 100.62 |
| Spectral Width | 24038.5 |
| Lowest Frequency | -2046.9 |
| Nucleus | 13C |
| Acquired Size | 32768 |
| Spectral Size | 65536 |

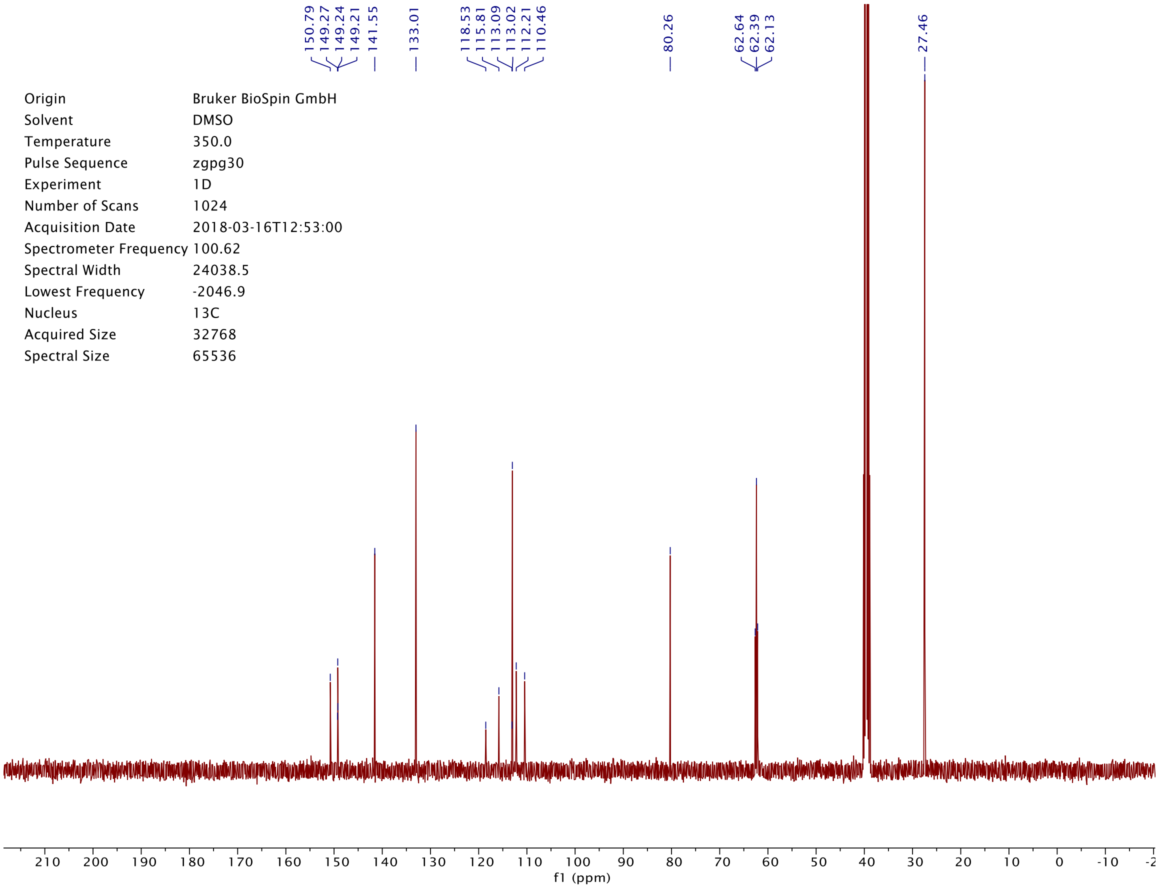

| | |
|---|---|
| Origin | Bruker BioSpin GmbH |
| Solvent | CDCl3 |
| Temperature | 300.0 |
| Pulse Sequence | zg30 |
| Experiment | 1D |
| Number of Scans | 16 |
| Acquisition Date | 2017-05-25T15:38:00 |
| Spectrometer Frequency | 400.13 |
| Spectral Width | 8012.8 |
| Lowest Frequency | -1546.8 |
| Nucleus | 1H |
| Acquired Size | 32768 |
| Spectral Size | 65536 |

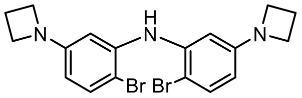


**S11**

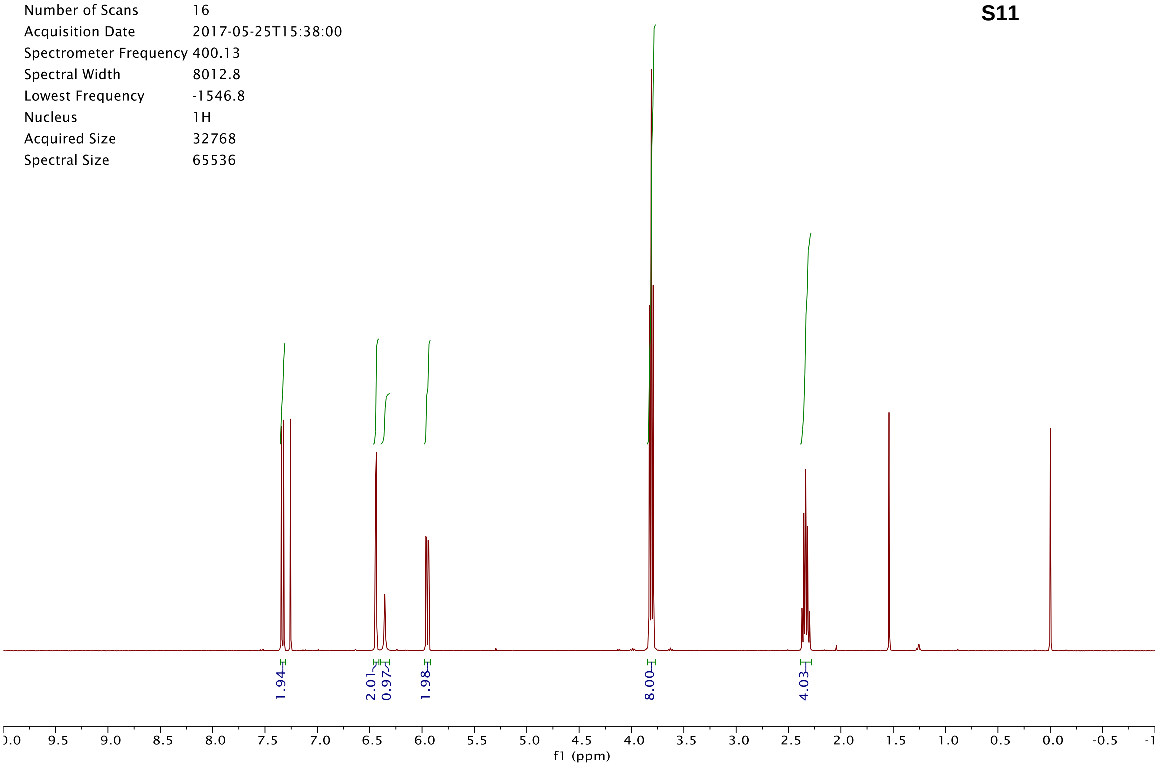


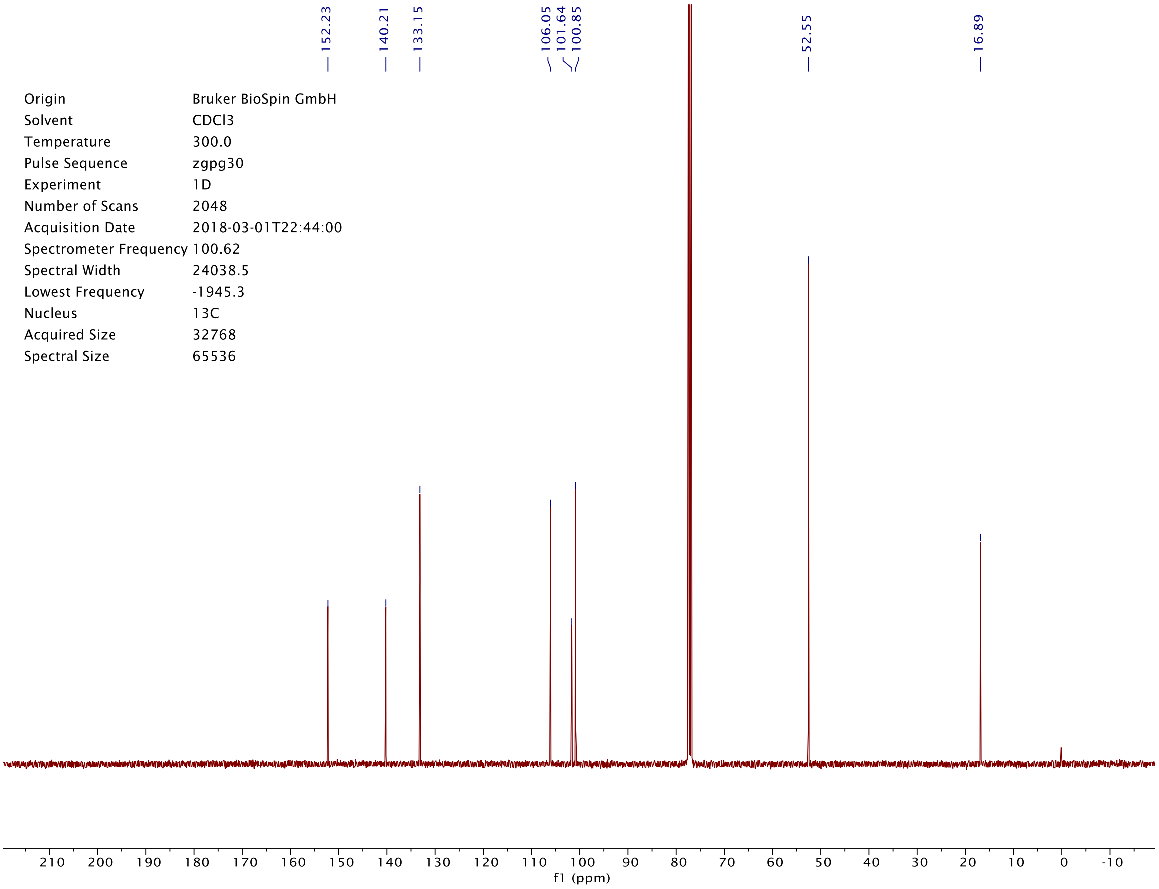


| | |
|---|---|
| Origin | Bruker BioSpin GmbH |
| Solvent | CDCl3 |
| Temperature | 300.0 |
| Pulse Sequence | zgpg30 |
| Experiment | 1D |
| Number of Scans | 2048 |
| Acquisition Date | 2018-03-01T22:44:00 |
| Spectrometer Frequency | 100.62 |
| Spectral Width | 24038.5 |
| Lowest Frequency | -1945.3 |
| Nucleus | 13C |
| Acquired Size | 32768 |
| Spectral Size | 65536 |

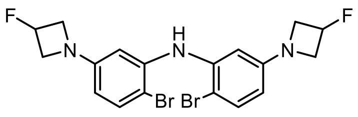
F
N
H
N
N
F
Br Br

**S12**

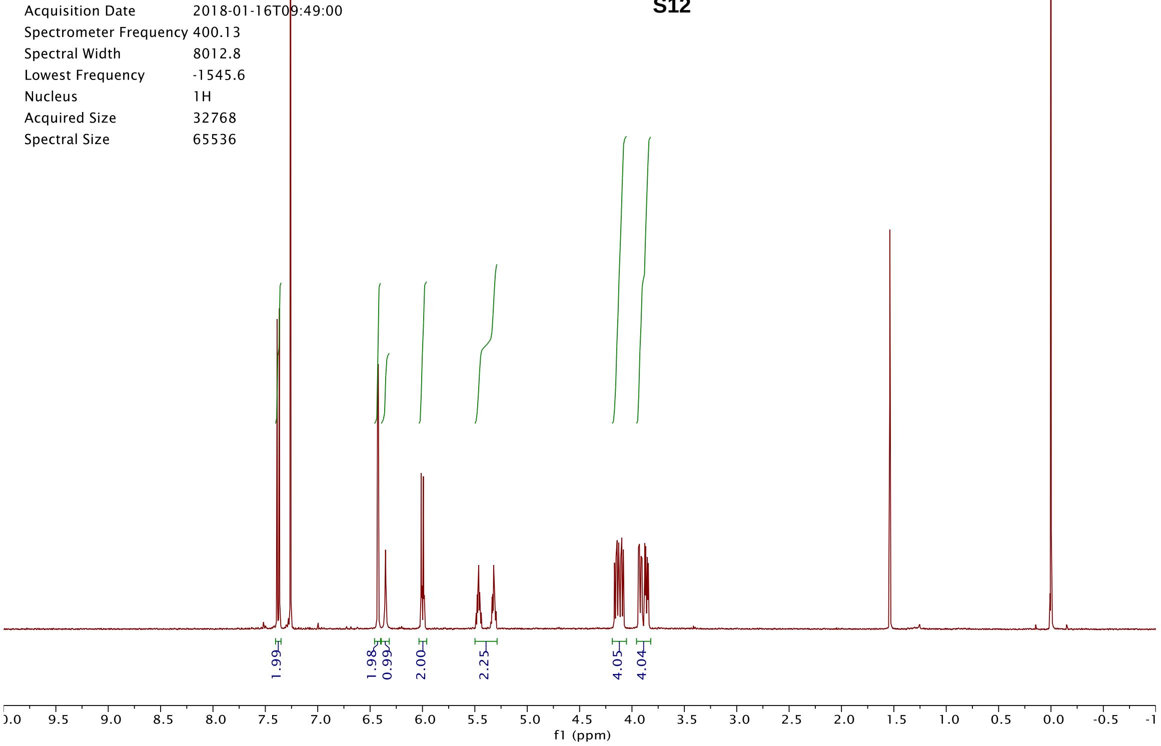
Origin Bruker BioSpin GmbH
Solvent CDCl3
Temperature 295.6
Pulse Sequence zg30
Experiment 1D
Number of Scans 16
Acquisition Date 2018-01-16T09:49:00
Spectrometer Frequency 400.13
Spectral Width 8012.8
Lowest Frequency -1545.6
Nucleus 1H
Acquired Size 32768
Spectral Size 65536
1.99
1.98
0.99
2.00
2.25
4.05
4.04
f1 (ppm)

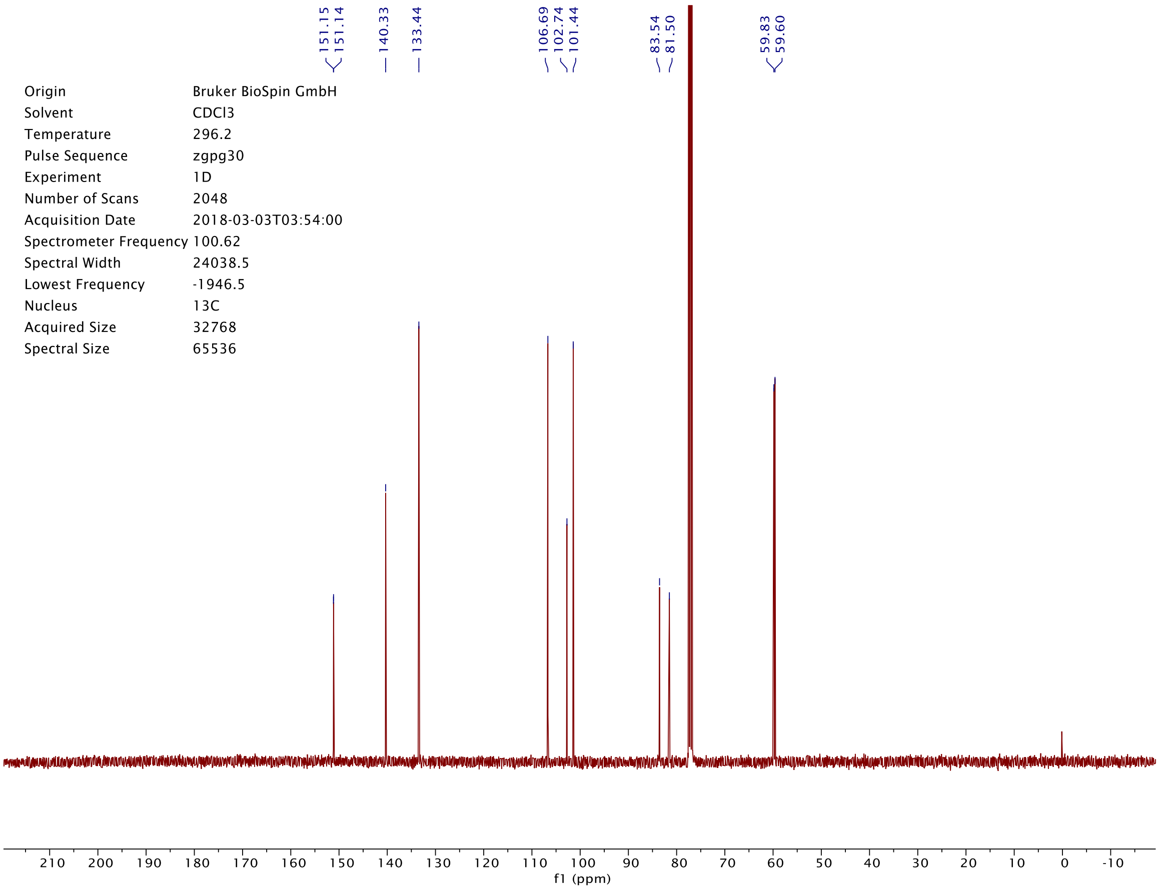
151.15
151.14
140.33
133.44
106.69
102.74
101.44
83.54
81.50
59.83
59.60
Origin Bruker BioSpin GmbH
Solvent CDCl3
Temperature 296.2
Pulse Sequence zgpg30
Experiment 1D
Number of Scans 2048
Acquisition Date 2018-03-03T03:54:00
Spectrometer Frequency 100.62
Spectral Width 24038.5
Lowest Frequency -1946.5
Nucleus 13C
Acquired Size 32768
Spectral Size 65536
f1 (ppm)

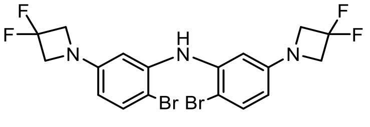

**S13**

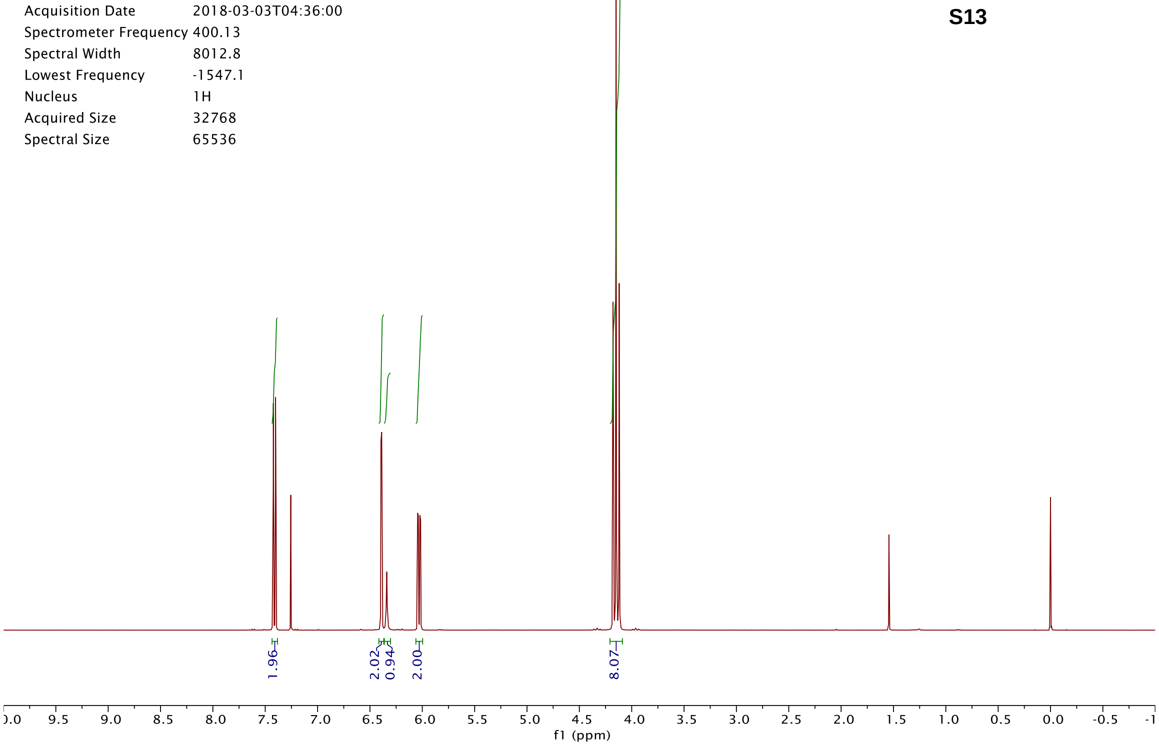

Origin Bruker BioSpin GmbH
Solvent CDCl3
Temperature 295.6
Pulse Sequence zg30
Experiment 1D
Number of Scans 16
Acquisition Date 2018-03-03T04:36:00
Spectrometer Frequency 400.13
Spectral Width 8012.8
Lowest Frequency -1547.1
Nucleus 1H
Acquired Size 32768
Spectral Size 65536
1.96
2.02
0.94
2.00
8.07
f1 (ppm)


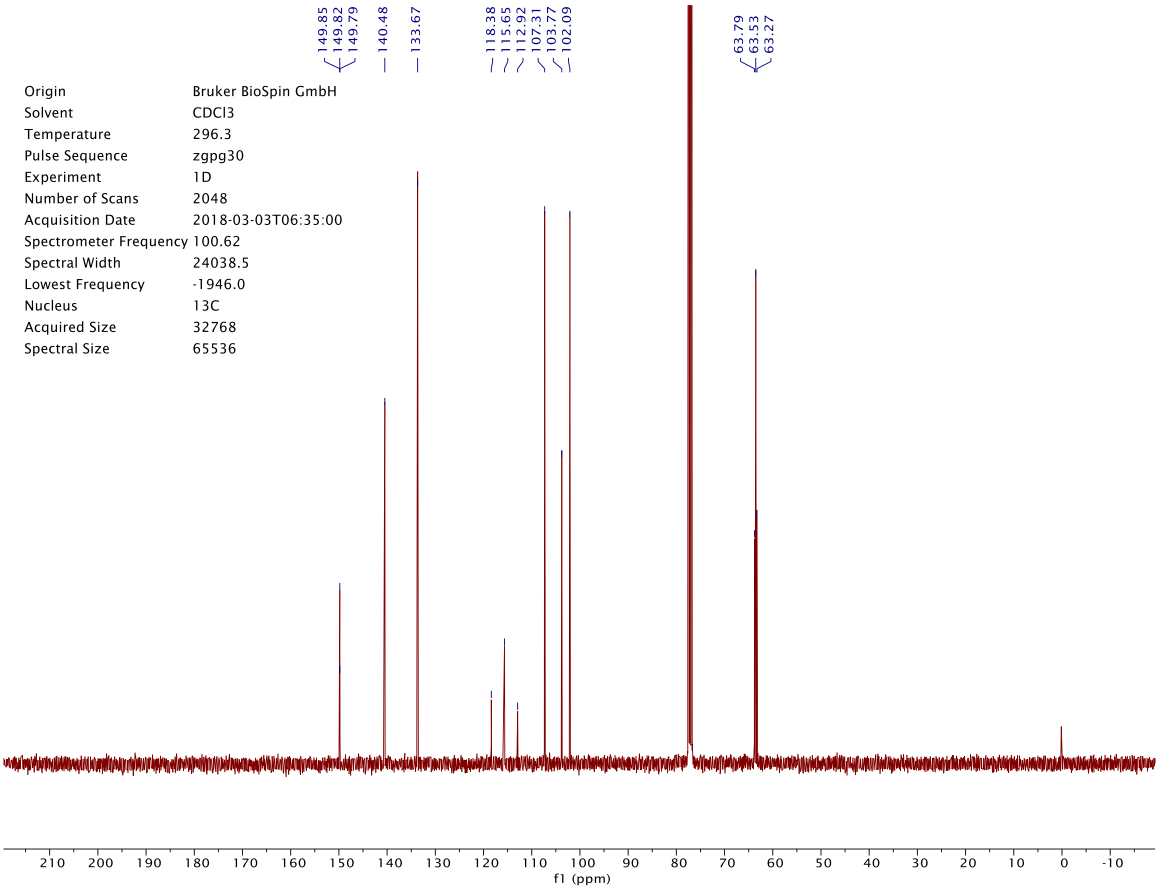

149.85
149.82
149.79
140.48
133.67
118.38
115.65
112.92
107.31
103.77
102.09
63.79
63.53
63.27
Origin Bruker BioSpin GmbH
Solvent CDCl3
Temperature 296.3
Pulse Sequence zgpg30
Experiment 1D
Number of Scans 2048
Acquisition Date 2018-03-03T06:35:00
Spectrometer Frequency 100.62
Spectral Width 24038.5
Lowest Frequency -1946.0
Nucleus 13C
Acquired Size 32768
Spectral Size 65536
f1 (ppm)

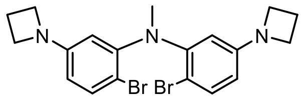


**S14**

| | |
|---|---|
| Origin | Bruker BioSpin GmbH |
| Solvent | CDCl3 |
| Temperature | 300.0 |
| Pulse Sequence | zg30 |
| Experiment | 1D |
| Number of Scans | 16 |
| Acquisition Date | 2017-05-30T15:55:00 |
| Spectrometer Frequency | 400.13 |
| Spectral Width | 8012.8 |
| Lowest Frequency | -1545.9 |
| Nucleus | 1H |
| Acquired Size | 32768 |
| Spectral Size | 65536 |

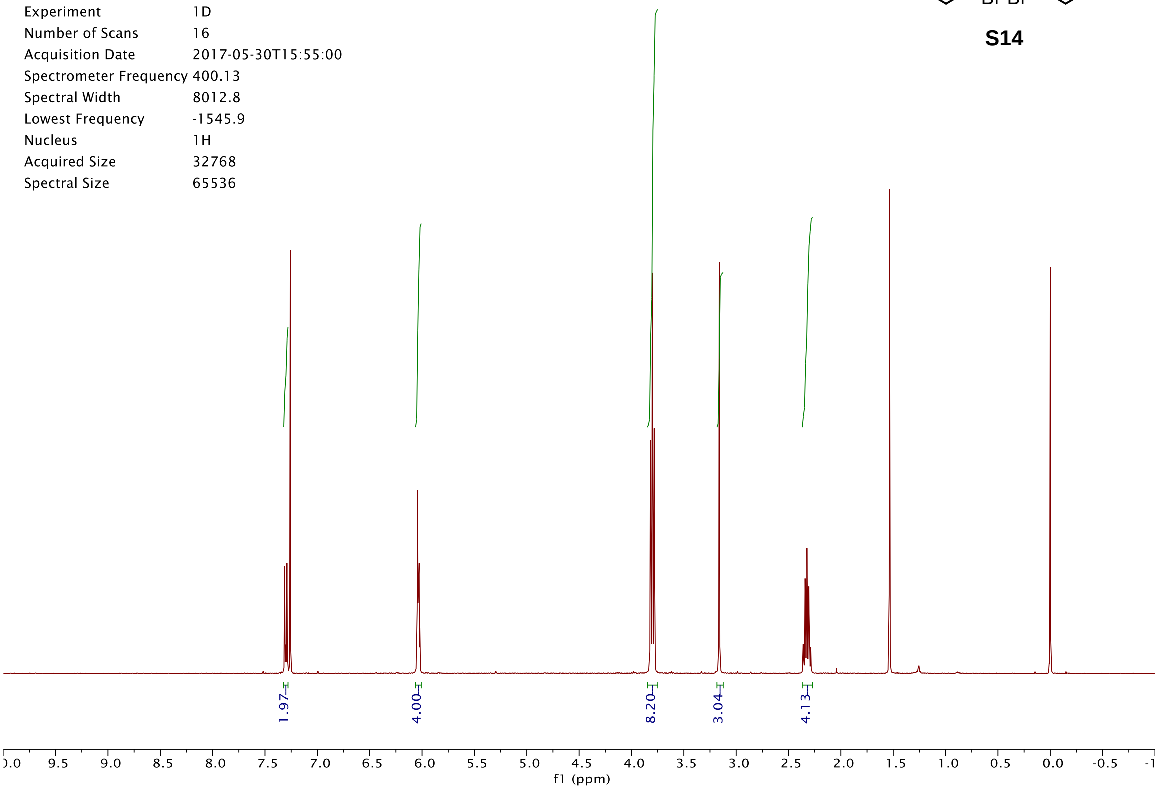


| | |
|---|---|
| Origin | Bruker BioSpin GmbH |
| Solvent | CDCl3 |
| Temperature | 296.2 |
| Pulse Sequence | zgpg30 |
| Experiment | 1D |
| Number of Scans | 2048 |
| Acquisition Date | 2018-03-02T20:02:00 |
| Spectrometer Frequency | 100.62 |
| Spectral Width | 24038.5 |
| Lowest Frequency | -1946.6 |
| Nucleus | 13C |
| Acquired Size | 32768 |
| Spectral Size | 65536 |

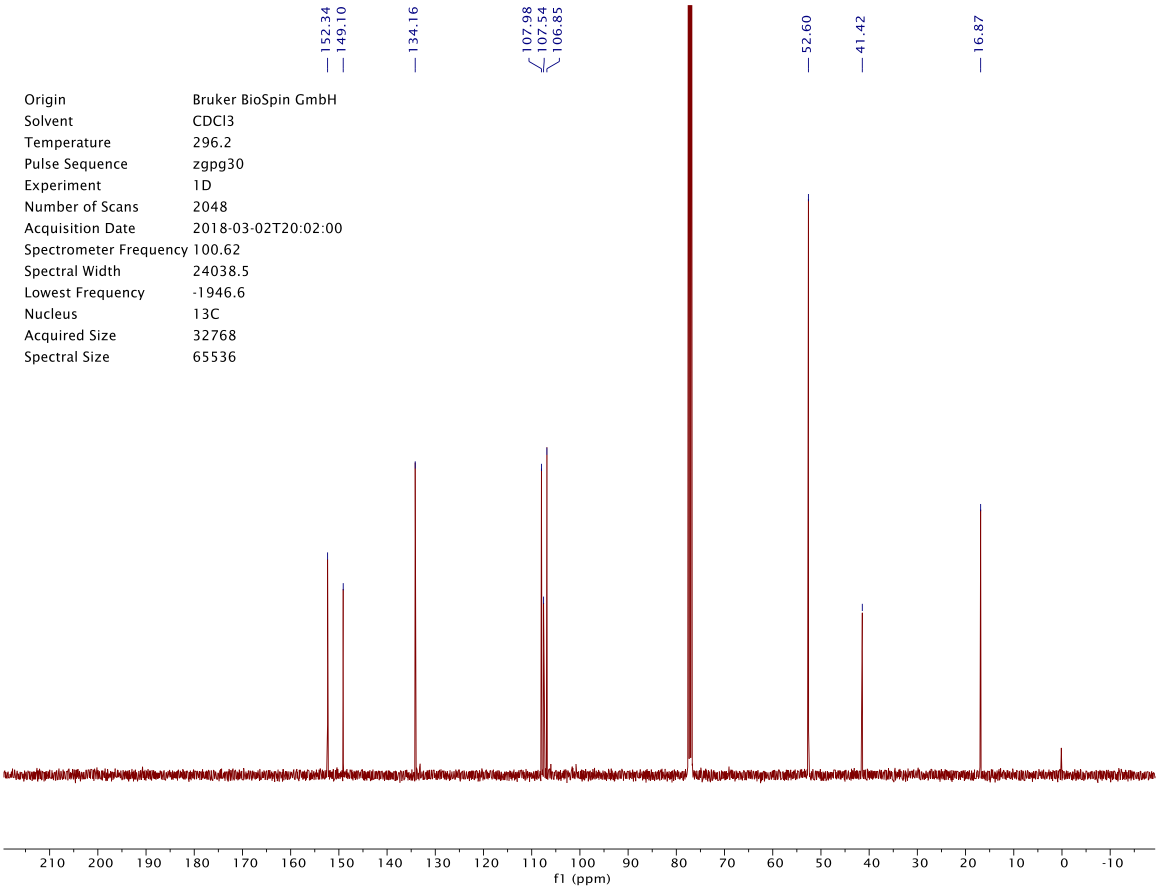

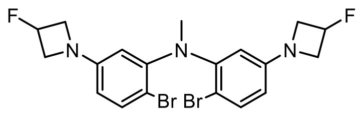

**S15**

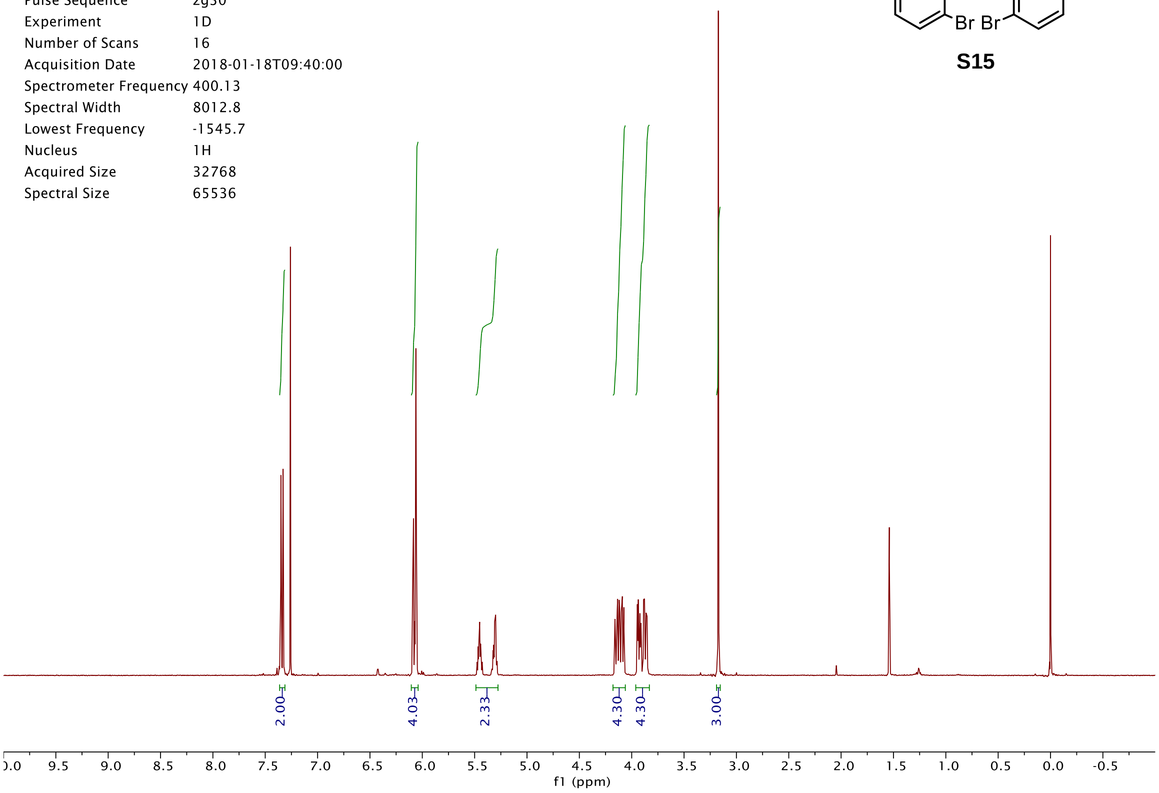
Origin Bruker BioSpin GmbH
Solvent CDCl3
Temperature 295.2
Pulse Sequence zg30
Experiment 1D
Number of Scans 16
Acquisition Date 2018-01-18T09:40:00
Spectrometer Frequency 400.13
Spectral Width 8012.8
Lowest Frequency -1545.7
Nucleus 1H
Acquired Size 32768
Spectral Size 65536
2.00
4.03
2.33
4.30
4.30
3.00
f1 (ppm)

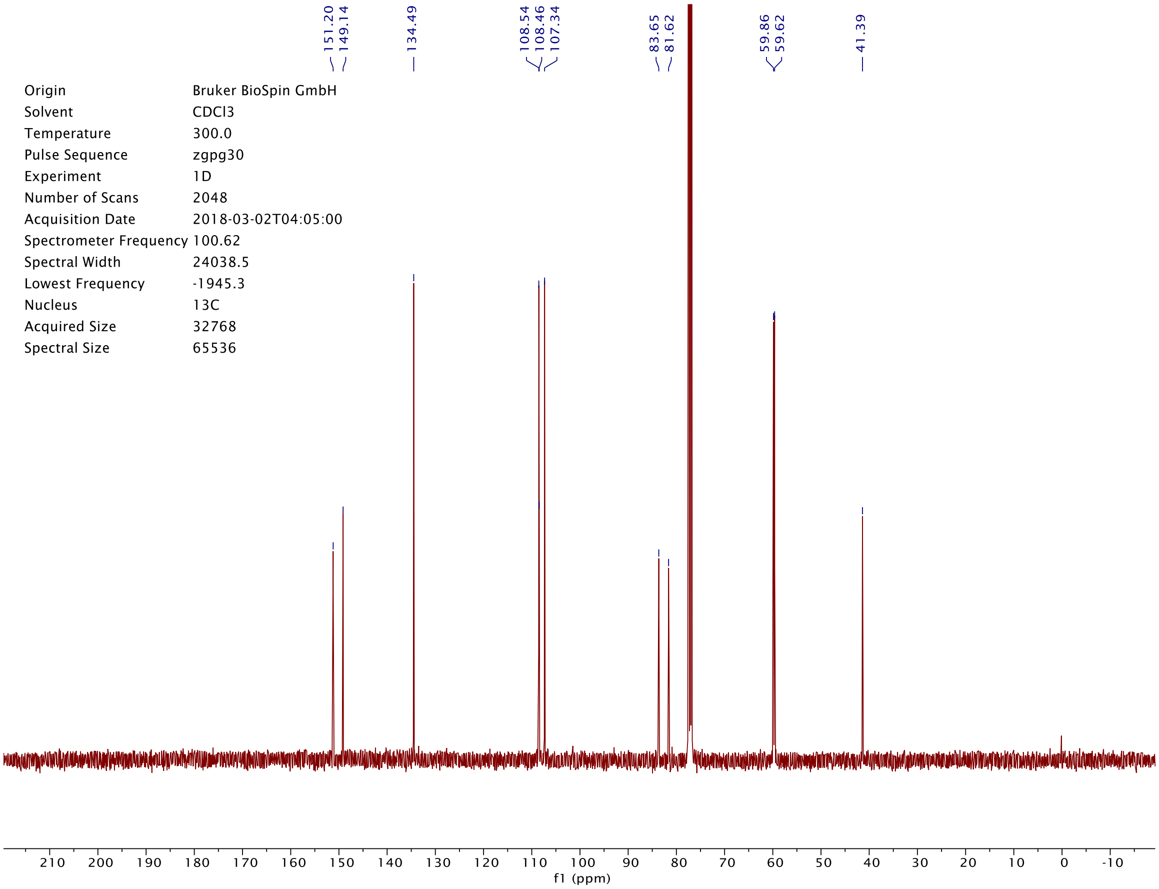
151.20
149.14
134.49
108.54
108.46
107.34
83.65
81.62
59.86
59.62
41.39
Origin Bruker BioSpin GmbH
Solvent CDCl3
Temperature 300.0
Pulse Sequence zgpg30
Experiment 1D
Number of Scans 2048
Acquisition Date 2018-03-02T04:05:00
Spectrometer Frequency 100.62
Spectral Width 24038.5
Lowest Frequency -1945.3
Nucleus 13C
Acquired Size 32768
Spectral Size 65536
f1 (ppm)

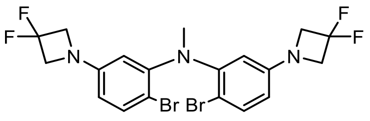
F
F
N
N
N
F
F
Br Br

**S16**

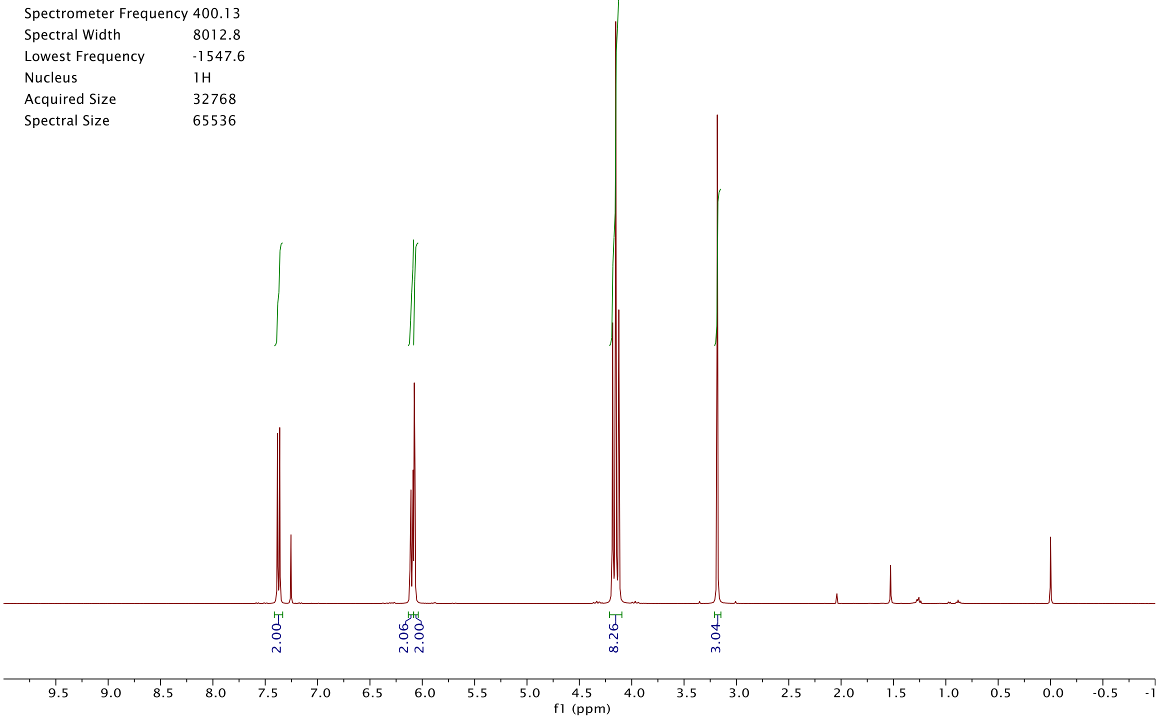
Origin Bruker BioSpin GmbH
Solvent CDCl3
Temperature 300.0
Pulse Sequence zg30
Experiment 1D
Number of Scans 16
Acquisition Date 2018-03-26T12:02:00
Spectrometer Frequency 400.13
Spectral Width 8012.8
Lowest Frequency -1547.6
Nucleus 1H
Acquired Size 32768
Spectral Size 65536
2.00
2.06
2.00
8.26
3.04
9.5 9.0 8.5 8.0 7.5 7.0 6.5 6.0 5.5 5.0 4.5 4.0 3.5 3.0 2.5 2.0 1.5 1.0 0.5 0.0 -0.5 -1
f1 (ppm)

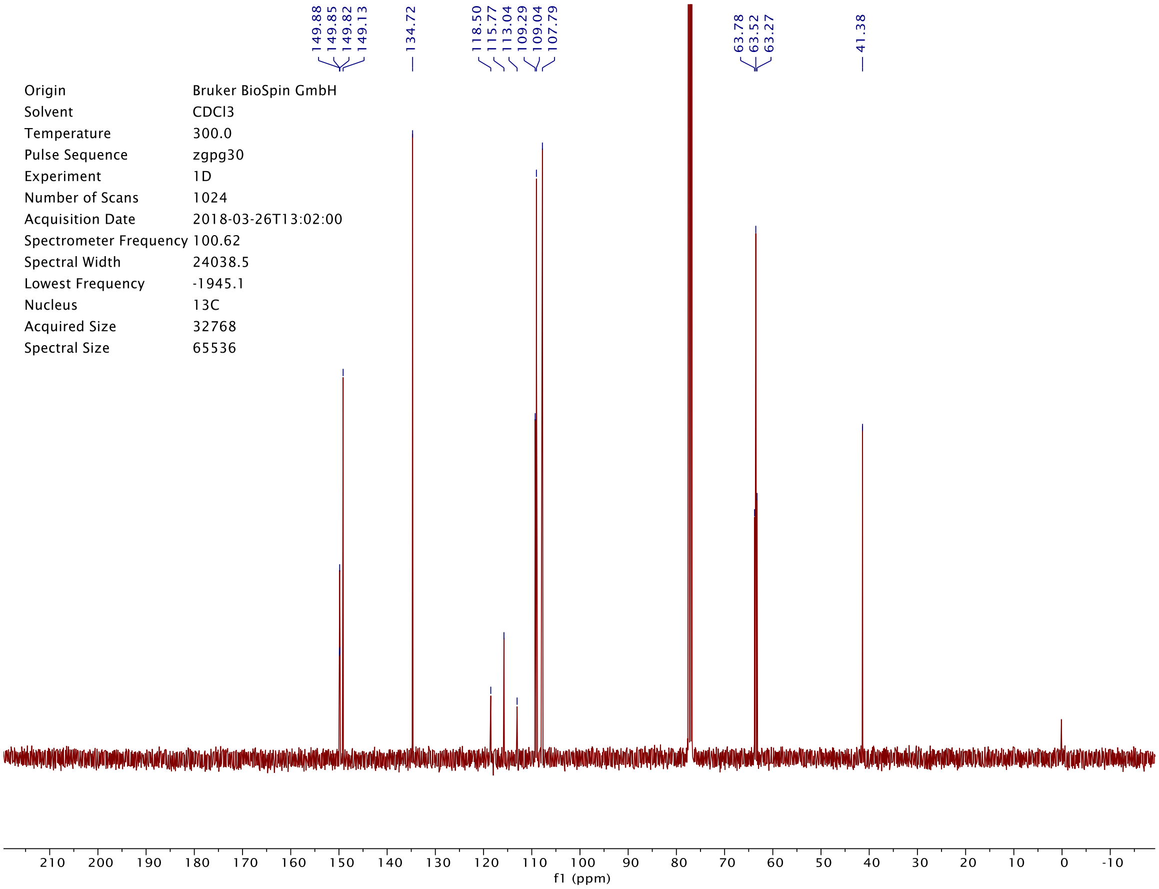
149.88
149.85
149.82
149.13
134.72
118.50
115.77
113.04
109.29
109.04
107.79
63.78
63.52
63.27
41.38
Origin Bruker BioSpin GmbH
Solvent CDCl3
Temperature 300.0
Pulse Sequence zgpg30
Experiment 1D
Number of Scans 1024
Acquisition Date 2018-03-26T13:02:00
Spectrometer Frequency 100.62
Spectral Width 24038.5
Lowest Frequency -1945.1
Nucleus 13C
Acquired Size 32768
Spectral Size 65536
210 200 190 180 170 160 150 140 130 120 110 100 90 80 70 60 50 40 30 20 10 0 -10
f1 (ppm)

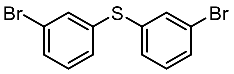


**S27**

| | |
|---|---|
| Origin | Bruker BioSpin GmbH |
| Solvent | CDCl3 |
| Temperature | 300.0 |
| Pulse Sequence | zg30 |
| Experiment | 1D |
| Number of Scans | 16 |
| Acquisition Date | 2018-03-19T19:06:00 |
| Spectrometer Frequency | 400.13 |
| Spectral Width | 8012.8 |
| Lowest Frequency | -1547.3 |
| Nucleus | 1H |
| Acquired Size | 32768 |
| Spectral Size | 65536 |

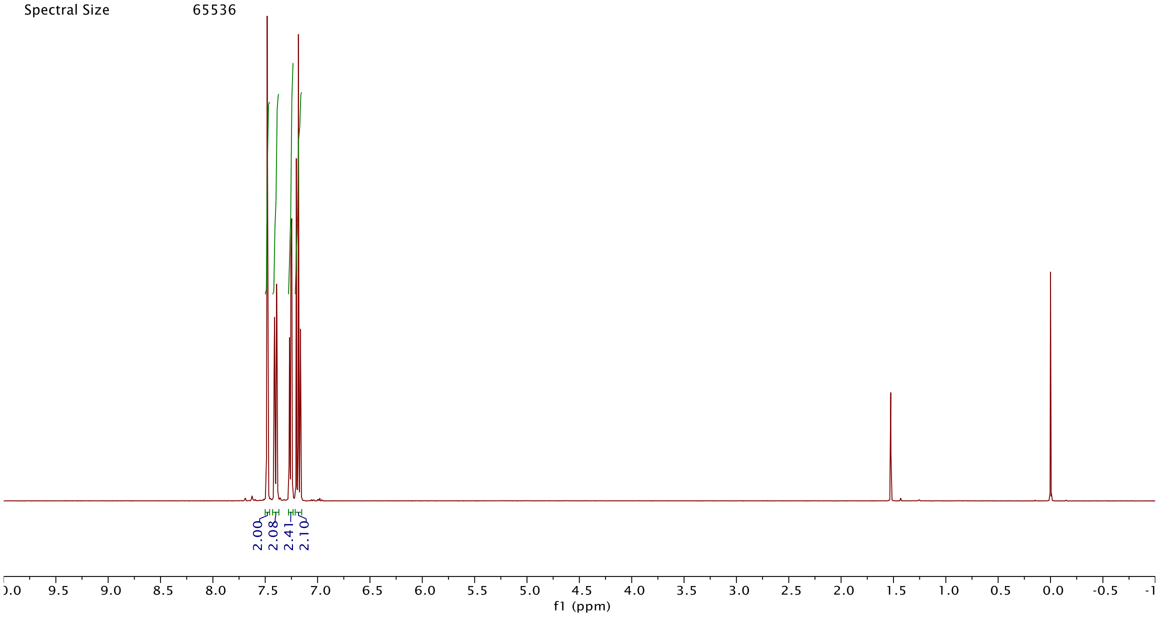


| | |
|---|---|
| Origin | Bruker BioSpin GmbH |
| Solvent | CDCl3 |
| Temperature | 300.0 |
| Pulse Sequence | zgpg30 |
| Experiment | 1D |
| Number of Scans | 2048 |
| Acquisition Date | 2018-03-19T21:05:00 |
| Spectrometer Frequency | 100.62 |
| Spectral Width | 24038.5 |
| Lowest Frequency | -1944.7 |
| Nucleus | 13C |
| Acquired Size | 32768 |
| Spectral Size | 65536 |

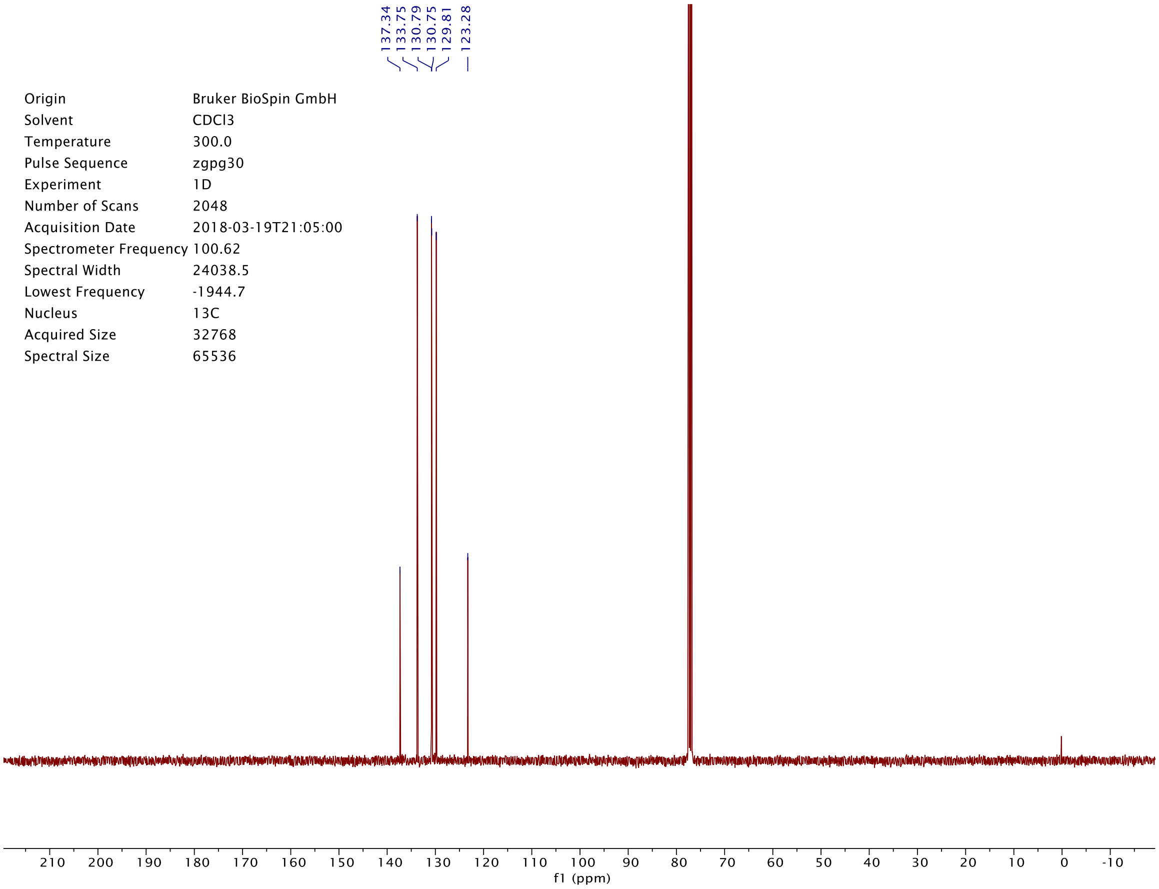

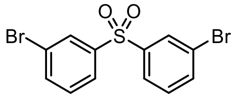

S28

| | |
|---|---|
| Origin | Bruker BioSpin GmbH |
| Solvent | CDCl3 |
| Temperature | 295.6 |
| Pulse Sequence | zg30 |
| Experiment | 1D |
| Number of Scans | 16 |
| Acquisition Date | 2018-03-05T18:05:00 |
| Spectrometer Frequency | 400.13 |
| Spectral Width | 8012.8 |
| Lowest Frequency | -1543.8 |
| Nucleus | 1H |
| Acquired Size | 32768 |
| Spectral Size | 65536 |

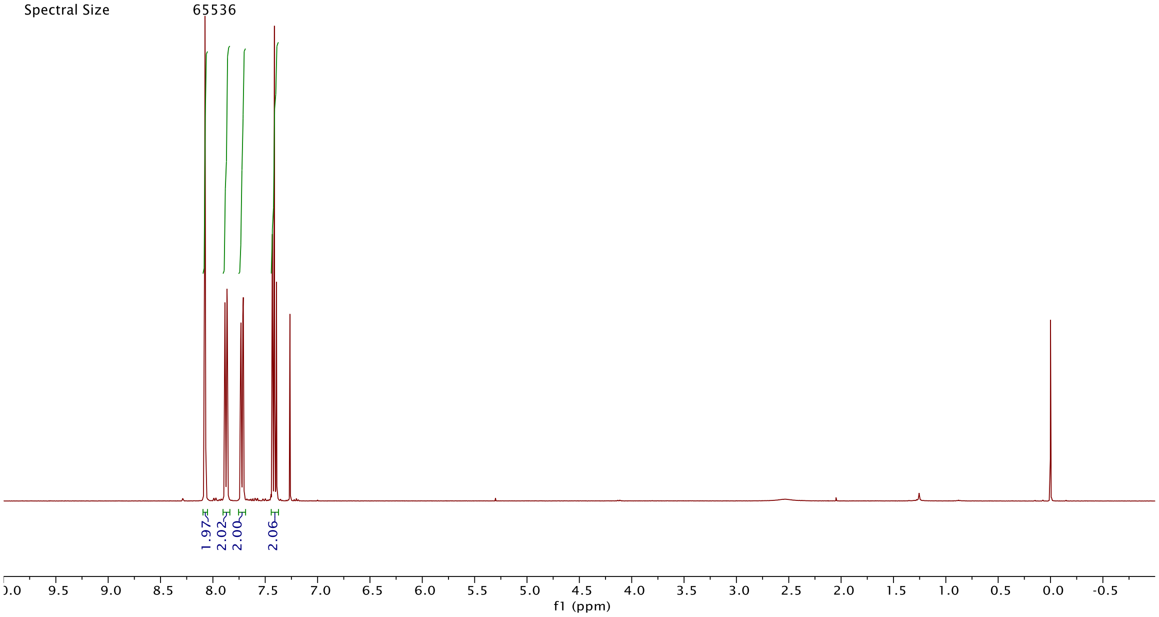


| | |
|---|---|
| Origin | Bruker BioSpin GmbH |
| Solvent | CDCl3 |
| Temperature | 296.3 |
| Pulse Sequence | zgpg30 |
| Experiment | 1D |
| Number of Scans | 2048 |
| Acquisition Date | 2018-03-05T20:04:00 |
| Spectrometer Frequency | 100.62 |
| Spectral Width | 24038.5 |
| Lowest Frequency | -1947.0 |
| Nucleus | 13C |
| Acquired Size | 32768 |
| Spectral Size | 65536 |

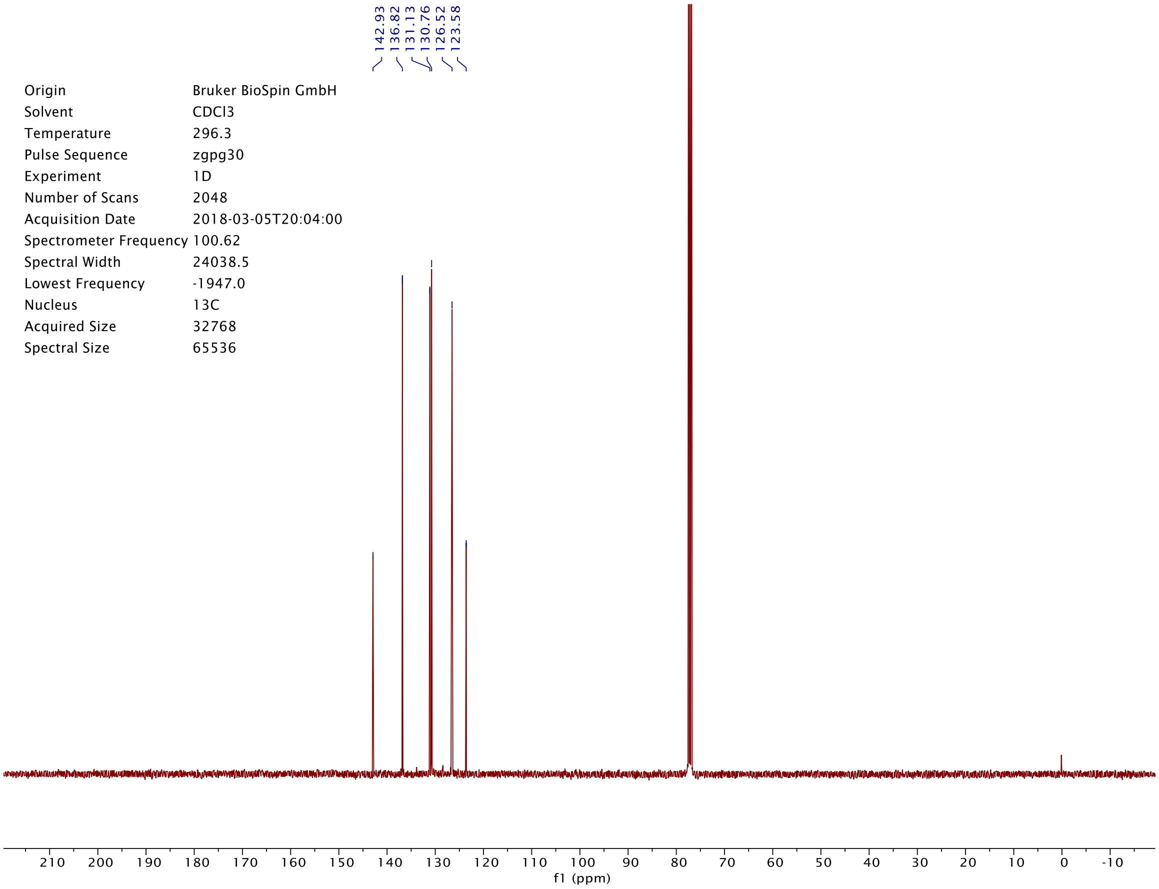

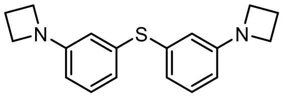


**S29**

| Origin | Bruker BioSpin GmbH |
| --- | --- |
| Solvent | CDCl3 |
| Temperature | 300.0 |
| Pulse Sequence | zg30 |
| Experiment | 1D |
| Number of Scans | 16 |
| Acquisition Date | 2017-08-30T09:56:00 |
| Spectrometer Frequency | 400.13 |
| Spectral Width | 8012.8 |
| Lowest Frequency | -1547.1 |
| Nucleus | 1H |
| Acquired Size | 32768 |
| Spectral Size | 65536 |

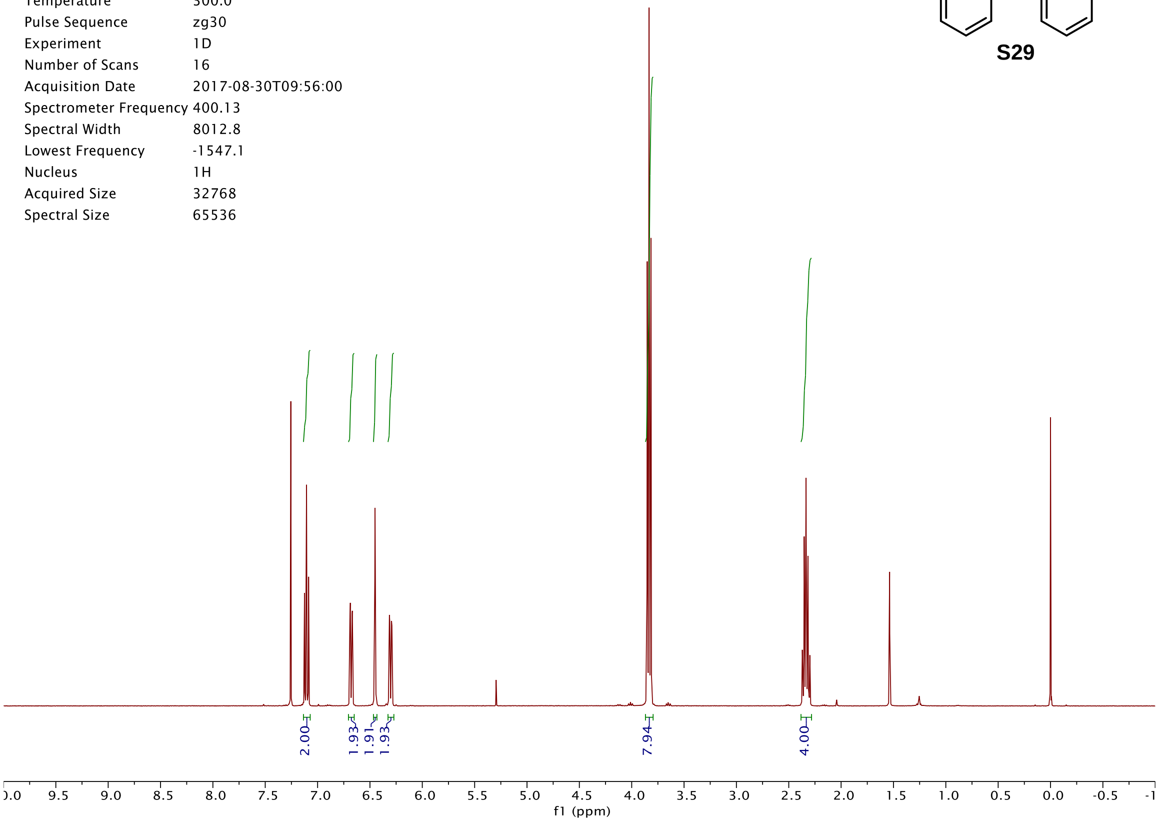


| Origin | Bruker BioSpin GmbH |
| --- | --- |
| Solvent | CDCl3 |
| Temperature | 300.0 |
| Pulse Sequence | zgpg30 |
| Experiment | 1D |
| Number of Scans | 512 |
| Acquisition Date | 2018-03-19T22:17:00 |
| Spectrometer Frequency | 100.62 |
| Spectral Width | 24038.5 |
| Lowest Frequency | -1947.3 |
| Nucleus | 13C |
| Acquired Size | 32768 |
| Spectral Size | 65536 |

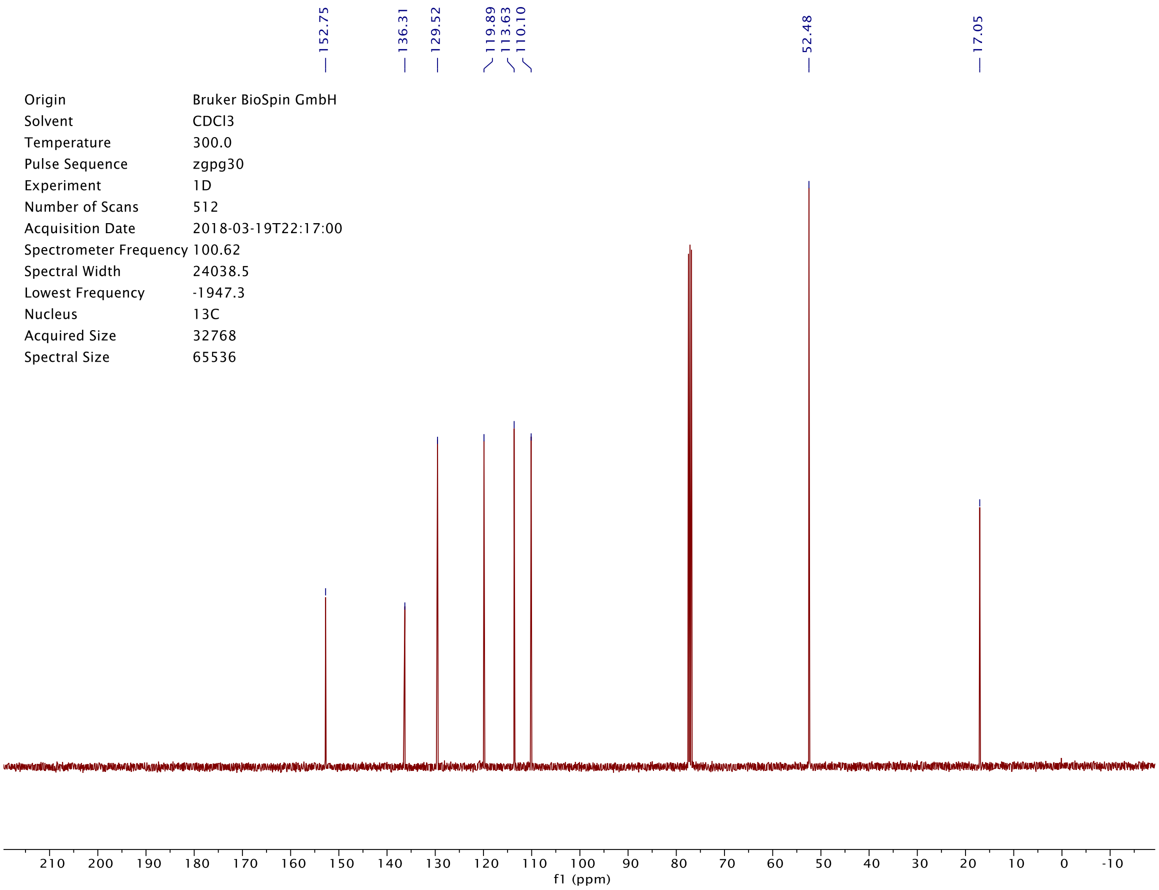

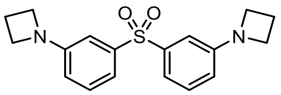

**S30**

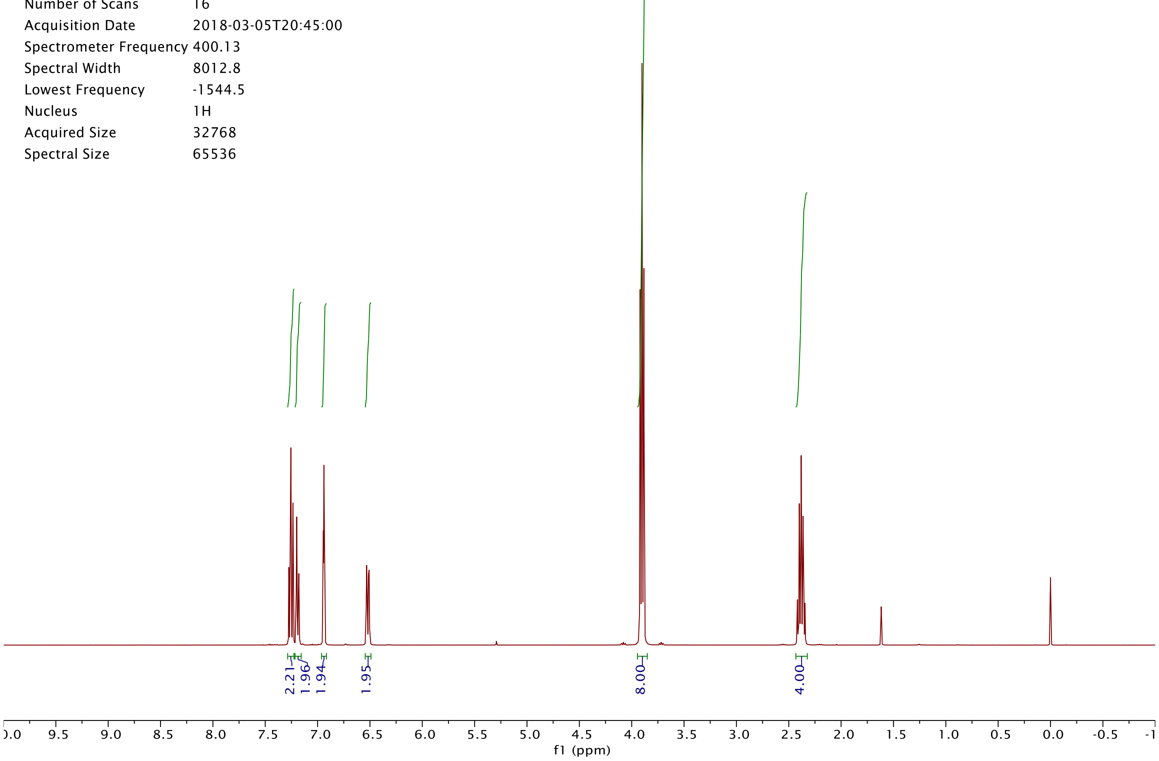

Origin Bruker BioSpin GmbH
Solvent CDCl3
Temperature 295.6
Pulse Sequence zg30
Experiment 1D
Number of Scans 16
Acquisition Date 2018-03-05T20:45:00
Spectrometer Frequency 400.13
Spectral Width 8012.8
Lowest Frequency -1544.5
Nucleus 1H
Acquired Size 32768
Spectral Size 65536
2.21
1.96
1.94
1.95
8.00
4.00
f1 (ppm)


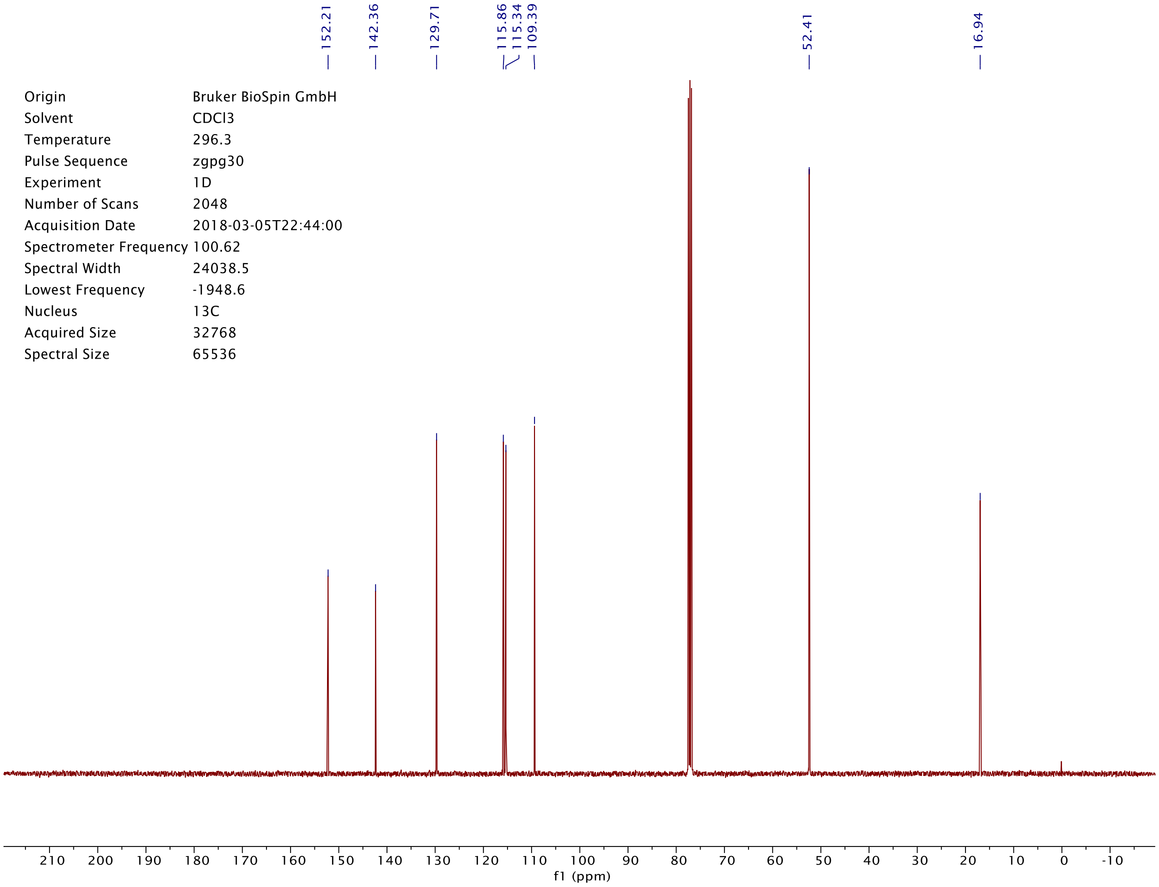

152.21
142.36
129.71
115.86
115.34
109.39
52.41
16.94
Origin Bruker BioSpin GmbH
Solvent CDCl3
Temperature 296.3
Pulse Sequence zgpg30
Experiment 1D
Number of Scans 2048
Acquisition Date 2018-03-05T22:44:00
Spectrometer Frequency 100.62
Spectral Width 24038.5
Lowest Frequency -1948.6
Nucleus 13C
Acquired Size 32768
Spectral Size 65536
f1 (ppm)

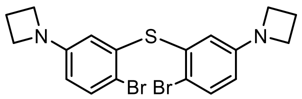


**S31**

| | |
|---|---|
| Origin | Bruker BioSpin GmbH |
| Solvent | CDCl3 |
| Temperature | 300.0 |
| Pulse Sequence | zg30 |
| Experiment | 1D |
| Number of Scans | 16 |
| Acquisition Date | 2017-08-31T15:09:00 |
| Spectrometer Frequency | 400.13 |
| Spectral Width | 8012.8 |
| Lowest Frequency | -1546.2 |
| Nucleus | 1H |
| Acquired Size | 32768 |
| Spectral Size | 65536 |

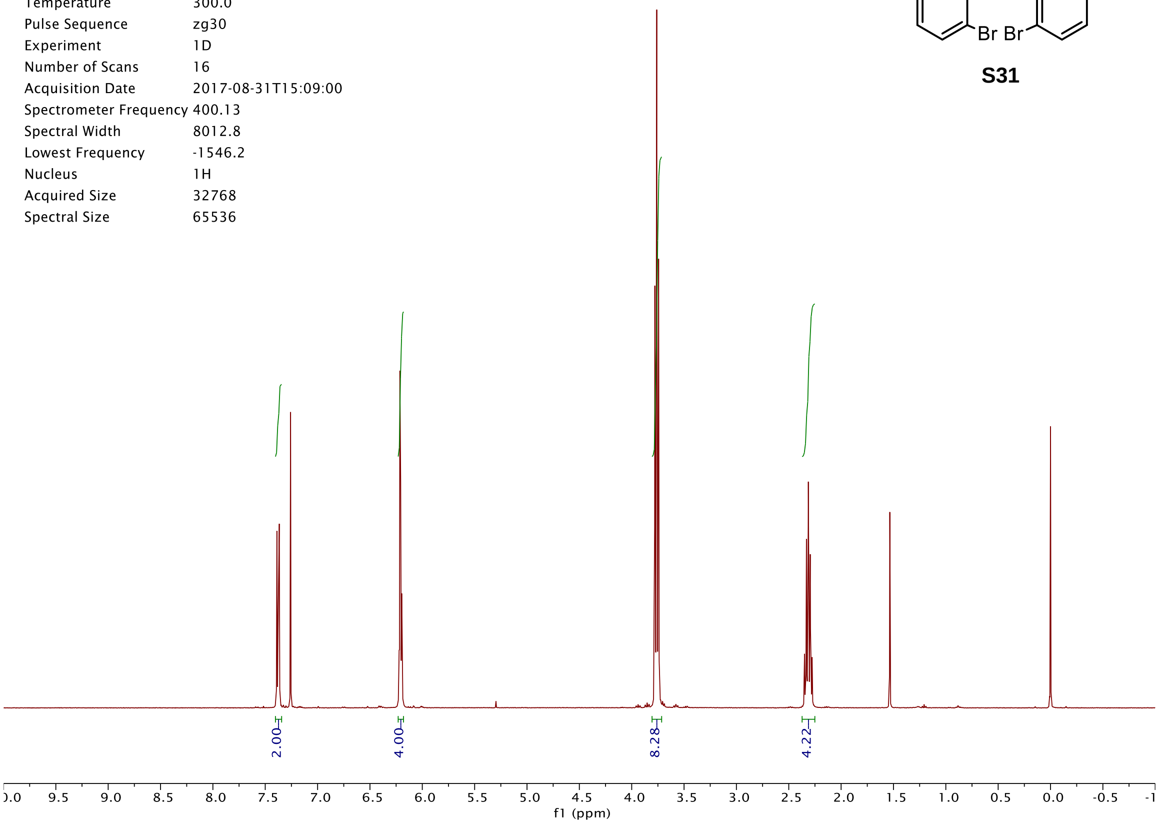


| | |
|---|---|
| Origin | Bruker BioSpin GmbH |
| Solvent | CDCl3 |
| Temperature | 296.3 |
| Pulse Sequence | zgpg30 |
| Experiment | 1D |
| Number of Scans | 2048 |
| Acquisition Date | 2018-03-06T01:24:00 |
| Spectrometer Frequency | 100.62 |
| Spectral Width | 24038.5 |
| Lowest Frequency | -1948.0 |
| Nucleus | 13C |
| Acquired Size | 32768 |
| Spectral Size | 65536 |

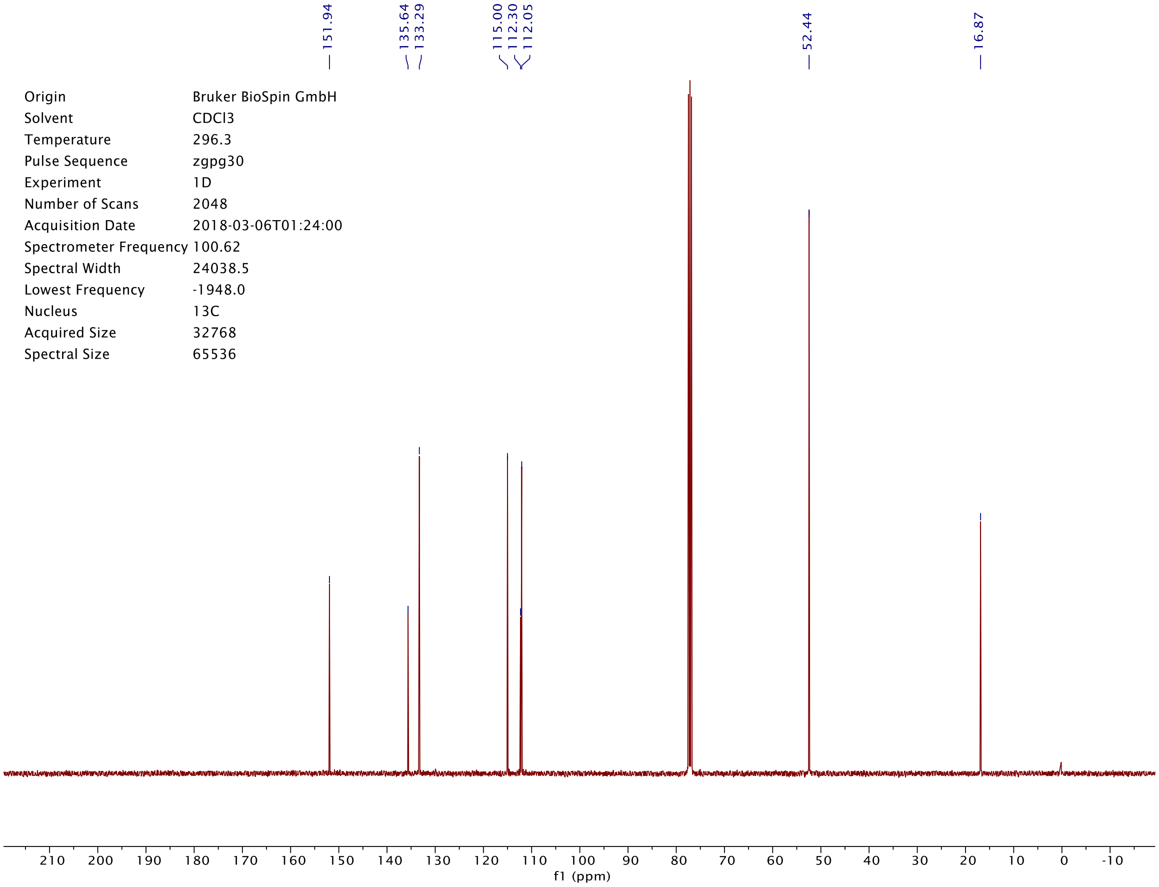

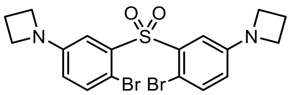
O O
S
N
N
Br Br

**S32**

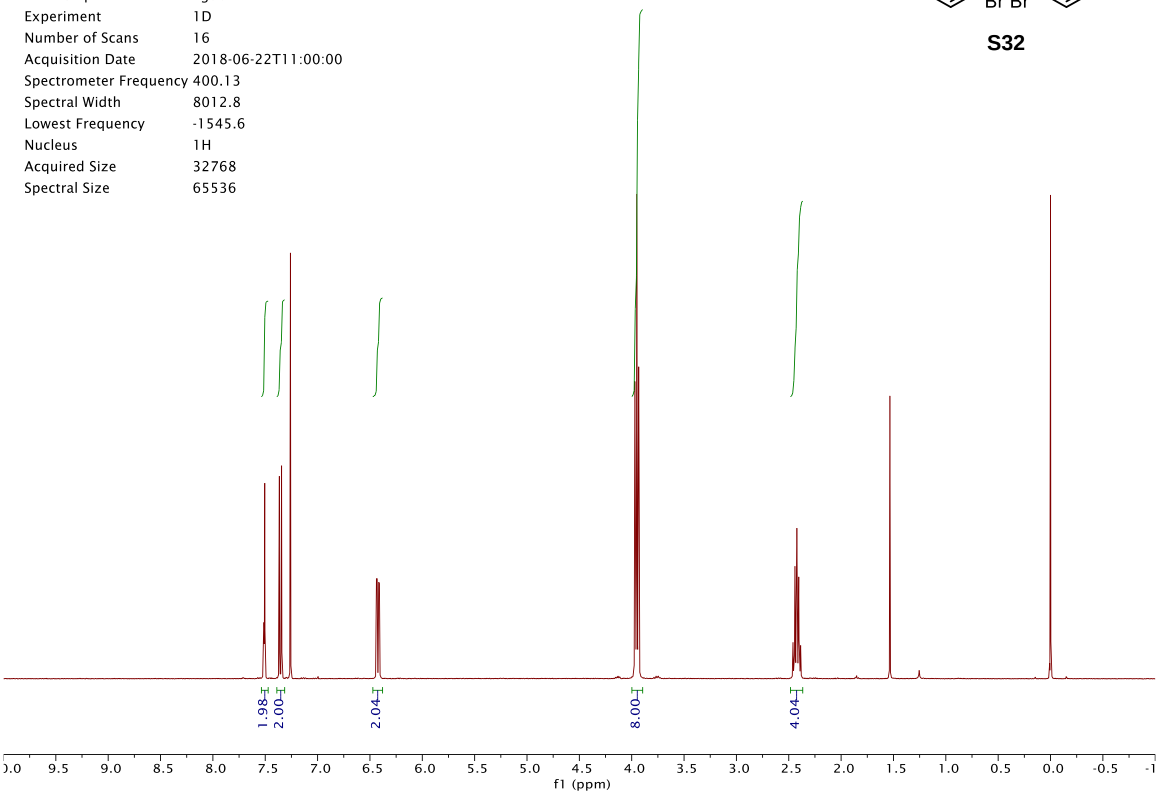
Origin Bruker BioSpin GmbH
Solvent CDCl3
Temperature 300.0
Pulse Sequence zg30
Experiment 1D
Number of Scans 16
Acquisition Date 2018-06-22T11:00:00
Spectrometer Frequency 400.13
Spectral Width 8012.8
Lowest Frequency -1545.6
Nucleus 1H
Acquired Size 32768
Spectral Size 65536
1.98
2.00
2.04
8.00
4.04
f1 (ppm)

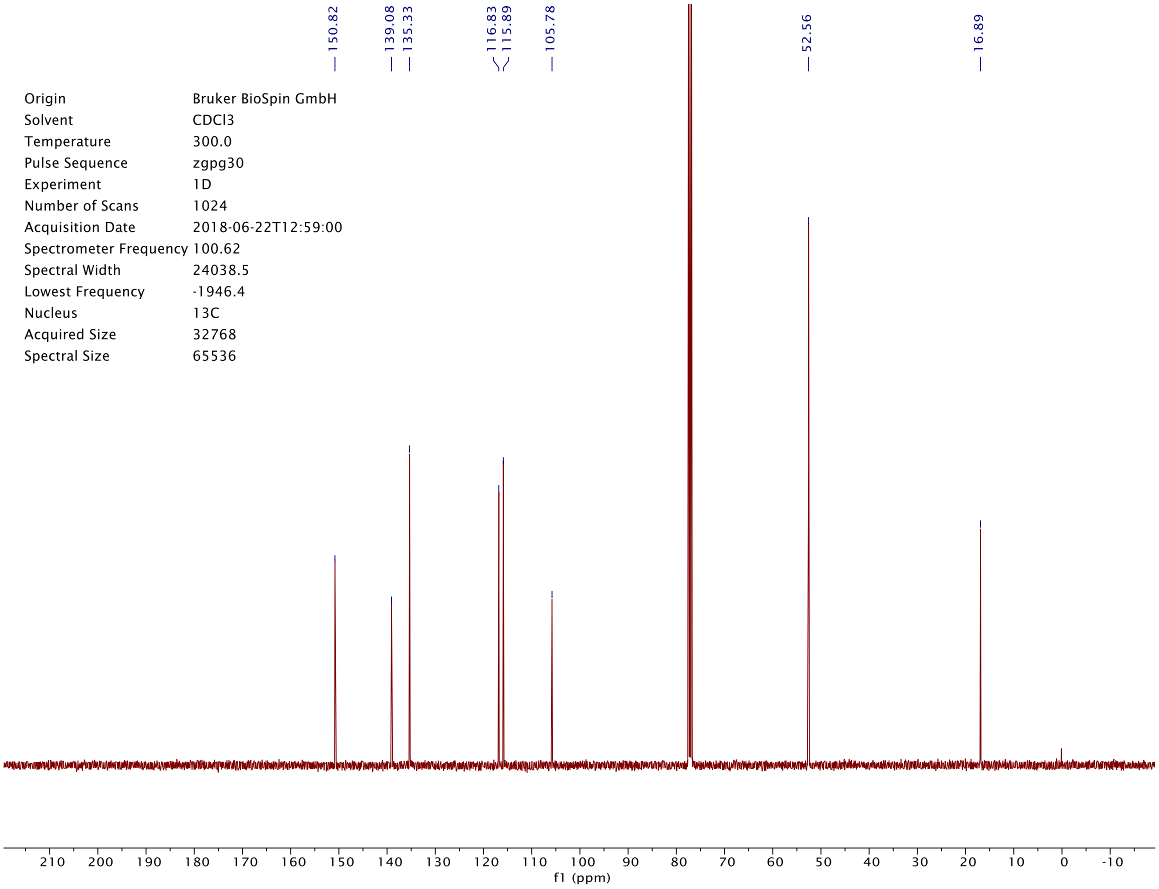
150.82
139.08
135.33
116.83
115.89
105.78
52.56
16.89
Origin Bruker BioSpin GmbH
Solvent CDCl3
Temperature 300.0
Pulse Sequence zgpg30
Experiment 1D
Number of Scans 1024
Acquisition Date 2018-06-22T12:59:00
Spectrometer Frequency 100.62
Spectral Width 24038.5
Lowest Frequency -1946.4
Nucleus 13C
Acquired Size 32768
Spectral Size 65536
f1 (ppm)

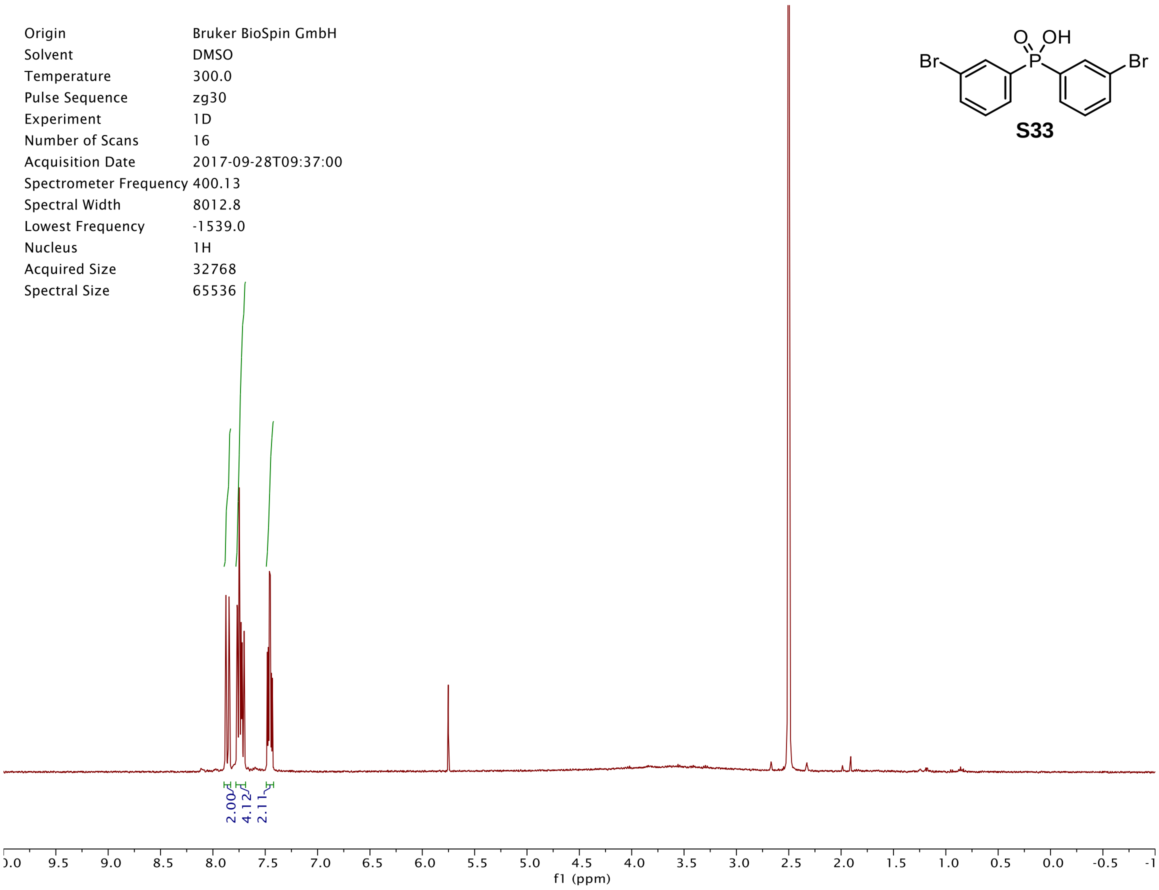
O
OH
Br
P
Br
S33
Origin Bruker BioSpin GmbH
Solvent DMSO
Temperature 300.0
Pulse Sequence zg30
Experiment 1D
Number of Scans 16
Acquisition Date 2017-09-28T09:37:00
Spectrometer Frequency 400.13
Spectral Width 8012.8
Lowest Frequency -1539.0
Nucleus 1H
Acquired Size 32768
Spectral Size 65536
2.00
4.12
2.11
9.5
9.0
8.5
8.0
7.5
7.0
6.5
6.0
5.5
5.0
4.5
4.0
3.5
3.0
2.5
2.0
1.5
1.0
0.5
0.0
-0.5
f1 (ppm)

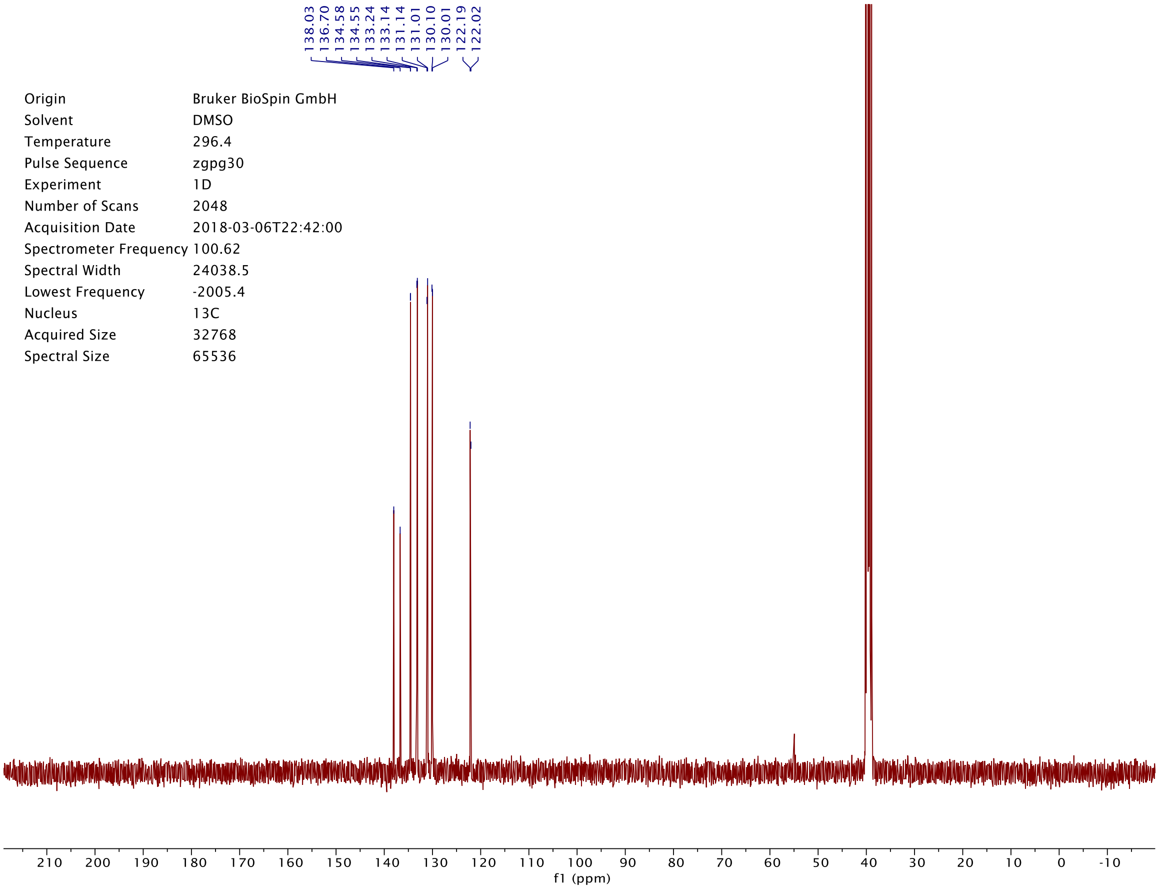
138.03
136.70
134.58
134.55
133.24
133.14
131.14
131.01
130.10
130.01
122.19
122.02
Origin Bruker BioSpin GmbH
Solvent DMSO
Temperature 296.4
Pulse Sequence zgpg30
Experiment 1D
Number of Scans 2048
Acquisition Date 2018-03-06T22:42:00
Spectrometer Frequency 100.62
Spectral Width 24038.5
Lowest Frequency -2005.4
Nucleus 13C
Acquired Size 32768
Spectral Size 65536
210
200
190
180
170
160
150
140
130
120
110
100
90
80
70
60
50
40
30
20
10
0
-10
f1 (ppm)

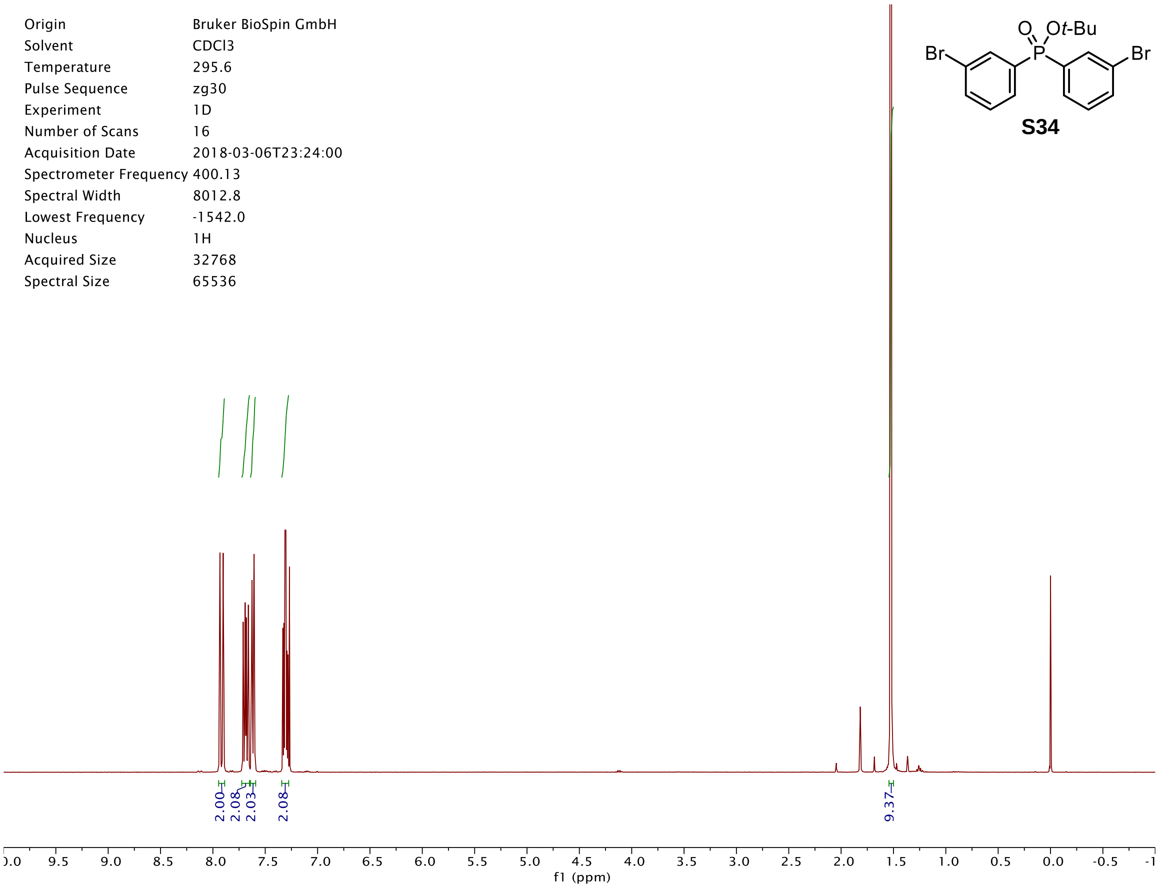
Origin Bruker BioSpin GmbH
Solvent CDCl3
Temperature 295.6
Pulse Sequence zg30
Experiment 1D
Number of Scans 16
Acquisition Date 2018-03-06T23:24:00
Spectrometer Frequency 400.13
Spectral Width 8012.8
Lowest Frequency -1542.0
Nucleus 1H
Acquired Size 32768
Spectral Size 65536
2.00
2.08
2.03
2.08
9.37
f1 (ppm)
O
O*t*-Bu
P
Br
Br

**S34**

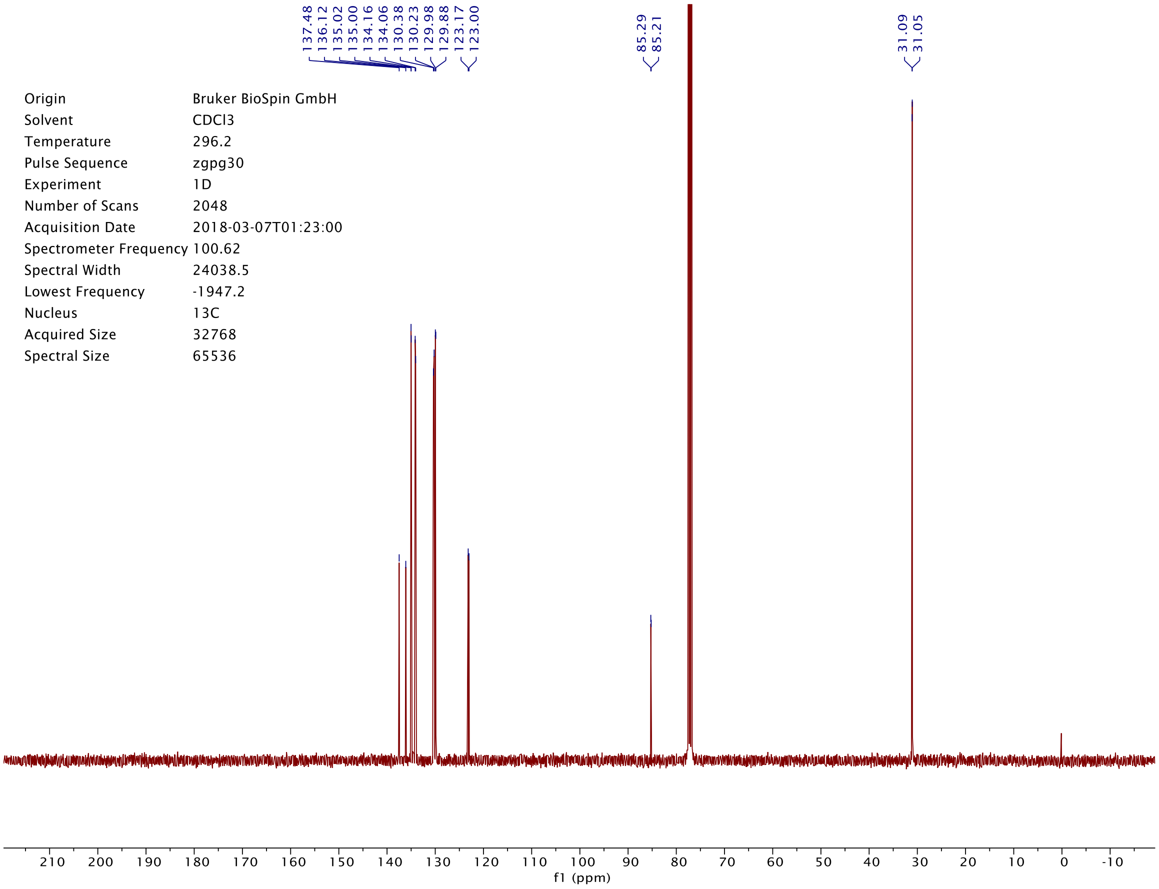
137.48
136.12
135.02
135.00
134.16
134.06
130.38
130.23
129.98
129.88
123.17
123.00
85.29
85.21
31.09
31.05
Origin Bruker BioSpin GmbH
Solvent CDCl3
Temperature 296.2
Pulse Sequence zgpg30
Experiment 1D
Number of Scans 2048
Acquisition Date 2018-03-07T01:23:00
Spectrometer Frequency 100.62
Spectral Width 24038.5
Lowest Frequency -1947.2
Nucleus 13C
Acquired Size 32768
Spectral Size 65536
f1 (ppm)

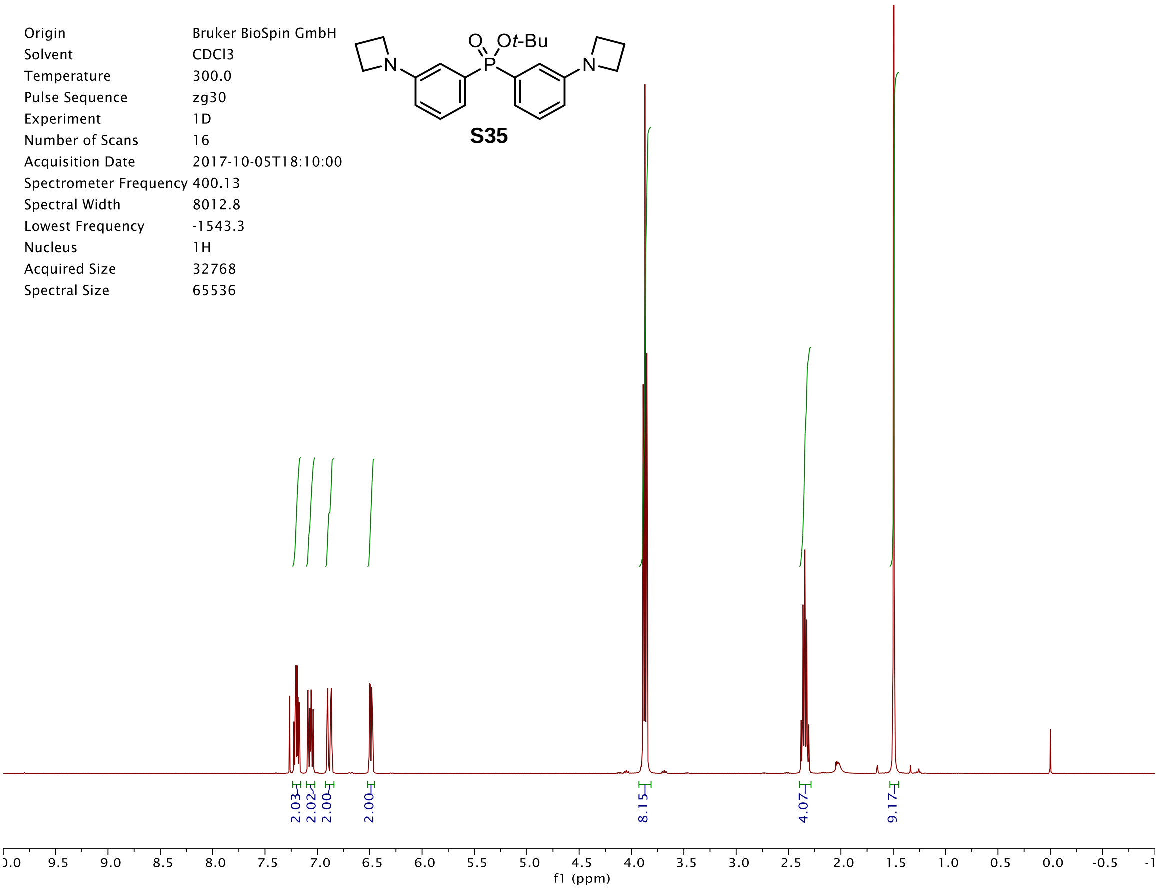
Origin Bruker BioSpin GmbH
Solvent CDCl3
Temperature 300.0
Pulse Sequence zg30
Experiment 1D
Number of Scans 16
Acquisition Date 2017-10-05T18:10:00
Spectrometer Frequency 400.13
Spectral Width 8012.8
Lowest Frequency -1543.3
Nucleus 1H
Acquired Size 32768
Spectral Size 65536
O
Ot-Bu
P
N
N
S35
2.03
2.02
2.00
2.00
8.15
4.07
9.17
f1 (ppm)

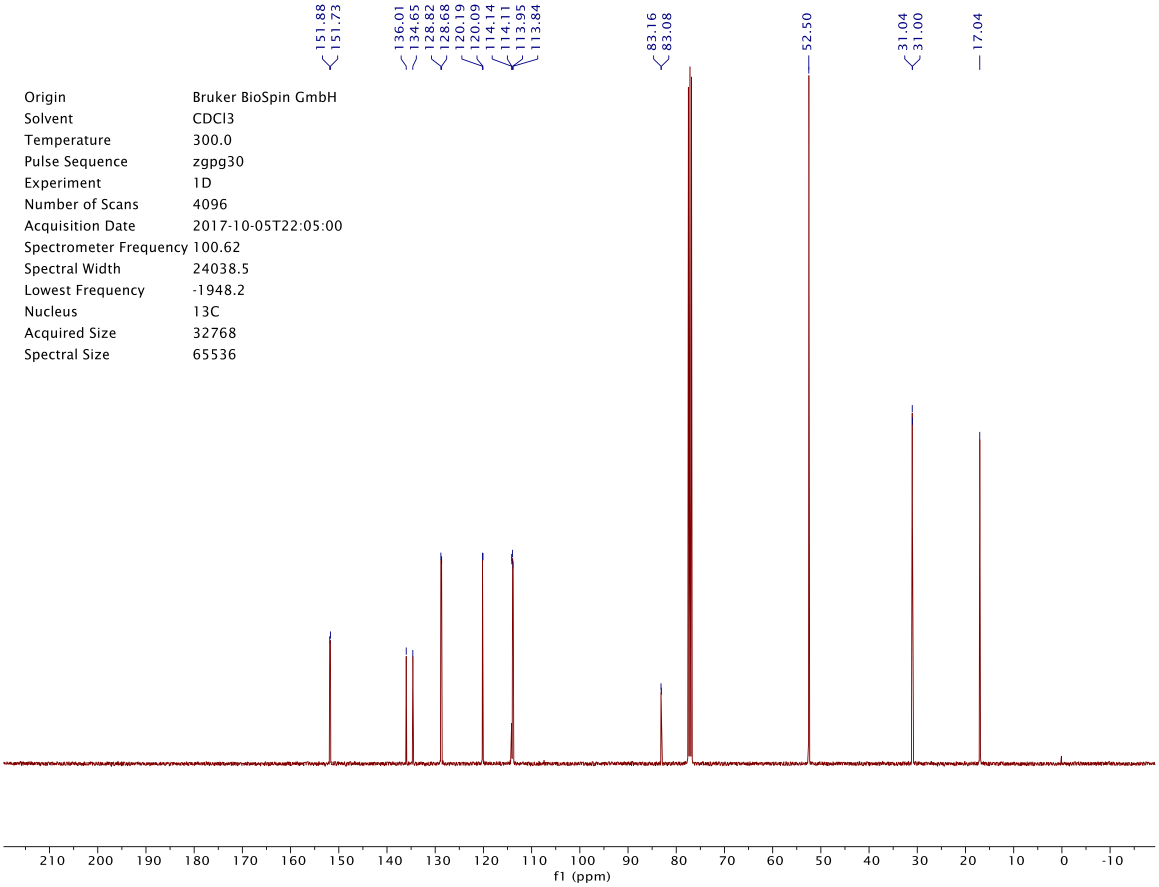
151.88
151.73
136.01
134.65
128.82
128.68
120.19
120.09
114.14
114.11
113.95
113.84
83.16
83.08
52.50
31.04
31.00
17.04
Origin Bruker BioSpin GmbH
Solvent CDCl3
Temperature 300.0
Pulse Sequence zgpg30
Experiment 1D
Number of Scans 4096
Acquisition Date 2017-10-05T22:05:00
Spectrometer Frequency 100.62
Spectral Width 24038.5
Lowest Frequency -1948.2
Nucleus 13C
Acquired Size 32768
Spectral Size 65536
f1 (ppm)

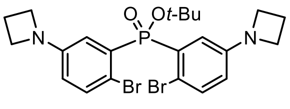
O
Ot-Bu
P
N
N
Br Br

**S36**

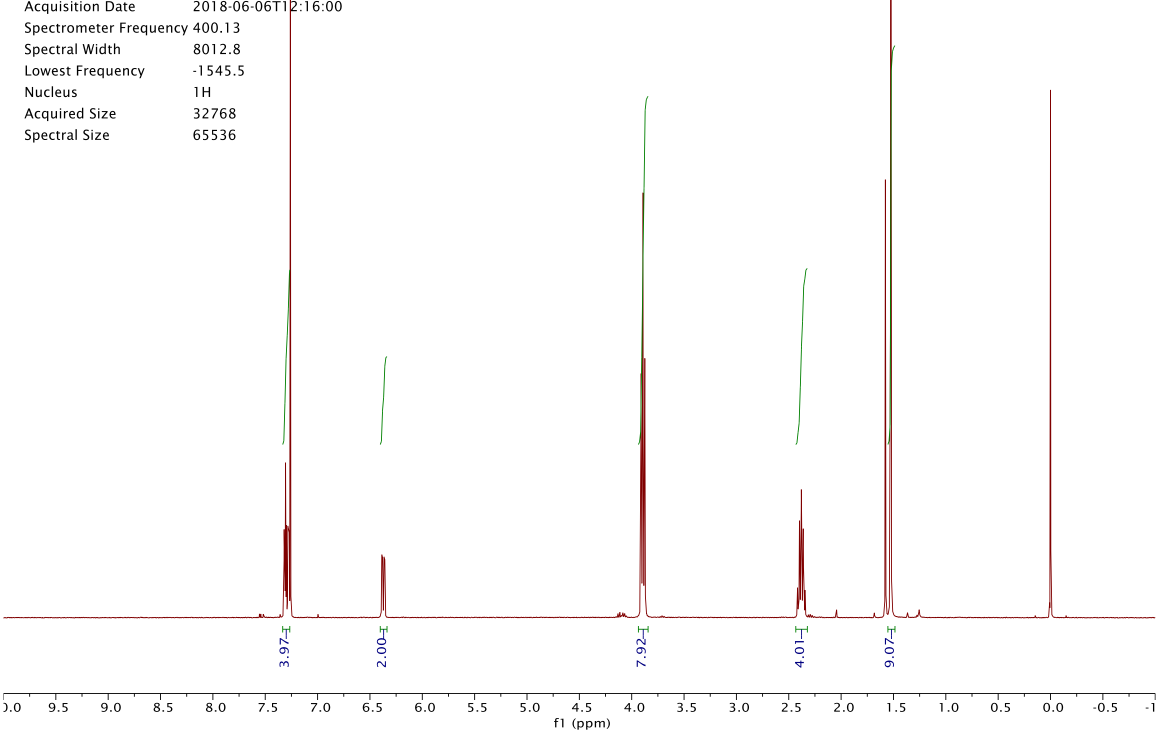
Origin Bruker BioSpin GmbH
Solvent CDCl3
Temperature 300.0
Pulse Sequence zg30
Experiment 1D
Number of Scans 16
Acquisition Date 2018-06-06T12:16:00
Spectrometer Frequency 400.13
Spectral Width 8012.8
Lowest Frequency -1545.5
Nucleus 1H
Acquired Size 32768
Spectral Size 65536
3.97
2.00
7.92
4.01
9.07
f1 (ppm)

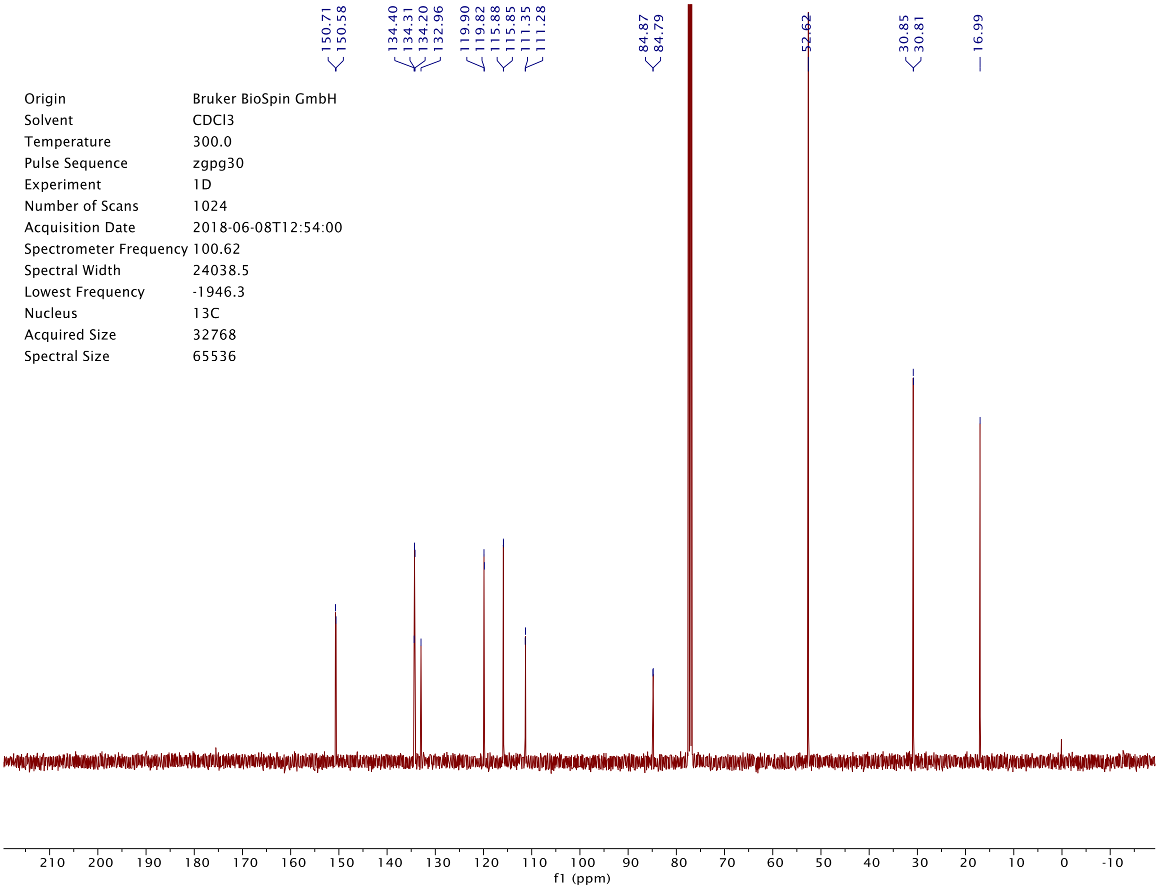
150.71
150.58
134.40
134.31
134.20
132.96
119.90
119.82
115.88
115.85
111.35
111.28
84.87
84.79
52.62
30.85
30.81
16.99
Origin Bruker BioSpin GmbH
Solvent CDCl3
Temperature 300.0
Pulse Sequence zgpg30
Experiment 1D
Number of Scans 1024
Acquisition Date 2018-06-08T12:54:00
Spectrometer Frequency 100.62
Spectral Width 24038.5
Lowest Frequency -1946.3
Nucleus 13C
Acquired Size 32768
Spectral Size 65536
f1 (ppm)

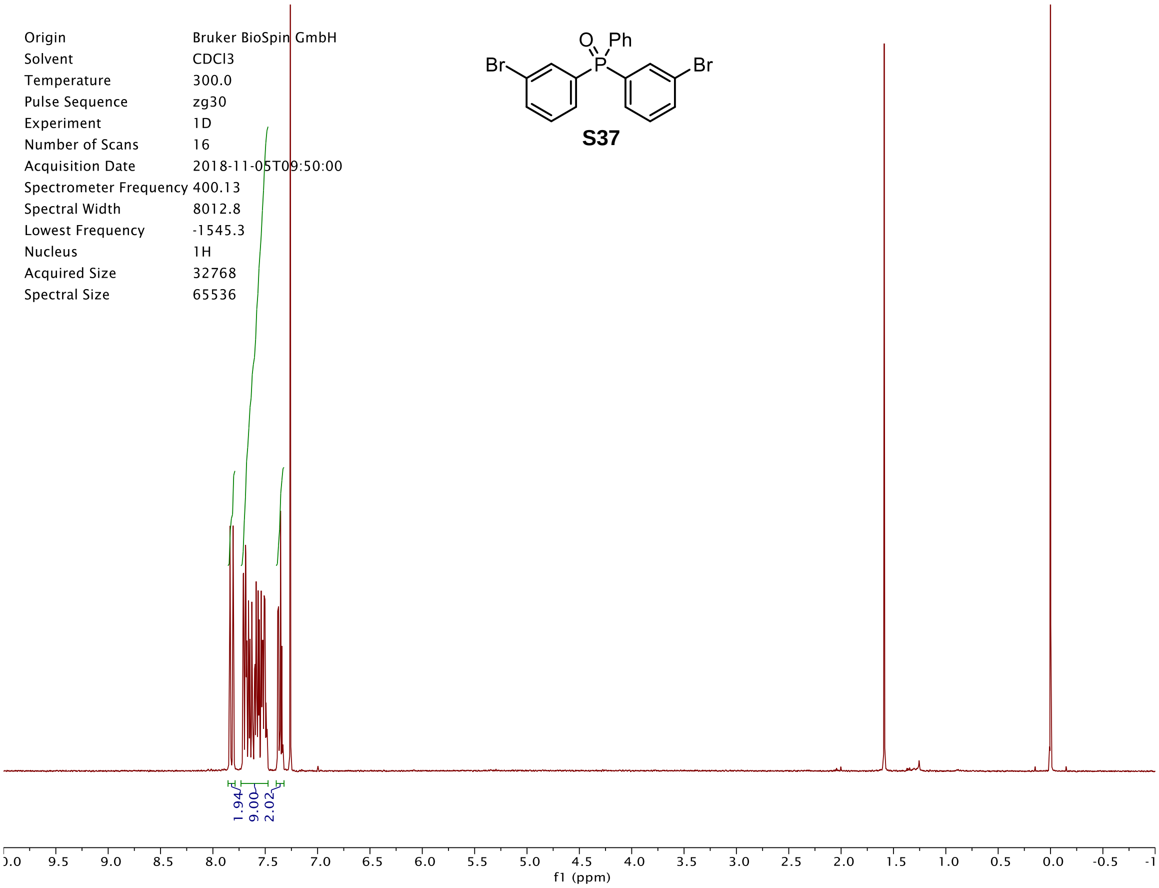
Origin Bruker BioSpin GmbH
Solvent CDCl3
Temperature 300.0
Pulse Sequence zg30
Experiment 1D
Number of Scans 16
Acquisition Date 2018-11-05T09:50:00
Spectrometer Frequency 400.13
Spectral Width 8012.8
Lowest Frequency -1545.3
Nucleus 1H
Acquired Size 32768
Spectral Size 65536
S37
1.94
9.00
2.02
f1 (ppm)

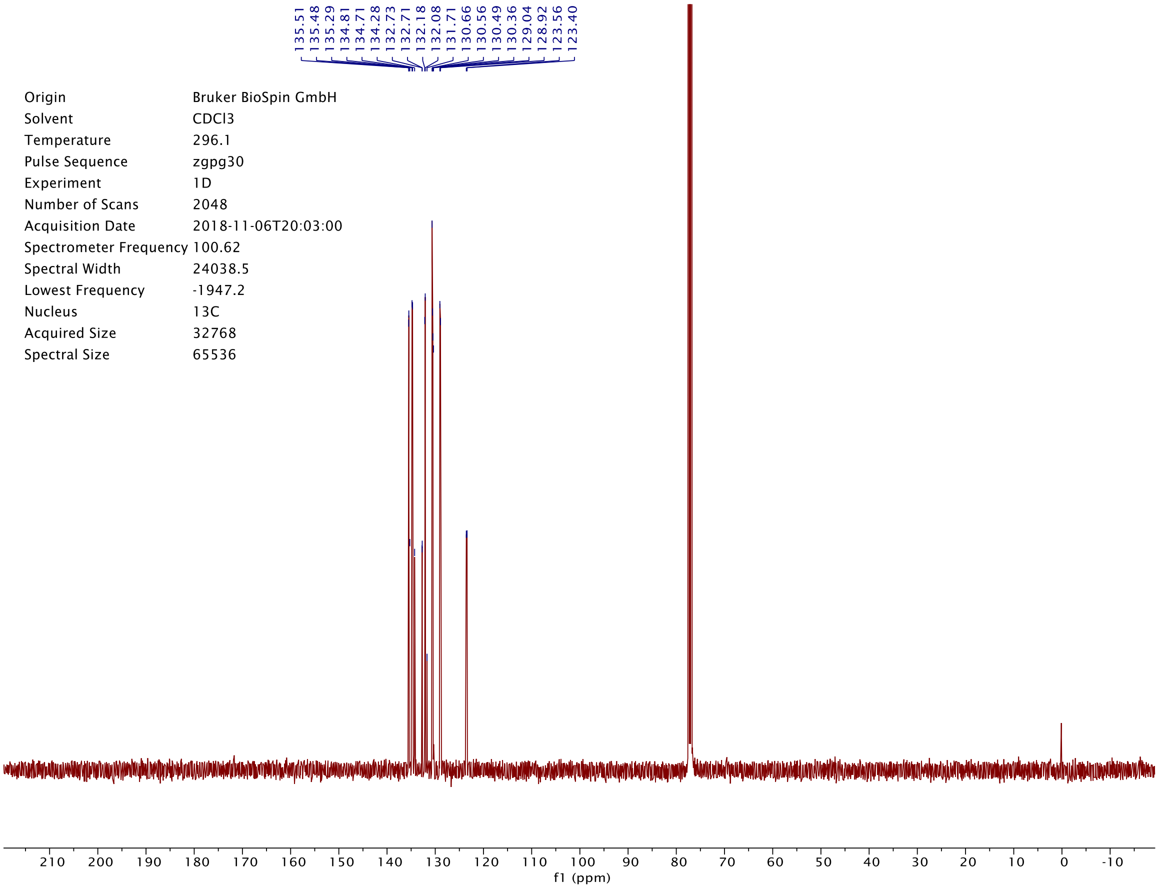
135.51
135.48
135.29
134.81
134.71
134.28
132.73
132.71
132.18
132.08
131.71
130.66
130.56
130.49
130.36
129.04
128.92
123.56
123.40
Origin Bruker BioSpin GmbH
Solvent CDCl3
Temperature 296.1
Pulse Sequence zgpg30
Experiment 1D
Number of Scans 2048
Acquisition Date 2018-11-06T20:03:00
Spectrometer Frequency 100.62
Spectral Width 24038.5
Lowest Frequency -1947.2
Nucleus 13C
Acquired Size 32768
Spectral Size 65536
f1 (ppm)

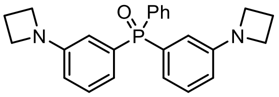

**S38**

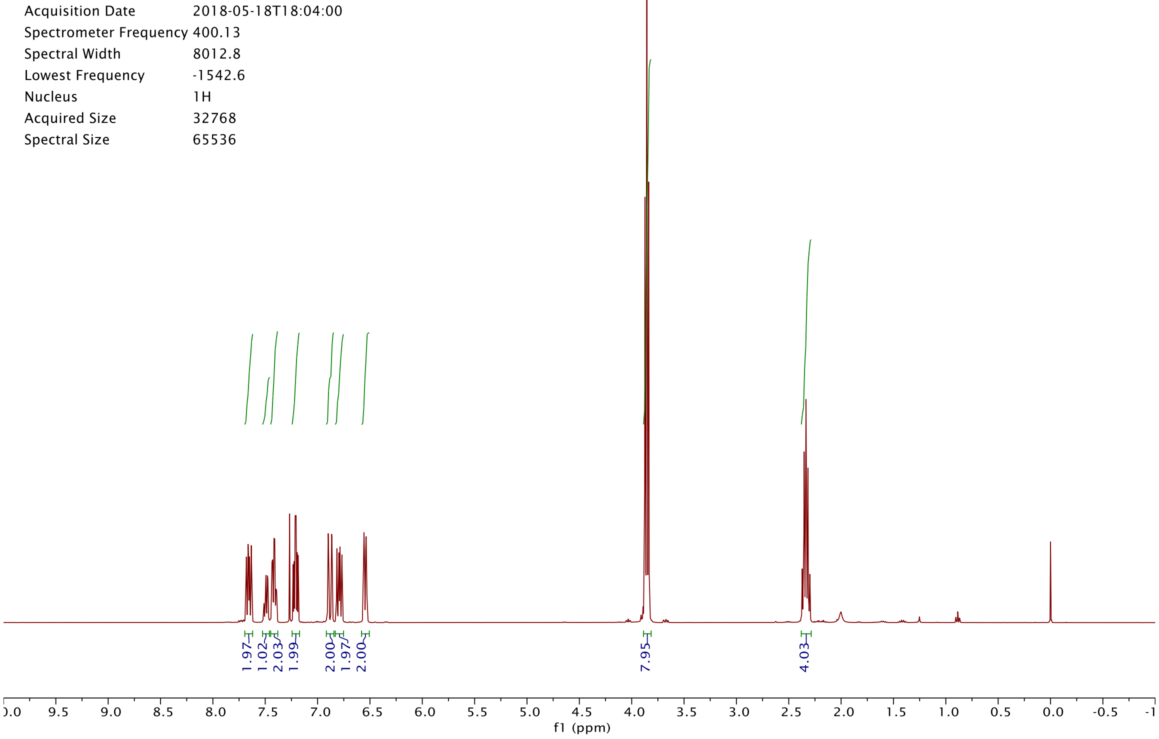

Origin Bruker BioSpin GmbH
Solvent CDCl3
Temperature 300.0
Pulse Sequence zg30
Experiment 1D
Number of Scans 16
Acquisition Date 2018-05-18T18:04:00
Spectrometer Frequency 400.13
Spectral Width 8012.8
Lowest Frequency -1542.6
Nucleus 1H
Acquired Size 32768
Spectral Size 65536
1.97
1.02
2.03
1.99
2.00
1.97
2.00
7.95
4.03
f1 (ppm)


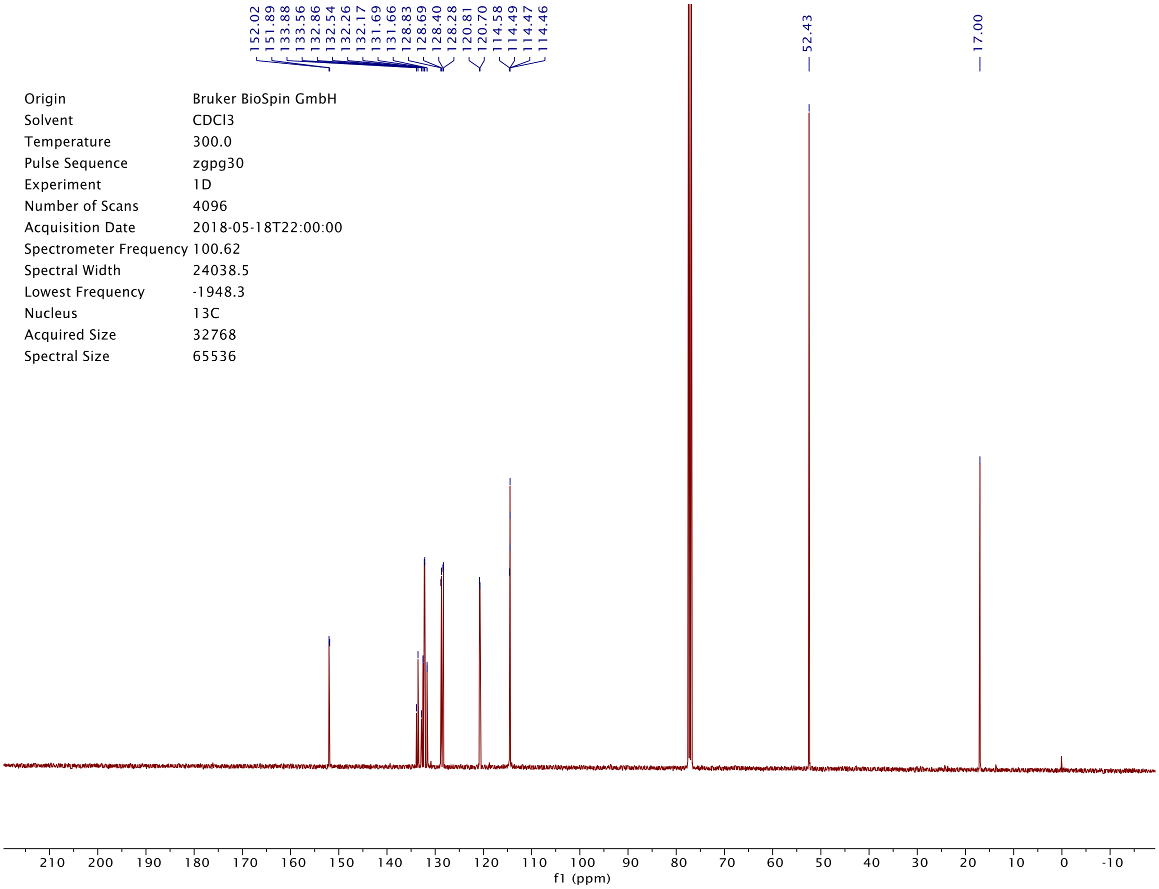

152.02
151.89
133.88
133.56
132.86
132.54
132.26
132.17
131.69
131.66
128.83
128.69
128.40
128.28
120.81
120.70
114.58
114.49
114.47
114.46
52.43
17.00
Origin Bruker BioSpin GmbH
Solvent CDCl3
Temperature 300.0
Pulse Sequence zgpg30
Experiment 1D
Number of Scans 4096
Acquisition Date 2018-05-18T22:00:00
Spectrometer Frequency 100.62
Spectral Width 24038.5
Lowest Frequency -1948.3
Nucleus 13C
Acquired Size 32768
Spectral Size 65536
f1 (ppm)

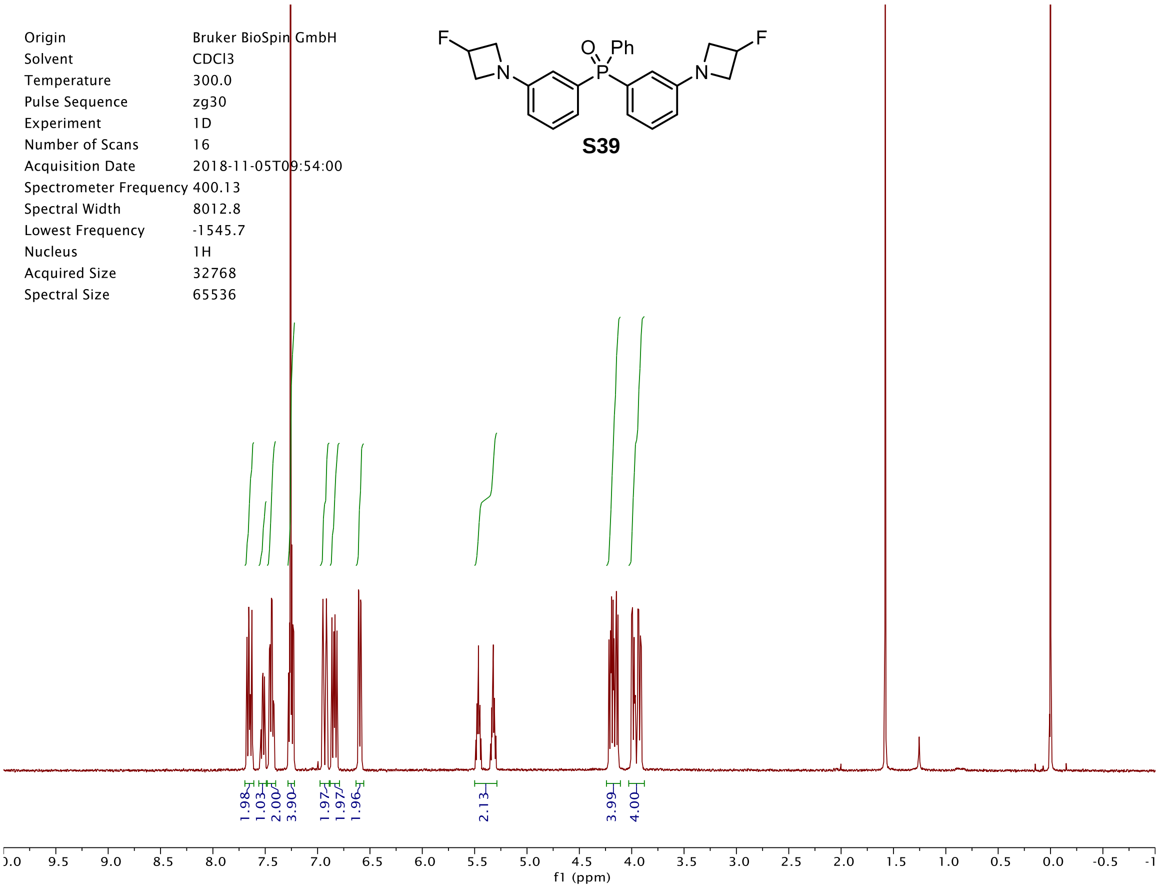
S39
Origin Bruker BioSpin GmbH
Solvent CDCl3
Temperature 300.0
Pulse Sequence zg30
Experiment 1D
Number of Scans 16
Acquisition Date 2018-11-05T09:54:00
Spectrometer Frequency 400.13
Spectral Width 8012.8
Lowest Frequency -1545.7
Nucleus 1H
Acquired Size 32768
Spectral Size 65536
1.98
1.03
2.00
3.90
1.97
1.97
1.96
2.13
3.99
4.00
f1 (ppm)

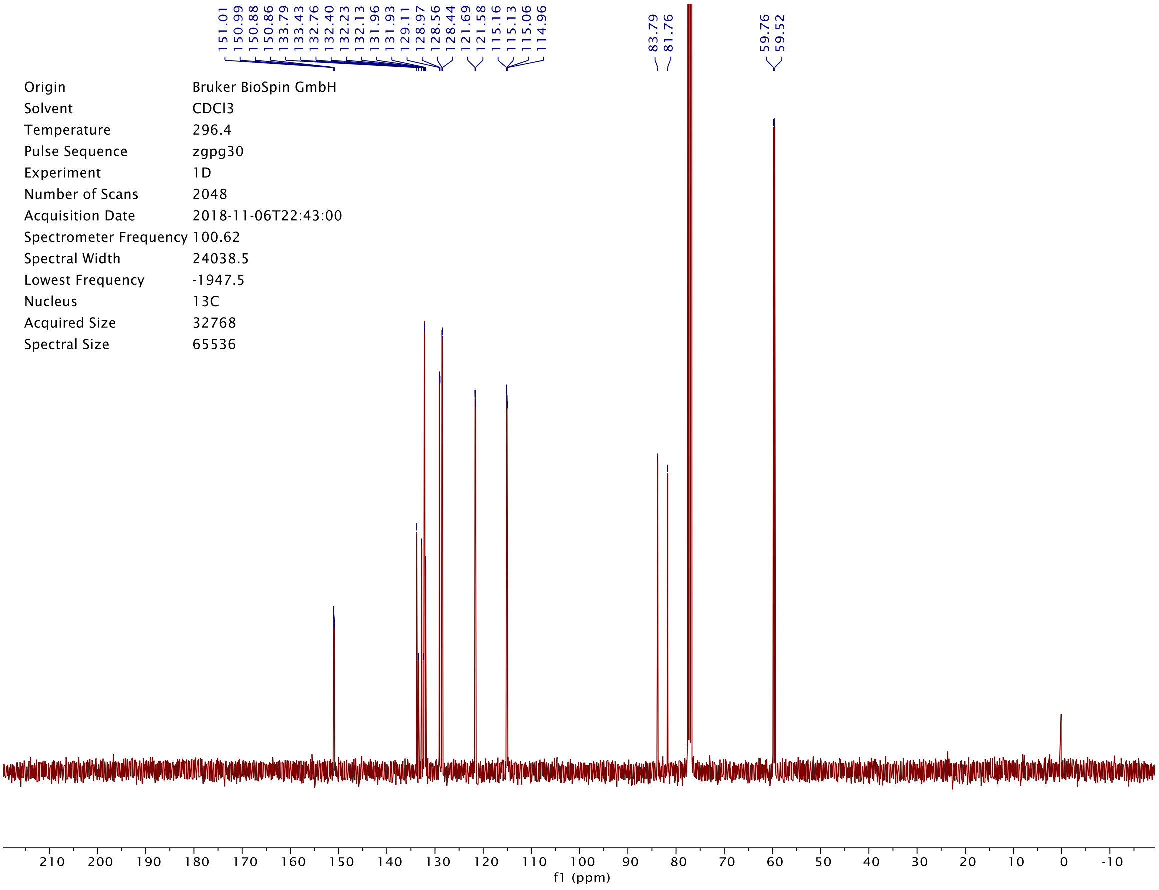
151.01
150.99
150.88
150.86
133.79
133.43
132.76
132.40
132.23
132.13
131.96
131.93
129.11
128.97
128.56
128.44
121.69
121.58
115.16
115.13
115.06
114.96
83.79
81.76
59.76
59.52
Origin Bruker BioSpin GmbH
Solvent CDCl3
Temperature 296.4
Pulse Sequence zgpg30
Experiment 1D
Number of Scans 2048
Acquisition Date 2018-11-06T22:43:00
Spectrometer Frequency 100.62
Spectral Width 24038.5
Lowest Frequency -1947.5
Nucleus 13C
Acquired Size 32768
Spectral Size 65536
f1 (ppm)

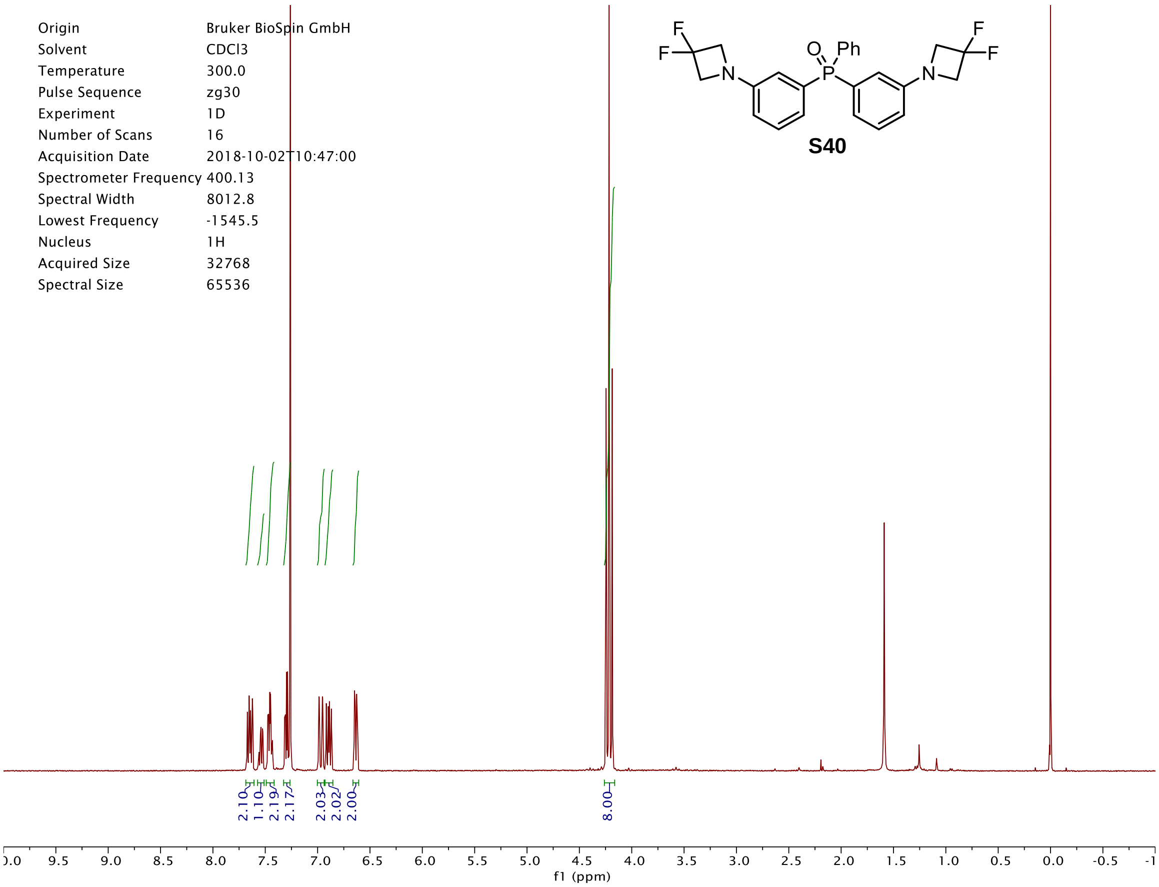
Origin Bruker BioSpin GmbH
Solvent CDCl3
Temperature 300.0
Pulse Sequence zg30
Experiment 1D
Number of Scans 16
Acquisition Date 2018-10-02T10:47:00
Spectrometer Frequency 400.13
Spectral Width 8012.8
Lowest Frequency -1545.5
Nucleus 1H
Acquired Size 32768
Spectral Size 65536
S40
2.10
1.10
2.19
2.17
2.03
2.02
2.00
8.00
f1 (ppm)

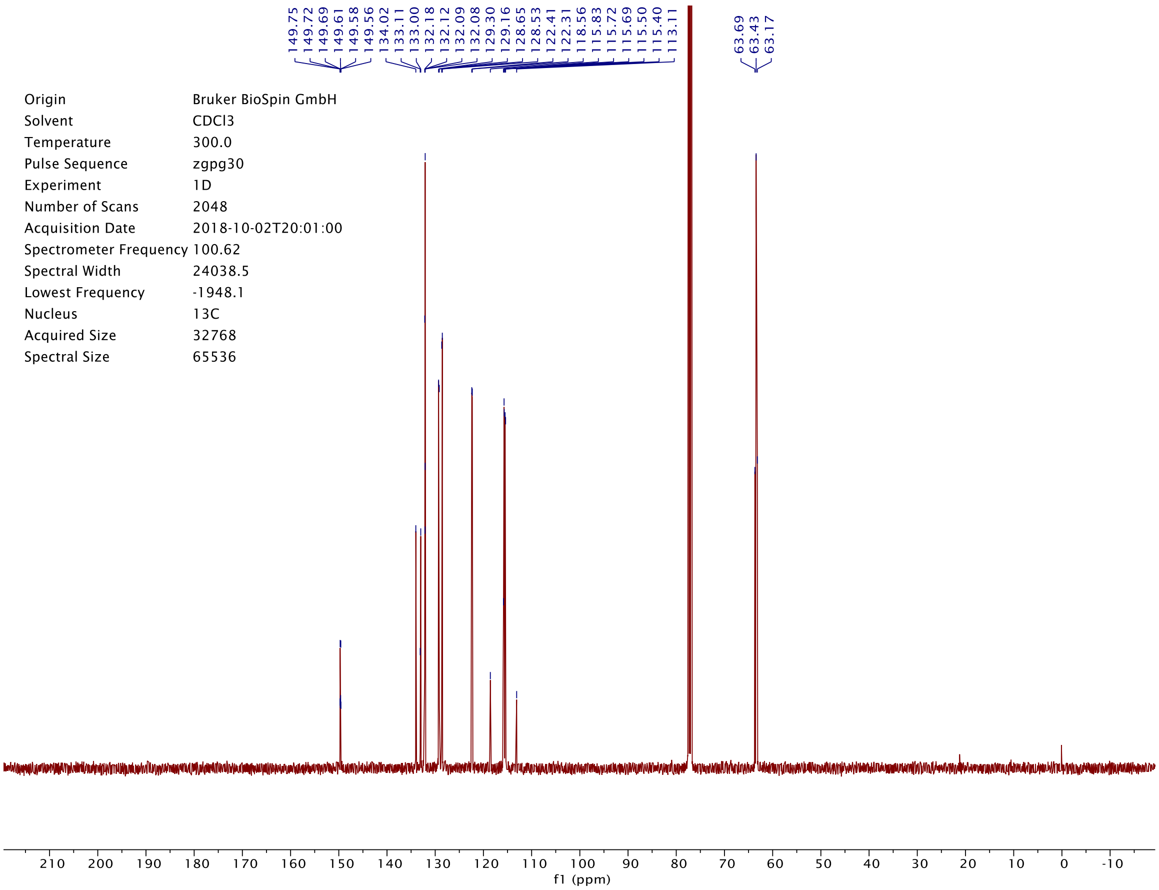
149.75
149.72
149.69
149.61
149.58
149.56
134.02
133.11
133.00
132.18
132.12
132.09
132.08
129.30
129.16
128.65
128.53
122.41
122.31
118.56
115.83
115.72
115.69
115.50
115.40
113.11
63.69
63.43
63.17
Origin Bruker BioSpin GmbH
Solvent CDCl3
Temperature 300.0
Pulse Sequence zgpg30
Experiment 1D
Number of Scans 2048
Acquisition Date 2018-10-02T20:01:00
Spectrometer Frequency 100.62
Spectral Width 24038.5
Lowest Frequency -1948.1
Nucleus 13C
Acquired Size 32768
Spectral Size 65536
f1 (ppm)

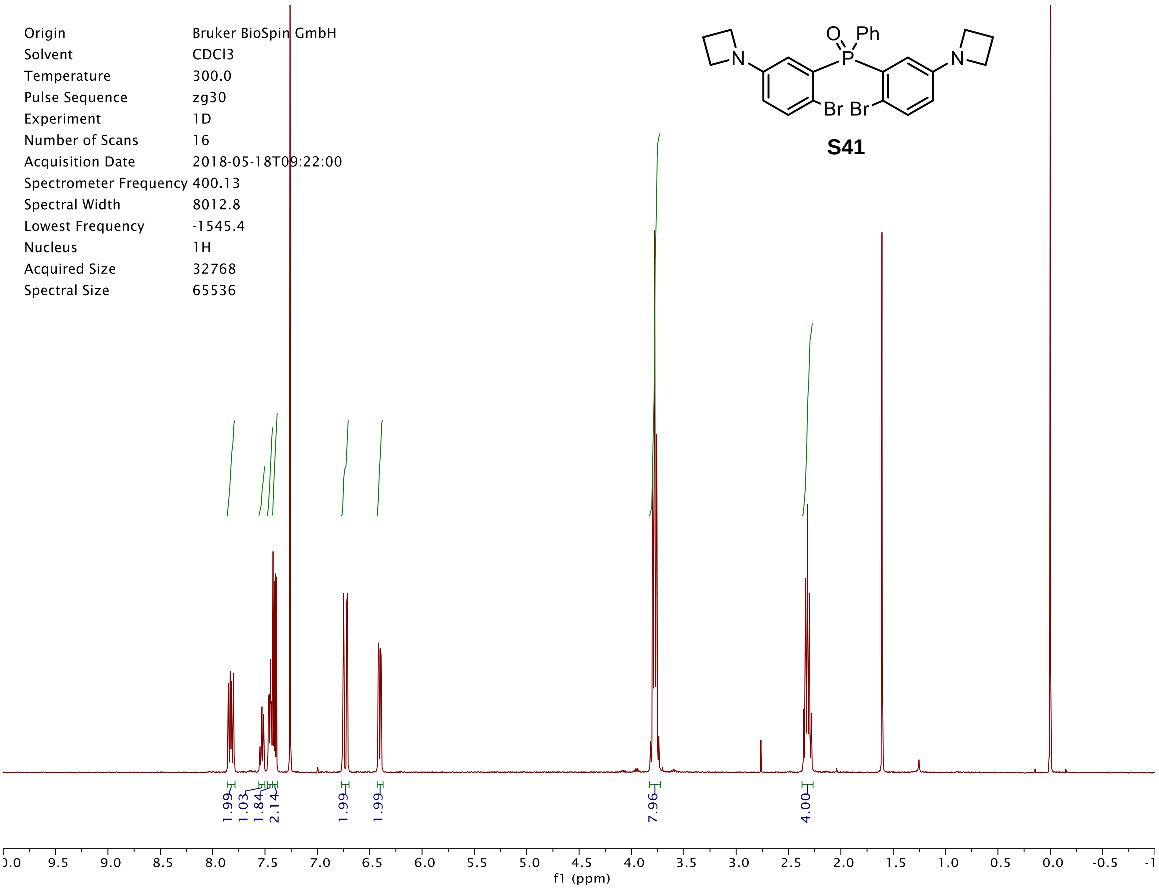
S41
O
Ph
P
N
Br Br
Origin Bruker BioSpin GmbH
Solvent CDCl3
Temperature 300.0
Pulse Sequence zg30
Experiment 1D
Number of Scans 16
Acquisition Date 2018-05-18T09:22:00
Spectrometer Frequency 400.13
Spectral Width 8012.8
Lowest Frequency -1545.4
Nucleus 1H
Acquired Size 32768
Spectral Size 65536
1.99
1.03
1.84
2.14
1.99
1.99
7.96
4.00
f1 (ppm)

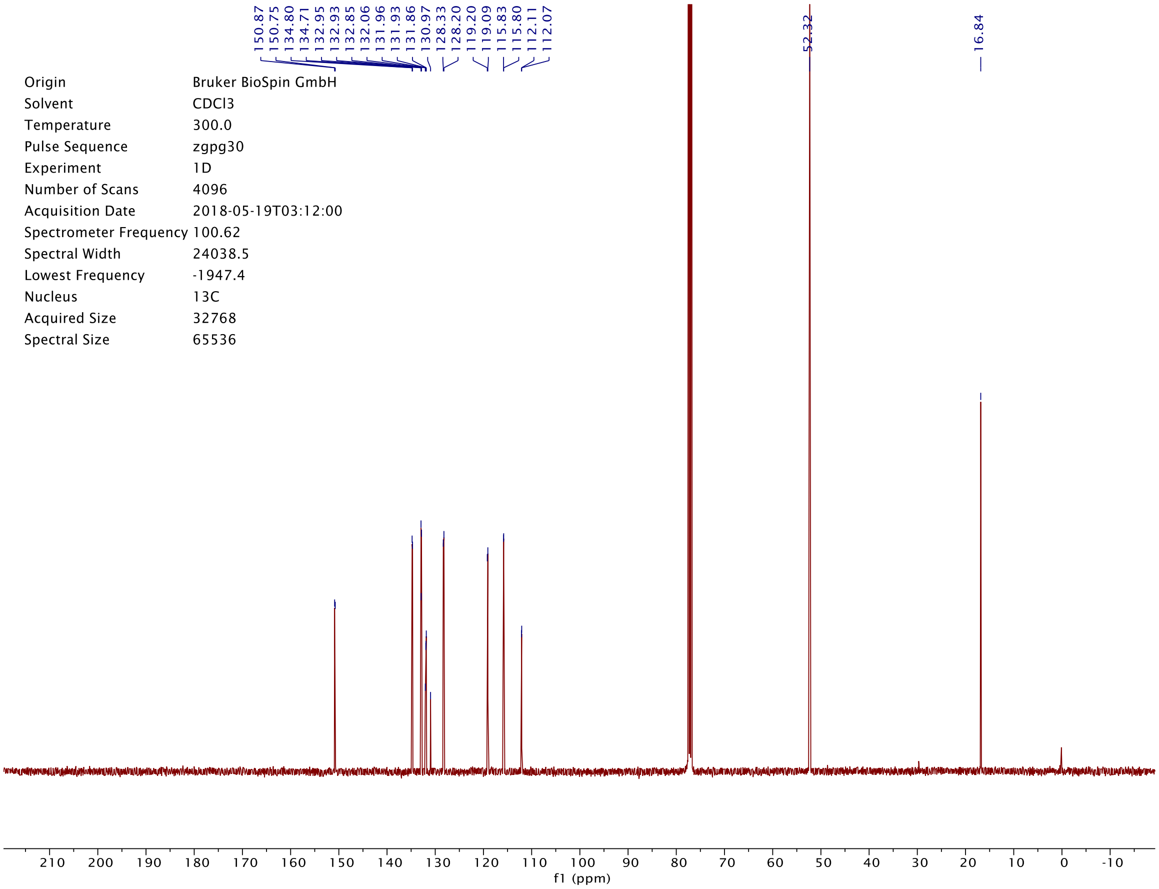
150.87
150.75
134.80
134.71
132.95
132.93
132.85
132.06
131.96
131.93
131.86
130.97
128.33
128.20
119.20
119.09
115.83
115.80
112.11
112.07
52.32
16.84
Origin Bruker BioSpin GmbH
Solvent CDCl3
Temperature 300.0
Pulse Sequence zgpg30
Experiment 1D
Number of Scans 4096
Acquisition Date 2018-05-19T03:12:00
Spectrometer Frequency 100.62
Spectral Width 24038.5
Lowest Frequency -1947.4
Nucleus 13C
Acquired Size 32768
Spectral Size 65536
f1 (ppm)

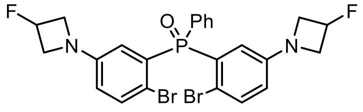
F
O
Ph
P
N
Br
Br
N
F

**S42**

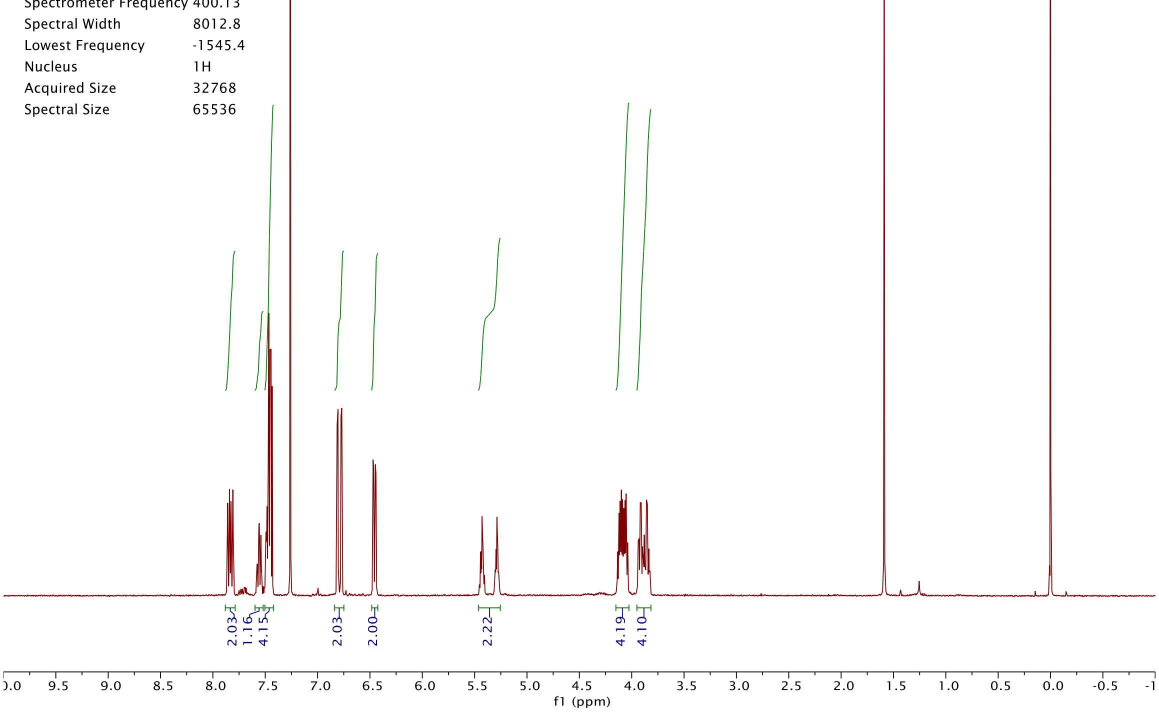
Origin Bruker BioSpin GmbH
Solvent CDCl3
Temperature 300.0
Pulse Sequence zg30
Experiment 1D
Number of Scans 16
Acquisition Date 2018-10-02T11:01:00
Spectrometer Frequency 400.13
Spectral Width 8012.8
Lowest Frequency -1545.4
Nucleus 1H
Acquired Size 32768
Spectral Size 65536
2.03
1.16
4.15
2.03
2.00
2.22
4.19
4.10
f1 (ppm)

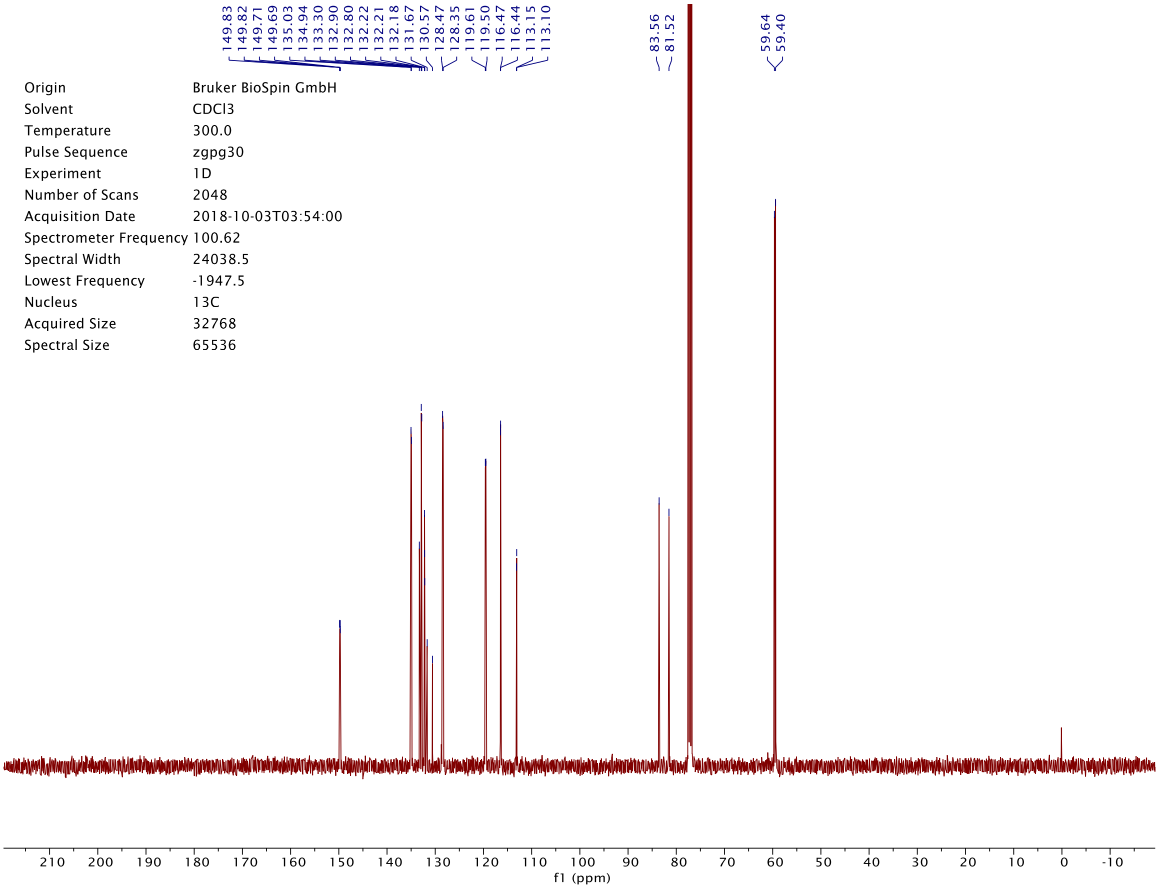
149.83
149.82
149.71
149.69
135.03
134.94
133.30
132.90
132.80
132.22
132.21
132.18
131.67
130.57
128.47
128.35
119.61
119.50
116.47
116.44
113.15
113.10
83.56
81.52
59.64
59.40
Origin Bruker BioSpin GmbH
Solvent CDCl3
Temperature 300.0
Pulse Sequence zgpg30
Experiment 1D
Number of Scans 2048
Acquisition Date 2018-10-03T03:54:00
Spectrometer Frequency 100.62
Spectral Width 24038.5
Lowest Frequency -1947.5
Nucleus 13C
Acquired Size 32768
Spectral Size 65536
f1 (ppm)

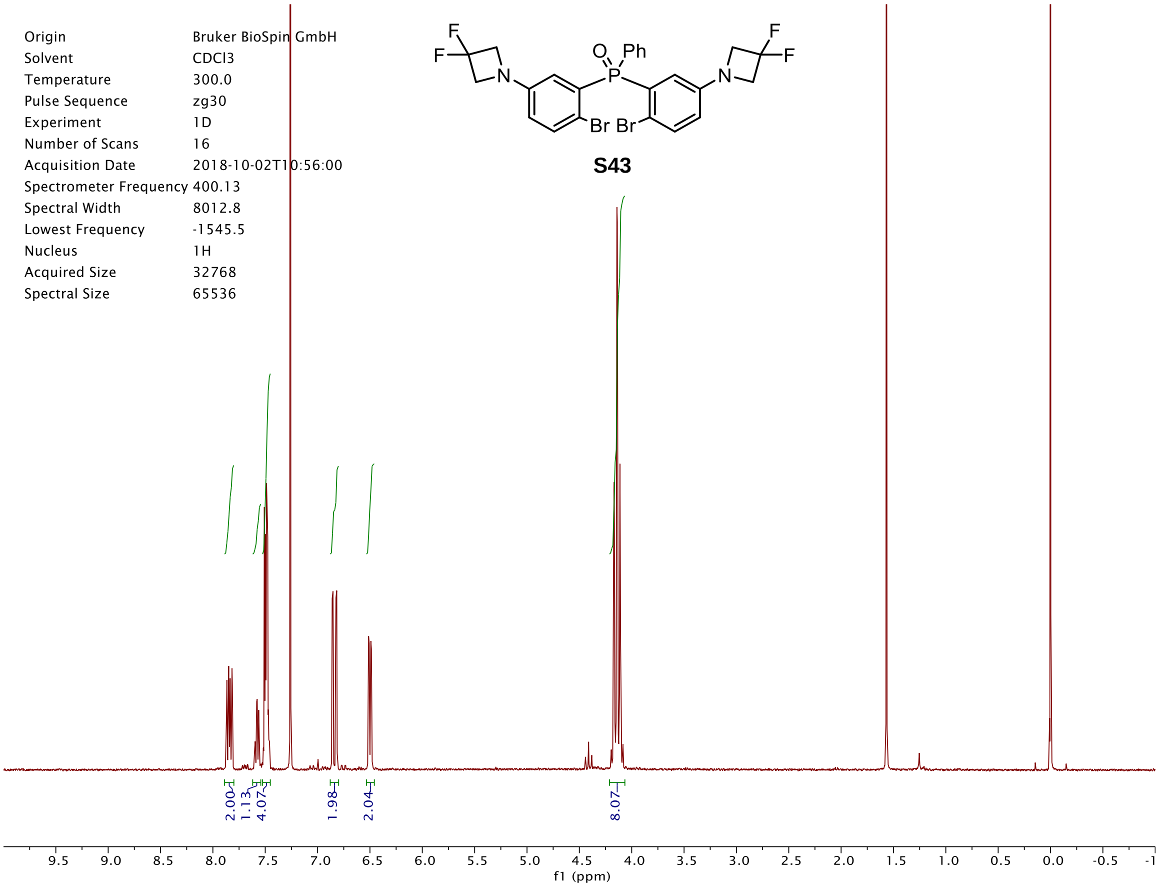
Origin Bruker BioSpin GmbH
Solvent CDCl3
Temperature 300.0
Pulse Sequence zg30
Experiment 1D
Number of Scans 16
Acquisition Date 2018-10-02T10:56:00
Spectrometer Frequency 400.13
Spectral Width 8012.8
Lowest Frequency -1545.5
Nucleus 1H
Acquired Size 32768
Spectral Size 65536
F
F
N
O
P
Ph
Br
Br
N
F
F
S43
2.00
1.13
4.07
1.98
2.04
8.07
f1 (ppm)

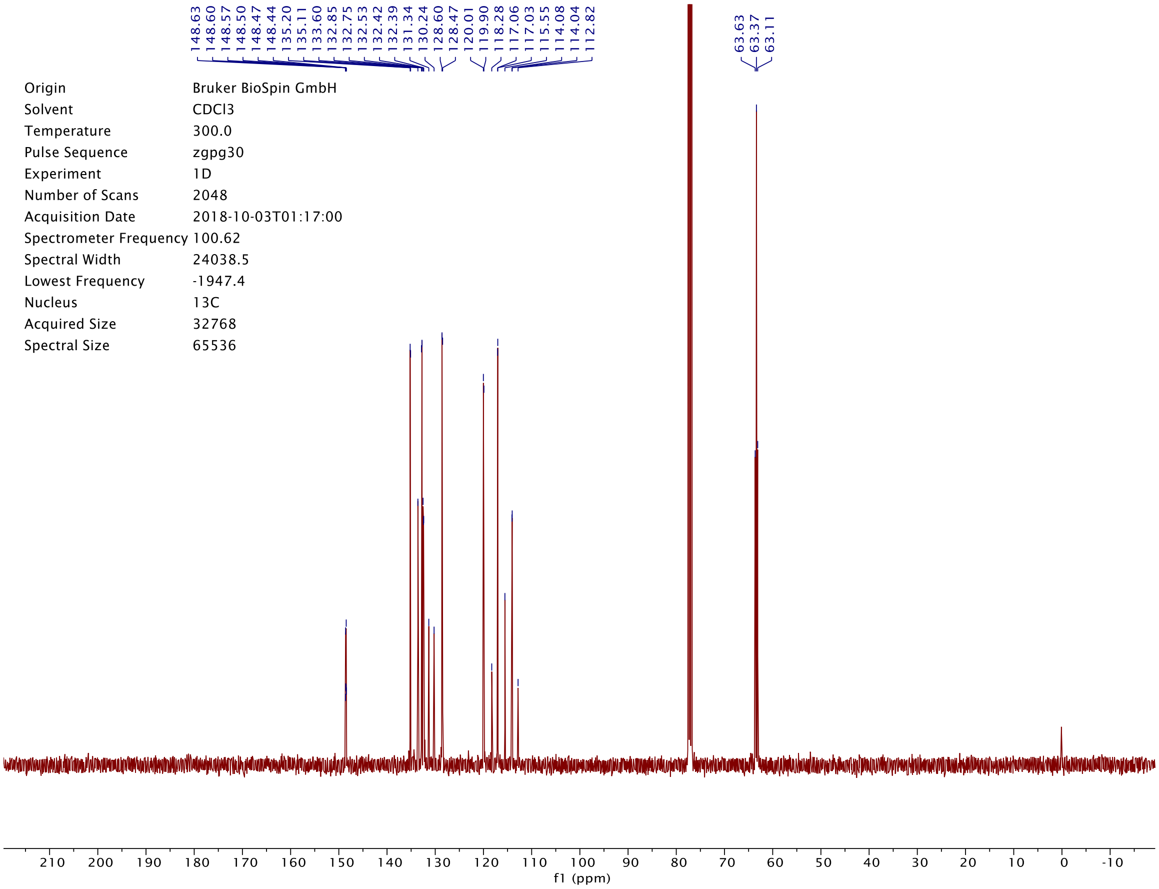
148.63
148.60
148.57
148.50
148.47
148.44
135.20
135.11
133.60
132.85
132.75
132.53
132.42
132.39
131.34
130.24
128.60
128.47
120.01
119.90
118.28
117.06
117.03
115.55
114.08
114.04
112.82
63.63
63.37
63.11
Origin Bruker BioSpin GmbH
Solvent CDCl3
Temperature 300.0
Pulse Sequence zgpg30
Experiment 1D
Number of Scans 2048
Acquisition Date 2018-10-03T01:17:00
Spectrometer Frequency 100.62
Spectral Width 24038.5
Lowest Frequency -1947.4
Nucleus 13C
Acquired Size 32768
Spectral Size 65536
f1 (ppm)

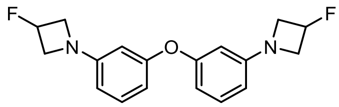
F
N
O
N
F

**S45**

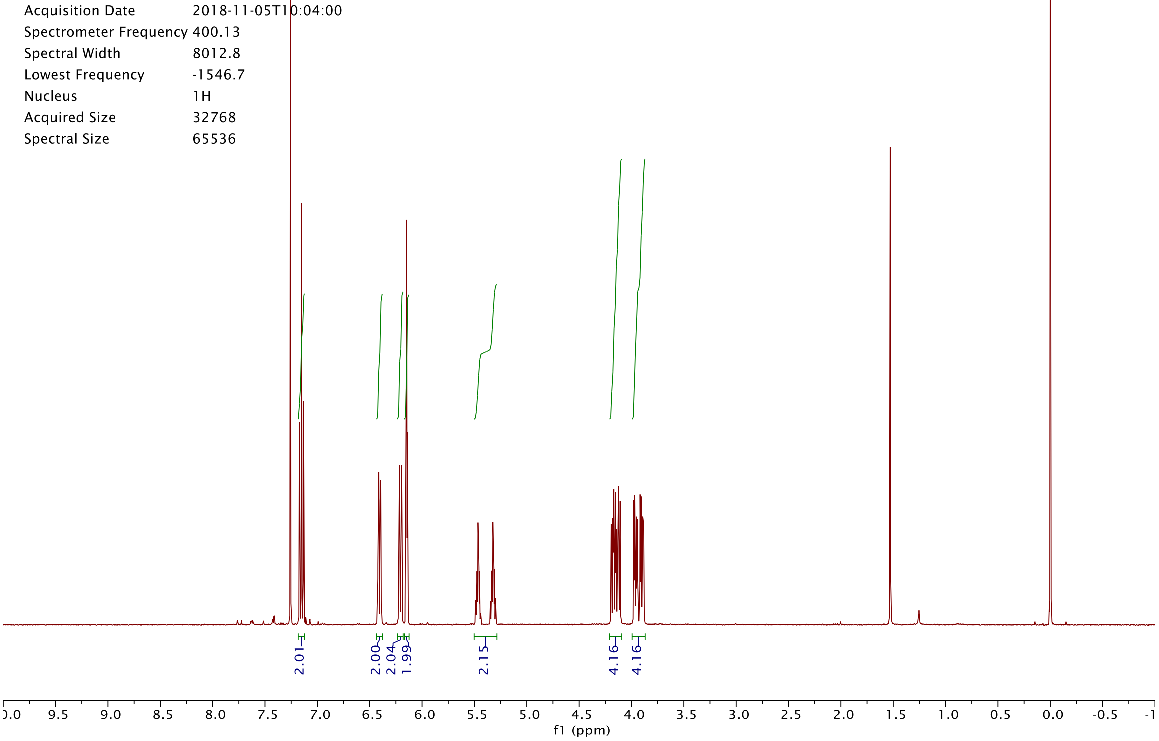
Origin Bruker BioSpin GmbH
Solvent CDCl3
Temperature 300.0
Pulse Sequence zg30
Experiment 1D
Number of Scans 16
Acquisition Date 2018-11-05T10:04:00
Spectrometer Frequency 400.13
Spectral Width 8012.8
Lowest Frequency -1546.7
Nucleus 1H
Acquired Size 32768
Spectral Size 65536
2.01
2.00
2.04
1.99
2.15
4.16
4.16
f1 (ppm)

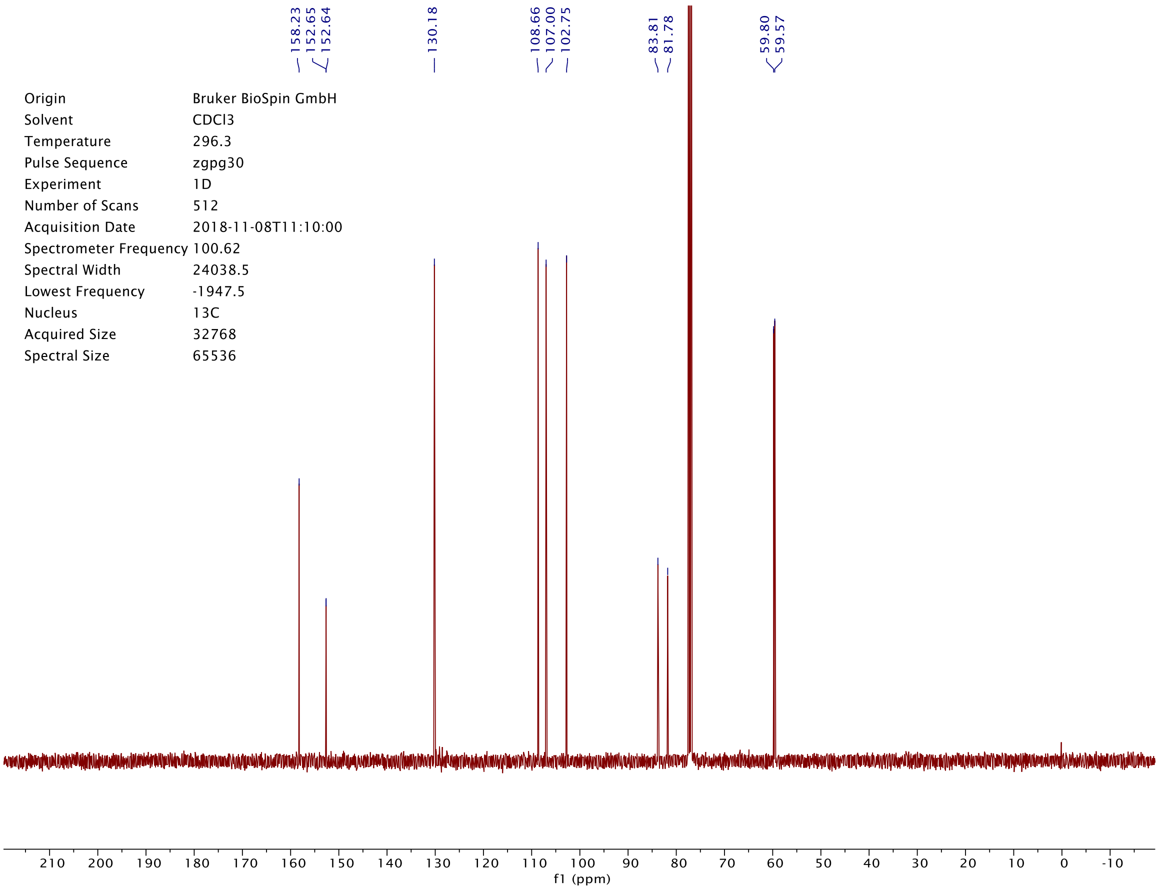
158.23
152.65
152.64
130.18
108.66
107.00
102.75
83.81
81.78
59.80
59.57
Origin Bruker BioSpin GmbH
Solvent CDCl3
Temperature 296.3
Pulse Sequence zgpg30
Experiment 1D
Number of Scans 512
Acquisition Date 2018-11-08T11:10:00
Spectrometer Frequency 100.62
Spectral Width 24038.5
Lowest Frequency -1947.5
Nucleus 13C
Acquired Size 32768
Spectral Size 65536
f1 (ppm)

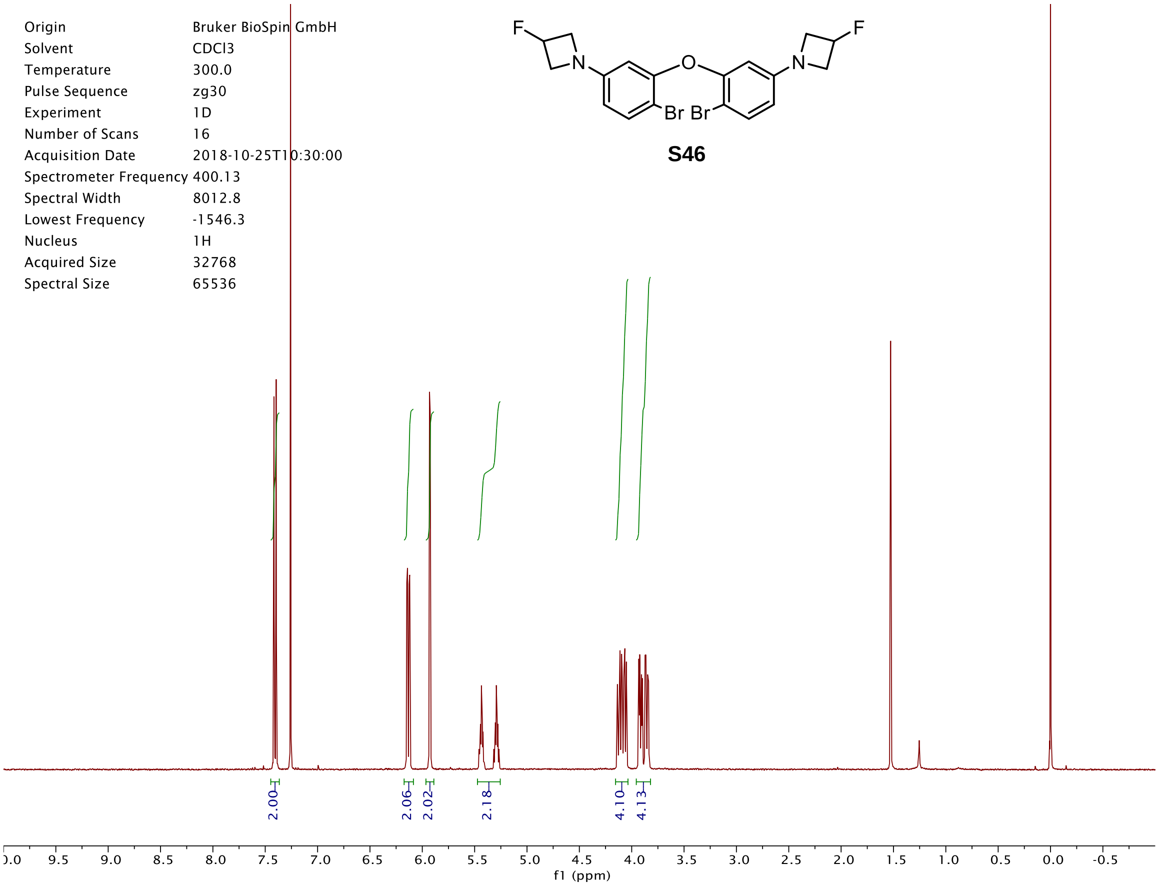

Origin Bruker BioSpin GmbH
Solvent CDCl3
Temperature 300.0
Pulse Sequence zg30
Experiment 1D
Number of Scans 16
Acquisition Date 2018-10-25T10:30:00
Spectrometer Frequency 400.13
Spectral Width 8012.8
Lowest Frequency -1546.3
Nucleus 1H
Acquired Size 32768
Spectral Size 65536
F
N
O
Br Br
S46
2.00
2.06
2.02
2.18
4.10
4.13
f1 (ppm)


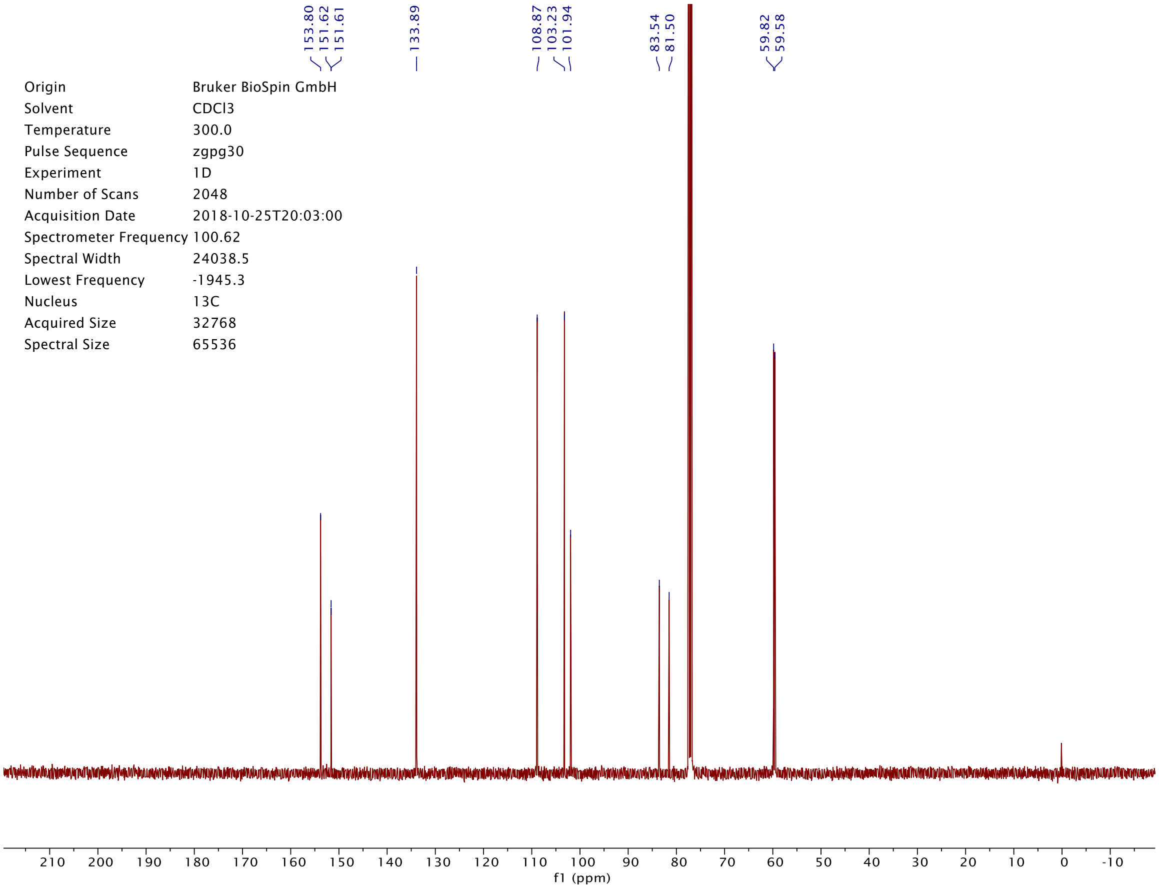

153.80
151.62
151.61
133.89
108.87
103.23
101.94
83.54
81.50
59.82
59.58
Origin Bruker BioSpin GmbH
Solvent CDCl3
Temperature 300.0
Pulse Sequence zgpg30
Experiment 1D
Number of Scans 2048
Acquisition Date 2018-10-25T20:03:00
Spectrometer Frequency 100.62
Spectral Width 24038.5
Lowest Frequency -1945.3
Nucleus 13C
Acquired Size 32768
Spectral Size 65536
f1 (ppm)

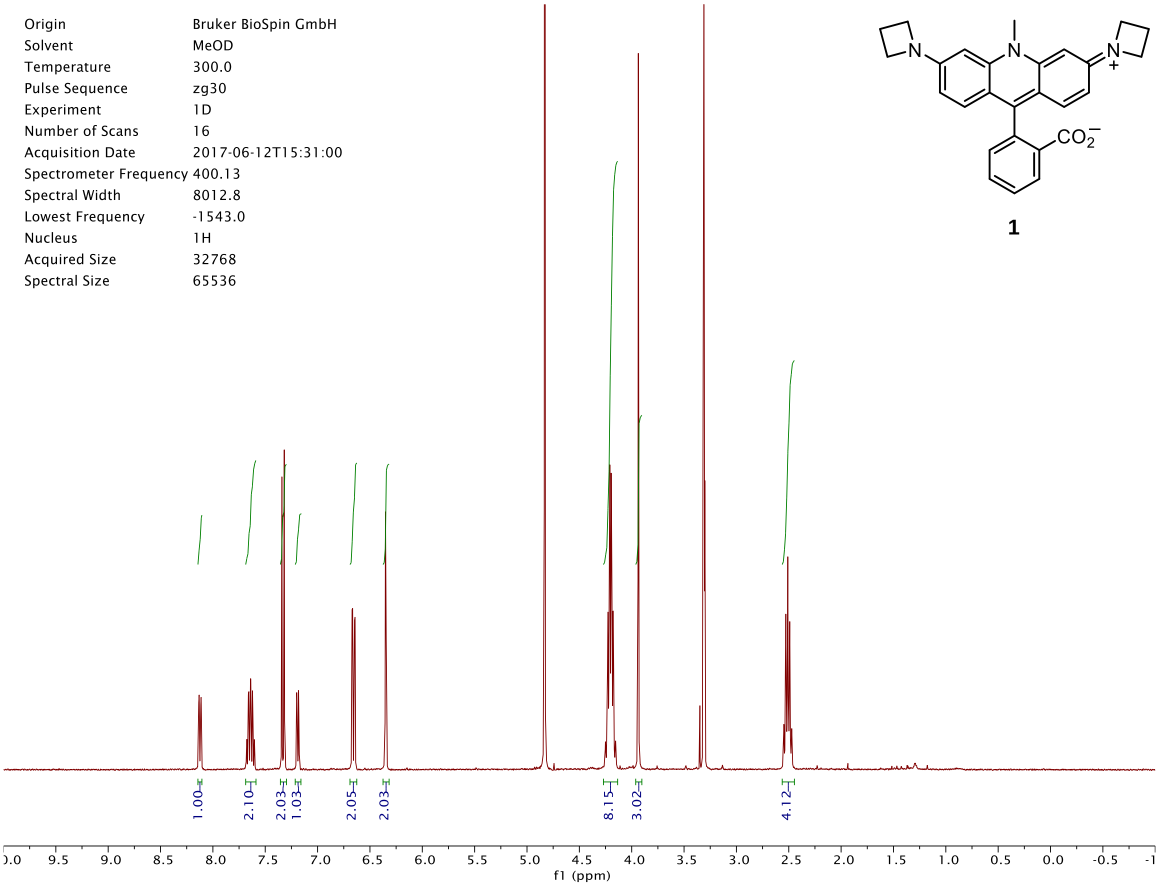

1
Origin Bruker BioSpin GmbH
Solvent MeOD
Temperature 300.0
Pulse Sequence zg30
Experiment 1D
Number of Scans 16
Acquisition Date 2017-06-12T15:31:00
Spectrometer Frequency 400.13
Spectral Width 8012.8
Lowest Frequency -1543.0
Nucleus 1H
Acquired Size 32768
Spectral Size 65536
1.00
2.10
2.03
1.03
2.05
2.03
8.15
3.02
4.12
f1 (ppm)


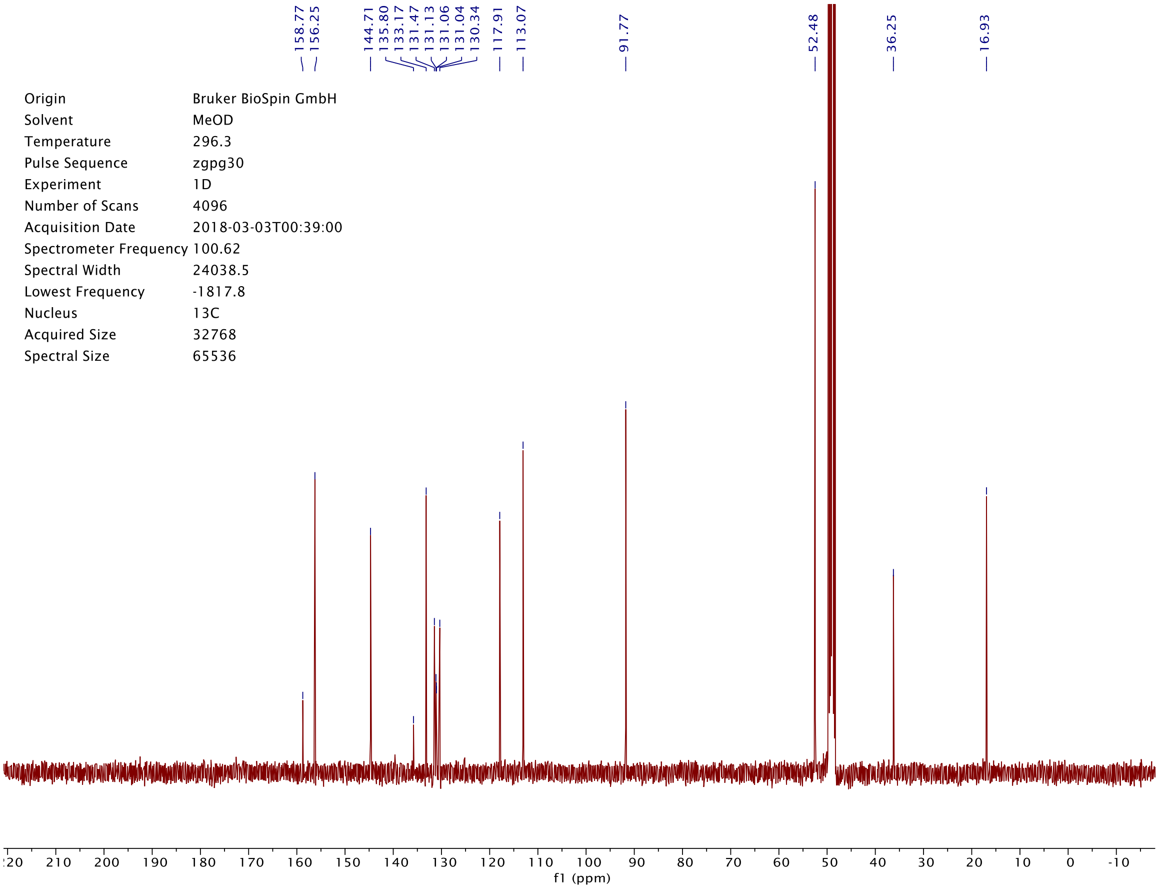

158.77
156.25
144.71
135.80
133.17
131.47
131.13
131.06
131.04
130.34
117.91
113.07
91.77
52.48
36.25
16.93
Origin Bruker BioSpin GmbH
Solvent MeOD
Temperature 296.3
Pulse Sequence zgpg30
Experiment 1D
Number of Scans 4096
Acquisition Date 2018-03-03T00:39:00
Spectrometer Frequency 100.62
Spectral Width 24038.5
Lowest Frequency -1817.8
Nucleus 13C
Acquired Size 32768
Spectral Size 65536
f1 (ppm)

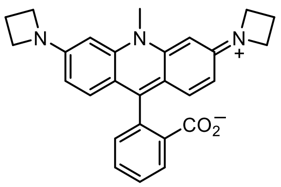

**1**

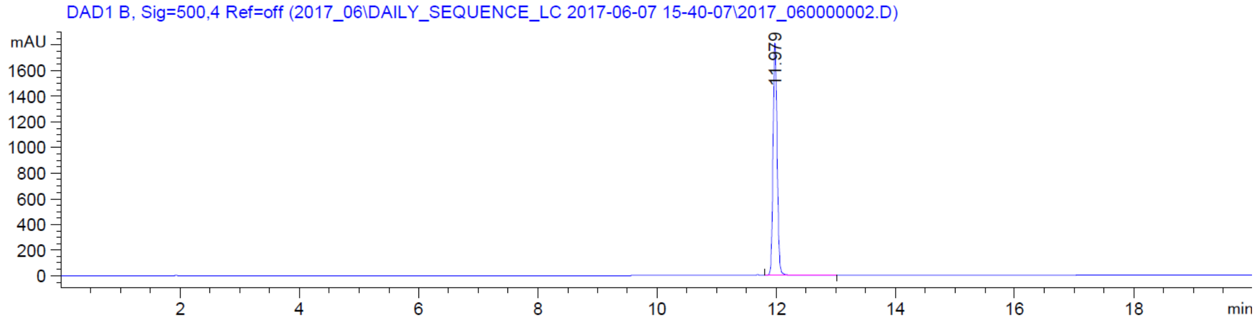

DAD1 B, Sig=500,4 Ref=off (2017_06\DAILY_SEQUENCE_LC 2017-06-07 15-40-07\2017_060000002.D)
mAU
11.979
min


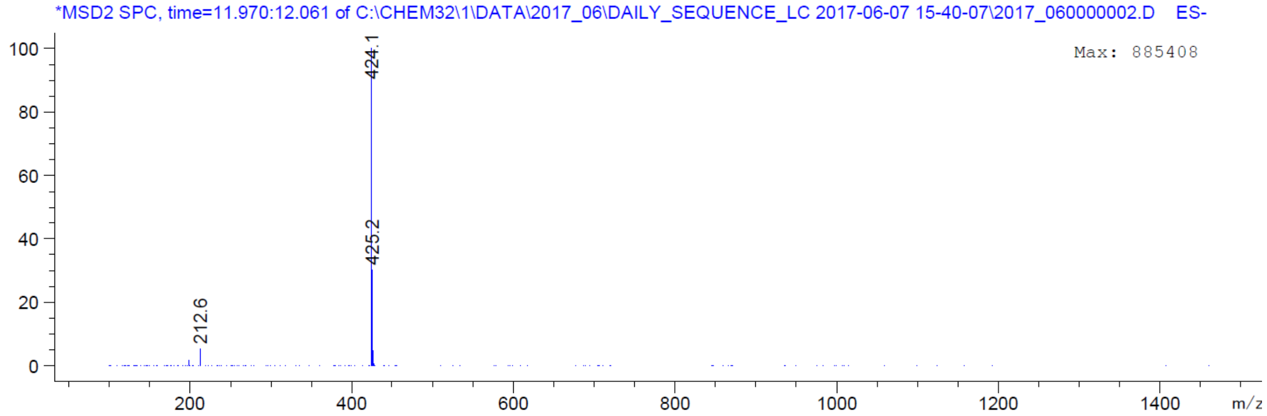

*MSD2 SPC, time=11.970:12.061 of C:\CHEM32\1\DATA\2017_06\DAILY_SEQUENCE_LC 2017-06-07 15-40-07\2017_060000002.D ES-
Max: 885408
424.1
425.2
212.6
m/z

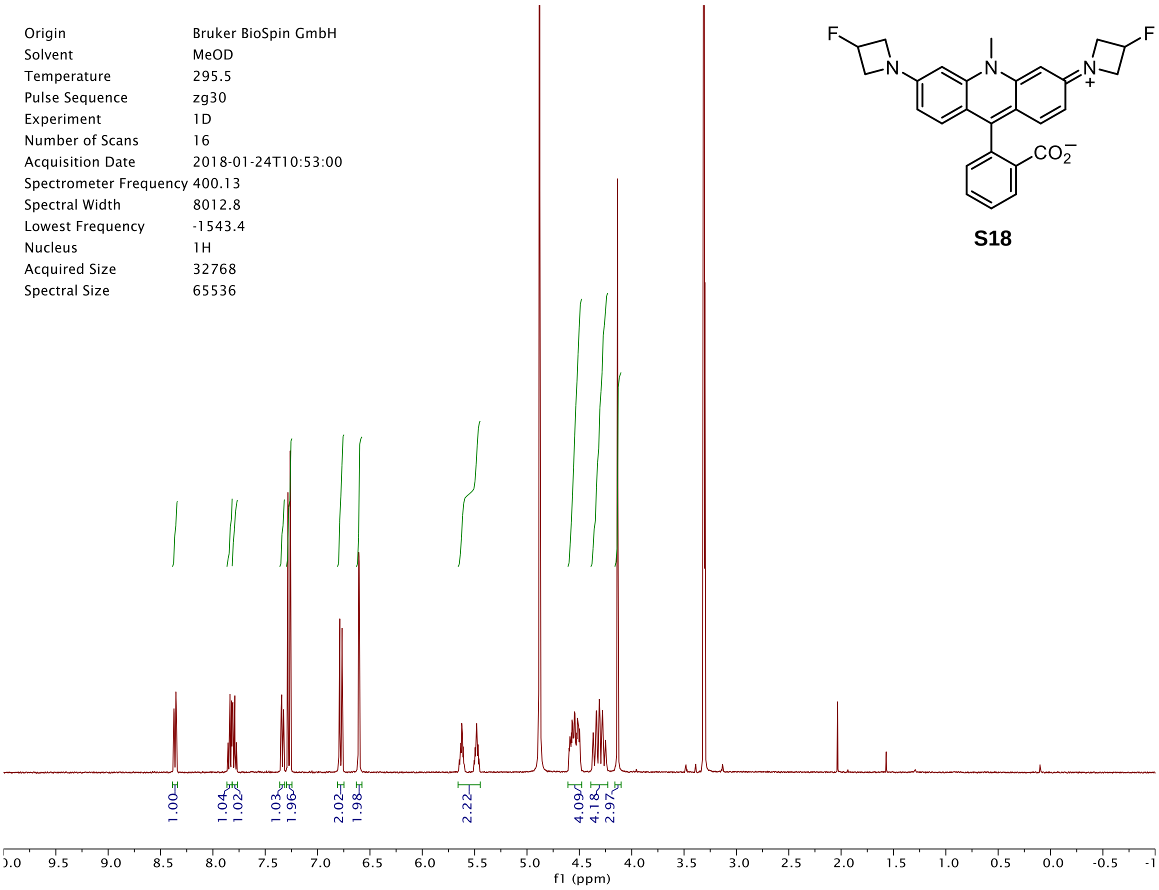
Origin Bruker BioSpin GmbH
Solvent MeOD
Temperature 295.5
Pulse Sequence zg30
Experiment 1D
Number of Scans 16
Acquisition Date 2018-01-24T10:53:00
Spectrometer Frequency 400.13
Spectral Width 8012.8
Lowest Frequency -1543.4
Nucleus 1H
Acquired Size 32768
Spectral Size 65536
1.00
1.04
1.02
1.03
1.96
2.02
1.98
2.22
4.09
4.18
2.97
f1 (ppm)
F
N
N
N
+
F
$CO_2^-$

**S18**

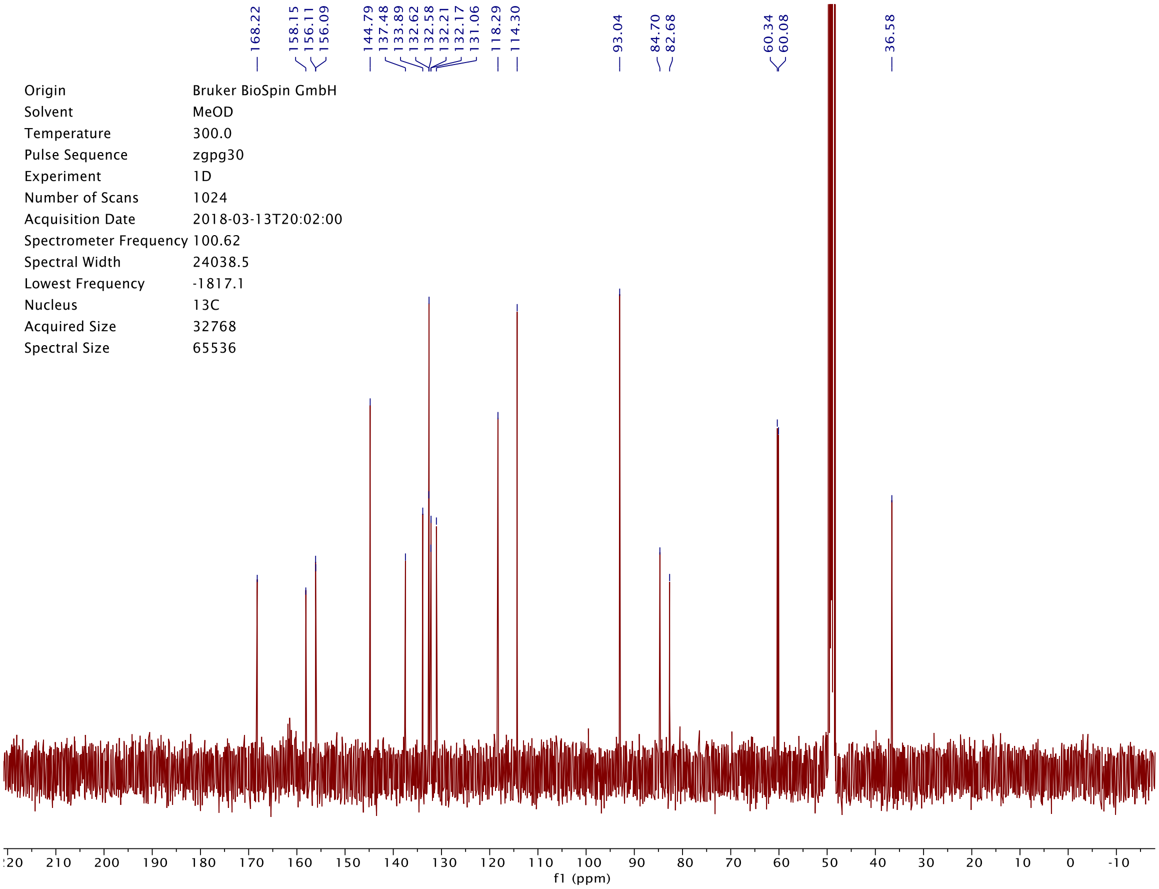
168.22
158.15
156.11
156.09
144.79
137.48
133.89
132.62
132.58
132.21
132.17
131.06
118.29
114.30
93.04
84.70
82.68
60.34
60.08
36.58
Origin Bruker BioSpin GmbH
Solvent MeOD
Temperature 300.0
Pulse Sequence zgpg30
Experiment 1D
Number of Scans 1024
Acquisition Date 2018-03-13T20:02:00
Spectrometer Frequency 100.62
Spectral Width 24038.5
Lowest Frequency -1817.1
Nucleus 13C
Acquired Size 32768
Spectral Size 65536
f1 (ppm)

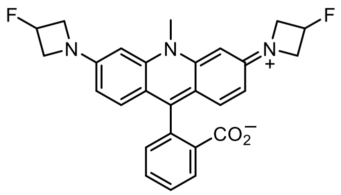
F
N
N
N
+
F
CO2−

**S18**

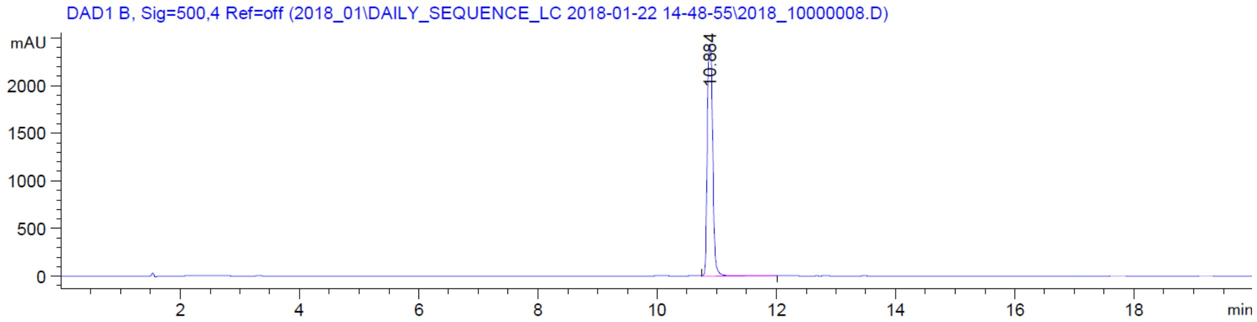
DAD1 B, Sig=500,4 Ref=off (2018_01\DAILY_SEQUENCE_LC 2018-01-22 14-48-55\2018_10000008.D)
mAU
2000
1500
1000
500
0
10.884
2
4
6
8
10
12
14
16
18
min

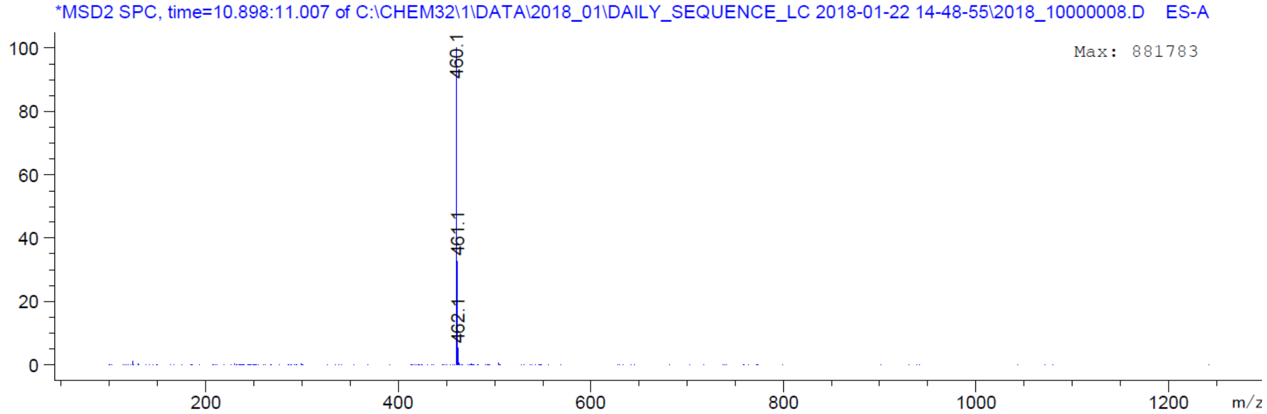
*MSD2 SPC, time=10.898:11.007 of C:\CHEM32\1\DATA\2018_01\DAILY_SEQUENCE_LC 2018-01-22 14-48-55\2018_10000008.D ES-A
Max: 881783
100
80
60
40
20
0
460.1
461.1
462.1
200
400
600
800
1000
1200
m/z

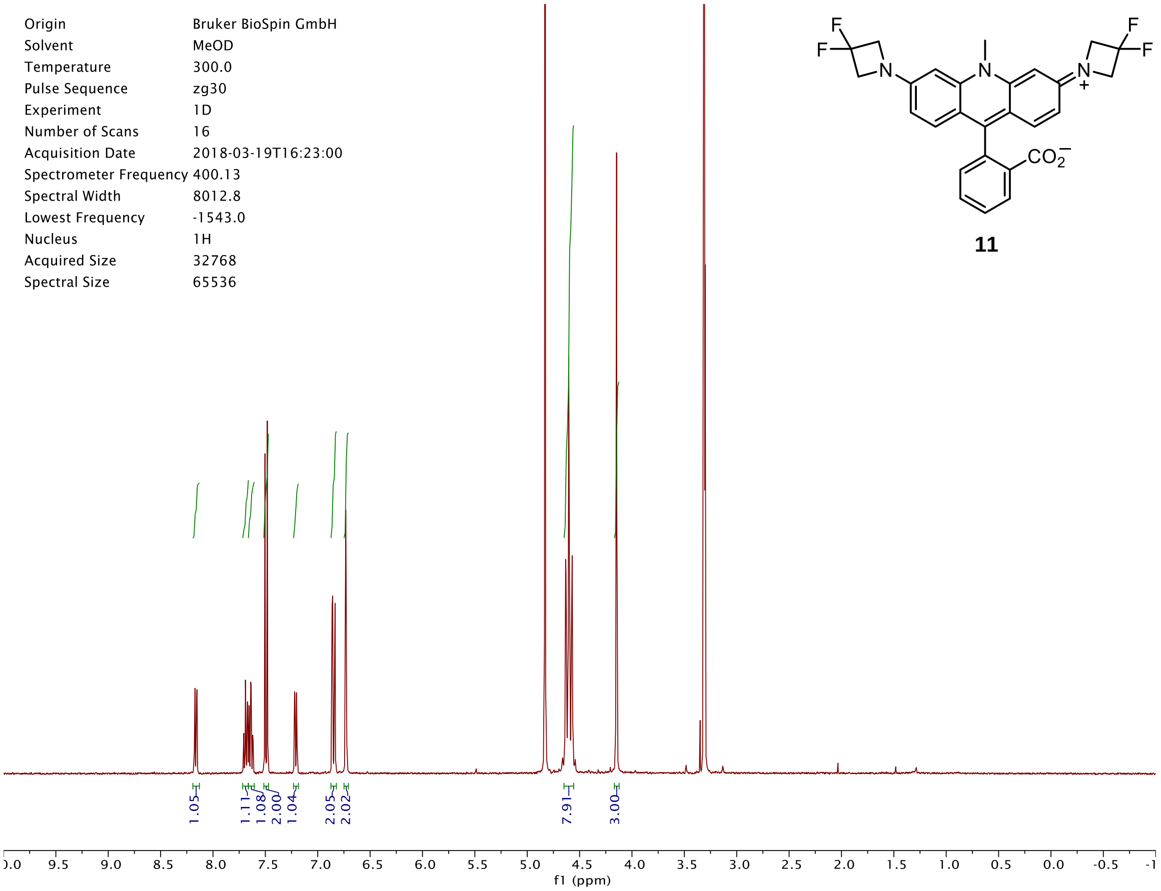

Origin Bruker BioSpin GmbH
Solvent MeOD
Temperature 300.0
Pulse Sequence zg30
Experiment 1D
Number of Scans 16
Acquisition Date 2018-03-19T16:23:00
Spectrometer Frequency 400.13
Spectral Width 8012.8
Lowest Frequency -1543.0
Nucleus 1H
Acquired Size 32768
Spectral Size 65536
1.05
1.11
1.08
2.00
1.04
2.05
2.02
7.91
3.00
f1 (ppm)
F
F
N
N
N
F
F
+
CO2−


**11**

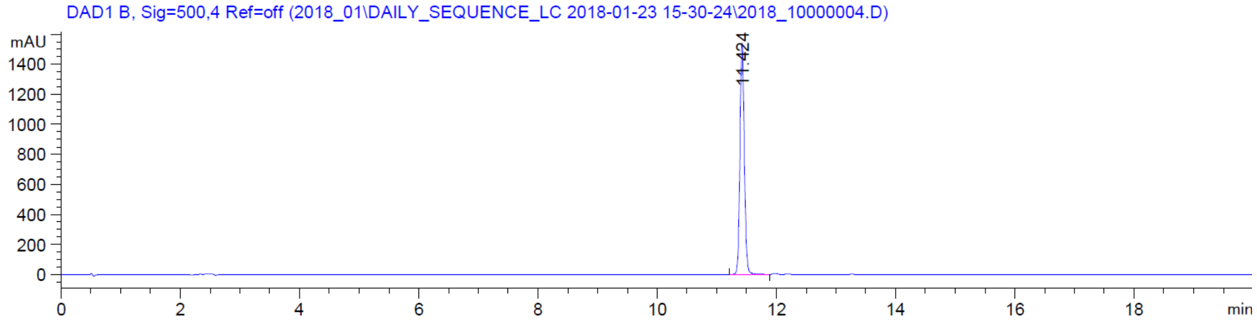

DAD1 B, Sig=500,4 Ref=off (2018_01\DAILY_SEQUENCE_LC 2018-01-23 15-30-24\2018_10000004.D)
mAU
11.424
min


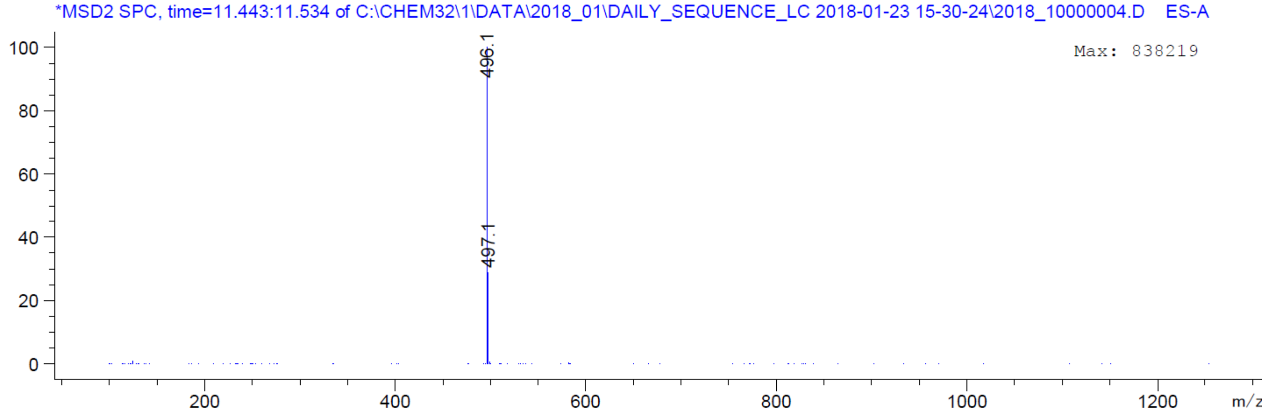

*MSD2 SPC, time=11.443:11.534 of C:\CHEM32\1\DATA\2018_01\DAILY_SEQUENCE_LC 2018-01-23 15-30-24\2018_10000004.D ES-A
Max: 838219
496.1
497.1
m/z

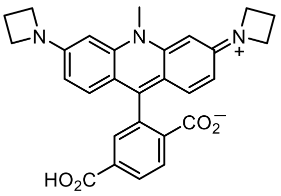

**S22**

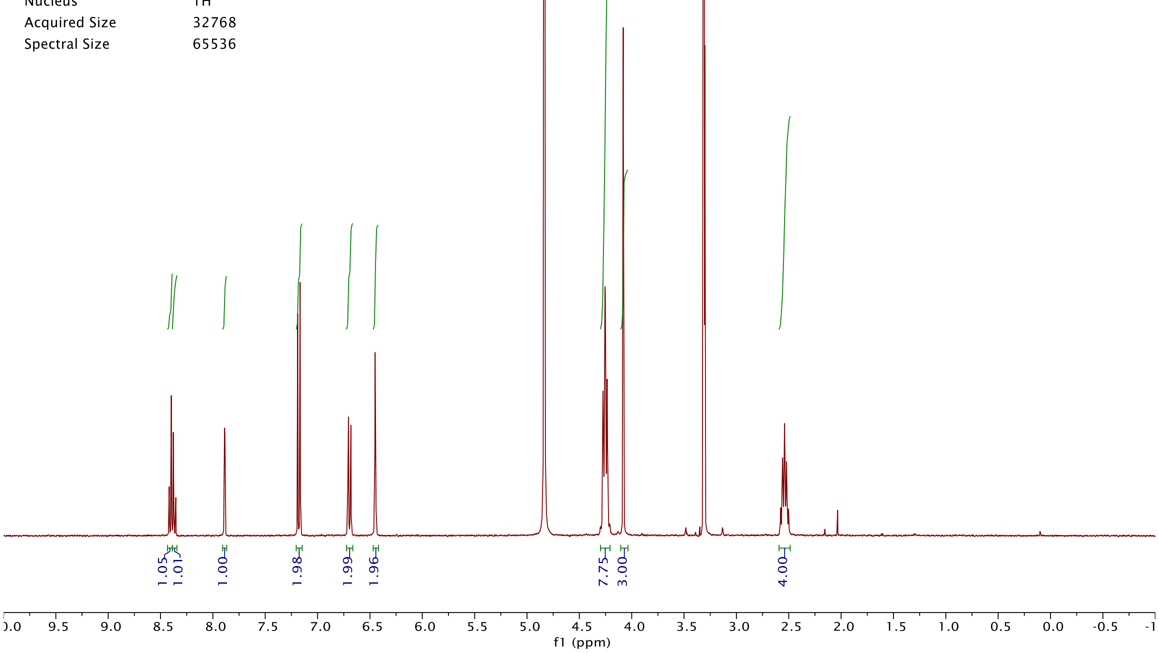

Origin Bruker BioSpin GmbH
Solvent MeOD
Temperature 300.0
Pulse Sequence zg30
Experiment 1D
Number of Scans 16
Acquisition Date 2018-05-29T11:26:00
Spectrometer Frequency 400.13
Spectral Width 8012.8
Lowest Frequency -1543.0
Nucleus 1H
Acquired Size 32768
Spectral Size 65536
1.05
1.01
1.00
1.98
1.99
1.96
7.75
3.00
4.00
f1 (ppm)


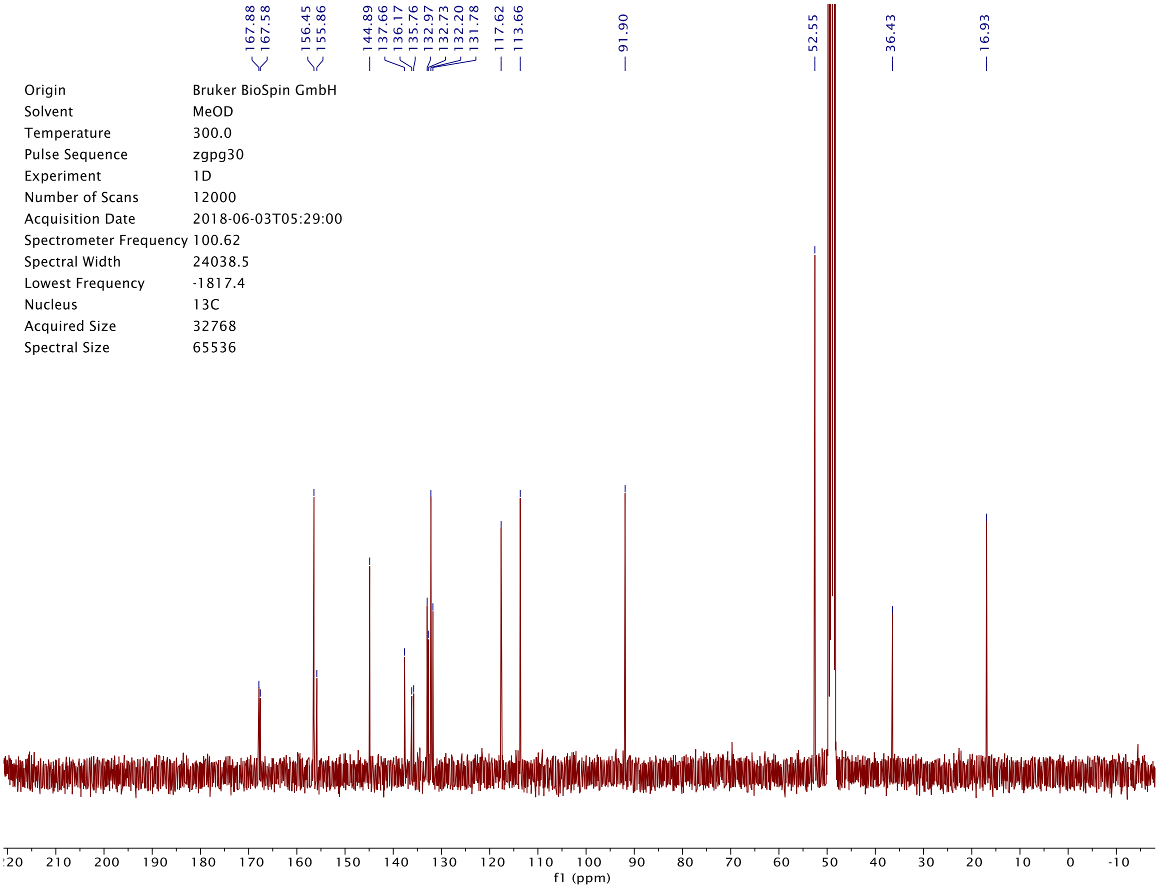

167.88
167.58
156.45
155.86
144.89
137.66
136.17
135.76
132.97
132.73
132.20
131.78
117.62
113.66
91.90
52.55
36.43
16.93
Origin Bruker BioSpin GmbH
Solvent MeOD
Temperature 300.0
Pulse Sequence zgpg30
Experiment 1D
Number of Scans 12000
Acquisition Date 2018-06-03T05:29:00
Spectrometer Frequency 100.62
Spectral Width 24038.5
Lowest Frequency -1817.4
Nucleus 13C
Acquired Size 32768
Spectral Size 65536
f1 (ppm)

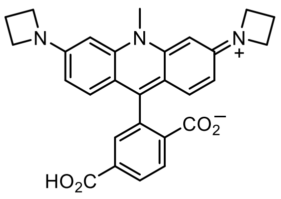

**S22**

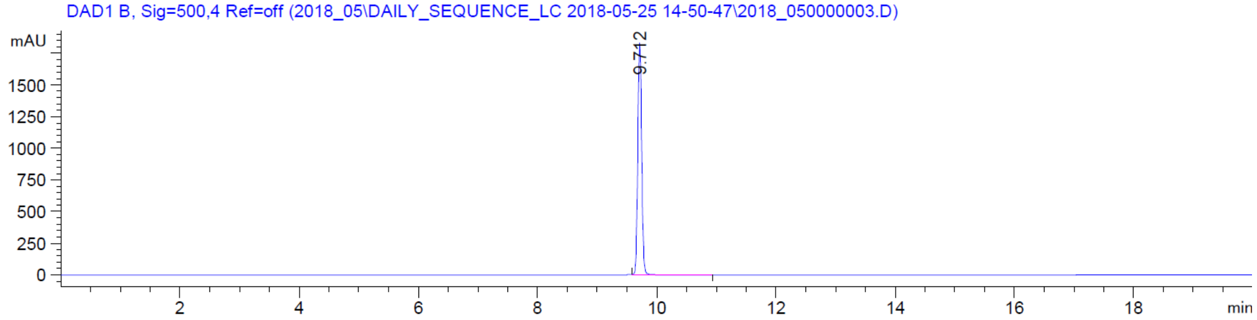

DAD1 B, Sig=500,4 Ref=off (2018_05\DAILY_SEQUENCE_LC 2018-05-25 14-50-47\2018_050000003.D)
mAU
9.712
min


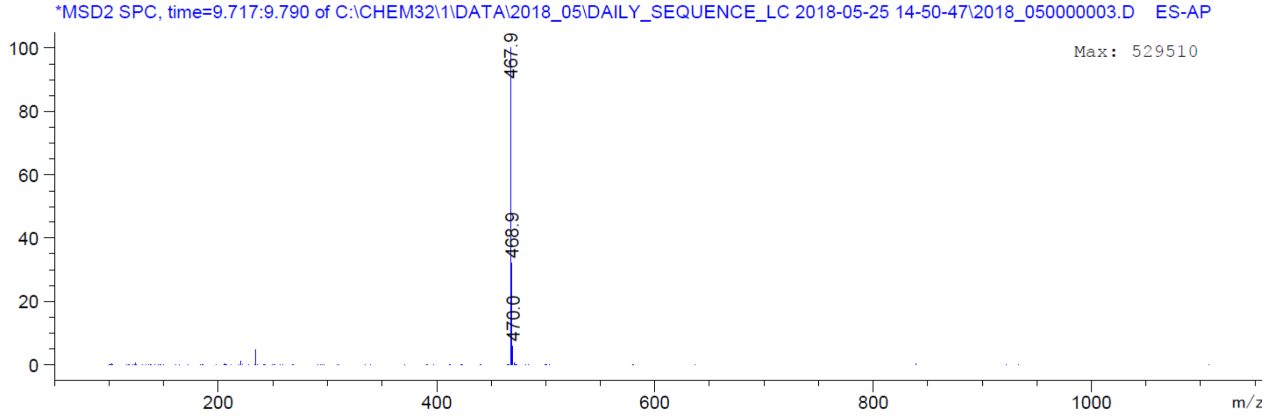

*MSD2 SPC, time=9.717:9.790 of C:\CHEM32\1\DATA\2018_05\DAILY_SEQUENCE_LC 2018-05-25 14-50-47\2018_050000003.D ES-AP
Max: 529510
467.9
468.9
470.0
m/z

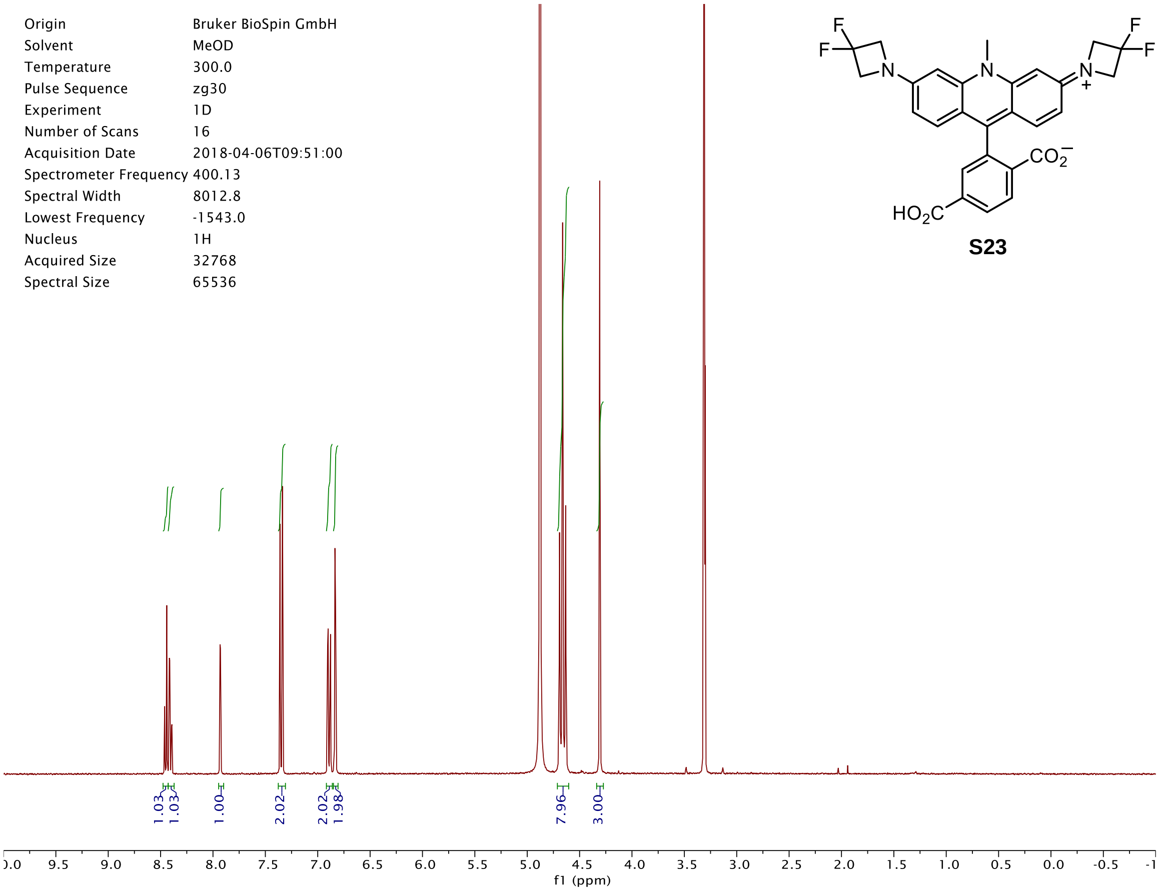
Origin Bruker BioSpin GmbH
Solvent MeOD
Temperature 300.0
Pulse Sequence zg30
Experiment 1D
Number of Scans 16
Acquisition Date 2018-04-06T09:51:00
Spectrometer Frequency 400.13
Spectral Width 8012.8
Lowest Frequency -1543.0
Nucleus 1H
Acquired Size 32768
Spectral Size 65536
1.03
1.03
1.00
2.02
2.02
1.98
7.96
3.00
f1 (ppm)

S23

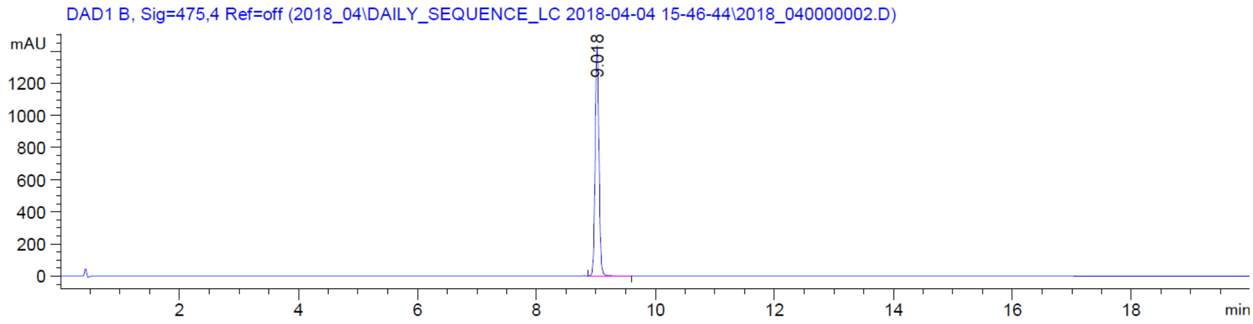
DAD1 B, Sig=475,4 Ref=off (2018_04\DAILY_SEQUENCE_LC 2018-04-04 15-46-44\2018_040000002.D)
mAU
9.018
min

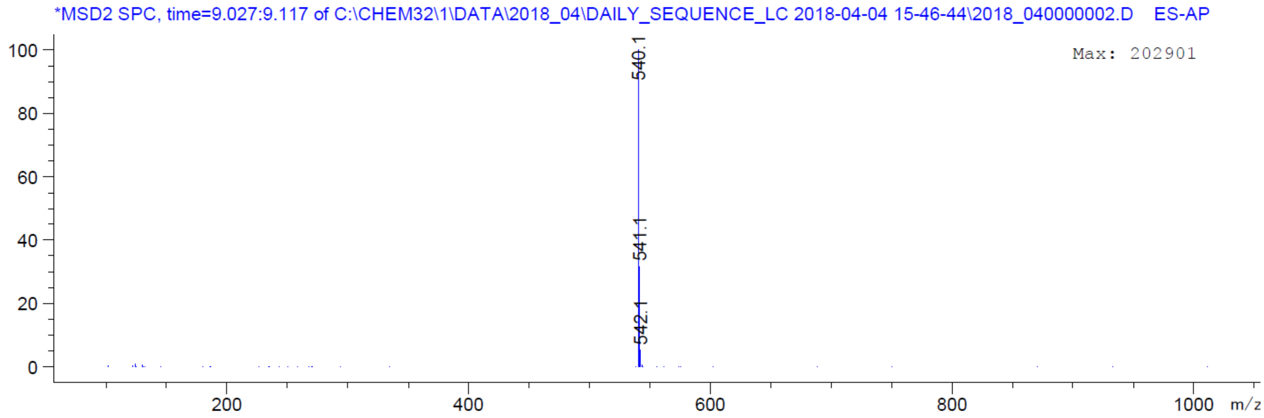
*MSD2 SPC, time=9.027:9.117 of C:\CHEM32\1\DATA\2018_04\DAILY_SEQUENCE_LC 2018-04-04 15-46-44\2018_040000002.D ES-AP
Max: 202901
540.1
541.1
542.1
m/z

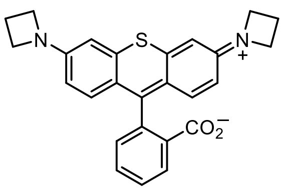


**3**

| | |
|---|---|
| Origin | Bruker BioSpin GmbH |
| Solvent | MeOD |
| Temperature | 295.6 |
| Pulse Sequence | zg30 |
| Experiment | 1D |
| Number of Scans | 16 |
| Acquisition Date | 2017-08-04T11:43:00 |
| Spectrometer Frequency | 400.13 |
| Spectral Width | 8012.8 |
| Lowest Frequency | -1543.4 |
| Nucleus | 1H |
| Acquired Size | 32768 |
| Spectral Size | 65536 |

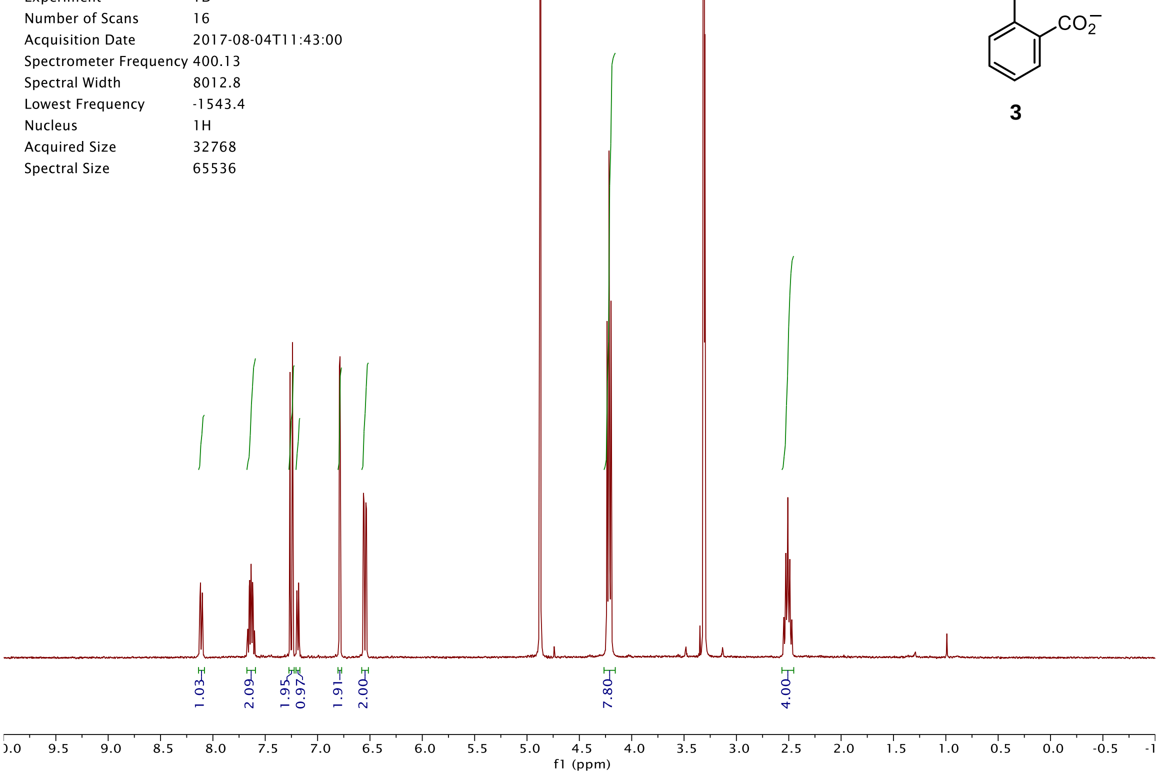


| | |
|---|---|
| Origin | Bruker BioSpin GmbH |
| Solvent | MeOD |
| Temperature | 296.3 |
| Pulse Sequence | zgpg30 |
| Experiment | 1D |
| Number of Scans | 2048 |
| Acquisition Date | 2018-03-06T20:03:00 |
| Spectrometer Frequency | 100.62 |
| Spectral Width | 24038.5 |
| Lowest Frequency | -1818.8 |
| Nucleus | 13C |
| Acquired Size | 32768 |
| Spectral Size | 65536 |

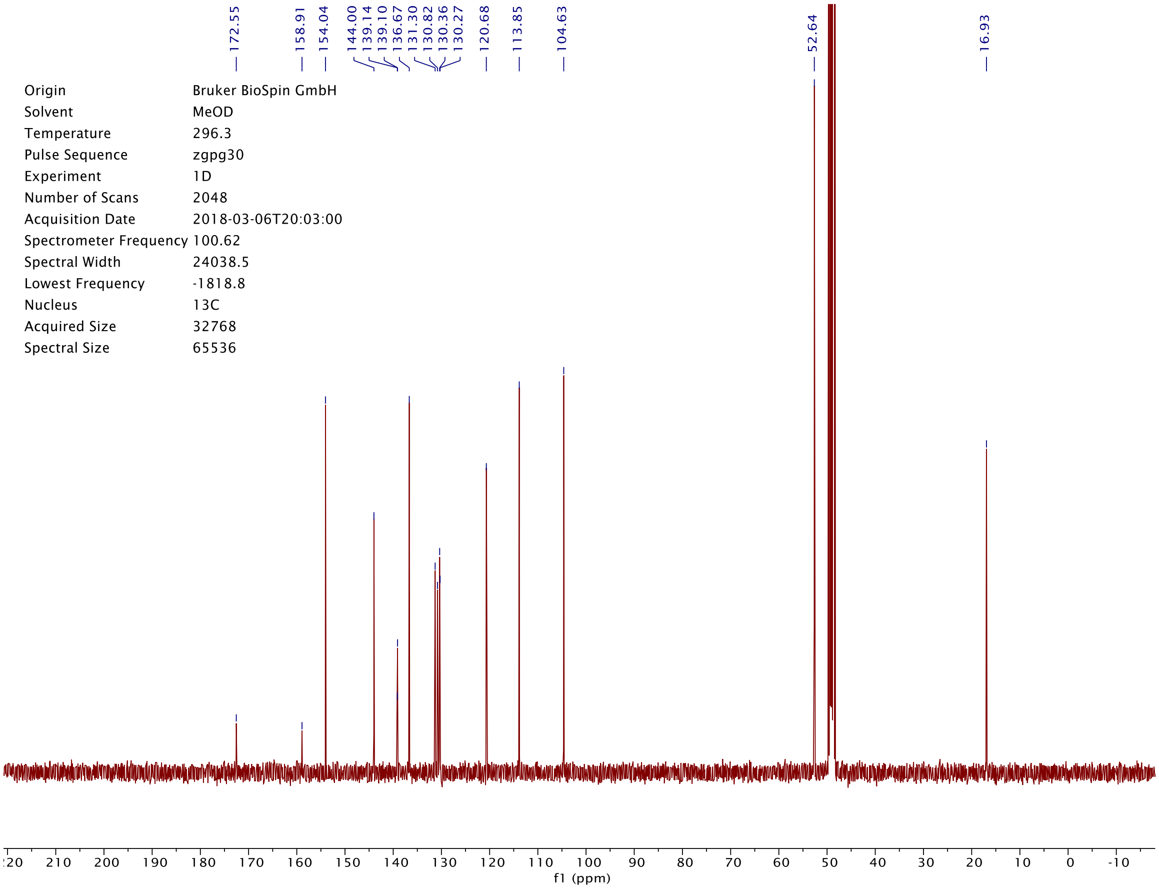

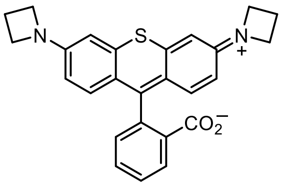

S
N
N
+
$CO_2^-$


**3**

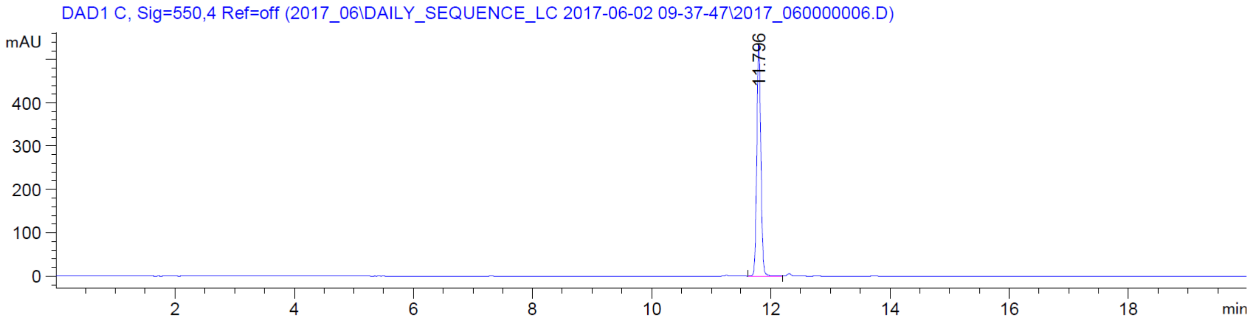

DAD1 C, Sig=550,4 Ref=off (2017_06\DAILY_SEQUENCE_LC 2017-06-02 09-37-47\2017_060000006.D)
mAU
400
300
200
100
0
11.796
2
4
6
8
10
12
14
16
18
min


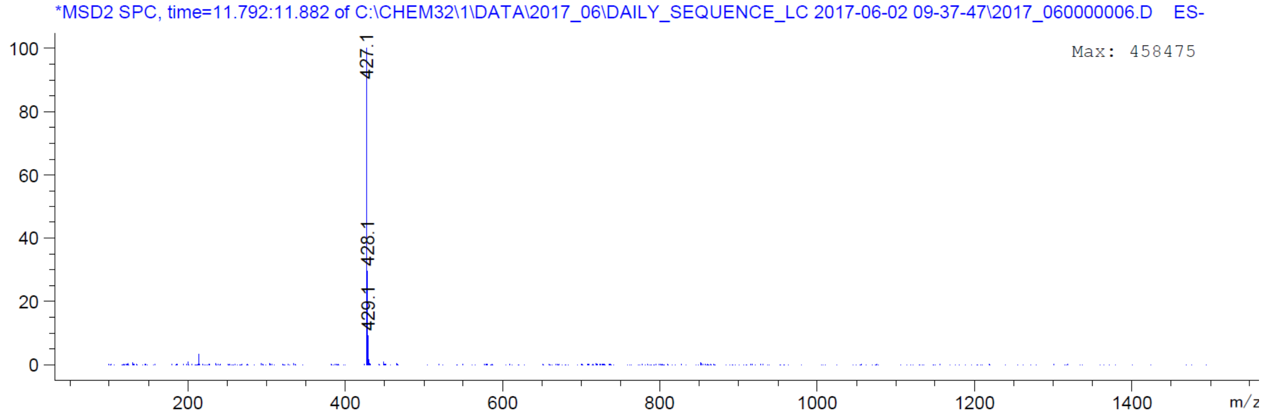

*MSD2 SPC, time=11.792:11.882 of C:\CHEM32\1\DATA\2017_06\DAILY_SEQUENCE_LC 2017-06-02 09-37-47\2017_060000006.D ES-
Max: 458475
100
80
60
40
20
0
427.1
428.1
429.1
200
400
600
800
1000
1200
1400
m/z

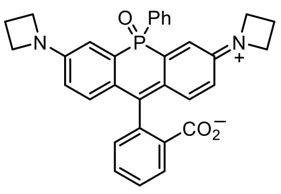


**7**

| | |
|---|---|
| Origin | Bruker BioSpin GmbH |
| Solvent | CDCl3 |
| Temperature | 300.0 |
| Pulse Sequence | zg30 |
| Experiment | 1D |
| Number of Scans | 16 |
| Acquisition Date | 2018-06-05T18:04:00 |
| Spectrometer Frequency | 400.13 |
| Spectral Width | 8012.8 |
| Lowest Frequency | -1542.6 |
| Nucleus | 1H |
| Acquired Size | 32768 |
| Spectral Size | 65536 |

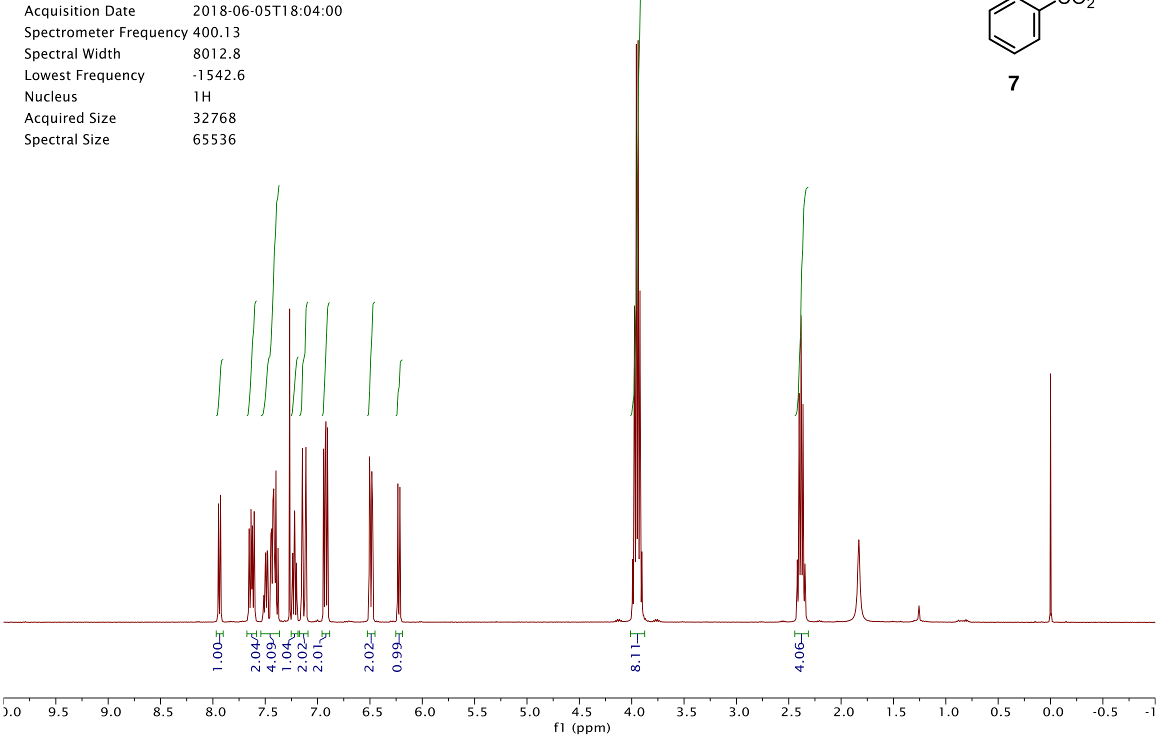


| | |
|---|---|
| Origin | Bruker BioSpin GmbH |
| Solvent | CDCl3 |
| Temperature | 300.0 |
| Pulse Sequence | zgpg30 |
| Experiment | 1D |
| Number of Scans | 8192 |
| Acquisition Date | 2018-06-06T01:54:00 |
| Spectrometer Frequency | 100.62 |
| Spectral Width | 24038.5 |
| Lowest Frequency | -1958.4 |
| Nucleus | 13C |
| Acquired Size | 32768 |
| Spectral Size | 65536 |

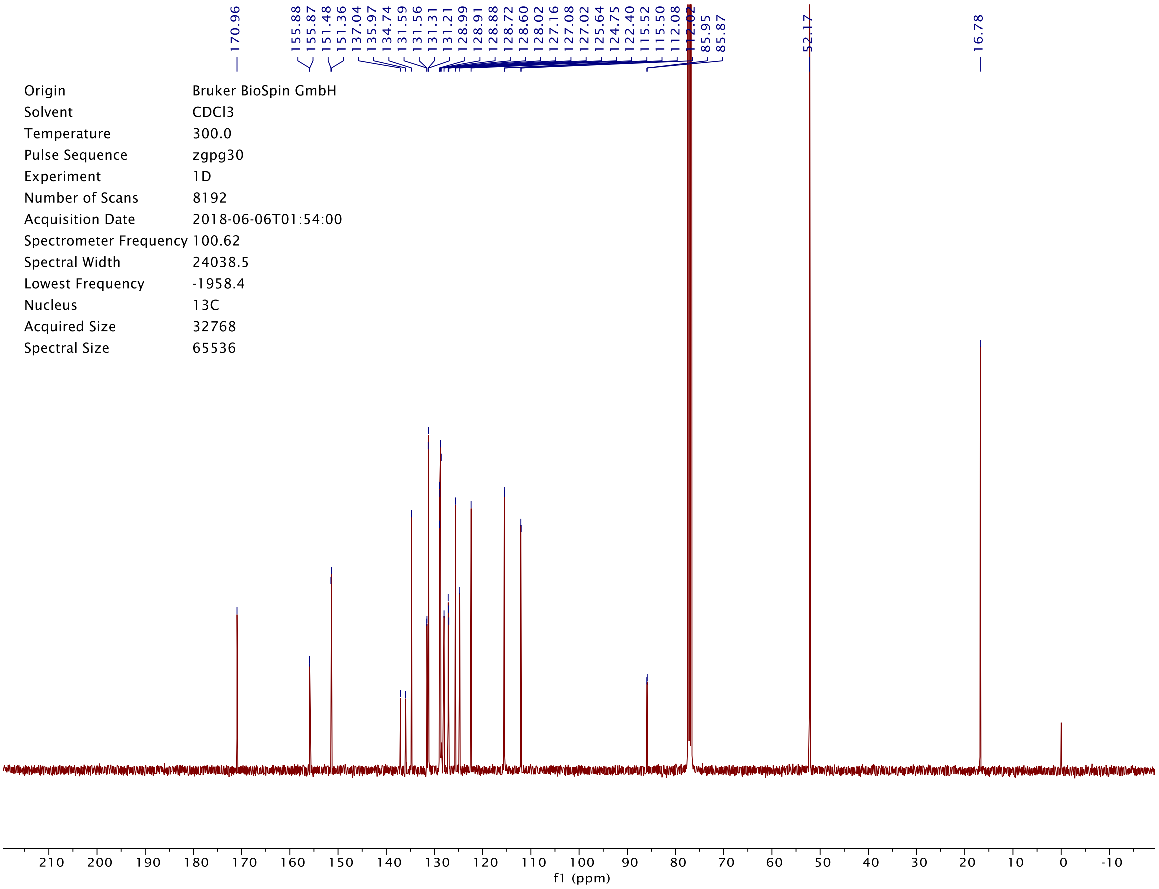

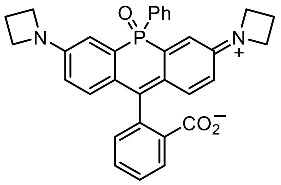
O
Ph
P
N
N
+
$CO_2^-$

**7**

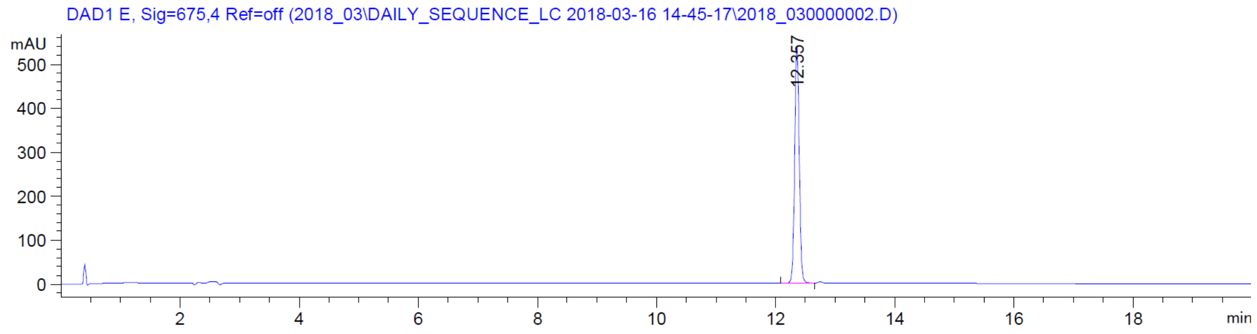
DAD1 E, Sig=675,4 Ref=off (2018_03\DAILY_SEQUENCE_LC 2018-03-16 14-45-17\2018_030000002.D)
mAU
12.357
min

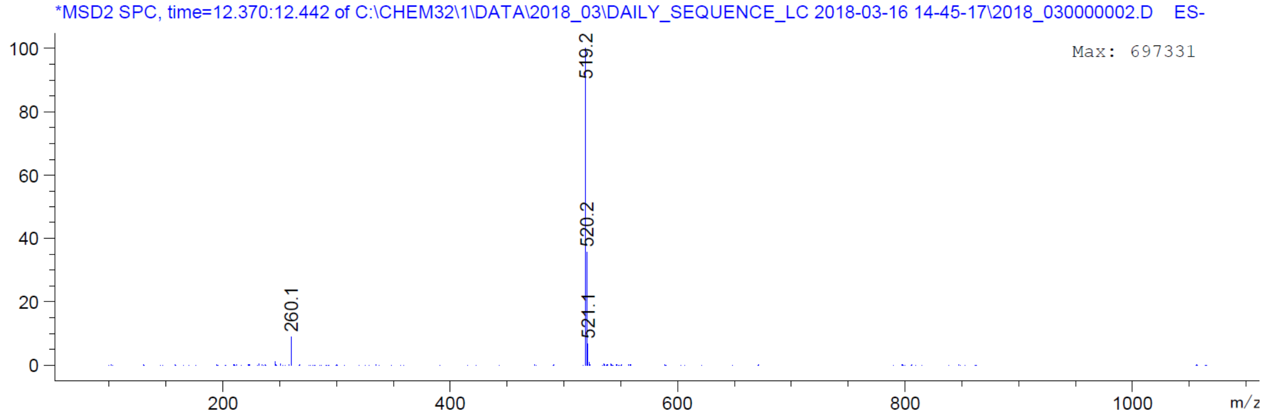
*MSD2 SPC, time=12.370:12.442 of C:\CHEM32\1\DATA\2018_03\DAILY_SEQUENCE_LC 2018-03-16 14-45-17\2018_030000002.D ES-
Max: 697331
519.2
520.2
521.1
260.1
m/z

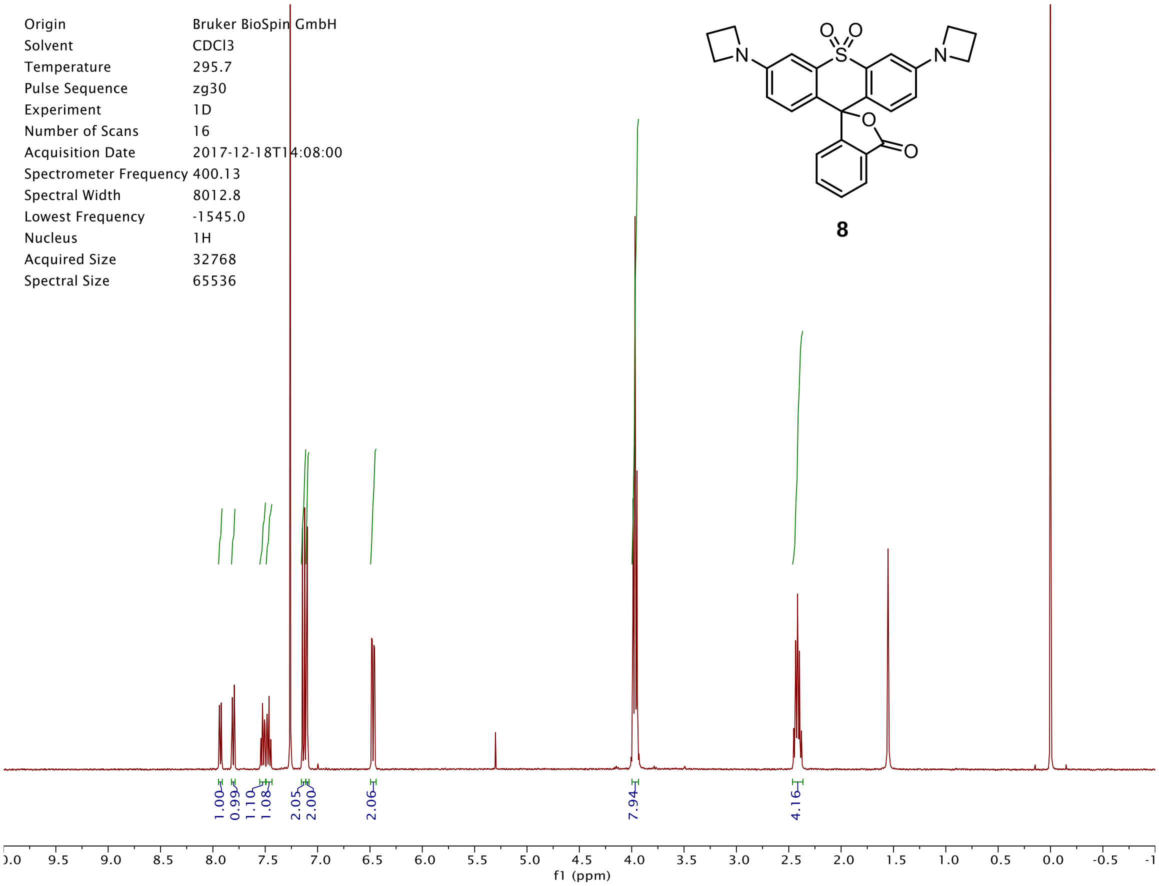

Origin Bruker BioSpin GmbH
Solvent CDCl3
Temperature 295.7
Pulse Sequence zg30
Experiment 1D
Number of Scans 16
Acquisition Date 2017-12-18T14:08:00
Spectrometer Frequency 400.13
Spectral Width 8012.8
Lowest Frequency -1545.0
Nucleus 1H
Acquired Size 32768
Spectral Size 65536
8
1.00
0.99
1.10
1.08
2.05
2.00
2.06
7.94
4.16
f1 (ppm)


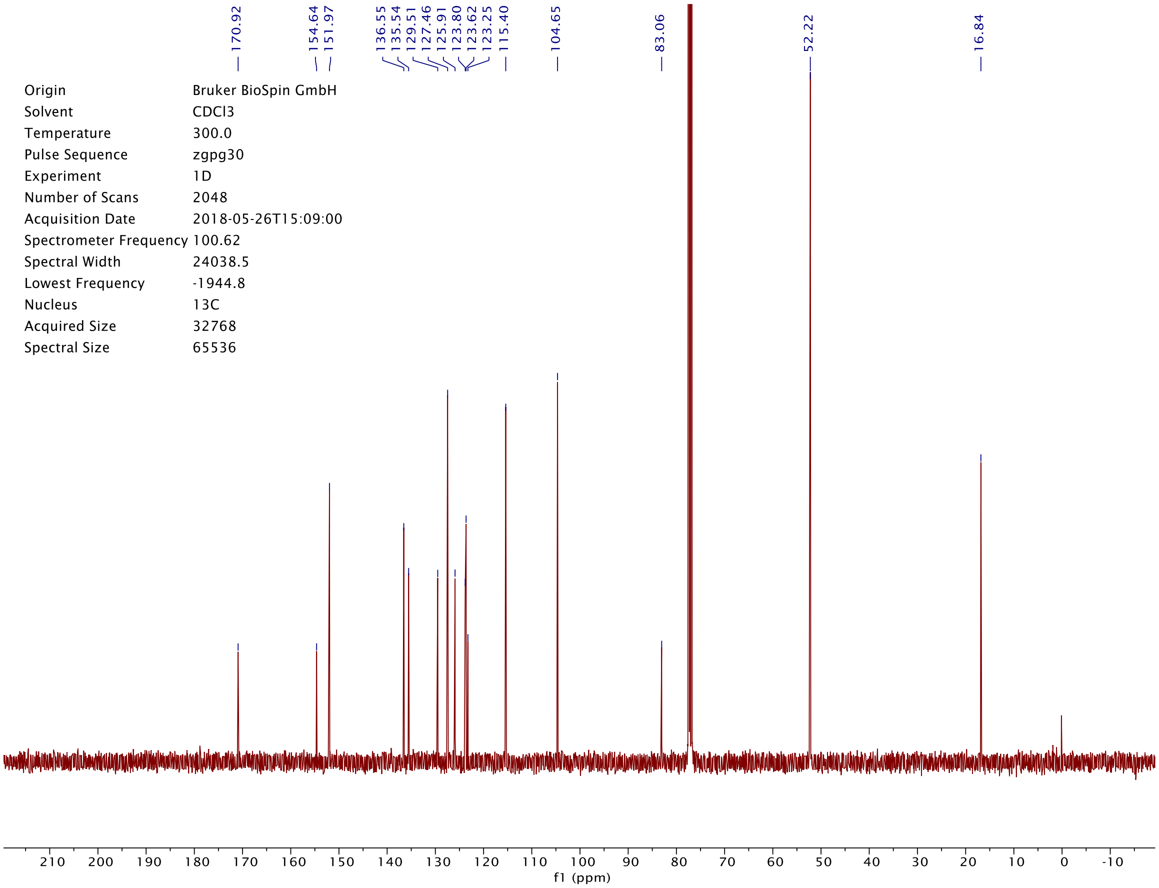

170.92
154.64
151.97
136.55
135.54
129.51
127.46
125.91
123.80
123.62
123.25
115.40
104.65
83.06
52.22
16.84
Origin Bruker BioSpin GmbH
Solvent CDCl3
Temperature 300.0
Pulse Sequence zgpg30
Experiment 1D
Number of Scans 2048
Acquisition Date 2018-05-26T15:09:00
Spectrometer Frequency 100.62
Spectral Width 24038.5
Lowest Frequency -1944.8
Nucleus 13C
Acquired Size 32768
Spectral Size 65536
f1 (ppm)

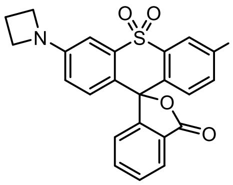

**8**

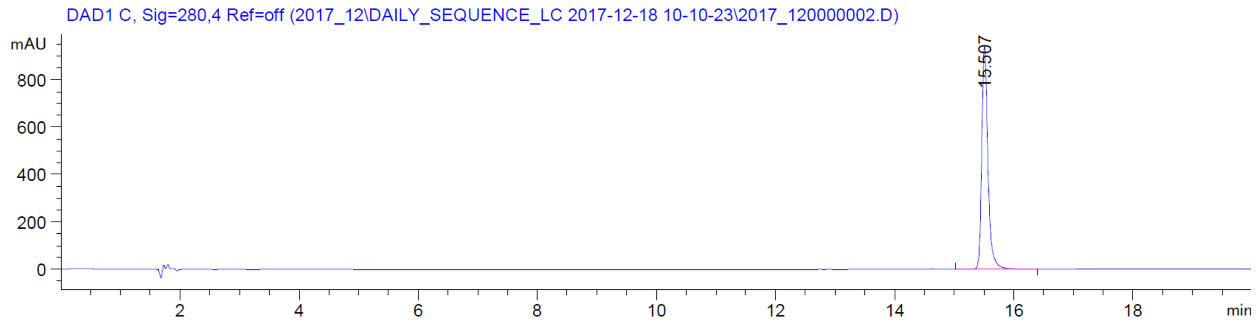

DAD1 C, Sig=280,4 Ref=off (2017_12\DAILY_SEQUENCE_LC 2017-12-18 10-10-23\2017_120000002.D)
mAU
15.507
min


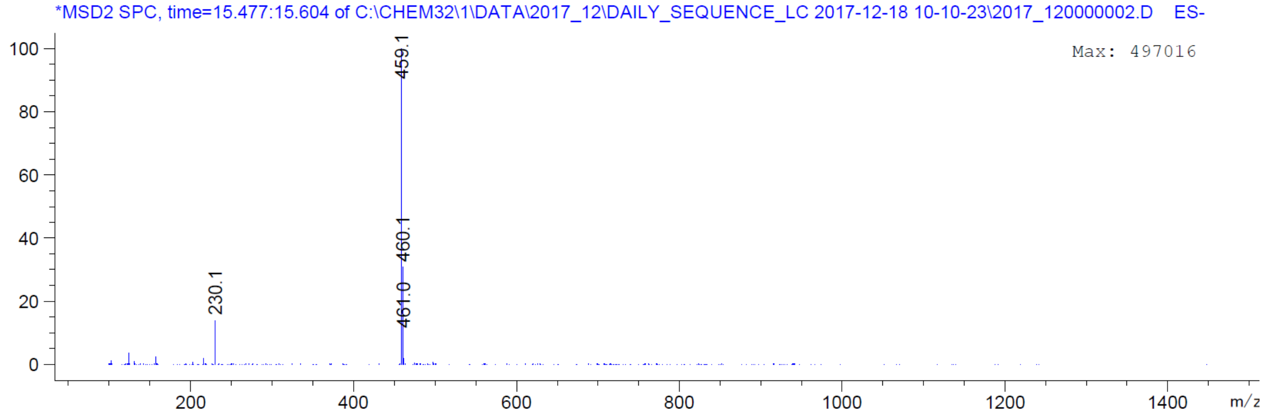

*MSD2 SPC, time=15.477:15.604 of C:\CHEM32\1\DATA\2017_12\DAILY_SEQUENCE_LC 2017-12-18 10-10-23\2017_120000002.D ES-
Max: 497016
459.1
460.1
461.0
230.1
m/z

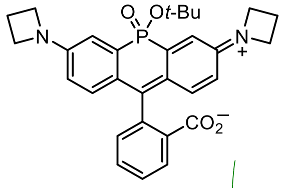
O
Ot-Bu
P
N
N
+
$CO_2^-$

**S48**

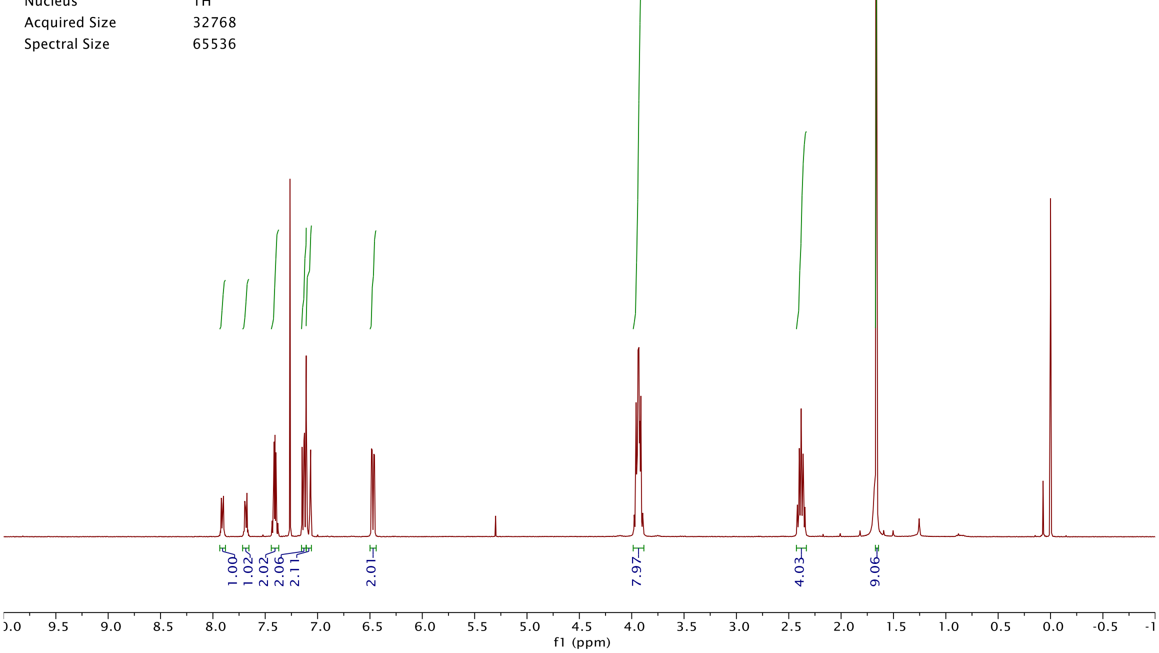
Origin Bruker BioSpin GmbH
Solvent CDCl3
Temperature 295.5
Pulse Sequence zg30
Experiment 1D
Number of Scans 16
Acquisition Date 2018-03-12T19:05:00
Spectrometer Frequency 400.13
Spectral Width 8012.8
Lowest Frequency -1544.2
Nucleus 1H
Acquired Size 32768
Spectral Size 65536
1.00
1.02
2.02
2.06
2.11
2.01
7.97
4.03
9.06
f1 (ppm)

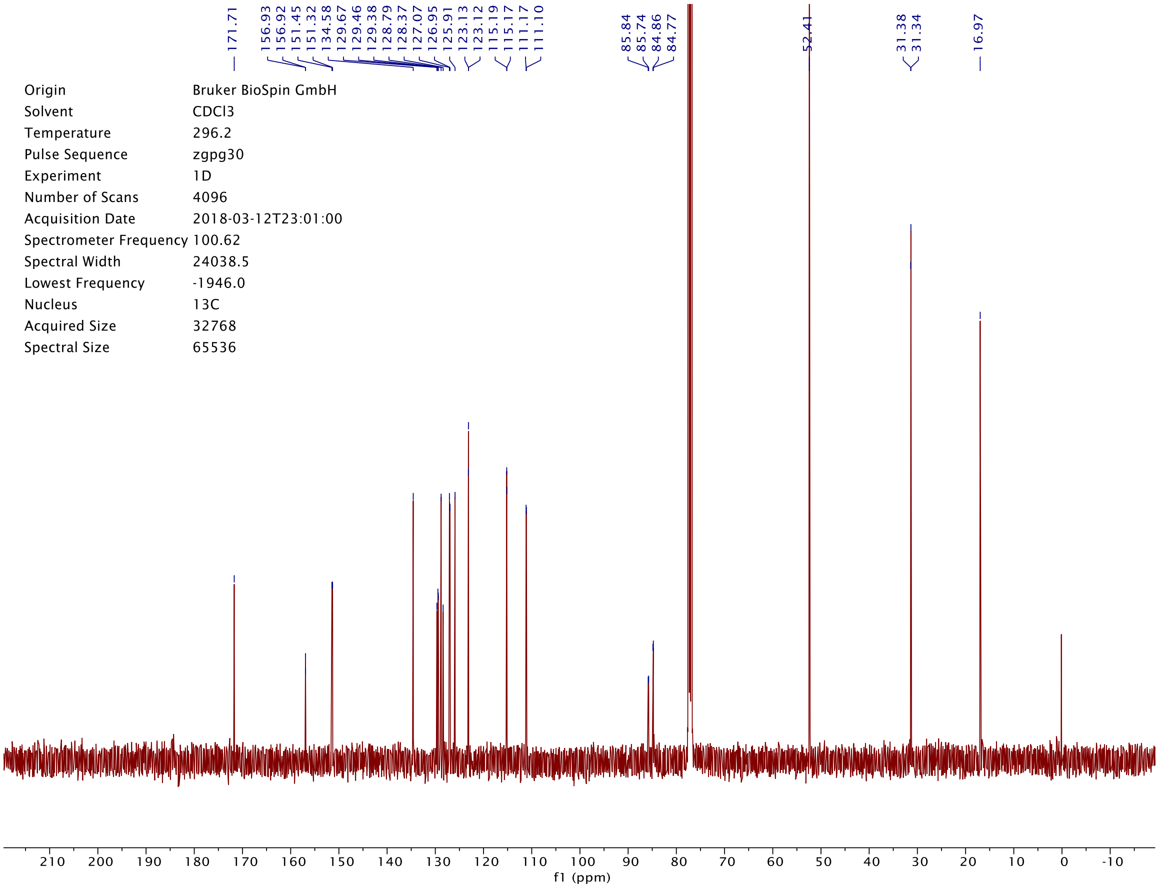
171.71
156.93
156.92
151.45
151.32
134.58
129.67
129.46
129.38
128.79
128.37
127.07
126.95
125.91
123.13
123.12
115.19
115.17
111.17
111.10
85.84
85.74
84.86
84.77
52.41
31.38
31.34
16.97
Origin Bruker BioSpin GmbH
Solvent CDCl3
Temperature 296.2
Pulse Sequence zgpg30
Experiment 1D
Number of Scans 4096
Acquisition Date 2018-03-12T23:01:00
Spectrometer Frequency 100.62
Spectral Width 24038.5
Lowest Frequency -1946.0
Nucleus 13C
Acquired Size 32768
Spectral Size 65536
f1 (ppm)

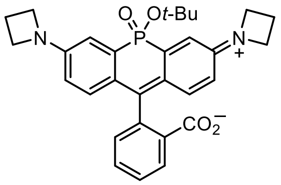

O
Ot-Bu
P
N
N
+
$CO_2^-$


**S48**

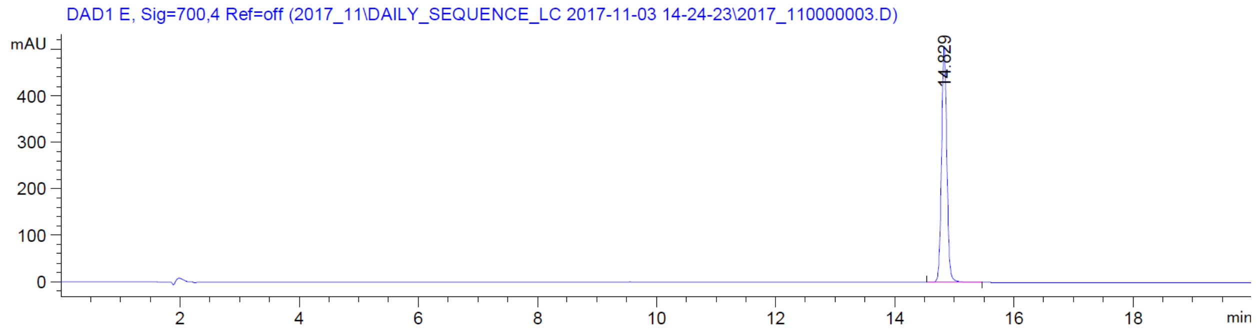

DAD1 E, Sig=700,4 Ref=off (2017_11\DAILY_SEQUENCE_LC 2017-11-03 14-24-23\2017_110000003.D)
mAU
400
300
200
100
0
14.829
2
4
6
8
10
12
14
16
18
min


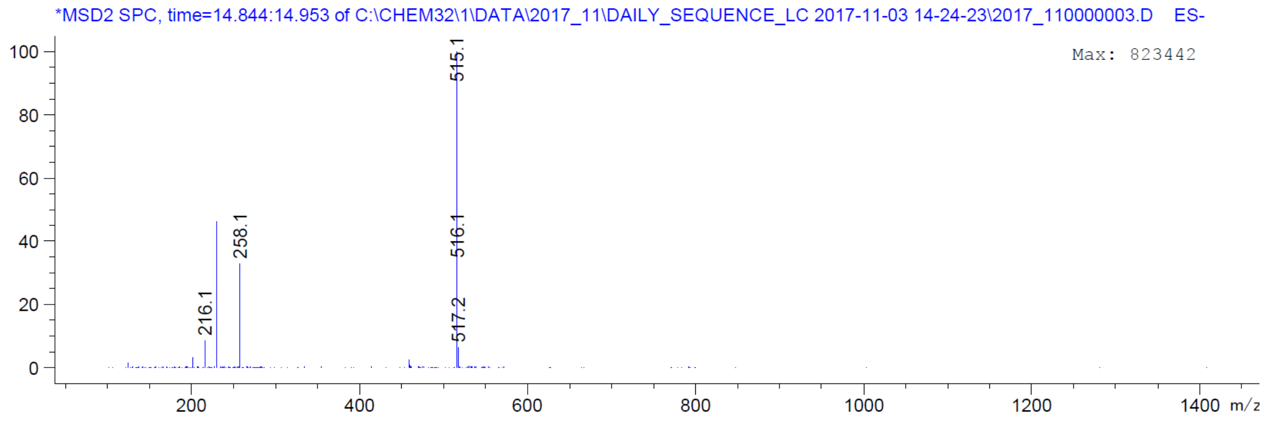

*MSD2 SPC, time=14.844:14.953 of C:\CHEM32\1\DATA\2017_11\DAILY_SEQUENCE_LC 2017-11-03 14-24-23\2017_110000003.D ES-
Max: 823442
100
80
60
40
20
0
515.1
516.1
517.2
258.1
216.1
200
400
600
800
1000
1200
1400
m/z

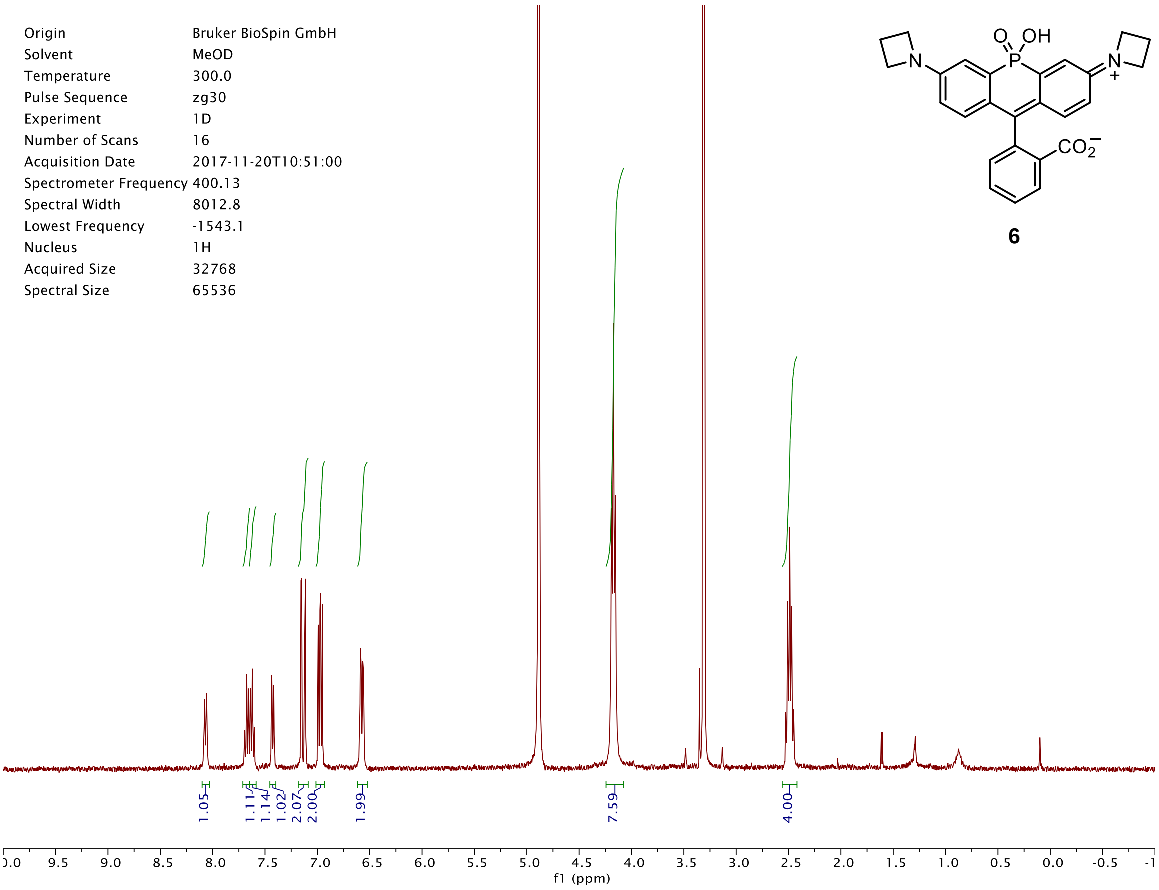
Origin Bruker BioSpin GmbH
Solvent MeOD
Temperature 300.0
Pulse Sequence zg30
Experiment 1D
Number of Scans 16
Acquisition Date 2017-11-20T10:51:00
Spectrometer Frequency 400.13
Spectral Width 8012.8
Lowest Frequency -1543.1
Nucleus 1H
Acquired Size 32768
Spectral Size 65536
1.05
1.11
1.14
1.02
2.07
2.00
1.99
7.59
4.00
f1 (ppm)
O
OH
P
N
N
+
$CO_2^-$

**6**

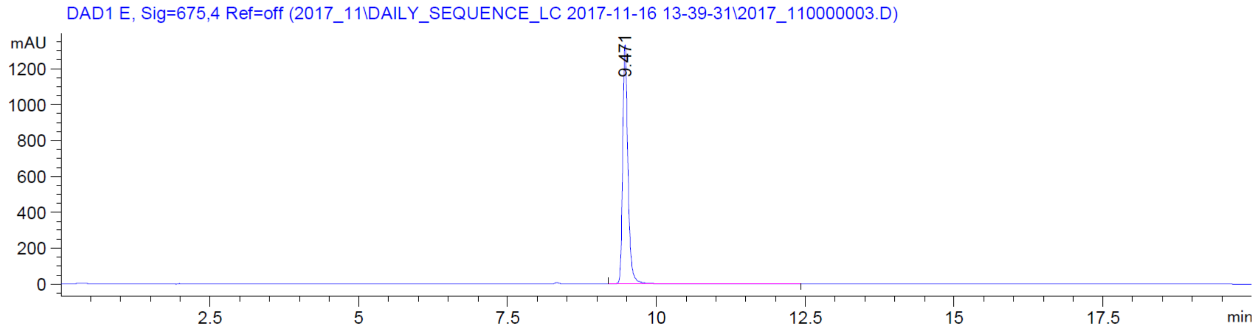
DAD1 E, Sig=675,4 Ref=off (2017_11\DAILY_SEQUENCE_LC 2017-11-16 13-39-31\2017_110000003.D)
mAU
9.471
min

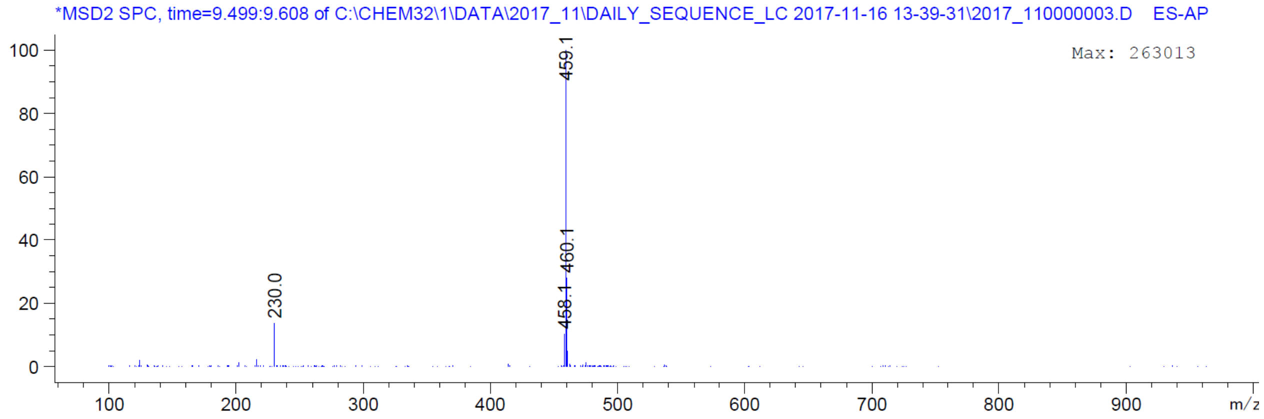
*MSD2 SPC, time=9.499:9.608 of C:\CHEM32\1\DATA\2017_11\DAILY_SEQUENCE_LC 2017-11-16 13-39-31\2017_110000003.D ES-AP
Max: 263013
459.1
460.1
458.1
230.0
m/z

| | |
|---|---|
| Origin | Bruker BioSpin GmbH |
| Solvent | CDCl3 |
| Temperature | 300.0 |
| Pulse Sequence | zg30 |
| Experiment | 1D |
| Number of Scans | 16 |
| Acquisition Date | 2017-11-20T09:57:00 |
| Spectrometer Frequency | 400.13 |
| Spectral Width | 8012.8 |
| Lowest Frequency | -1545.6 |
| Nucleus | 1H |
| Acquired Size | 32768 |
| Spectral Size | 65536 |

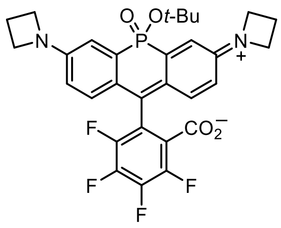


**S49**

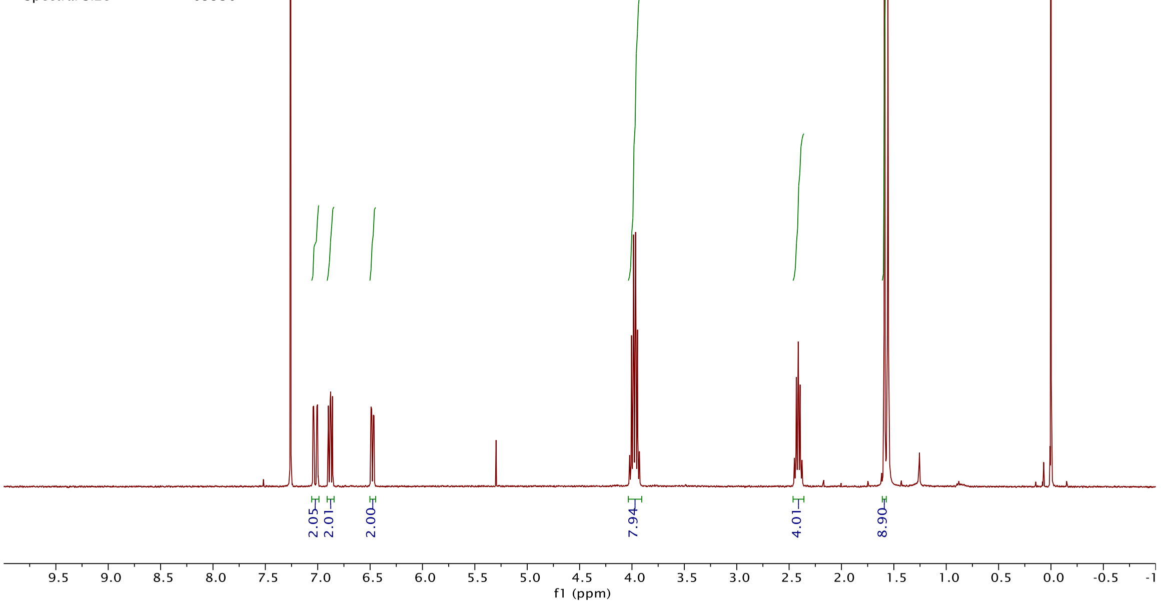


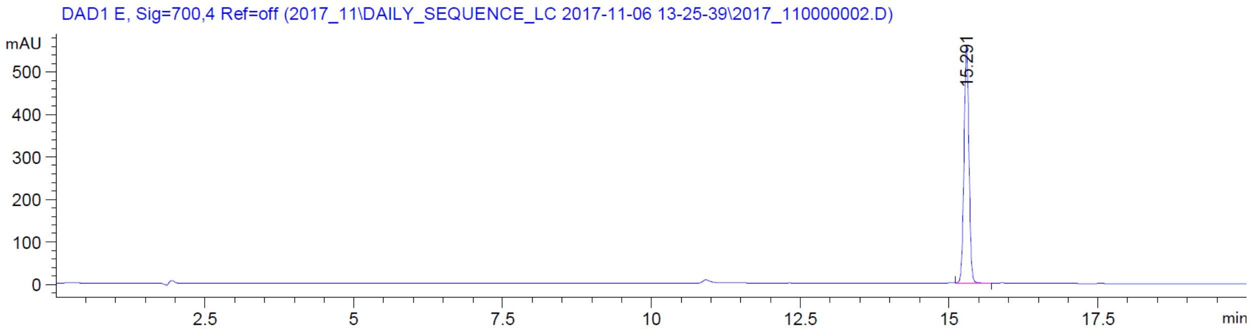


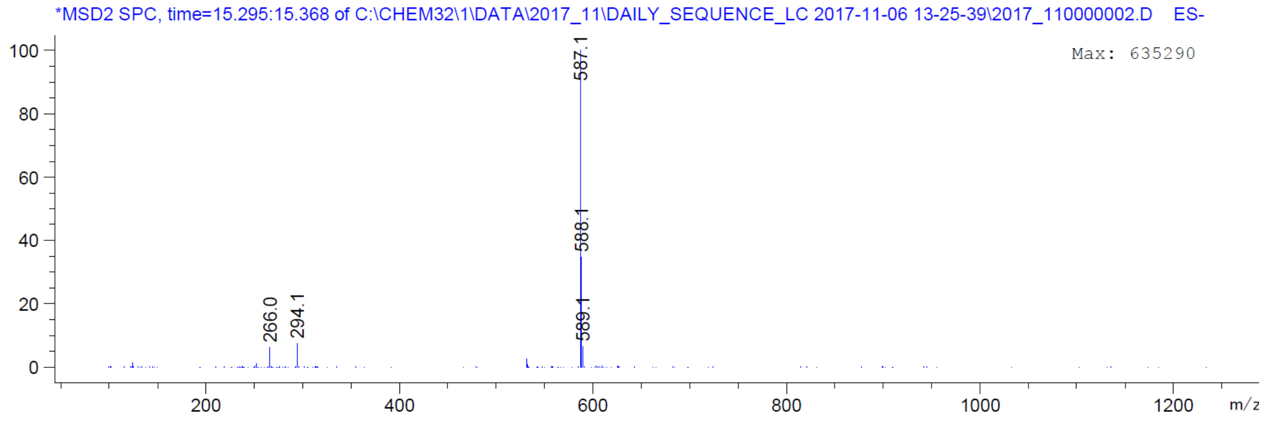

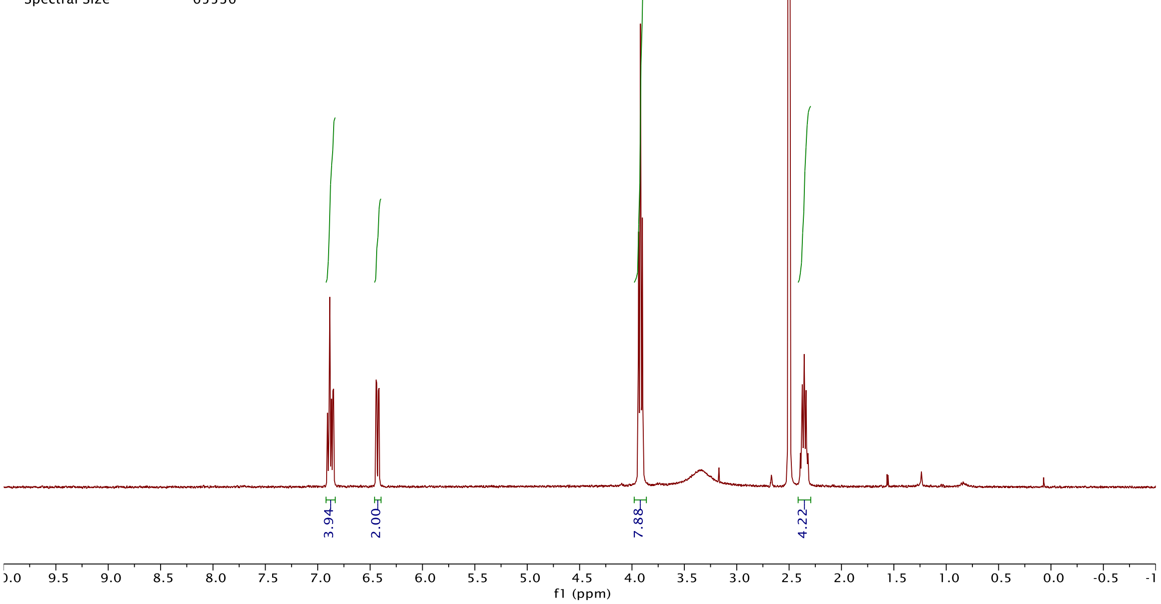
Origin Bruker BioSpin GmbH
Solvent DMSO
Temperature 300.0
Pulse Sequence zg30
Experiment 1D
Number of Scans 16
Acquisition Date 2017-11-20T10:10:00
Spectrometer Frequency 400.13
Spectral Width 8012.8
Lowest Frequency -1538.8
Nucleus 1H
Acquired Size 32768
Spectral Size 65536
3.94
2.00
7.88
4.22
f1 (ppm)

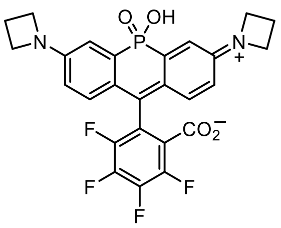
O
OH
P
N
N
+
F
CO2−
F
F
F

**16**

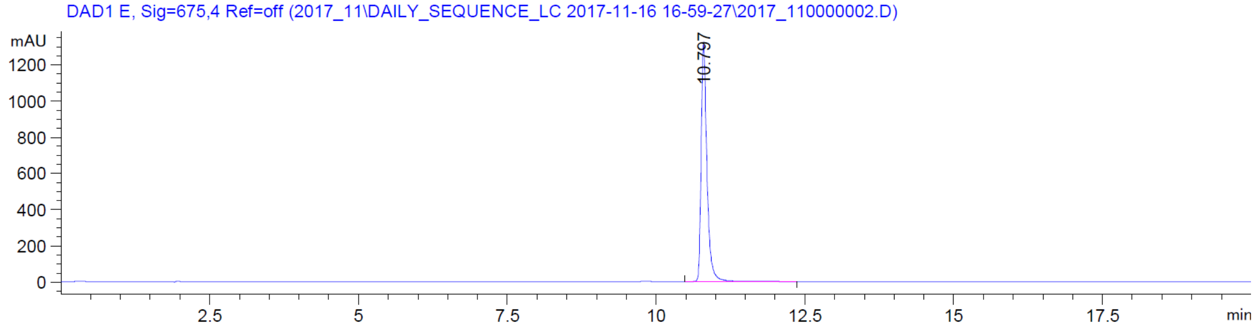
DAD1 E, Sig=675,4 Ref=off (2017_11\DAILY_SEQUENCE_LC 2017-11-16 16-59-27\2017_110000002.D)
mAU
10.797
min

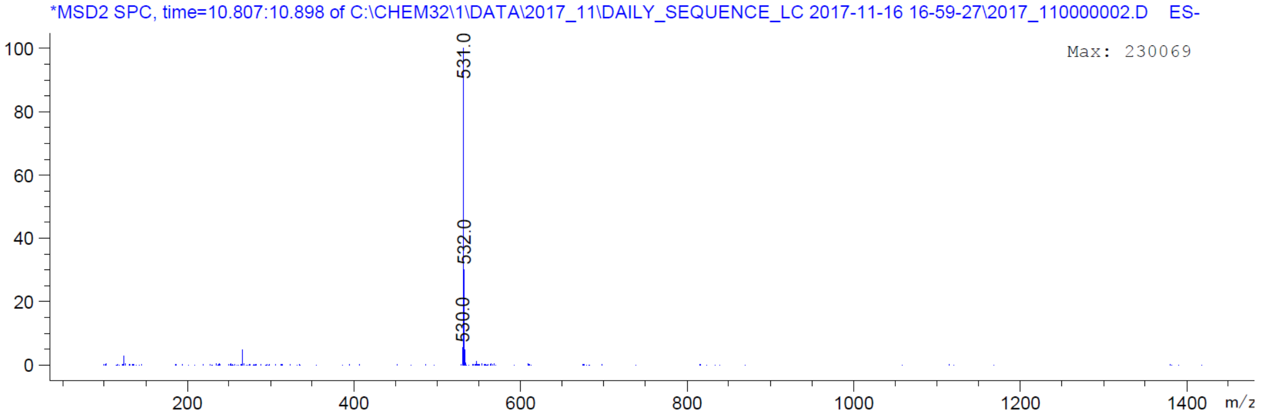
*MSD2 SPC, time=10.807:10.898 of C:\CHEM32\1\DATA\2017_11\DAILY_SEQUENCE_LC 2017-11-16 16-59-27\2017_110000002.D ES-
Max: 230069
531.0
532.0
530.0
m/z

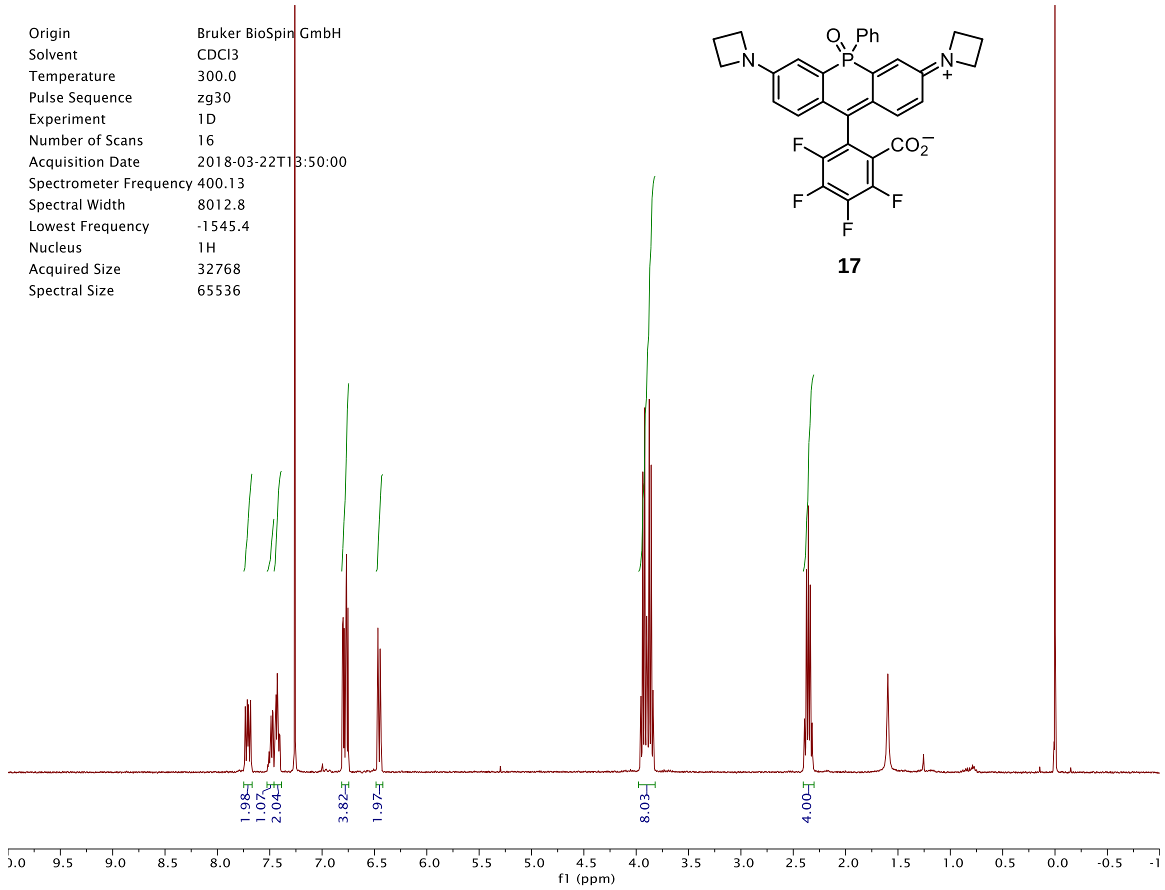
Origin Bruker BioSpin GmbH
Solvent CDCl3
Temperature 300.0
Pulse Sequence zg30
Experiment 1D
Number of Scans 16
Acquisition Date 2018-03-22T13:50:00
Spectrometer Frequency 400.13
Spectral Width 8012.8
Lowest Frequency -1545.4
Nucleus 1H
Acquired Size 32768
Spectral Size 65536
O Ph P N N + F CO2− F F F
17
1.98 1.07 2.04 3.82 1.97 8.03 4.00
f1 (ppm)

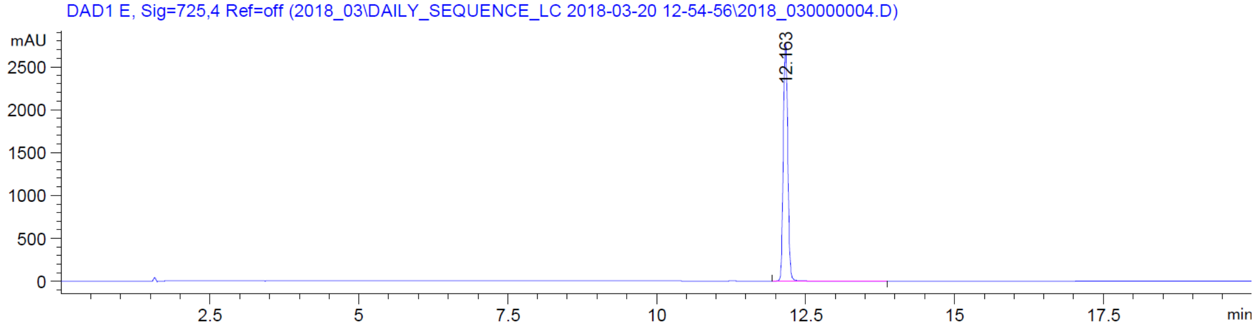
DAD1 E, Sig=725,4 Ref=off (2018_03\DAILY_SEQUENCE_LC 2018-03-20 12-54-56\2018_030000004.D)
mAU
12.163
min

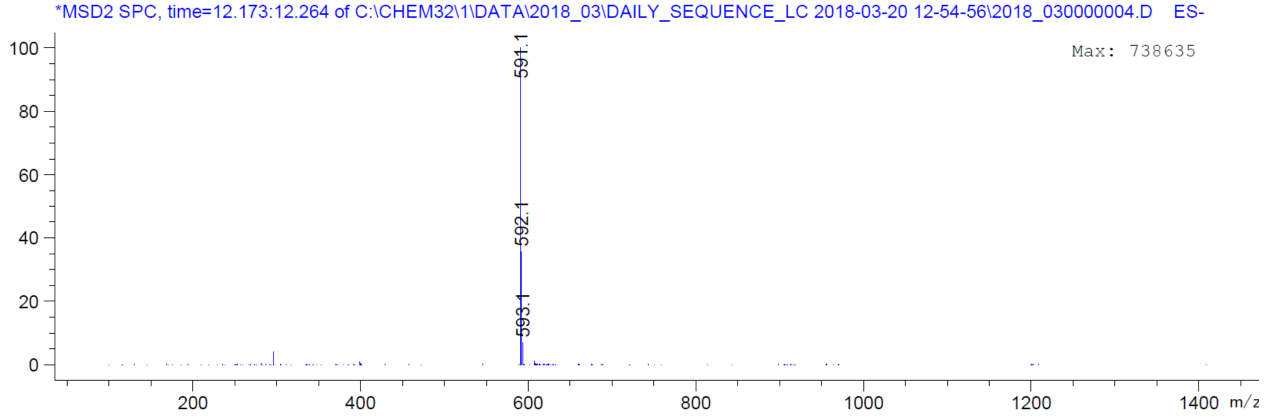
*MSD2 SPC, time=12.173:12.264 of C:\CHEM32\1\DATA\2018_03\DAILY_SEQUENCE_LC 2018-03-20 12-54-56\2018_030000004.D ES-
Max: 738635
591.1
592.1
593.1
m/z

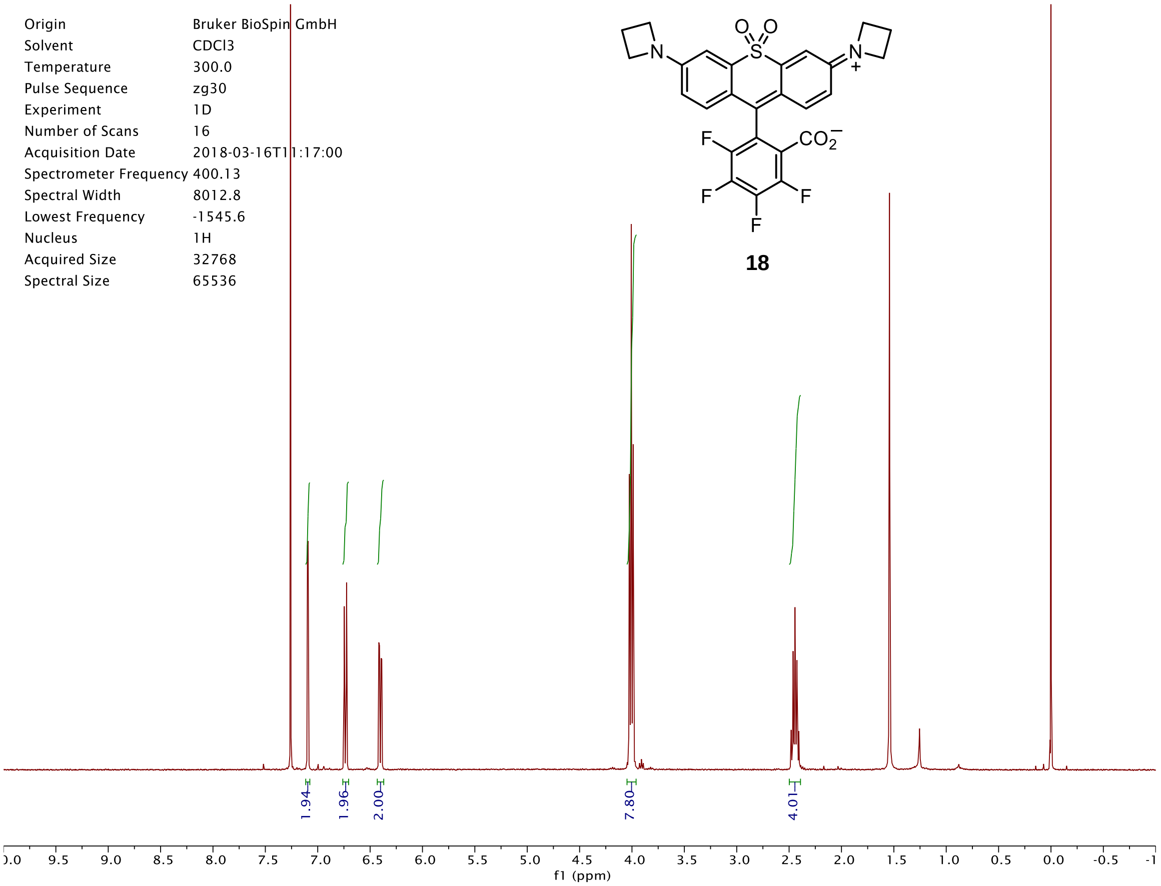
Origin Bruker BioSpin GmbH
Solvent CDCl3
Temperature 300.0
Pulse Sequence zg30
Experiment 1D
Number of Scans 16
Acquisition Date 2018-03-16T11:17:00
Spectrometer Frequency 400.13
Spectral Width 8012.8
Lowest Frequency -1545.6
Nucleus 1H
Acquired Size 32768
Spectral Size 65536
1.94
1.96
2.00
7.80
4.01
f1 (ppm)

**18**

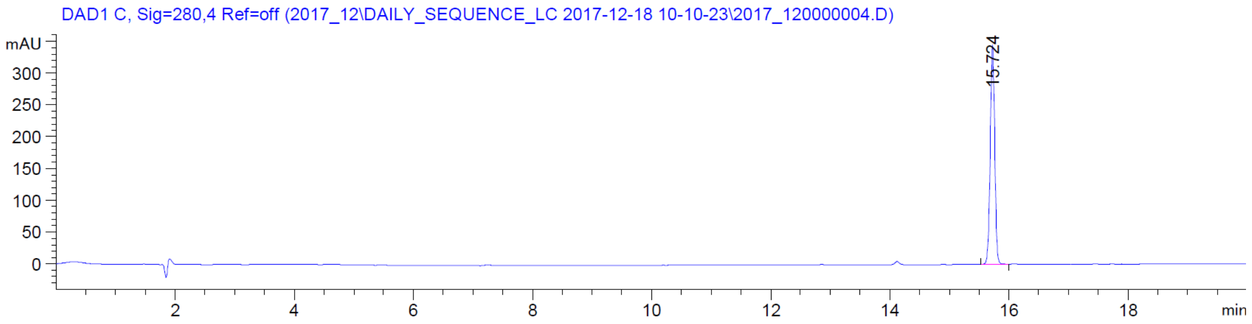
DAD1 C, Sig=280,4 Ref=off (2017_12\DAILY_SEQUENCE_LC 2017-12-18 10-10-23\2017_120000004.D)
mAU
15.724
min

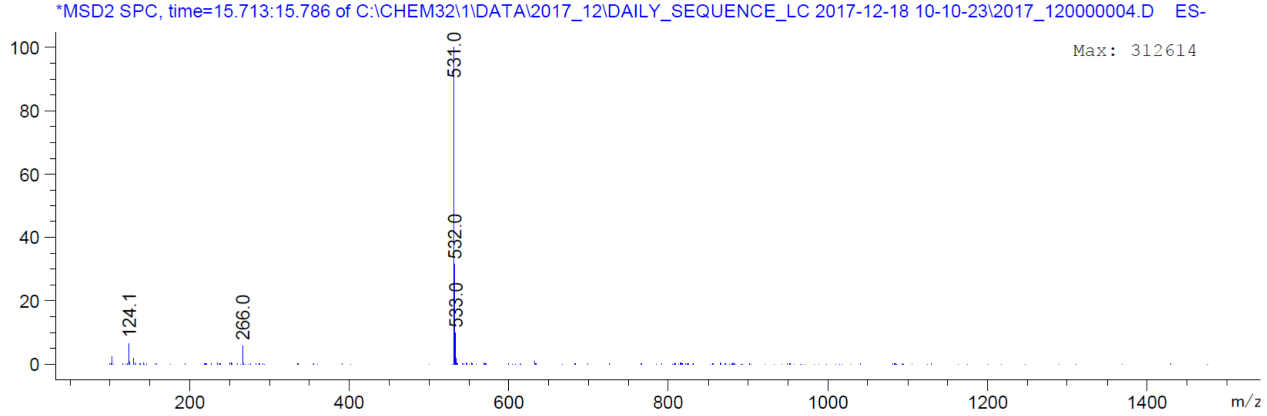
*MSD2 SPC, time=15.713:15.786 of C:\CHEM32\1\DATA\2017_12\DAILY_SEQUENCE_LC 2017-12-18 10-10-23\2017_120000004.D ES-
Max: 312614
531.0
532.0
533.0
124.1
266.0
m/z

| | |
|---|---|
| Origin | Bruker BioSpin GmbH |
| Solvent | MeOD |
| Temperature | 300.0 |
| Pulse Sequence | zg30 |
| Experiment | 1D |
| Number of Scans | 16 |
| Acquisition Date | 2018-04-23T09:42:00 |
| Spectrometer Frequency | 400.13 |
| Spectral Width | 8012.8 |
| Lowest Frequency | -1543.2 |
| Nucleus | 1H |
| Acquired Size | 32768 |
| Spectral Size | 65536 |

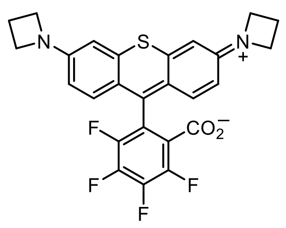

**20**

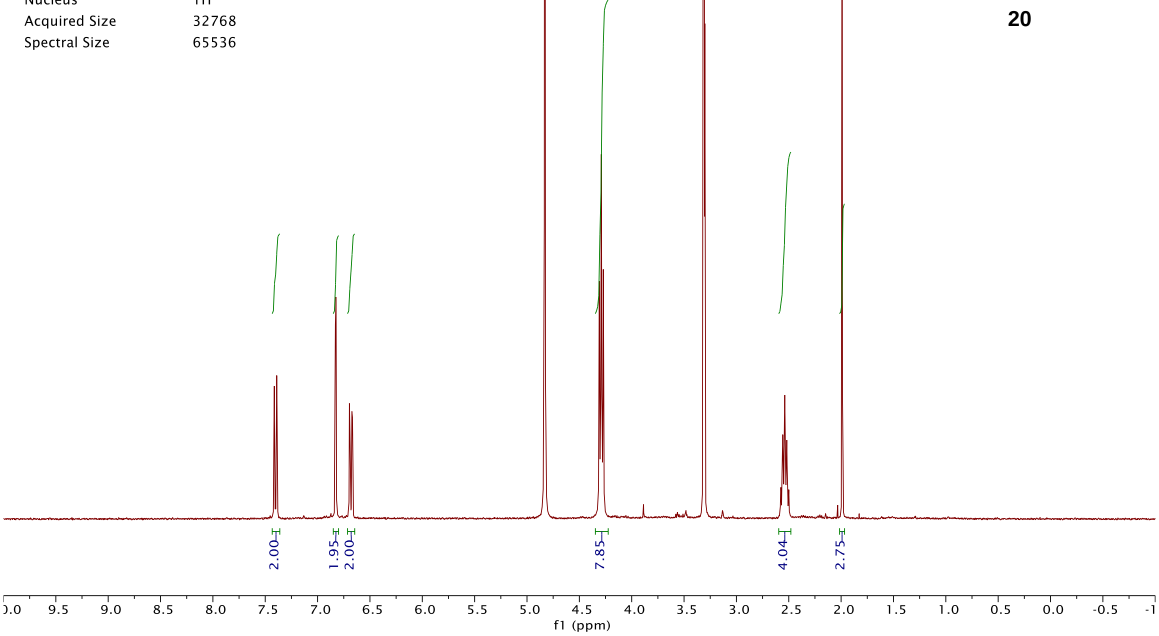


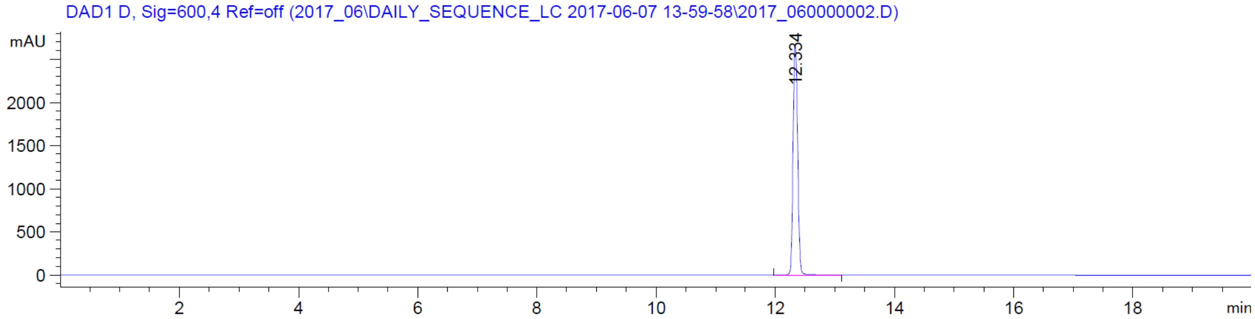


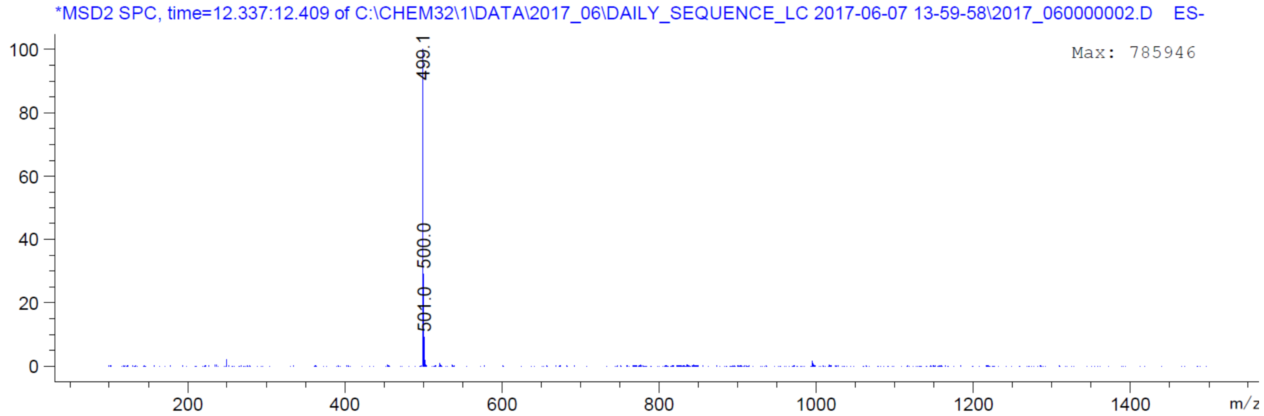

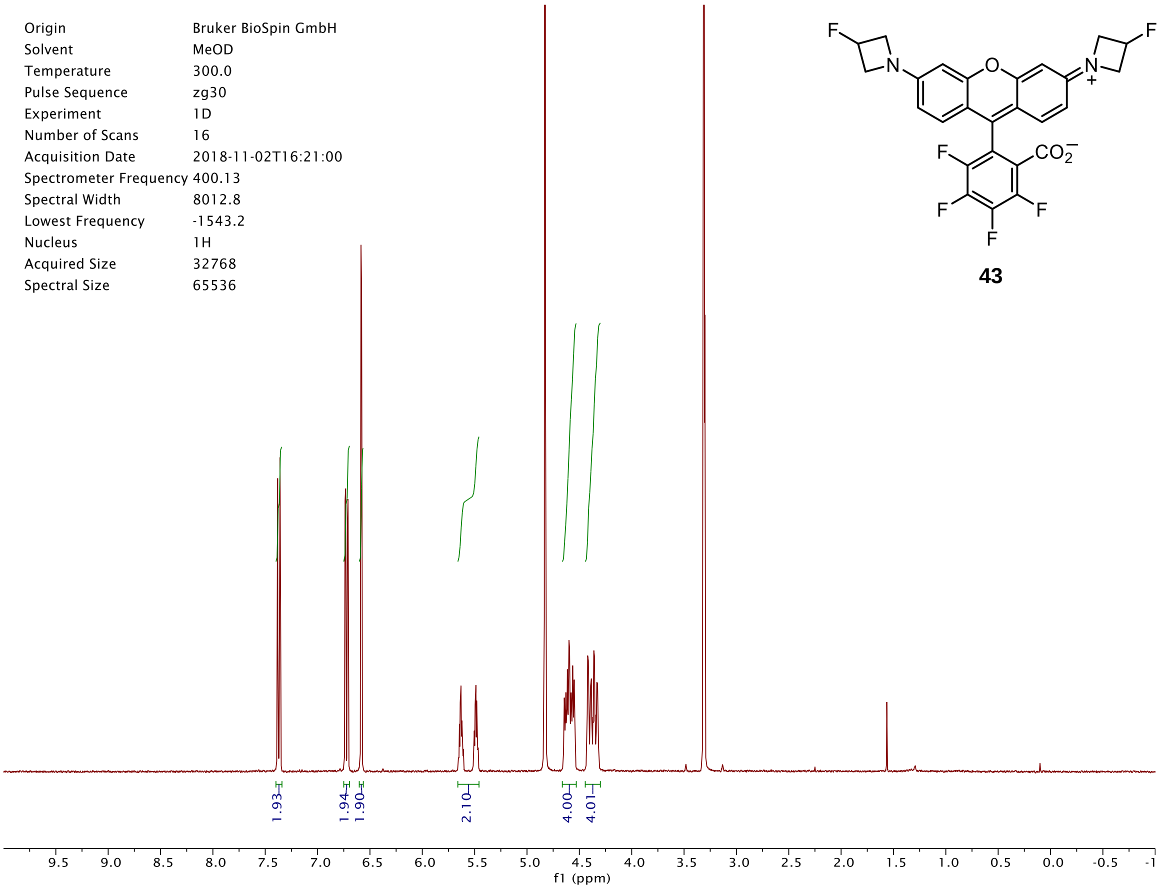
Origin Bruker BioSpin GmbH
Solvent MeOD
Temperature 300.0
Pulse Sequence zg30
Experiment 1D
Number of Scans 16
Acquisition Date 2018-11-02T16:21:00
Spectrometer Frequency 400.13
Spectral Width 8012.8
Lowest Frequency -1543.2
Nucleus 1H
Acquired Size 32768
Spectral Size 65536
1.93
1.94
1.90
2.10
4.00
4.01
f1 (ppm)

**43**

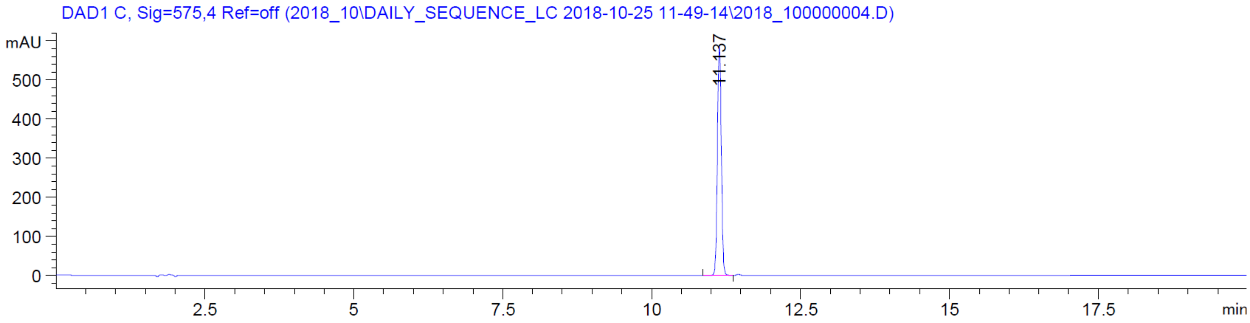
DAD1 C, Sig=575,4 Ref=off (2018_10\DAILY_SEQUENCE_LC 2018-10-25 11-49-14\2018_100000004.D)
mAU
11.137
min

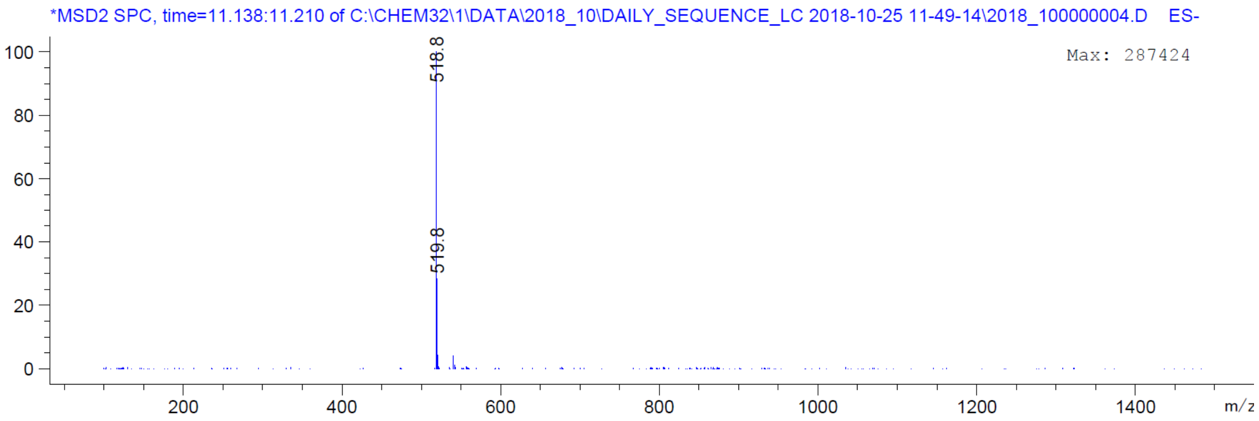
*MSD2 SPC, time=11.138:11.210 of C:\CHEM32\1\DATA\2018_10\DAILY_SEQUENCE_LC 2018-10-25 11-49-14\2018_100000004.D ES-
Max: 287424
518.8
519.8
m/z

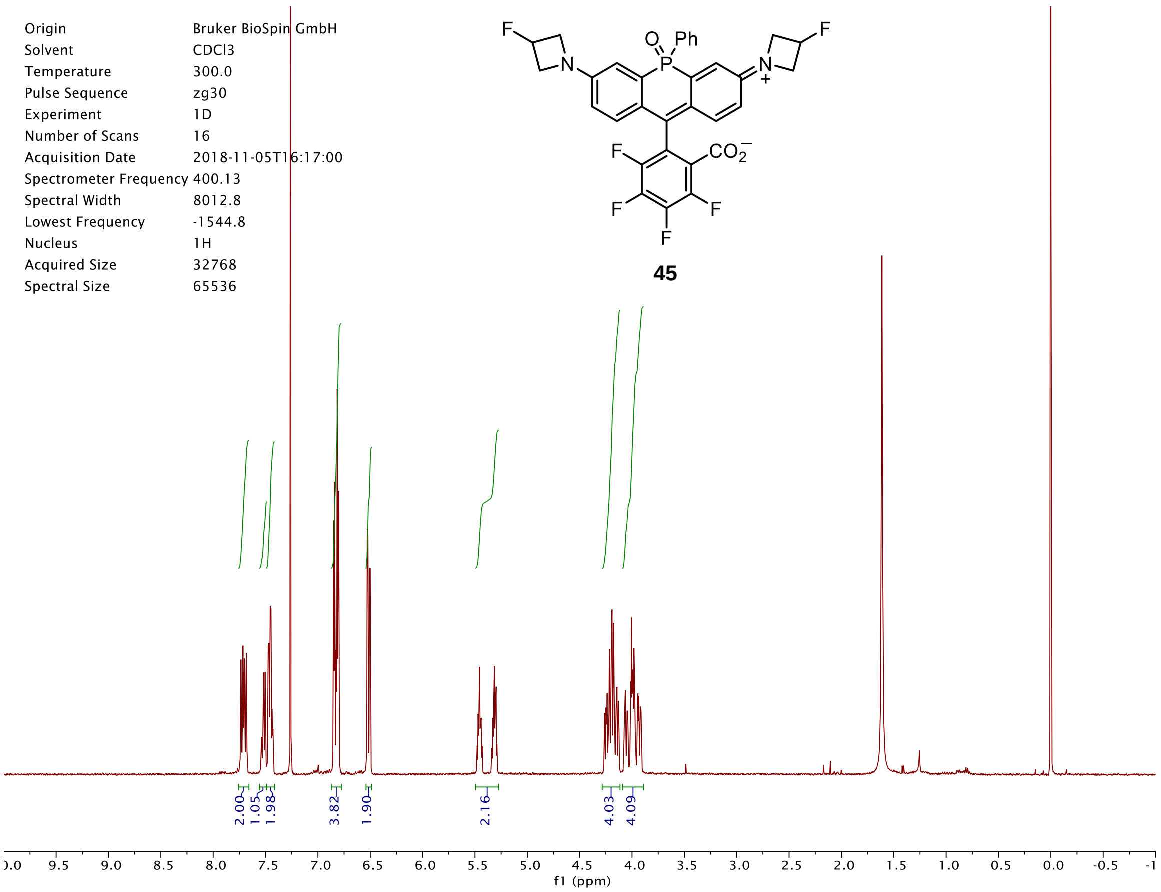
Origin Bruker BioSpin GmbH
Solvent CDCl3
Temperature 300.0
Pulse Sequence zg30
Experiment 1D
Number of Scans 16
Acquisition Date 2018-11-05T16:17:00
Spectrometer Frequency 400.13
Spectral Width 8012.8
Lowest Frequency -1544.8
Nucleus 1H
Acquired Size 32768
Spectral Size 65536
45
2.00
1.05
1.98
3.82
1.90
2.16
4.03
4.09
f1 (ppm)

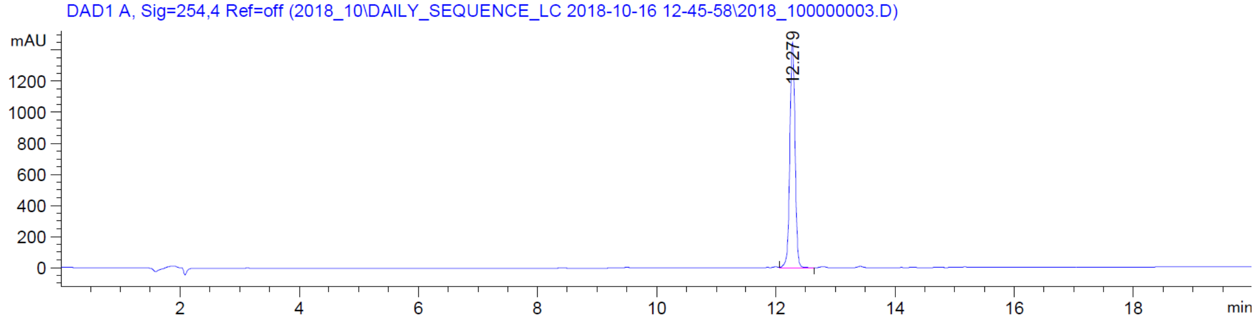
DAD1 A, Sig=254,4 Ref=off (2018_10\DAILY_SEQUENCE_LC 2018-10-16 12-45-58\2018_100000003.D)
mAU
12.279
min

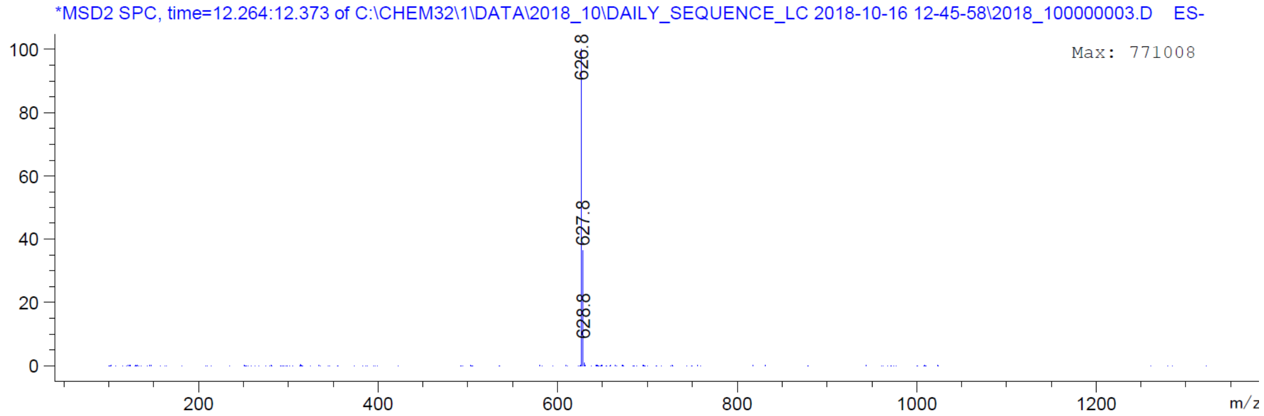
*MSD2 SPC, time=12.264:12.373 of C:\CHEM32\1\DATA\2018_10\DAILY_SEQUENCE_LC 2018-10-16 12-45-58\2018_100000003.D ES-
Max: 771008
626.8
627.8
628.8
m/z

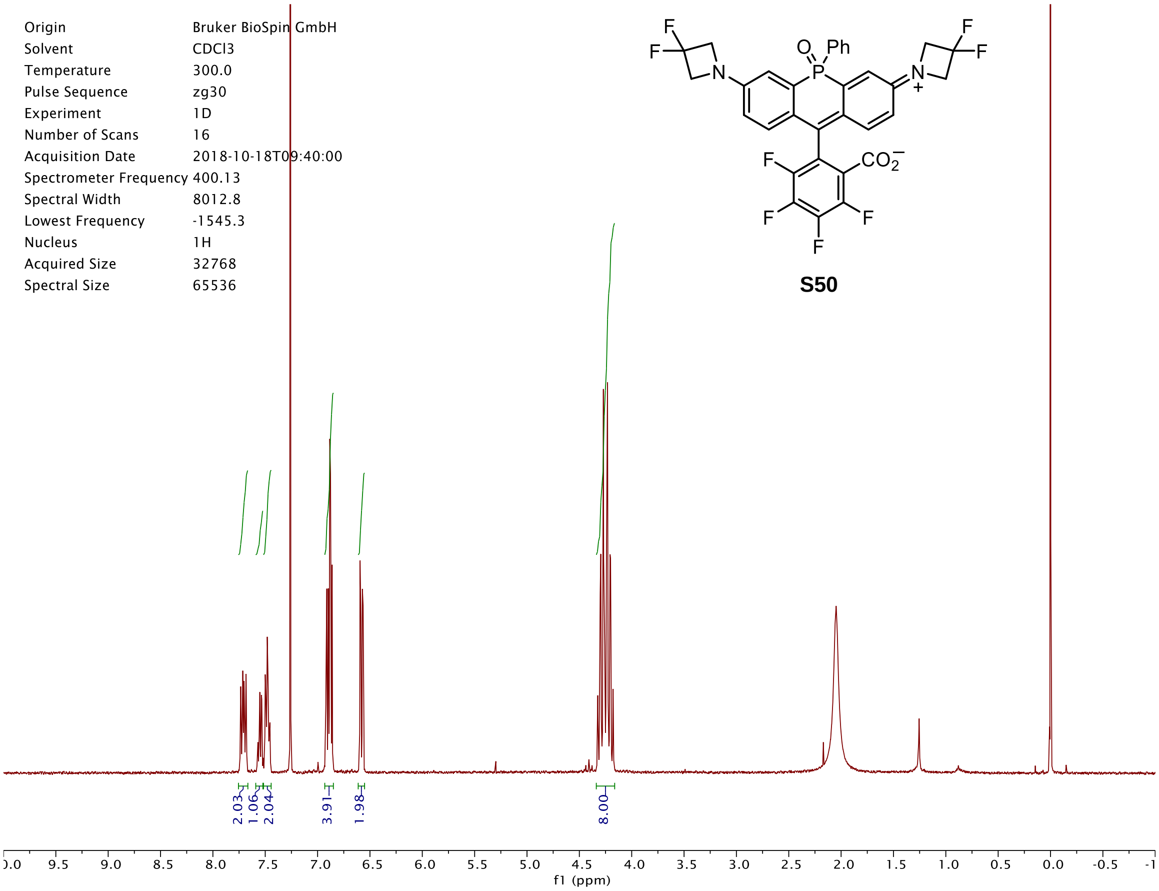

Origin Bruker BioSpin GmbH
Solvent CDCl3
Temperature 300.0
Pulse Sequence zg30
Experiment 1D
Number of Scans 16
Acquisition Date 2018-10-18T09:40:00
Spectrometer Frequency 400.13
Spectral Width 8012.8
Lowest Frequency -1545.3
Nucleus 1H
Acquired Size 32768
Spectral Size 65536
S50
2.03
1.06
2.04
3.91
1.98
8.00
f1 (ppm)


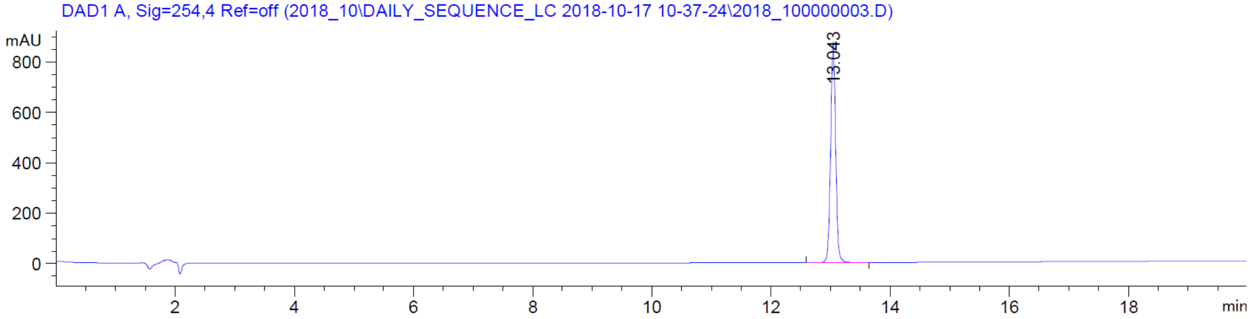

DAD1 A, Sig=254,4 Ref=off (2018_10\DAILY_SEQUENCE_LC 2018-10-17 10-37-24\2018_100000003.D)
mAU
13.043
min


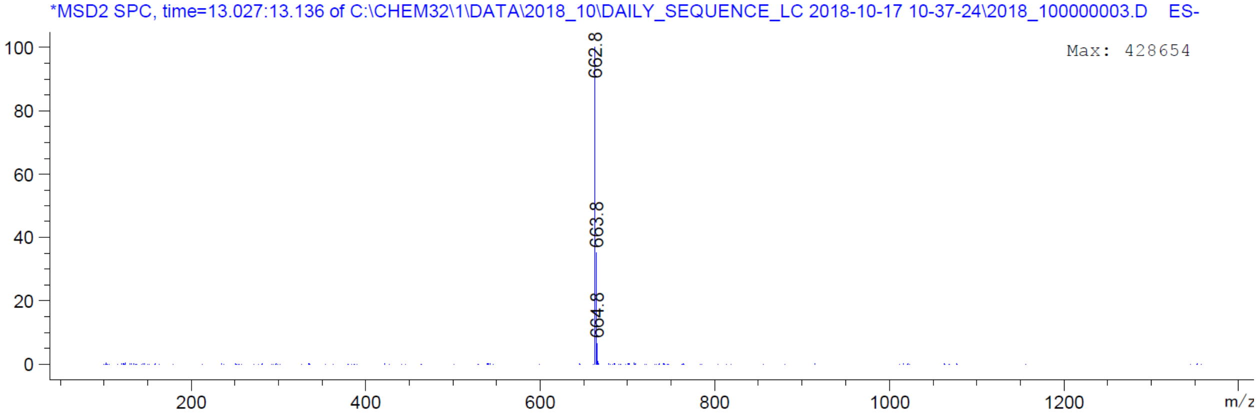

*MSD2 SPC, time=13.027:13.136 of C:\CHEM32\1\DATA\2018_10\DAILY_SEQUENCE_LC 2018-10-17 10-37-24\2018_100000003.D ES-
Max: 428654
662.8
663.8
664.8
m/z

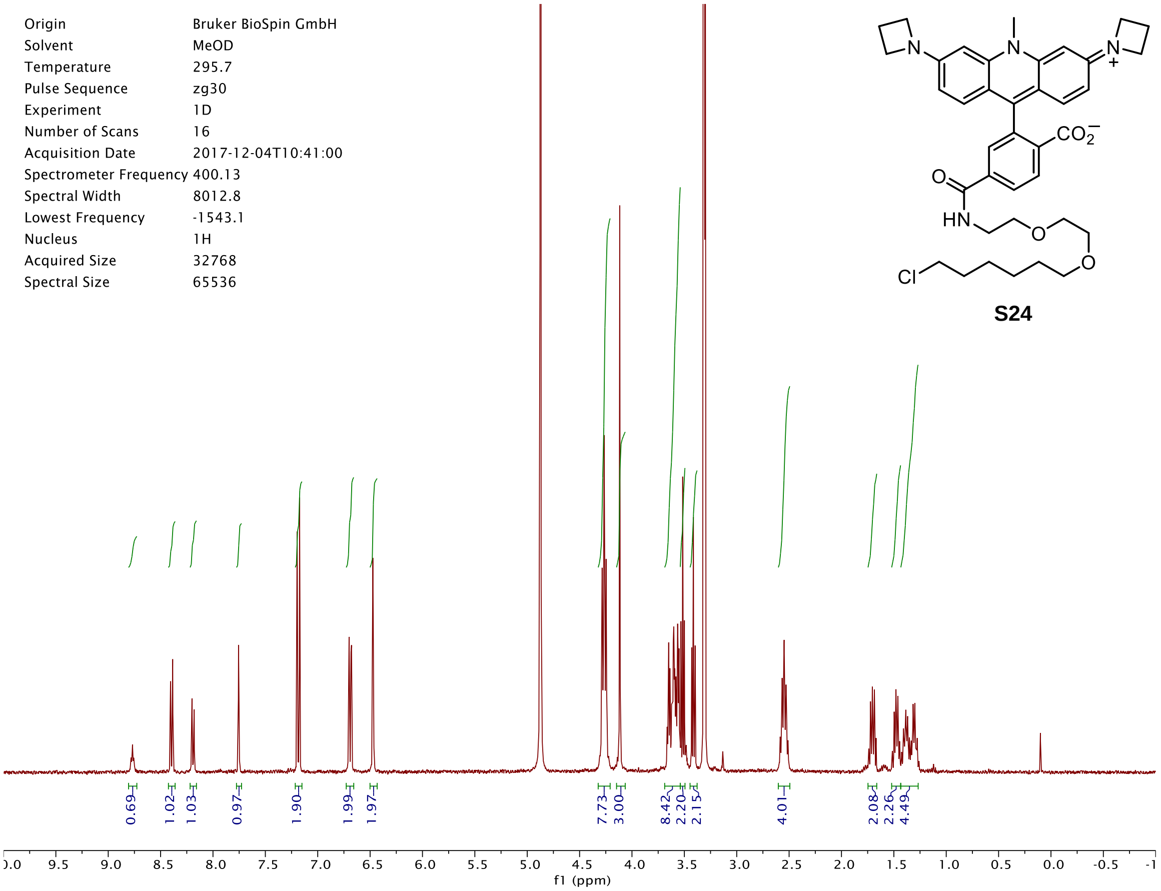

Origin Bruker BioSpin GmbH
Solvent MeOD
Temperature 295.7
Pulse Sequence zg30
Experiment 1D
Number of Scans 16
Acquisition Date 2017-12-04T10:41:00
Spectrometer Frequency 400.13
Spectral Width 8012.8
Lowest Frequency -1543.1
Nucleus 1H
Acquired Size 32768
Spectral Size 65536
0.69 1.02 1.03 0.97 1.90 1.99 1.97 7.73 3.00 8.42 2.20 2.15 4.01 2.08 2.26 4.49
f1 (ppm)


**S24**

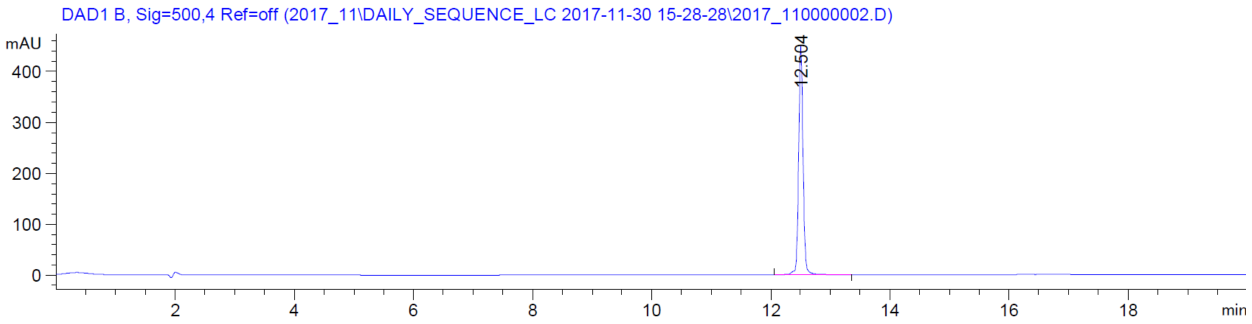

DAD1 B, Sig=500,4 Ref=off (2017_11\DAILY_SEQUENCE_LC 2017-11-30 15-28-28\2017_110000002.D)
mAU
12.504
min


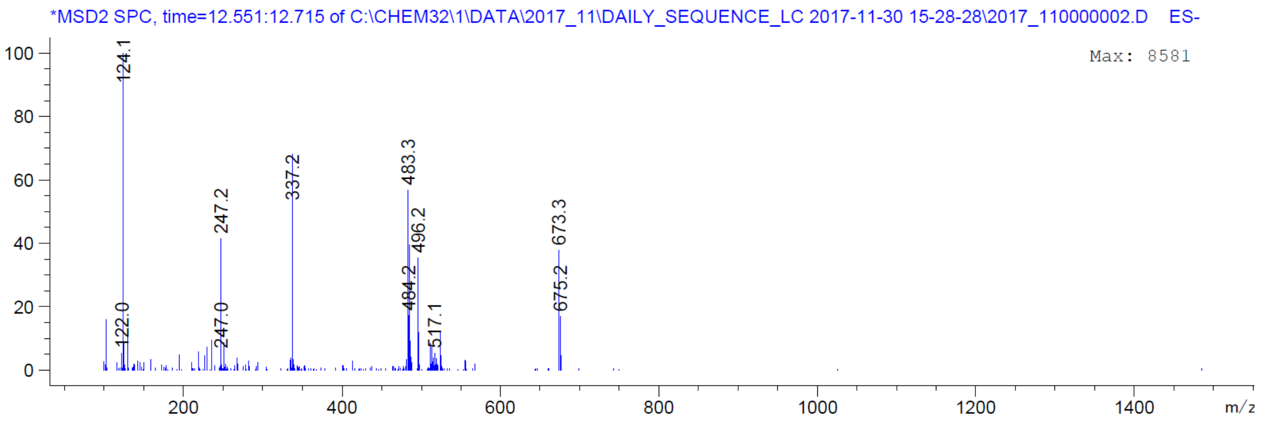

*MSD2 SPC, time=12.551:12.715 of C:\CHEM32\1\DATA\2017_11\DAILY_SEQUENCE_LC 2017-11-30 15-28-28\2017_110000002.D ES-
Max: 8581
124.1
122.0
247.2
247.0
337.2
483.3
484.2
496.2
517.1
673.3
675.2
m/z

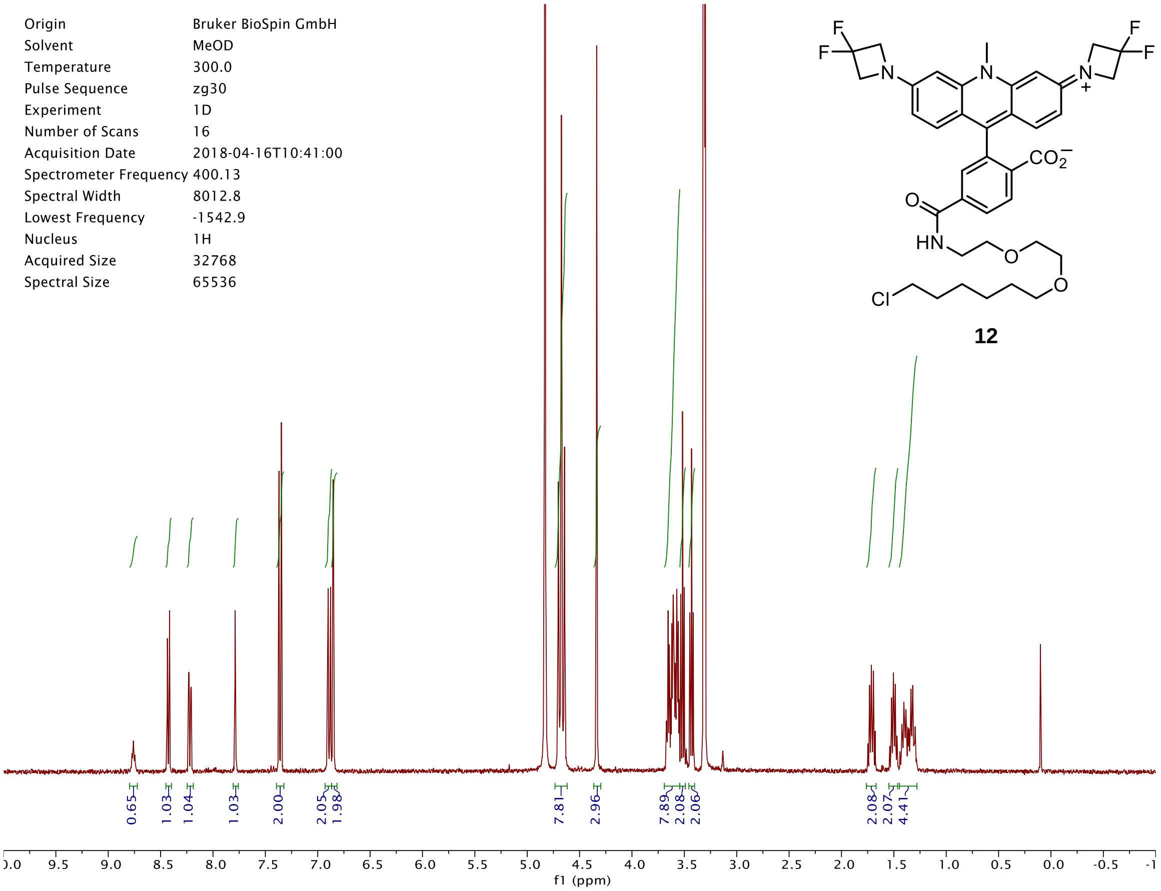

Origin Bruker BioSpin GmbH
Solvent MeOD
Temperature 300.0
Pulse Sequence zg30
Experiment 1D
Number of Scans 16
Acquisition Date 2018-04-16T10:41:00
Spectrometer Frequency 400.13
Spectral Width 8012.8
Lowest Frequency -1542.9
Nucleus 1H
Acquired Size 32768
Spectral Size 65536
0.65 1.03 1.04 1.03 2.00 2.05 1.98 7.81 2.96 7.89 2.08 2.06 2.08 2.07 4.41
f1 (ppm)
F
F
N
N
N
+
F
F
$CO_2^-$
O
HN
O
O
Cl


**12**

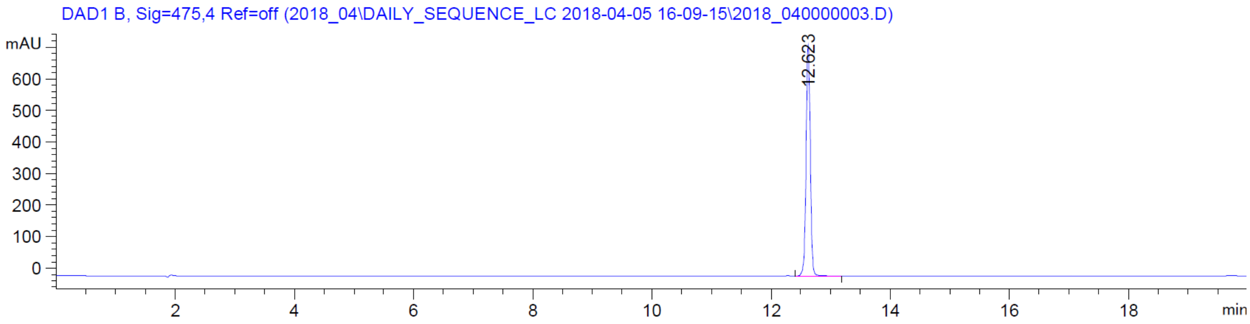

DAD1 B, Sig=475,4 Ref=off (2018_04\DAILY_SEQUENCE_LC 2018-04-05 16-09-15\2018_040000003.D)
mAU
12.623
min


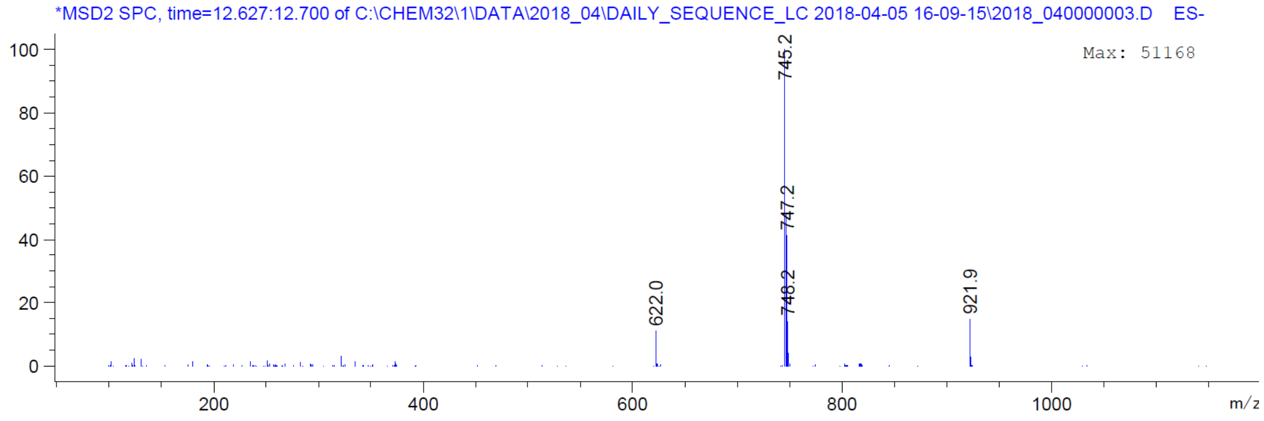

*MSD2 SPC, time=12.627:12.700 of C:\CHEM32\1\DATA\2018_04\DAILY_SEQUENCE_LC 2018-04-05 16-09-15\2018_040000003.D ES-
Max: 51168
745.2
747.2
748.2
622.0
921.9
m/z

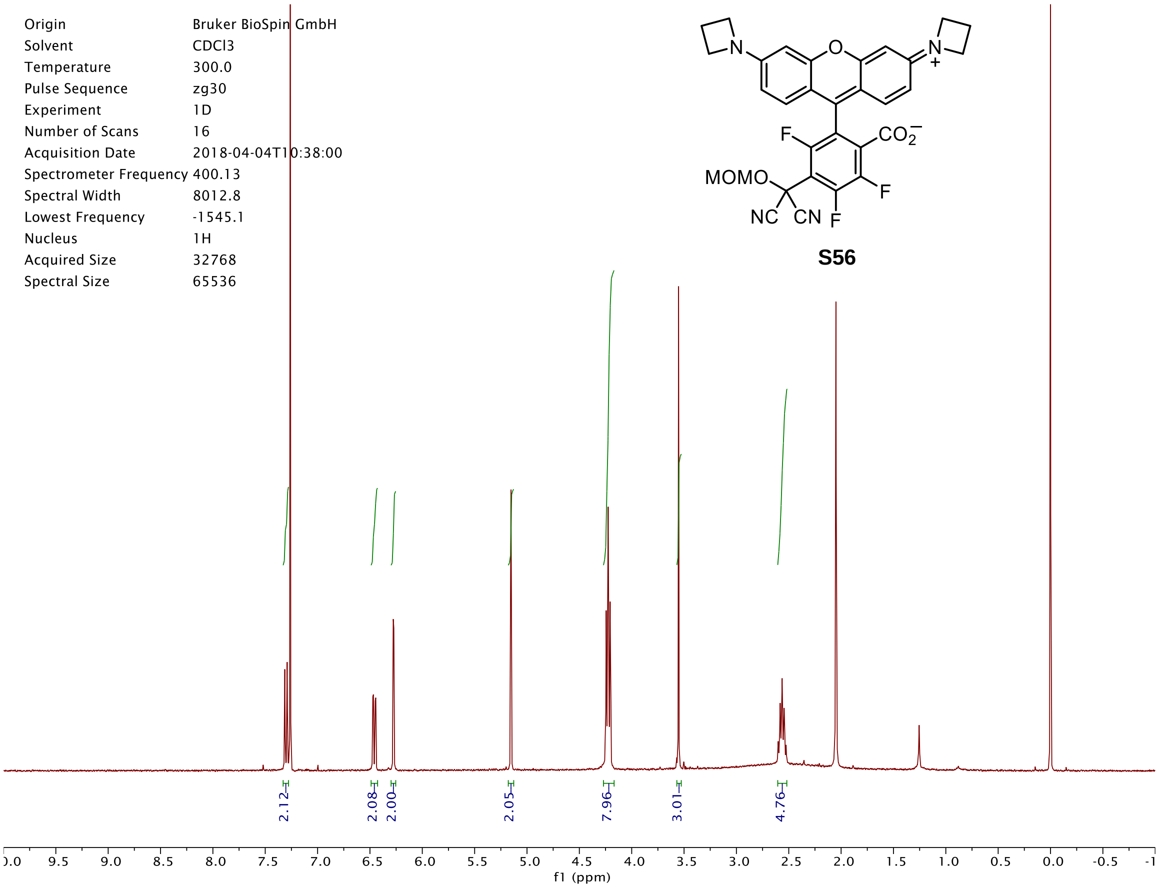

Origin Bruker BioSpin GmbH
Solvent CDCl3
Temperature 300.0
Pulse Sequence zg30
Experiment 1D
Number of Scans 16
Acquisition Date 2018-04-04T10:38:00
Spectrometer Frequency 400.13
Spectral Width 8012.8
Lowest Frequency -1545.1
Nucleus 1H
Acquired Size 32768
Spectral Size 65536
MOMO
NC CN F
F
F
CO2−
N
N+
O
S56
2.12
2.08
2.00
2.05
7.96
3.01
4.76
f1 (ppm)


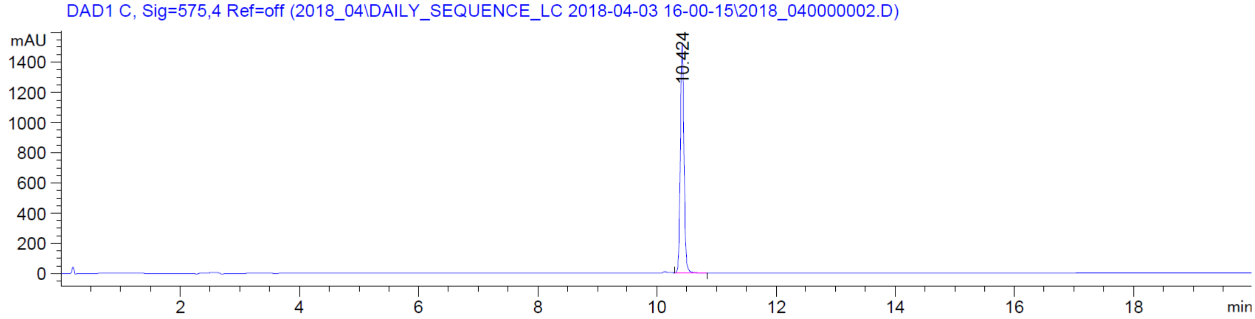

DAD1 C, Sig=575,4 Ref=off (2018_04\DAILY_SEQUENCE_LC 2018-04-03 16-00-15\2018_040000002.D)
mAU
10.424
min


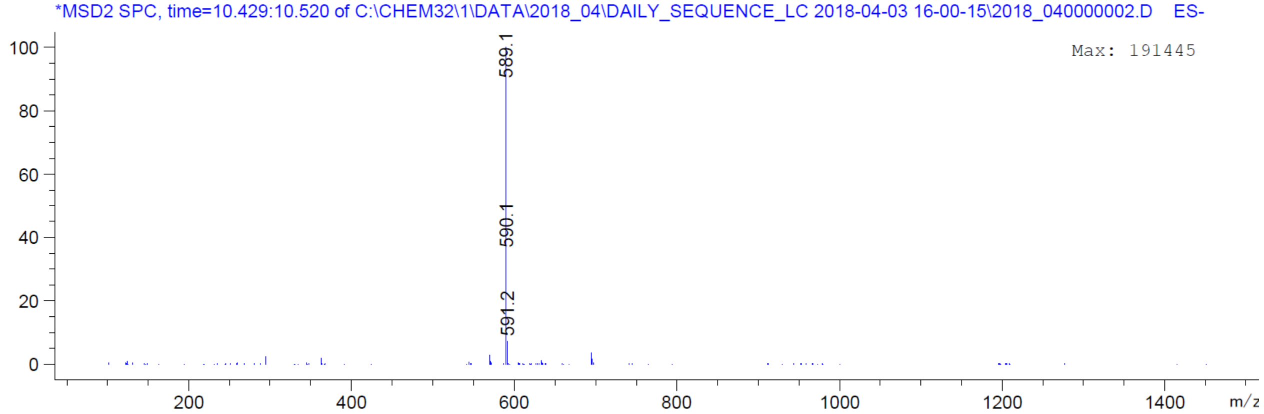

*MSD2 SPC, time=10.429:10.520 of C:\CHEM32\1\DATA\2018_04\DAILY_SEQUENCE_LC 2018-04-03 16-00-15\2018_040000002.D ES-
Max: 191445
589.1
590.1
591.2
m/z

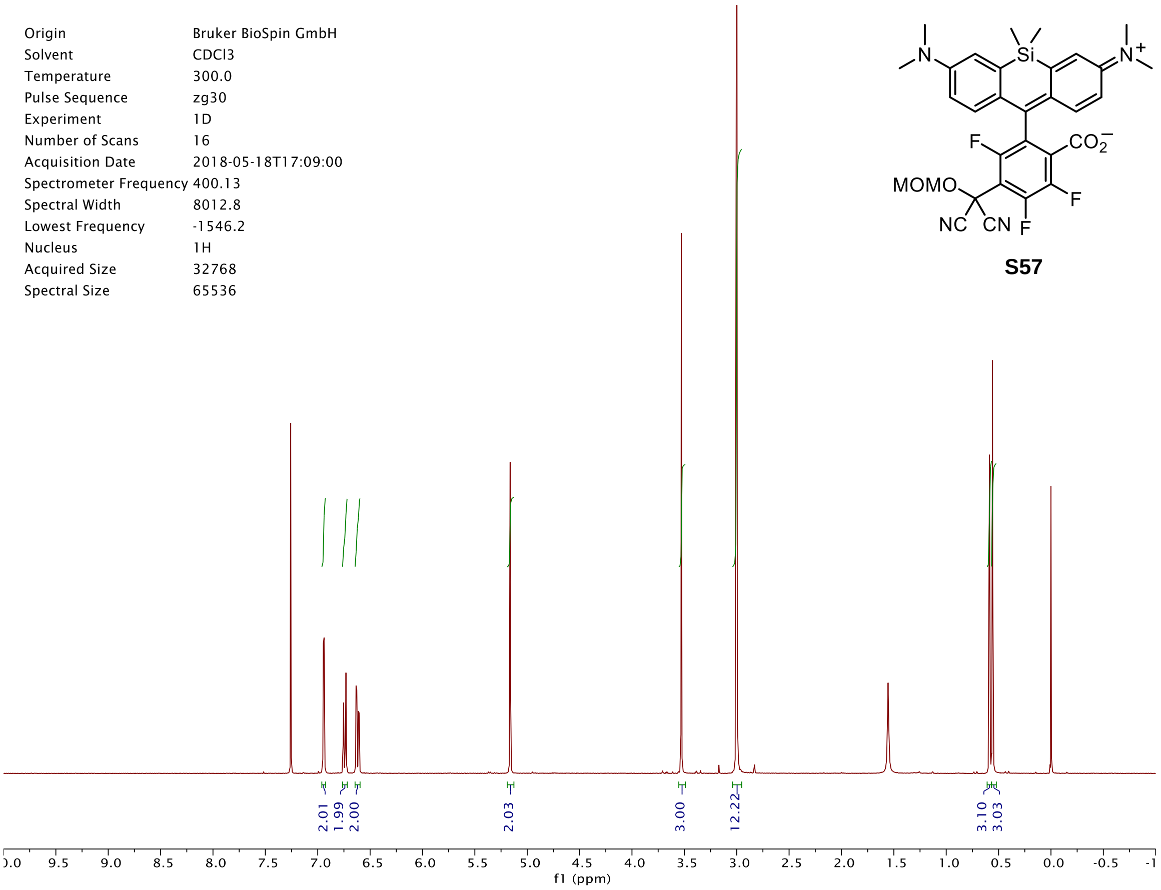
Origin Bruker BioSpin GmbH
Solvent CDCl3
Temperature 300.0
Pulse Sequence zg30
Experiment 1D
Number of Scans 16
Acquisition Date 2018-05-18T17:09:00
Spectrometer Frequency 400.13
Spectral Width 8012.8
Lowest Frequency -1546.2
Nucleus 1H
Acquired Size 32768
Spectral Size 65536
S57
2.01
1.99
2.00
2.03
3.00
12.22
3.10
3.03
f1 (ppm)

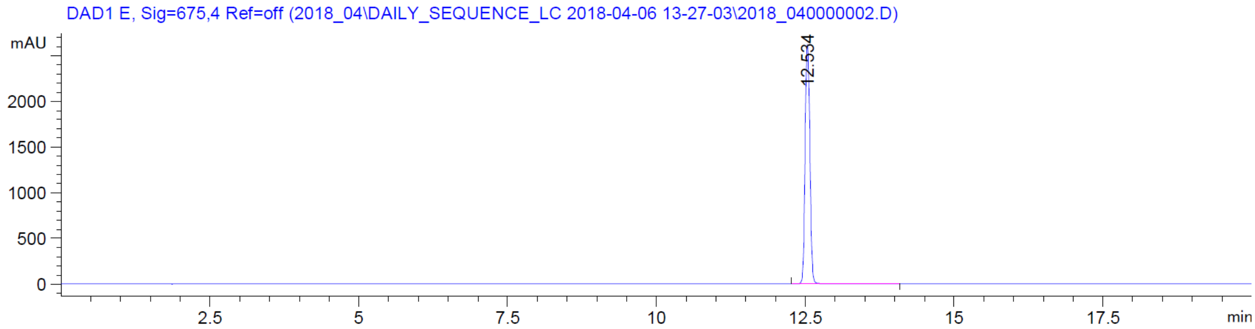
DAD1 E, Sig=675,4 Ref=off (2018_04\DAILY_SEQUENCE_LC 2018-04-06 13-27-03\2018_040000002.D)
mAU
12.534
min

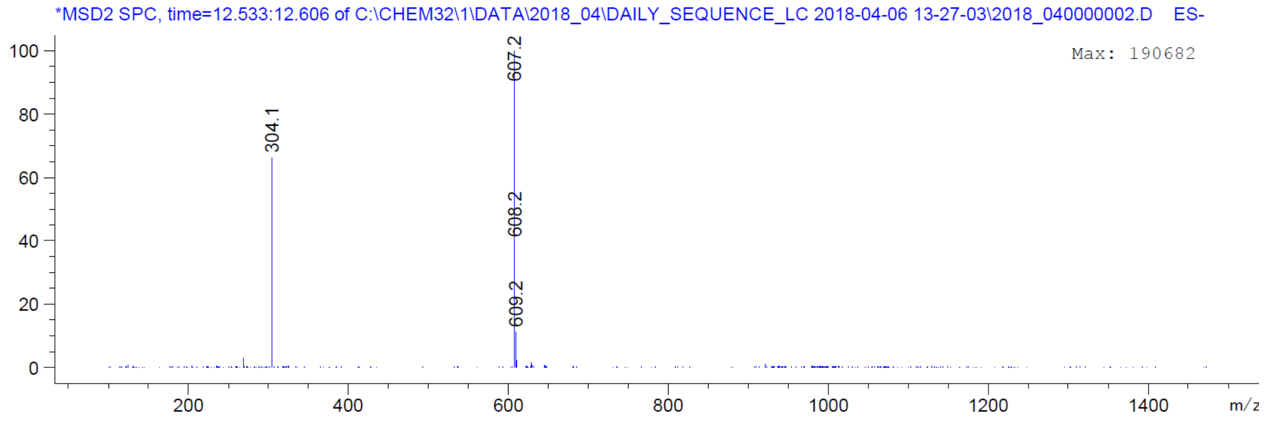
*MSD2 SPC, time=12.533:12.606 of C:\CHEM32\1\DATA\2018_04\DAILY_SEQUENCE_LC 2018-04-06 13-27-03\2018_040000002.D ES-
Max: 190682
304.1
607.2
608.2
609.2
m/z

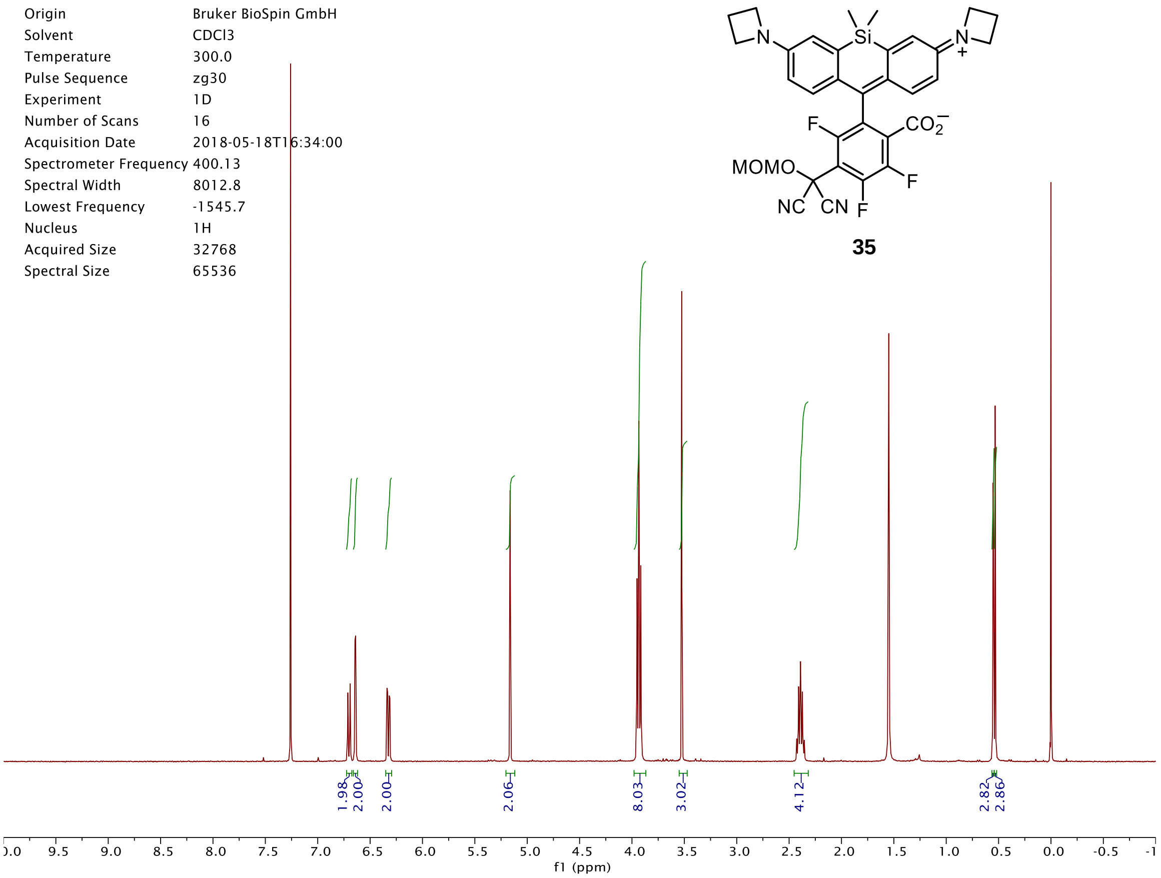

Origin Bruker BioSpin GmbH
Solvent CDCl3
Temperature 300.0
Pulse Sequence zg30
Experiment 1D
Number of Scans 16
Acquisition Date 2018-05-18T16:34:00
Spectrometer Frequency 400.13
Spectral Width 8012.8
Lowest Frequency -1545.7
Nucleus 1H
Acquired Size 32768
Spectral Size 65536
1.98
2.00
2.00
2.06
8.03
3.02
4.12
2.82
2.86
f1 (ppm)
Si
N
N
+
F
$CO_2^-$
MOMO
F
NC
CN
F


**35**

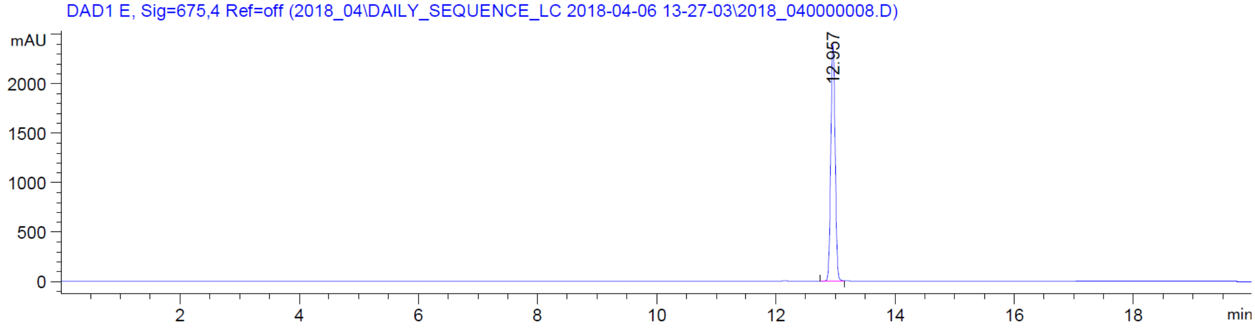

DAD1 E, Sig=675,4 Ref=off (2018_04\DAILY_SEQUENCE_LC 2018-04-06 13-27-03\2018_040000008.D)
mAU
12.957
min


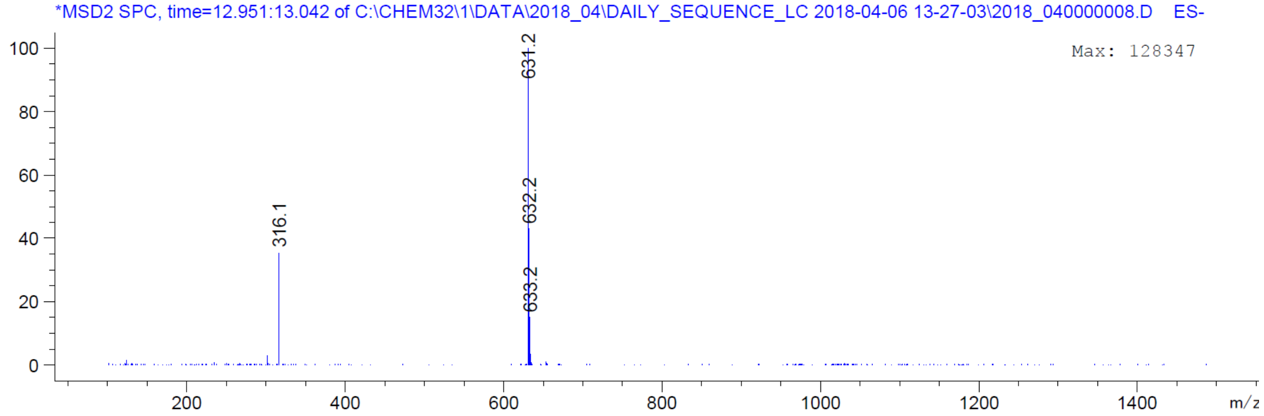

*MSD2 SPC, time=12.951:13.042 of C:\CHEM32\1\DATA\2018_04\DAILY_SEQUENCE_LC 2018-04-06 13-27-03\2018_040000008.D ES-
Max: 128347
631.2
632.2
633.2
316.1
m/z

| | |
|---|---|
| Origin | Bruker BioSpin GmbH |
| Solvent | MeOD |
| Temperature | 300.0 |
| Pulse Sequence | zg30 |
| Experiment | 1D |
| Number of Scans | 16 |
| Acquisition Date | 2018-05-29T11:31:00 |
| Spectrometer Frequency | 400.13 |
| Spectral Width | 8012.8 |
| Lowest Frequency | -1543.2 |
| Nucleus | 1H |
| Acquired Size | 32768 |
| Spectral Size | 65536 |

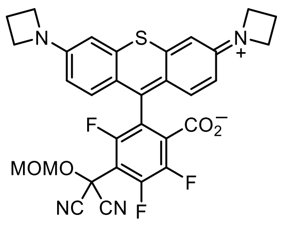


**S67**

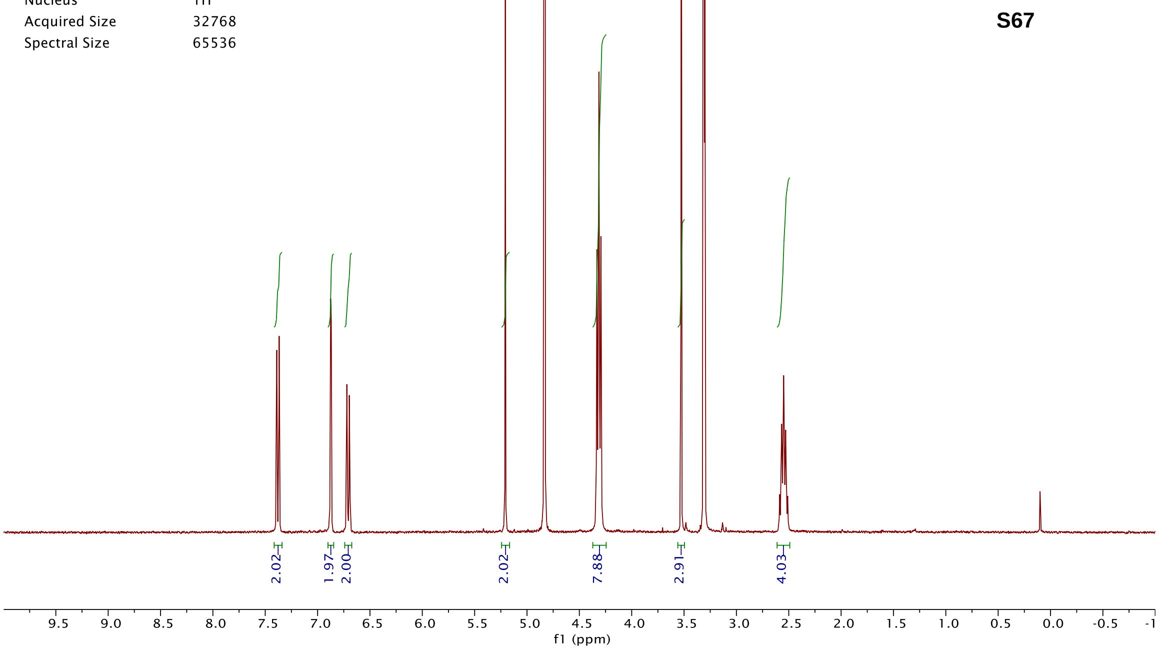


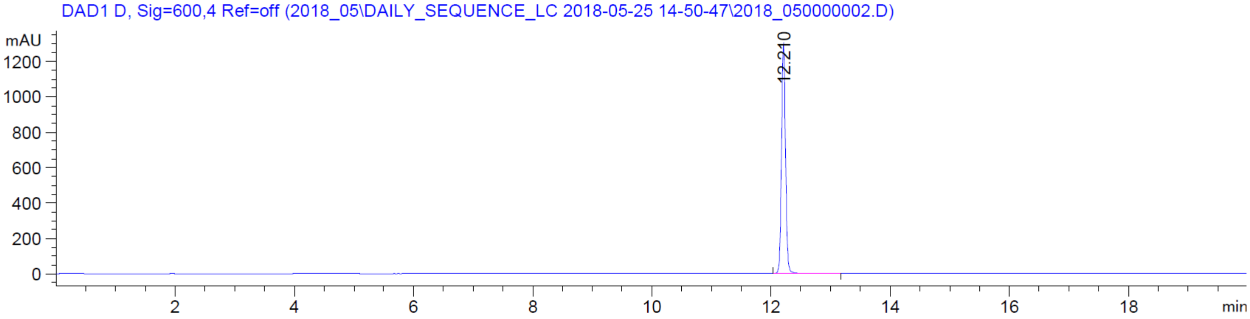


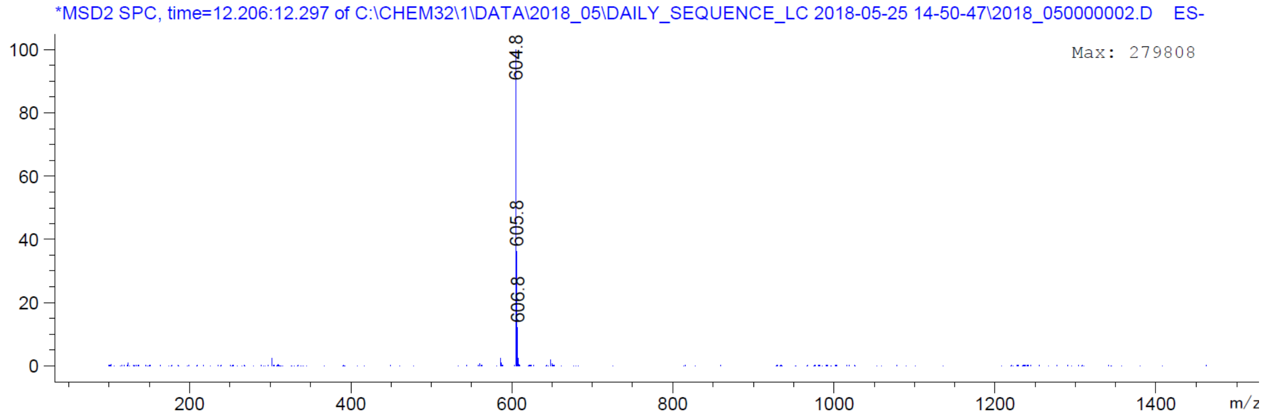

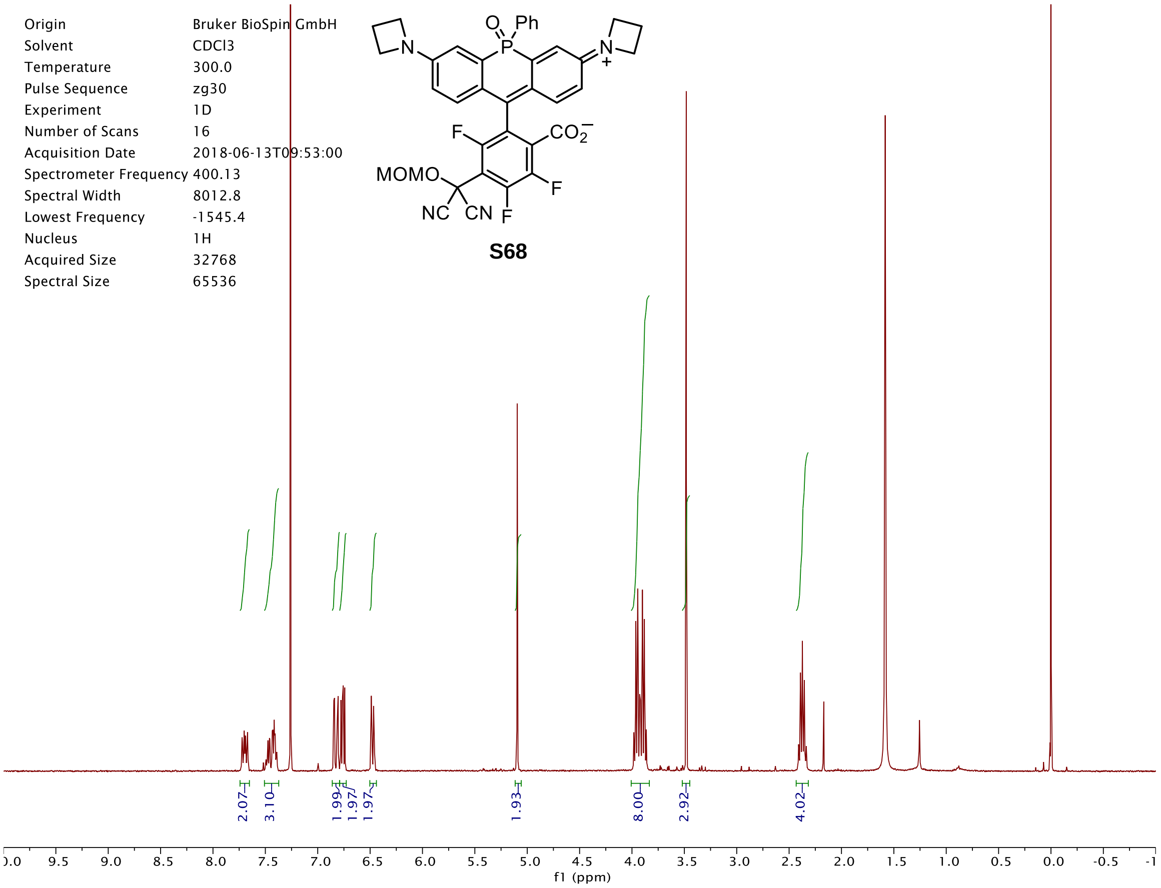
Origin Bruker BioSpin GmbH
Solvent CDCl3
Temperature 300.0
Pulse Sequence zg30
Experiment 1D
Number of Scans 16
Acquisition Date 2018-06-13T09:53:00
Spectrometer Frequency 400.13
Spectral Width 8012.8
Lowest Frequency -1545.4
Nucleus 1H
Acquired Size 32768
Spectral Size 65536
MOMO
NC
CN
F
Ph
S68
2.07
3.10
1.99
1.97
1.97
1.93
8.00
2.92
4.02
f1 (ppm)

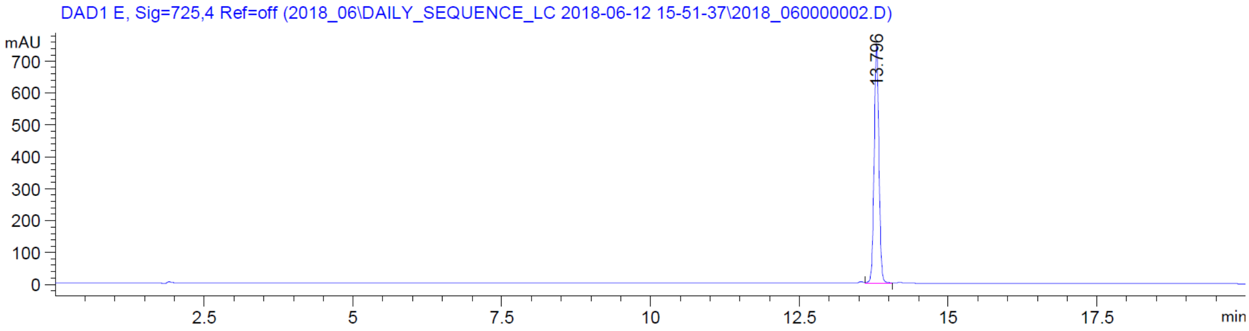
DAD1 E, Sig=725,4 Ref=off (2018_06\DAILY_SEQUENCE_LC 2018-06-12 15-51-37\2018_060000002.D)
mAU
13.796
min

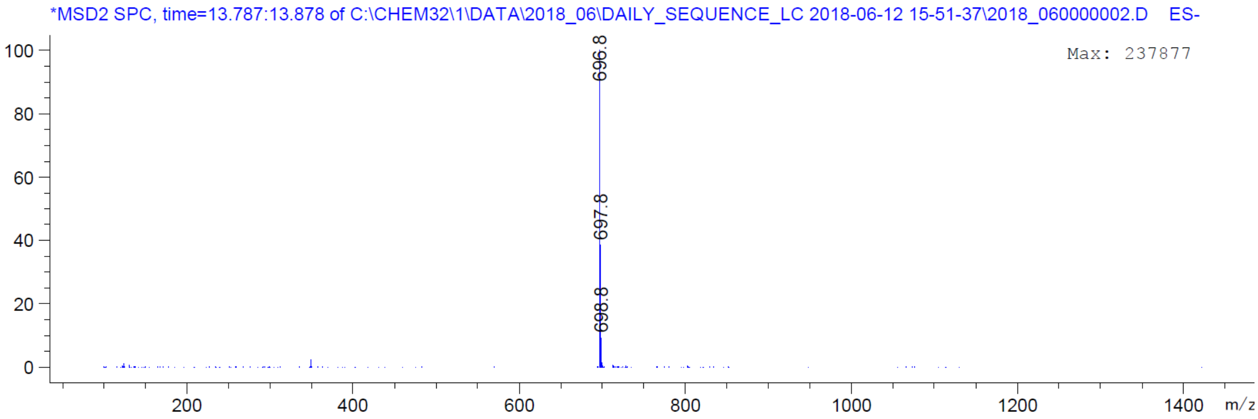
*MSD2 SPC, time=13.787:13.878 of C:\CHEM32\1\DATA\2018_06\DAILY_SEQUENCE_LC 2018-06-12 15-51-37\2018_060000002.D ES-
Max: 237877
696.8
697.8
698.8
m/z

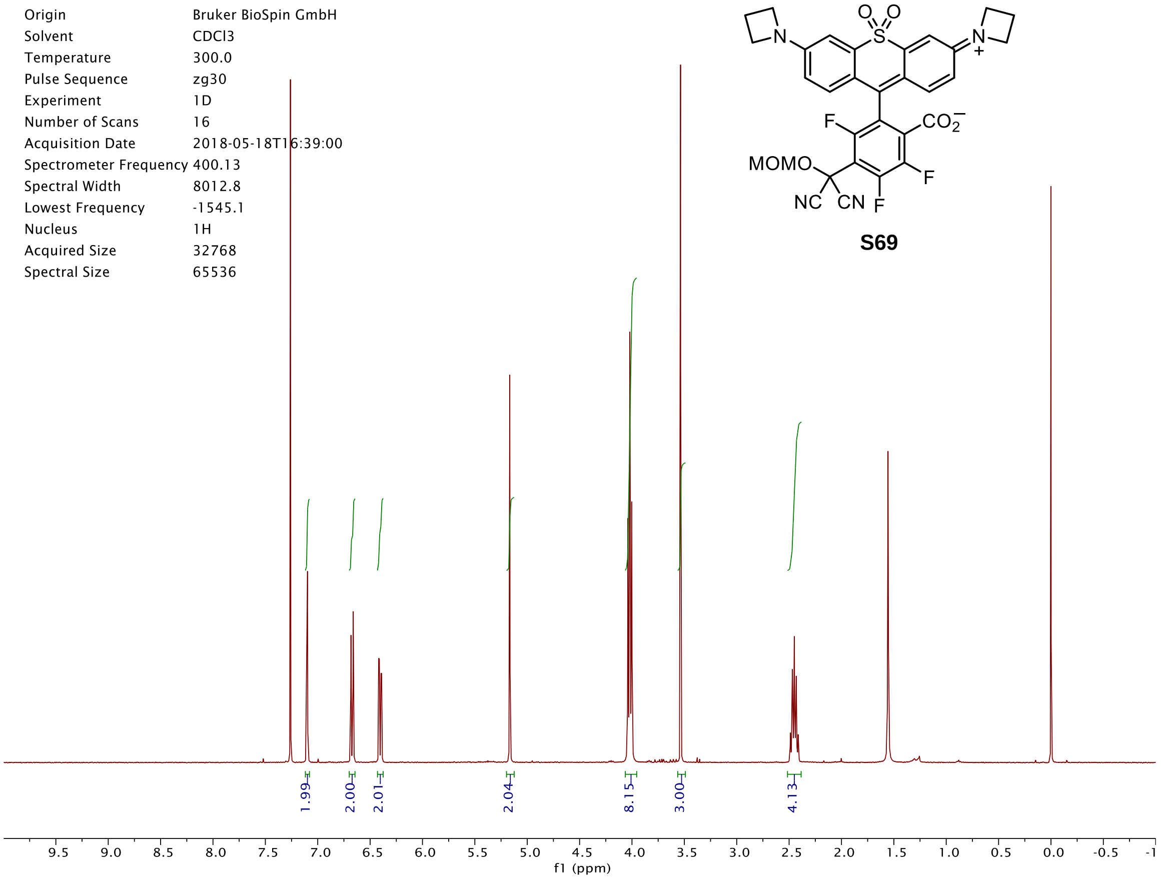
Origin Bruker BioSpin GmbH
Solvent CDCl3
Temperature 300.0
Pulse Sequence zg30
Experiment 1D
Number of Scans 16
Acquisition Date 2018-05-18T16:39:00
Spectrometer Frequency 400.13
Spectral Width 8012.8
Lowest Frequency -1545.1
Nucleus 1H
Acquired Size 32768
Spectral Size 65536
1.99
2.00
2.01
2.04
8.15
3.00
4.13
f1 (ppm)
MOMO
NC
CN
F
CO2
N
S
O

**S69**

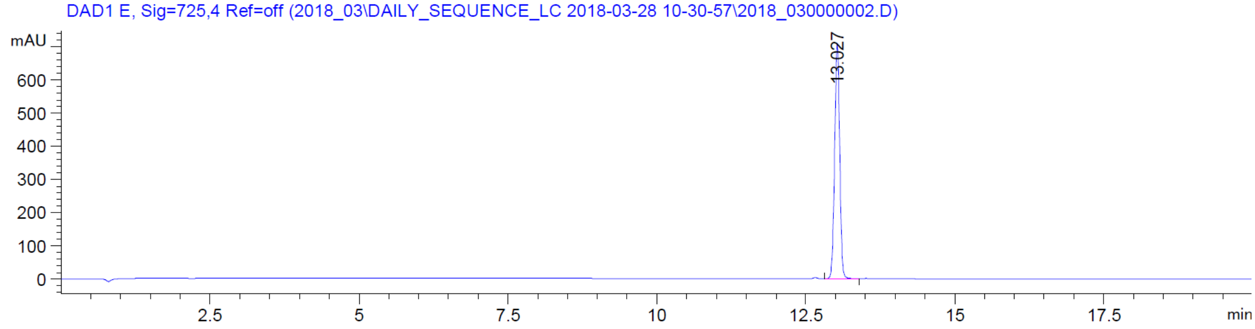
DAD1 E, Sig=725,4 Ref=off (2018_03\DAILY_SEQUENCE_LC 2018-03-28 10-30-57\2018_030000002.D)
mAU
13.027
min

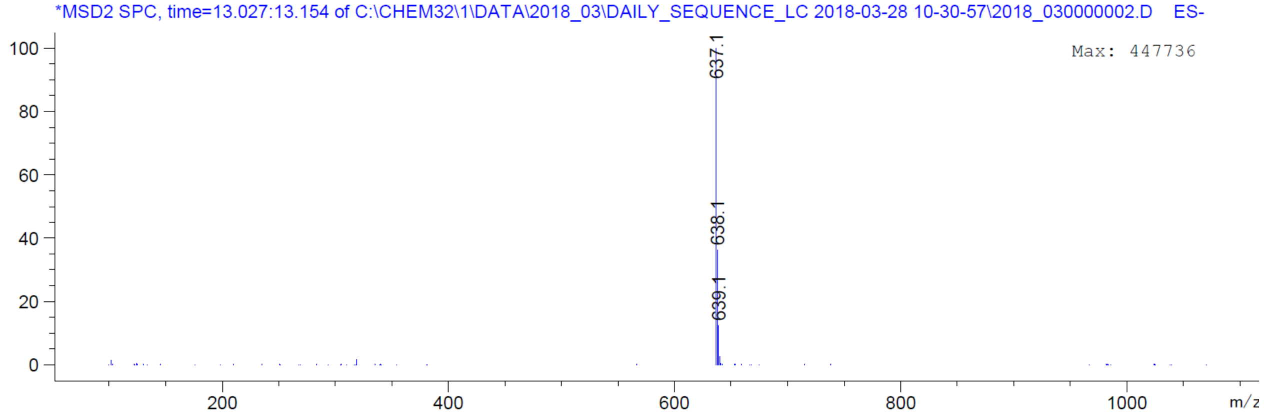
*MSD2 SPC, time=13.027:13.154 of C:\CHEM32\1\DATA\2018_03\DAILY_SEQUENCE_LC 2018-03-28 10-30-57\2018_030000002.D ES-
Max: 447736
637.1
638.1
639.1
m/z

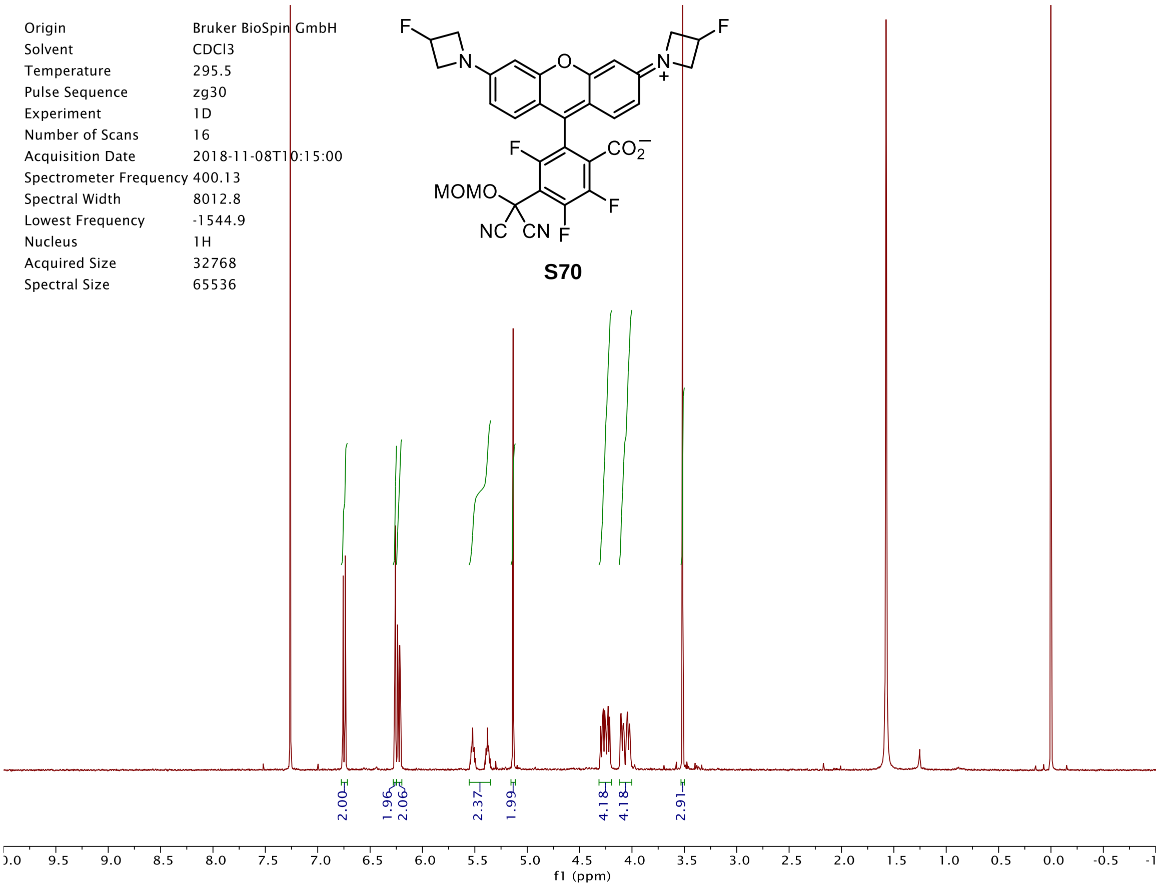
Origin Bruker BioSpin GmbH
Solvent CDCl3
Temperature 295.5
Pulse Sequence zg30
Experiment 1D
Number of Scans 16
Acquisition Date 2018-11-08T10:15:00
Spectrometer Frequency 400.13
Spectral Width 8012.8
Lowest Frequency -1544.9
Nucleus 1H
Acquired Size 32768
Spectral Size 65536
MOMO
NC CN
S70
2.00 1.96 2.06 2.37 1.99 4.18 4.18 2.91
f1 (ppm)

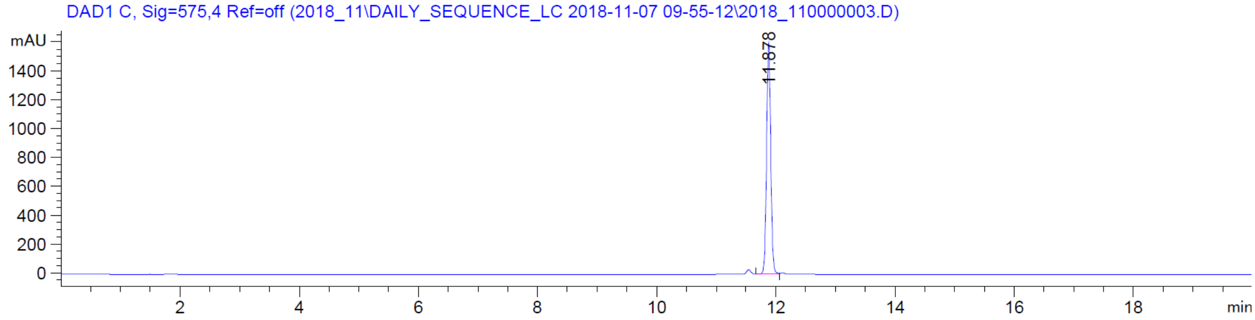
DAD1 C, Sig=575,4 Ref=off (2018_11\DAILY_SEQUENCE_LC 2018-11-07 09-55-12\2018_110000003.D)
mAU
11.878
min

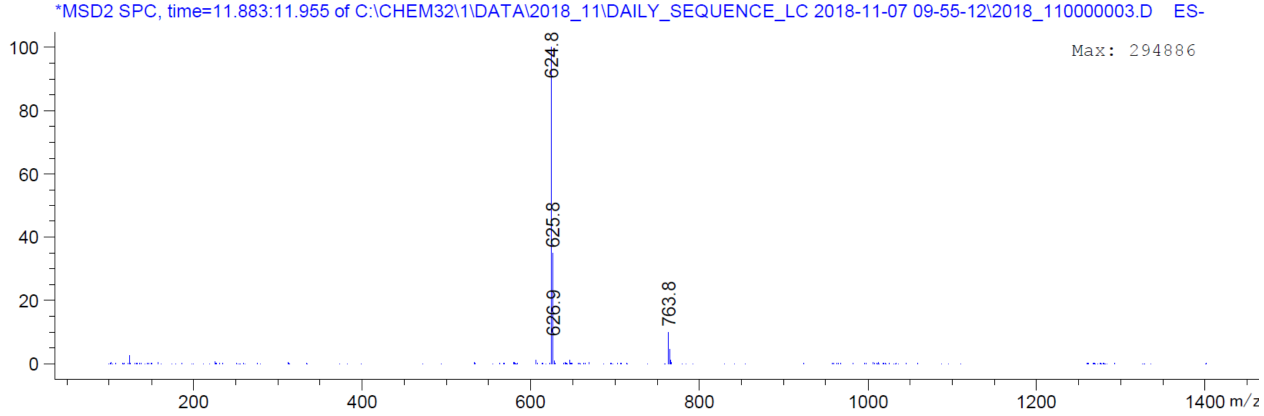
*MSD2 SPC, time=11.883:11.955 of C:\CHEM32\1\DATA\2018_11\DAILY_SEQUENCE_LC 2018-11-07 09-55-12\2018_110000003.D ES-
Max: 294886
624.8
625.8
626.9
763.8
m/z

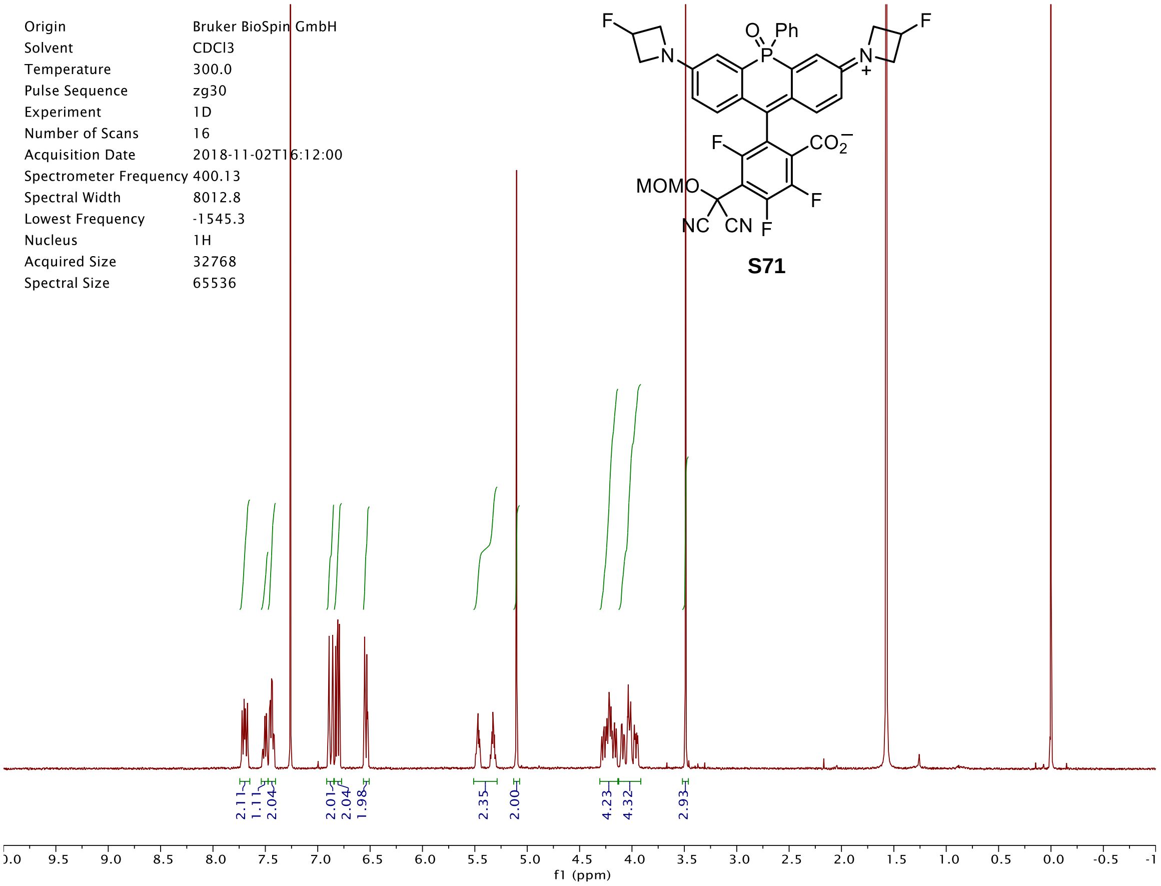

Origin Bruker BioSpin GmbH
Solvent CDCl3
Temperature 300.0
Pulse Sequence zg30
Experiment 1D
Number of Scans 16
Acquisition Date 2018-11-02T16:12:00
Spectrometer Frequency 400.13
Spectral Width 8012.8
Lowest Frequency -1545.3
Nucleus 1H
Acquired Size 32768
Spectral Size 65536
S71
2.11 1.11 2.04 2.01 2.04 1.98 2.35 2.00 4.23 4.32 2.93
f1 (ppm)


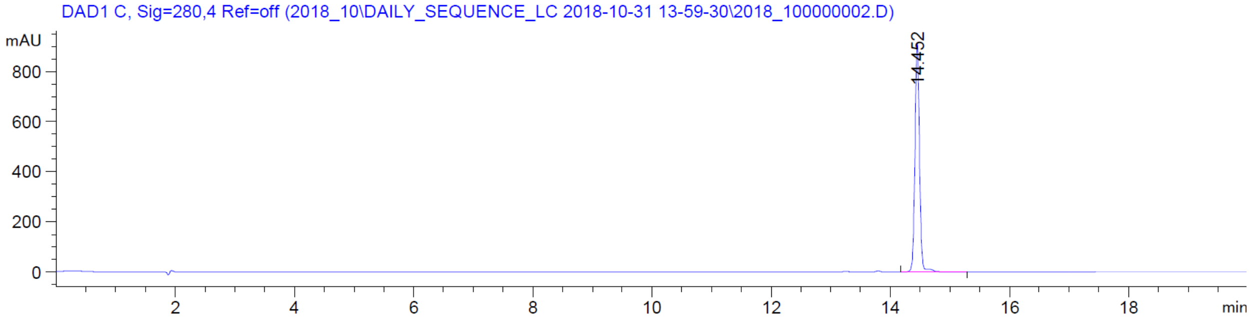

DAD1 C, Sig=280,4 Ref=off (2018_10\DAILY_SEQUENCE_LC 2018-10-31 13-59-30\2018_100000002.D)
mAU
14.452
min


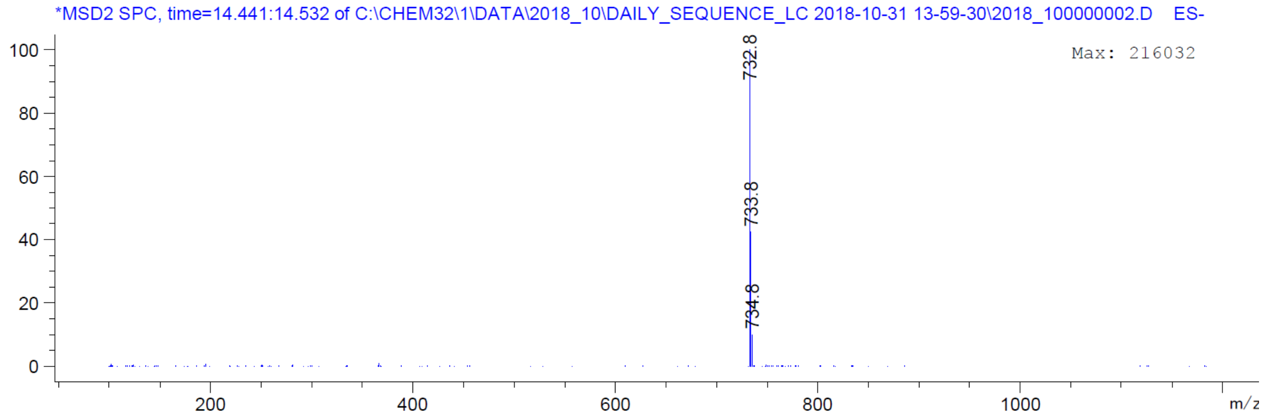

*MSD2 SPC, time=14.441:14.532 of C:\CHEM32\1\DATA\2018_10\DAILY_SEQUENCE_LC 2018-10-31 13-59-30\2018_100000002.D ES-
Max: 216032
732.8
733.8
734.8
m/z

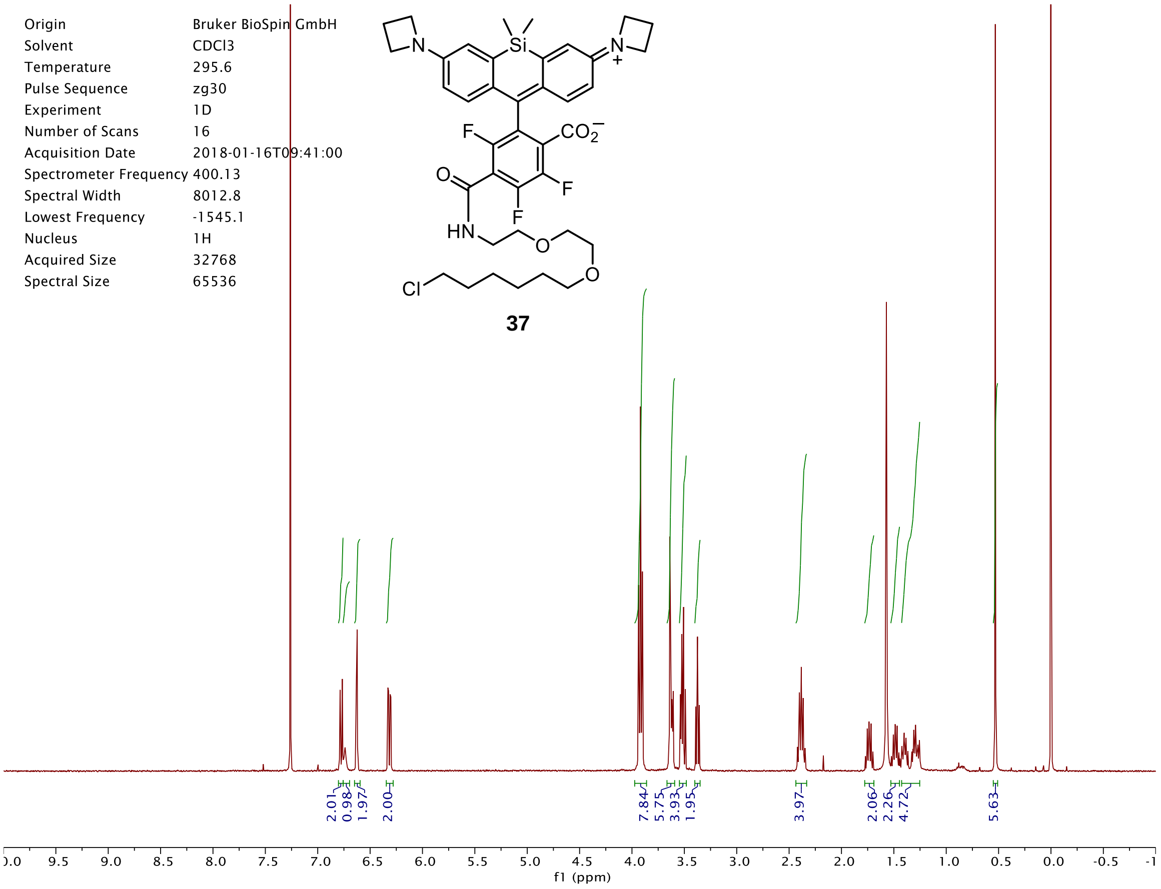
Origin Bruker BioSpin GmbH
Solvent CDCl3
Temperature 295.6
Pulse Sequence zg30
Experiment 1D
Number of Scans 16
Acquisition Date 2018-01-16T09:41:00
Spectrometer Frequency 400.13
Spectral Width 8012.8
Lowest Frequency -1545.1
Nucleus 1H
Acquired Size 32768
Spectral Size 65536
37
2.01 0.98 1.97 2.00 7.84 5.75 3.93 1.95 3.97 2.06 2.26 4.72 5.63
f1 (ppm)

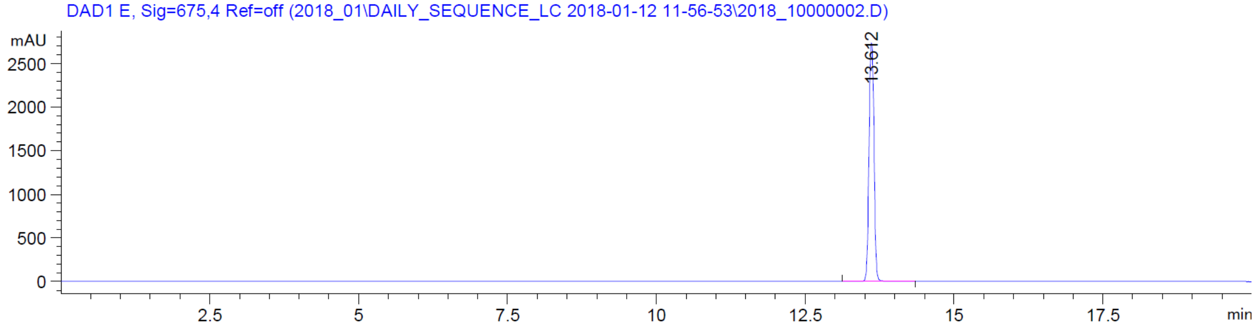
DAD1 E, Sig=675,4 Ref=off (2018_01\DAILY_SEQUENCE_LC 2018-01-12 11-56-53\2018_10000002.D)
mAU
13.612
min

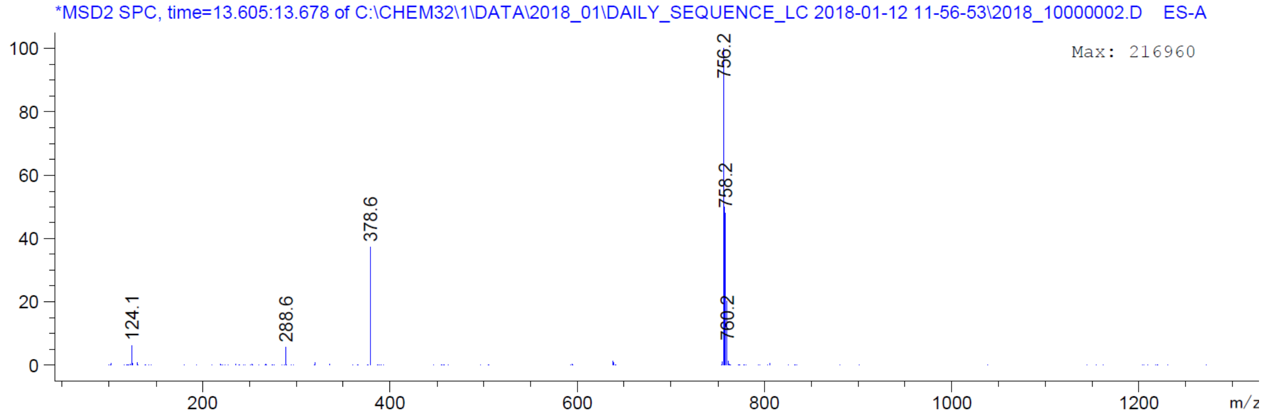
*MSD2 SPC, time=13.605:13.678 of C:\CHEM32\1\DATA\2018_01\DAILY_SEQUENCE_LC 2018-01-12 11-56-53\2018_10000002.D ES-A
Max: 216960
756.2
758.2
760.2
378.6
288.6
124.1
m/z

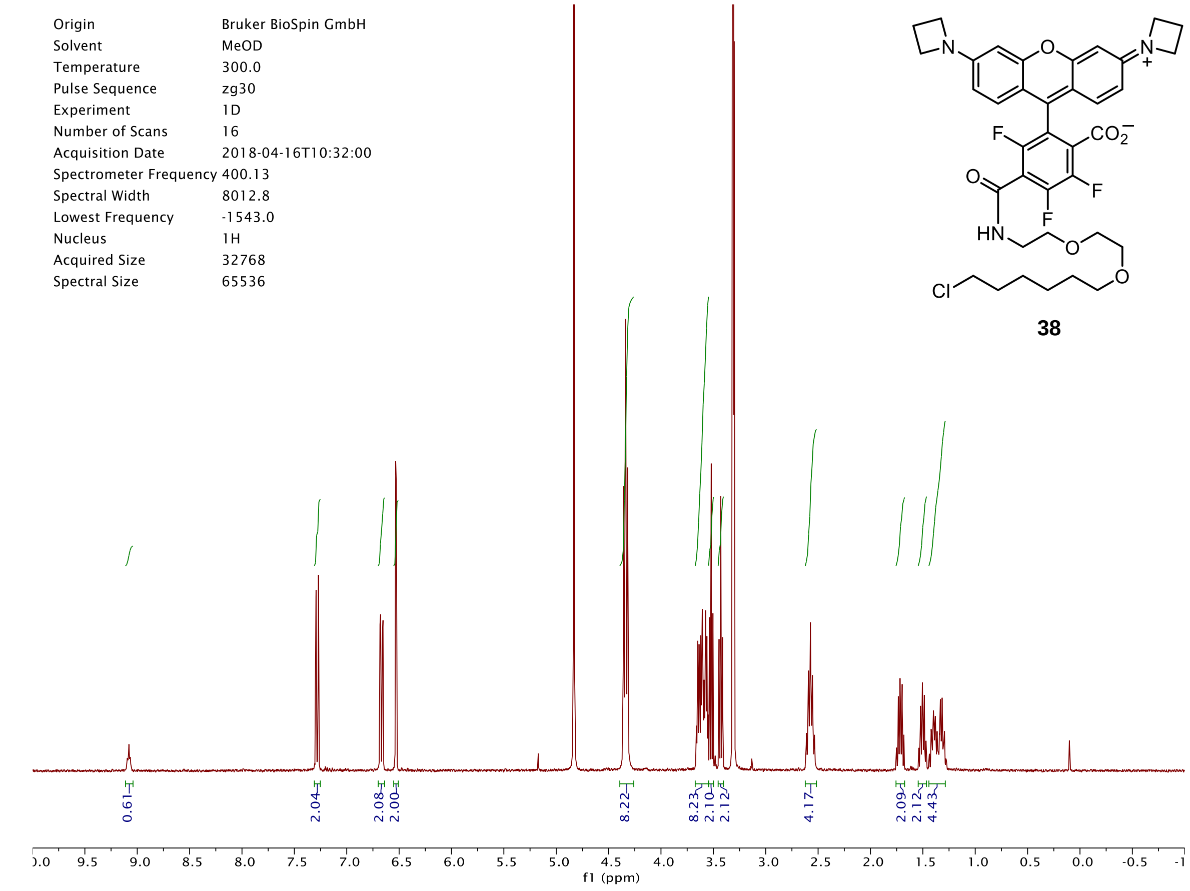
Origin Bruker BioSpin GmbH
Solvent MeOD
Temperature 300.0
Pulse Sequence zg30
Experiment 1D
Number of Scans 16
Acquisition Date 2018-04-16T10:32:00
Spectrometer Frequency 400.13
Spectral Width 8012.8
Lowest Frequency -1543.0
Nucleus 1H
Acquired Size 32768
Spectral Size 65536
38
0.61
2.04
2.08
2.00
8.22
8.23
2.10
2.12
4.17
2.09
2.12
4.43
f1 (ppm)

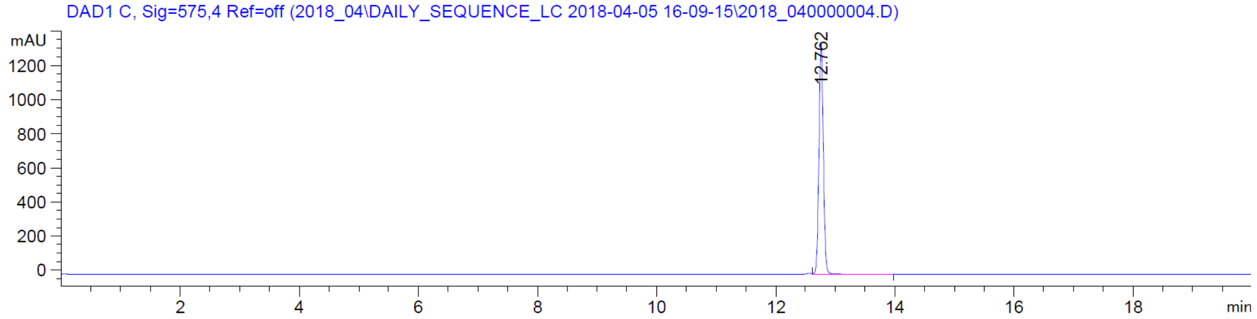
DAD1 C, Sig=575,4 Ref=off (2018_04\DAILY_SEQUENCE_LC 2018-04-05 16-09-15\2018_040000004.D)
mAU
12.762
min

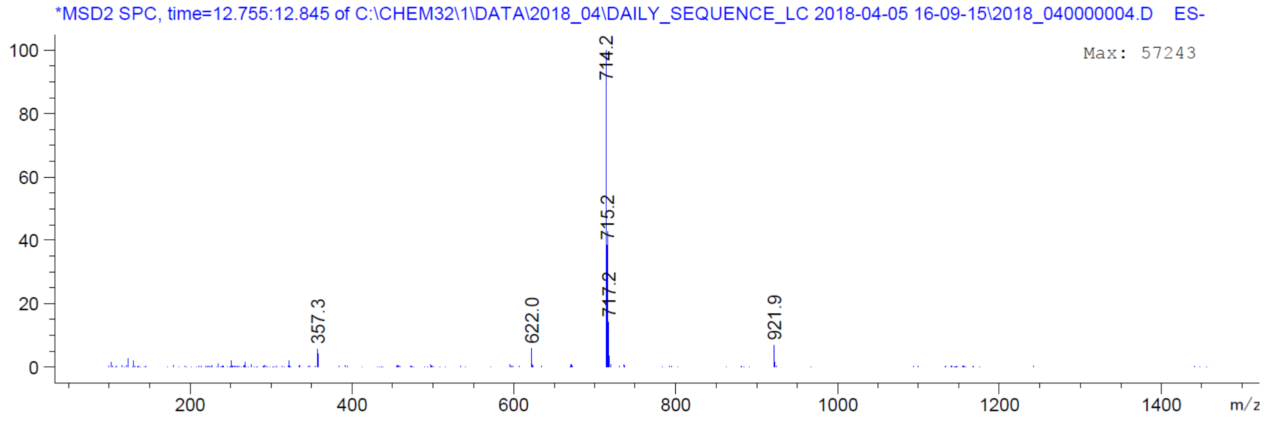
*MSD2 SPC, time=12.755:12.845 of C:\CHEM32\1\DATA\2018_04\DAILY_SEQUENCE_LC 2018-04-05 16-09-15\2018_040000004.D ES-
Max: 57243
714.2
715.2
717.2
357.3
622.0
921.9
m/z

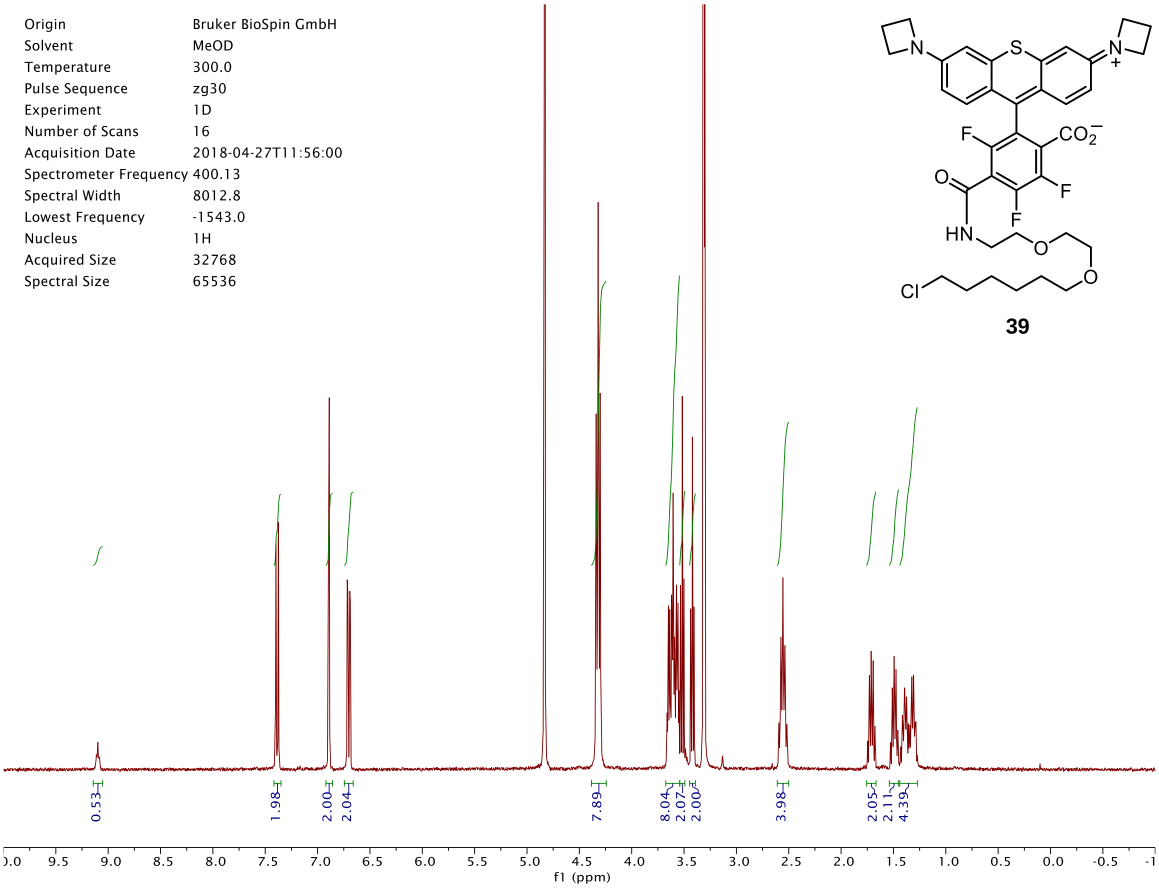

Origin Bruker BioSpin GmbH
Solvent MeOD
Temperature 300.0
Pulse Sequence zg30
Experiment 1D
Number of Scans 16
Acquisition Date 2018-04-27T11:56:00
Spectrometer Frequency 400.13
Spectral Width 8012.8
Lowest Frequency -1543.0
Nucleus 1H
Acquired Size 32768
Spectral Size 65536
0.53
1.98
2.00
2.04
7.89
8.04
2.07
2.00
3.98
2.05
2.11
4.39
f1 (ppm)
S
N
N
+
F
$CO_2^-$
O
F
F
HN
O
O
Cl


**39**

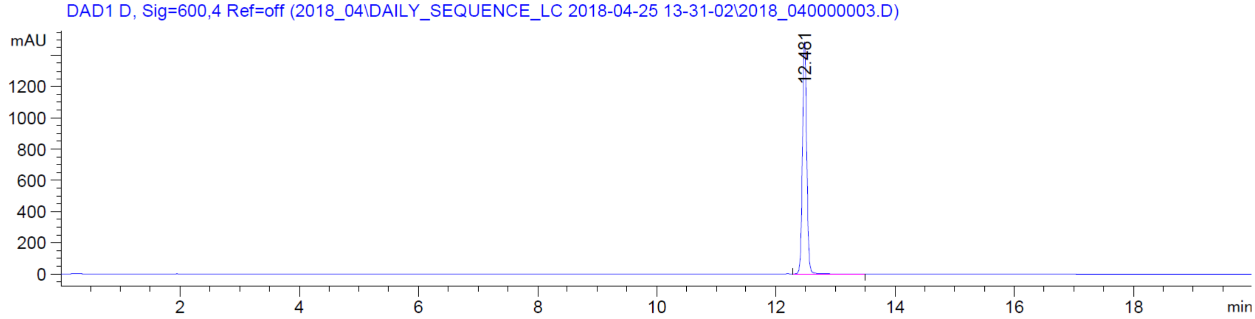

DAD1 D, Sig=600,4 Ref=off (2018_04\DAILY_SEQUENCE_LC 2018-04-25 13-31-02\2018_040000003.D)
mAU
12.481
min


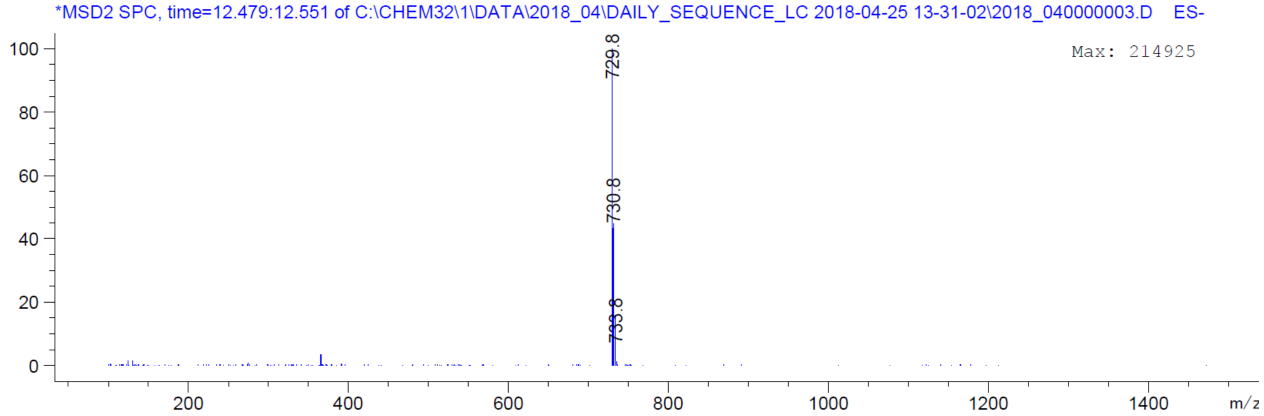

*MSD2 SPC, time=12.479:12.551 of C:\CHEM32\1\DATA\2018_04\DAILY_SEQUENCE_LC 2018-04-25 13-31-02\2018_040000003.D ES-
Max: 214925
729.8
730.8
733.8
m/z

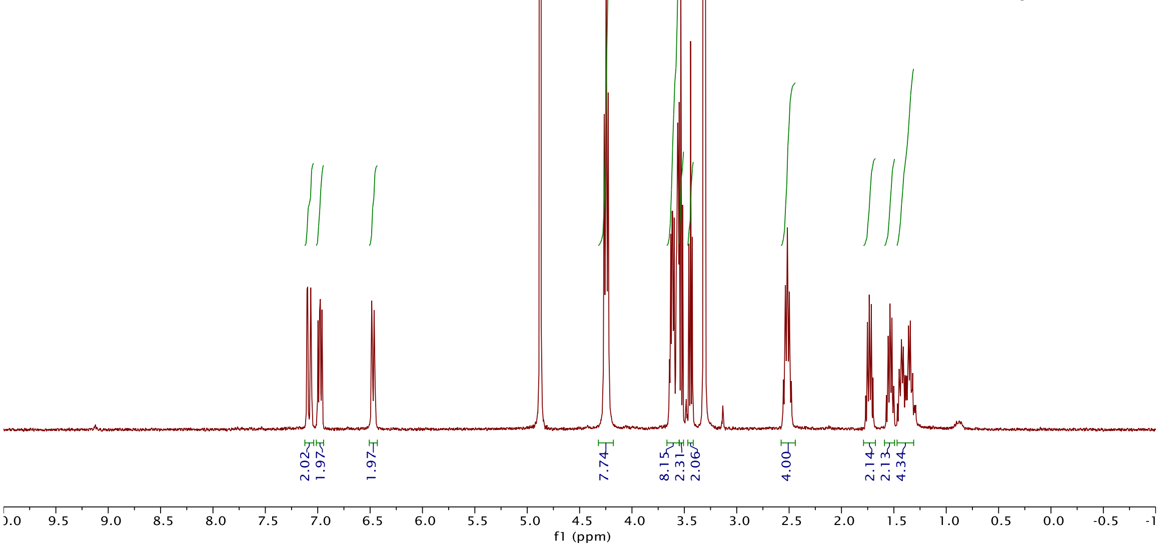
Origin Bruker BioSpin GmbH
Solvent MeOD
Temperature 295.7
Pulse Sequence zg30
Experiment 1D
Number of Scans 16
Acquisition Date 2018-11-08T16:25:00
Spectrometer Frequency 400.13
Spectral Width 8012.8
Lowest Frequency -1543.2
Nucleus 1H
Acquired Size 32768
Spectral Size 65536
f1 (ppm)

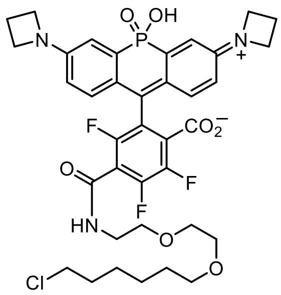

**40**

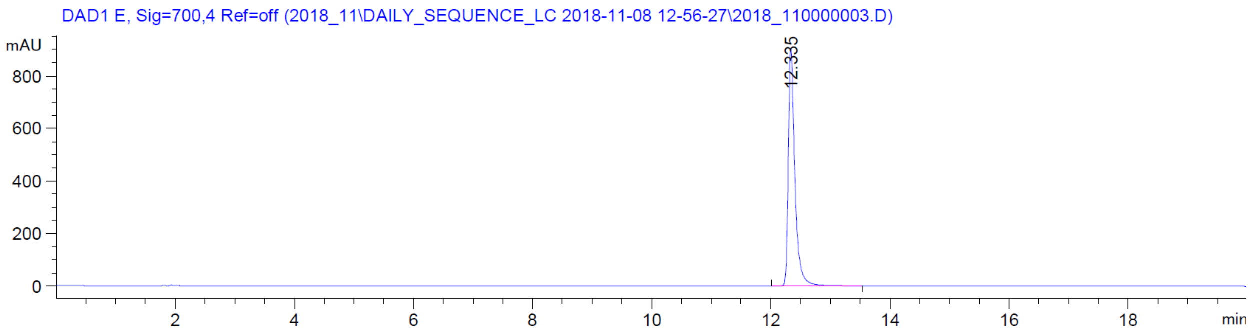
DAD1 E, Sig=700,4 Ref=off (2018_11\DAILY_SEQUENCE_LC 2018-11-08 12-56-27\2018_110000003.D)
mAU
12.335
min

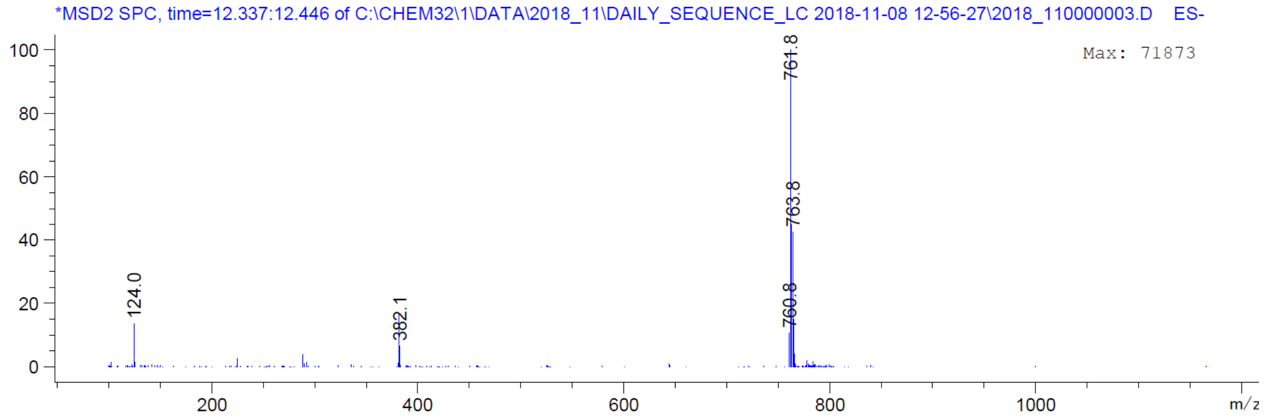
*MSD2 SPC, time=12.337:12.446 of C:\CHEM32\1\DATA\2018_11\DAILY_SEQUENCE_LC 2018-11-08 12-56-27\2018_110000003.D ES-
Max: 71873
761.8
763.8
760.8
124.0
382.1
m/z

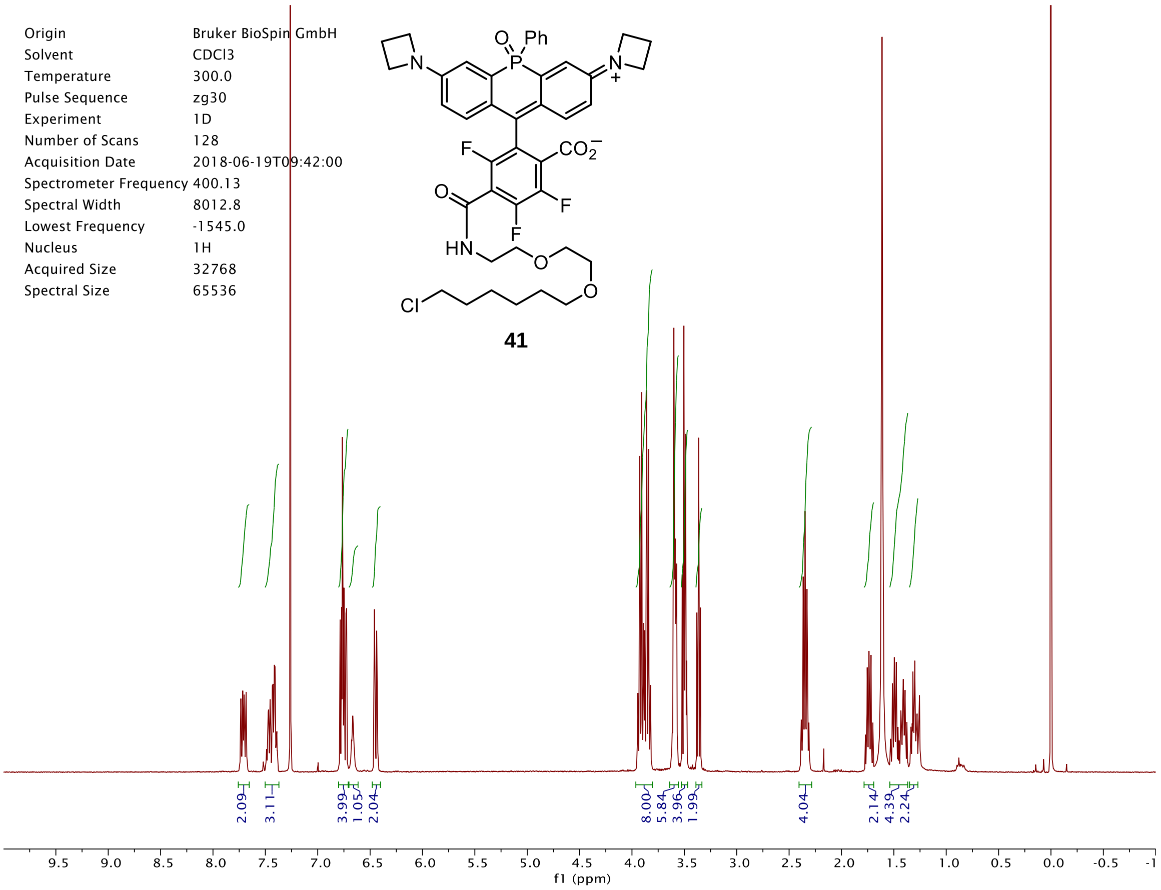
Origin Bruker BioSpin GmbH
Solvent CDCl3
Temperature 300.0
Pulse Sequence zg30
Experiment 1D
Number of Scans 128
Acquisition Date 2018-06-19T09:42:00
Spectrometer Frequency 400.13
Spectral Width 8012.8
Lowest Frequency -1545.0
Nucleus 1H
Acquired Size 32768
Spectral Size 65536
41
2.09
3.11
3.99
1.05
2.04
8.00
5.84
3.96
1.99
4.04
2.14
4.39
2.24
f1 (ppm)

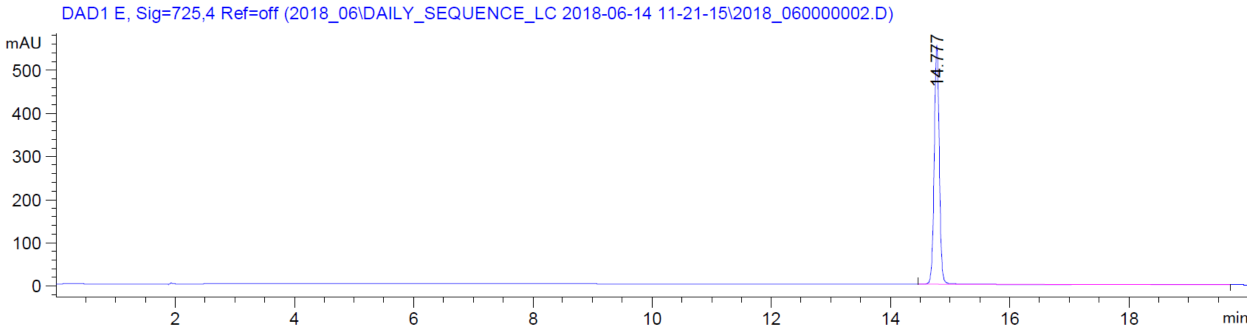
DAD1 E, Sig=725,4 Ref=off (2018_06\DAILY_SEQUENCE_LC 2018-06-14 11-21-15\2018_060000002.D)
mAU
14.777
min

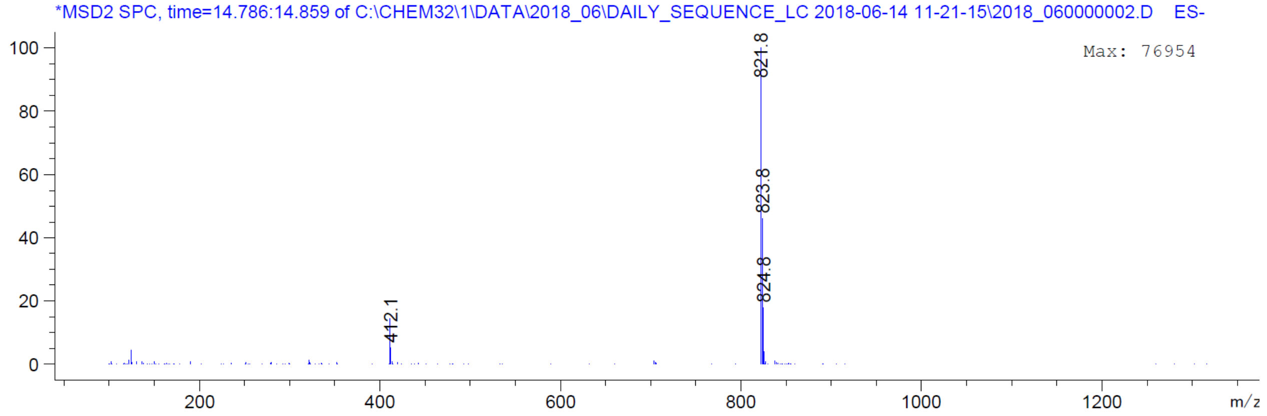
*MSD2 SPC, time=14.786:14.859 of C:\CHEM32\1\DATA\2018_06\DAILY_SEQUENCE_LC 2018-06-14 11-21-15\2018_060000002.D ES-
Max: 76954
821.8
823.8
824.8
412.1
m/z

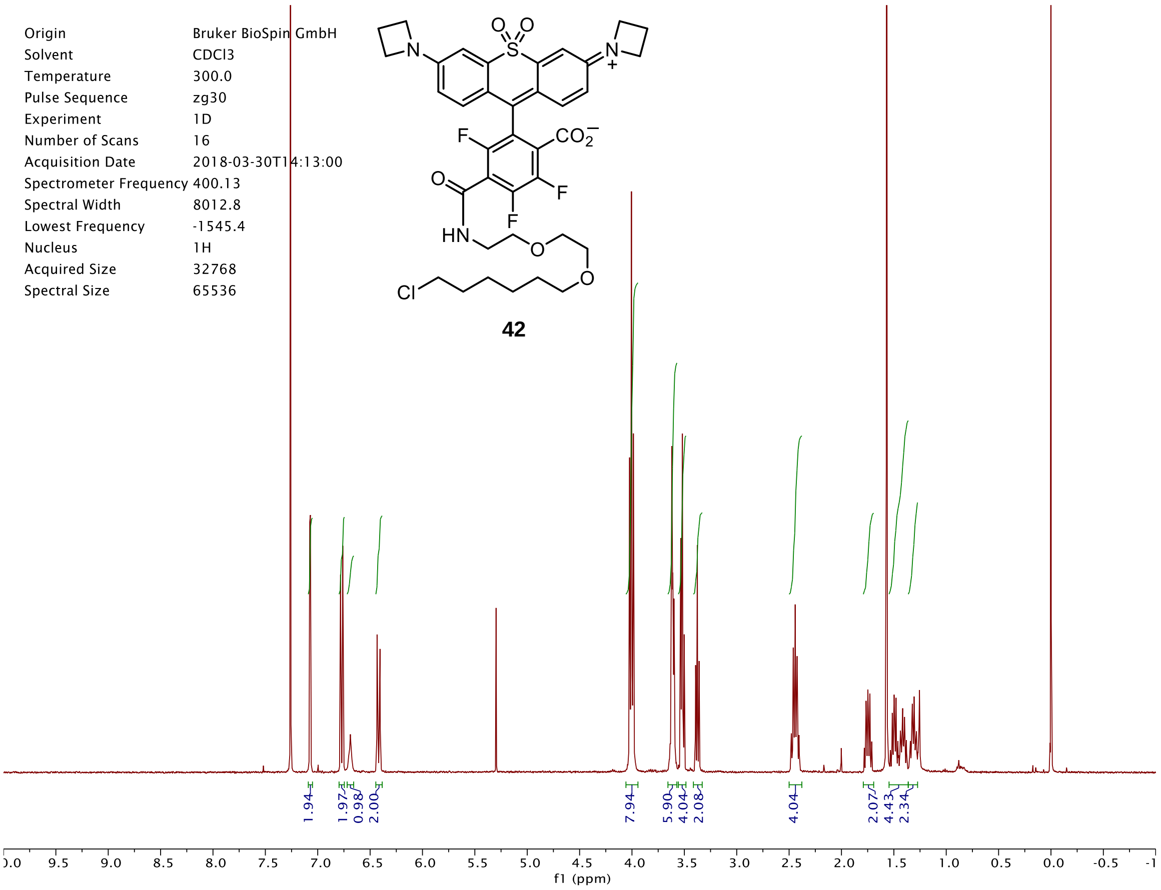

Origin Bruker BioSpin GmbH
Solvent CDCl3
Temperature 300.0
Pulse Sequence zg30
Experiment 1D
Number of Scans 16
Acquisition Date 2018-03-30T14:13:00
Spectrometer Frequency 400.13
Spectral Width 8012.8
Lowest Frequency -1545.4
Nucleus 1H
Acquired Size 32768
Spectral Size 65536
42
1.94
1.97
0.98
2.00
7.94
5.90
4.04
2.08
4.04
2.07
4.43
2.34
f1 (ppm)


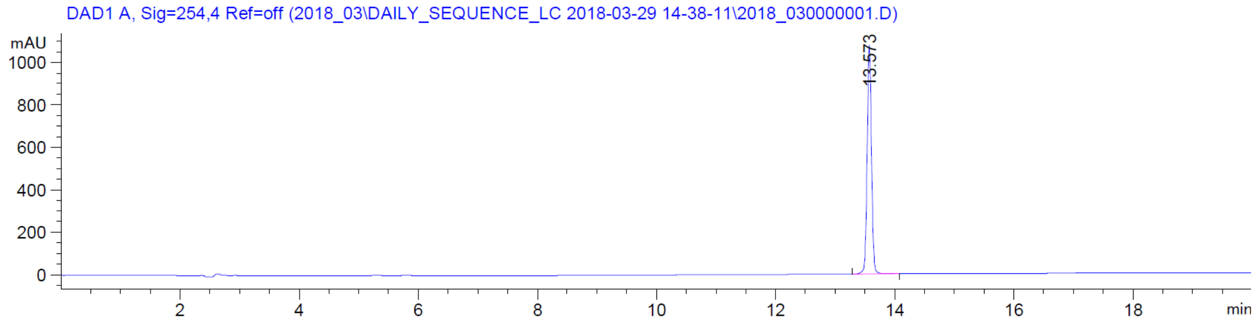

DAD1 A, Sig=254,4 Ref=off (2018_03\DAILY_SEQUENCE_LC 2018-03-29 14-38-11\2018_030000001.D)
mAU
13.573
min


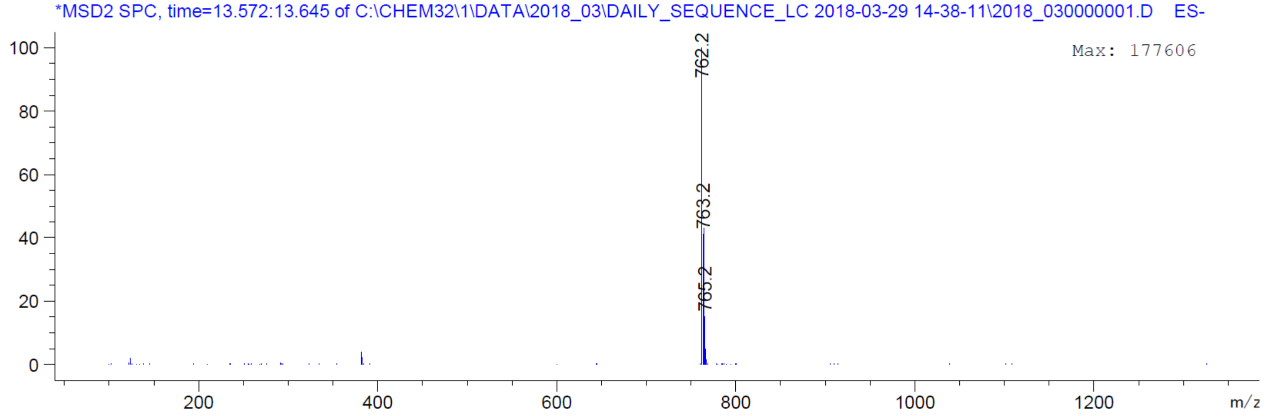

*MSD2 SPC, time=13.572:13.645 of C:\CHEM32\1\DATA\2018_03\DAILY_SEQUENCE_LC 2018-03-29 14-38-11\2018_030000001.D ES-
Max: 177606
762.2
763.2
765.2
m/z

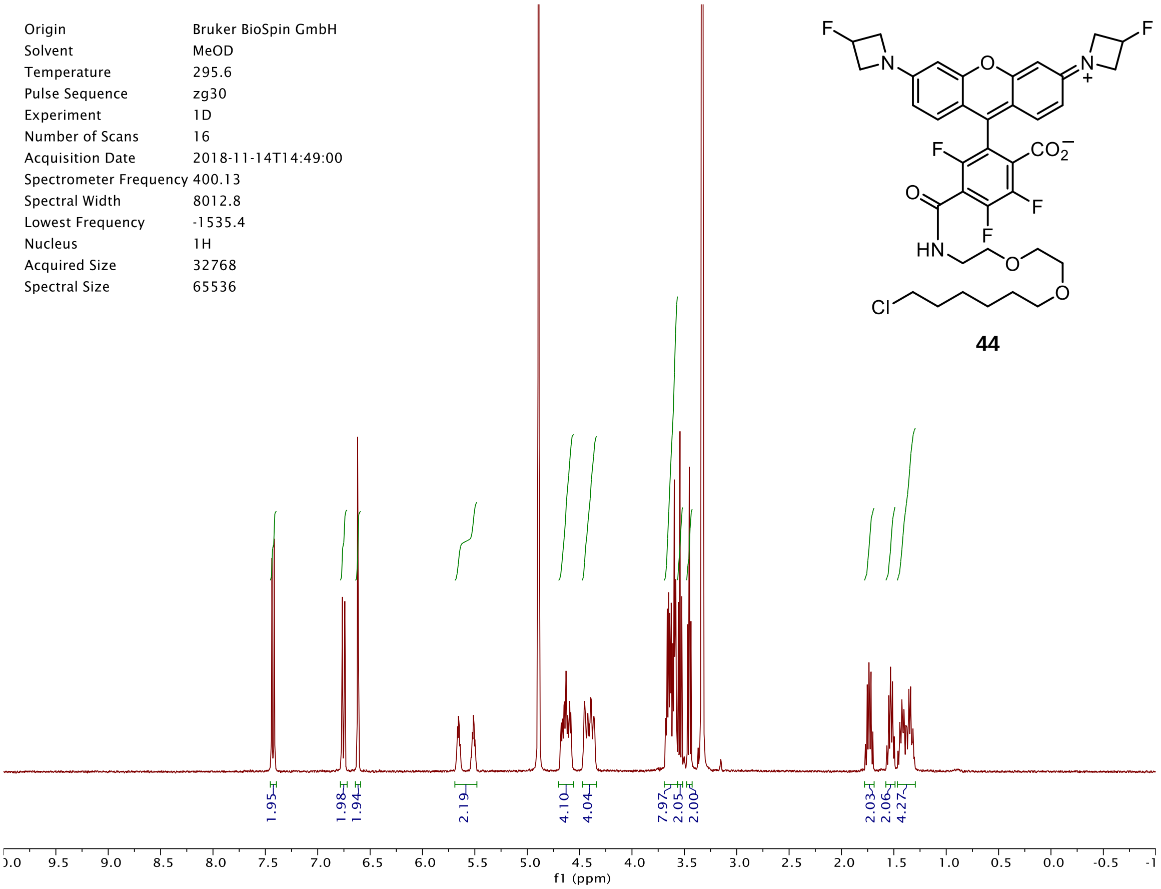
Origin Bruker BioSpin GmbH
Solvent MeOD
Temperature 295.6
Pulse Sequence zg30
Experiment 1D
Number of Scans 16
Acquisition Date 2018-11-14T14:49:00
Spectrometer Frequency 400.13
Spectral Width 8012.8
Lowest Frequency -1535.4
Nucleus 1H
Acquired Size 32768
Spectral Size 65536
1.95
1.98
1.94
2.19
4.10
4.04
7.97
2.05
2.00
2.03
2.06
4.27
f1 (ppm)

**44**

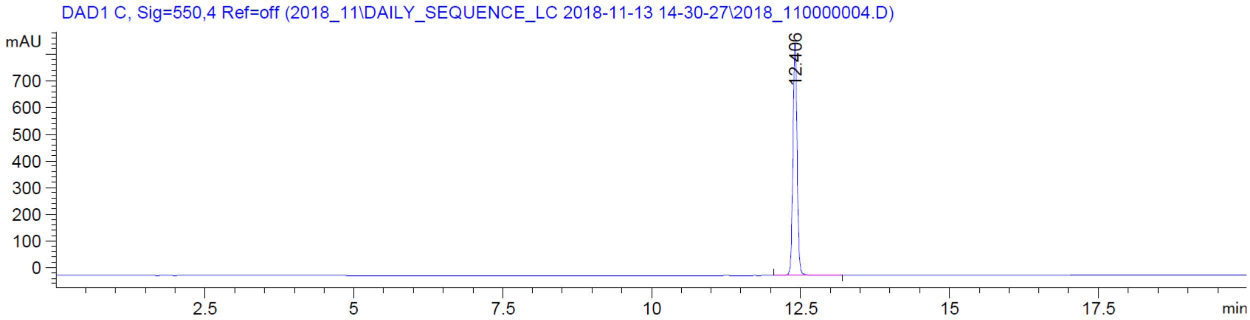
DAD1 C, Sig=550,4 Ref=off (2018_11\DAILY_SEQUENCE_LC 2018-11-13 14-30-27\2018_110000004.D)
mAU
12.406
min

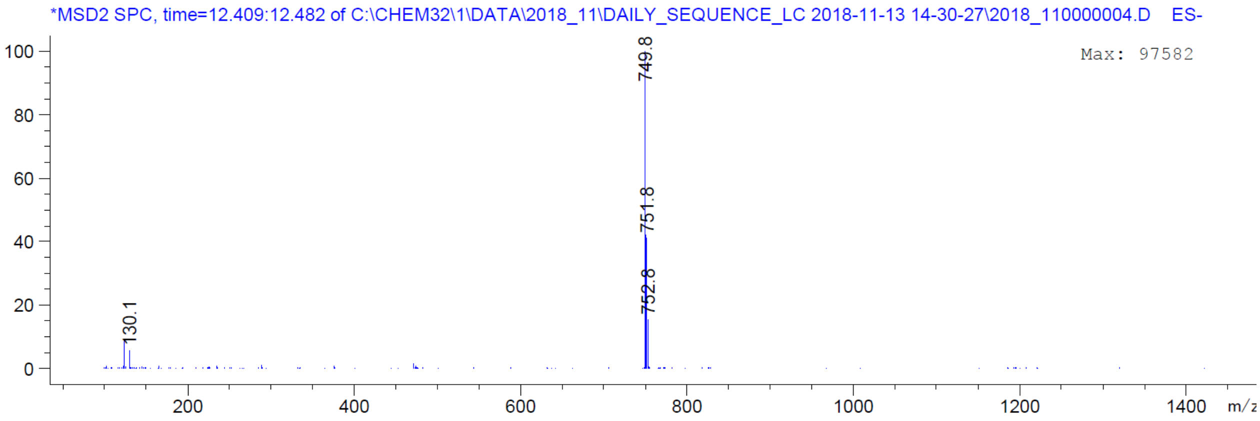
*MSD2 SPC, time=12.409:12.482 of C:\CHEM32\1\DATA\2018_11\DAILY_SEQUENCE_LC 2018-11-13 14-30-27\2018_110000004.D ES-
Max: 97582
749.8
751.8
752.8
130.1
m/z

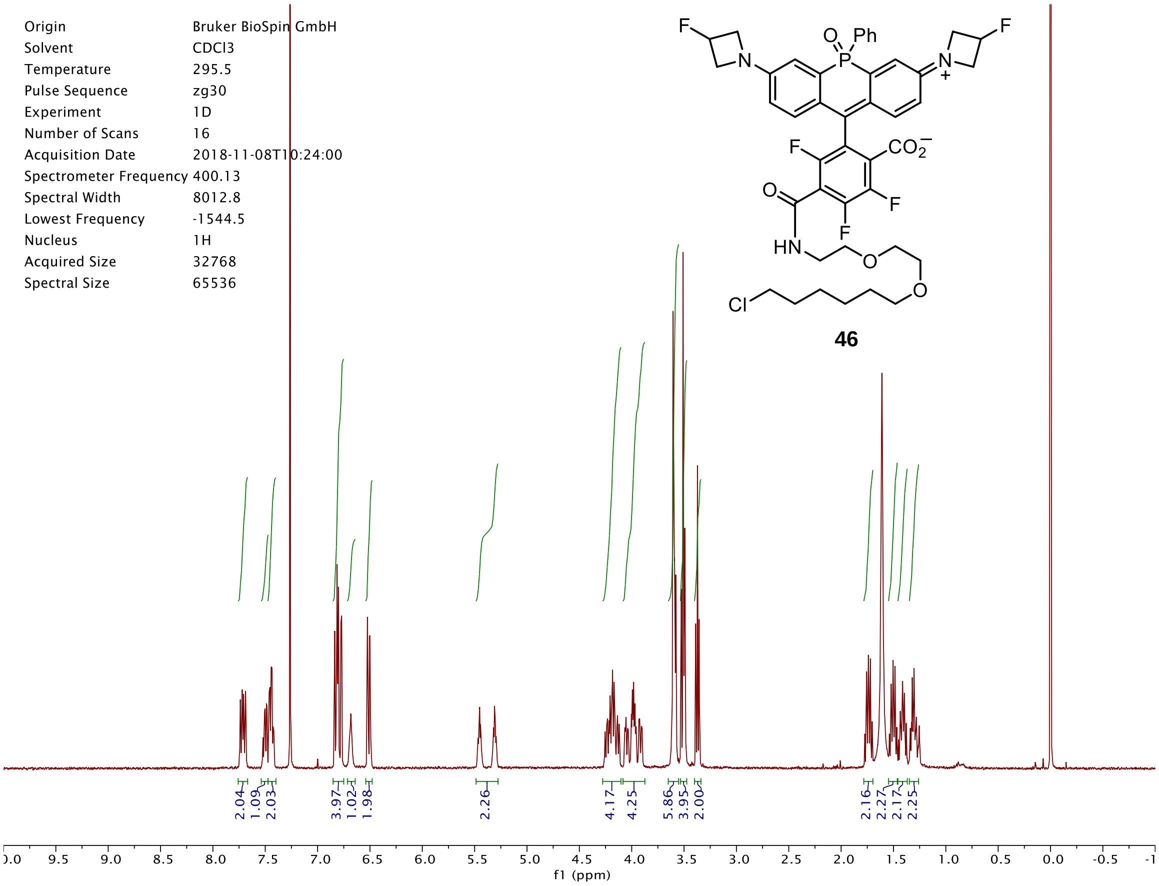
Origin Bruker BioSpin GmbH
Solvent CDCl3
Temperature 295.5
Pulse Sequence zg30
Experiment 1D
Number of Scans 16
Acquisition Date 2018-11-08T10:24:00
Spectrometer Frequency 400.13
Spectral Width 8012.8
Lowest Frequency -1544.5
Nucleus 1H
Acquired Size 32768
Spectral Size 65536
46
2.04 1.09 2.03 3.97 1.02 1.98 2.26 4.17 4.25 5.86 3.95 2.00 2.16 2.27 2.17 2.25
f1 (ppm)

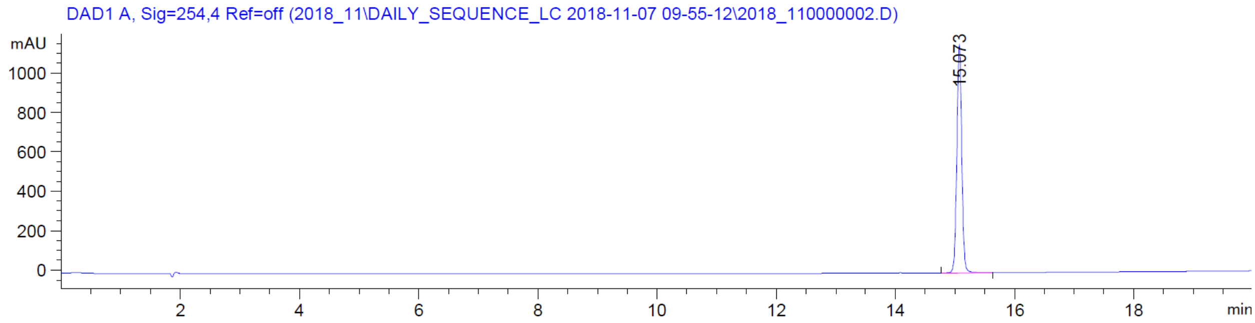
DAD1 A, Sig=254,4 Ref=off (2018_11\DAILY_SEQUENCE_LC 2018-11-07 09-55-12\2018_110000002.D)
mAU
15.073
min

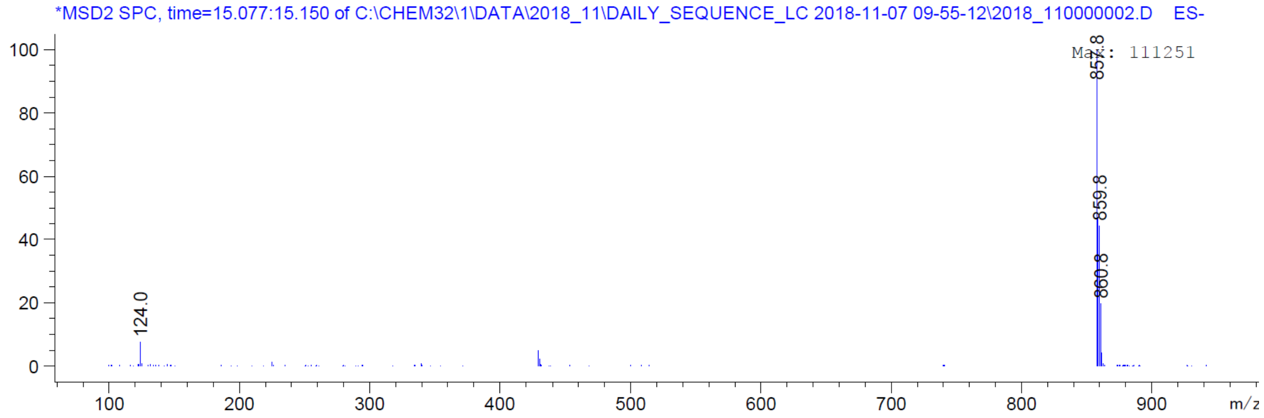
*MSD2 SPC, time=15.077:15.150 of C:\CHEM32\1\DATA\2018_11\DAILY_SEQUENCE_LC 2018-11-07 09-55-12\2018_110000002.D ES-
Max: 111251
857.8
859.8
860.8
124.0
m/z

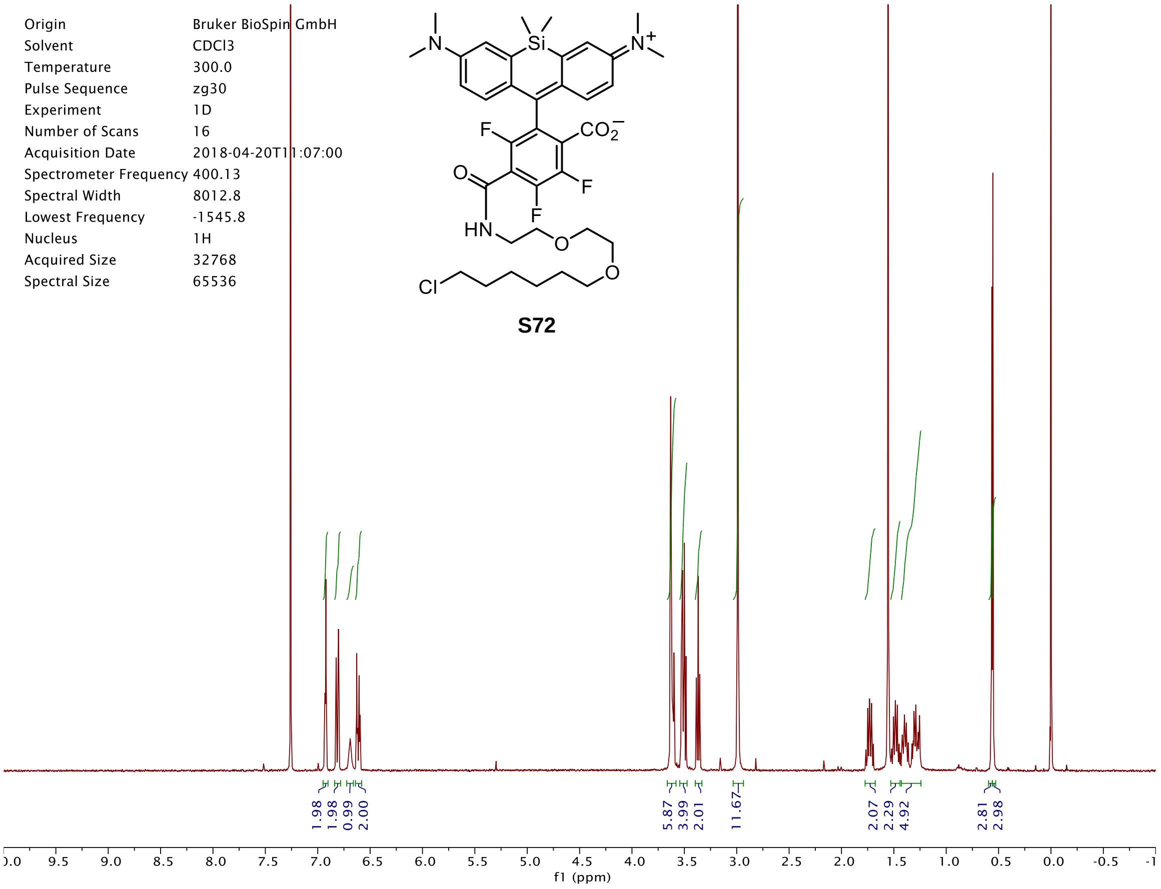

Origin Bruker BioSpin GmbH
Solvent CDCl3
Temperature 300.0
Pulse Sequence zg30
Experiment 1D
Number of Scans 16
Acquisition Date 2018-04-20T11:07:00
Spectrometer Frequency 400.13
Spectral Width 8012.8
Lowest Frequency -1545.8
Nucleus 1H
Acquired Size 32768
Spectral Size 65536
S72
1.98 1.98 0.99 2.00
5.87 3.99 2.01
11.67
2.07 2.29 4.92
2.81 2.98
f1 (ppm)


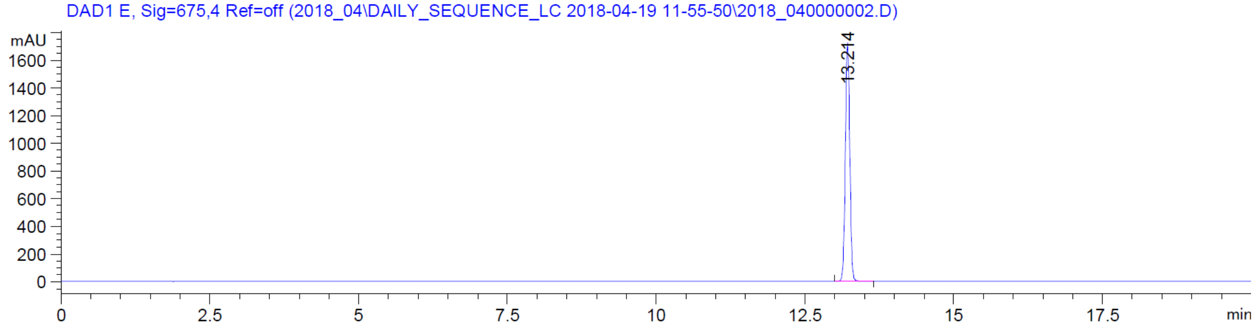

DAD1 E, Sig=675,4 Ref=off (2018_04\DAILY_SEQUENCE_LC 2018-04-19 11-55-50\2018_040000002.D)
mAU
13.214
min


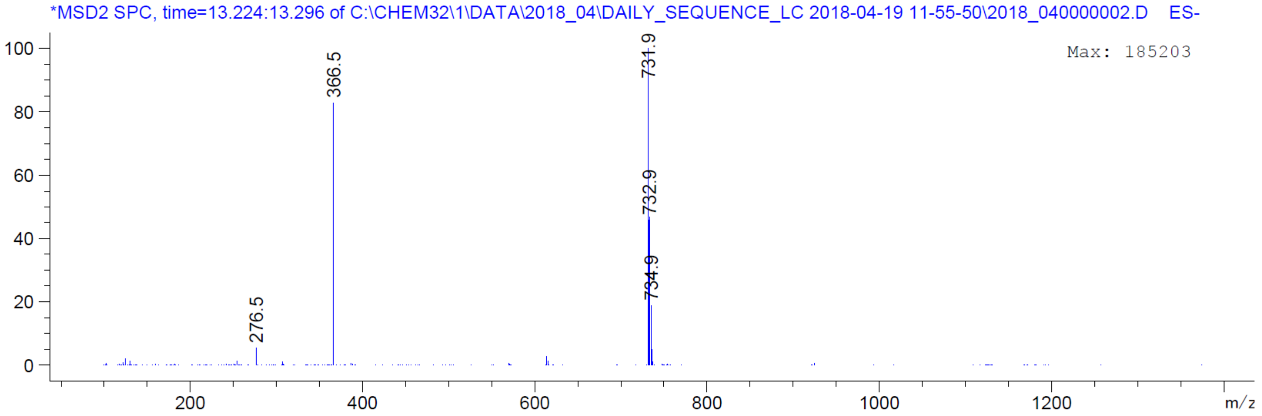

*MSD2 SPC, time=13.224:13.296 of C:\CHEM32\1\DATA\2018_04\DAILY_SEQUENCE_LC 2018-04-19 11-55-50\2018_040000002.D ES-
Max: 185203
276.5
366.5
731.9
732.9
734.9
m/z

| | |
|---|---|
| Origin | Bruker BioSpin GmbH |
| Solvent | MeOD |
| Temperature | 300.0 |
| Pulse Sequence | zg30 |
| Experiment | 1D |
| Number of Scans | 16 |
| Acquisition Date | 2018-04-20T10:58:00 |
| Spectrometer Frequency | 400.13 |
| Spectral Width | 8012.8 |
| Lowest Frequency | -1543.0 |
| Nucleus | 1H |
| Acquired Size | 32768 |
| Spectral Size | 65536 |

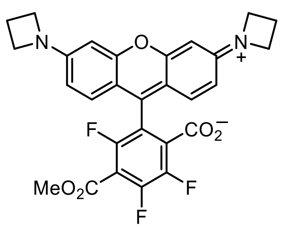


**S73**

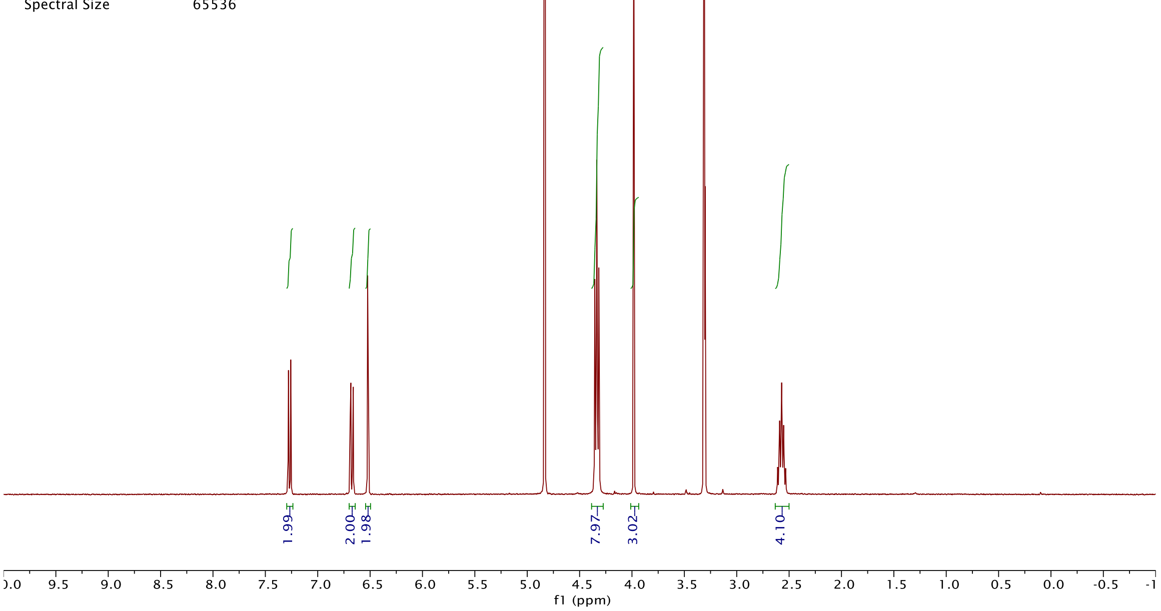


DAD1 C, Sig=575,4 Ref=off (2018_04\DAILY_SEQUENCE_LC 2018-04-18 14-26-38\2018_040000007.D)

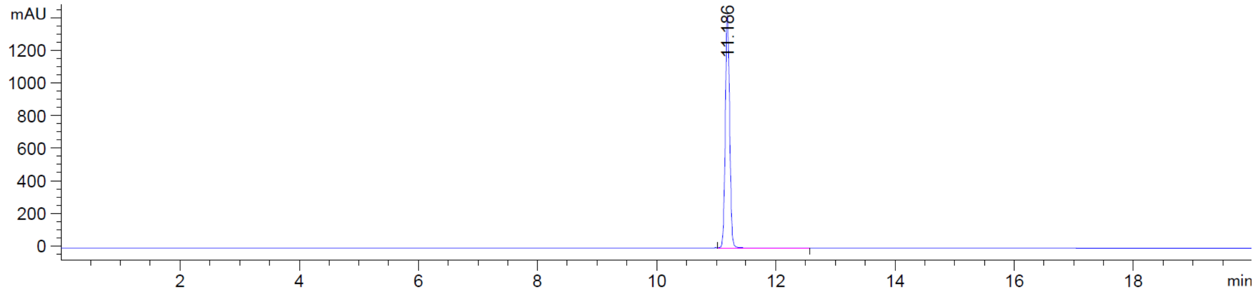


*MSD2 SPC, time=11.192:11.265 of C:\CHEM32\1\DATA\2018_04\DAILY_SEQUENCE_LC 2018-04-18 14-26-38\2018_040000007.D ES-

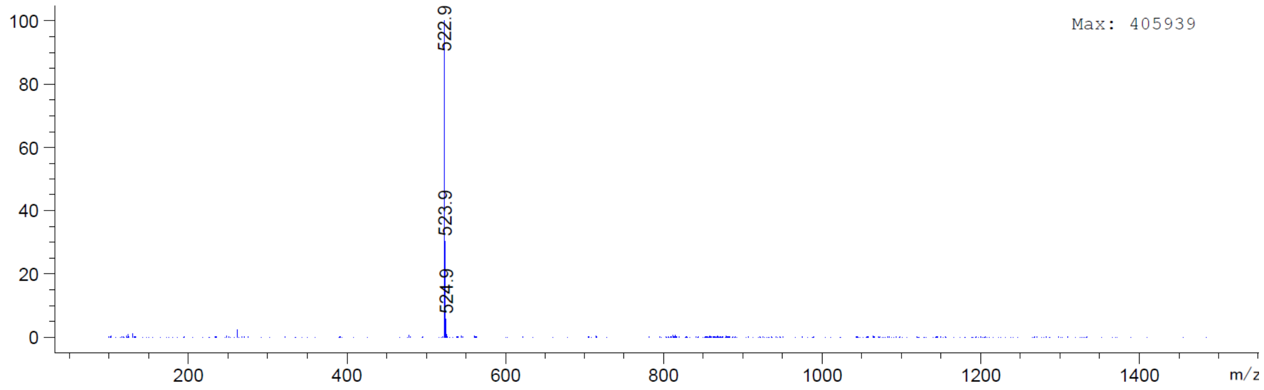

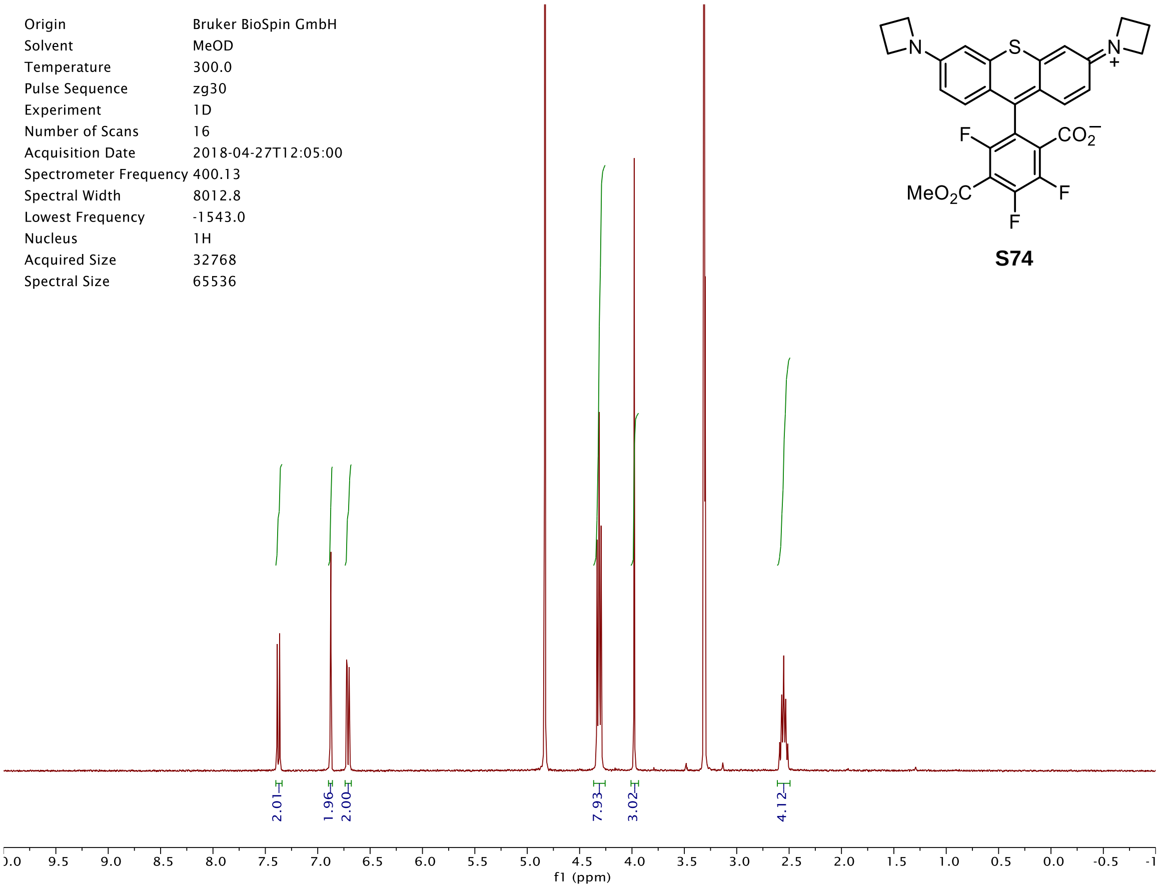
N
S
N
+
F
$CO_2^-$
$MeO_2C$
F
F
S74
Origin Bruker BioSpin GmbH
Solvent MeOD
Temperature 300.0
Pulse Sequence zg30
Experiment 1D
Number of Scans 16
Acquisition Date 2018-04-27T12:05:00
Spectrometer Frequency 400.13
Spectral Width 8012.8
Lowest Frequency -1543.0
Nucleus 1H
Acquired Size 32768
Spectral Size 65536
2.01
1.96
2.00
7.93
3.02
4.12
f1 (ppm)

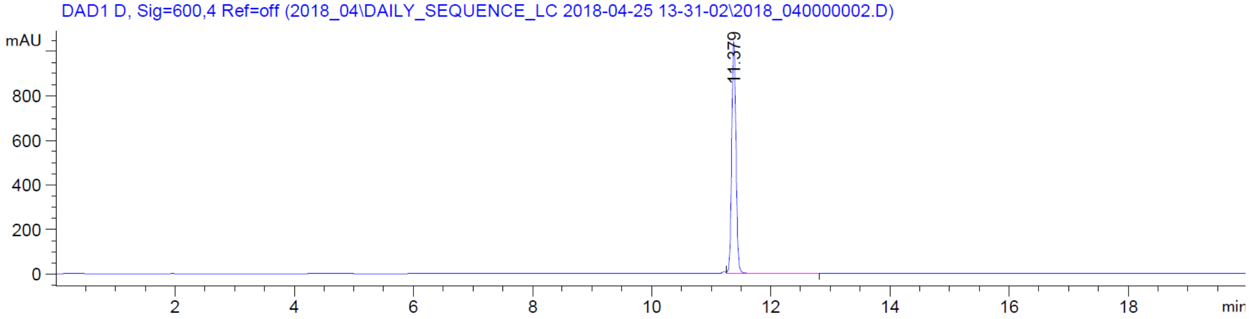
DAD1 D, Sig=600,4 Ref=off (2018_04\DAILY_SEQUENCE_LC 2018-04-25 13-31-02\2018_040000002.D)
mAU
11.379
min

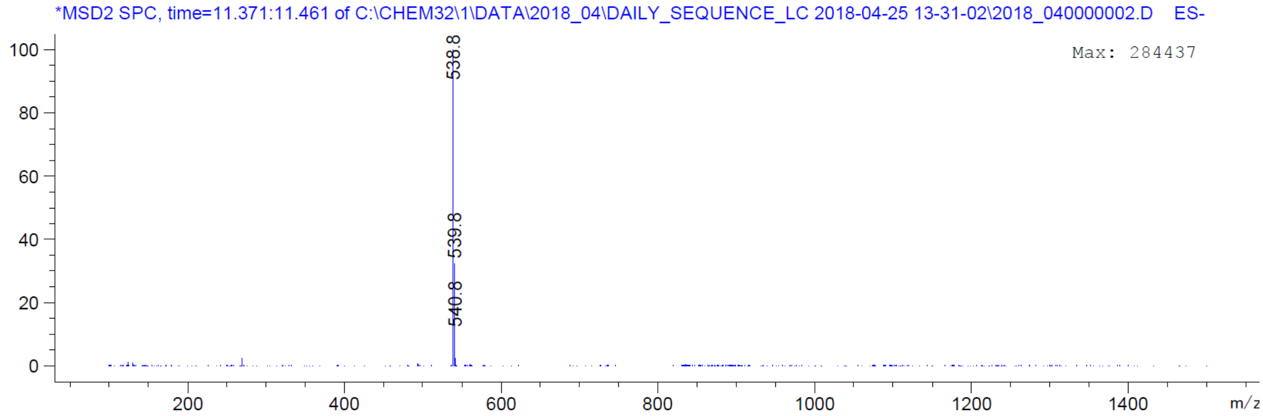
*MSD2 SPC, time=11.371:11.461 of C:\CHEM32\1\DATA\2018_04\DAILY_SEQUENCE_LC 2018-04-25 13-31-02\2018_040000002.D ES-
Max: 284437
538.8
539.8
540.8
m/z

| | |
|---|---|
| Origin | Bruker BioSpin GmbH |
| Solvent | CDCl3 |
| Temperature | 300.0 |
| Pulse Sequence | zg30 |
| Experiment | 1D |
| Number of Scans | 16 |
| Acquisition Date | 2018-02-20T13:50:00 |
| Spectrometer Frequency | 400.13 |
| Spectral Width | 8012.8 |
| Lowest Frequency | -1546.1 |
| Nucleus | 1H |
| Acquired Size | 32768 |
| Spectral Size | 65536 |

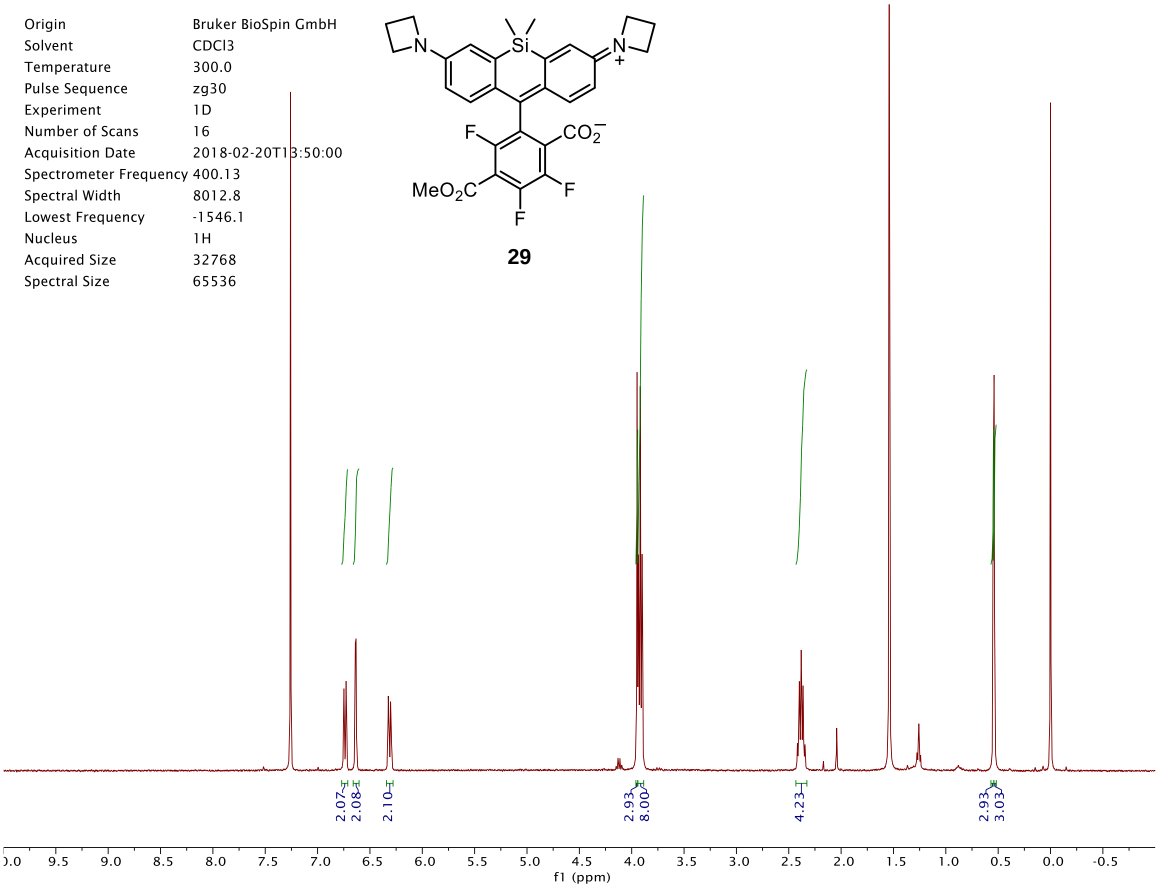


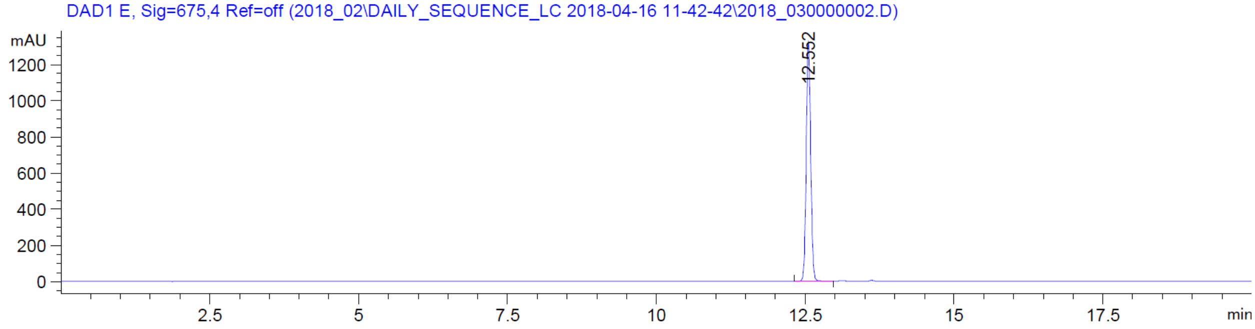


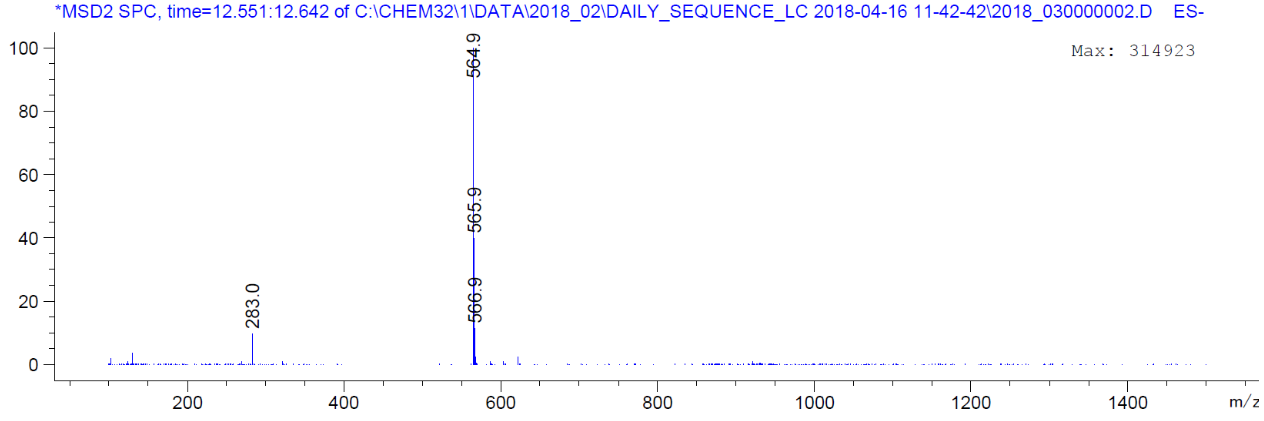

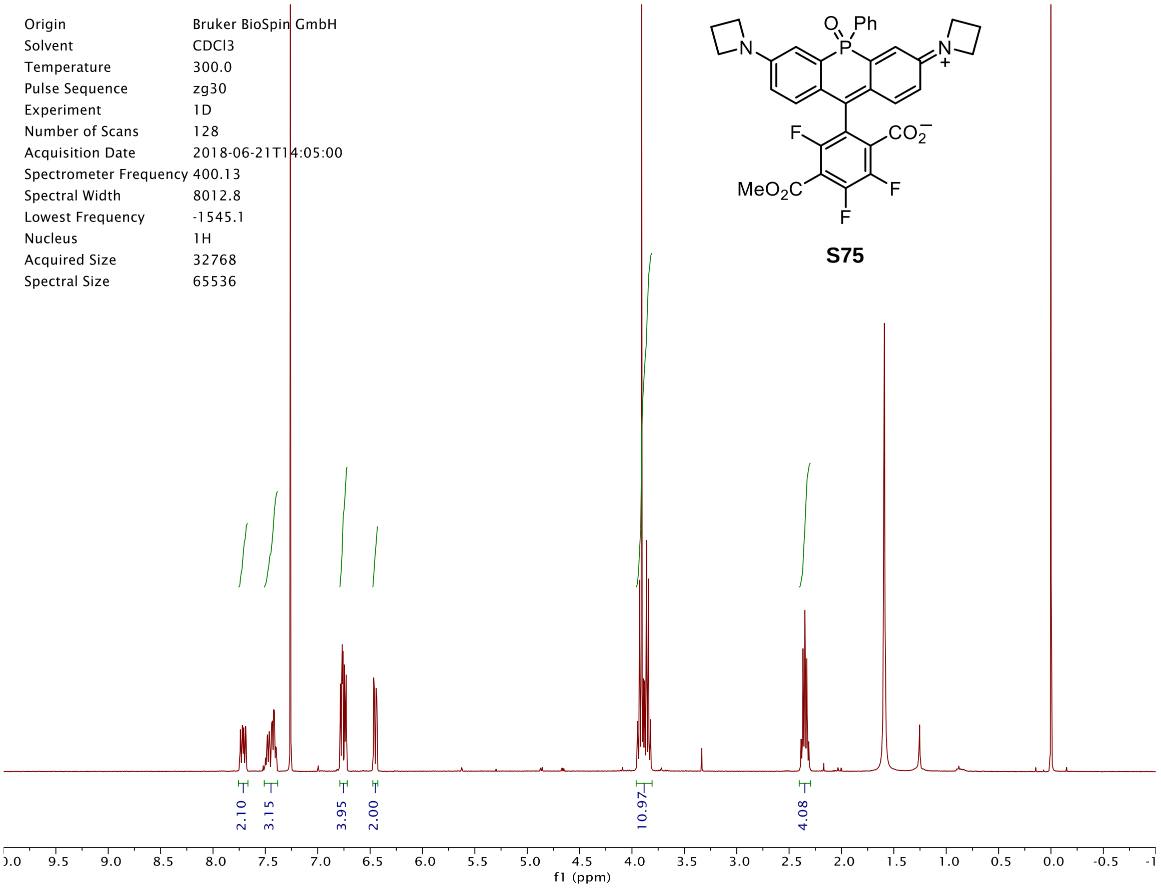
Origin Bruker BioSpin GmbH
Solvent CDCl3
Temperature 300.0
Pulse Sequence zg30
Experiment 1D
Number of Scans 128
Acquisition Date 2018-06-21T14:05:00
Spectrometer Frequency 400.13
Spectral Width 8012.8
Lowest Frequency -1545.1
Nucleus 1H
Acquired Size 32768
Spectral Size 65536
S75
2.10
3.15
3.95
2.00
10.97
4.08
f1 (ppm)

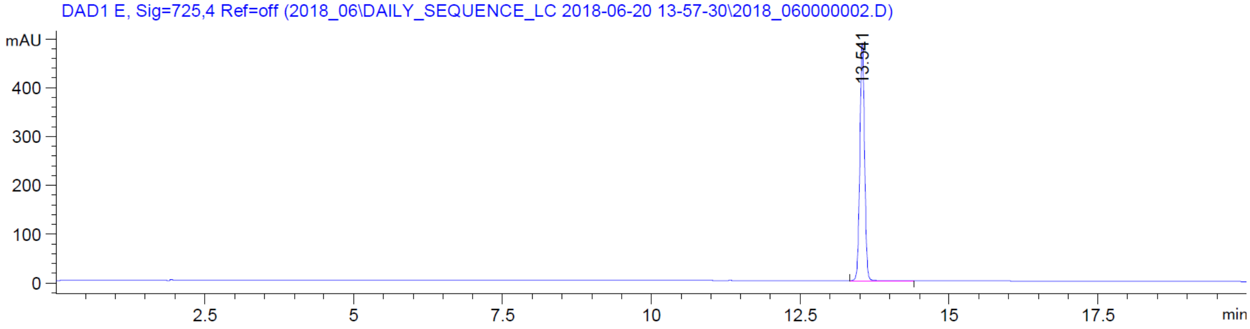
DAD1 E, Sig=725,4 Ref=off (2018_06\DAILY_SEQUENCE_LC 2018-06-20 13-57-30\2018_060000002.D)
mAU
13.541
min

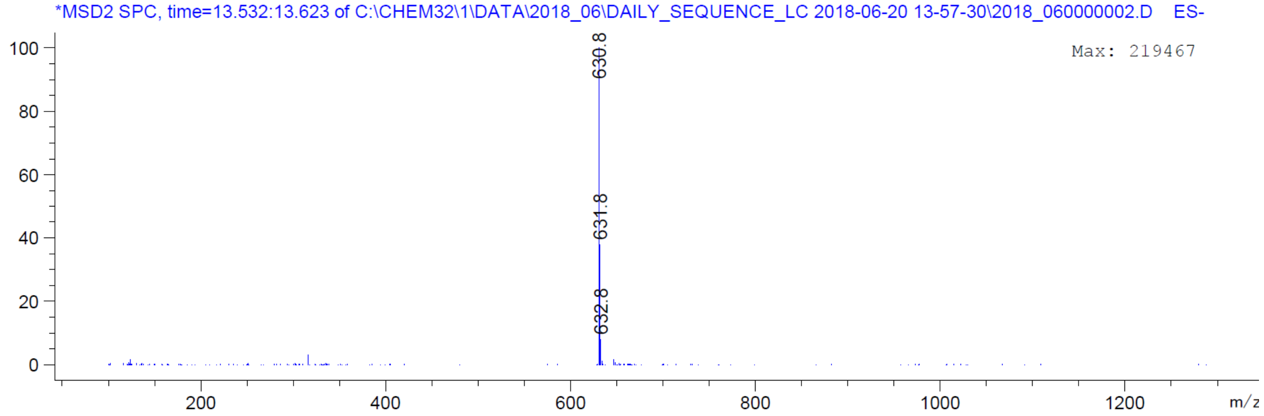
*MSD2 SPC, time=13.532:13.623 of C:\CHEM32\1\DATA\2018_06\DAILY_SEQUENCE_LC 2018-06-20 13-57-30\2018_060000002.D ES-
Max: 219467
630.8
631.8
632.8
m/z

| | |
|---|---|
| Origin | Bruker BioSpin GmbH |
| Solvent | CDCl3 |
| Temperature | 300.0 |
| Pulse Sequence | zg30 |
| Experiment | 1D |
| Number of Scans | 128 |
| Acquisition Date | 2018-06-21T13:51:00 |
| Spectrometer Frequency | 400.13 |
| Spectral Width | 8012.8 |
| Lowest Frequency | -1545.1 |
| Nucleus | 1H |
| Acquired Size | 32768 |
| Spectral Size | 65536 |

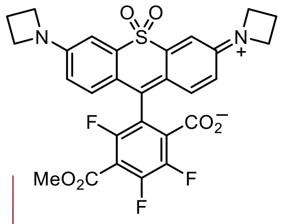


**S76**

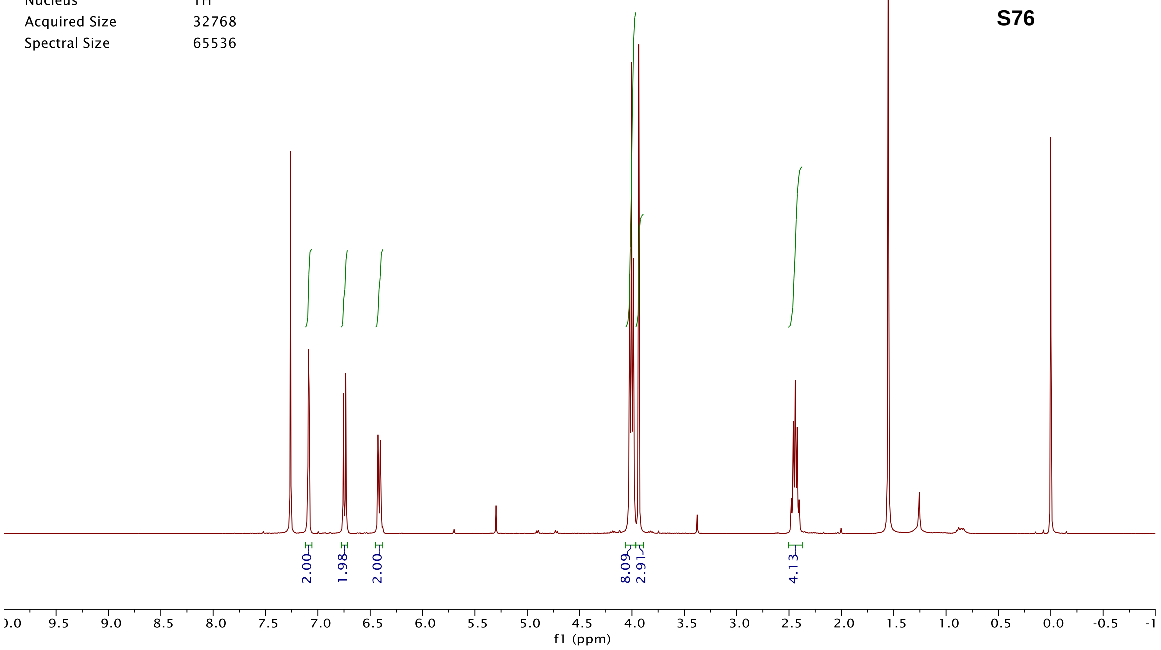


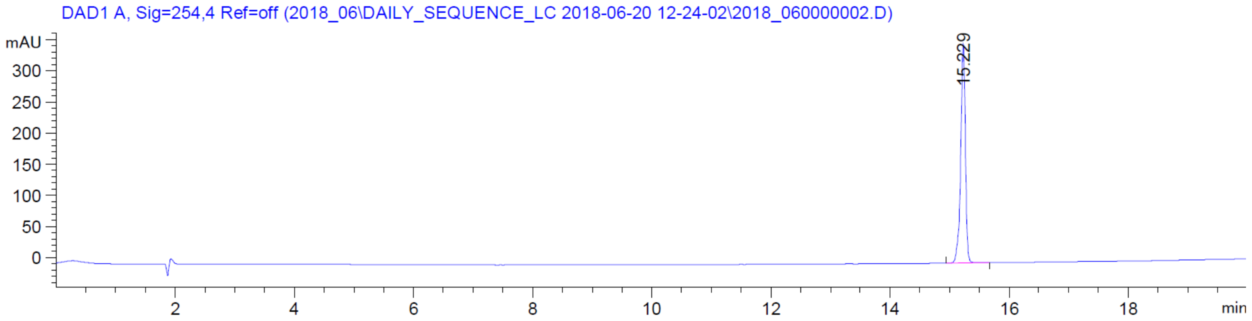


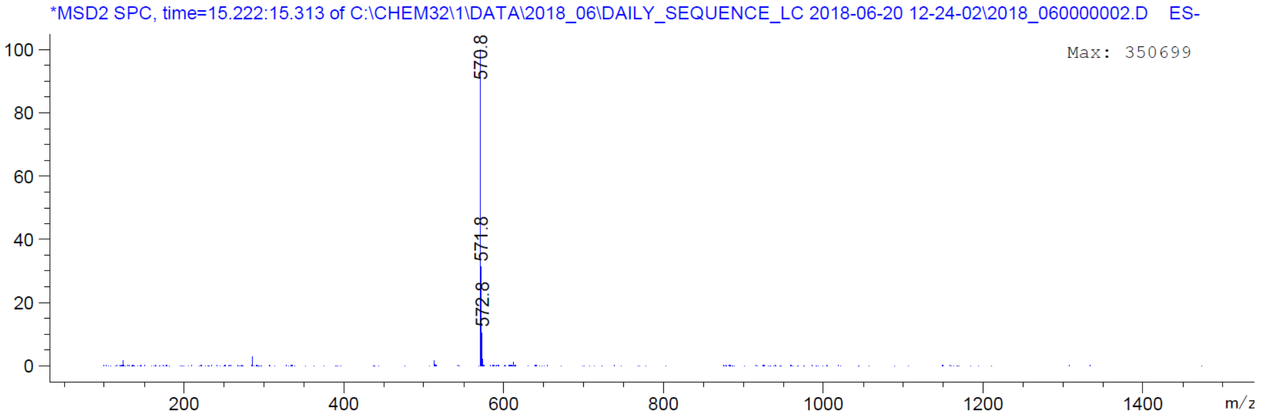

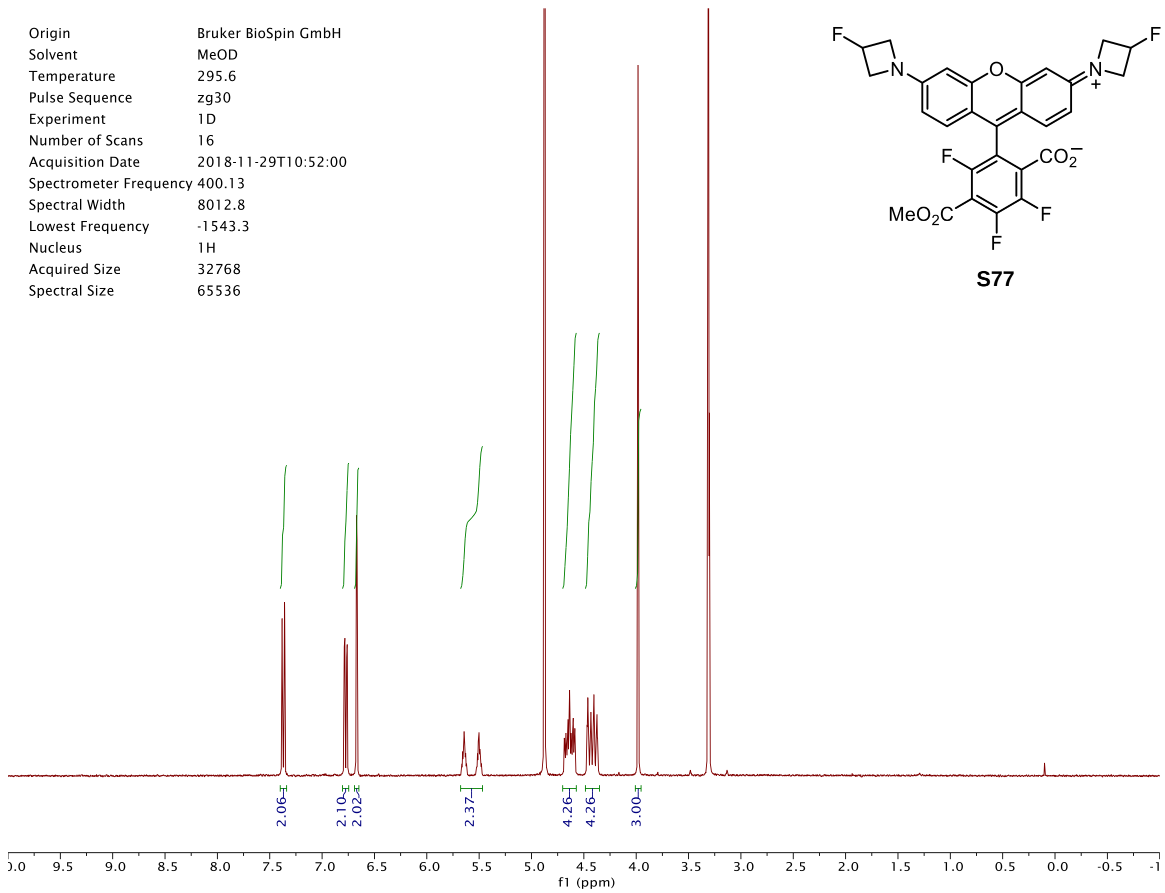

Origin Bruker BioSpin GmbH
Solvent MeOD
Temperature 295.6
Pulse Sequence zg30
Experiment 1D
Number of Scans 16
Acquisition Date 2018-11-29T10:52:00
Spectrometer Frequency 400.13
Spectral Width 8012.8
Lowest Frequency -1543.3
Nucleus 1H
Acquired Size 32768
Spectral Size 65536
2.06
2.10
2.02
2.37
4.26
4.26
3.00
f1 (ppm)


S77

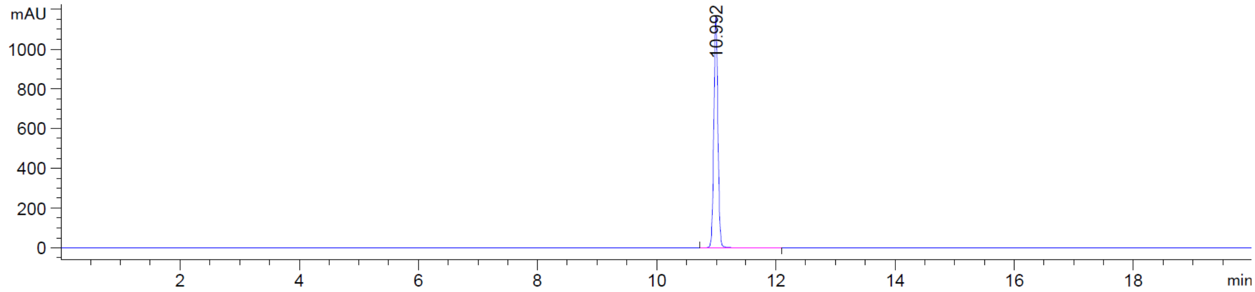

DAD1 C, Sig=550,4 Ref=off (2018_11\DAILY_SEQUENCE_LC 2018-11-27 10-13-33\2018_110000003.D)
mAU
10.992
min


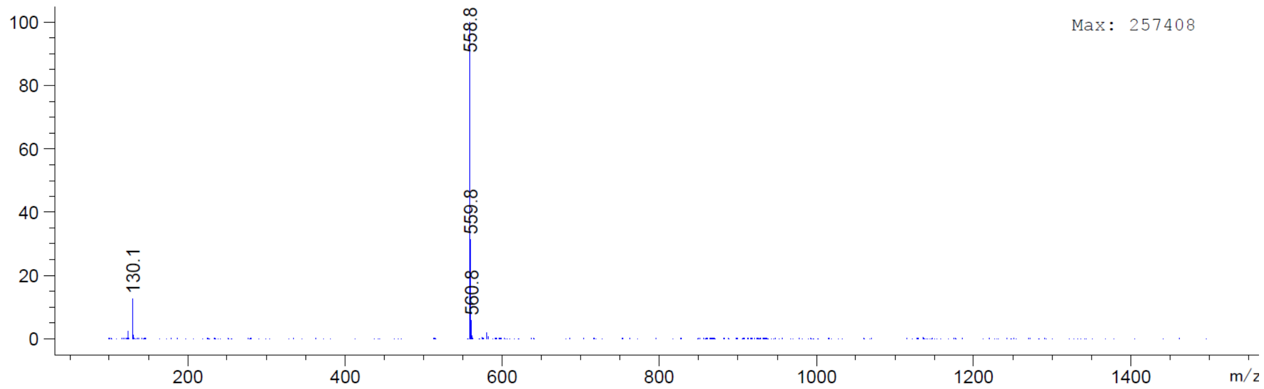

*MSD2 SPC, time=10.989:11.080 of C:\CHEM32\1\DATA\2018_11\DAILY_SEQUENCE_LC 2018-11-27 10-13-33\2018_110000003.D ES-
Max: 257408
558.8
559.8
560.8
130.1
m/z

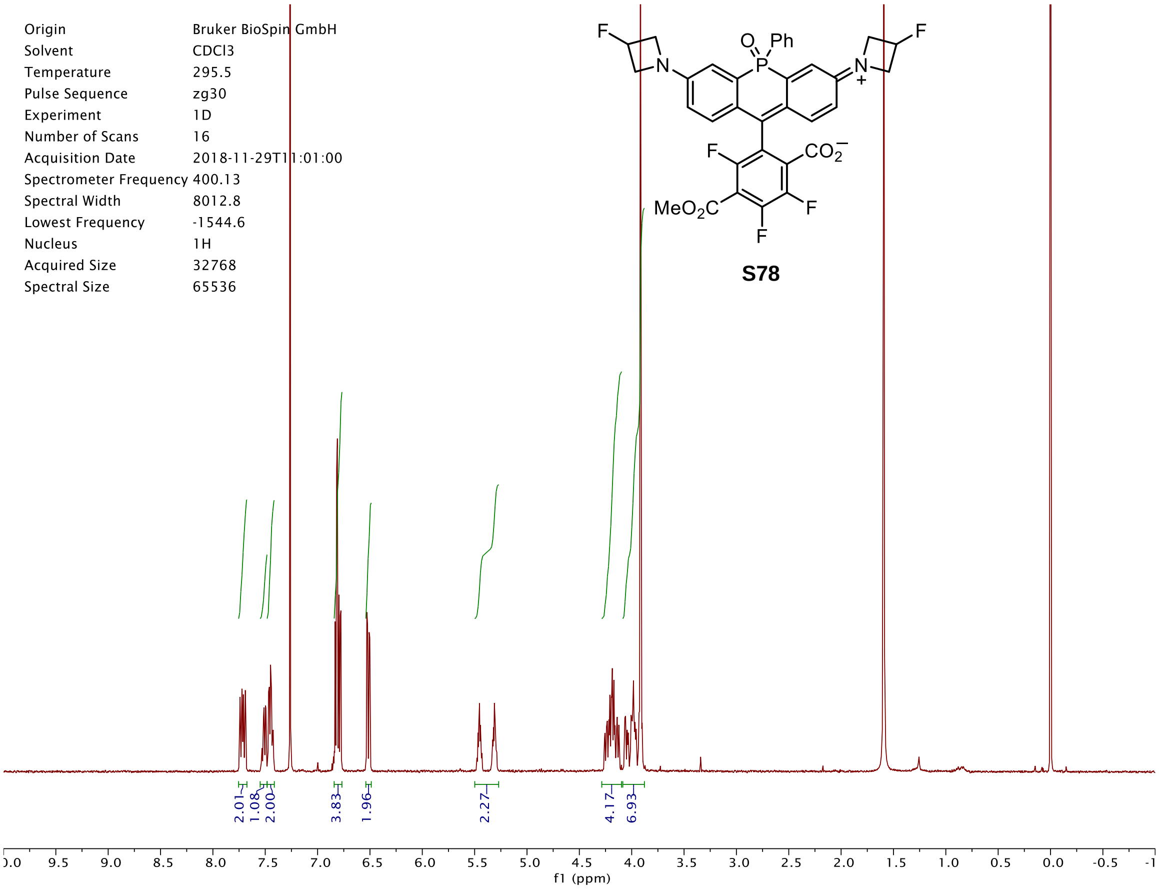

Origin Bruker BioSpin GmbH
Solvent CDCl3
Temperature 295.5
Pulse Sequence zg30
Experiment 1D
Number of Scans 16
Acquisition Date 2018-11-29T11:01:00
Spectrometer Frequency 400.13
Spectral Width 8012.8
Lowest Frequency -1544.6
Nucleus 1H
Acquired Size 32768
Spectral Size 65536
S78
2.01 1.08 2.00 3.83 1.96 2.27 4.17 6.93
f1 (ppm)


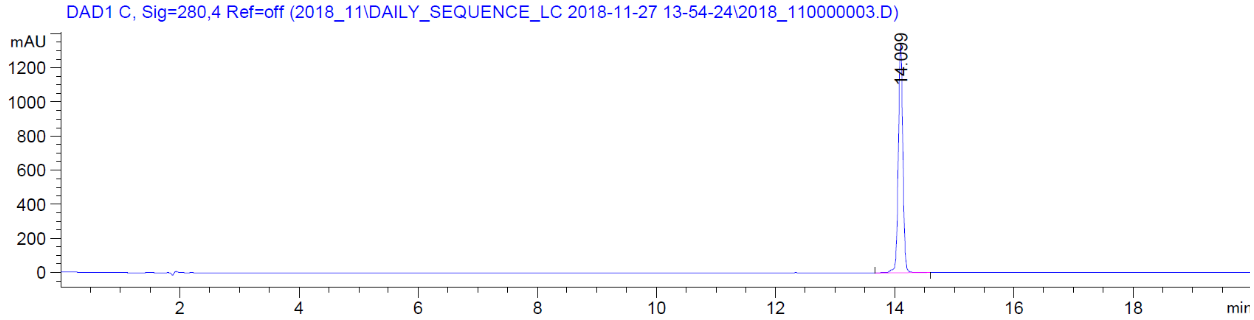

DAD1 C, Sig=280,4 Ref=off (2018_11\DAILY_SEQUENCE_LC 2018-11-27 13-54-24\2018_110000003.D)
mAU
14.099
min


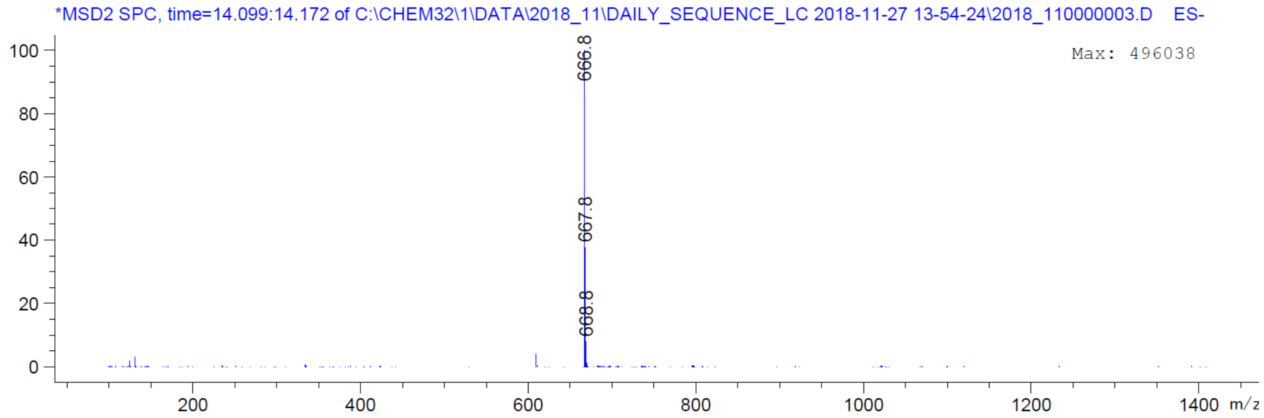

*MSD2 SPC, time=14.099:14.172 of C:\CHEM32\1\DATA\2018_11\DAILY_SEQUENCE_LC 2018-11-27 13-54-24\2018_110000003.D ES-
Max: 496038
666.8
667.8
668.8
m/z

| | |
|---|---|
| Origin | Bruker BioSpin GmbH |
| Solvent | CDCl3 |
| Temperature | 300.0 |
| Pulse Sequence | zg30 |
| Experiment | 1D |
| Number of Scans | 16 |
| Acquisition Date | 2018-05-18T16:46:00 |
| Spectrometer Frequency | 400.13 |
| Spectral Width | 8012.8 |
| Lowest Frequency | -1546.2 |
| Nucleus | 1H |
| Acquired Size | 32768 |
| Spectral Size | 65536 |

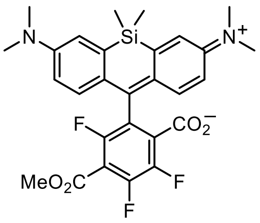


**S79**

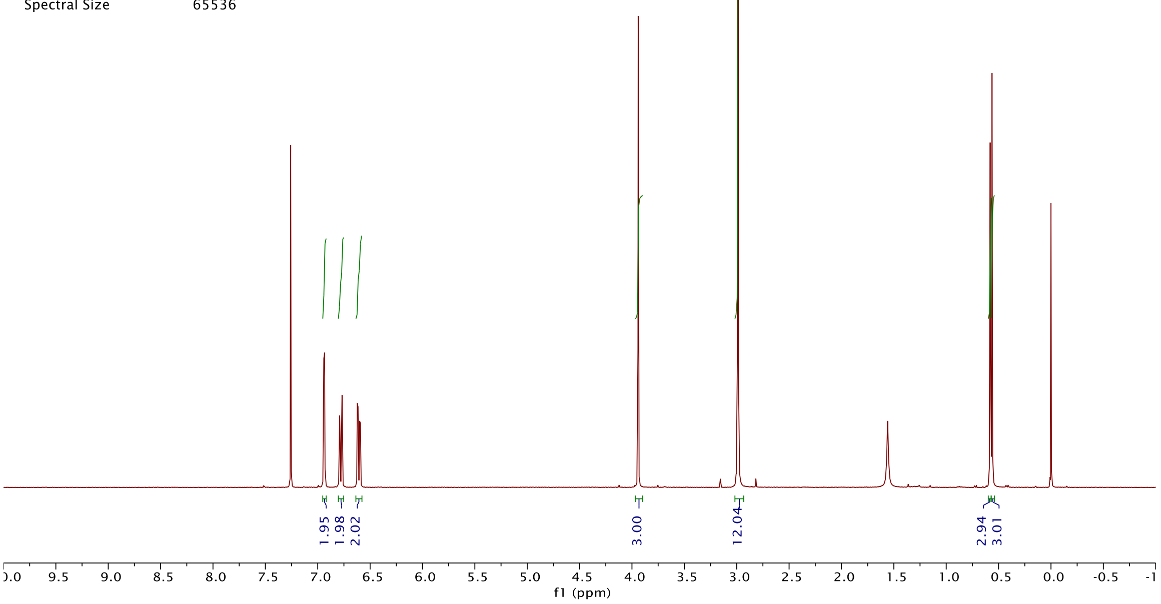


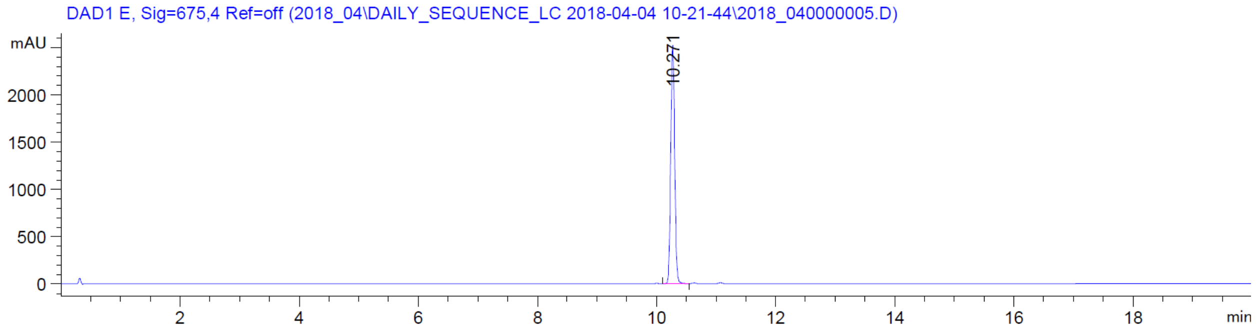


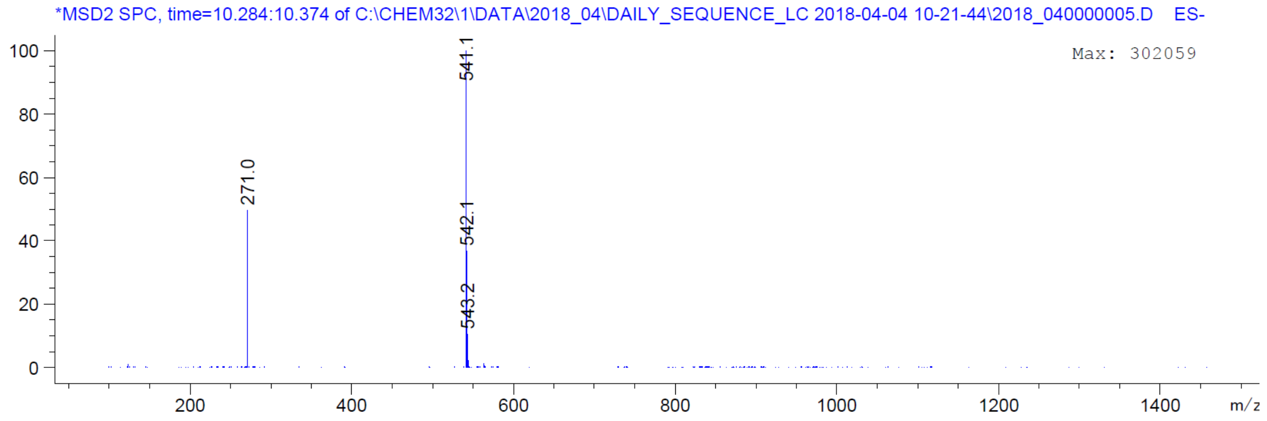

| | |
|---|---|
| Origin | Bruker BioSpin GmbH |
| Solvent | CDCl3 |
| Temperature | 300.0 |
| Pulse Sequence | zg30 |
| Experiment | 1D |
| Number of Scans | 16 |
| Acquisition Date | 2018-10-26T15:43:00 |
| Spectrometer Frequency | 400.13 |
| Spectral Width | 8012.8 |
| Lowest Frequency | -1546.2 |
| Nucleus | 1H |
| Acquired Size | 32768 |
| Spectral Size | 65536 |

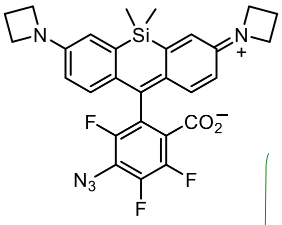


**21**

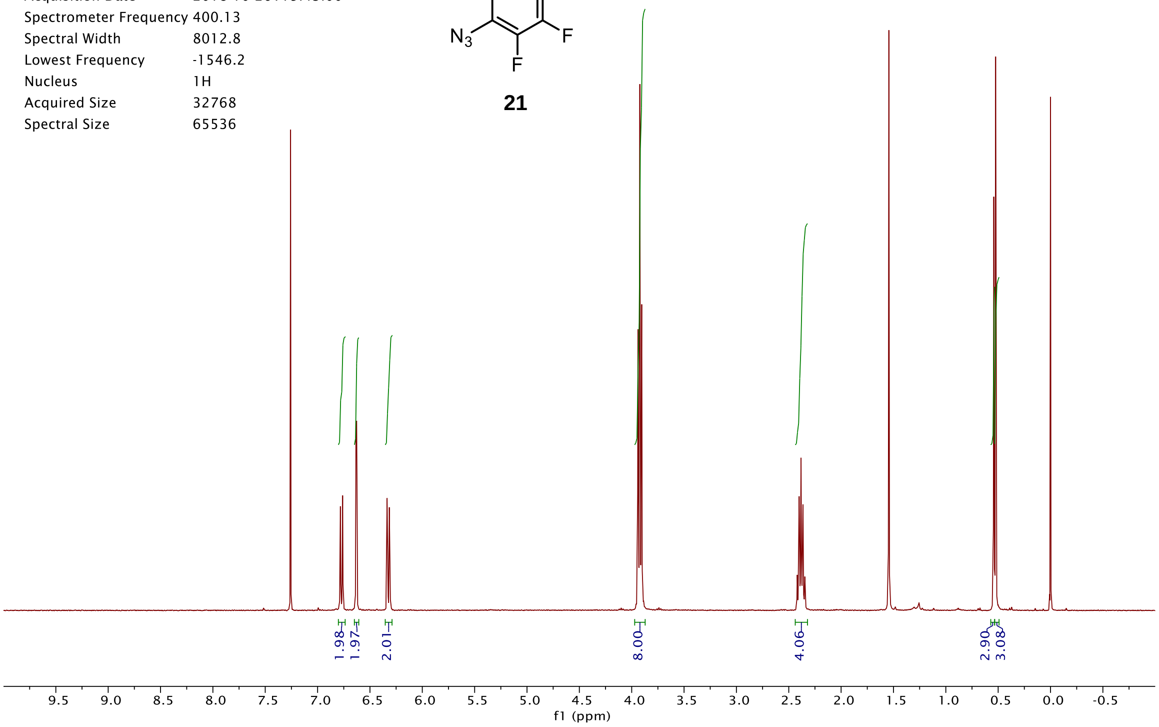


DAD1 E, Sig=675,4 Ref=off (2018_10\DAILY_SEQUENCE_LC 2018-10-18 14-12-26\2018_100000002.D)

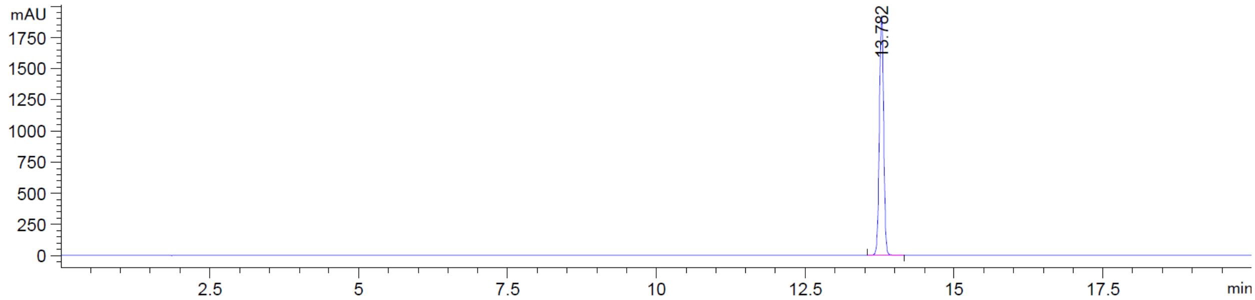


*MSD2 SPC, time=13.769:13.860 of C:\CHEM32\1\DATA\2018_10\DAILY_SEQUENCE_LC 2018-10-18 14-12-26\2018_100000002.D ES-

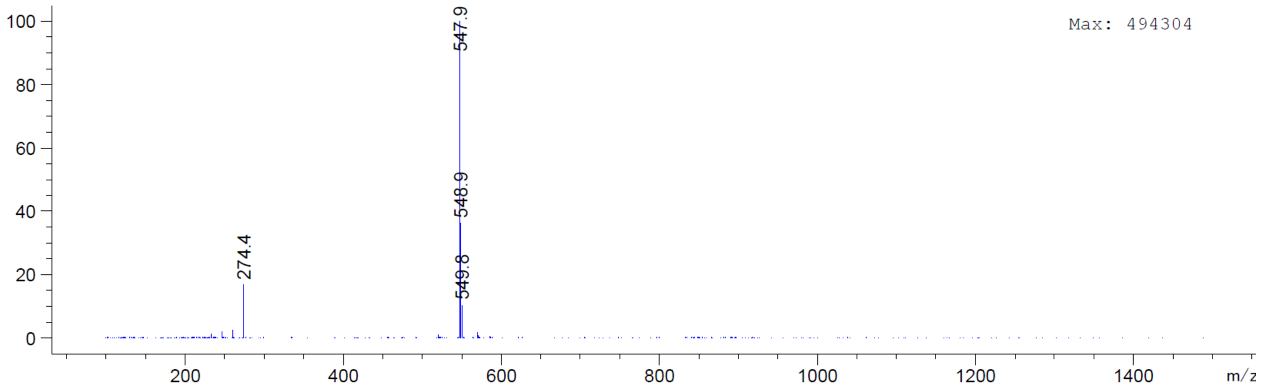

| | |
|---|---|
| Origin | Bruker BioSpin GmbH |
| Solvent | CDCl3 |
| Temperature | 300.0 |
| Pulse Sequence | zg30 |
| Experiment | 1D |
| Number of Scans | 16 |
| Acquisition Date | 2019-05-09T12:02:00 |
| Spectrometer Frequency | 400.13 |
| Spectral Width | 8012.8 |
| Lowest Frequency | -1546.0 |
| Nucleus | 1H |
| Acquired Size | 32768 |
| Spectral Size | 65536 |

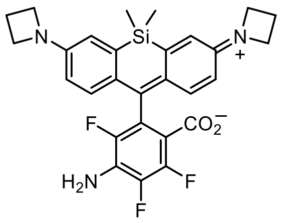


**22**

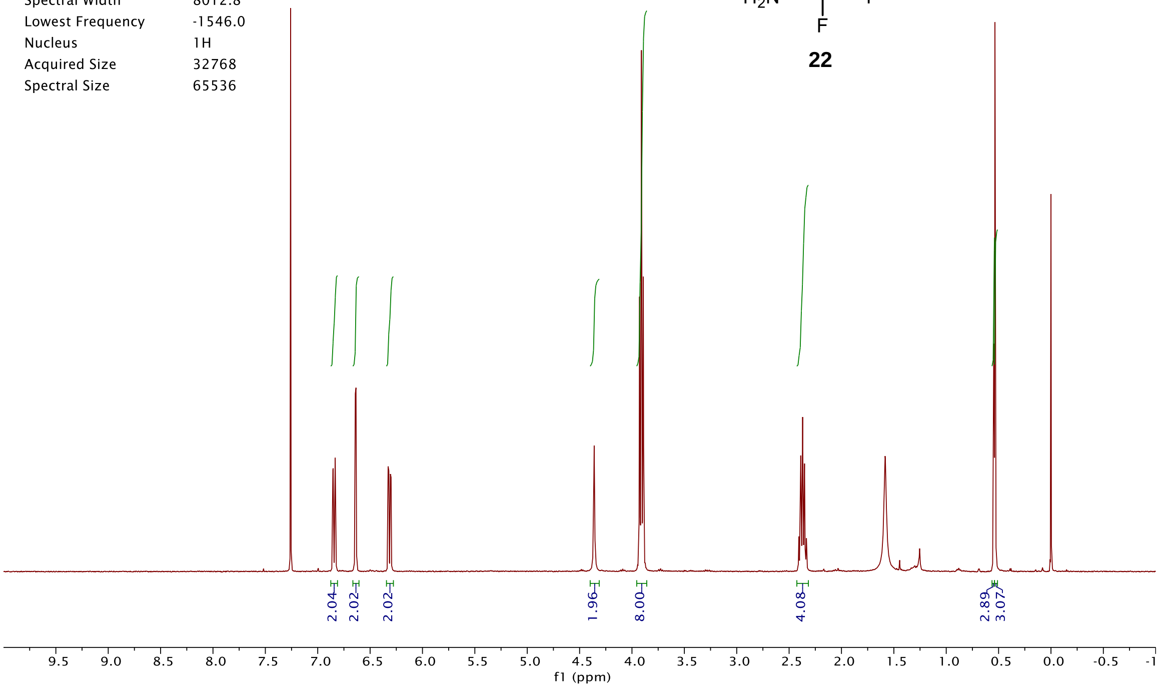


DAD1 E, Sig=675,4 Ref=off (2019_02\DAILY_SEQUENCE_LC 2019-02-19 11-46-41\2019_020000002.D)

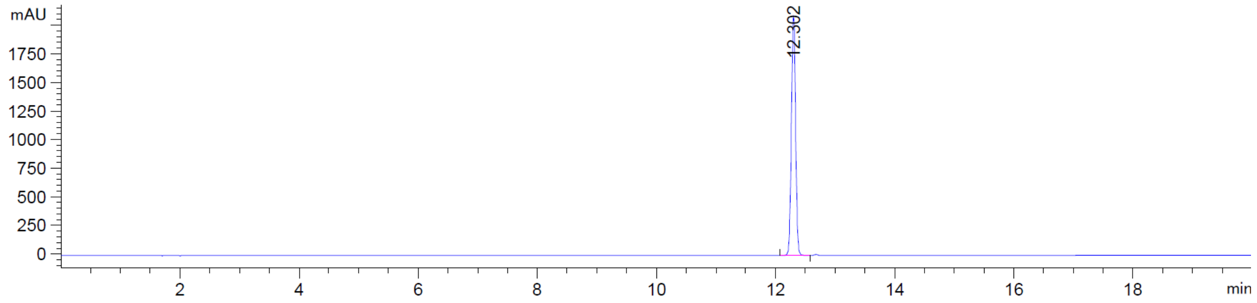


*MSD2 SPC, time=12.282:12.428 of C:\CHEM32\1\DATA\2019_02\DAILY_SEQUENCE_LC 2019-02-19 11-46-41\2019_020000002.D ES-

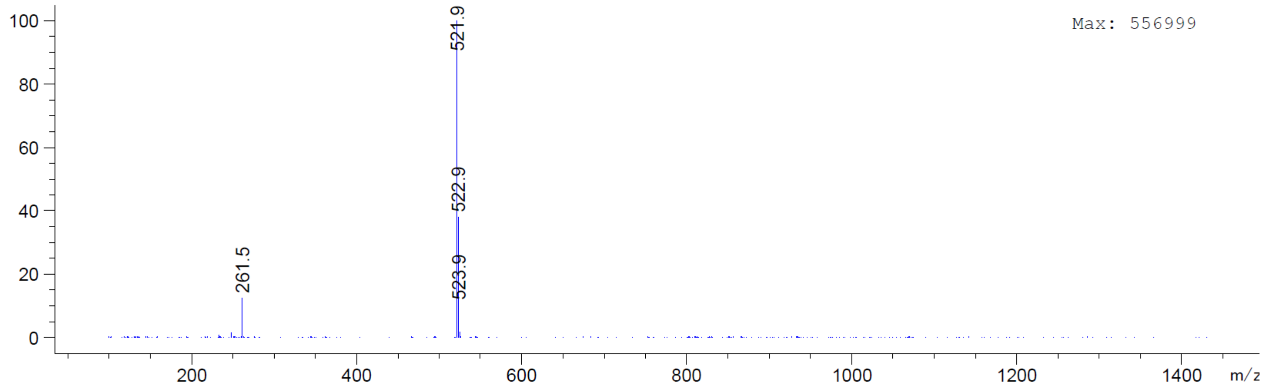

| | |
|---|---|
| Origin | Bruker BioSpin GmbH |
| Solvent | CDCl3 |
| Temperature | 300.0 |
| Pulse Sequence | zg30 |
| Experiment | 1D |
| Number of Scans | 16 |
| Acquisition Date | 2018-11-05T16:26:00 |
| Spectrometer Frequency | 400.13 |
| Spectral Width | 8012.8 |
| Lowest Frequency | -1546.2 |
| Nucleus | 1H |
| Acquired Size | 32768 |
| Spectral Size | 65536 |

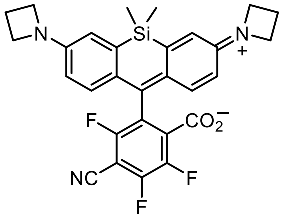


**23**

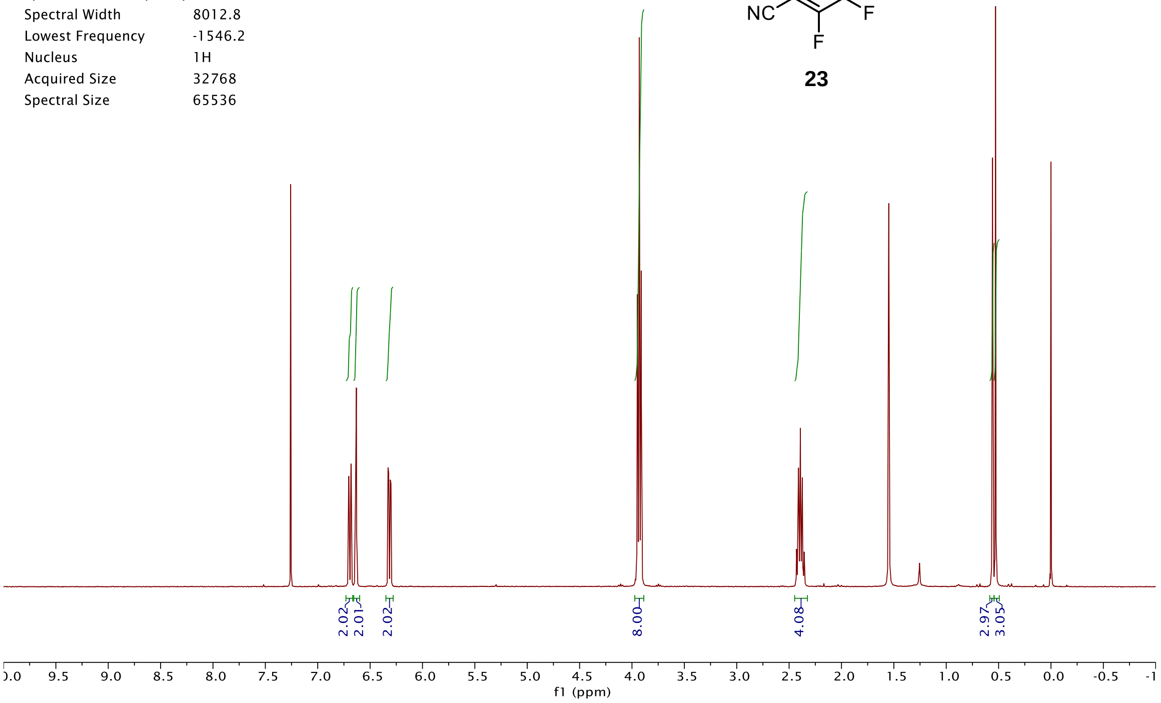


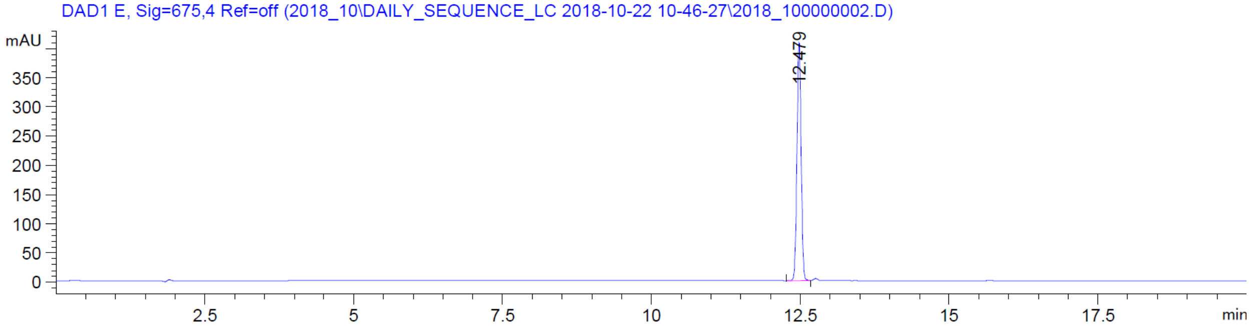


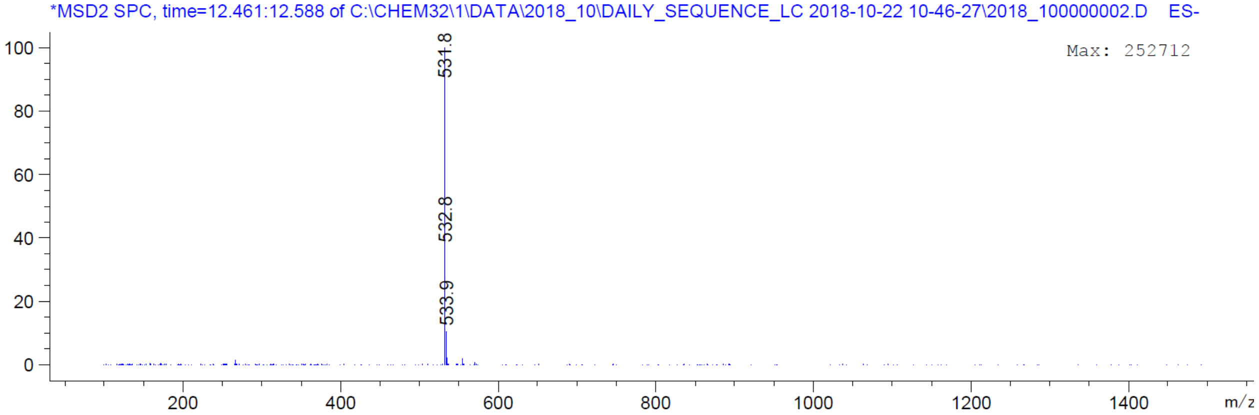

| | |
|---|---|
| Origin | Bruker BioSpin GmbH |
| Solvent | CDCl3 |
| Temperature | 300.0 |
| Pulse Sequence | zg30 |
| Experiment | 1D |
| Number of Scans | 16 |
| Acquisition Date | 2019-03-26T15:45:00 |
| Spectrometer Frequency | 400.13 |
| Spectral Width | 8012.8 |
| Lowest Frequency | -1546.1 |
| Nucleus | 1H |
| Acquired Size | 32768 |
| Spectral Size | 65536 |

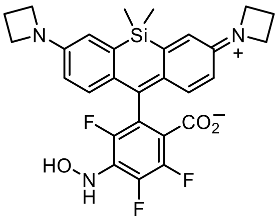


**24**

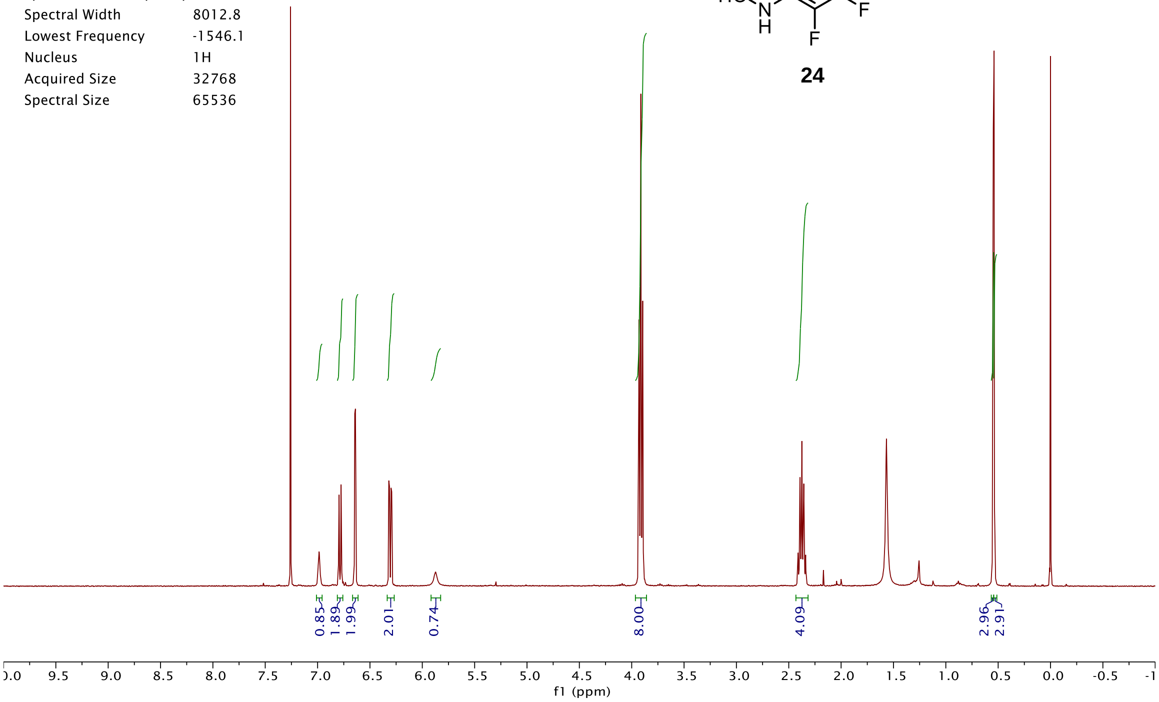


DAD1 E, Sig=675,4 Ref=off (2019_05\DAILY_SEQUENCE_LC 2019-05-10 13-40-26\2019_050000002.D)

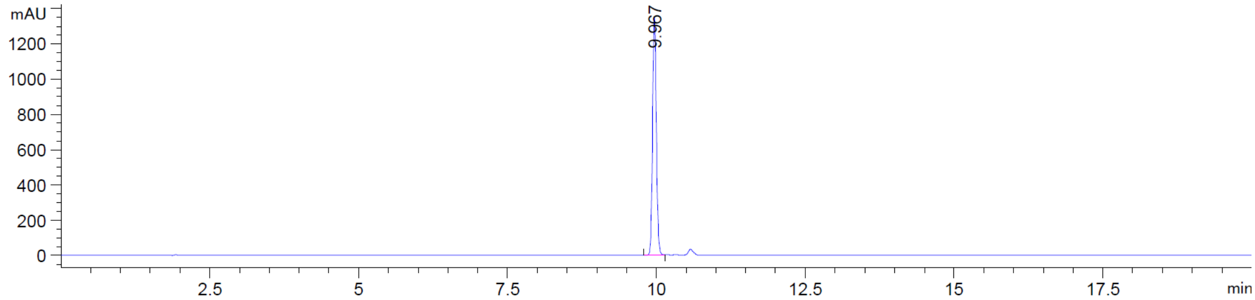


*MSD2 SPC, time=9.990:10.062 of C:\CHEM32\1\DATA\2019_05\DAILY_SEQUENCE_LC 2019-05-10 13-40-26\2019_050000002.D ES-A

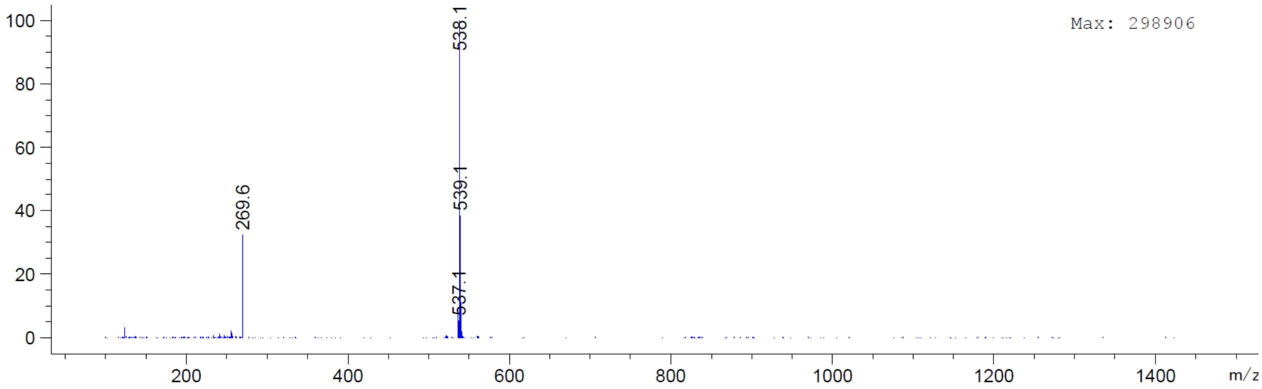

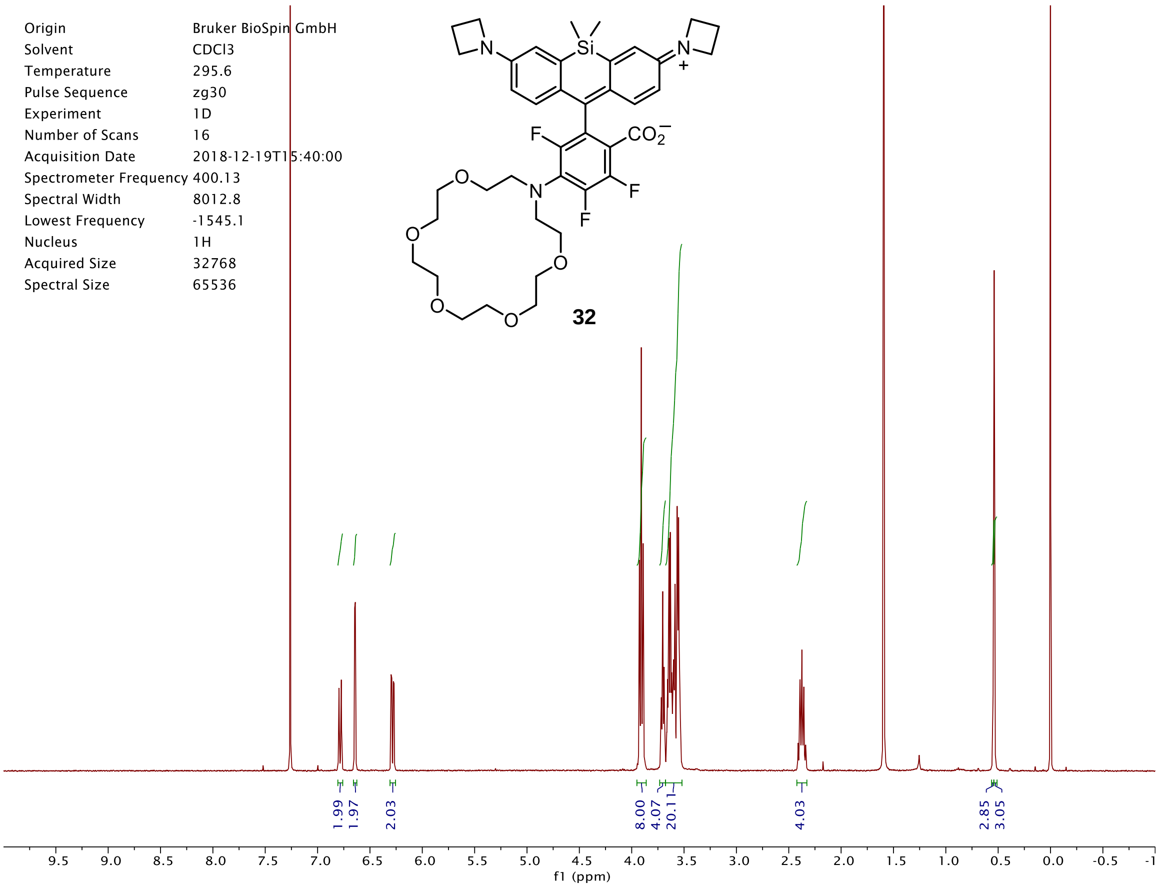


DAD1 E, Sig=675,4 Ref=off (2018_12\DAILY_SEQUENCE_LC 2018-12-13 13-28-22\2018_120000005.D)

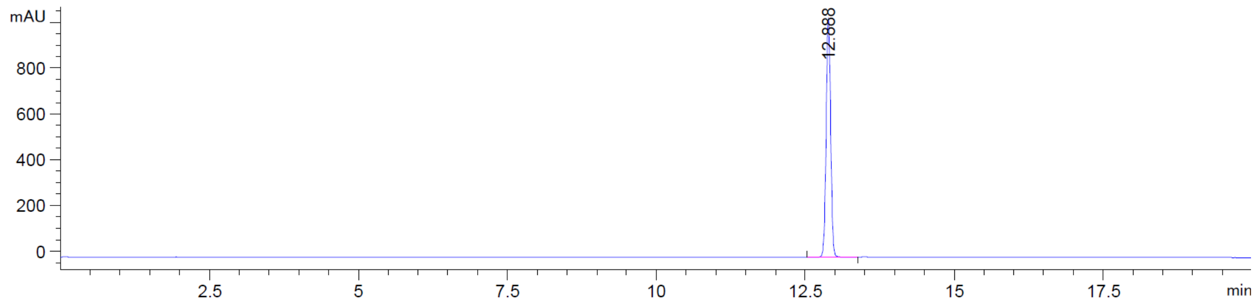


*MSD2 SPC, time=12.882:12.973 of C:\CHEM32\1\DATA\2018_12\DAILY_SEQUENCE_LC 2018-12-13 13-28-22\2018_120000005.D ES-

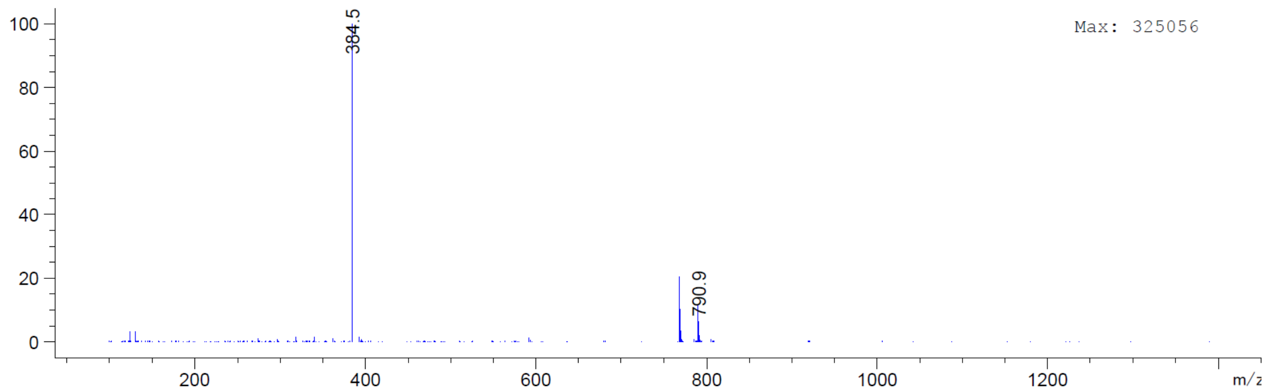

| | |
|---|---|
| Origin | Bruker BioSpin GmbH |
| Solvent | MeOD |
| Temperature | 300.0 |
| Pulse Sequence | zg30 |
| Experiment | 1D |
| Number of Scans | 16 |
| Acquisition Date | 2019-03-14T15:13:00 |
| Spectrometer Frequency | 400.13 |
| Spectral Width | 8012.8 |
| Lowest Frequency | -1543.2 |
| Nucleus | 1H |
| Acquired Size | 32768 |
| Spectral Size | 65536 |

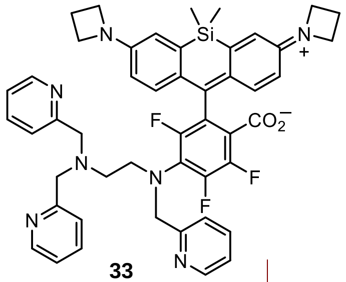


**33**

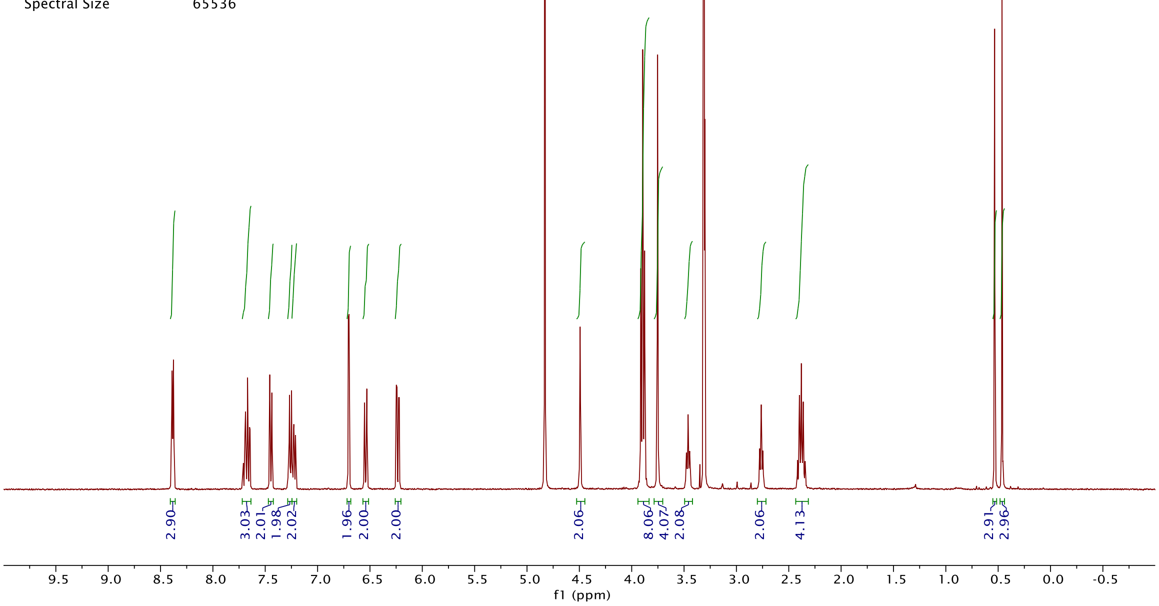


DAD1 E, Sig=675,4 Ref=off (2019_04\DAILY_SEQUENCE_LC 2019-04-08 14-40-21\2019_040000002.D)

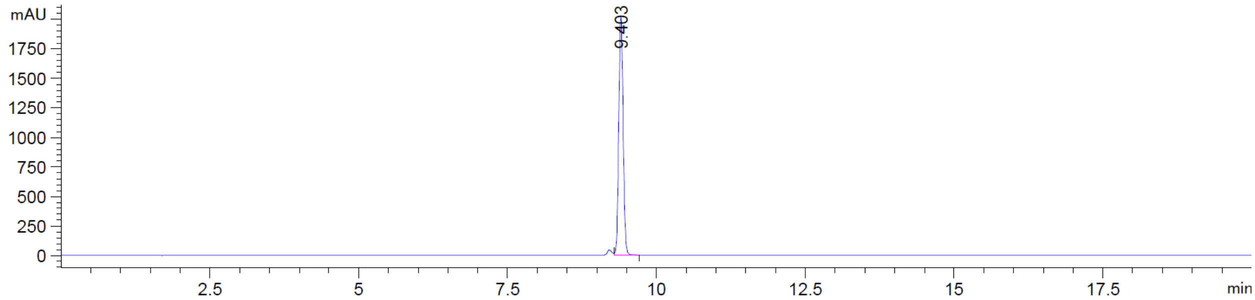


*MSD2 SPC, time=9.408:9.499 of C:\CHEM32\1\DATA\2019_04\DAILY_SEQUENCE_LC 2019-04-08 14-40-21\2019_040000002.D ES-AP

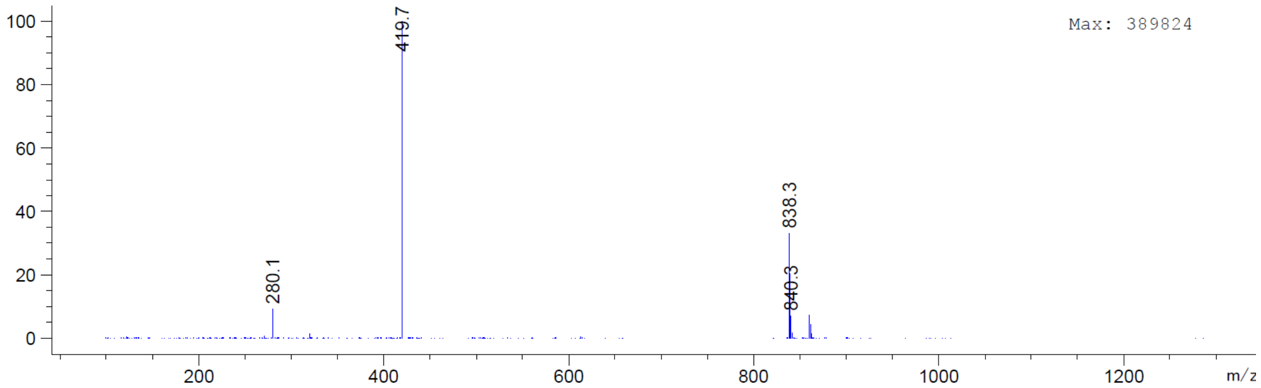

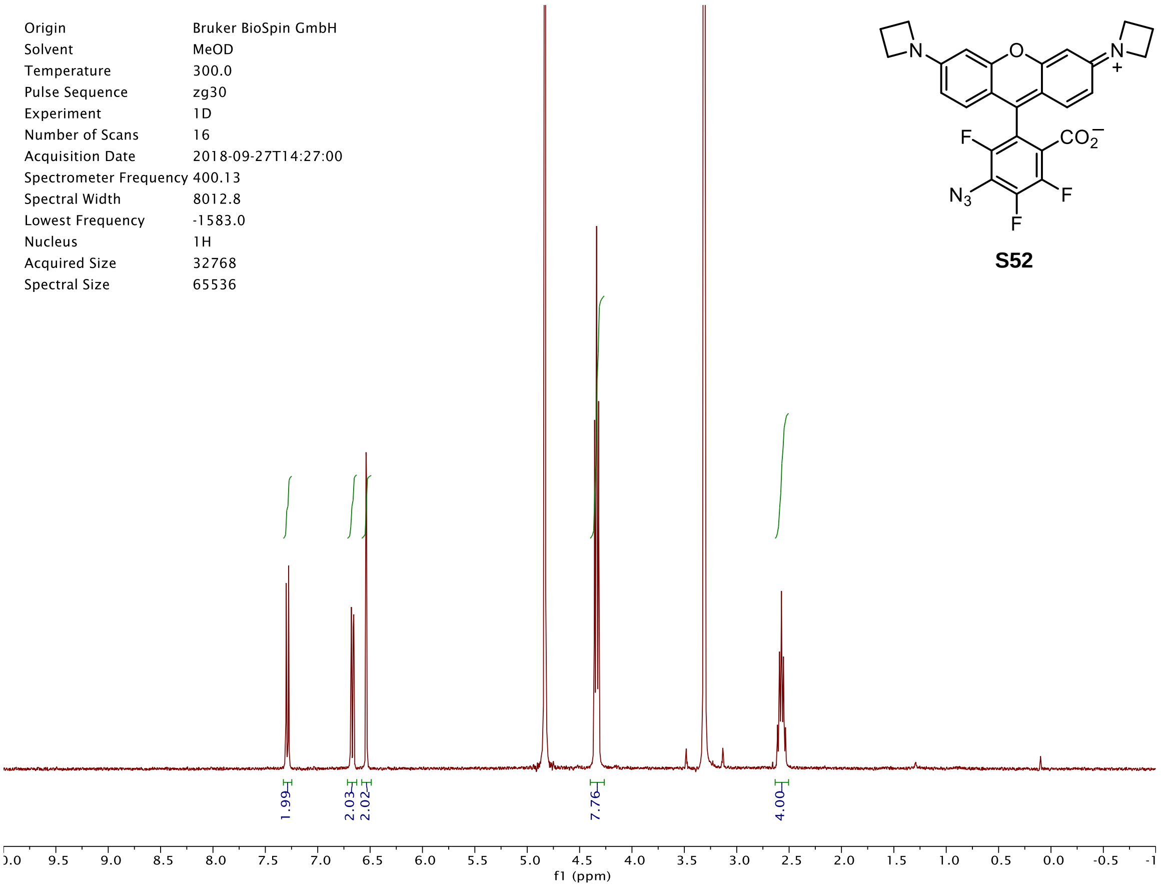

Origin Bruker BioSpin GmbH
Solvent MeOD
Temperature 300.0
Pulse Sequence zg30
Experiment 1D
Number of Scans 16
Acquisition Date 2018-09-27T14:27:00
Spectrometer Frequency 400.13
Spectral Width 8012.8
Lowest Frequency -1583.0
Nucleus 1H
Acquired Size 32768
Spectral Size 65536
N
O
N
+
F
$CO_2^-$
$N_3$
F
F
1.99
2.03
2.02
7.76
4.00
f1 (ppm)


**S52**

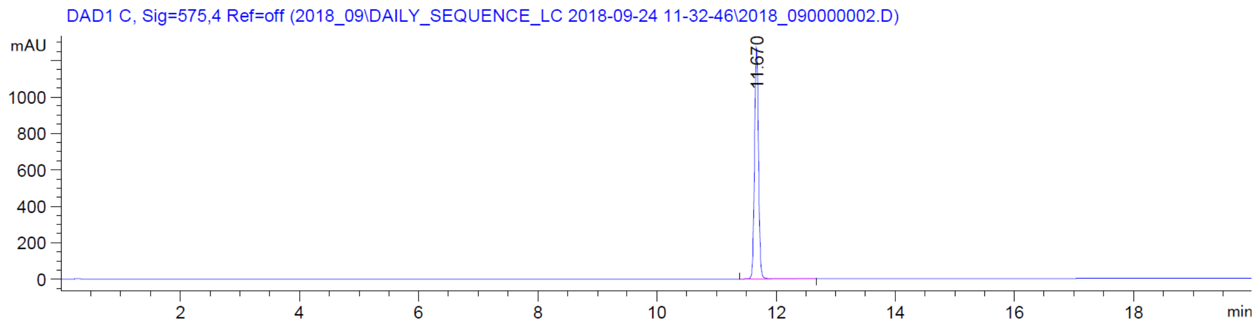

DAD1 C, Sig=575,4 Ref=off (2018_09\DAILY_SEQUENCE_LC 2018-09-24 11-32-46\2018_090000002.D)
mAU
11.670
min


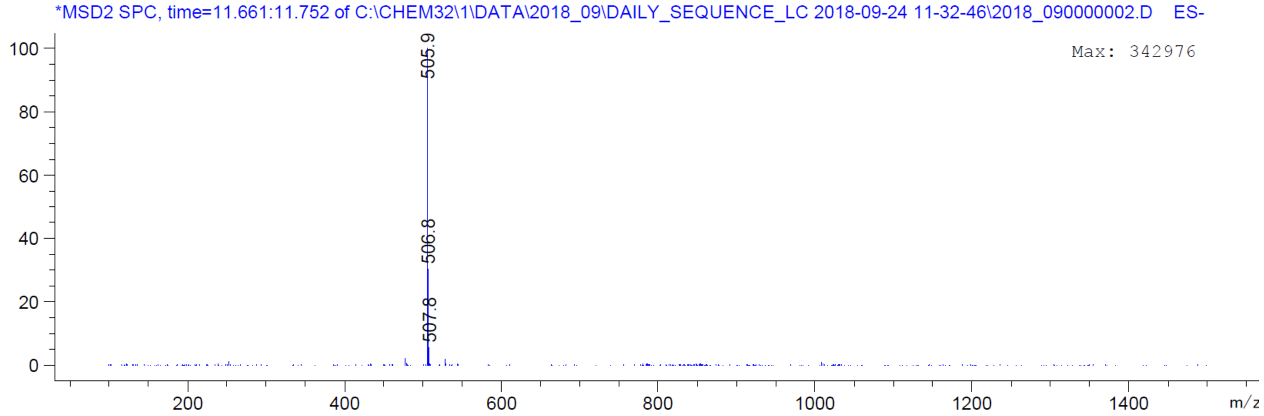

*MSD2 SPC, time=11.661:11.752 of C:\CHEM32\1\DATA\2018_09\DAILY_SEQUENCE_LC 2018-09-24 11-32-46\2018_090000002.D ES-
Max: 342976
505.9
506.8
507.8
m/z

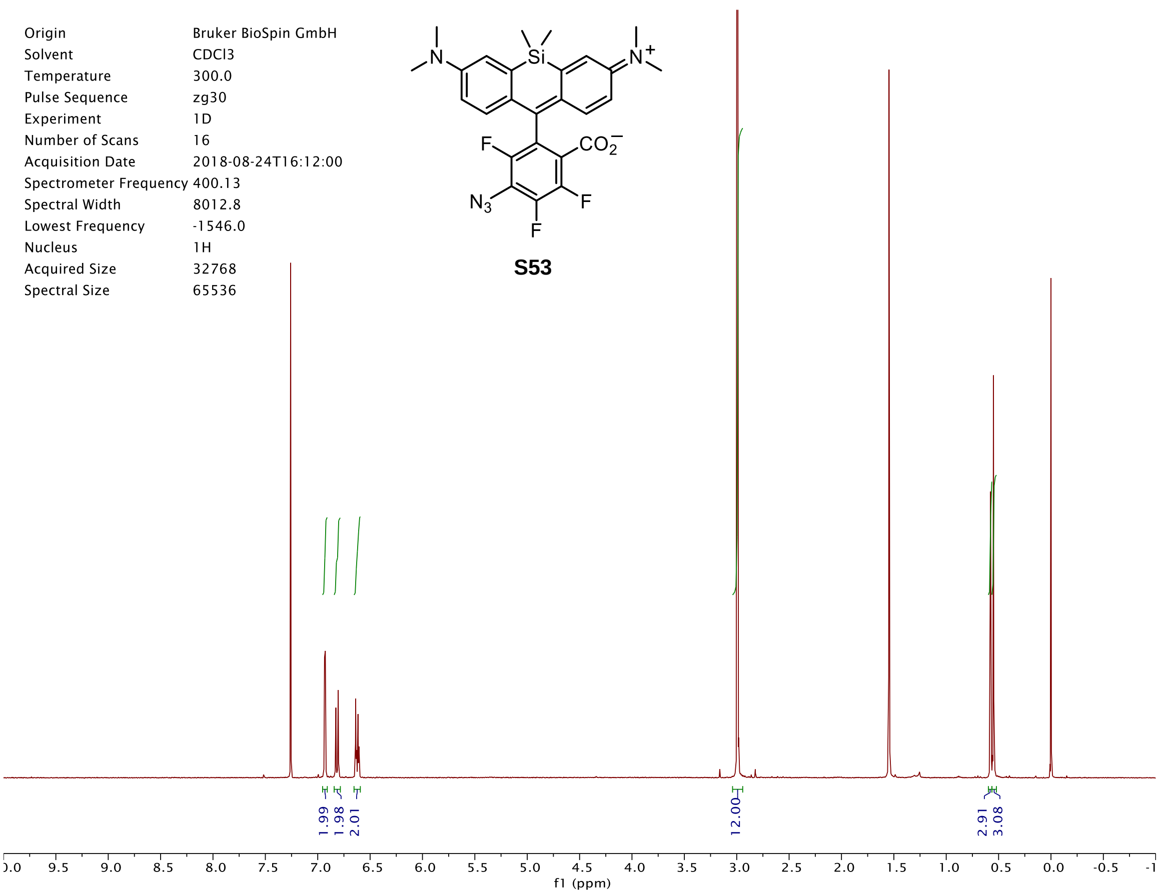
Origin Bruker BioSpin GmbH
Solvent CDCl3
Temperature 300.0
Pulse Sequence zg30
Experiment 1D
Number of Scans 16
Acquisition Date 2018-08-24T16:12:00
Spectrometer Frequency 400.13
Spectral Width 8012.8
Lowest Frequency -1546.0
Nucleus 1H
Acquired Size 32768
Spectral Size 65536
1.99
1.98
2.01
12.00
2.91
3.08
9.5 9.0 8.5 8.0 7.5 7.0 6.5 6.0 5.5 5.0 4.5 4.0 3.5 3.0 2.5 2.0 1.5 1.0 0.5 0.0 -0.5
f1 (ppm)

**S53**

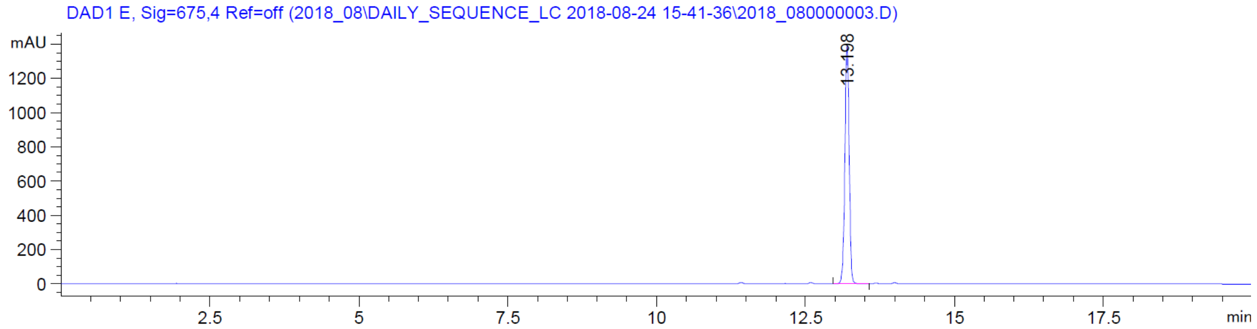
DAD1 E, Sig=675,4 Ref=off (2018_08\DAILY_SEQUENCE_LC 2018-08-24 15-41-36\2018_080000003.D)
mAU
1200
1000
800
600
400
200
0
13.198
2.5 5 7.5 10 12.5 15 17.5
min

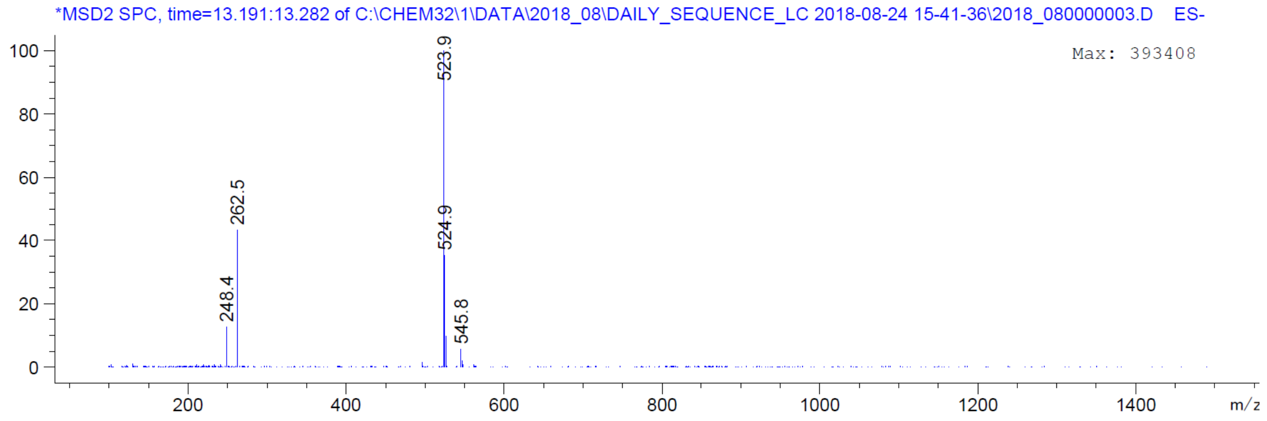
*MSD2 SPC, time=13.191:13.282 of C:\CHEM32\1\DATA\2018_08\DAILY_SEQUENCE_LC 2018-08-24 15-41-36\2018_080000003.D ES-
Max: 393408
100
80
60
40
20
0
523.9
262.5
524.9
248.4
545.8
200 400 600 800 1000 1200 1400
m/z

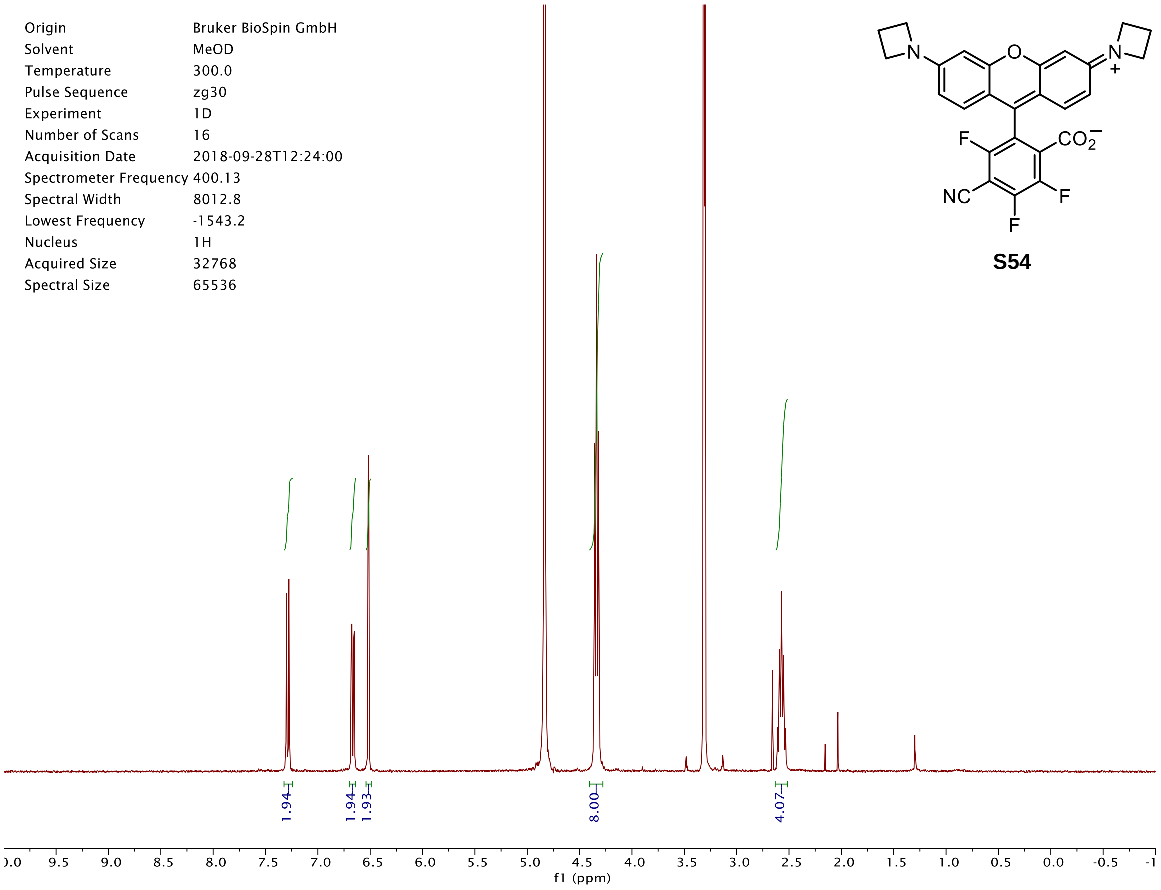
Origin Bruker BioSpin GmbH
Solvent MeOD
Temperature 300.0
Pulse Sequence zg30
Experiment 1D
Number of Scans 16
Acquisition Date 2018-09-28T12:24:00
Spectrometer Frequency 400.13
Spectral Width 8012.8
Lowest Frequency -1543.2
Nucleus 1H
Acquired Size 32768
Spectral Size 65536
1.94
1.94
1.93
8.00
4.07
f1 (ppm)
N
O
N
+
F
$CO_2^-$
NC
F
F

**S54**

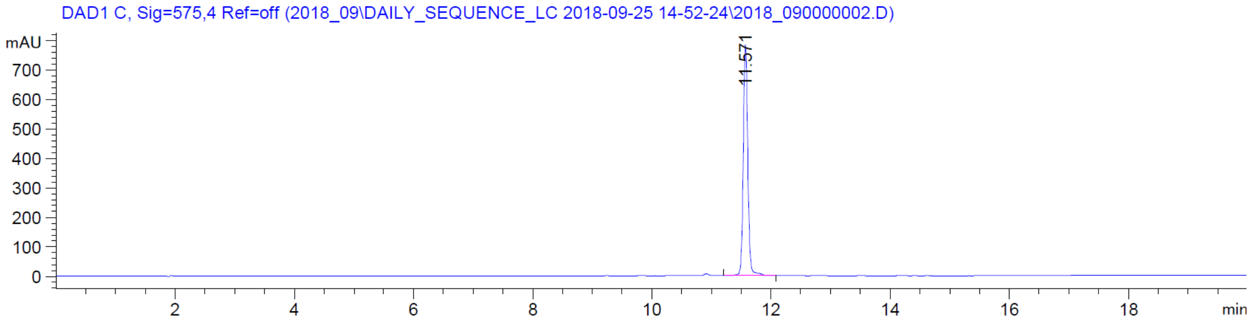
DAD1 C, Sig=575,4 Ref=off (2018_09\DAILY_SEQUENCE_LC 2018-09-25 14-52-24\2018_090000002.D)
mAU
11.571
min

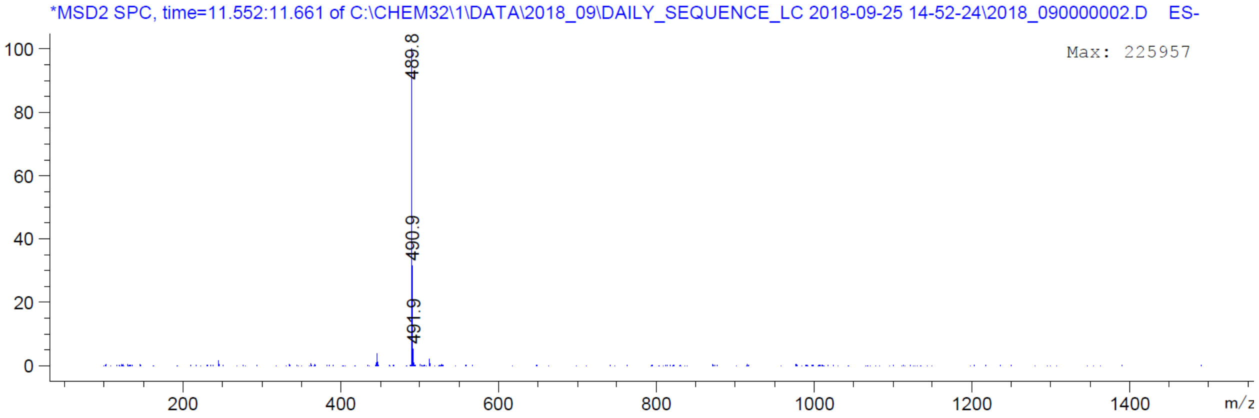
*MSD2 SPC, time=11.552:11.661 of C:\CHEM32\1\DATA\2018_09\DAILY_SEQUENCE_LC 2018-09-25 14-52-24\2018_090000002.D ES-
Max: 225957
489.8
490.9
491.9
m/z

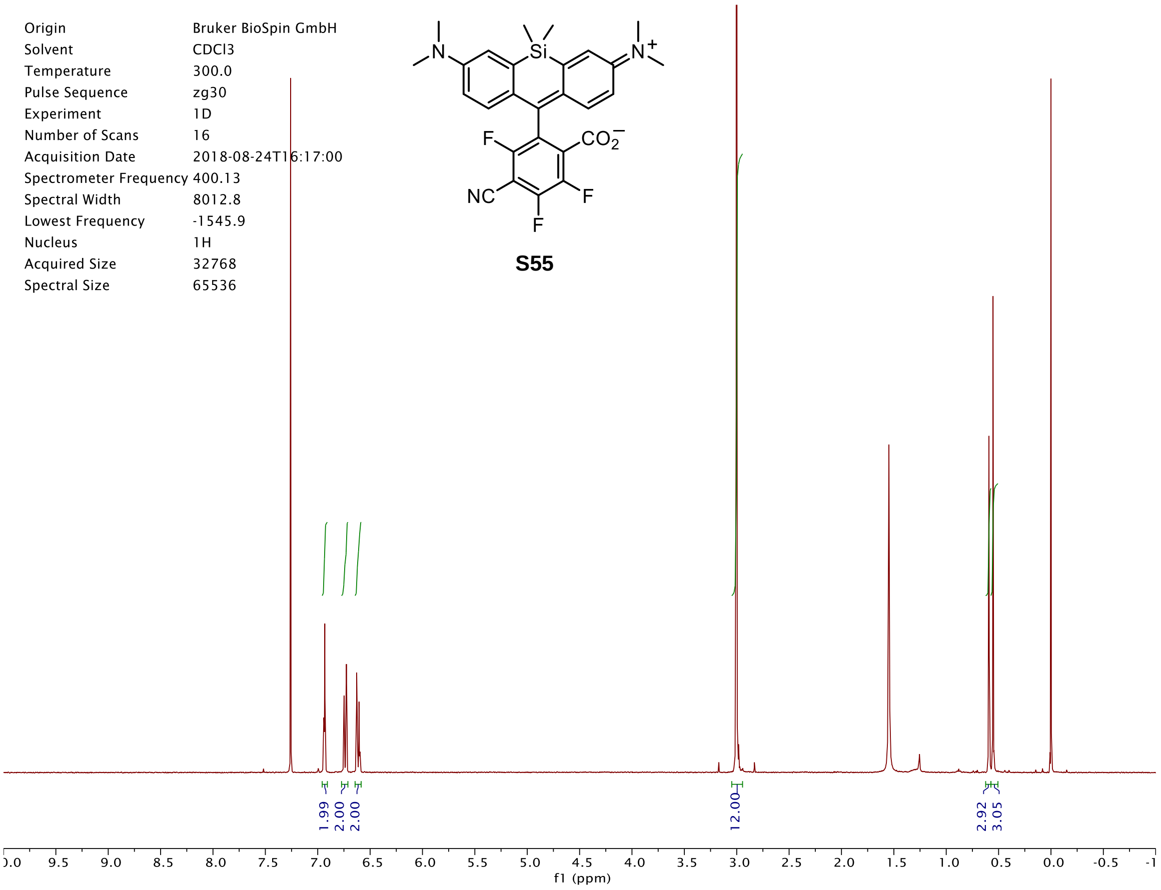
Origin Bruker BioSpin GmbH
Solvent CDCl3
Temperature 300.0
Pulse Sequence zg30
Experiment 1D
Number of Scans 16
Acquisition Date 2018-08-24T16:17:00
Spectrometer Frequency 400.13
Spectral Width 8012.8
Lowest Frequency -1545.9
Nucleus 1H
Acquired Size 32768
Spectral Size 65536
S55
1.99
2.00
2.00
12.00
2.92
3.05
f1 (ppm)

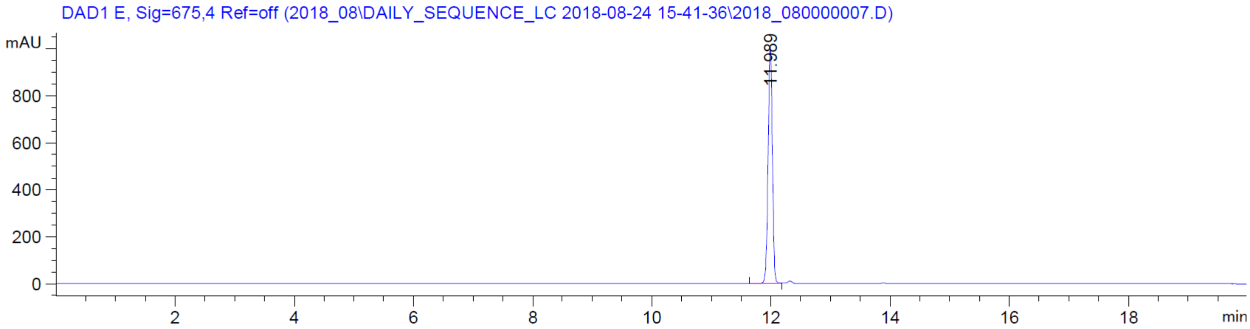
DAD1 E, Sig=675,4 Ref=off (2018_08\DAILY_SEQUENCE_LC 2018-08-24 15-41-36\2018_080000007.D)
mAU
11.989
min

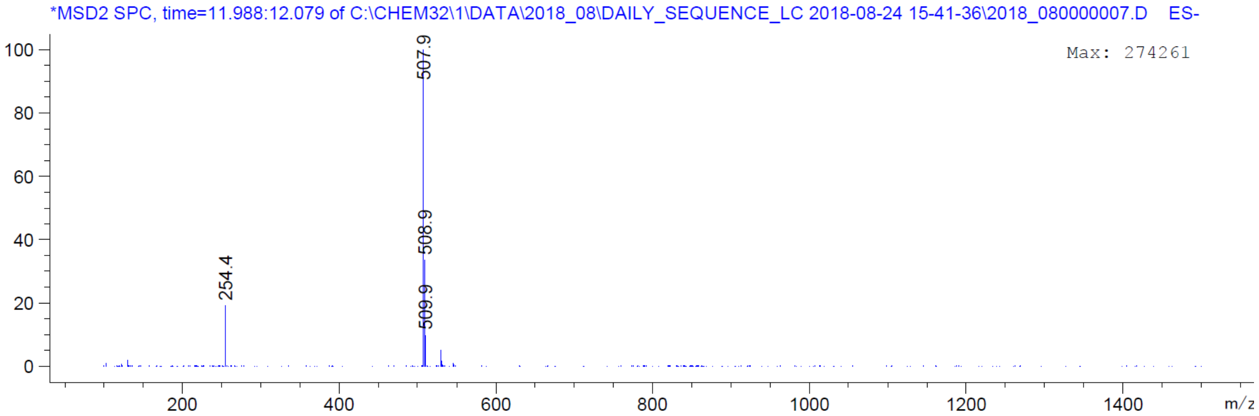
*MSD2 SPC, time=11.988:12.079 of C:\CHEM32\1\DATA\2018_08\DAILY_SEQUENCE_LC 2018-08-24 15-41-36\2018_080000007.D ES-
Max: 274261
507.9
508.9
509.9
254.4
m/z

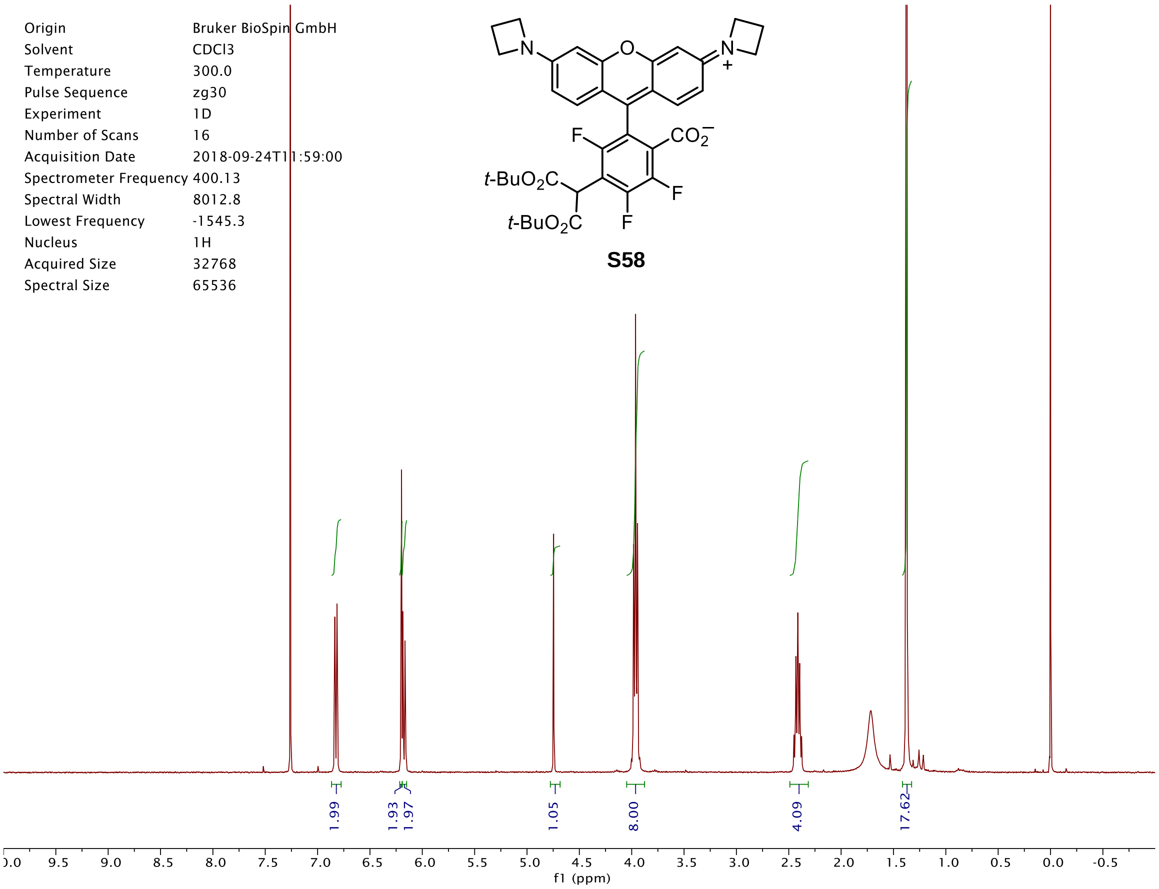
Origin Bruker BioSpin GmbH
Solvent CDCl3
Temperature 300.0
Pulse Sequence zg30
Experiment 1D
Number of Scans 16
Acquisition Date 2018-09-24T11:59:00
Spectrometer Frequency 400.13
Spectral Width 8012.8
Lowest Frequency -1545.3
Nucleus 1H
Acquired Size 32768
Spectral Size 65536
t-BuO2C
t-BuO2C
CO2−
S58
1.99
1.93
1.97
1.05
8.00
4.09
17.62
f1 (ppm)

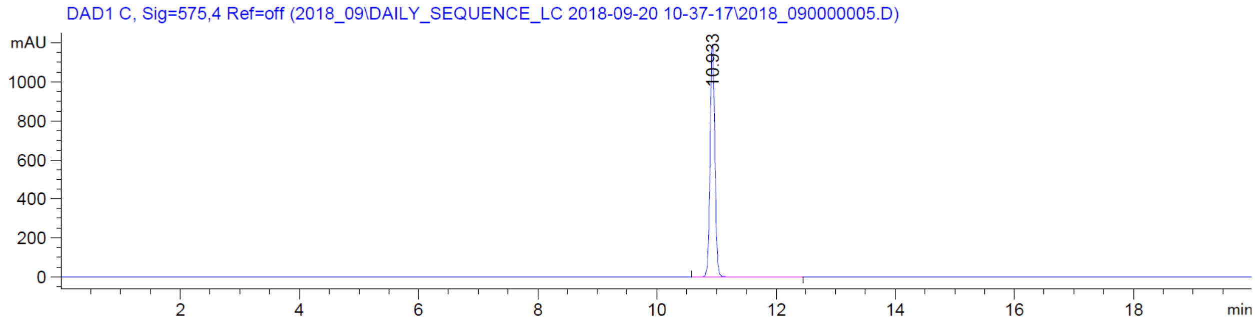
DAD1 C, Sig=575,4 Ref=off (2018_09\DAILY_SEQUENCE_LC 2018-09-20 10-37-17\2018_090000005.D)
mAU
10.933
min

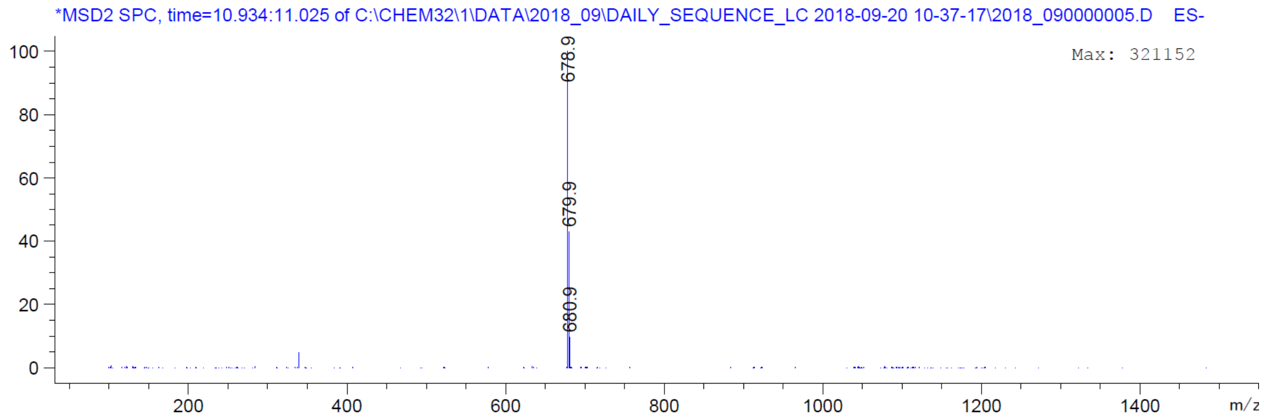
*MSD2 SPC, time=10.934:11.025 of C:\CHEM32\1\DATA\2018_09\DAILY_SEQUENCE_LC 2018-09-20 10-37-17\2018_090000005.D ES-
Max: 321152
678.9
679.9
680.9
m/z

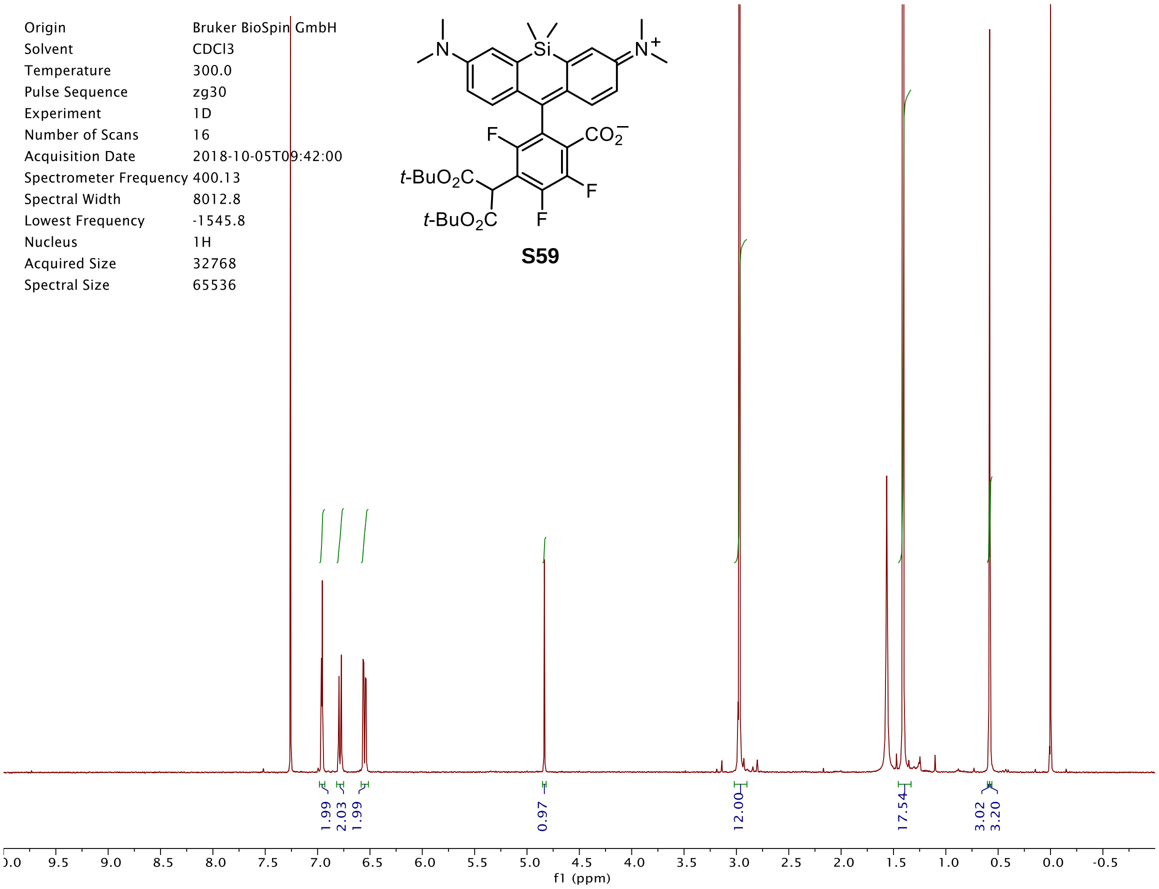

t-BuO2C
t-BuO2C
F
F
F
CO2−
Si
N
N+
S59
Origin Bruker BioSpin GmbH
Solvent CDCl3
Temperature 300.0
Pulse Sequence zg30
Experiment 1D
Number of Scans 16
Acquisition Date 2018-10-05T09:42:00
Spectrometer Frequency 400.13
Spectral Width 8012.8
Lowest Frequency -1545.8
Nucleus 1H
Acquired Size 32768
Spectral Size 65536
1.99
2.03
1.99
0.97
12.00
17.54
3.02
3.20
f1 (ppm)


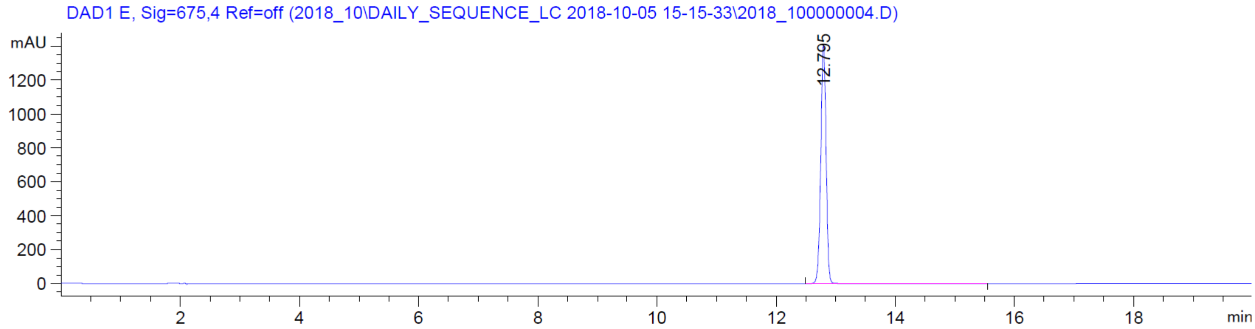

DAD1 E, Sig=675,4 Ref=off (2018_10\DAILY_SEQUENCE_LC 2018-10-05 15-15-33\2018_100000004.D)
mAU
12.795
min


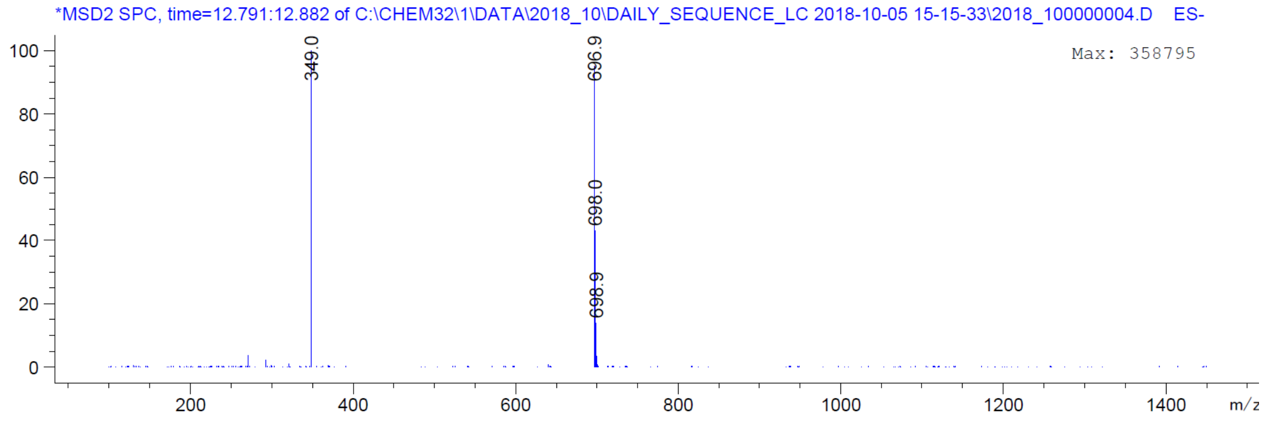

*MSD2 SPC, time=12.791:12.882 of C:\CHEM32\1\DATA\2018_10\DAILY_SEQUENCE_LC 2018-10-05 15-15-33\2018_100000004.D ES-
Max: 358795
349.0
696.9
698.0
698.9
m/z

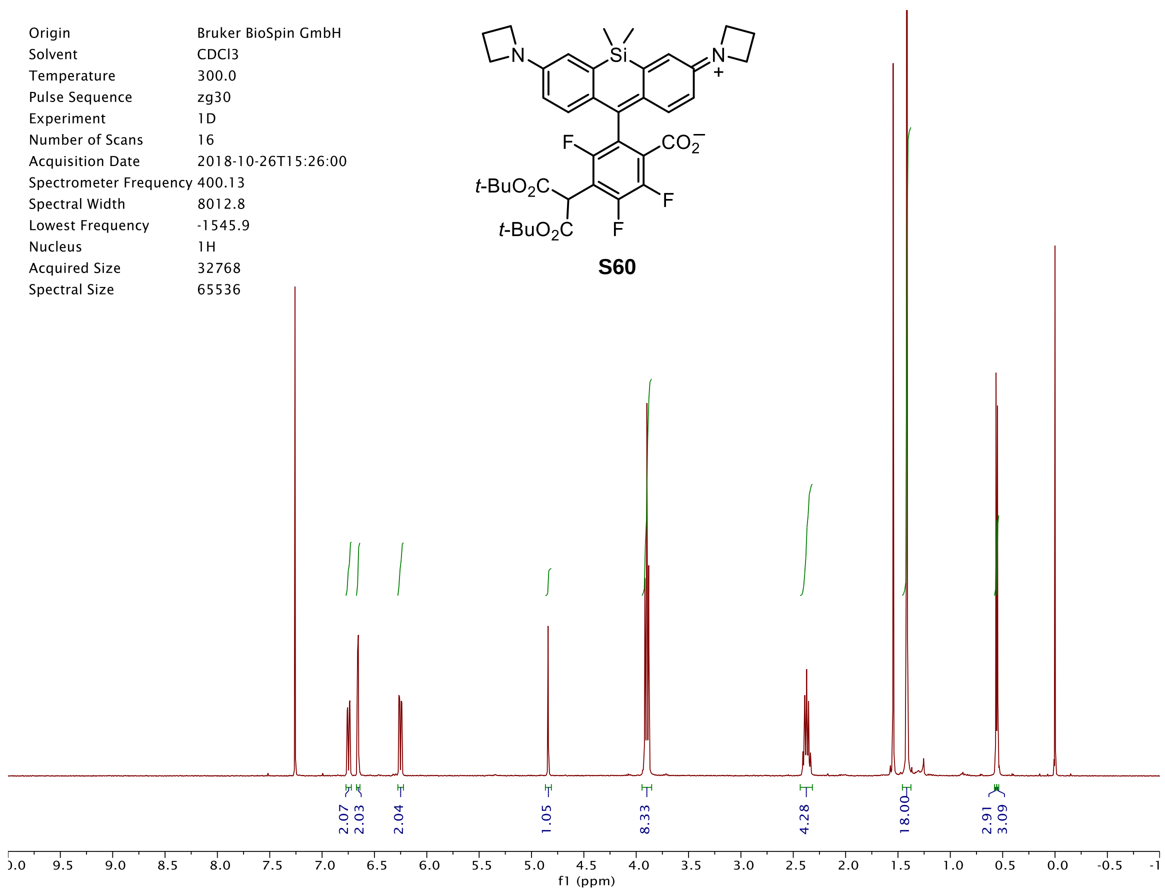
Origin Bruker BioSpin GmbH
Solvent CDCl3
Temperature 300.0
Pulse Sequence zg30
Experiment 1D
Number of Scans 16
Acquisition Date 2018-10-26T15:26:00
Spectrometer Frequency 400.13
Spectral Width 8012.8
Lowest Frequency -1545.9
Nucleus 1H
Acquired Size 32768
Spectral Size 65536
S60
2.07
2.03
2.04
1.05
8.33
4.28
18.00
2.91
3.09
f1 (ppm)

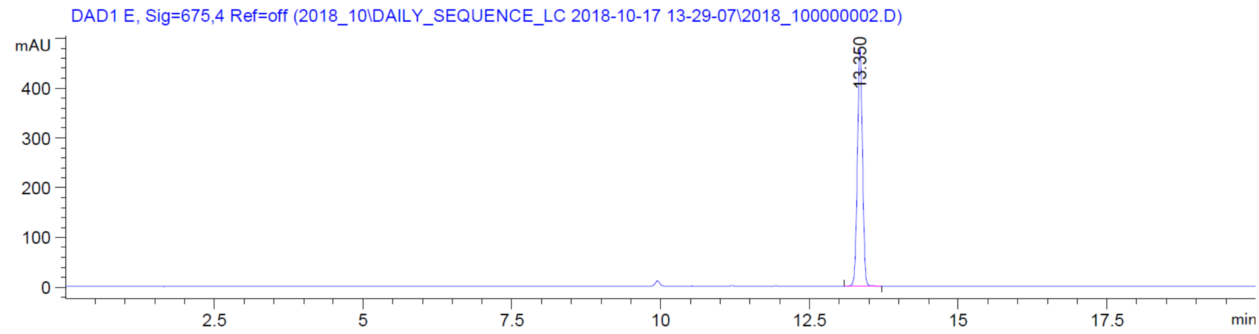
DAD1 E, Sig=675,4 Ref=off (2018_10\DAILY_SEQUENCE_LC 2018-10-17 13-29-07\2018_100000002.D)
mAU
13.350
min

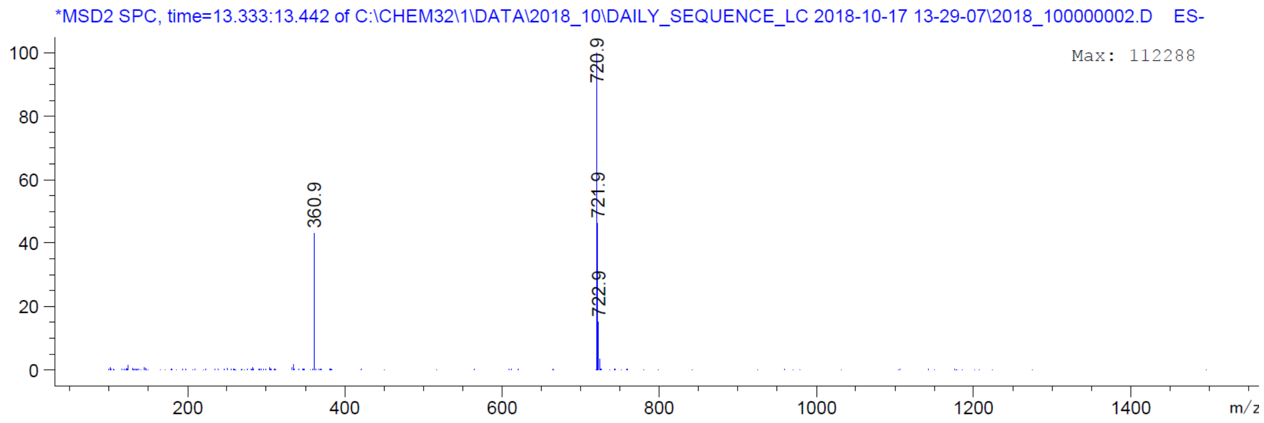
*MSD2 SPC, time=13.333:13.442 of C:\CHEM32\1\DATA\2018_10\DAILY_SEQUENCE_LC 2018-10-17 13-29-07\2018_100000002.D ES-
Max: 112288
360.9
720.9
721.9
722.9
m/z

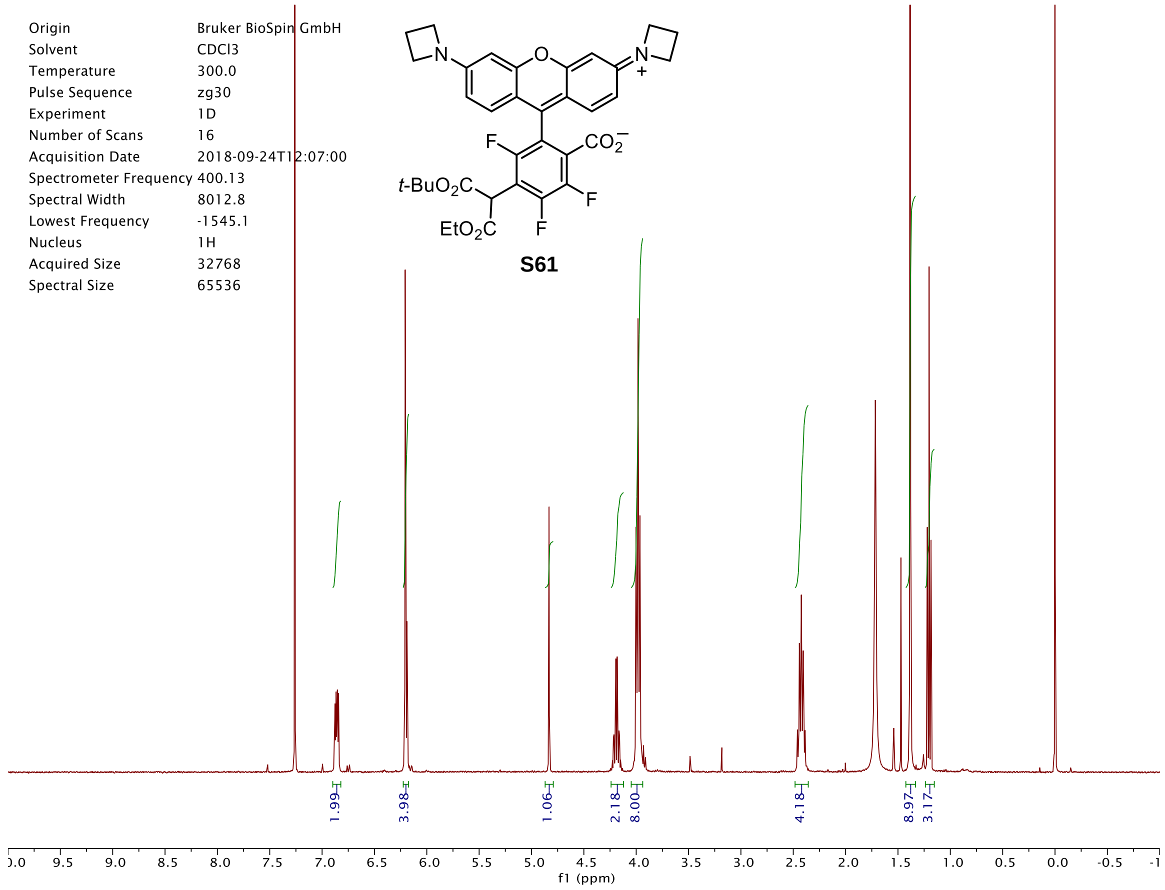
Origin Bruker BioSpin GmbH
Solvent CDCl3
Temperature 300.0
Pulse Sequence zg30
Experiment 1D
Number of Scans 16
Acquisition Date 2018-09-24T12:07:00
Spectrometer Frequency 400.13
Spectral Width 8012.8
Lowest Frequency -1545.1
Nucleus 1H
Acquired Size 32768
Spectral Size 65536
t-BuO2C
EtO2C
F
CO2−
N
S61
1.99
3.98
1.06
2.18
8.00
4.18
8.97
3.17
f1 (ppm)

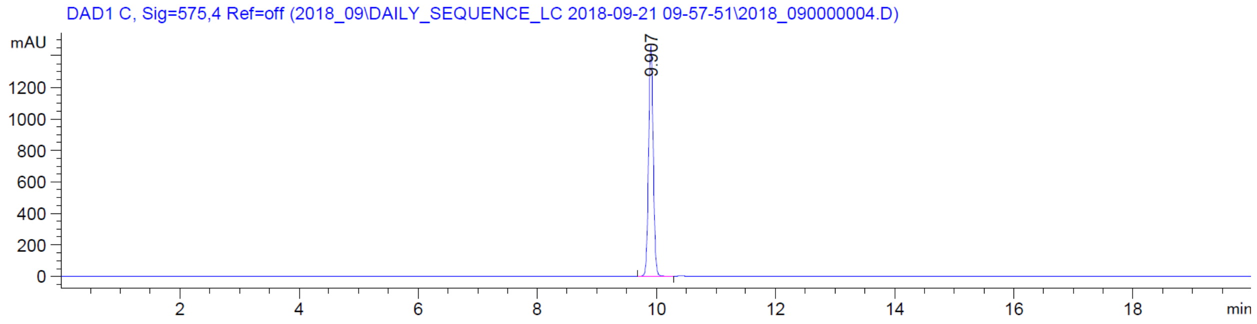
DAD1 C, Sig=575,4 Ref=off (2018_09\DAILY_SEQUENCE_LC 2018-09-21 09-57-51\2018_090000004.D)
mAU
9.907
min

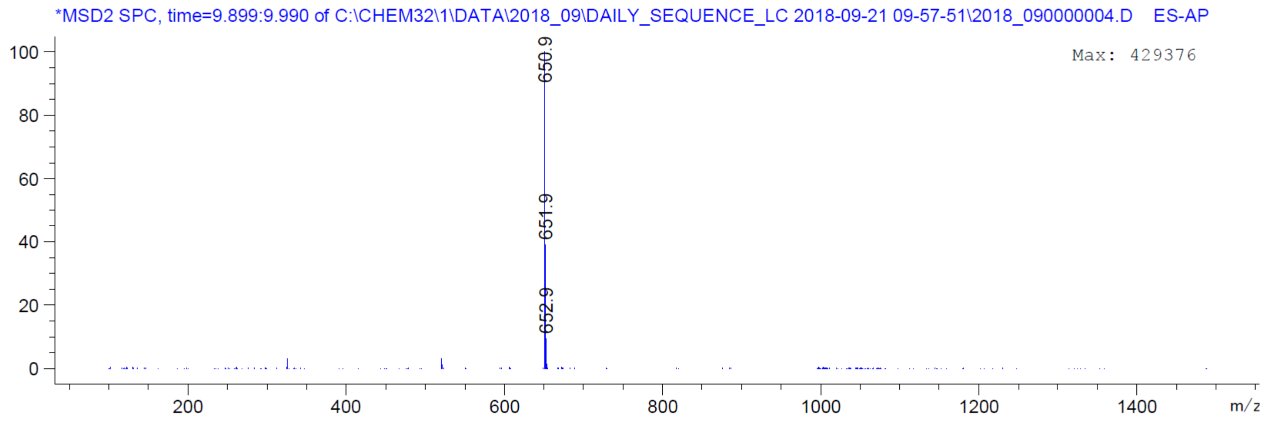
*MSD2 SPC, time=9.899:9.990 of C:\CHEM32\1\DATA\2018_09\DAILY_SEQUENCE_LC 2018-09-21 09-57-51\2018_090000004.D ES-AP
Max: 429376
650.9
651.9
652.9
m/z

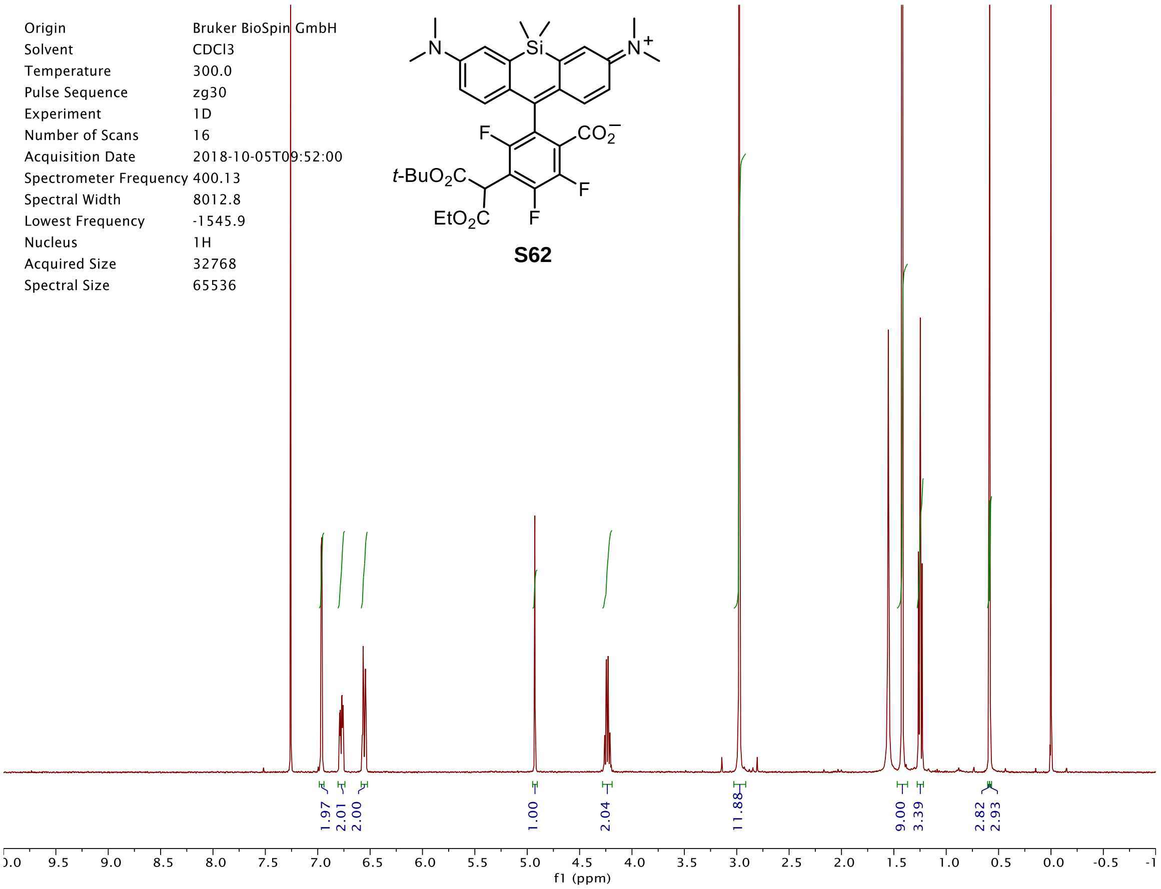
Origin Bruker BioSpin GmbH
Solvent CDCl3
Temperature 300.0
Pulse Sequence zg30
Experiment 1D
Number of Scans 16
Acquisition Date 2018-10-05T09:52:00
Spectrometer Frequency 400.13
Spectral Width 8012.8
Lowest Frequency -1545.9
Nucleus 1H
Acquired Size 32768
Spectral Size 65536
S62
1.97
2.01
2.00
1.00
2.04
11.88
9.00
3.39
2.82
2.93
f1 (ppm)

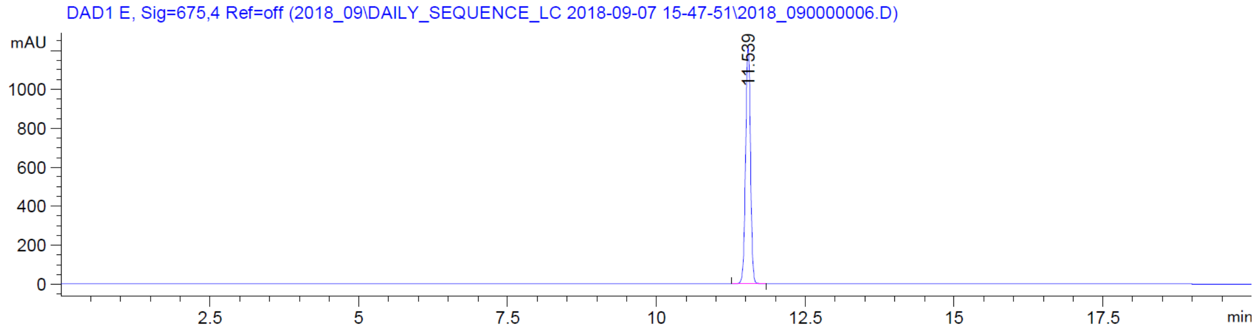
DAD1 E, Sig=675,4 Ref=off (2018_09\DAILY_SEQUENCE_LC 2018-09-07 15-47-51\2018_090000006.D)
mAU
11.539
min

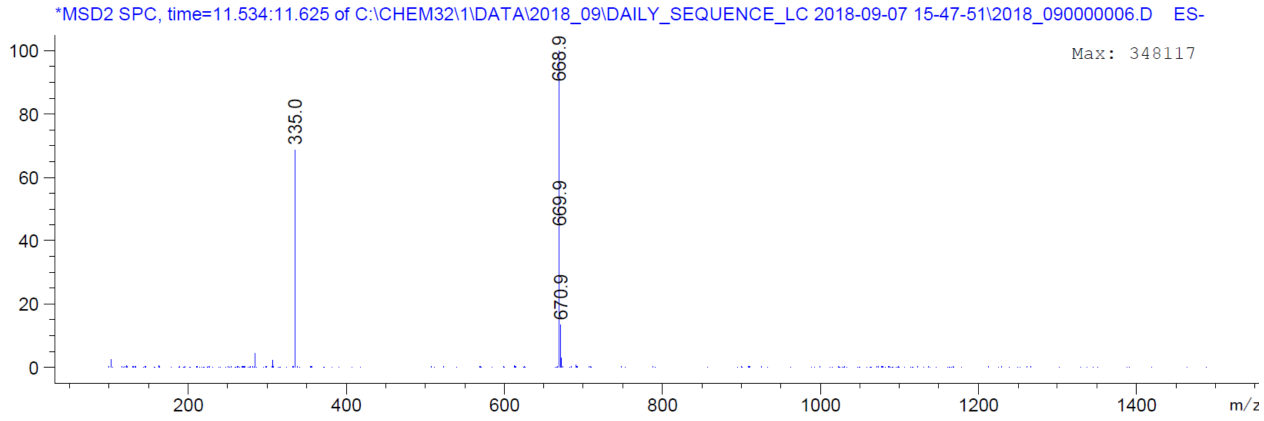
*MSD2 SPC, time=11.534:11.625 of C:\CHEM32\1\DATA\2018_09\DAILY_SEQUENCE_LC 2018-09-07 15-47-51\2018_090000006.D ES-
Max: 348117
335.0
668.9
669.9
670.9
m/z

| | |
|---|---|
| Origin | Bruker BioSpin GmbH |
| Solvent | CDCl3 |
| Temperature | 300.0 |
| Pulse Sequence | zg30 |
| Experiment | 1D |
| Number of Scans | 16 |
| Acquisition Date | 2018-10-26T15:35:00 |
| Spectrometer Frequency | 400.13 |
| Spectral Width | 8012.8 |
| Lowest Frequency | -1545.9 |
| Nucleus | 1H |
| Acquired Size | 32768 |
| Spectral Size | 65536 |

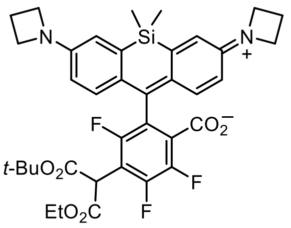


**S63**

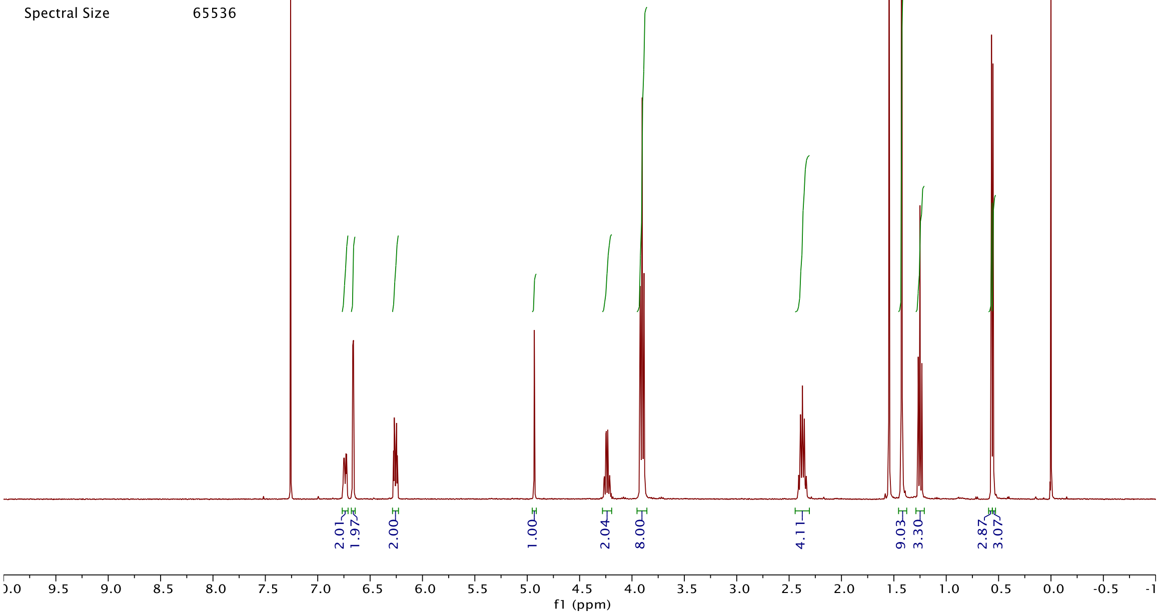


DAD1 E, Sig=675,4 Ref=off (2018_10\DAILY_SEQUENCE_LC 2018-10-16 15-48-57\2018_100000002.D)

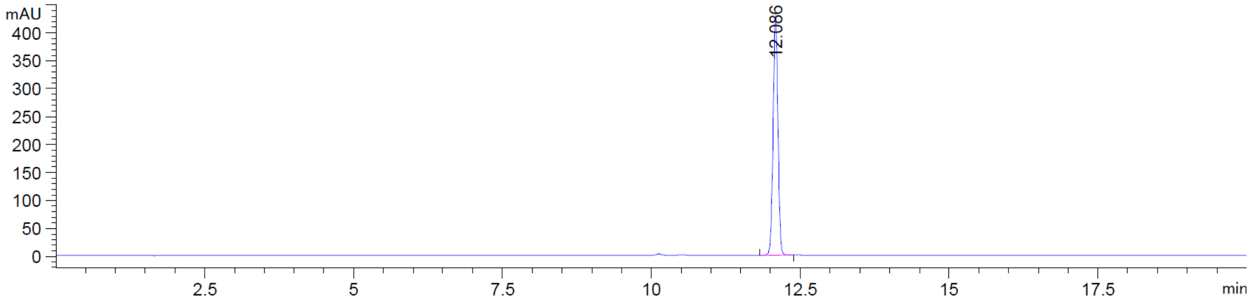


*MSD2 SPC, time=12.079:12.170 of C:\CHEM32\1\DATA\2018_10\DAILY_SEQUENCE_LC 2018-10-16 15-48-57\2018_100000002.D ES-

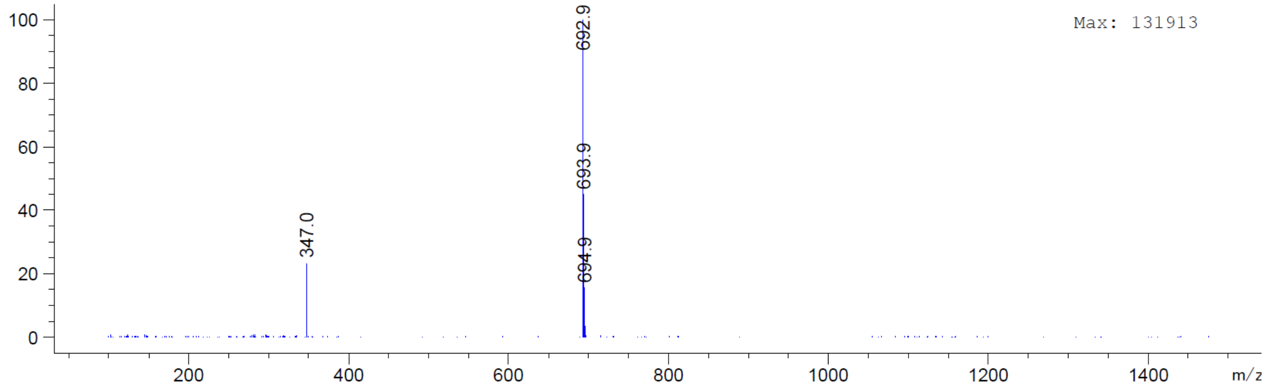

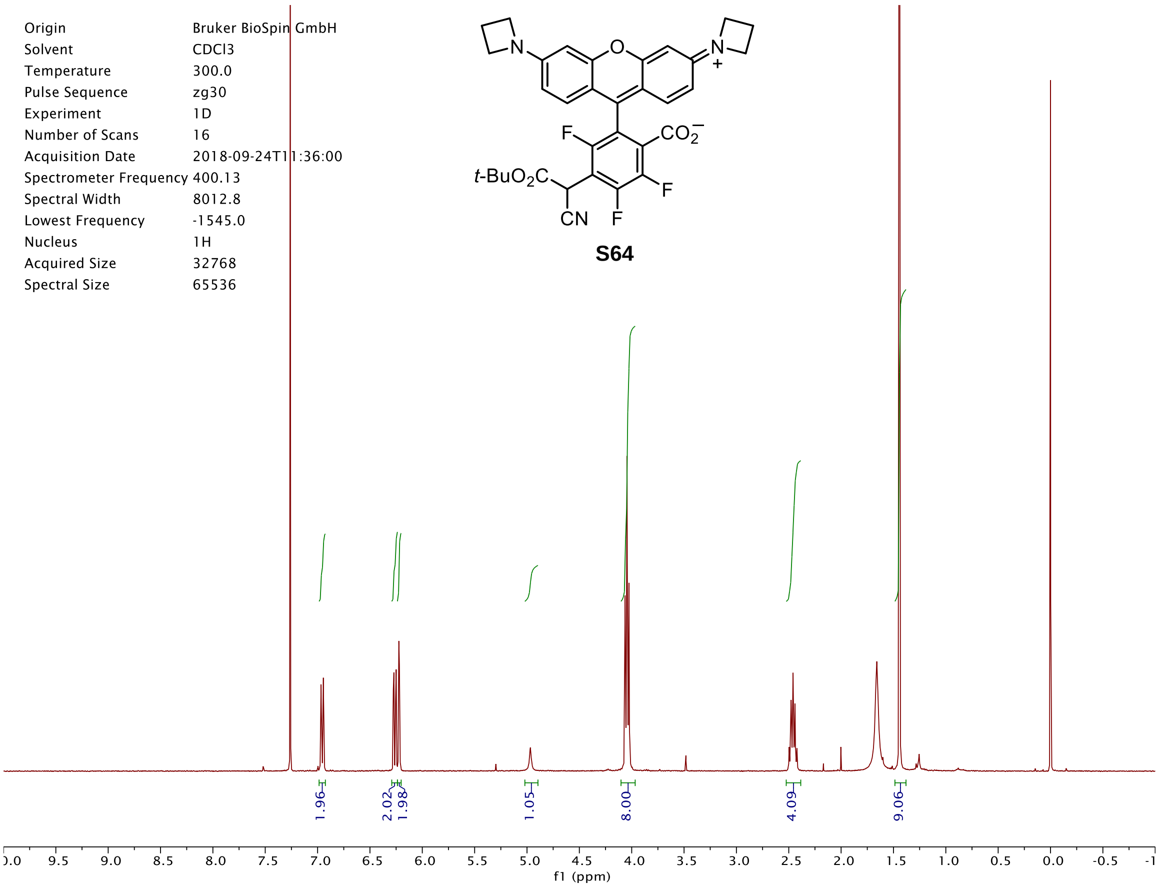
Origin Bruker BioSpin GmbH
Solvent CDCl3
Temperature 300.0
Pulse Sequence zg30
Experiment 1D
Number of Scans 16
Acquisition Date 2018-09-24T11:36:00
Spectrometer Frequency 400.13
Spectral Width 8012.8
Lowest Frequency -1545.0
Nucleus 1H
Acquired Size 32768
Spectral Size 65536
t-BuO2C
CN
F
CO2−
N+
S64
1.96
2.02
1.98
1.05
8.00
4.09
9.06
f1 (ppm)

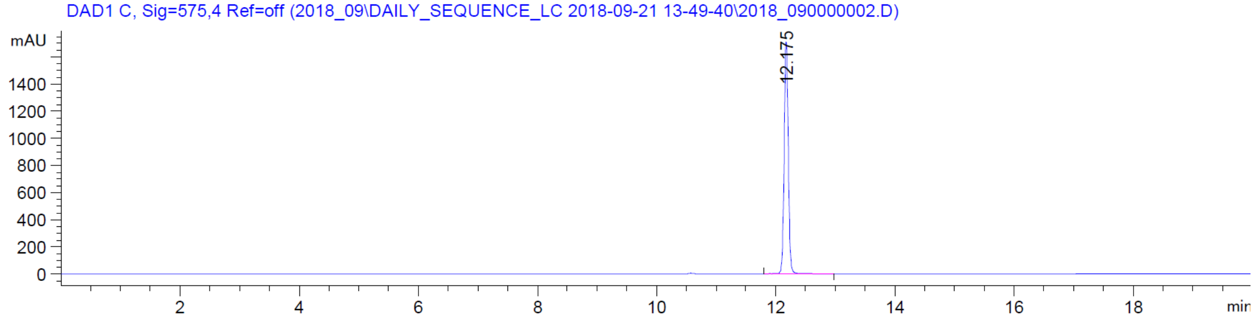
DAD1 C, Sig=575,4 Ref=off (2018_09\DAILY_SEQUENCE_LC 2018-09-21 13-49-40\2018_090000002.D)
mAU
12.175
min

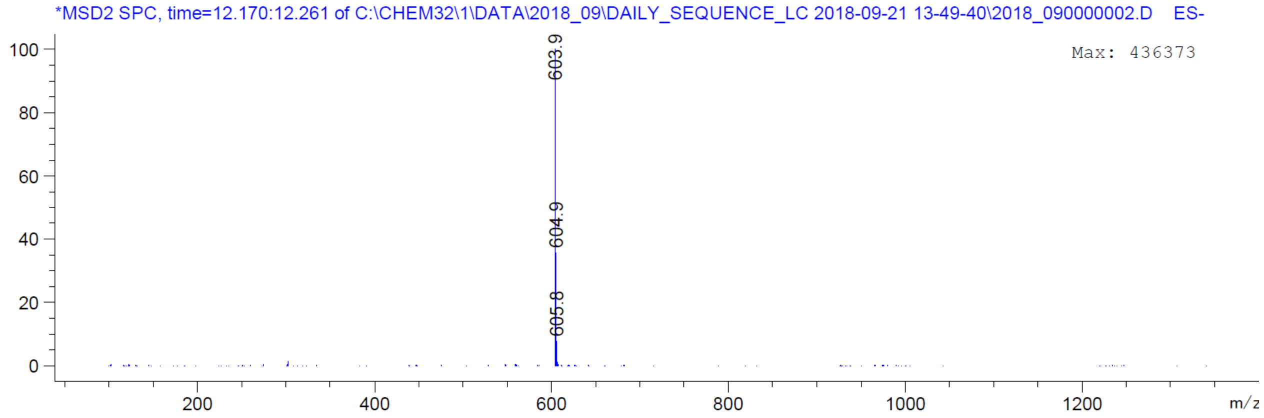
*MSD2 SPC, time=12.170:12.261 of C:\CHEM32\1\DATA\2018_09\DAILY_SEQUENCE_LC 2018-09-21 13-49-40\2018_090000002.D ES-
Max: 436373
603.9
604.9
605.8
m/z

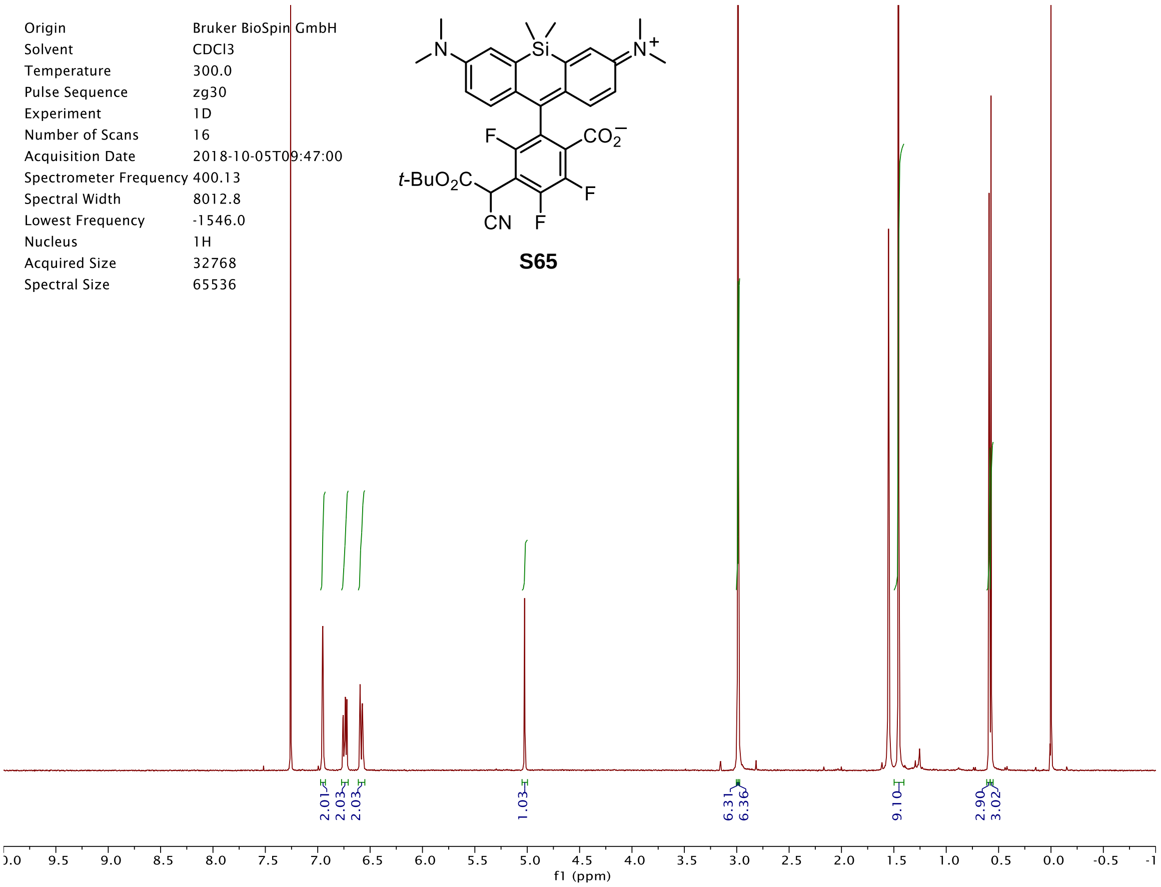
Origin Bruker BioSpin GmbH
Solvent CDCl3
Temperature 300.0
Pulse Sequence zg30
Experiment 1D
Number of Scans 16
Acquisition Date 2018-10-05T09:47:00
Spectrometer Frequency 400.13
Spectral Width 8012.8
Lowest Frequency -1546.0
Nucleus 1H
Acquired Size 32768
Spectral Size 65536
S65
2.01
2.03
2.03
1.03
6.31
6.36
9.10
2.90
3.02
f1 (ppm)

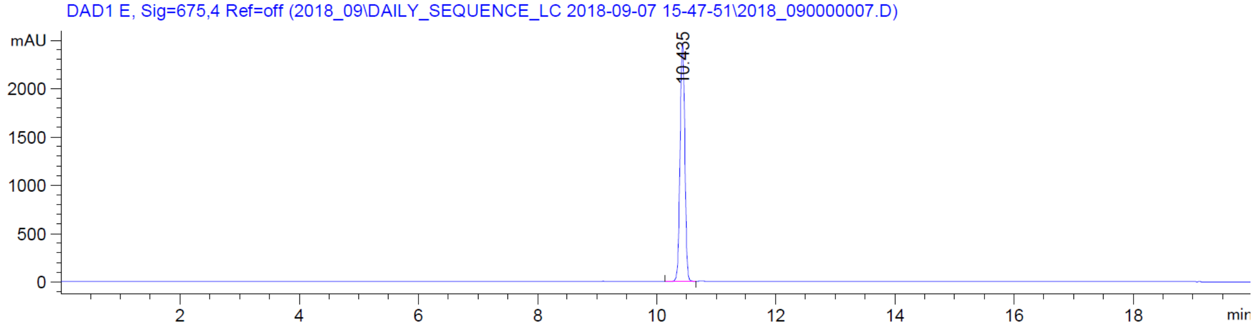
DAD1 E, Sig=675,4 Ref=off (2018_09\DAILY_SEQUENCE_LC 2018-09-07 15-47-51\2018_090000007.D)
mAU
10.435
min

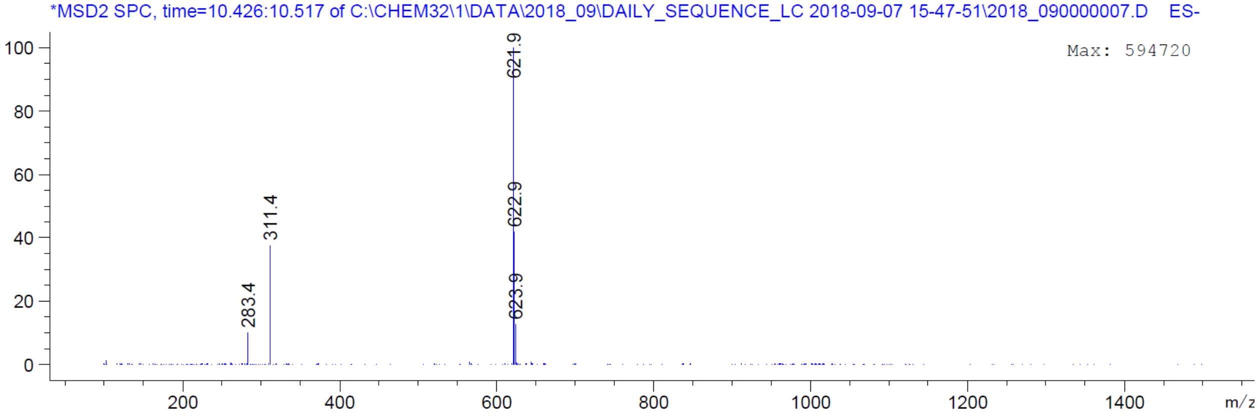
*MSD2 SPC, time=10.426:10.517 of C:\CHEM32\1\DATA\2018_09\DAILY_SEQUENCE_LC 2018-09-07 15-47-51\2018_090000007.D ES-
Max: 594720
621.9
622.9
623.9
311.4
283.4
m/z

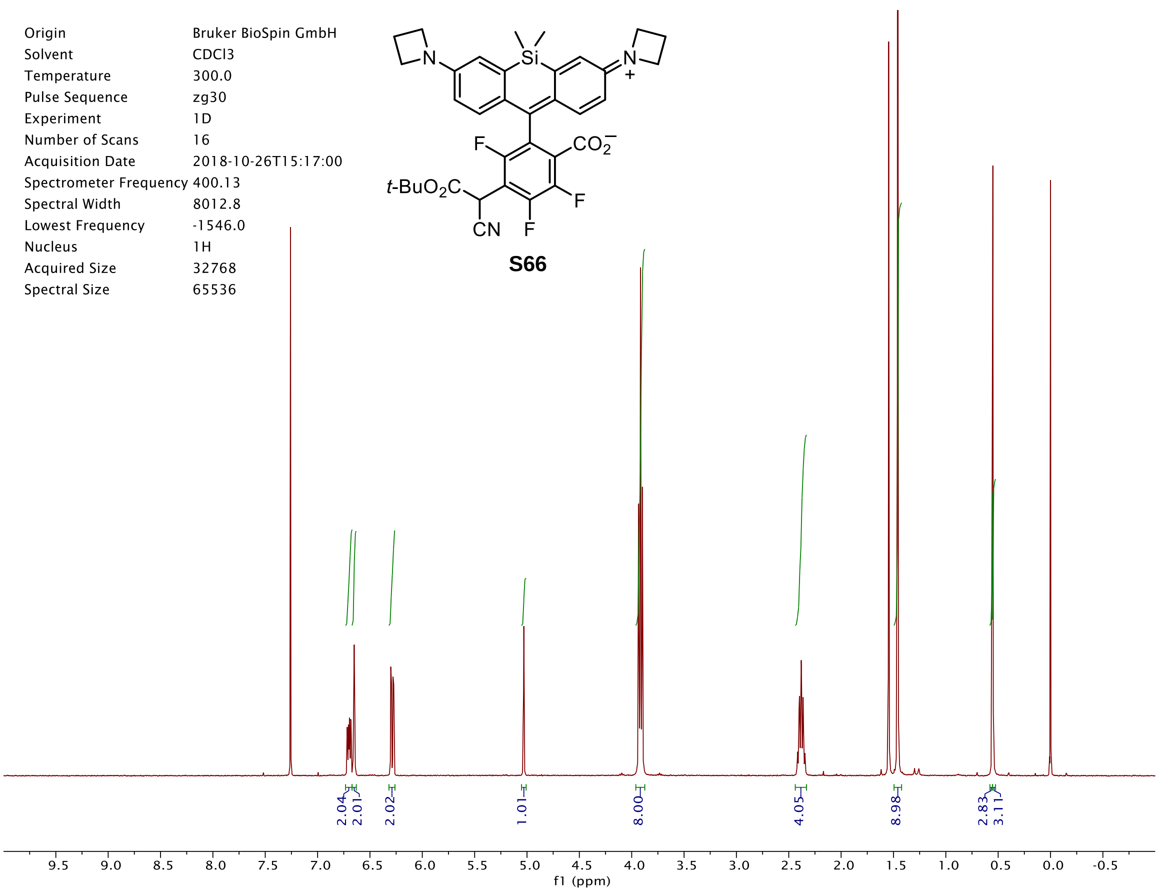

Origin Bruker BioSpin GmbH
Solvent CDCl3
Temperature 300.0
Pulse Sequence zg30
Experiment 1D
Number of Scans 16
Acquisition Date 2018-10-26T15:17:00
Spectrometer Frequency 400.13
Spectral Width 8012.8
Lowest Frequency -1546.0
Nucleus 1H
Acquired Size 32768
Spectral Size 65536
t-BuO2C
CN
F
CO2−
Si
N
+
S66
2.04
2.01
2.02
1.01
8.00
4.05
8.98
2.83
3.11
f1 (ppm)


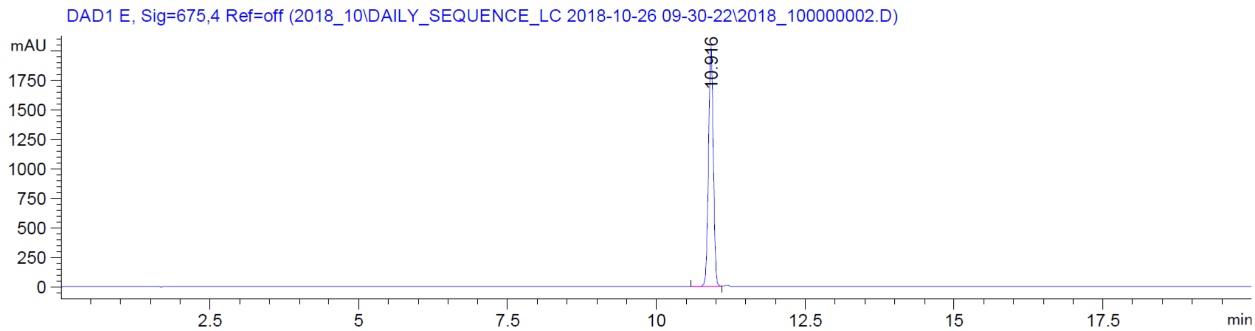

DAD1 E, Sig=675,4 Ref=off (2018_10\DAILY_SEQUENCE_LC 2018-10-26 09-30-22\2018_100000002.D)
mAU
10.916
min


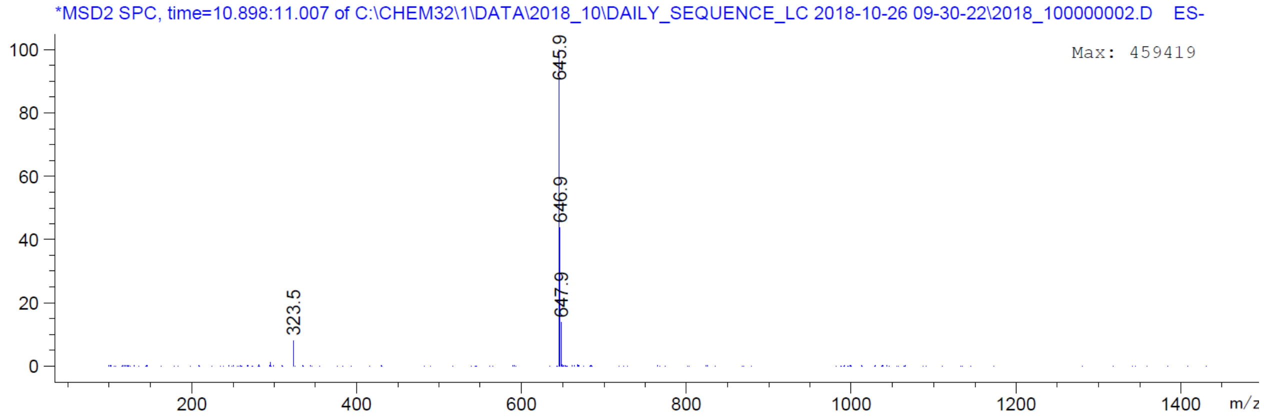

*MSD2 SPC, time=10.898:11.007 of C:\CHEM32\1\DATA\2018_10\DAILY_SEQUENCE_LC 2018-10-26 09-30-22\2018_100000002.D ES-
Max: 459419
645.9
646.9
647.9
323.5
m/z

| | |
|---|---|
| Origin | Bruker BioSpin GmbH |
| Solvent | DMSO |
| Temperature | 350.2 |
| Pulse Sequence | zg30 |
| Experiment | 1D |
| Number of Scans | 16 |
| Acquisition Date | 2019-01-28T16:13:00 |
| Spectrometer Frequency | 400.13 |
| Spectral Width | 8012.8 |
| Lowest Frequency | -1538.6 |
| Nucleus | 1H |
| Acquired Size | 32768 |
| Spectral Size | 65536 |

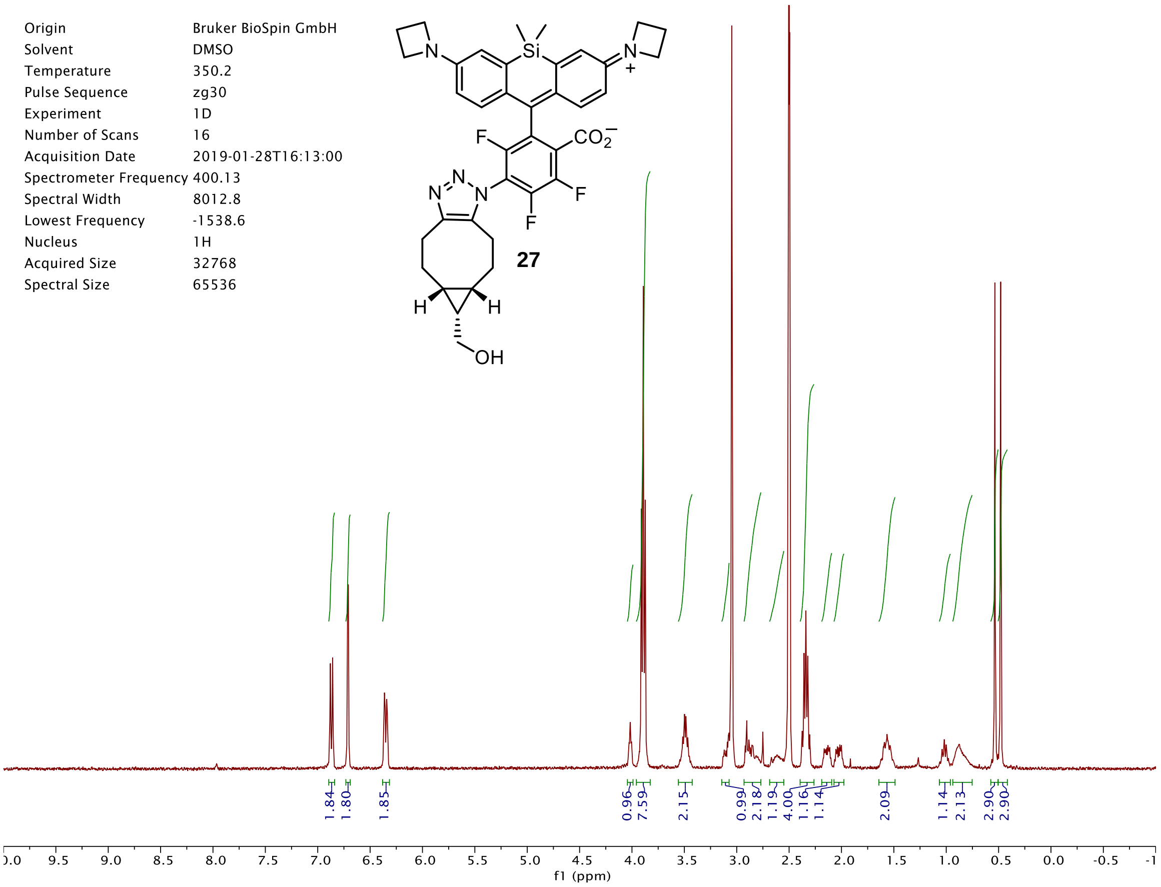


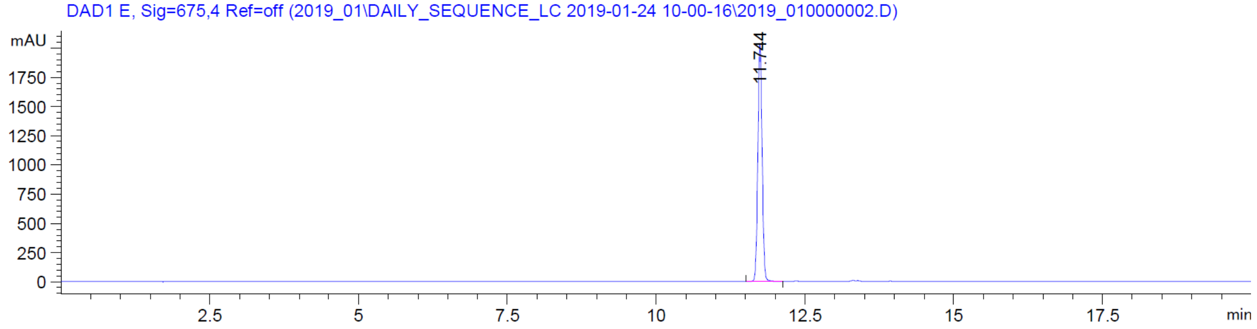


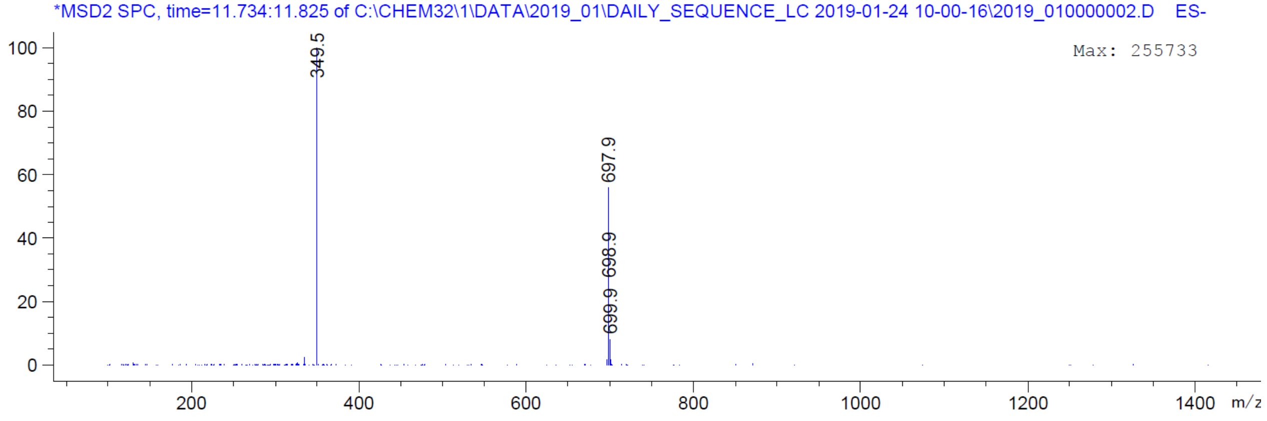

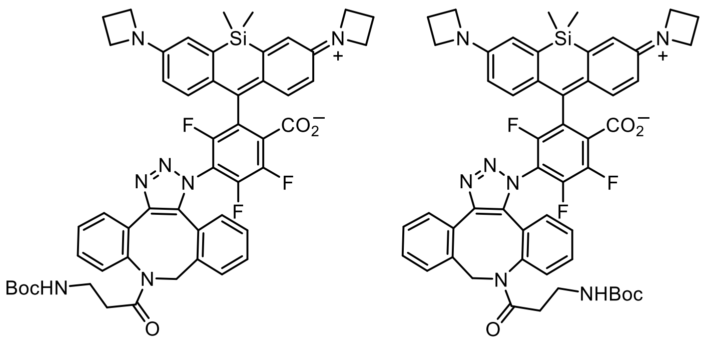

**28**

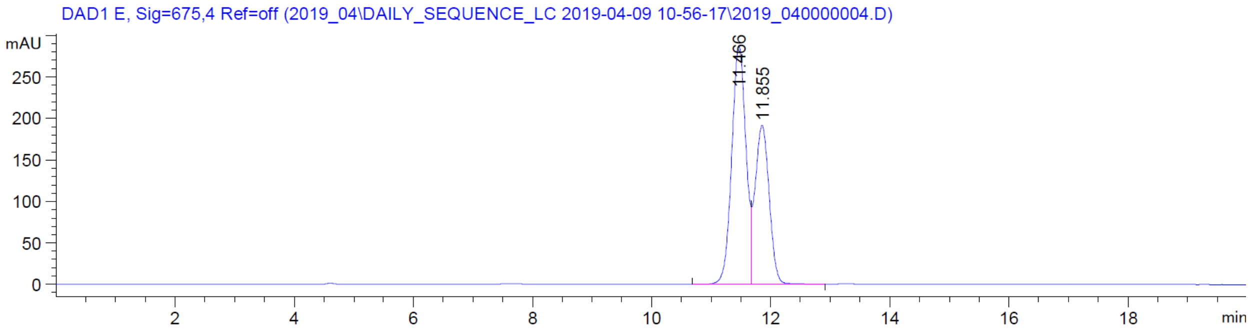
DAD1 E, Sig=675,4 Ref=off (2019_04\DAILY_SEQUENCE_LC 2019-04-09 10-56-17\2019_040000004.D)
mAU
11.466
11.855
min

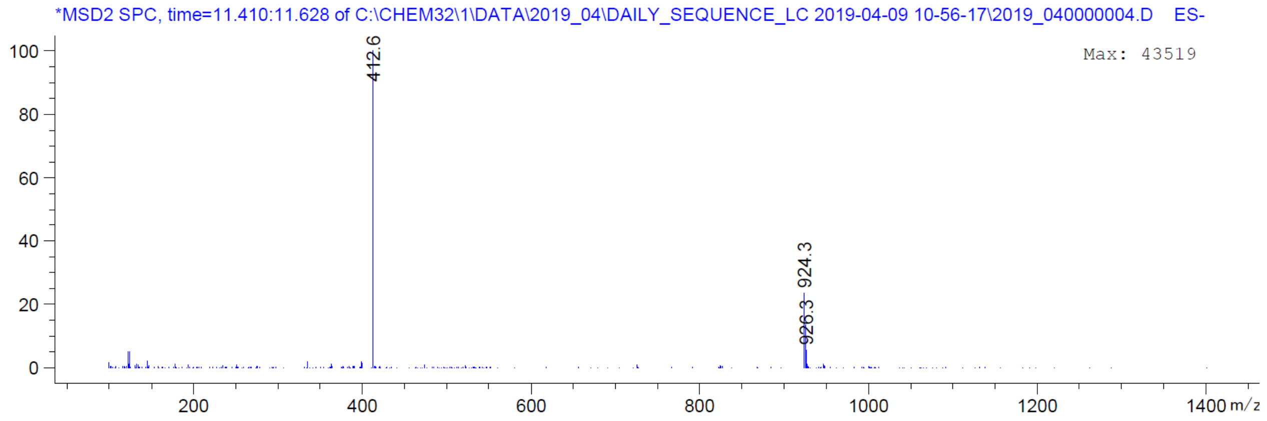
*MSD2 SPC, time=11.410:11.628 of C:\CHEM32\1\DATA\2019_04\DAILY_SEQUENCE_LC 2019-04-09 10-56-17\2019_040000004.D ES-
Max: 43519
412.6
924.3
926.3
m/z

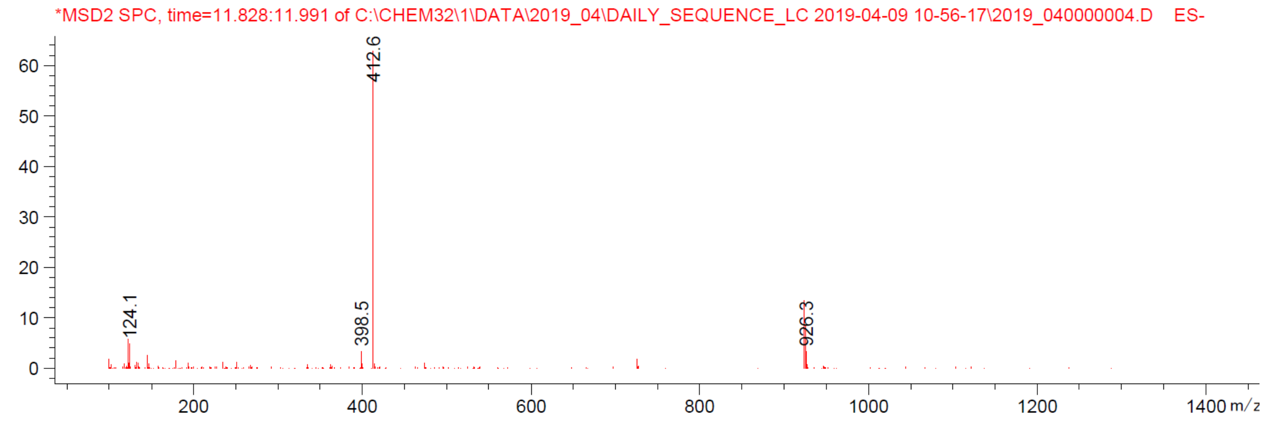
*MSD2 SPC, time=11.828:11.991 of C:\CHEM32\1\DATA\2019_04\DAILY_SEQUENCE_LC 2019-04-09 10-56-17\2019_040000004.D ES-
412.6
124.1
398.5
926.3
m/z
